# Evolved for success in novel environments: The round goby genome

**DOI:** 10.1101/708974

**Authors:** Irene Adrian-Kalchhauser, Anders Blomberg, Tomas Larsson, Zuzana Musilova, Claire R Peart, Martin Pippel, Monica Hongroe Solbakken, Jaanus Suurväli, Jean-Claude Walser, Joanna Yvonne Wilson, Magnus Alm Rosenblad, Demian Burguera, Silvia Gutnik, Nico Michiels, Mats Töpel, Kirill Pankov, Siegfried Schloissnig, Sylke Winkler

## Abstract

Since the beginning of global trade, hundreds of species have colonized territories outside of their native range. Some of these species proliferate at the expense of native ecosystems, i.e., have become invasive. Invasive species constitute powerful *in situ* experimental systems to study fast adaptation and directional selection on short ecological timescales. They also present promising case studies for ecological and evolutionary success in novel environments.

We seize this unique opportunity to study genomic substrates for ecological success and adaptability to novel environments in a vertebrate. We report a highly contiguous long-read based genome assembly for the most successful temperate invasive fish, the benthic round goby (*Neogobius melanostomus*), and analyse gene families that may promote its impressive ecological success.

Our approach provides novel insights from the large evolutionary scale to the small species-specific scale. We describe expansions in specific cytochrome P450 enzymes, a remarkably diverse innate immune system, an ancient duplication in red light vision accompanied by red skin fluorescence, evolutionary patterns in epigenetic regulators, and the presence of genes that may have contributed to the round goby’s capacity to invade cold and salty waters.

A recurring theme across all analyzed gene families are gene expansions. This suggests that gene duplications may promote ecological flexibility, superior performance in novel environments, and underlie the impressive colonization success of the round goby. *Gobiidae* generally feature fascinating adaptations and are excellent colonizers. Further long-read genome approaches across the goby family may reveal whether the ability to conquer new habitats relates more generally to gene copy number expansions.

## Introduction

Since the beginning of global trade and the colonial period, hundreds of species have colonized territories outside their native range. A fraction of those species proliferates at the expense of native species and ecosystems, i.e., they are invasive. While invasive species present challenges for biodiversity and ecosystem conservation, they also constitute exciting eco-evolutionary models for adaptation and ecological success in novel or changing environments (1–4).

The benthic round goby *Neogobius melanostomus* (Figure 1A) is one of the most widespread invasive fish species. Since 1990, round gobies have been detected in over 20 countries outside their native Ponto-Caspian range. In some regions of Europe and North America, they have become the most common fish species (5–7) (Figure 1B). Lasting impacts on biodiversity and on ecosystems have been observed (see (8) for a summary of the impacts). In recent years, the round goby has therefore become a novel model for ecology, behavior and evolution (Figure 1C).

**Figure 1.**
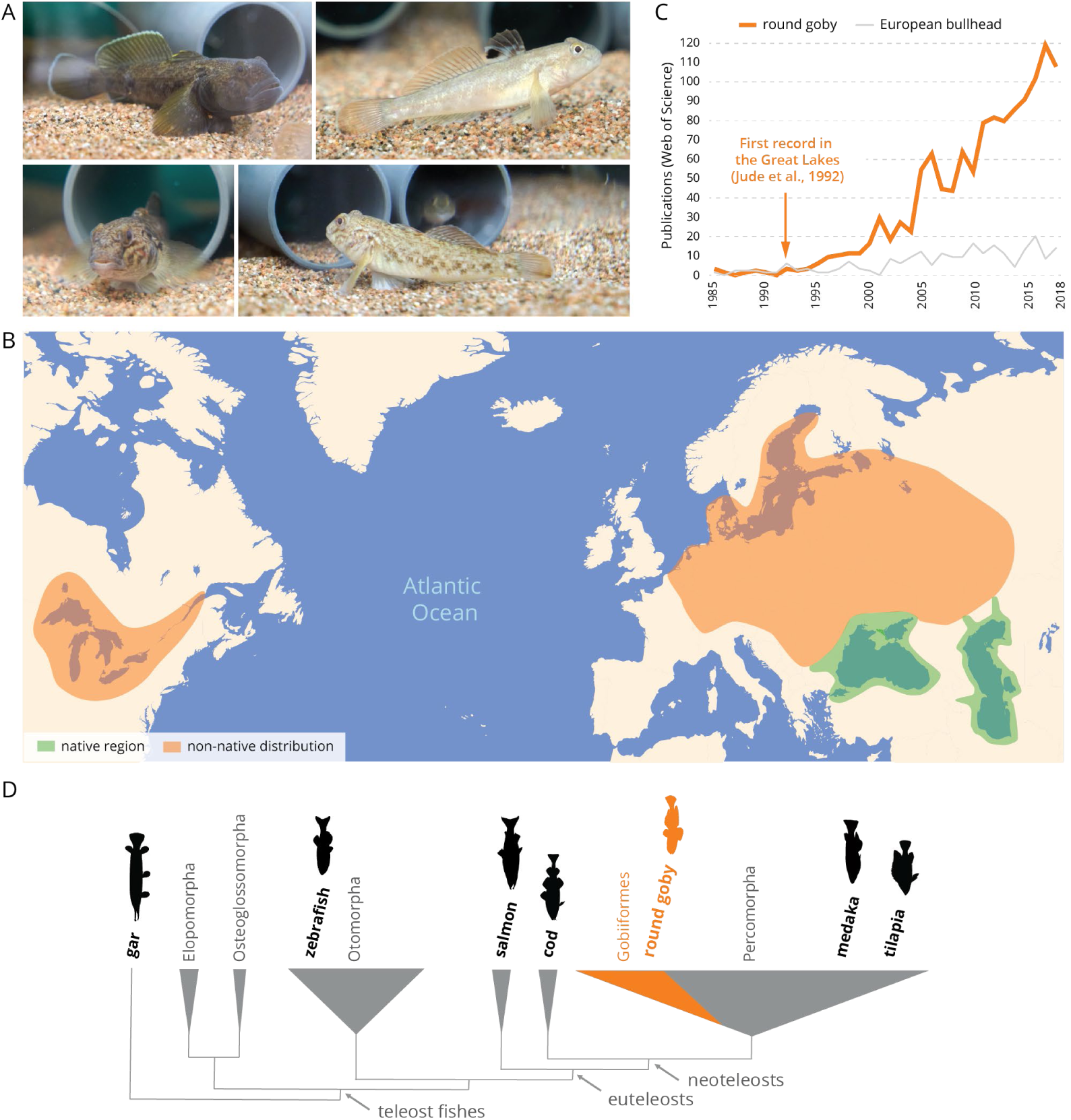
The round goby, a benthic invasive fish species. **A** Wild-caught round goby in aquaria. Individuals are usually brightly colored or spotted with a characteristic black dot on the first dorsal fin. During the reproductive season, territorial males develop a black body color (first panel). **B** The growing scientific relevance of the species is reflected by records in Web of Science (orange) when compared to a non-invasive fish with similar ecology (European bullhead, Cottus gobio; grey). **C** Current distribution of round goby. The round goby has spread from its native region (green) to many European rivers and lakes, the Baltic Sea, the Great Lakes and their tributaries (orange). **D** Phylogenetic position of the round goby among fishes.

The round goby effortlessly outcompetes native species with similar ecology, and is therefore a promising candidate to study fundamental questions on long-term ecological and evolutionary success. Round goby sequence data are presently restricted to a handful of phylogenetic markers (9–15). However, genome analyses have previously provided significant insights into fish ecology and evolution. Examples are genome compaction (16), the transition from fin to limb (17), loss of major parts of adaptive immunity (18), or effects of genome duplication (19). We therefore expect relevant insights into round goby biology and into success in novel environments from the round goby genome sequence.

The survival of an individual in a novel environment depends on how well it can perceive, react to, and accommodate to its new surroundings. In this study, we therefore explore a high quality and contiguous genome assembly of the round goby for genes related to three categories: environmental perception, reaction to environmental conditions, and long-term accommodation to novel environments. We focus on gene families that have been hypothesized to play a role in the colonization of novel environments and on gene families that may relate to round goby invasion ecology.

For environmental perception, we investigated genes responsible for sensory perception in fishes. We specifically focused on the opsin genes for visual perception, as well as on the olfactory receptors for odor perception. Vision in fishes is often specifically adapted to environmental conditions, such as darkness in deep water (20), modified color spectrum in turbid water (21, 22), habitat color (23), or specific light regimes or light compositions (24–26). The overall spectral sensitivity range of teleost fishes exceeds the human visual range and, in many cases, includes the UV (23) and far-red (27) spectrum. Similarly, olfaction is an essential chemoreception sense for fish, allowing for fast responses to predators and alarm cues as well as for intra-species communication. Pheromones have play an important role in the round goby (28–30), and males attract females into their nests by releasing them (31). A particularly specialized sense of smell therefore may provide an advantage during initial population establishment in novel environments.

We further investigated genes that may mediate responses to novel environments, namely genes involved in detoxification, ion transport and the immune system. The round goby occurs even in chemically contaminated harbors (32–34) and appears to tolerate xenobiotic compounds well. This suggests that the round goby may be particularly well equipped to degrade and eliminate chemical pollutants. We therefore analyze the cytochrome P450 gene superfamily, which is a particularly important and conserved part of the xenobiotic response (35). The round goby also tolerates a wide range of salinities (0 to 25 PSU / ‰) and temperatures (0°C-30°C) and occurs at latitudes ranging from <40° N in the Ponto-Caspian region to >60° N in the Baltic Sea. Most fishes tolerate only a narrow range of salinities (36); the round goby however belongs to a specialized group, the euryhaline fish species, which thrive in fresh and brackish environments and includes estuarine species and migratory species such as salmon. We study the genetic basis of osmoregulation and osmolyte production in round goby to gain insights into the evolution of salinity and cold tolerance, and to possibly predict future range expansions. Invasive species encounter an array of previously unknown pathogens when they colonize a habitat, and invasion success may be related to a species’ ability to tackle novel immune challenges (37). Intriguingly, the round goby displays a low parasite load at the invasion front (38). We therefore characterize key factors of the innate and the adaptive immune system.

We also investigated conserved gene regulators because such genes might be involved in long-term adaptation to a novel environment. Mechanisms such as DNA methylation and histone modifications promote long- and short-term gene expression regulation and therefore mediate adaptations to altered conditions at the cellular level (39), but also regulate genome-scale evolutionary processes such as the distribution of meiotic recombination events (40) or transposon activity (41), and provide stochastic variability as basis for selection (42). Epigenetic variants have been proposed to cause fitness-relevant differences in gene expression and phenotype (43, 44). The ecological flexibility of the round goby has been linked to enhanced gene expression plasticity in response to environmental stimuli (45) and to their ability to pass information on water temperature to their offspring through maternal RNA (46). To understand the features of core epigenetic regulators in the round goby, we focused on two widely conserved and well characterized parts of the epigenetic machinery: the histone-methylating PRC2 complex and the DNA methylases. Both mechanisms are thought to restrict developmental plasticity, downregulate gene expression (at least in mammals), and have been linked to plastic responses, behavioral changes, and environmental memory (47–51).

Finally, we take advantage of the high genome contiguity to investigate sex determination using RAD sequencing data. Fish display a wide variety of sex determination mechanisms, ranging from sex chromosomes to multilocus genetic sex determination to environmental sex determination (52), and sex determination in the round goby has not previously been investigated.

## Results

### 1. The round goby genome

The round goby genome assembly (“RGoby_Basel_V2”, BioProject accession PRJNA549924, BioSample SAMN12099445, GenBank genome accession VHKM00000000, release date July 22 2019) consists of 1364 contigs with a total length of 1.00 Gb (1’003’738’563 bp), which is within the expected size range (53–55). It is assembled to high contiguity (NG50 at 1’660’458 bp and N50 at 2’817’412 bp). GC content is 41.60%. An automated Maker gene annotation predicts a total of 38,773 genes and 39,166 proteins, of which 30,698 are longer than 100 amino acids (Table 1; annotation track available as **Supplemental_Material_S1**). The genome does not appear to contain a sex chromosome or a large sex determining region, since a RAD-tag dataset from 40 females and 40 males with an estimated resolution of 25,000 – 45,000 bp does not contain any sex-specific loci.

**Table 1.**
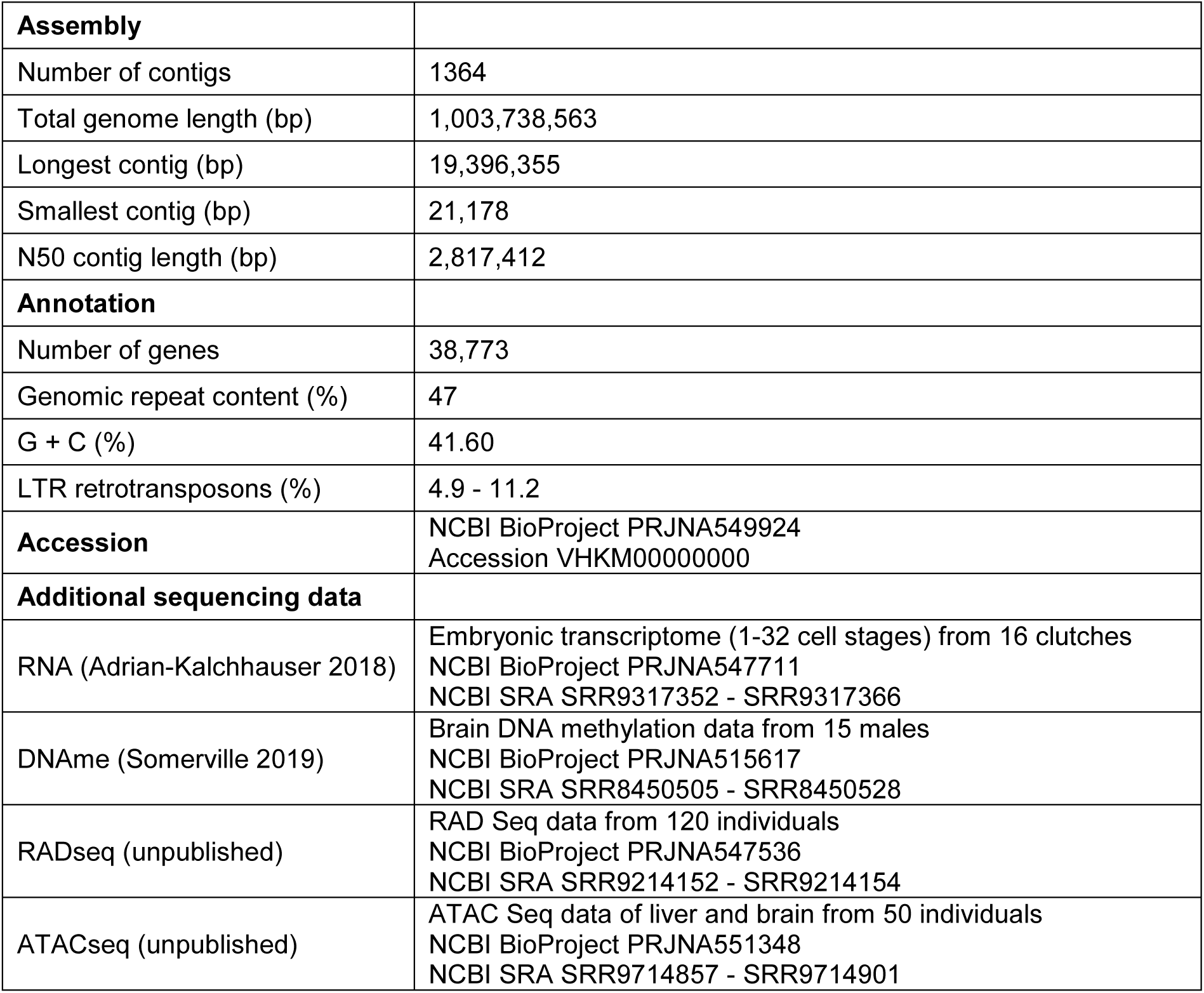
Round goby genome assembly and annotation statistics.

Approximately 47% of the genome assembly is masked as various types of repetitive sequences by RepeatMasker in the Maker annotation pipeline. The genome consists of approximately 9% predicted interspersed repeats (**Supplemental_Table_S1**), which is much lower than for zebrafish (*Danio rerio*, total genome size 1427.3Mb, 46% predicted as interspersed repeats) but higher than for the more closely related three-spined stickleback (*Gasterosteus aculeatus*, total genome size 446.6Mb, 3.2% predicted as interspersed repeats). Among interspersed repeats, the long terminal repeat (LTR) retrotransposon family is the most common in many species including fish (Repbase, https://www.girinst.org/repbase/). RepeatMasker identifies 0.9% LTR retrotransposons in the round goby genome, but separately run *de novo* predictions with LTRfinder and LTRharvest (**Supplemental_Table_S1)** indicate an underestimation of LTR retrotransposons and interspersed repeats by this approach. The latter approaches estimate that the proportion of LTR retrotransposons in the round goby genome is 11.2% (3.8% LTRs with target-site-repeats; LTRfinder) or 4.9% (LTRharvest), respectively.

In addition to the genome sequence, we provide raw short read sequencing data from various published and ongoing projects. They include RNA sequencing data from early cleavage embryos (46), DNA methylation capture data from adult male brains (47), as well as RAD tags from two local Swiss populations and ATAC seq reads from brain and liver (unpublished; Table 1).

### 2. Sensory perception genes: Vision

Vertebrates perceive color with cone cells expressing one of four types of opsin proteins (usually sensitive to the red, green, blue, and ultraviolet part of the spectrum) and dim light with rod cells expressing the rod opsin. The UV and blue light is detected by the short-wavelength sensitive SWS1 and SWS2 opsins, the green part of the spectrum is perceived mostly by the rhodopsin-like RH2 opsins, and the red color by the long-wavelength sensitive LWS opsins. Rod cells are active in the dim-light conditions and contain the rod opsin RH1 (56). Gene duplications and losses of the opsin genes during fish evolution correlate to certain extent with adaptations to specific environments (20, 57).

We identified two cone opsin gene duplications in the round goby genome. Firstly, the genome features a recent duplication of the green-sensitive RH2 gene. RH2 duplications are a common phenomenon in fish (**Figure 2**). Secondly, the genome features an ancient duplication of the long-wave red-sensitive LWS gene. The event can be traced most likely to the ancestor of all teleosts, or possibly even to the ancestor of Neopterygii (**Figure 2**; see **Supplemental_Fig_S1** for full tree). As expected, the round goby genome further contains one dim-light rod opsin (RH1) gene, two blue-sensitive SWS2 genes (57), and as previously reported for gobies, lacks the UV/violet-sensitive SWS1 gene (20, 25, 57).

**Figure 2.**
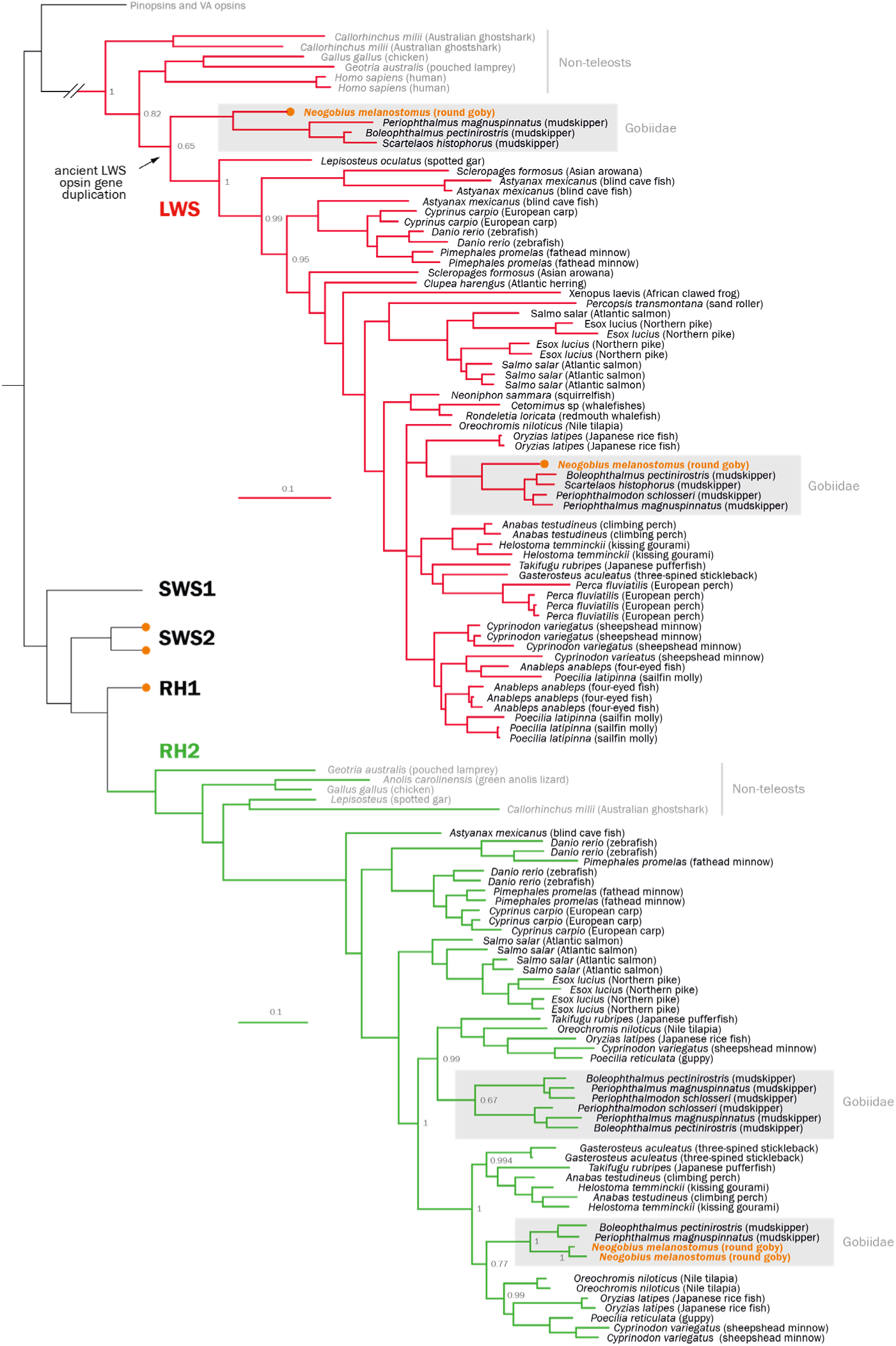
Phylogenetic tree of vertebrate opsin gene sequences. Maximum-likelihood phylogenetic tree based on the cone and rod visual opsins and using VA opsins and pinopsins as outgroup. The round goby genome contains two LWS gene copies, which seem to be the results of an ancient gene duplication event, and two more recently duplicated RH2 gene copies. Round goby is indicated in orange. Red opsin branches are indicated in red. Green opsin branches are indicated in green. Non-teleost species and the outgroup (VA opsins and pinopsins) are indicated in grey. Grey boxes highlight Gobiidae.

The proposed ancestral position of the red opsin gene duplication is supported by three lines of evidence. First, the monophyly of all other teleost + gar LWS genes is strongly supported by the Bayesian analysis (Bayesian posterior probability value = 1). Second, the distant phylogenetic position is supported by trees based on individual exons, which indicate a low probability of a compromised phylogenetic signal, e.g. due to the partial gene conversion. Three of four exons cluster at the same position as the whole gene, while the fourth exon (exon 4) cluster with the genes resulting from a more recent teleost-specific LWS duplication specific to *Astyanax* and *Scleropages* (58) (**Supplemental_Fig_S2**). Third, the choice of outgroup (parietopsin or pinopsin) does not affect the position of the LWS2 gene. Together, these analyses suggest either (1) the presence of an ancient gene duplication event of the LWS gene in the ancestor of teleost and holostean fishes (i.e. *Neopterygii*) which was retained only in the goby family, or (2) a teleost-specific event, possibly identical to that reported for characins and bony tongues (58), with a subsequent concerted goby-specific sequence diversification in exons 2, 3 and 5.

The spectral sensitivity of photopigments, i.e. their excitation wavelength can be modified by substitutions in certain key amino acids (59). We find that round goby LWS1 and LWS2 differ in the key spectral tuning site at amino acid 277 (position 261 of bovine rhodopsin, Table 2) suggesting a sensitivity shift of 10 nm.

**Table 2.** Amino acid analysis of key tuning sites in Gobiidae red opsins proteins.

| species | ecology | gene | key tuning amino acid site in round goby |  |  |  |  | max. spectral sensitivity (wavelength) | reference |
| --- | --- | --- | --- | --- | --- | --- | --- | --- | --- |
|  |  |  | 180 | 197 | 277 | 285 | 308 |  |  |
|  |  | *bovine rhodopsin equivalent site:* | 164 | 181 | 261 | 269 | 292 |  |  |
| *Boleophthalmus pectinirostris* | terrestrial mudskipper | LWS1 | A | H | Y | T | A | 553 nm | (25) |
|  |  | LWS2 | A | H | F | A | A | 531 nm |  |
| *Periophthalmus magnuspinnatus* | terrestrial mudskipper | LWS1 | S | H | Y | T | A | 560 nm |  |
|  |  | LWS2 | A | H | F | T | A | 546 nm |  |
| *Neogobius melanostomus* | freshwater temperate rivers and lakes | LWS1 | S | H | Y | T | A | 560 nm | this study |
|  |  | LWS2 | S | H | F | T | A | 550 nm* | this study |
| * = predicted by the key tuning sites, and Y261F shift of 10 nm; Yokoyama, 2008. |  |  |  |  |  |  |  |  |  |

To find a possible link to the ecological significance of the red opsin duplication, we checked for the presence of red skin fluorescence in the round goby. Interestingly, round goby individuals of both sexes and of all sizes (n=10) feature weakly red fluorescent crescents above the eyes (**Figure 3**). Whether such pattern has any relevance for the putatively enhanced vision in the red spectrum remains elusive.

**Figure 3.**
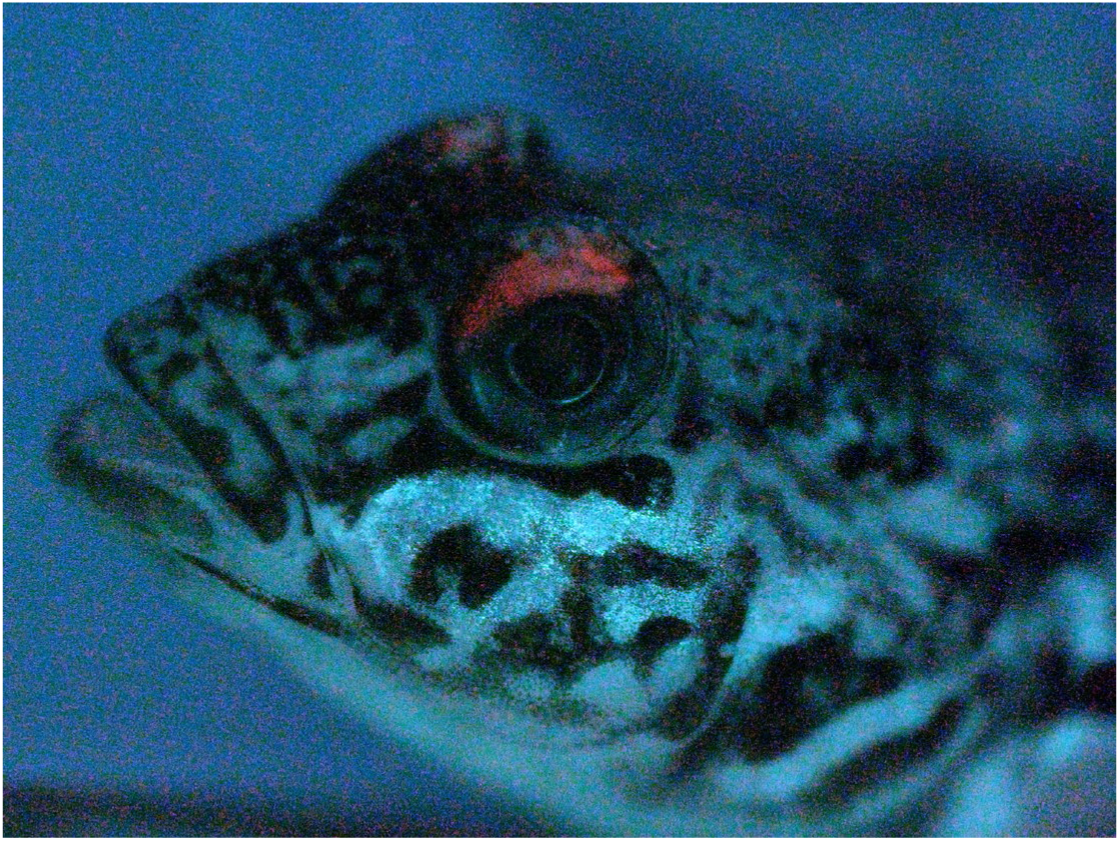
Red fluorescence in the round goby. Round gobies exhibit red fluorescence above the eyes when exposed to green light.

### 3. Sensory perception genes: Olfaction

Olfactory receptors (OR) in vertebrates are 7-transmembrane-domain G-protein coupled transmembrane proteins. They are expressed in neurons embedded in membranes of the olfactory lamellae. Mammals usually have several hundred OR genes that cluster in two major types (∼400 in human, and ∼1000 genes in mouse) (60). Teleost fishes possess fewer OR genes but feature a higher diversity (5 kinds of type 2 ORs in teleosts as compared to 2 kinds of type 2 ORs in mammals) (61). The binding properties of individual ORs, especially in fishes, are virtually unexplored.

We identified 112 putative olfactory receptor genes in the round goby genome. To put this result into evolutionary context, all analyses were carried out in comparison with two Gobiidae species (blue-spotted mudskipper and giant mudskipper) and two percomorph species (three-spined stickleback, *Gasterosteus aculeatus* and Nile tilapia, *Oreochromis niloticus*; **Figure 4A**). The round goby presented similar number of ORs (112) to the giant mudskipper (106) and stickleback (115), notably less than the blue-spotted mudskipper (154) and near half the amount compared to Nile tilapia (214). We find that all ORs belong to one of two transmembrane domain subtypes according to the Pfam database (7tm4 or 7tm1; **Figure 4B**; **Supplemental_Fig_S3**). This matches a previous large-scale phylogenetic analysis which identified two main types of olfactory receptor genes in vertebrates (61). The functional differences between the domain subtypes are unclear, but their different consensus sequences may confer distinct biochemical properties.

**Figure 4.**
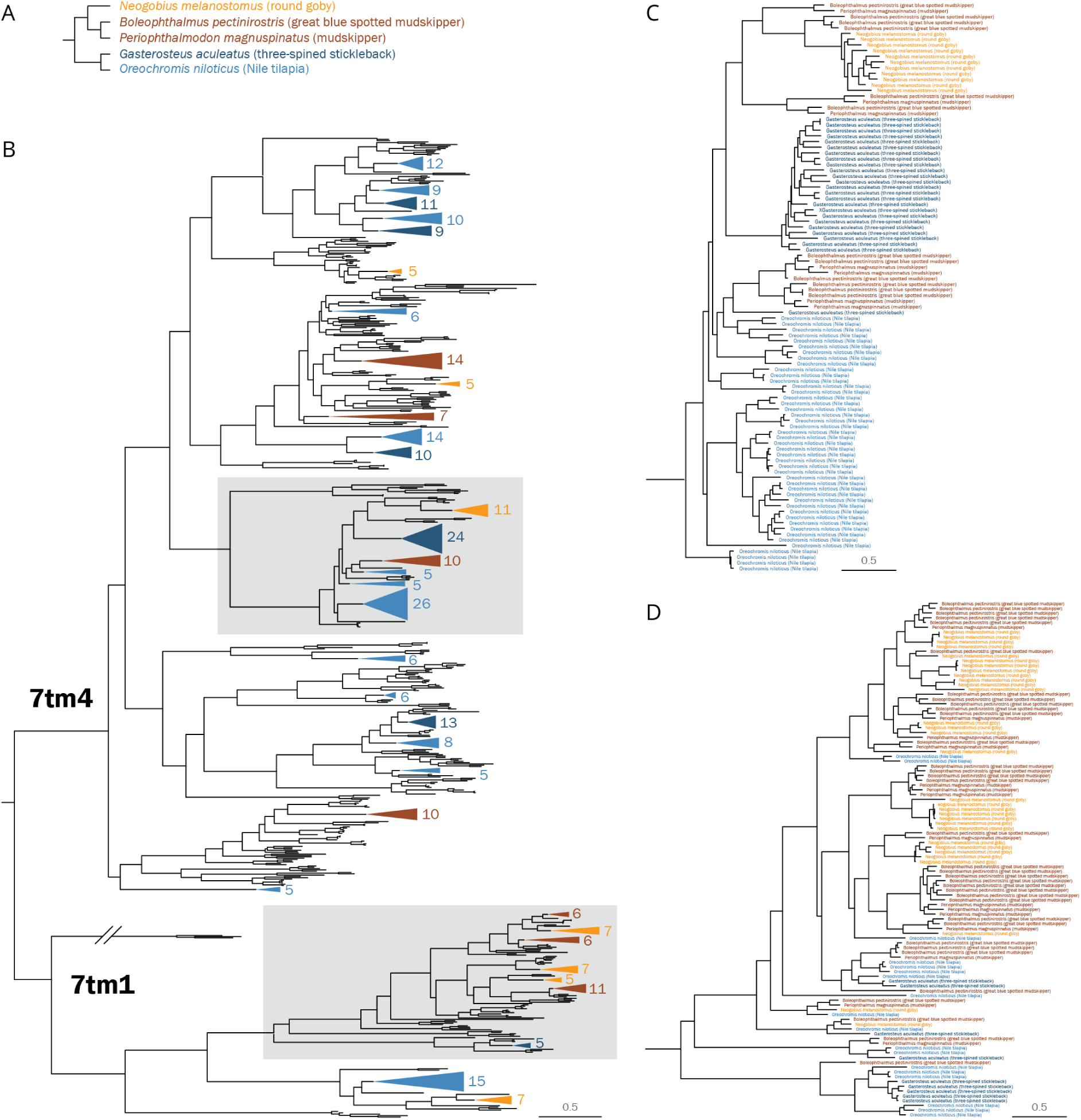
Phylogenetic tree of percomorph olfactory receptor protein sequences. **A** Phylogenetic relationship among five analyzed percomorph species, i.e. three gobiids, one cichlid and one stickleback. **B** Maximum-likelihood phylogenetic tree constructed with adrenergic receptors as outgroup. Sequences were identified de novo except for nile tilapia (blue). Branches magnified in panels C and D are highlighted with grey boxes. **C** Branch of the 7tm4 family featuring large independent expansions in all species analyzed. **D** Branch of the 7tm1 family featuring several expansions in Gobiidae (red, orange) that are not paralleled in other percomorph species (blue).

Our analyses identify several cases of clade-specific gene expansions. Certain OR genes are expanded in parallel in several lineages (**Figure 4C**). Likely, those expansion events are the result of clade-restricted gene duplications, although a secondary role for gene conversion after species divergence cannot be ruled out. While the Nile tilapia features the greatest overall amount of expansions, the round goby presents the highest number of genes and expansions within the 7tm1 subfamily, a trend that is consistent in the other Gobiidae species (**Figure 4D**).

### 4. Response to the environment: Detoxification

The CYP gene superfamily is an essential part of the defensome, a collection of genes that provide protection against harmful chemicals (35). Vertebrate genomes contain between 50-100 CYP genes. The genomes of fugu and zebrafish, for example, encode 54 (62) and 94 (63) CYP genes respectively. Expansions of individual CYP families occur in both mammals and fish. For example, zebrafish have three times as many CYP2 family members (40) as most other vertebrate species (13–15), and similar expansions of CYP2 genes have been observed in mice and rats (64).

We find that the round goby genome contains few CYP genes. We identify 25 complete or partial CYP genes, as well as 21 gene fragments. Pseudogenes are common for CYP genes (62, 63, 65), which is why strict annotation criteria are applied first before smaller fragments are considered. In total, the genome contains approximately 50 CYP genes (**Supplemental_Table_S2**).

When including gene fragments, all expected CYP families are present in the round goby, and the phylogenetic analyses show the expected relationships between gene families and between vertebrates (**Figure 5**). Fish and most vertebrates have CYP genes from 17 families (CYP 1-5, 7, 8, 11, 17, 19, 20, 21, 24, 26, 27, 46 and 51) (62), while the CYP39 family occurs in humans and zebrafish, but not in fugu (62, 63). In the round goby, the complete or partial genes could be assigned to 9 CYP families (CYP 1-4, 8, 19, 26, 27 and 51). The families CYP7, CYP11, CYP17 and CYP21 were present among the sequence fragments.

**Figure 5.**
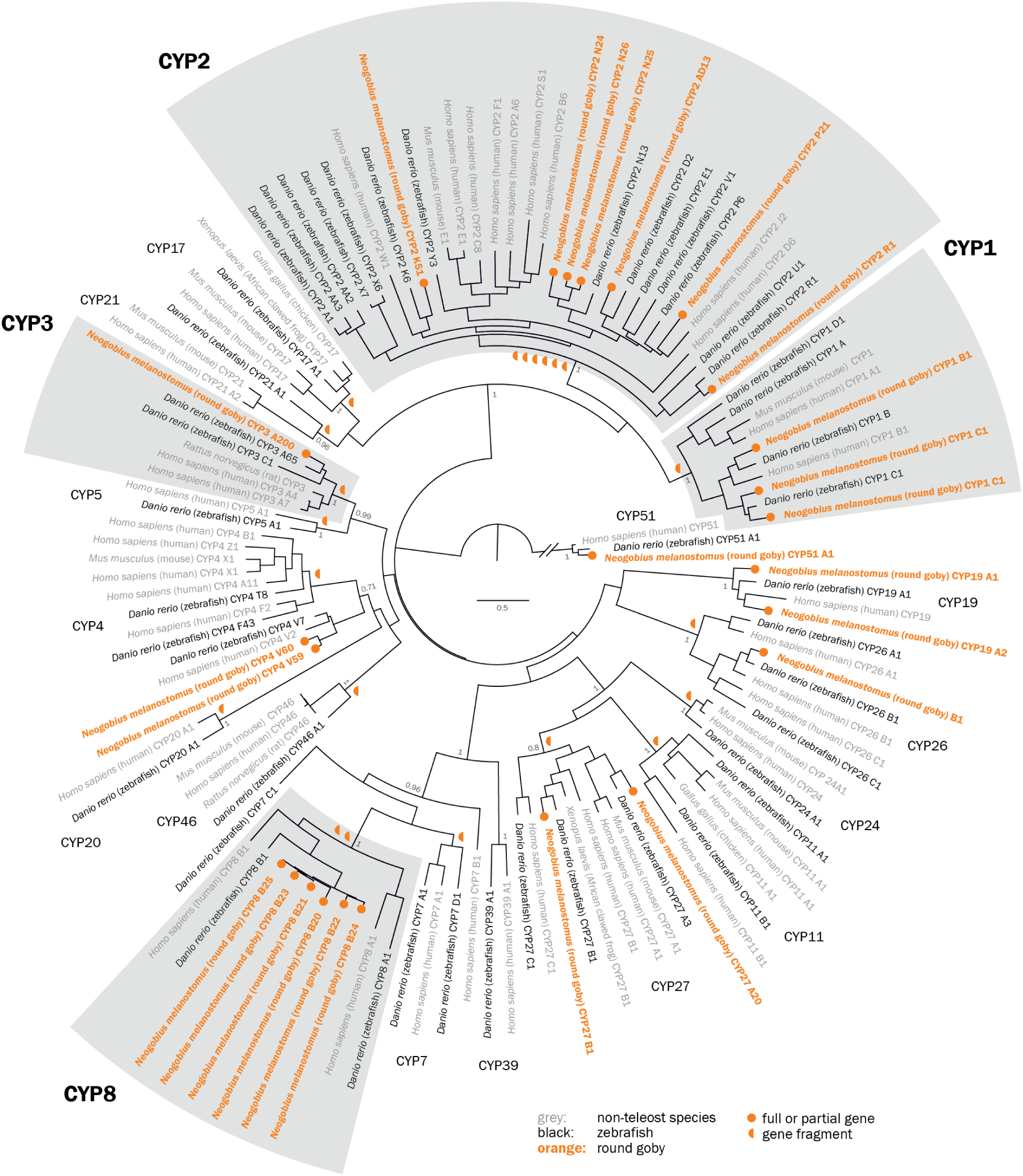
Phylogenetic tree of vertebrate CYP protein sequences. Maximum likelihood phylogenetic tree with 100 bootstraps, rooted with the CYP51 family. Detoxification genes CYP1-3 do not feature unusual expansions, while the CYP8 family is expanded to six members (grey boxes). Non-fish vertebrates are printed in grey. Fragments too short for tree building but attributable to a certain family are indicated by orange half circles next to the root of the respective family.

Contrary to expectations, the classical detoxification families CYP1-3 were not expanded (**Figure 5**). CYP1, 2, 3 and to a lesser extent CYP4 proteins are responsible for the oxidative metabolism of xenobiotic compounds (pollutants, drugs, etc.). In rodents and humans, the CYP1 family metabolizes planar cyclic aromatic hydrocarbon compounds (reviewed in (66)), the CYP2 family metabolizes structurally diverse drugs, steroids and carcinogens, the CYP4 family catalyzes the ω-hydroxylation of the terminal carbon of fatty acids and xenobiotics, and CYP3 genes metabolize a range of structurally different compounds in the liver and intestines. Over 50% of all pharmaceutical compounds are metabolized by CYP3A genes in human. The goby genome contains three or four CYP1 genes: one CYP1B gene, two CYP1C genes, and one CYP1A fragment. The latter lacks two main characteristics (I- and K-helix) and could therefore be a pseudogene. As expected for a vertebrate (64), the genome contains many CYP2 genes. The most important fish CYP2 families were represented, including CYP2J, CYP2N, CYP2Y and CYP2AD. Finally, the round goby had a single CYP3A gene and a potential CYP3A fragment. This is somewhat unusual because fish often feature species-specific CYP3 subfamilies in addition to CYP3A. For example, medaka also contains CYP3B genes, zebrafish CYP3C genes, and Acanthopterygii fish CYP3D genes (67).

We find that the round goby genome contains six CYP8 genes, which is more than expected based on observations from the other gobies. The closely related large blue-spotted mudskipper has only two CYP8 genes (XM_020924471 and XM_020919000.1; about 73-85% identity); no sequences were found in other mudskipper species. Accordingly, we assume that the CYP8B genes have undergone species-specific tandem duplications in the round goby, as is also known for the subfamilies CYP2AA, CYP2X and CYP2K in zebrafish (64). Five round goby CYP8 genes locate to the same contig with high sequence similarity (∼90%), which is similar to zebrafish CYP8B1-3 that also colocalize on the same chromosome (63). Misidentification of closely related CYP7, CYP8, and CYP39 genes as CYP8 is unlikely given the colocalization and high sequence similarity. The function of the expansion is presently unclear, although expression patterns in zebrafish suggest a role in the early embryo (63). In humans, CYP8 genes act as prostacyclin synthases that mediate steroid metabolic pathways in bile acid production or prostaglandin synthesis (68). Based on structural similarities with yeast proteins, CYP8 genes might also have E3 ubiquitin ligase activity. The almost identical crystal structures of zebrafish and human CYP8A1 suggest similar functions in fish and mammals (69).

### 5. Response to the environment: Osmoregulation

Osmotic homeostasis depends on passive ion and water uptake through cell membranes and the intercellular space, on the active uptake or excretion of ions, and on the production and accumulation of osmolytes. To understand the ability of round goby to tolerate a wide range of salinities, we therefore compared the round goby repertoire of osmoregulatory genes to those of a stenohaline freshwater species (zebrafish) and of euryhaline species (Nile tilapia, blue-spotted mudskipper and three-spine stickleback).

Passive ion and water transport across membranes (transcellular permeability) depends on the superfamily of aquaporin proteins. Aquaporins transport water (classical aquaporins), water and glycerol (aquaglyceroporins), ammonia (aquaammoniaporins), or additional undescribed molecules (unorthodox aquaporins; **Figure 6**). Primary sequences are only moderately conserved between the classes (approximately 30% identity), but all aquaporins share six membrane-spanning segments and five connecting loops. We find 15 aquaporin genes in the round goby, which compares to the number in human (n=13) or zebrafish (n=20) and is lower than in the euryhaline Atlantic salmon (n=42) (70, 71). With 5 classical water aquaporins, 6 aquaglyceroporins, 2 aquaammonioporins, and 2 unorthodox aquaporins, the round goby features the same types of aquaporins as freshwater stenohaline fish (e.g., zebrafish) and highly euryhaline fish (e.g., tilapia; **Figure 6**).

**Figure 6.**
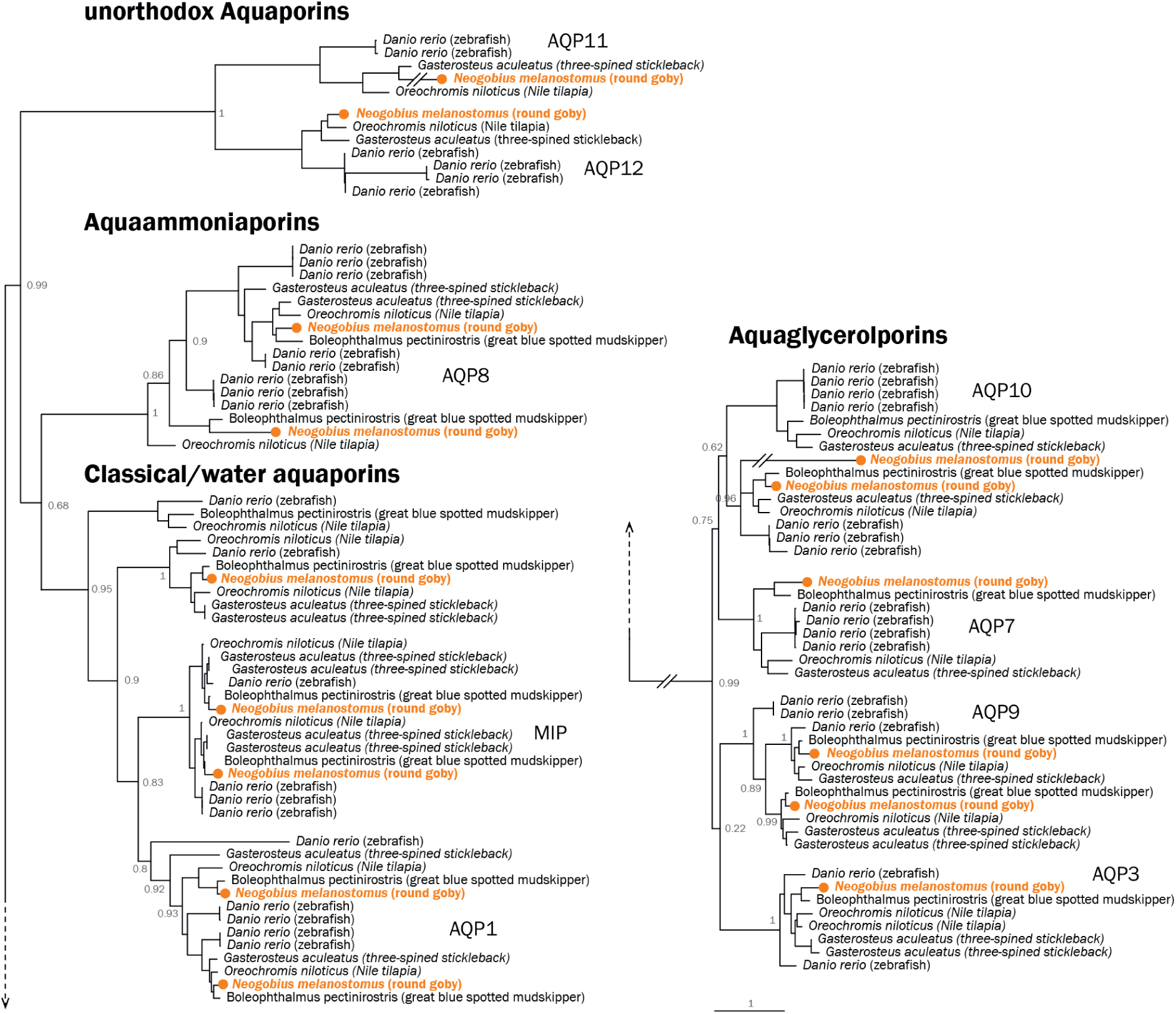
Phylogenetic tree of fish aquaporin proteins. Maximum-likelihood tree with 100 bootstraps of round goby (Neogobius melanostomus, orange) in relation to cyprinid zebrafish (Danio rerio) and percomorph three spine stickleback (Gasterosteus aculeatus), nile tilapia (Oreochromis niloticus), and great blue-spotted mudskipper (Boleophthalmus pectinirostris). Zebrafish was used as outgroup in each aquaporin subfamiliy. The main classes of aquaporins are labeled with human genes names.

Ion and water flow between cells in epithelia (paracellular permeability) is regulated by tight junctions, of which claudin and occludin proteins are the most important components. Mammalian genomes contain ∼ 20 claudin genes, invertebrates such as *Caenorhabditis elegans* or *Drosophila melanogaster* contain 4-5 genes, and fish often feature large expansions. For example, the fugu genome contains 56 claudins, of which some occur in clusters of > 10 genes (72). The round goby genome features 40 claudin paralogues, which is in line with numbers known from other fish. All human claudin genes were represented as homologues (**Supplemental_Fig_S4**), and the round goby genome contains one occludin gene in each of the two known subclades of the protein family (**Supplemental_Fig_S5**).

In the kidney, intestine and gills, fish use active ion transport (mostly sodium transporters) to maintain osmotic balance. Mechanisms mediating sodium uptake include electroneutral Na+/H+ exchange via the NHE3b protein, Na+/Cl-cotransport via the NCC protein, and coupling of Na+ absorption with H+ secretion by a V-type H+-ATPase (73). We find 12 Na+/H+ exchanger genes, 5 Na+-K+-ATPase catalytic alpha subunits and 6 Na+-K+-ATPase regulatory beta subunits in the round goby genome.

The round goby thus contains the same types of genes, but less copies, than either zebrafish or tilapia (**Supplemental_Fig_S6**). We find that round goby, and also mudskippers, feature an interesting distribution of Na+/Cl-co-transporters to subgroups; while most zebrafish and tilapia Na+/Cl-co-transporters belong to the NKCC1 subgroup, Gobiidae feature more genes in the NKCC2 subgroup (**Figure 7**).

**Figure 7.**
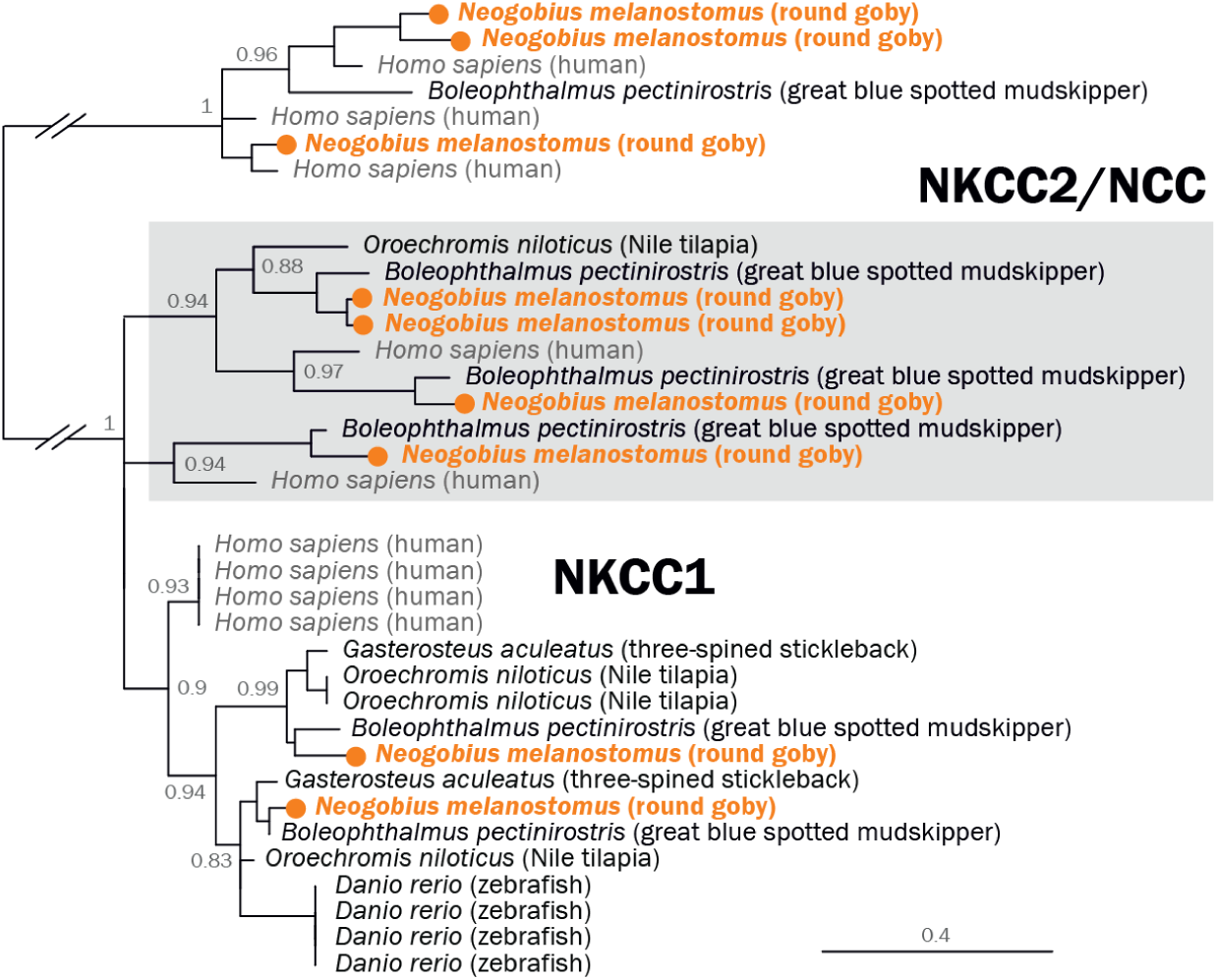
Phylogenetic tree of human and fish sodium/potassium/chloride co-transporter proteins (NKCC). Maximum likelihood tree with 100 bootstraps of round goby (Neogobius melanostomus, orange), zebrafish (Danio rerio), three spine stickleback (Gasterosteus aculeatus), nile tilapia (Oreochromis niloticus), great blue-spotted mudskipper (Boleophthalmus pectinirostris) and as contrast humans (Homo sapiens, grey). Potassium/chloride co-transporters (KCC) are used as outgroup.

Finally, fish produce osmolytes to actively take up and retain water. In particular, the cyclic polyol myo-inositol is used by euryhaline teleosts to acclimate to high salinity. Two enzymes are required for its production: myo-D inositol 3-phosphate synthase (MIPS) and inositol monophosphatase (IMPA). In addition, some fish actively accumulate myo-inositol with a sodium/myo-inositol cotransporter (SMIT) (74, 75). This transporter is of particular importance for marine fish exposed to high salt concentrations (76, 77), while freshwater fish lack a SMIT gene (e.g. the freshwater stenohaline zebrafish lacks the SMIT gene). The presence of SMIT has therefore been proposed to be a critical prerequisite for high salinity tolerance in fish (78). We find that the round goby genome contains MIPS and IMPA, and importantly, also a SMIT gene (**Supplemental_Fig_S7**).

### 6. Response to the environment: Immune System

It has been speculated that invasion success may relate to the ability to fight novel immune challenges (37). We therefore characterized key genes related to the immune system, focusing on genes that span both the innate and adaptive immune system such as pattern recognition receptors, selected cytokines and chemokines, antigen presentation, T-cell surface receptors and antibodies (**Supplemental_Table_S3; Supplemental_Table_S4**).

We find that the round goby genome features a classical adaptive immunity setup (Table 3). Vertebrate adaptive immunity is characterized by the Major Histocompatibility Complex (MHC) class I and MHC class II proteins and their regulators. MHCI presents antigens derived from a cell’s intracellular environment, while MHCII presents antigens derived from material engulfed by macrophages, B-cells or dendritic cells (79). We find 26 full length MHCI sequences from the classic U-lineage and one sequence from the teleost-specific Z-lineage (80) (**Supplemental_Table_S5**). MHCII is represented by 8 alpha (2 fragments) and 9 beta copies (**Supplemental_Table_S6)**. The uneven numbers may be attributed to assembly issues, but also, additional small fragments were not further investigated (data not shown). We also identify the key MHC-supporting peptides Beta-2-Microglobulin, *CD74*, *TAP1/2* and *tapasin*. Beta-2-Microglobulin (*B2M*) is present in two copies, one of which contains several indels, a diverged region, and no stop codon and thus may be a pseudogene. The round goby has two copies of *TAP2*, which promotes the delivery of peptides to MHCI (annotated as *TAP2* and *TAP2T;* **Supplemental_Table_S4; Supplemental_Fig_S8**). Two *TAP2* genes have also been described in zebrafish, and our results thus suggest this is conserved feature among teleosts (81). In addition, we identify the MHC transcriptional regulators *CIITA* and *NLRC5* (**Supplemental_Table_S3**). The presence of the thymus transcription factor *AIRE* and the T-cell receptors *CD4* and *CD8* confirms the presence of helper T cells and cytotoxic T cells in the round goby.

**Table 3.** Overview of manually annotated key adaptive immune genes

| Gene | NEME annotation | Contig annotation | Start | End | Strand |
| --- | --- | --- | --- | --- | --- |
| **CIITA** | NEME_493 | Contig_2585 | 3 985 719 | 3 993 128 | Antisense |
| **AICDA** | NEME_58 | Contig_447 | 597 424 | 599 014 | sense |
| **AIRE** | NEME_9 | Contig_79 | 14 106 230 | 14 113 573 | antisense |
| **B2M** | NEME_421 | Contig_2242 | 363 050 | 363 352 | antisense |
| **B2M_pseudo** | NEME_421 | Contig_2242 | 368 352 | 368 721 | antisense |
| **CD4** | NEME_213 | Contig_1334 | 340 445 | 348 248 | sense |
| **CD74** | NEME_71 | Contig_593 | 791 743 | 796 652 | antisense |
| **CD8a** | NEME_729 | Contig_3231 | 634 222 | 648 487 | antisense |
| **CD8b** | NEME_729 | Contig_3231 | 656 030 | 660 462 | antisense |
| **RAG1** | NEME_106 | Contig_787 | 4 690 414 | 4 695 142 | sense |
| **RAG2** | NEME_106 | Contig_787 | 4 699 042 | 4 700 651 | antisense |
| **TAP1** | NEME_582 | Contig_2864 | 694 776 | 722 339 | sense |
| **TAP2** | NEME_387 | Contig_2107 | 2 987 106 | 2 993 287 | antisense |
| **TAP2T** | NEME_299 | Contig_1786 | 3 697 645 | 3 704 089 | sense |
| **Tapasin** | NEME_387 | Contig_2107 | 3 111 989 | 3 119 308 | sense |

Similarly, the humoral adaptive immune response (also termed the B-cell mediated immune response) is intact in the round goby. Humoral immunity in fish is characterized by three antibody isotypes consisting of immunoglobulin heavy chains delta (IgD), mu (IgM), and tau (IgT). We identify a contig-spanning immunoglobulin heavy chain locus (**Supplemental_Fig_S9**) containing 8 delta constant domains, and 4 constant mu domains, as well as genes responsible for heavy chain recombination and immunoglobulin hypermutation (*RAG1/2* and *AID(AICDA)*; Table 3; **Supplemental_Table_S3**). There is no evidence for the presence of immunoglobulin tau constant domains, which are commonly found in carps and salmonids (82).

While round goby adaptive immunity conforms to vertebrate standards, its innate immune repertoire displays remarkable and unusual features. We find that all components of the inflammasome (a signaling pathway involved in inflammatory responses; **Figure 8**) are expanded. Inflammasome assembly is activated through pathogen pattern recognition receptors (83), and ultimately triggers a local or systemic acute phase response by producing IL-1 family cytokines (83, 84) and/or promotes cell death via pyroptosis (84). In the round goby genome, components of the entire cascade (pattern recognition receptors, ASC adaptor proteins, IL-1, and acute phase proteins) are present in unexpectedly large numbers (**Figure 8; Supplemental_Table_S8**). In the following, our findings are described step-by-step from the cell surface down to the acute phase response.

**Figure 8.**
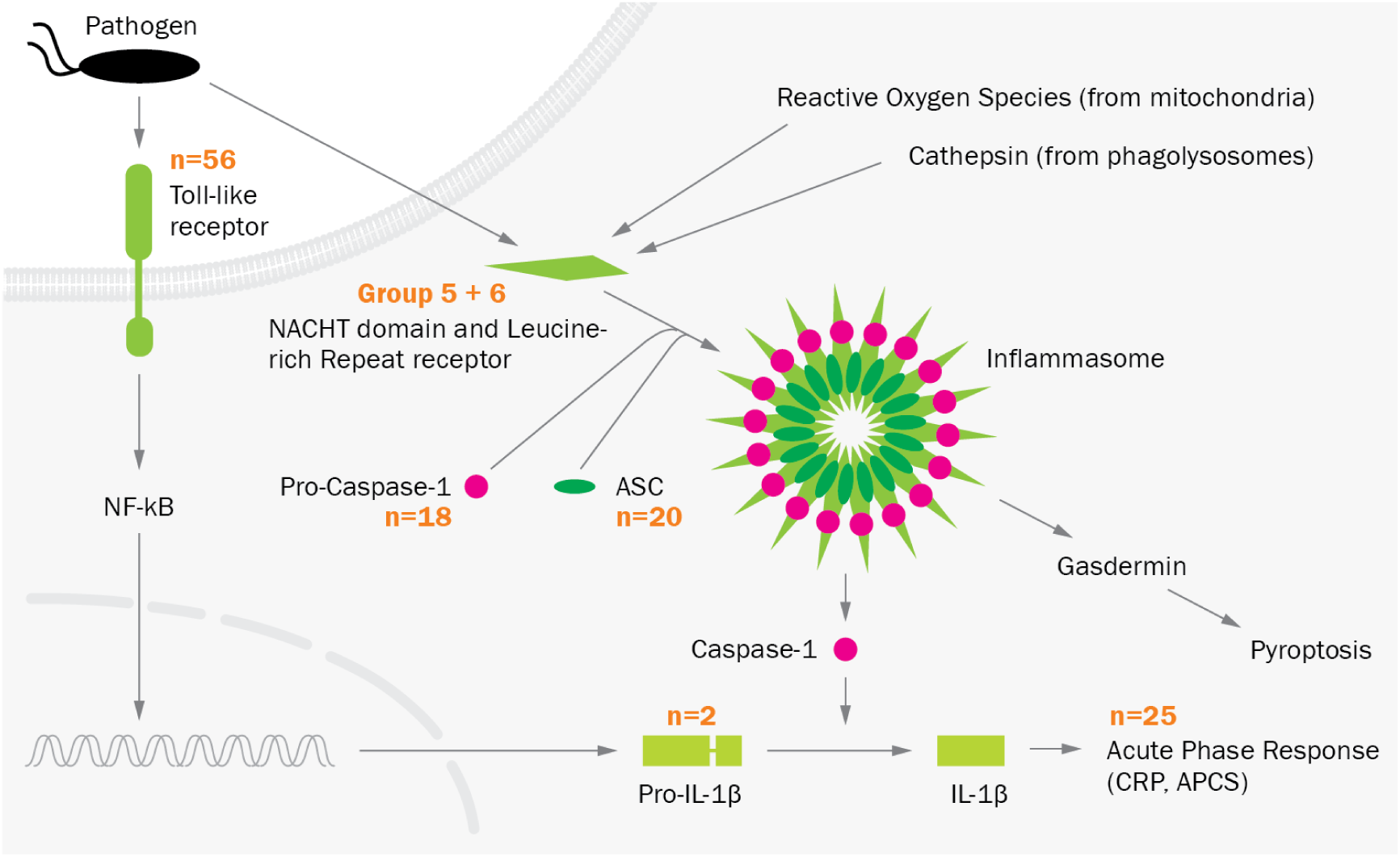
The inflammasome pathway. Several components of the pathway are expanded in the round goby (gene numbers in round goby, or novel groups for NLRs, are indicated in orange). Pathogen-associated patterns are recognized by pattern recognition receptors such as Toll-like receptors at the cell surface, or NLRs in the cytoplasm. This interaction triggers the transcription of cytokine precursors via NFkB, and the activation and assembly of inflammasome components (NLRs, Pro-Caspase-1, and ASC). Inflammasome-activated Caspase-1 then initiates the maturation of cytokines and an acute phase inflammatory response (CRP, APCS proteins), and / or pyroptosis through gasdermin.

Perhaps the best studied pattern recognition receptors are the Toll-like Receptors (TLRs), pathogen-recognizing molecules that are generally expressed either at the plasma membrane or on the endosomal membranes. Sixteen TLR types with slightly differing ligand binding activities are conserved across vertebrates, and most vertebrate genomes contain one to three copies of each type. As expected for a teleost, the round goby genome does not contain the LPS-detecting TLR4 genes. However, in total we find 56 TLRs, of which 40 appear to originate from an expansion of Toll-Like Receptor 23-like genes (**Figure 9**). Small expansions of specific *TLRs* are somewhat common in fish (85), and indeed, we find minor *TLR22* and *TLR23* expansions to 6-13 copies in the genomes of other *Gobiidae*. However, the extent of the expansion of *TLR23* exceeds even what is observed for *TLR22* in the relatives of cod (*Gadiformes*) (86). Phylogenetically, the identified TLR23 sequences form three clades, of which two are specific to *Gobiidae*, while the third contains TLR23 sequences from other teleosts as well (**Supplemental_Fig_S10**). In terms of genomic location, round goby TLRs 22 and 23 were distributed across several contigs with some copies arranged in tandem, which suggests several independent duplication events.

**Figure 9.**
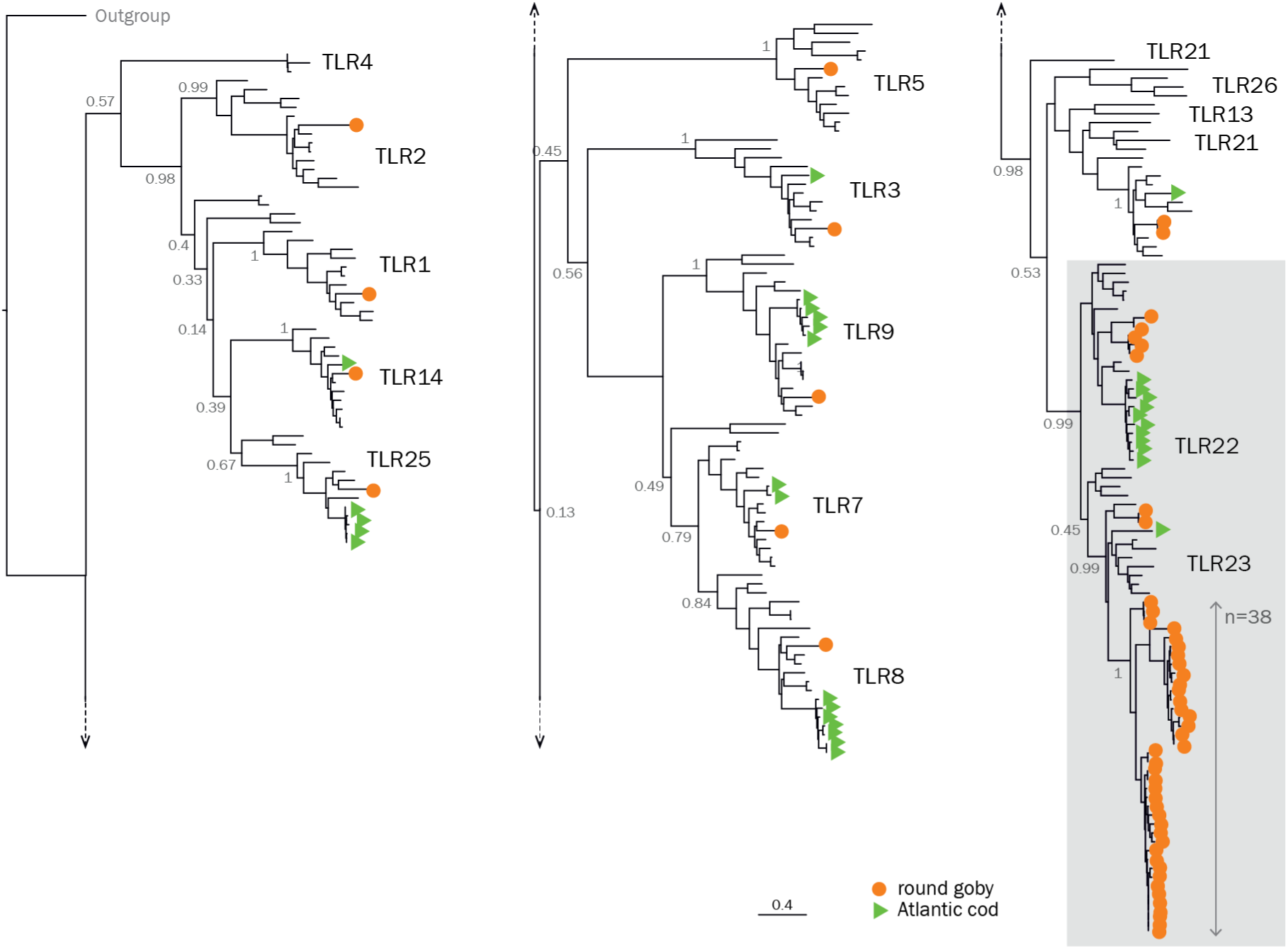
Phylogenetic tree of teleost Toll Like Receptor protein sequences. A maximum likelihood phylogenetic tree run with the JTT substitution model and 500 bootstrap replicates on the transmembrane, linker and TIR domain of all TLRs found in a selected set of teleosts in the Ensembl database, the Atlantic cod genome version 2, and all manually investigated Gobiiformes. A TLR sequence from the lancelet Branchiostoma belcheri was used as an outgroup and the root was placed upon its corresponding branch. Green triangles, Atlantic cod. Orange circles, round goby. Grey box, TLR22 and TLR23.

For intracellular pathogen recognition receptors of the NACHT domain and Leucine-rich Repeat containing receptor (NLR) family, we identify two new, previously undescribed families (Group 5 and 6) present in the round goby and also in the mudskipper *B. pectinirostris* (**Figure 10**).

**Figure 10.**
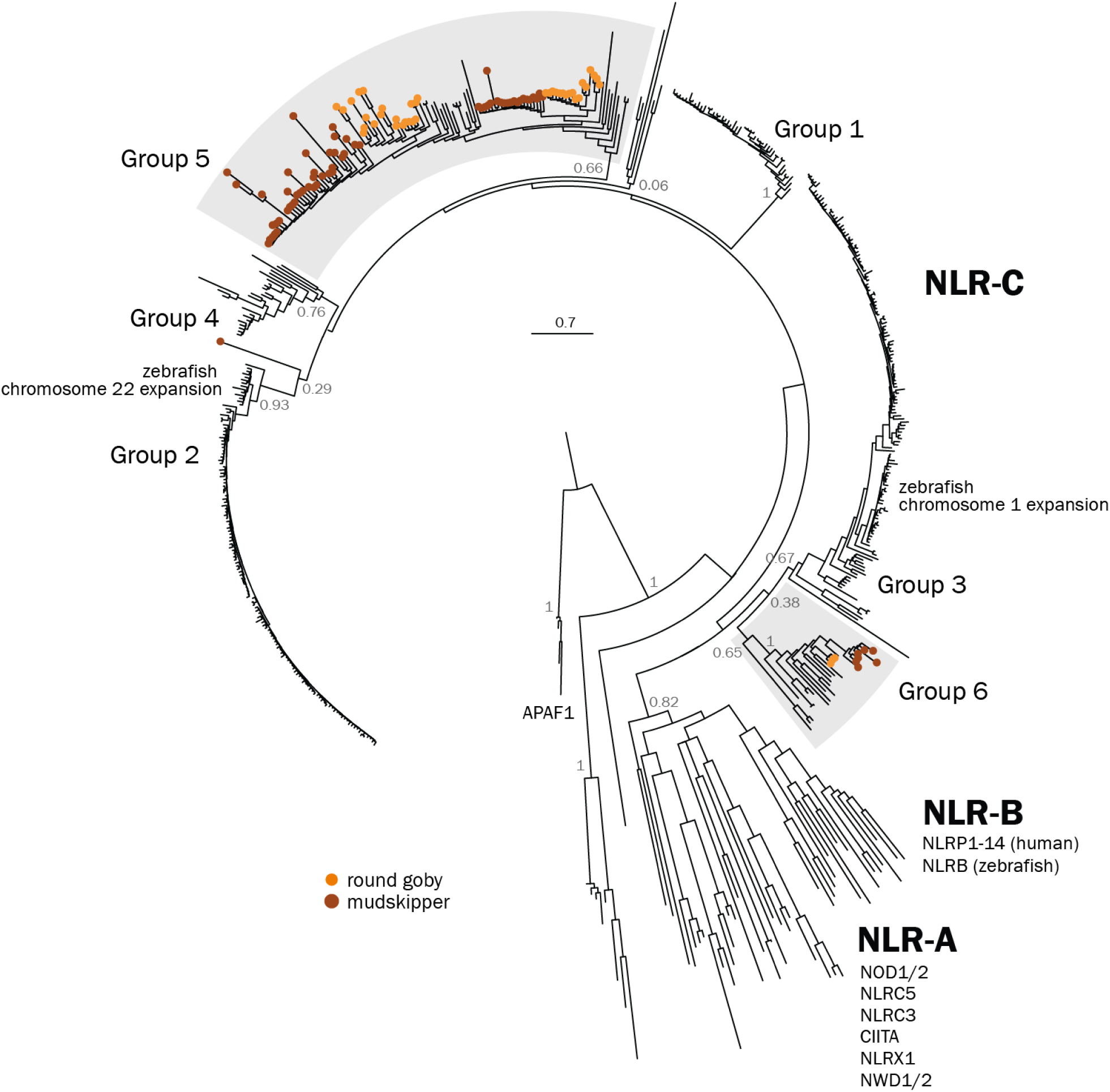
Phylogenetic tree of the NLR nucleotide-binding domain sequences in round goby, great blue dotted mudskipper, zebrafish and human. Maximum Likelihood phylogenetic tree with 500 bootstraps rooted at the split between NB-ARC (found in APAF) and NACHT domains (present in all the other NLRs). NB-ARC domains from APAF1 orthologs were used as an outgroup. Bootstrap values are shown for nodes that determine an entire cluster. The tree resolves all three major classes of vertebrate NLRs (NLR-A, NLR-B, NLR-C). NLR-A genes were conserved in all analyzed species, no NLR-B genes were found from the gobies. Six groups of NLR-C genes were identified, four of which are exclusive to zebrafish (Group 1-4) and two contain only sequences from gobies (Groups 5 and 6). Within the goby-specific groups, lineage-specific expansions can be seen for both round goby (orange) and mudskipper (brown).

NLRs have diverse roles from direct pathogen recognition to transcriptional regulation of the MHC (NLRs CIITA and NLRC5) and contribute to inflammasome activation. Mammalian genomes display 20-40 NLRs in families NLR-A and NLR-B, while fish also feature a fish-specific subfamily (NLR-C) (87) and a much expanded NLR repertoire (e.g. 405 NLR-C genes in zebrafish) (88, 89). The round goby genome contains at least 353 NLRs (**Supplemental_Table_S8**), which include 9 highly conserved vertebrate NLRs (*NOD1*, *NOD2*, *NLRC3*, *NLRC5*, *NLRX1*, *NWD1*, *NWD2*, *APAF1*, *CIITA*) as well as 344 NLR-C genes. Fish NLRs cluster into 6 groups of which 2 represent novel NLR-C clades (groups 5 and 6, **Figure 10**). The novel groups are supported by phylogenetic analyses as well as motif presence/absence (Table 4). NLR-C groups are characterized by highly conserved versions of the sequence motif Walker A. The most common sequence for Walker A observed in both goby NLR-C groups, GVAGVGKT, is not associated with any of the four major NLR-C groups in zebrafish (88). Also, NLR subtypes often carry group-specific combinations of the protein-protein interaction domain PYD and/or B30.2 domain. This holds true for *Gobiidae* NLR-C groups, since only group 5 NLRs can carry an N-terminal PYD domain and/or a C-terminal B30.2 domain (88), similar to the zebrafish Group 1 and 2 NLRs (Table 4). In contrast, some group 6 NLRs have C-terminal CARD domains, which in both human and zebrafish are attached to specific inflammasome-associated NLR-B genes (90). The round goby C-terminal CARD-containing NLRs are found on the same few scaffolds and share a high degree of sequence similarity, indicative of a recent expansion. This expansion is absent from mudskipper and thus restricted to the round goby lineage. Many other Group 6 NLRs are fragmented, with large insertions in the middle of their conserved 2 kb exon (**Supplemental_Table_S8**).

**Table 4.**
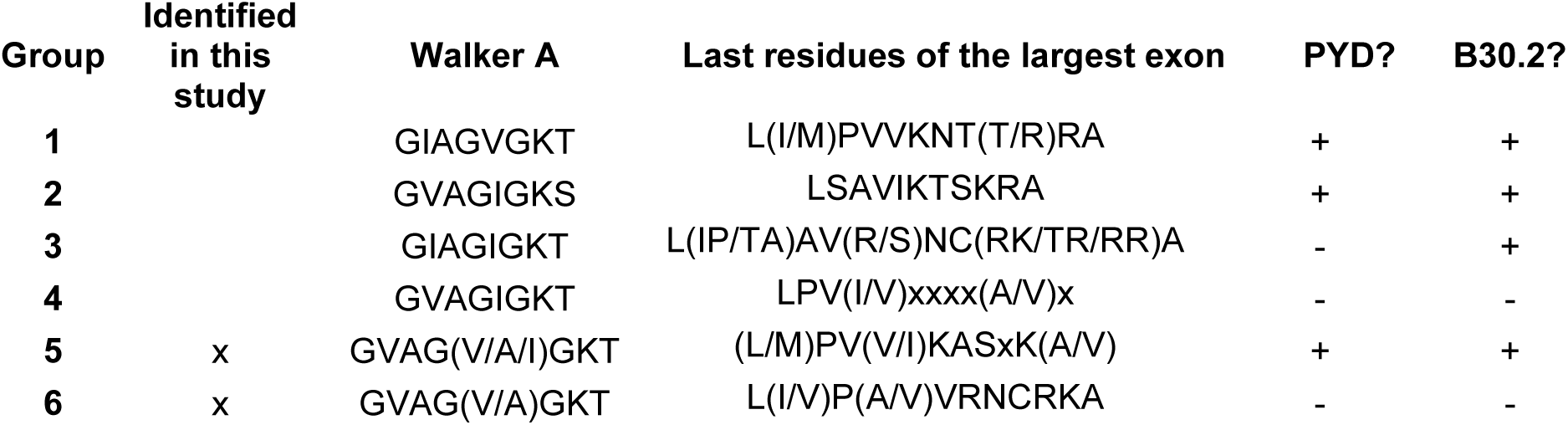
Key features of each of the six NLR-C subgroups. x denotes a variable amino acid.

| Group | Identified<br>in this<br>study | Walker A | Last residues of the largest exon | PYD? | B30.2? |
| --- | --- | --- | --- | --- | --- |
| 1 |  | GIAGVGKT | L(I/M)PVVKNT(T/R)RA | + | + |
| 2 |  | GVAGIGKS | LSAVIKTSKRA | + | + |
| 3 |  | GIAGIGKT | L(IP/TA)AV(R/S)NC(RK/TR/RR)A | - | + |
| 4 |  | GVAGIGKT | LPV(I/V)xxxx(A/V)x | - | - |
| 5 | x | GVAG(V/A/I)GKT | (L/M)PV(V/I)KASxK(A/V) | + | + |
| 6 | x | GVAG(V/A)GKT | L(I/V)P(A/V)VRNCRKA | - | - |

Once activated, some NLRs (including those with a C-terminal CARD) can oligomerize and form an inflammasome in order to activate specific caspases (usually Caspase 1, **Figure 8**). The interaction between NLRs and the caspase are mediated by the adaptor protein ASC (also known as PYCARD), which itself oligomerizes into large structures known as “specks” (91). Vertebrates have 1-2 copies of ASC, which are characterized by a characteristic combination of a single PYD and CARD domain. In the round goby genome, we find 20 cases of this domain combination. Since the genomes of other gobies contain 1-2 PYD-ASC combinations, the expansion appears to be specific to the round goby (Figure 11A). The effector protein Caspase 1 is present as one gene in humans and as two genes in zebrafish. We find that the round goby genome features an expansion to 18 copies. Interestingly, some of those genes appear to contain a CARD domain (as seen in mammals and several species of fish) while others have PYD (as seen in zebrafish). This suggests that a caspase with both domains may have existed in the common ancestor of fish and tetrapods, with most lineages having retained only one of the two. However, phylogenetic analyses reveal that all round goby Caspase 1 genes are the result of a single expansion event specific to this species (Figure 11B). An alternative explanation for the presence of both PYD- and CARD-caspases 1 genes would be a recurrent acquisition of PYD in different lineages. In any case, in addition to Caspase 1 genes, caspase 3 (a key component of apoptosis which may be activated by Caspase 1) is also expanded to 5 copies. Caspase 4 and 5, on the other hand, appear to be absent.

**Figure 11.**
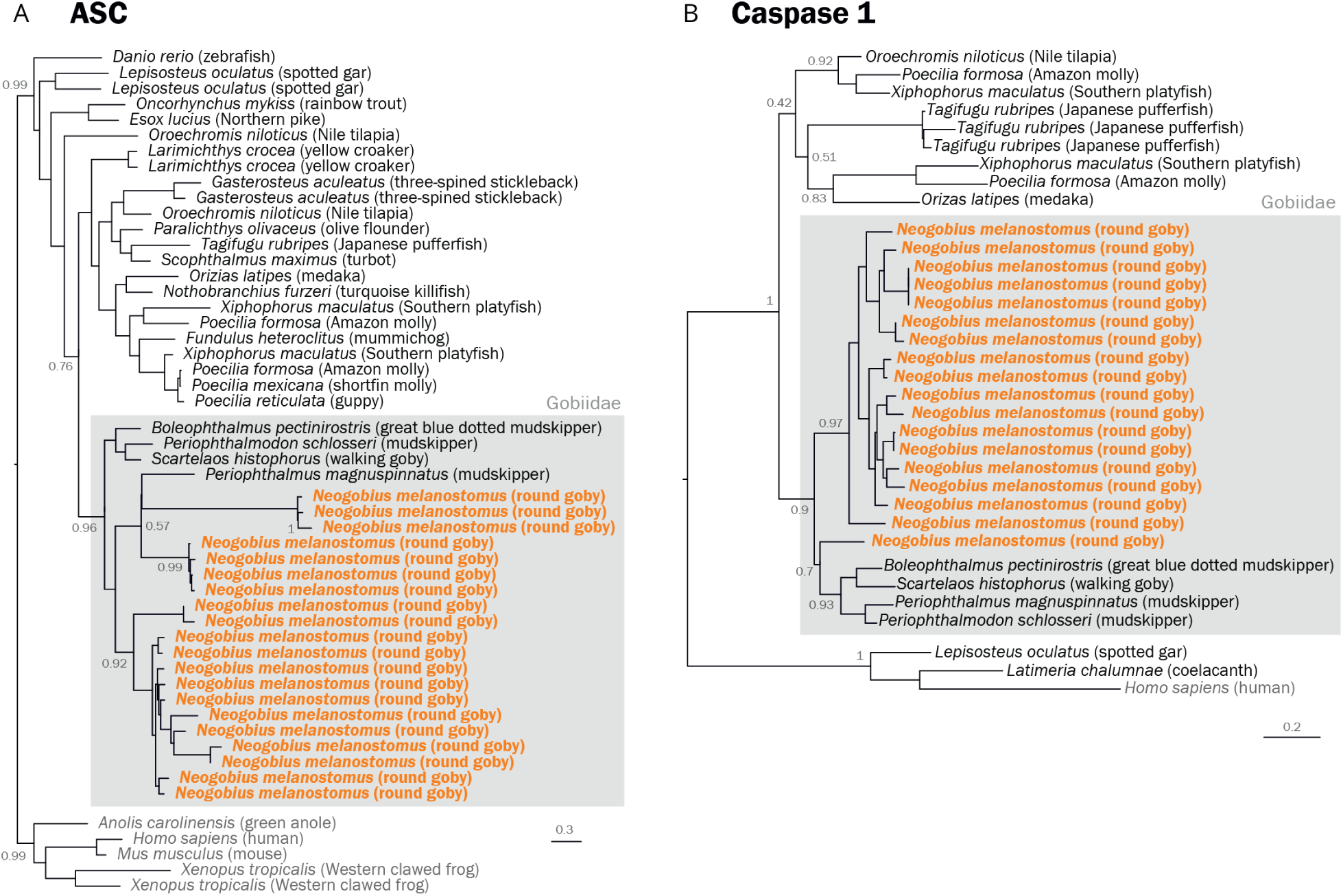
**A Phylogenetic tree of gnathostome ASC protein sequences**. Maximum Likelihood phylogenetic tree with 500 bootstraps rooted at the split between tetrapods and ray-finned fish. Tetrapods were used as outgroup. Round goby is indicated in orange. Gobiidae are highlighted with a grey box. The goby sequences form a clear separate cluster (marked with the box), with a large expansion apparent in the round goby. **B Phylogenetic tree of gnathostome Caspase 1 protein sequences** The Caspase 1 tree comprises all protein sequences annotated as CASP1 in the investigated Gobiiformes genomes aligned together with reference sequences from Ensembl and GenBank. The root was placed on the branch containing the mammalian sequences.

Finally, we find that genes encoding for two peptides produced in the course of inflammation, the acute phase reactants C-reactive protein (CRP) and serum amyloid component P (APCS), are expanded to a total of 25 copies (compared to 2-7 in fish, and 5-19 in the other *Gobiidae*). In fish, CRP and APCS are closely related and cannot be distinguished based on BLAST scores or phylogeny. As seen in other fish species, all investigated CRP/APCS sequences resolve into two major phylogenetic clades, with the mammalian sequences in a third (**Supplemental_Figure_S11**).

### 7. Adaptation to novel environments: Epigenetic regulators

The PRC2 complex establishes and maintains gene repression (92) and thus represents a plasticity-restricting mechanism. The complex mediates di- and trimethylation of lysine 27 on histone H3 and contains four proteins: a catalytic subunit (either *enhancer of zeste* EZH1 or EZH2), *suppressor of zeste* SUZ12, *embryonic ectoderm development* EED and *RB Binding Protein 4* RBBP4 (50). In mammals, the alternative catalytic subunits EZH1 and EZH2 have partially complementary roles (93, 94), and requirements for the two alternative catalytic subunits differ between species – in contrast to mammals, zebrafish develop in the absence of either catalytic subunit (95, 96). We find that the round goby genome contains the usual complement of PRC2 components: two copies of SUZ12 (of which one appears quite diverged), one copy of EED, one copy of RBBP4, and two copies of EZH (with multiple isoforms determined by RACE experiments). For SUZ12, EED, and RBBP4, sequence-based identification was straightforward, and phylogenetic analyses followed the known phylogenetic relationships of fish, mammals, and other vertebrates (**Supplemental_Fig_S12**). The catalytically active subunits EZH1 and EZH2 do cluster with the closest species in the phylogeny, the mudskipper *B. pectinirostris* (**Figure 12**), but the deeper relationships within EZH2 are poorly supported and may suggest a complex evolutionary history.

**Figure 12.**
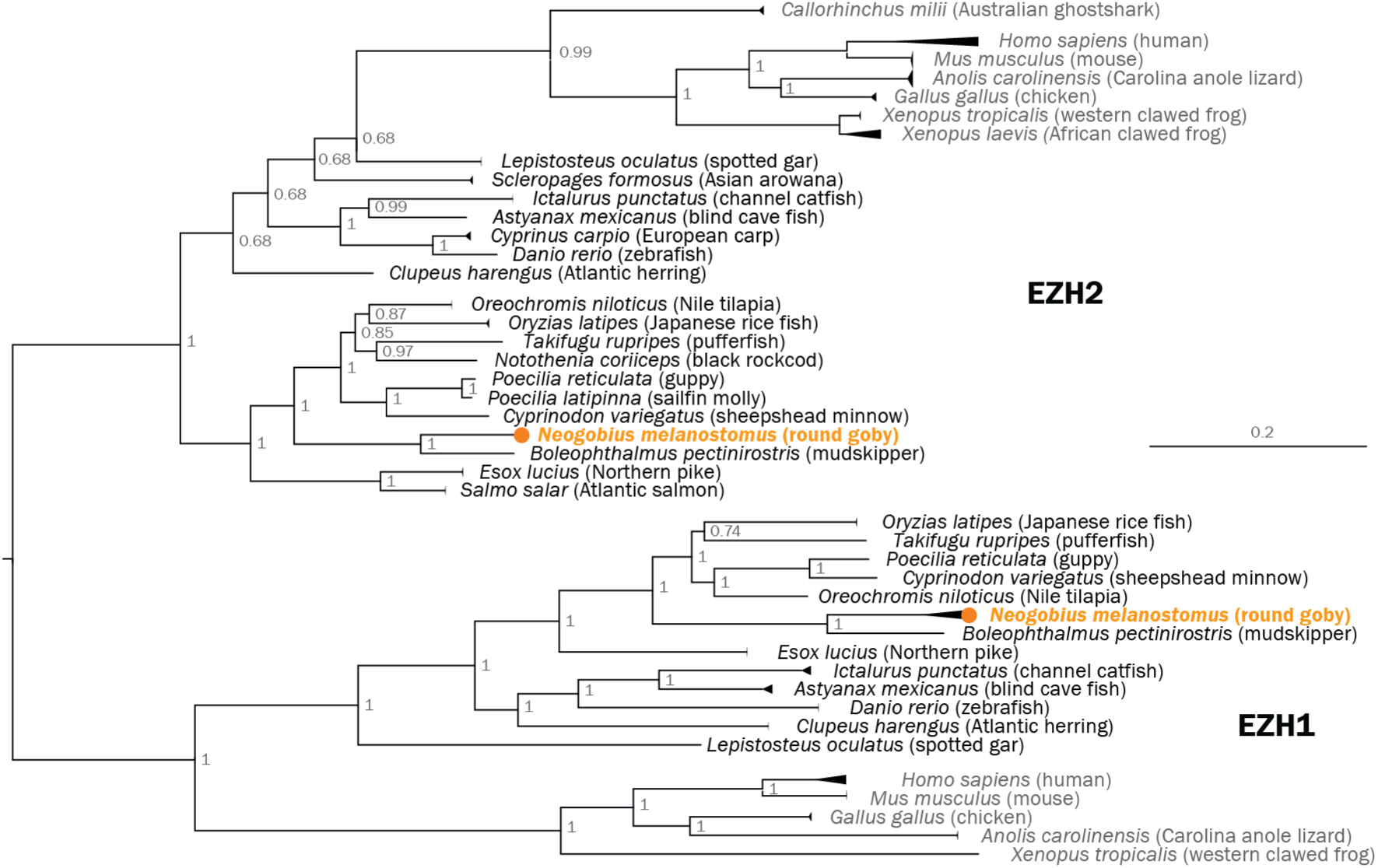
Phylogenetic tree of vertebrate EZH proteins. Midpoint-rooted Bayesian phylogenetic tree. Note the position of the Australian ghost shark (potential outgroup) within the poorly supported EZH2 branch. When rooting with Australian ghost shark, teleost EZH2 genes cluster with EZH1 (data not shown). Round goby is indicated in orange.

Methylation marks similarly regulate gene expression and are deposited by conserved enzymes called DNA methyltransferases (DNMTs). Mammals feature two types of DNMTs, DNMT3 (three genes A, B, and L) and DNMT1 (one gene). The two types perform both *de novo* and maintenance methylation, respectively, in a dynamic division of labor (97). Interestingly, fish feature a variable repertoire of DNMT3 genes. Medaka, fugu, zebrafish, and carp have three, five, six, and twelve DNMT3 genes, respectively (98). We find that the round goby genome features one DNMT1 that follows the expected phylogenies, and five DNMT3 genes, of which two cluster with vertebrate DNMT3A sequences, and three with vertebrate DNMT3B sequences (**Figure 13**). The number of DNMT3 genes in round goby corresponds to that seen in in stickleback, fugu and tilapia (99). In general, the DNMT3 phylogeny is not well supported, which limits conclusions about the evolution of specific DNMT3 genes.

**Figure 13.**
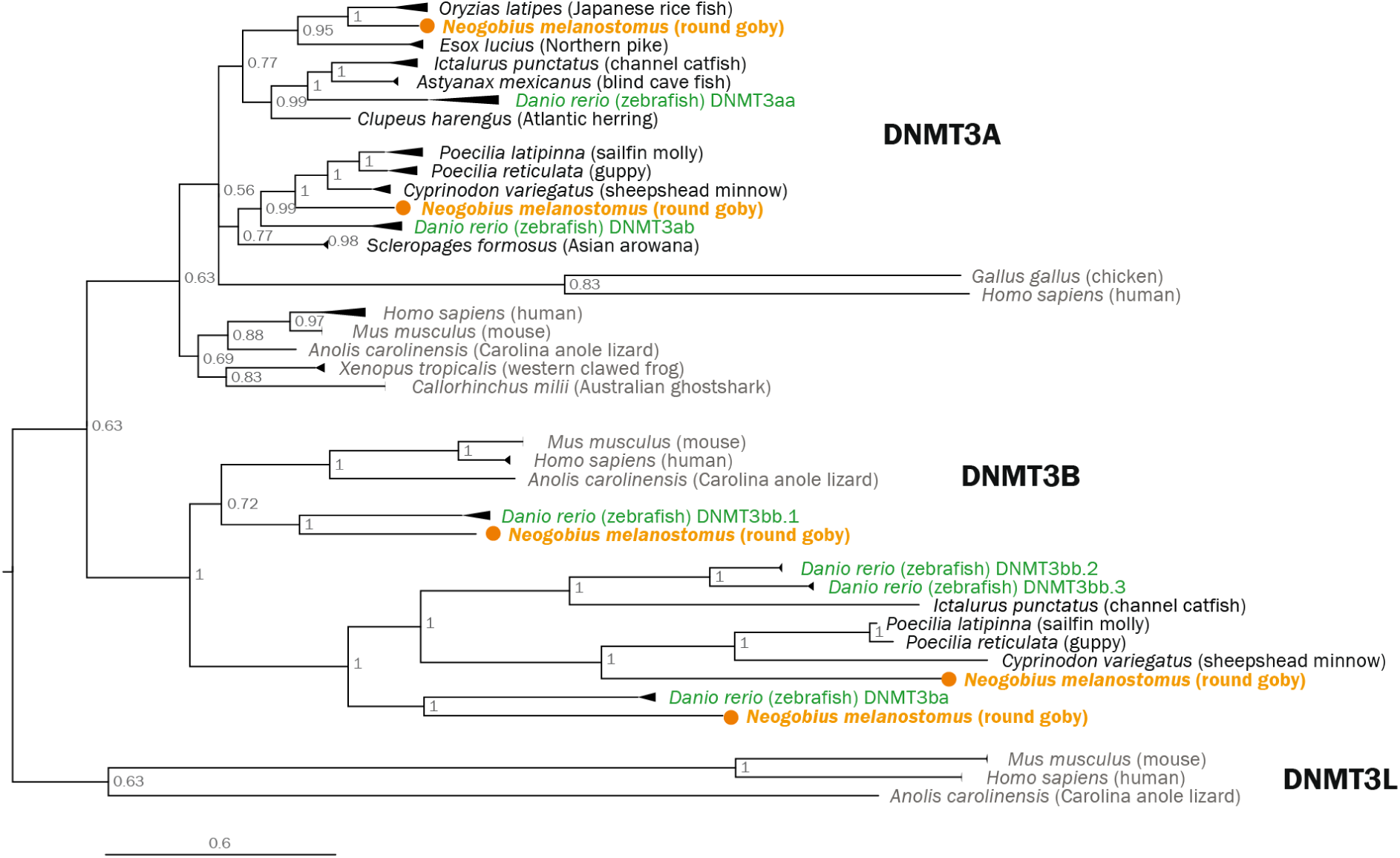
Phylogenetic tree of vertebrate DNMT3 proteins. Midpoint-rooted Bayesian phylogenetic tree. The Australian ghost shark (potential outgroup) is positioned among DNMT3A genes. Round goby is indicated in orange. Zebrafish, the only other fish with well-annotated DNMT3 genes, is indicated in green.

## Discussion

### General observations

Our analyses depict a genome that, in many respects, is similar to other teleost genomes. There is no evidence for recent genome duplications, and genome size, gene content and GC content are within the ordinary range. Transposable elements can create genetic variation and mediate ecological success (100), but repeat analyses do not reveal unusual transposon activities in the round goby. Small genome size has been proposed to foster invasiveness (101), but the round goby genome is not particularly small. Phylogenetic analyses reveal that many of the analyzed gene families conform to expectations. For example, green opsin gene duplications and the loss of the UV opsin are observed in many fish lineages (20). Similarly, the expected gene families and overall gene complements are found for olfactory receptors, cytochrome P450, and osmoregulatory proteins, for adaptive immunity and epigenetic regulators. Multilocus sex determination has previously been suggested for freshwater gobies (102), and indeed our data suggest a multigenic and/or environmental sex determination system for the round goby, rather than a large sex-determining region or a sex chromosome. Overall, these findings support the validity of the sequencing and assembly approach, and suggest that selected findings of interest are not artefacts. In addition, the round goby genome sequence also reveals several novel and interesting findings of which some pertain to teleost genomes in general, some to *Gobiidae*, and some to specific gene families, with possible implications for invasive potential.

### Environmental perception

We find that the visual system of *Gobiidae* may be more efficient in the red parts of the light spectrum. This is intriguing considering the benthic life style of gobies and their occurrence in turbid areas. In clear waters, red light from the sun is the least abundant part of the spectrum (and virtually absent below 15m of depth) because red light penetrates least through water, but many organisms convert the deeply penetrating green and blue wavelengths into red. Indeed, the eyes of gammarids, a common prey of round goby, strongly reflect red light (103). An enhanced red perception through an additional red opsin gene may thus be relevant for round goby predation success. In turbid waters, red is the most common part of the light spectrum because long wavelengths experience least scattering (21). Round gobies readily establish populations in turbid environments. The retention of two red opsin genes may thus possibly relate to the remarkable ability of the round goby to colonize turbid habitats. Our predictions based on the key amino-acid substitution suggest that LWS1 is expected to be most sensitive at 560 nm (same as one of the mudskipper gobies) (25), while LWS2 is expected to be most sensitive at 550 nm (59). Similar small differences in the sensitivity maximum can indeed result in functionally different spectral tuning palettes (e.g. during development or in different environmental conditions) (104).

The presence of red fluorescence on top of the eye in round goby is the first unequivocal description of fluorescence in a freshwater fish and might be interpreted as being associated with the ability to discriminate different shades of red colors. However, the fluorescence in the specimens investigated was quite weak. Unless fluorescence expression is stronger under natural conditions or in the ancestral population from which the invading populations stem, a visual function of the weak fluorescence observed here seems unlikely (see warnings by (105)). Fluorescence is, however, widespread and stronger among several marine gobies (106). Although the fluorescent “eyebrows” of the round goby show a striking similarity to those of some marine gobies, their function will remain unclear until properly tested. Social functions are possible – for example, in sand gobies, dark eyes indicate female readiness to spawn (107). Alternatively, they may simply provide camouflage for individuals buried in bright sand up to the eyes. Functional hypotheses for fluorescence, such as communication, camouflage and improved prey detection have been extensively reviewed by (108). The genetic tools now available for the round goby may allow for experimental manipulation of fluorescence expression, once the actual fluorophores that produce the fluorescent signal have been identified.

### Response to the environment

With respect to ecological and physiological aspects of success and invasiveness, some findings on CYP genes, on osmoregulation, and on innate immunity call for further attention. The mostly minimal complement of cytochrome P450 proteins present in the round goby is unexpected considering the occurrence of round goby in polluted areas (109, 110). The CYP1-3 gene complement for xenobiotic metabolism is similar to other teleost genomes, and the ability of the round goby to survive in contaminated environments must therefore have other reasons. Round goby may cope with contaminations at the level of gene expression, either through higher basal expression values or by a particularly rapid response to exposure (45). Alternatively, this species may have peculiarities in other, not yet analyzed areas of the defensome (e.g. transporters). Analyses of the tissue expression of CYP families 1, 2 and 3, and also the study of other defensome gene families, including the nuclear receptors regulating CYP gene expression, transporters and conjugating enzyme families, may be useful in this respect.

Another potentially relevant finding is the ability of the round goby to not only produce, but also accumulate osmolytes. Species distribution constraints often arise from physiological limitations. The round goby is one of the most geographically wide-ranging invasive fish species in Europe and North America, and the ability to accumulate osmolytes may impact its range expansion in three ways. Firstly, 0-25 PSU (common for coastal waters, but lower than the ocean) is the species’ current limit for unperturbed osmoregulation (111). However, the round goby’s repertoire of key-genes in myo-inositol production and accumulation might bestow the species with the potential to eventually tolerate higher salinities, for example through the evolution of altered gene regulation patterns, and colonize higher PSUs. Secondly, osmolytes improve water retention and thus desiccation tolerance. In this context, myo-inositol accumulation may have contributed to overland dispersal. Overland dispersal of eggs or larvae with boats or fishing gear involves air exposure, and indeed, round goby eggs withstand desiccation for up to 48 hours (112). Finally, osmolytes essentially act as anti-freeze agents and molecular chaperones, and contribute to cryoprotection in diverse organisms from bacteria (113) to flies (114). Osmolytes may thus enable the round goby to combat a number of environmental conditions and to colonize new areas. The surprising and unexpected ability of the round goby to colonize cold areas well below its temperature optimum of 22°C, such as the Northern Baltic Sea, may be linked to osmolyte production.

Lastly, the “strike fast, strike hard” innate immune system and the impressively large inflammation machinery of the round goby may enhance the species’ colonization potential. Fish immunity appears to be quite plastic. For example, cod have disposed of some core adaptive immunity components (18), yellow croaker feature an expanded TNF repertoire (115), and channel catfish retain a high number of recent duplications and SNPs in immune genes (116). Meanwhile, in salmonids, genes specifically retained after the 4^th^ whole genome duplication are not immune genes (117).

We find that the round goby genome contains multiple copies of genes for inflammasome assembly, activation, and function. This is interesting because the fish inflammasome complex is much more poorly characterized than that of mammals. Maturation of IL-1 by inflammasome-activated Caspase 1 cleavage in fish is a matter of debate, since teleost IL-1 proteins lack the conserved caspase cleavage site present in mammalian IL-1b and IL-18 (118). However, as has been shown for zebrafish, Caspase 1 can also utilize an alternative site to cleave and mature IL-1 (90, 119). In any case, the caspases also mediate cell death via pyroptosis and the presence of other components such as ASC, caspases and pro-IL1 and pro-IL18 supports a role for inflammasomes in fish. Zebrafish ASC oligomerize and form “specks” as seen in mammals (90). The molecular dynamics of inflammasome activation therefore represent a potential future research avenue in the round goby.

In terms of ecological success, the round goby’s expanded repertoire of pathogen recognition receptors may broaden the scope of its immune response and increase the range of detectable ligands and pathogens. The expanded acute phase repertoire may also contribute to a fast response, or inversely, may limit excessive cell damage. In humans, the acute phase protein CRP contains inflammation as part of a negative feedback loop (120). Thus, the round goby may re-enter homeostasis faster compared to other fish species with smaller CRP/APCS repertoires. The larger acute phase repertoire may also function to limit the cellular damage caused by the potentially large amount of inflammasome combinations the round goby can generate. In this context, we suggest systematic investigations into a potential relation between inflammasome expansions and invasiveness in *Gobiidae*, in combination with immune challenge experiments.

### Long-term adaptation

We identify a potentially interesting evolutionary history for the conserved PRC2 component EZH in fish, and add to the previous observation that the conserved *de novo* DNA methylation machinery features a surprising diversity in fish. These results underscore the need for in-depth investigations into the role and relevance of epigenetic regulation and transgenerational inheritance in teleosts. Our findings support the emerging idea that epigenetic regulation in fish follows somewhat different rules than in mammals. For the histone methylating complex PRC2, our results suggest interesting phylogenetic relationships of EZH proteins in fish. EZH proteins act in tissue specific complexes comprised of core SUZ12, EED, and RBBP4, but also AEBP2, PCL proteins and JARID2. These proteins enhance PRC2 efficiency, contribute to recruitment to target sites, or inhibit the complex (50, 92). Small sequence changes can have strong effects on the entire complex, since the precise interactions among the components and with other gene regulators impact its function and localization (121–124). For example, species-specific insertions (125) are thought to regulate PRC2 recruitment and/or exclusion from target genes (126). We suggest that the future incorporation of more sequences of both EZH1 and EZH2 from a greater range of taxa and the inclusion of currently unannotated versions of the genes associated with both the teleost specific whole genome duplication and lineage specific duplications (96) would aid understanding of the evolutionary history of the entire complex. We expect that studying PRC2 in non-mammalian vertebrates may reveal ancestral or less abundant interactions, functions or also complex associations of PRC2.

Similarly, our results warrant an in-depth exploration of DNA methylation in fish. Originally, DNA methylation evolved to distinguish own (methylated) DNA from foreign (non-methylated) DNA such as introduced by viruses. Therefore, cytosines in CG base contexts are by default methylated. In mammals, DNA methylation in CG dense regions (CG islands) is associated with gene repression. However, DNA methylation also features species- and taxon-specific differences, even among vertebrates, which are still greatly underappreciated. For example, non-methylated genome regions in fish are unexpectedly CG-poor (127), fish differ from mammals with respect to the distribution of methylated CpGs in the genome (128), algorithms developed on mammals fail to identify CpG islands in fish (129), genome-wide CpG island predictions in cold-blooded animals consist primarily of false positives (130), and fish CG methylation occurs mainly in coding regions, where it correlates positively with gene expression levels (131). These curious differences are further enhanced by the seemingly random copy number variation in the *de novo* DNA methyltransferase DNMT3 in teleosts, which do not reflect genome duplication events in teleosts (99). DNMT3 genes display highly spatiotemporal expression patterns particularly during development (132–135), and an in-depth and species-aware exploration of the role of DNA methylation in fish is clearly warranted.

### Gene expansions

A general theme across several of the analyzed gene families is gene expansions. Gene expansions have been linked to invasive potential before (136) and are recurrent in fish genomes, both within (117, 137) and outside (116, 138, 139) the context of whole genome duplications. Many duplicated genes are known to experience rapid neofunctionalization rather than subfunctionalization (137), and have the potential to compensate against mutation even after divergence (140). The round goby and its relatives are definitely strong candidates for further and systematic investigation of a link between gene expansions and colonization or invasion potential. The *Benthophilinae* group is recently diversified crowd of fish with many members inexplicably on the move (141), and *Gobiidae* in general share a remarkable colonization potential (10, 142). Importantly, recent gene expansions can be difficult to resolve with short reads, and genomes based on long read sequencing (as presented here) will be instrumental in this regard.

Among the receptor families analyzed, the NLRs, TLRs, and olfactory receptors, we identify a couple of particularly beautiful case studies for recent expansions and repeated radiations. Our identification of two previously undescribed NLR-C gene families (88), here termed group 5 and group 6, indicates substantial diversification of NLRs in fish. Different teleost lineages appear to feature different NLR-C subfamilies with large lineage-specific expansions reminiscent of olfactory receptor repertoires. Similarly, we identify interesting cases of parallel expansions across families, and also family-specific expansions, among olfactory receptors. Both cases warrant investigations into the evolution of ligand binding repertoires. For example, 7tm1 subfamily members may be involved in the detection of distinctive types of odors relevant for round goby, and possibly, *Gobiidae* ecology (28–30). Which types of odorants are detected by parallel expanded ORs, and whether these expansions serve to detect similar or different types of odorant molecules in different species, remains to be studied. Finally, the massively expanded TLR22 and TLR23 families warrant an exploration of their ligand binding properties. TLR22 and TLR23 have been suggested to recognize nucleic acid ligands (85), but some also react to protein or lipid pathogen-associated patterns (143–145), and their role in fish is currently unclear.

In summary, this work provides a solid basis for future research on the genomic, genetic, and epigenetic basis of ecological success. Clearly, many more gene families or pathways may contribute to the round goby’s invasion success. For example, the presented analyses barely scratch the surface of epigenetic regulation, innate immunity and transporters (e.g. of toxins). We did not investigate endocrine pathways (which govern growth and reproductive success) nor antimicrobial peptides (which contribute to innate immune defense), areas which may yield fruitful information of the success of this invader. We welcome future research using this novel genomic resource, and encourage experts on those pathways to contribute their knowledge.

## Methods

A relevant note upfront is that this manuscript is the product of a long-standing collaboration of leading experts in their respective fields. The gene families analyzed differ widely with regard to sequence conservation, the number and similarity of genes within and between species, the scope of questions in the field, etc. Compare, for example, the *de novo* identification of hundreds of virtually identical NLR receptors with the manual annotation of a handful of extremely conserved DNA methyltransferases, or the phylogenetic analysis of the conserved vertebrate CYP gene family with a fish-centered comparison of osmotic balance regulators. Accordingly, each collaborator applied methods that were suited for the respective situation. As a common theme, genes were identified by blast, sequences were extracted and aligned with other fish and/or other vertebrates, trees were constructed with either bayesian or maximum-likelihood methods, and findings were always verified against the mudskipper genomes.

### Genomic DNA library preparation and PacBio sequencing

Genomic DNA was extracted from the liver of one male individual of round goby caught in Basel, Switzerland (47° 35′ 18″ N, 7° 35′ 26″ E). At the Genome Center Dresden, Germany, 300 mg of liver tissue were ground by mortar and pestle in liquid nitrogen and lysed in Qiagen G2 lysis buffer with Proteinase K. RNA was digested by RNase A treatment. Proteins and fat were removed with two cycles of phenol-chloroform extraction and two cycles of chloroform extraction. Then, DNA was precipitated in 100% ice cold ethanol, spooled onto a glass hook, eluted in 1x TE buffer, and stored at 4 °C. 10 μg of DNA was cleaned using AMPure beads. From this DNA, five long insert libraries were prepared for PacBio sequencing according to the manufacturer’s protocols. Genomic DNA was sheared to 30-40 kb using the Megaruptor device. The PacBio libraries were size selected for fragments larger than 15-17.5 kb using the BluePippin device. PacBio SMRT sequencing was performed with the P6/C4 chemistry using 240 min sequencing runs. Average read length was 11-12 kb. In total, 86 SMRT cells were sequenced on the PacBio RSII instrument resulting in 46 gigabases (Gb; an estimated 46x coverage for a putative ∼1 Gb genome) polymerase reads.

### Assembly of the round goby genome

The round goby genome was assembled at the Heidelberg Institute for Theoretical Studies HITS gGmbH. Raw PacBio reads were assembled using the Marvel (146, 147) assembler with default parameters unless mentioned otherwise. Marvel consisted of three major steps, namely the setup phase, patch phase and the assembly phase. In the setup phase, reads were filtered by choosing only the best read of each Zero-Mode Waveguide as defined by the H5dextract tool (146) and requiring subsequently a minimum read length of 4k. The resulting 3.2 million reads were stored in an internal Marvel database. The patch phase detected and fixed read artefacts including missed adapters, polymerase strand jumps, chimeric reads and long low-quality segments that were the primary impediments to long contiguous assemblies (146). To better resolve those artefacts only low complexity regions were masked (DBdust) and no further repeat masking was done. The resulting patched reads longer than 3k (41x coverage) were then used for the final assembly phase. The assembly phase stitched short alignment artefacts from bad sequencing segments within overlapping read pairs. This step was followed by repeat annotation and the generation of the overlap graph, which was subsequently toured in order to generate the final contigs. By using an alignment-based approach, the final contigs were separated into a primary set and an alternative set containing bubbles and spurs in an overlap graph. To correct base errors, we first used the correction module of Marvel, which made use of the final overlap graph and corrected only the reads that were used to build the contigs. After tracking the raw reads to contigs, PacBio’s Quiver (148) algorithm was applied twice to further polish contigs as previously described (146).

### Automated annotation of the round goby genome

The round goby genome assembly was annotated using Maker v2.31.8 (149, 150). Two iterations were run with assembled transcripts from round goby embryonic tissue (46) and data from eleven other actinopterygian species available in the ENSEMBL database (downloaded the 15th February 2016, http://www.ensembl.org, see Table 5) as well as the SwissProt protein set from the uniprot database as evidence (downloaded March 2, 2016; https://www.uniprot.org/downloads). In addition, an initial set of reference sequences obtained from a closely related species, the sand goby *Pomatoschistus minutus*, sequenced by the CeMEB consortium at University of Gothenburg, Sweden (https://cemeb.science.gu.se), was included. The second maker iteration was run after first training the gene modeler SNAP version 2006-07-28 (151) based on the results from the first run. Augustus v3.2.2 (152) was run with initial parameter settings from Zebrafish. Repeat regions in the genome were masked using RepeatMasker known elements (153) and repeat libraries from Repbase (154) as well as *de novo* identified repeats from the round goby genome assembly obtained from a RepeatModeler analysis (153).

**Table 5.** Summary of reference data from Ensembl used for the annotation.

| Reference species | Number of protein sequences | Assembly version from ENSEMBL (downloaded 15th Feb 2016) |
| --- | --- | --- |
| *Astyanax mexicanus* | 23698 | AstMex102 |
| *Danio rerio* | 44487 | GRCz10 |
| *Gadus morhua* | 22100 | gadMor1 |
| *Gasterosteus aculeatus* | 27576 | BROADS1 |
| *Lepisosteus oculatus* | 22483 | LepOcu1 |
| *Oreochromis niloticus* | 26763 | Orenil1.0 |
| *Oryzias latipes* | 24674 | MEDAKA1 |
| *Poecilia formosa* | 30898 | PoeFor_5.1.2 |
| *Takifugu rubripes* | 47841 | FUGU4 |
| *Tetraodon nigroviridis* | 23118 | TETRAODON8 |
| *Xiphophorus maculatus* | 20454 | Xipmac4.4.2 |

In order to ensure the completeness and quality of the current assembly and the associated gene models, the assembly and the predicted protein sequences were run against reference sets at two different taxonomical levels (303 eukaryotic and 4584 actinopterygian single copy orthologues) using the BUSCO pipeline v2.0 (155, 156).

The maker annotation results were used to generate a database for JBrowse/Webapollo using the script “maker2jbrowse” included with JBrowse (157, 158). Predicted protein and transcript sequences were used to query the uniprot database, using blastp and blastn respectively, and the best hit descriptions were transferred to the fasta headers with scripts bundled with Maker as described in (150). The annotated genome is currently hosted on a WebApollo genome browser and Blast server at the University of Gothenburg, Sweden at http://albiorix.bioenv.gu.se/.

Our analyses reveal that some degree of care is warranted regarding gene models. *De novo* annotation without transcriptome data tends to be biased towards known and conserved genes, homopolymer sequencing errors may cause annotation errors, and fish proteins have diverged faster than mammalian homologs (159). For example, 25% of human genes cannot be identified in the pufferfish (16). Even in the well-characterized zebrafish, targeted approaches have the potential to reveal additional novel genes (160). We therefore encourage researchers to consider genome-wide blast searches in addition to a consultation of round goby gene models, and hope that extensive RNA sequencing data can be generated in the future to improve the predictions.

### Sex determining regions

To investigate whether the round goby genome features large sex determining regions, we analyzed own available RAD sequencing data. We prepared restriction site–associated DNA (RAD) (161) libraries following the protocol used by (162, 163), which is largely based on (166). In short, we used the Sbf1 enzyme on DNA extracted from 57 females, 56 males, and 5 juveniles caught in Basel, Switzerland, and pooled 39-40 individuals per library for SR 100bp sequencing with Illumina. 45 females and 47 males retained sufficient numbers of reads (>150000) per sample after cleaning and demultiplexing, were processed with the Stacks pipeline using the genome independent approach (164), and were analyzed for sex-specific loci present exclusively in males or females. Considering a genome size of ∼1GB, the presence of 23 chromosomes (165), and a calling success of 21877 loci in 95 or 96 individuals (49220 loci in at least 40 individuals), we expected an average density of one RAD locus every 45710 (20316) bp and an average number of 951 (2140) markers for an average sized chromosome. The presence of a sex chromosome should thus be indicated by hundreds of sex-specific RAD loci, while a contiguous sex determining region larger than 45000 bp would be indicated by one or more sex specific RAD loci. Read numbers per locus for each sample were extracted from the *.matches.tsv file output from Stacks and analyzed for sex-specific loci with standard R table manipulations.

### Vision

Opsin genes were extracted from the genome assembly using the Geneious software (http://www.geneious.com) (167) by mapping the genomic scaffolds (Medium Sensitivity, 70% identity threshold) against individual opsin exons of Nile tilapia (*Oreochromis niloticus*; GenBank Acc. no.: MKQE00000000.1). This led to capturing of all scaffolds containing any visual opsin. The genes were then annotated by mapping back of the single exons of tilapia against each scaffold separately (High Sensitivity; 50% identity threshold) combined with the Live Annotate & Predict function as implemented in Geneious, based on the Nile tilapia and mudskipper (25) opsin gene annotation. All regions upstream and downstream from every opsin gene, as well as the intergenic regions were separately tested for presence of any further opsin gene or its fragment (pseudogene). The annotated genes were checked for the reading frame and the putative protein product was predicted.

We next performed phylogenetic analysis on the visual opsin genes (i.e. SWS1, SWS2, RH2, RH1 and LWS opsins) across vertebrates, with focus on selected model species of teleost fishes. We further specifically focused on the LWS genes from the fish species or lineages known to possess multiple LWS copies, such as livebearers and pupfishes (Cyprinodontiformes) (168), zebrafish (*Danio rerio*) (169), salmon (*Salmo salar*) (170), common carp (*Cyprinus carpio*) (170), cavefish (*Astyanax mexicanus*) (171), Northern pike (*Esox lucius*) (170), labyrinth fishes (*Anabas testudineus*) (20), Asian arowana (*Scleropages* formosus) (170) as well as other gobies, such as mudskippers (25) and reef gobies (20). The opsin gene sequences from round goby and other fish species, including outgroup of non-visual opsins (pinopsin, parietopsin, vertebrate-ancestral opsins and opn3 opsin; **Supplemental_Material_S2**) were aligned using the MAFFT (172) plugin (v1.3.5) under the L-ins-i algorithm as implemented in Geneious. Exon 5 (exon 6 in case of LWS) and part of exon 1 (or entire exon 1 in case of LWS), which provided ambiguous alignment due to their higher variability, were discarded. We estimated the model parameters by jModeltest 2.1.6 (173, 174), and subsequently used the bayesian inference to calculate single-gene phylogeny using the MrBayes 3.2.6 (175) software as implemented on the CIPRES Science gateway (176).

### Olfaction

Olfactory receptor (OR) peptide sequences to be used as a reference were extracted from a publicly available *Oreochromis niloticus* protein dataset (177). Those references were blasted (tblastn) against the genomes of the round goby (*Neogobius melanostomus*), the blue-spotted mudskipper (*Boleophthalmus pectinirostris*) (25), the giant mudskipper (*Periophthalmodon magnuspinatus*) (25) and the three-spined stickleback (*Gasterosteus aculeatus*) (178), using an e-value threshold of 10e^-50^. Only the hit with highest bit-score for each genomic position with more than one alignment was employed in subsequent steps. Mapped hits belonging to contiguous positions of the protein (maximum overlap of 15 aminoacids) and with a genomic distance smaller than 10kb were joined as exons of the same CDS-gene model. Obtained sequences were translated to proteins using TransDecoder (http://transdecoder.github.io), filtering all models that produce peptides smaller than 250 aminoacids. While many ORs are usually around 300 aminoacids long in total, 250 is close to the average size of their main transmembrane domain, which is centrally located in the protein and more suitable to interspecific alignment compared to N-terminal and C-terminal ends. We acknowledge that this method might introduce a reduced proportion of recent pseudogenes that could lead to a small overestimation of OR genes with coding capacity, although all species should be affected equivalently.

Next, an hmmscan (http://hmmer.org/) was produced against Pfam database to identify the domain with highest score for each obtained protein sequence. We also filtered against false positive detection using blast against confident OR and non-OR protein datasets. For phylogenetic analysis, sequences (**Supplemental_Material_S3**) were aligned with MAFFT (https://mafft.cbrc.jp/alignment/server/) and a Maximum Likelihood methodology was employed to build the tree using W-IQ-TREE software (179) with standard parameters and Ultrafast bootstrap (180). Four adrenergic receptor sequences from *Oreochromis niloticus* were used as an outgroup. Monophyletic groups formed by five or more genes of the same species were considered as lineage-specific gene expansions. Because of the phylogenetic proximity of the two mudskippers and the differences in their genome assembly statistics, only *B. pectinirostris* was considered and *P. magnuspinatus* sequences were allowed to be included in their lineage-specific expansion groups.

### Detoxification

The Basic Local Alignment Search Tool (BLAST, v. 2.2.31) (181) was used to identify local alignments between the round goby genome and a query including all annotated CYPs in humans and zebrafish (vertebrate) and the most dissimilar invertebrate CYPs from *Drosophila melanogaster* (arthropod), *Caenorhabditis elegans* (nematode) and *Capitella teleta* (annelid; **Supplemental_Material_S4**). Only BLAST high scoring pairs with Expect values of 1.0 x 10-10 or smaller were considered significant.

The JBrowse genome viewer (v1.12.1) was used to manually annotate the significant regions of each genome from the BLAST search, identifying start (ATG) and stop (TGA/TAA/TAG) codons, exon number, and splice site signals (GT/AG) at intron-exon boundaries. The lengths of the potential CYPs were identified and considered full length at ∼500 amino acid residues long. Potential genes were matched to the well-curated cytochrome P450 HMM in the Pfam protein family database (182) to confirm identity. The ScanProsite tool (183) was used to verify the presence of four largely conserved CYP motifs: the I-helix, K-helix, meander coil and heme loop. Each gene was classified as ‘complete’ (proper length with start and stop codon, all motifs present, and match to the HMM) or ‘partial’ (presence of at least the entire ∼120 amino acid region that contains all motifs but clearly less than full length). Any potential CYP that was missing at least one of the motifs was considered a gene ‘fragment’ (**Supplemental_Table_S9**).

All of the ‘complete’ and ‘partial’ round goby CYPs (**Supplemental_Table_S9**) were included in further analyses. Clustal Omega (v1.2.4) (184) was used to generate a multiple sequence alignment of the round goby sequences and a variety of well-known vertebrate CYPs from humans, *Danio rerio*, *Mus musculus*, *Xenopus laevis*, *Gallus gallus*, and *Rattus norvegicus* (125 sequences in total; **Supplemental_Material_S5**). Mesquite (v3.10) (185) was utilized to trim the alignment, especially at the termini of the protein sequences where significant variation is typically observed, leaving only the portion of the alignment representative of the homology of the sequences. The final ‘masked’ alignment (**Supplemental_Material_S6**) was used as input for the Randomized Axelerated Maximum Likelihood program (RAxML v8.2.10) (186). 100 bootstrap trees were generated with the rapid generation algorithm (-x) and a gamma distribution. The JTT substitution matrix with empirical frequencies was implemented in tree generation. The final maximum likelihood phylogenetic tree was visualized with Figtree (v1.4.3) (187) and rooted with the CYP51 family of enzymes.

### Osmoregulation

Protein sequences for aquaporins, tight junction proteins, ion transporters, and enzymes in osmolyte production pathways were retrieved from the round goby genome by BLASTing well-characterized proteins from zebrafish, downloaded from Uniprot (March 2018), against the round goby gene models/proteins. Only round goby gene-models/proteins for which the predicted protein covered at least 70%, with a sequence identity of at least 40% and with E-value < 10^-20^ of the corresponding protein in zebrafish were used for the phylogenetic analyses. Well-established paralogues belonging to different subclasses of the respective protein family, based on either literature search or from initial phylogenetic analysis of that particular protein family, were used as additional query sequences to minimize the risk of missing relevant round goby sequences. Osmoregulatory genes from human and zebrafish were used for overall classification of clades in the respective protein family. Some modifications were made to the retrieved round goby sequences before analysis: i) For NHE ion transporters, a 780 aa long non-homologous N-terminus from one of the *Neogobius* sequences was removed before the phylogenetic analysis. ii) Some of the claudin genes were subjected to manual curation of Maker predicted proteins. The claudin genes in fish consist of several tandem arrays, which in some cases results in merging of 2-4 claudin genes by the Maker software. Claudins have a typical trans-membrane (TM) pattern with four distinct TM domains. All manually curated claudin genes from round goby were examined to have the expected four TM domains by TMHMM searches. Round goby protein sequences after manual curation are available in the supplement (**Supplemental_Material_S7**).

No myo-inositol phosphate synthase (MIPS) and sodium/inositol cotransporter (SMIT) proteins from zebrafish was found in Uniprot. To confirm that there are truly no MIPS and SMIT genes in zebrafish, the zebrafish genome at NCBI was also searched for homologies using blastp and tblastn using as query the MIPS and SMIT protein sequences from tilapia as query, and no hits were found. Thus, in the case of MIPS and SMIT, tilapia sequences were used for searching for round goby homologues. For the phylogenetic analyses, protein sequences from zebrafish (*Danio rerio*), three spine stickleback (*Gasterosteus aculeatus*), tilapia (*Oreochromis niloticus*), mudskipper (*Boleophthalmus pectinirostris*) and *Homo sapiens* (exception for human NKA-beta) were used in comparison to round goby, and were obtained from Uniprot (zebrafish, stickleback, tilapia, human) or RefSeq (mudskipper; **Supplemental_Material_S7**). Phylogenetic analyses of osmoregulatory proteins in round goby were performed using maximum likelihood with PhyML v3.0 with 100 bootstraps and using Gblocks to eliminate poorly aligned positions and highly divergent regions. PhyML analyses were performed at the Phylogeny.fr website (http://www.phylogeny.fr) using default settings.

### Immune system

To perform an overall characterization of key genes related to the immune system, protein queries representing core components of innate and acquired immunity from several fish species as well as mammalian reference sequences were downloaded from UniProt and Ensembl. The protein queries were aligned prior to usage to ensure sequence homology. We also added previously extracted protein sequences from the Toll-like receptor family, reported by (85), and MHCI sequences reported by (80). All queries are listed in **Supplemental_Table_S4**. To enable comparative analyses between sequenced Gobiiformes, the genomes of *Periophthalmodon schlosseri* (GCA_000787095.1), *Periophthalmus magnuspinatus* (GCA_000787105.1), *Scartelaos histophorus* (GCA_000787155.1) and *Boleophthalmus pectinirostris* (GCA_000788275.1) were additionally downloaded from NCBI.

All protein queries were used in a tblastn (blast+ v. 2.6.0) towards the round goby genome assembly using default parameters and a e-value cutoff of 1e-10 (188). Some queries (*caspase-1*, *TLRs*, *IL1* and *IL8*) were also used in an identical tblastn towards the other Gobiiformes genomes. Genomic hit regions were extracted using BEDtools (v. 2.17.0) extending both up- and downstream as needed to obtain full length gene sequences (189). The extracted genomic regions were imported into MEGA7, the reading frame was adjusted for each exon and aligned as proteins to the corresponding translated coding sequence of queries using MUSCLE with default parameters. Intronic sequences were removed leaving an in-frame coding sequence (190, 191). All alignments were subjected to manual evaluation before subsequent analysis.

To generate phylogenetic trees, protein alignments were made and model tested using the ProtTest3 server (http://darwin.uvigo.es/software/prottest_server.html) specifying BIC and no tree optimization (server has been disabled but ProtTest is available for download from GitHub) (192). All alignments reported the JTT model as best hit. Maximum likelihood trees were produced by using RAxML-PTHREADS (v 8.0.26), PROTCATJTT, rapid bootstrap and 500 bootstrap replicates (193). The final trees were imported into FigTree (http://tree.bio.ed.ac.uk/software/figtree/), and subsequently Adobe Illustrator, for presentation purposes.

In order to identify members of the large multigenic family of fish-specific NACHT and Leucine-Rich Repeats containing genes (NLRs; the fish-specific subset is also known as NLR-C) (87), an alignment of 368 zebrafish NLR-C proteins was obtained from (88). A combination of tblastn, HMMER3 searches (194) and alignments with MAFFT v7.310 (195) was used to generate first an initial list of “candidate regions” potentially containing an NLR (**Supplemental_Material_S8**) and then an annotation of the characteristic domains in round goby NLR-C family members (see **Supplemental_Material_S9)** and **Supplemental_Table_S8** for details), consisting of 25 PYRIN, 1 N-terminal CARD, 12 C-terminal CARD, 343 FISNA-NACHT and 178 B30.2 domains. Custom HMM models for major NLR exons (FISNA-NACHT, and PRY-SPRY/B30.2) were generated and utilized during this process (Supplementary Methods, **Supplemental_Material_S10**). The majority of identified FISNA-NACHT exons contained frameshifts or a large insertion, indicating either pseudogenization, acquisition of new introns, problems with the assembly, or a combination of the three (196). In any case, for the subsequent phylogenetic analysis, only the 61 clearly intact NLRs were used. These were aligned with NLRs from human, zebrafish and the mudskipper goby using MAFFT (**Supplemental_Material_S9**; **Supplemental_Material_S11**); Maximum Likelihood trees were produced with RAxML-PTHREADS, PROTCATJTT, rapid bootstrap and 500 bootstrap replicates (193). The final trees were imported into FigTree (http://tree.bio.ed.ac.uk/software/figtree/), and subsequently Adobe Illustrator. The alignments were inspected manually for presence of the conserved Walker A motifs and sequence logos for these were generated with WebLogo (197). Finally, we performed a survey of the PYD domains, Peptidase_C14 domains (Caspases) and CARD. All cases of a PYD domain followed by an adjacent CARD in the round goby (putative apoptosis-associated speck-like protein containing a CARD (ASC), also known as PYD-CARD or PYCARD) were identified from the HMMER3 dataset. The open reading frames containing these were translated, concatenated, and aligned with similarly structured proteins from human, mouse, lizard, frog and all the fish in Ensembl, and with PYD-CARDs identified from the other available goby assemblies (**Supplemental_Material_S12**). A phylogenetic tree was generated as described above. The annotation for NLR-C genes consists of predicted positions for all of the major conserved NLR-associated domains (PYD, CARD, FISNA-NACHT-helices, LRRs, B30.2; **Supplemental_Table_S8**).

### Epigenetic regulators

We focused on two gene expression regulators which are conserved among all eukaryotes: the Polycomb Repressive Complex 2 (PRC2), which deposits repressive histone methylation marks, and the DNA methylases, which methylate cytosine in CpG contexts. The presence of both marks is commonly associated with a downregulation of gene expression. The protein sequences of zebrafish orthologues of PRC2 components RBBP2, EED, EZH1-2, and SUZ12 (50) and of DNA methylases DNMT1 and DNMT3 (198) were blasted against the round goby genome using default parameters of the Albiorix Blast server. The protein sequence of predicted proteins at the hit site was extracted manually in the round goby genome browser and aligned with mouse, human, and zebrafish protein sequences. When the first and/or last exon sequences as predicted in the round goby genome differed significantly from the mouse, human, and zebrafish sequences, we attempted confirmation by 3’ and 5’RACE on RNA extracted from whole juvenile animals (see **Supplemental_Material_S13** for primer sequences and PCR conditions). A putative CDS was combined from automated annotation and RACE results, and aligned to sequences extracted from a variety of fish taxa, shark, chicken, frog, lizard, and human (**Supplemental_Material_S14**). Given the high conservation of these proteins in eukaryotes, and the absence of major unexpected differences between round goby and other vertebrates, additional Gobiidae were not included in the analyses. In order to perform codon aware alignment MACSE (199) was used. The model and partitioning scheme used for each phylogenetic analysis was estimated using PartitionFinder2 (200) using PhyML (201) with corrected AIC scores (AICc) used for model selection. Phylogenetic analyses were performed using MrBayes 3.2.6 (175, 202) with three independent runs for each gene. Analyses were run for 2,000,000 generations or until the standard deviation of split frequencies was below 0.01 up to a maximum of 20,000,000 generations. In order to aid convergence in the EZH analyses the temperature parameter was set to 0.05.

### Transposable elements

A number of different applications were used for the repeat annotation of the genome. They are described in the repeat annotation report (**Supplemental_Material_S15**). In summary, in addition to the identification of repeats with RepeatModeler (as described above), we used TRF (203) to predict tandem repeats. RepeatMasker (153), a homology-based approach was used to produce a genome-wide overview of interspersed repeats. LTR Finder (204) and LTRharvest (205) in combination with LTRdigest (206), both de novo approaches, were used to predict LTRs.

## Availability of data and materials

The Whole Genome Shotgun project has been deposited at DDBJ/ENA/GenBank under the accession VHKM00000000. The version described in this paper is version VHKM01000000.

Various Illumina reads are available under the accessions indicated in Table 1.

Other datasets supporting the conclusions of this article are included within the article and its additional files.

## Additional files

Supplemental Figure S1: .pdf; full opsin phylogenetic tree

Supplemental Figure S2: .pdf; opsin tree constructed with individual exons

Supplemental Figure S3: .pdf; full olfactory receptor phylogenetic tree

Supplemental Figure S4: .pdf; claudin phylogenetic tree

Supplemental Figure S5: .pdf; occludin phylogenetic tree

Supplemental Figure S6: .pdf; sodium transporter phylogenetic trees

Supplemental Figure S7: .pdf; myo-inositol production and accumulation gene phylogenetic trees

Supplemental Figure S8: .pdf; TAP gene phylogenetic tree

Supplemental Figure S9: .pdf; Ig locus schematic

Supplemental Figure S10: .pdf; TLR phylogenetic tree of Gobiidae

Supplemental Figure S11: .pdf; CRP / APCS phylogenetic tree

Supplemental Figure S12: .pdf; SUZ12, EED, and RBBP4 phylogenetic trees

Supplemental Material S1: .gz; annotation tracks for genome

Supplemental Material S2: .txt; opsin sequences used for tree building

Supplemental Material S3: .txt; olfactory receptor sequences used for tree building

Supplemental Material S4: .txt; CYP sequences used as query

Supplemental Material S5: .txt; CYP sequences used for tree building

Supplemental Material S6: .phy; CYP alignment

Supplemental Material S7: .doc; sequences for osmoregulatory proteins

Supplemental Material S8: .fas; NLR candidate regions

Supplemental Material S9: .doc; detailed methods for NLR annotation

Supplemental Material S10: zip of .hmm; hmm models used to identify NLRs

Supplemental Material S11: .fas; NLR sequences used for vertebrate tree building

Supplemental Material S12: .fas; NLR sequences used for Gobiidae tree building

Supplemental Material S13: .doc; detailed methods for RACE

Supplemental Material S14: zip of .nex; dnmt1, dnmt3, eed, ezh, rbbp4, and suz12 alignments

Supplemental Material S15: .txt; detailed methods for repeat annotation

Supplemental Table S1: .xls; overview and results for repeat sequences analysed

Supplemental Table S2: .xls; results for CYP genes (location, sequence, length)

Supplemental Table S3: .doc; overview table of immune genes analysed

Supplemental Table S4: .xls; immune gene sequences used as query

Supplemental Table S5: .doc; results for MHCI

Supplemental Table S6: .doc; results for MHCII

Supplemental Table S7: .txt; results for other immune genes analysed

Supplemental Table S8: .xls; results for NLRs

Supplemental Table S9: .xls; results for CYPs

## Acknowledgements

We are grateful to Prof. Patricia Burkhardt-Holm for her continuous support and encouragement. We thank Bernd Egger, Astrid Böhne, Philipp Hirsch and Patricia Burkhardt-Holm for critically reading the manuscript. We thank Fabio Cortesi for his insightful comments and the Center for Marine Evolutionary Biology for hosting a Blast server and a genome browser. Computational resources were provided by the CESNET LM2015042 and the CERIT Scientific Cloud LM2015085, provided under the programme “Projects of Large Research, Development, and Innovations Infrastructures”. We thank Maria Leptin for her helpful comments and advice during annotation of the inflammasome components.

## Author contributions

Sylke Winkler isolated DNA and generated PacBio reads, Martin Pippel and Siegfried Schloissnig assembled the genome sequence, and Tomas Larsson, Mats Tölpel and Magnus Alm Rosenblad performed automated annotation and provided the genome browser and Blast server.

Jean-Claude Walser provided transposable element analyses, Silvia Gutnik, Claire Peart, and Irene Adrian-Kalchhauser provided DNA methyltransferase and PRC2 analyses, Anders Blomberg provided osmoregulation analyses, Monica Hongroe Solbakken and Jaanus Suurväli provided immune gene analyses, Zuzana Musilova and Demian Burguera provided vision and olfaction analyses, Joanna Yvonne Wilson and Kirill Pankov provided CYP gene analyses, Nico Michiels investigated red fluorescence.

Irene Adrian-Kalchhauser initiated, designed, and supervised the project, acquired the necessary funding, coordinated annotation efforts, compiled the manuscript and handled the submission process.

## Permissions

Fish used in this work were caught in accordance with permission 2-3-6-4-1 from the Cantonal Office for Environment and Energy, Basel Stadt.

---

**Supplemental_Fig_S1.**
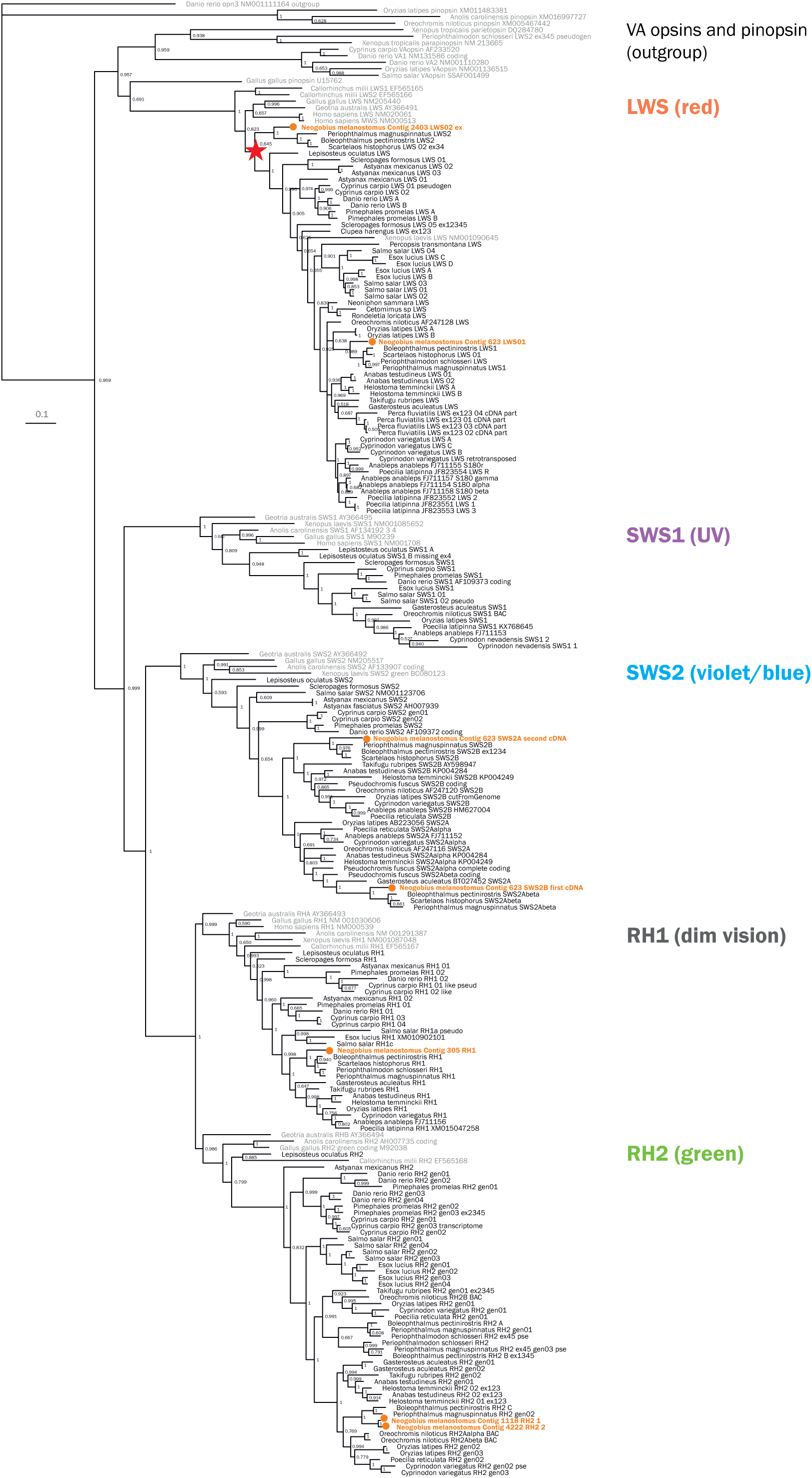
Phylogenetic tree of vertebrate opsin protein sequences. Maximum-likelihood phylogenetic tree with VA opsins and pinopbsins as outgroup. Round goby is indicated in orange. Non-teleost species and the outgroup (VA opsins and pinopsins) are indicated in grey.

**Supplemental_Fig_S2.**
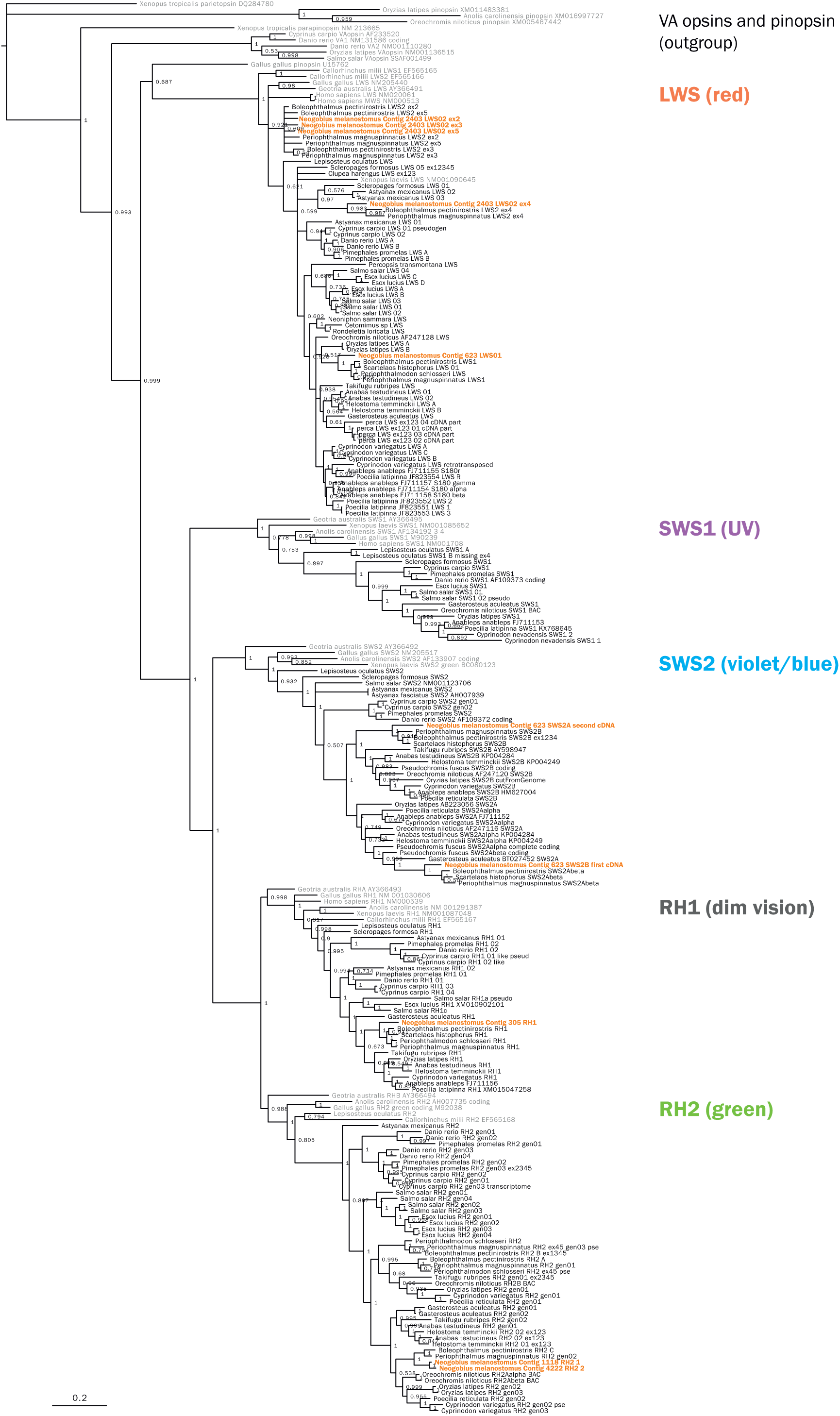
Phylogenetic tree of vertebrate opsin protein sequences, using single exons. Maximum-likelihood phylogenetic tree with VA opsins and pinopbsins as outgroup. Round goby is indicated in orange. Non-teleost species and the outgroup (VA opsins and pinopsins) are indicated in grey.

**Supplemental_Fig_S3.**
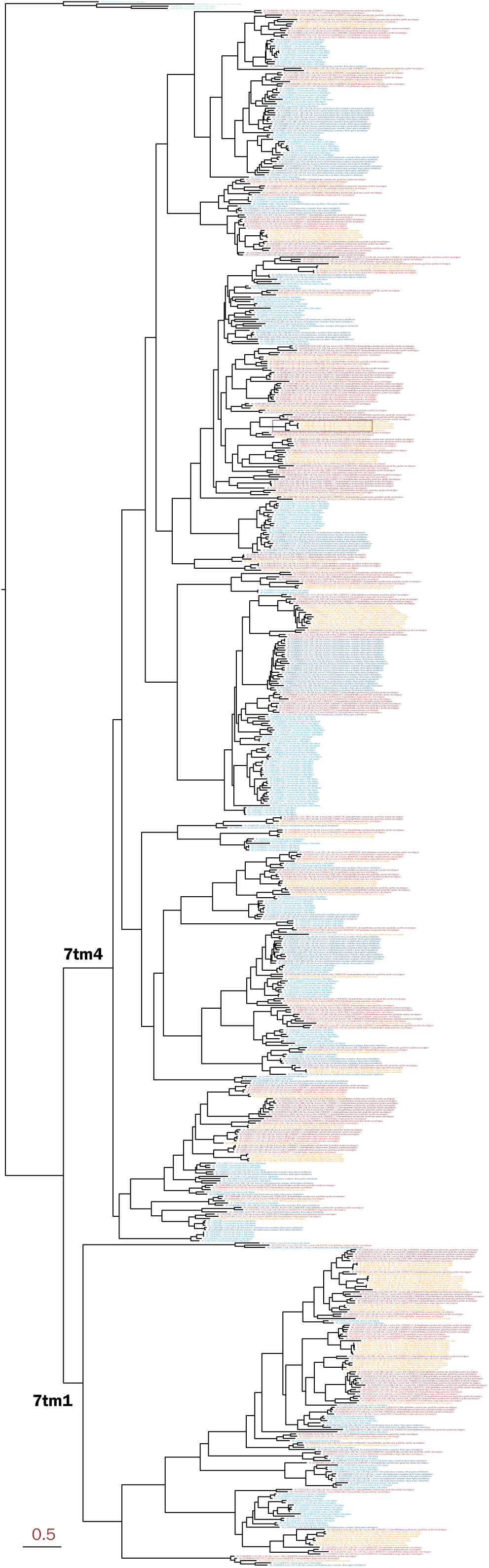
Maximum-likelihood phylogenetic tree of percomorph olfactory receptor protein sequences.

**Supplemental_Fig_S4.**
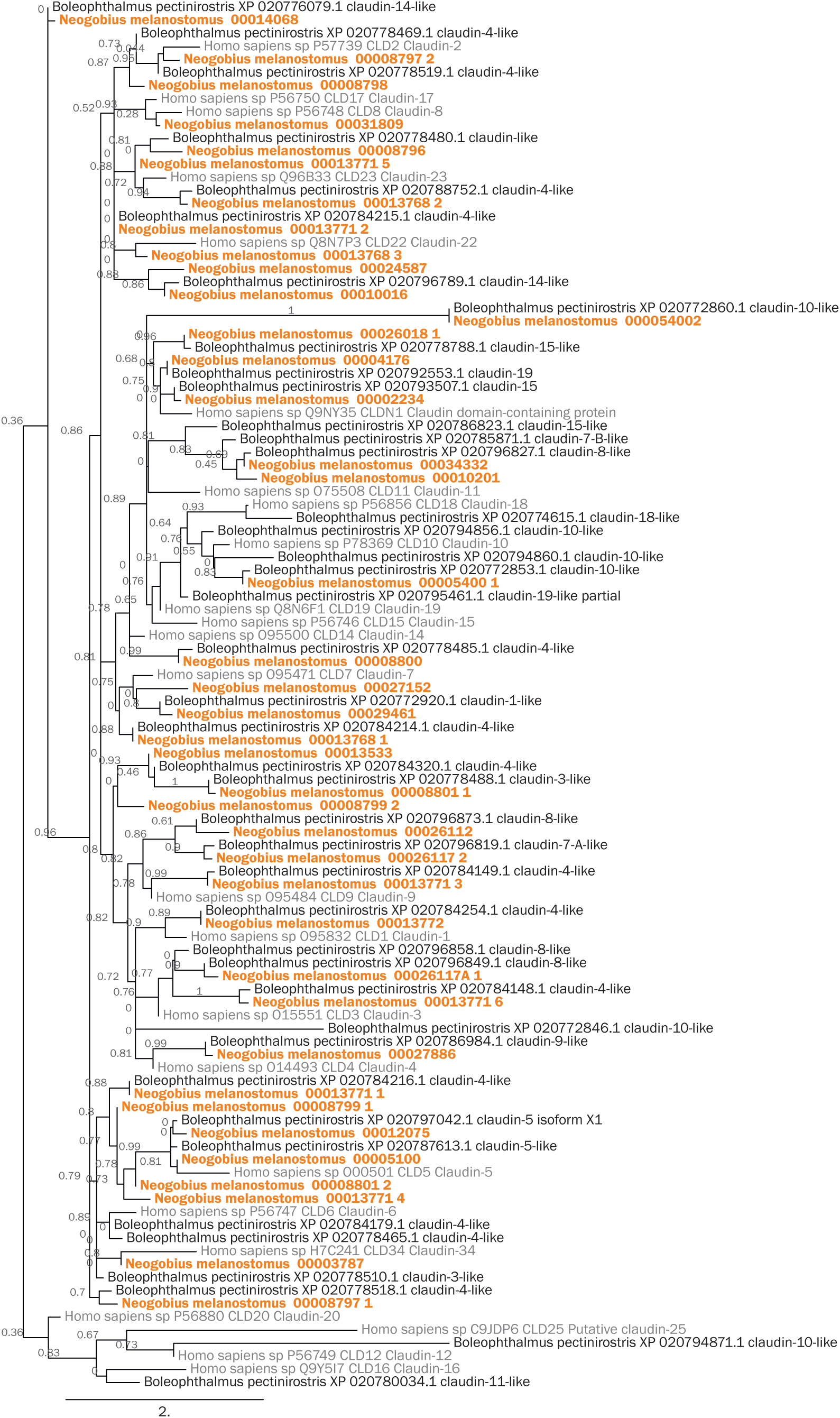
Phylogenetic tree of vertebrate claudin genes. Maximum-likelihood tree with 100 bootstraps of round goby (Neogobius melanostomus, orange) in relation to great blue-spotted mudskipper (Boleophthalmus pectinirostris) and human (Homo sapiens, grey).

**Supplemental_Fig_S5.**
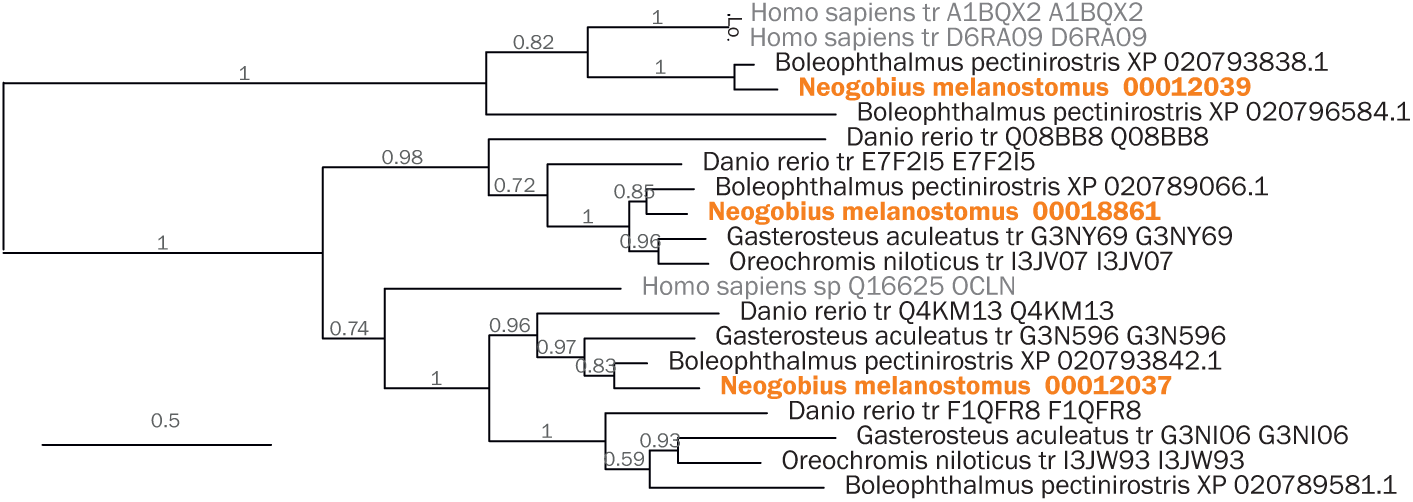
Phylogenetic tree of vertebrate occludin genes. Maximum-likelihood tree with 100 bootstraps of round goby (Neogobius melanostomus, orange) in relation to great blue-spotted mudskipper (Boleophthalmus pectinirostris), stickleback (Gasterosteus aculeatus), nile tilapia (Oreochromis niloticus), zebrafish (Danio rerio), and human (Homo sapiens, grey).

**Supplemental_Fig_S6.**
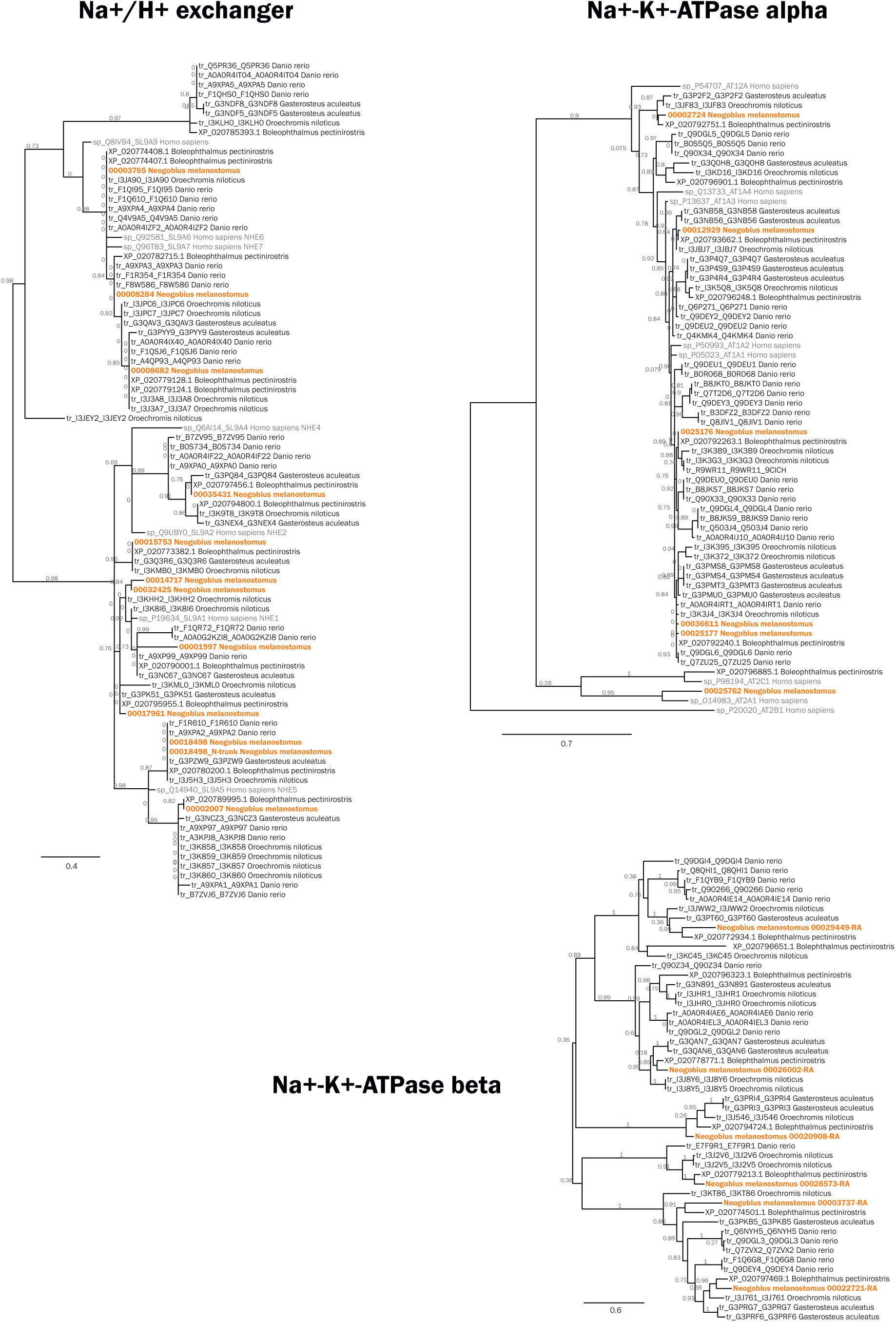
Phylogenetic tree of vertebrate ion transporters. Maximum-likelihood tree with 100 bootstraps of round goby (Neogobius melanostomus, orange) in relation to great blue-spotted mudskipper (Boleophthalmus pectinirostris), stickleback (Gasterosteus aculeatus), nile tilapia (Oreochromis niloticus), zebrafish (Danio rerio), and human (Homo sapiens, grey)

**Supplemental_Fig_S7.**
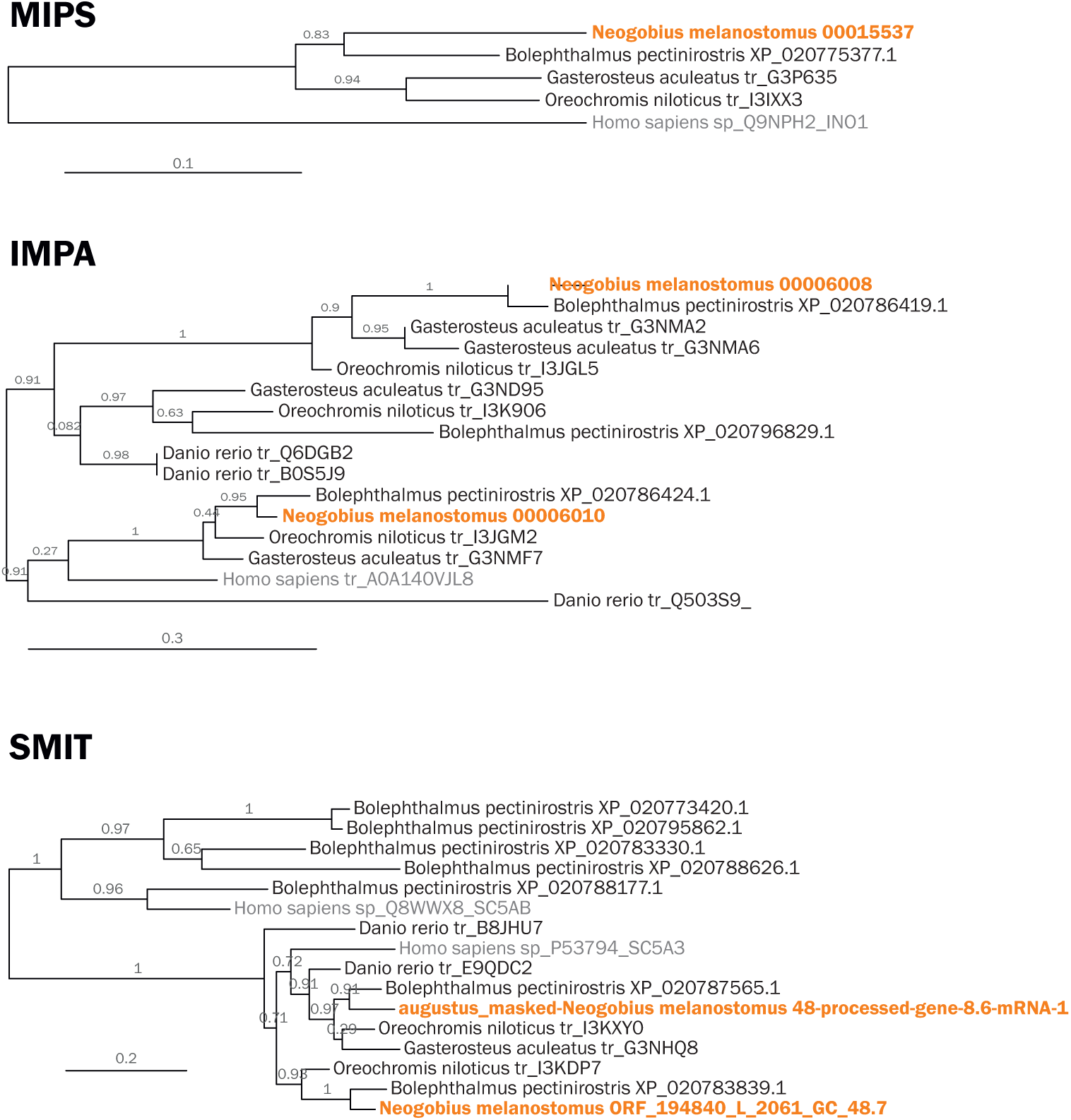
Phylogenetic tree of vertebrate genes promoting osmolyte production. Maximum-likelihood tree with 100 bootstraps of round goby (Neogobius melanostomus, orange) in relation to great blue-spotted mudskipper (Boleophthalmus pectinirostris), stickleback (Gasterosteus aculeatus), nile tilapia (Oreochromis niloticus), zebrafish (Danio rerio), and human (Homo sapiens, grey)

**Supplemental_Fig_S8.**
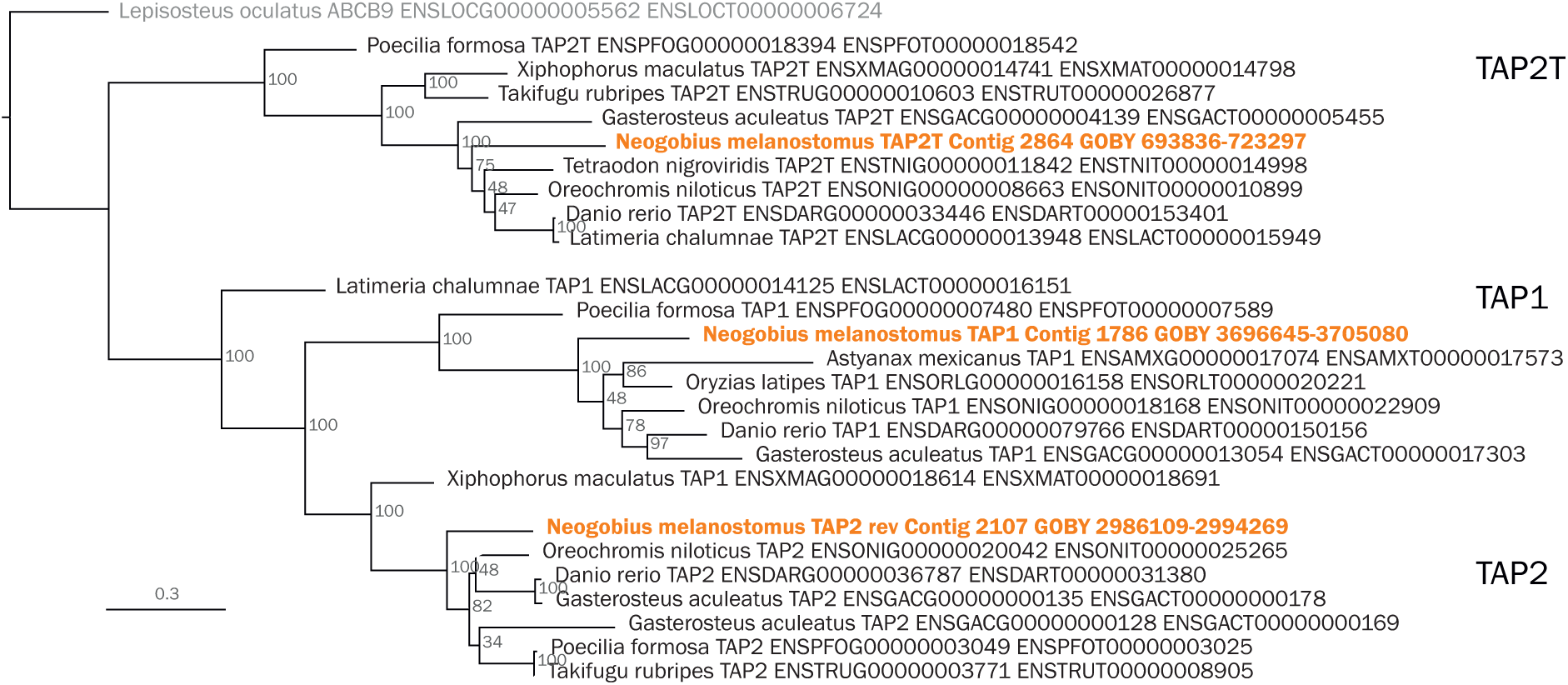
Phylogenetic tree of fish TAP genes. Maximum-likelihood tree with 500 bootstraps. Round goby (Neogobius melanostomus) is labeled orange. Outgroup: Lepistosteus oculatus ABC transporter.

**Supplemental_Fig_S9.**
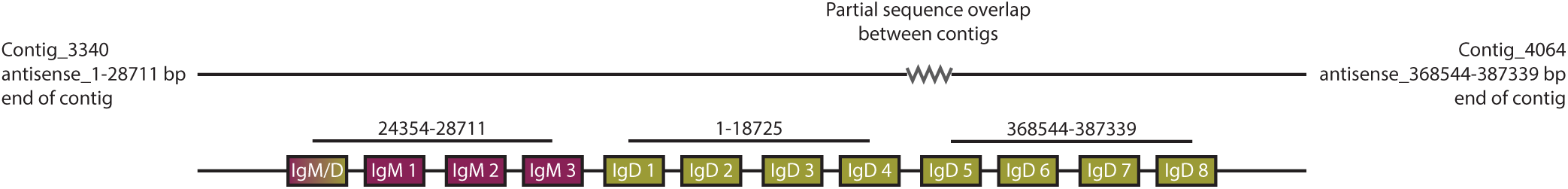
The Ig locus spans two contigs and contains IgM and IgD domains.

**Supplemental_Fig_S10.**
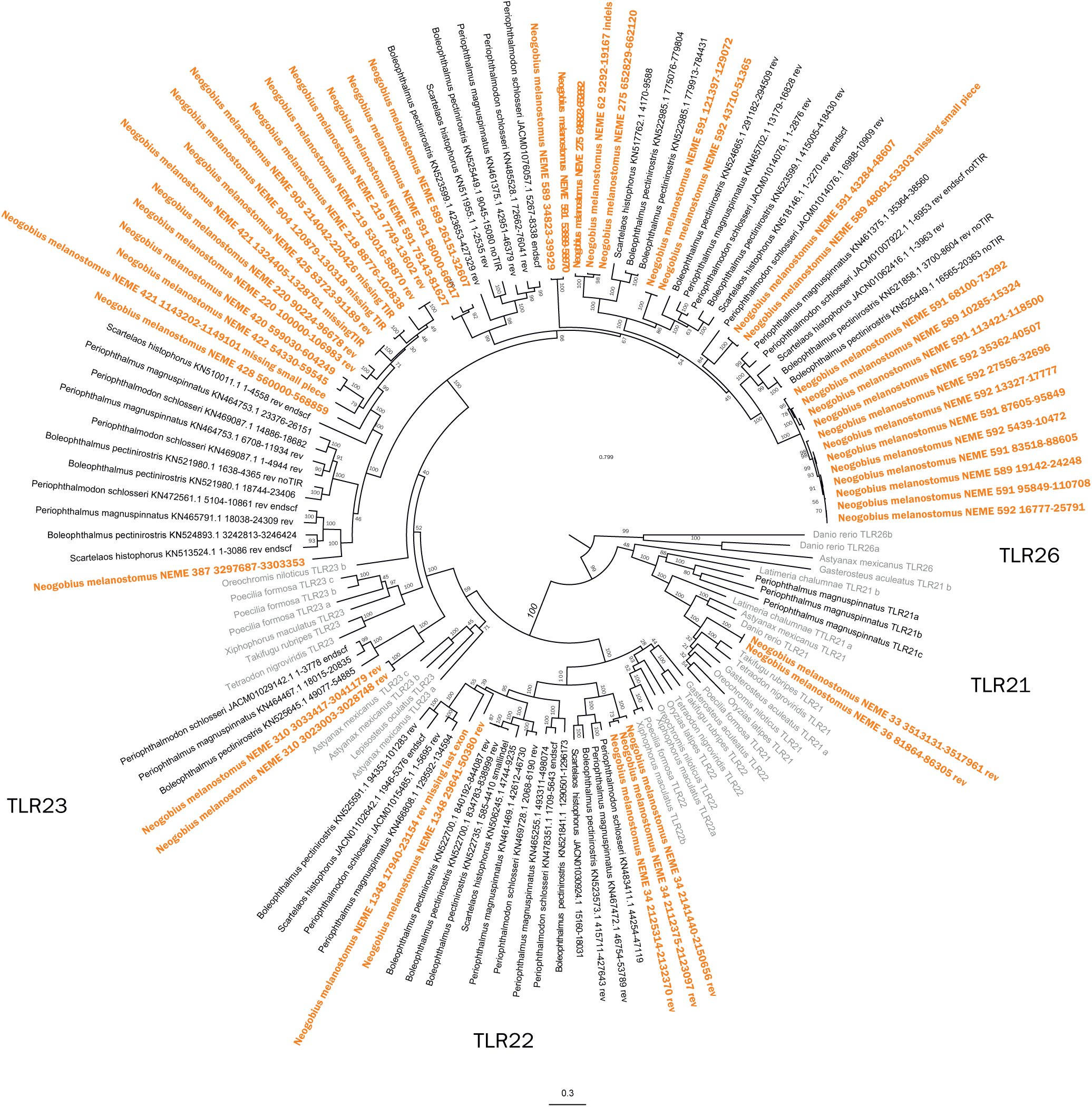
Phylogenetic tree of Gobiidae Toll Like Receptor protein sequences. A maximum likelihood phylogenetic tree run with the JTT substitution model and 500 bootstrap replicates on the transmembrane, linker and TIR domain of all TLRs found in Gobiiformes (Neogobius melanostomus, orange; other Gobiidae, black) and selected other fish (grey) for context.

**Supplemental_Fig_S11.**
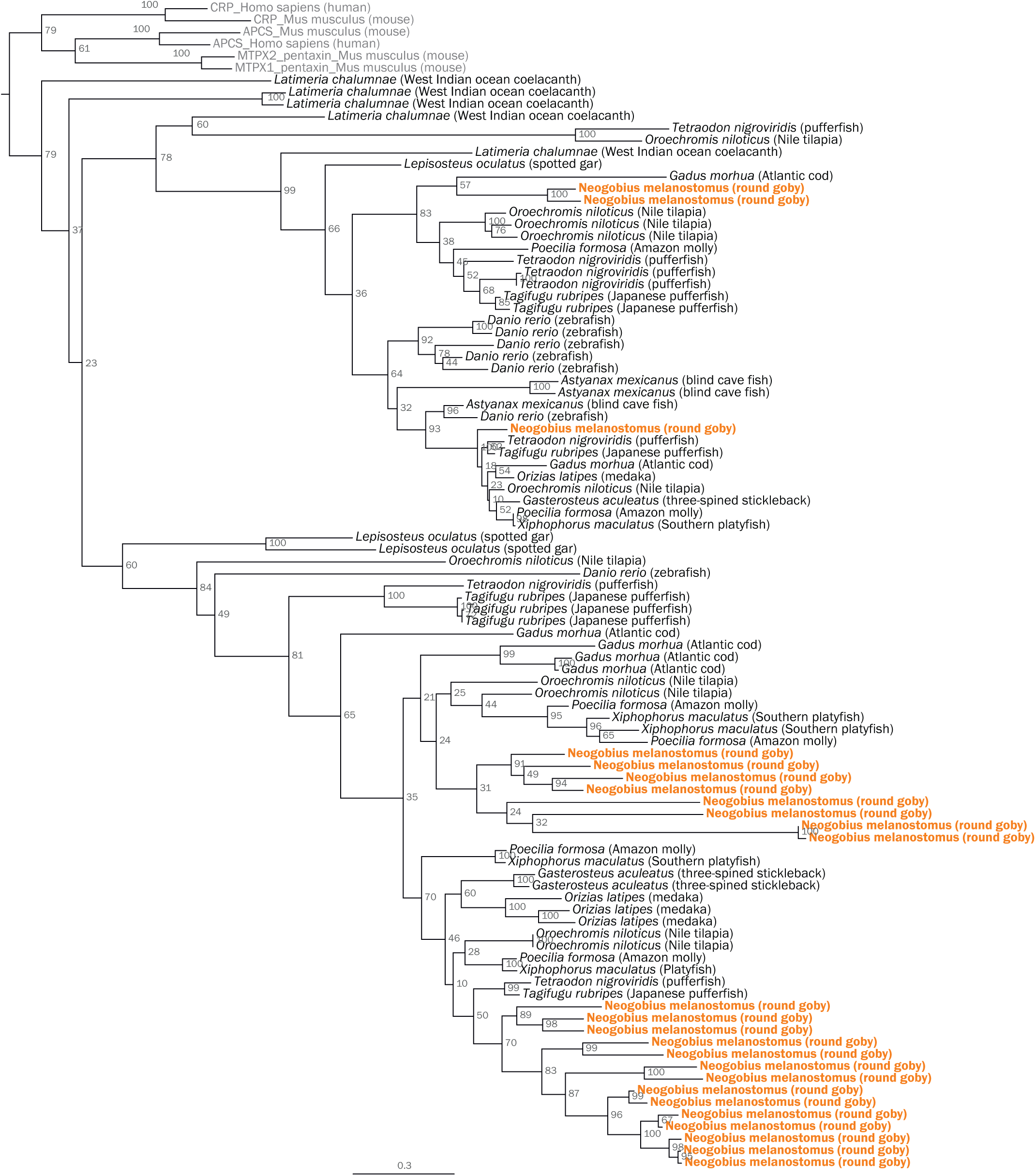
Phylogenetic tree of fish CRP/APCS sequences. Maximum Likelihood phylogenetic tree with 500 bootstraps rooted at the split between tetrapods and ray-finned fish. Tetrapods were used as outgroup and are indicated in grey. Round goby is indicated in orange.

**Supplemental_Fig_S12.**
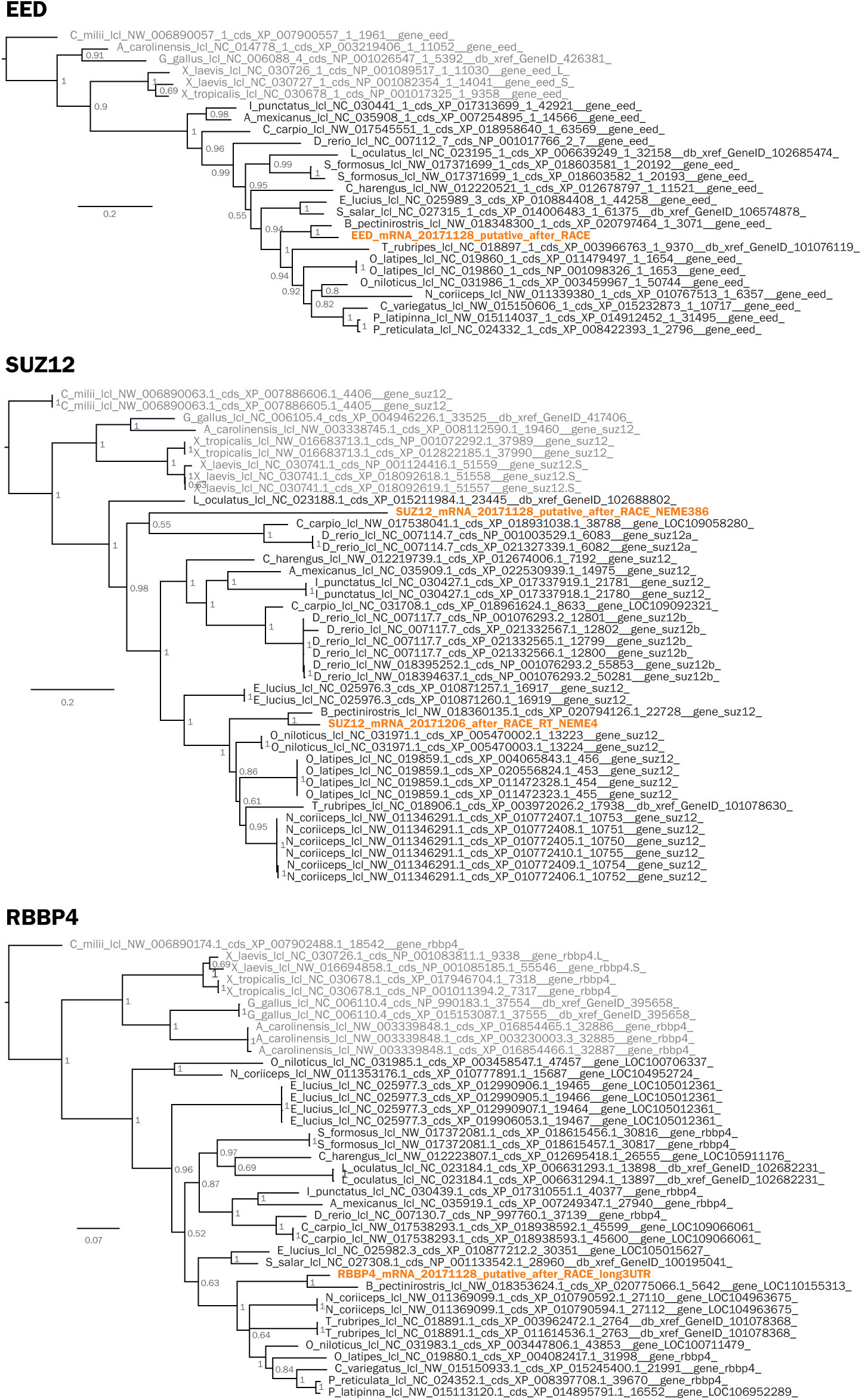
Phylogenetic tree of vertebrate PRC2 components EED, SUZ12, and RBBP4. Bayesian phylogenetic tree rooted with Australian ghostshark (C. milii). Round goby is indicated in orange.

