## Supplementary material for "Evolved for success in novel environments: The round goby genome": Supp Materials: Supplemental_Material_S7.docx

*Supplementary Material Osmoregulation*

Fasta sequences for round goby (*Neogobius melanostomus*), zebrafish (*Danio rerio*), three spine stickleback (*Gasterosteus aculeatus*), tilapia (*Oreochromis niloticus*), mudskipper (*Boleophthalmus pectinirostris*) and *Homo sapiens* the following protein families:

AQUAPORINS	page 1-20

CLAUDINS	page 21-55

OCCLUDINS	page 56-61

INOSITOL MONOPHOSPHATASES (IMPA)	page 62-65

MYO-INOSITOL PHOSPHATE SYNTHASES (MIPS)	page 66-67

SODIUM/INOSITOL COTRANSPORTERS (SMIT)	page 68-73

Na+/H+ EXCHANGERS (NHE)	page 74-103

Na+/K+-ATPase alfa-subunits (NKA)	page 104-132

Na/K-ATPase beta-subunits (NKB)	page 133-142

Na+/K+/Cl- COTRANSPORTERS (NKCC)	page 143-158

**AQUAPORINS**

***Round goby (Neogobius melanostomus)***

>NEME_00028054_Neogobius

MMLFIFLSLVTAIGNKNNANPDQEVKVALGFGLAIATLAQSLGHISGAHLNPAVTLGMLA

SCQISVLKAVMYIIAQMLGSALASGIVYGVRPNTTDALGLNSLSGVTPSQGVGIELLATF

QLVLCVIAVTDKRRRDVTGSAPLAIGLSVCLGHLAAISYTGCGINPARSFGPAVILNDFT

NHWVYWVGPMCGGVVAALIYDFLLSPKLDDFPERMKVLVSGPVGDYDVNGRNDVTNMEMS

SK

>NEME_00028052_Neogobius

MRELKSWPFWRAVAAEFIGMLLFIVFGIASAIGSPGNAAQEVKVSLAFGLAIATLVQTLG

HVSGAHLNPAVSIGLLVSCQISLFRFLCYTVAQMLGAVAASALLDGFRARKIGLAVNQLT

GVSAGQGFAIEFFATLQLVLCVIAVTDKRRTDVQGSAPLAIGLSVTLGHLVAISYTGCGI

NPARSFGPAIIRSQMANHWVYWLGPMCGGITAAFIYDLILAPRSPSLSARRDILLNGPVE

DTNDNLIRDGGSPGPSQWPKQ

>NEME_00001894_Neogobius

MERILRKCRIQNQLVRECLAECLGVYILILFGCGSVAQVTTTRNQKGEYLSINLGFALGV

TLGVYVSRGVSGAHLNPAVSLSLCALGRHSWFKLPFYIFFQLLGAFLAAATVALQYYEAI

QSYSGGELTVSGPNATAGIFCTFPDEYLSLLGGFIDQVIGTAALLLCILALGDSGQGSIP

EGVQPVLVGAVVMVIGVSMGSNSGYALNPARDFGPRLFTYIAGWGVDVFTAGGGWWWVPL

VAPCVGALLGTLVYELMIEVHHDPGLKHRRPVR

>NEME_00000693_Neogobius

MVAFKGVWTKDFWRAVTAEYLATLIFVFLGVGSTINWHAGTDNPPPADLVLISLCFGLSI

ATMVQCFGHISGGHINPAVTAAMVVTRKLSLAKGMFYVVAQCLGAITGAGILYMVTPAAV

RGGFGVNDVSSSVSLGHGLVVEIIITFELVFTVFATCDPKRTDLGGSAALAIGVAVAIGH

LFGIYWLGPILGGLLAAGLYEYLYCPDPDLKKRLKQVFQKDQSGRYKEVDAGDITIKPGS

IDAEKAEKKDSYNDMGDVLSSE

>NEME_00011098_Neogobius

MASAMEIDLQQFQGFNSFKDLRIFKDFRNFKDFSSFKLRSRQFWRAVLAEMLGTLVVVST

VLGSSVPGPGDVPVGPLYPALAVGVAIVTVATCFGEISGAQVNPSVTLALLATRRMDVTR

AMVPLELNAGQALGIEVLITFQMVFTLFSVEDQRRRDSTEPGNLALGLTHTAGVFIGARF

SGACMNPARALAPAVITGFWENHWVYWIGPILGAVLAGLSHEFLFAPSASRQKLVACLTC

KDIEIVESTSMTGSSFHSQIVTMWEFRSMNFWRAVFAEFFGTMFFVFFGMGAALRWTTGP

HHVLHVALCFGLAAATLIQSIGHISGGHINPAVTFAYLVGSQMSLFRAVFYMAAQCLGAV

AGAAVLYGVTPGNMRGNMAMNTLQPGISLGMATTVEVFLTMQLVVCIFAVTDERRNGRLG

SAALSIGFSVTIGHLMGMYYTGAGMNPARSFAPAVLFRNFINHWVYWVGPMIGGAMGALL

YDFMLFPRMRGLSERLATLKGSRPSESETQQDPAESPSSSRHKPYKPVPGQTEETRNRPT

SPPICITHTSTSLHHAPNGHS

>NEME_00003357_Neogobius

MEENKRRMKEKFGLRRDIFKEFLAEFLGIFVLILFGCGSVAQTVLSKGALGEPLTIHIGF

SLGVMMAVYMAGGVSVGSFCPTTSLKMIIGIGLMSFTGAWISGEEGSGGHVNPAVSLAMV

ILGKLPIKKFPVYVTAQFLGAFAGSCAVFGLYYGKCLKLITLQKTLPYHIYSLDALMEYT

NGEFVVTGPNATANIFASYPAKHLSVLNGFVDQVIGTGALILCILAITDKRNIGAPKGME

PLCIGLAITAIAVSMGLNCGYPINPARDLGPRFFTAVAGWGLDVFSAGGCWWWIPVAGPM

VGGAVGAGIYFLCVELHHPEPEKQDEKNIPDKYEMVTMS

>NEME_00007642_Neogobius

LERDTQRQPKRRSMRRRCALKHGICREFLAEFLGTFVLVLFGCGSVAQSVLSRSQLGEPL

TVHIGFSVGLMMGVYVSGGVSGGHVNPAVSLAMVILGKLKIWKFPIYVLAQFLGAFAGAA

TVFGLYYDAFMDFTSGILSVTGINATGHIFASYPARHLSVLGGFIDQVVGTGMLVLCILA

IIDGGNIGAPKGIEPLAIGLIIMAIGVSMGLNCGYPLNPARDLGPRLFTAVAGWGWEVFS

TAGYWWWIPVAGPMVGAVVAAALYFLLVEMHHKQADPQKPTEEEEEDDDEDDEDEDSSLK

DKYEMIIMN

>NEME_00005163_Neogobius

MGLHKVGLERVSRFFQIRNLLLRQAMAECLGTLILVMFGTGAVAQLVLSGGSHGMFLTVN

FAFGFAATLGILVCGQISGGHLNPAVTFALCLLGRERWRKFPTFFLFQTIGAFCGAAIIF

GMYYDALWDFPGCFNMTGKTATAGIFATYPGDHLTIVNGFFDQINICTCVTLFSLTMADH

WDSLSDSLYLGYCGSTQPCRPPGLEAFTVGFVVLVIGLSMGFNSGYAVNPARDLGPRIFS

AMAGWGTTVFSFRNGWFLVPTFVPFIGSFIGVVVYQVMVGFHVEGEVRDRKAAEHENERI

RLTNVKTNNDN

>NEME_00026274_Neogobius

MVQSMELGSEVQQKTLRSRVRIHNQLLRLGLCETLCTYVMMAFGLCSVAQVVTGRGEFGH

YISINLSFGLAVALGVHVGGGVSGDPELLRRKPDSDRARSHGGNLCHVPRPYLSITGGFI

DQPVVVGLLVLLIGMSAGSNSGYAINPTRDIAPRIFTAIAGWGPHVFRAGNGWWWVPLVA

PCVGAVMGTALYKLLVQLHHPPPPPGEEAEERPGESRGESVERAPLDKHQNYCV

>NEME_00015375_Neogobius

MGVEKMEMEGDKLMGKKKAVPKPKSKYERVFQPCLAELIGTMFFVFIGCVSVIENVPAAG

RLQPALVHGLAVAVLVAVMDNIRSSLRHPEVTKPALRRPLREVAMTCLITMVVLLVAVNS

KTKTPLAPFLVGCTVIINVLAGGDISGTCLNPARAFGPAVIANYWTYHWVYWAGPIGGGV

VGAILLRLILGDEKMRVILKT

>NEME_00015374_Neogobius

MWSQPREQNSLVNSIGSFVQCTNVYEQYVQPVLAEFYGSTLFIFVGCACVVGNTGTGAIQ

PAVAHGLALGVVTVSSCSLDRGHFNPAVTLSVYLCGGMSLRLVVLPNVLAQMAGGMAGAG

LVKAMFPPEVYQASSGGAFKPQALAKPWESHTCRTCDDHVPHHRVTMGAVNRKTSSQMAP

FGIGLTVAANILAGRALSGACMNPARAFGPAVAANHWDHHWIYWVGPLSGALSIVIYVRY

SSHGCIMQLFMREKFDICE

>NEME_00001205_Neogobius

MYPAPAKWYDRRDAVFIEFCVEDSRDVKVIFDKTKLDFSCISGTDNVKHHNEVELFEPGK

TWVRLTKAKAKCNWLSVDFNNWKDWENDSDEDLSGFDKFSEIMKNMGGDDLDIDEISSAN

MWEFRSMSFWRAVFAEFYGTMFFVFFGLGAALRWTTGPINVLHVAFCFGLAAATLIQSIG

HISGGHINPAVTFAYLIASQMSLFRAFFYIFAQCLGALAGAAVLYGVTPTNMRGNLALNM

LKPGISLGMATTMEIFLTVQLVICIFAVTDERRNGRLGSAALAIGFSVLMGHLLGMYYTG

AGMNPARSFAPAVLVRNFVNHWVYWVGPMIGGAIGALLYDFMLFPRVRGLSERLATLKGI

RPREAEGQSETRGEPIELKTQAL

>NEME_00035108_Neogobius

LPGGGGNGSSRTTARTHVAPLRPVSQVEPPARCWVCGYRGRRGRSQRMEAAELAQWAADR

ADVAVSLAVLVGLVLLGEGVRRILGRVLAHSGLEAHAAELMSTLQLCCCTHELRVLSDVG

RIERRLAHTLTYVAAVVHALTFRGAIGNPACTLVHVYRRRLVLGAALRLIVCQFAAAAAA

RTIMRFAWSLGLSGMHVHHQARGFQCTSPIHAPLTATAVELVCAFAVQTAMTHTRNVEDK

YRVHAVAAVITTAVYAGGPSTGAVFNPALAFSTQFRCSGPSLAEYCLVYWLGPMLGMLGS

VLLSDWVEGQKPVPWR

>NEME_00023043_Neogobius

MSGLNASLGYFVTVVIFAVLSRALIRKWPRLSFLLEFASSFMLVGCLLEVQTIVEVGEWA

GGLGPDVTLTMLFAVLLSHGVISGTASGNPSVSLQKFLQLDASALSTLLALVAQFVGSHL

AIMAAVYYWSLELTDMHMIKNLMASECSTSLKVSLVQGFITEFVCALVFHLLLLGLRRRS

TLIRVPLIAVLLTFLSHTARNYTSGFVNPSLAYGLTFHCPGFSHLQYAVVYWLSSVTGMM

LALVLFMGHIPKLFATNLLYSQKTRFRSRKAEKGEKKKK

>NEME_00028479_Neogobius

MSAPVLSAVGVSVGVVVSVVLLCELLRWFRRFLSGDSGRVLLEGASTLQLCWCALELKLL

GERSSDLPLSWRLSLLYAVTACTCRASAGQLLSDWPSGARVSGRLTLRAALGLVSVQLSA

ALVARALAVLVWSHGWSEVHEQHRRFGYRCFDPLGGTLLEAAANELALSFIVQGVWAVSG

AVLNPVLAVSLQFPCSGHSLLDYLFVYWLGPILVNNSQDLIH

***Zebrafish (Danio rerio)***

>AQP1a_tr|Q6NZ72|Q6NZ72_DANRE_Danio

MNELKSKAFWRAVLAELLGMTLFIFLSITAAVGNTNTQNPDQEIKVALAFGLSIATLAQS

LGHISGAHLNPAVTLGLLASCQISLLRAVMYILAQMIGATVASAIVLGVSKGDALGLNQI

HTDISAGQGVGIELLATFQLVLCVLATTDKRRRDVSGSAPLAIGLSVCLGHLTAISFTGC

GINPARTFGPAMIRLDFANHWVYWVGPMCGGVAAALIYDFLLYPKMDDFPERVRVLVSGP

ATDYEVNGTDDPPAVEMSSK

>AQP3_tr|Q803U6|Q803U6_DANRE_Danio

MGWQKSVLDKLAQTFQIRNKLLRQGLAECLGTLILVMFGCGSLAQLKLSEGSHGLFLTAN

LAFGFGATLGILVCGQVSGGHLNPAVTFALCLLGREKWRKFPVYFLFQTLGSFLGAAIIF

AEYHDAIYDYAGESNELLVLGEKETAGIFATYPSKYLTPLNGFFDQVIGTASLIVCILAI

VDPYNNPIPQGLEAFTVGFSVLIIGLSMGFNSGYAVNPARDFGPRLFTAMAGWGSEVFTA

RDYWFLVPIFAPFIGAVIGVIVYQLMVGWHVEGEARDKKAKAREEVMNLNDVASKE

>AQP8a.1_tr|Q66I64|Q66I64_Danio

MTSAESKSELFTVATGDGGDNHQNQPKKLPFFEHYIQPCLAEVVGSFLFMFVGCVSVMGN

VGISGSIQPALAHGLALAIAIAIFGEISGGHFNPAVSVCVYLIGGMEVILLVPYIISQML

GGVIAASLAKAVTTNDAFSNATGAAFNAIPSSDGIGAATMAEMIMTLFLTIVVSMGAVNG

RTKSQLAPFCIGLTVTANILAGGGISGACMNPARAFGPAVVSGHWTHHWIYWVGPLTGAL

VTVSIVRLVMGDKKVRVIFK

>AQP0a_tr|Q6DEI6|Q6DEI6_DANRE_Danio

MWEFRSMSFWRAVFAEFYGTMFFVFFGLGAALRWTTGPHNVLQVAFCFGLAAATFIQSIG

HISGGHINPAVTFAYLIGSQMSLFRAFFYICAQCLGALAGAAVLYGVTPTNMRGNLALNT

LQPGISMGMATTIEIFLTLQLVVCVFAVTDERRNGRLGSAALSIGFSVLVGHLLGMYYTG

AGMNPARSFAPAVLYRNFINHWVYWVGPMIGAAMGALLYDFMLFPRVRGLSERLAVLKGN

KPTEPEAQQETRGEPIELKTQAL

>AQP3b_tr|D3TI80|D3TI80_DANRE_Danio

MGRQKVILEKMARIFQIRNMLMRQALAECLGTLILVMFGCGALAQHILSGGSHGMFLTVN

FAFGFAATLGILVCGQVSGGHINPTVTFSLCLLGREPWRKFPVYFLAQTVGAFLGAGIIF

GMYFDAIWKFGQGSLDVDGVNATAGIFATYPSKHLTLLNGFFDQMIGTAALIVCILAIVD

PYNNPIPQGLEAFTVGFVVLVIGLSMGFNSGYAVNPARDLGPRIFTAIAGWGSKVFSAES

YWSFVPVFAPFIGAVFGVMVYQLMVGCHVKGEERDKREAVEREEKERLKLSAVSDKDAA

>AQP12_tr|Q0IIQ6|Q0IIQ6_DANRE_Danio

MSGLNSSLGFFLAVVGLSVSGRLLLRRWTVLLELVSAFALCACRLEVDTIAEVGQWAGAL

GPDVAVTMLFLSIAVHTAVMQDVSGNPAVTLLRLLQRDVSVVVAVLSIAAQLIGAFLALE

VAGRFWAMELSDMHMIKNLMMSECSTSLRVSTALGVSTEALAALLLLLLHLVLKNRSQML

KVPALSVALTLIAYTANNYTSGYVNPALAYAVTFTCPGHSFLVYSLVYWLGPLIGVFLAV

FLYLGNIPLLFSKNLLYSKKNRFRLPKGKTNDEKSS

>AQP8a.2_tr|D3TI82|D3TI82_DANRE_Danio

MPVEAEKLELEELDKTLLNRDKPKPQSKYERIFQPCIAELVGTTFFVFIGCVSVIENVEA

AGRLQPALVHGLAVAVLVACMAEISGSHFNPPFTIAIWLCGGMQLTMVVPYLISQLIGGV

LGAAMSKVMTSDENYANATGAAFAVLKSDEQLGKVVFAEMAMTCLVTLVVLMGAVNGKSK

SPMVPFMVGCTVIVNILAGGDVSGTCLNPARAFGPALMANHWTYHWVYWVGPLGGGLVAA

ALMRLLLGDEKLRVVMK

>AQP10a_tr|Q6DHQ6|Q6DHQ6_DANRE_Danio

MKRMKVKNELARQIMGEILGTFVLLLFGCAAAAQVKTSRETKGQFLSGNIAFSVGVMSAM

YLCRAVSGAHLNPAVSLSFCVLGDLAWIKLLPYSLAQILGAYLASGLVYLIYHDAIMEFS

GGVLTVFGPNETASIFATYPTDVVSVQTNFLDQVVGTAMLMLCILPLNDKRNAPAPEALL

PPIVATVVLGISISMSANCGAAINPARDLGPRLFTFTAGWGTEVFTCYDYFFWIPLVAPM

VGGVLGSIIYLVFIQWHLPELEDESESGEMNDQTKVMEHNNKKDEIYLKMSSI

>AQP9a_tr|F1QA20|F1QA20_DANRE_Danio

MEYLENIRNLRGRCVLRRDIIREFLAELLGTFVLILFGCGSVAQTVLSREAKGQLLTIHF

GFTLGVMLAVYMAGGVSGGHVNPAVSLAMVVLRKLPLKKFPVYVLAQFLGAFFGSCAVYC

LYYDAFTEFANGELAVTGPNATAGIFASYPREGLSLLNGFIDQVIGAGALVLCILAVVDK

KNIGAPKGMEPLLVGLSILAIGVSMALNCGYPINPARDLGPRLFTAIAGWGLTVFSAGNG

WWWVPVVGPMVGGVVGAAIYFLMIEMHHPENDKNLEDDNSLKDKYELNTVN

>AQP10a_tr|F1Q9W1|F1Q9W1_DANRE_Danio

MSQMKKIMKRMKVKNELARQIMGEILGTFVLLLFGCAAAAQVKTSRETKGQFLSGNIAFS

VGVMSAMYLCRAVSGAHLNPAVSLSFCVLGDLAWIKLLPYSLAQILGAYLASGLVYLIYH

DAIMEFSGGVLTVFGPNETASIFATYPTDVVSVQTNFLDQVVGTAMLMLCILPLNDKRNA

PAPEALLPPIVATVVLGISISMSANCGAAINPARDLGPRLFTFTAGWGTEVFTCYDYFFW

IPLVAPMVGGVLGSIIYLVFIQWHLPELEDESESGEMNDQTKVMEHNNKKDEIYLKMSSI

>AQP9a_tr|Q498W2|Q498W2_DANRE_Danio

MKQHCALKQRLFKEFLAEFLGTFVLVLFGCGSVAQTVLSRNTLGEPLTIHIGFSTGLMMG

VYVSGGVSGGHLNPAVSLAMVILGKLKIWKFPVYVIAQMLGAFAGAAAVFGLYYDAFMEF

TSGILSVTGINATGHIFSSYPGRHLTVLGGFVDQVVGTGMLVLCILAIVDGRNIGAPRGV

EPLAVGVVLLGISVSMGLNCGYPLNPARDLGPRLFTALAGWGMEVFSTADYWWWIPVAGP

LVGGVVGAVIYFLLIELHHSNHNDTPQEEPEEEEDEDEEEDSSLKDKYEMINMS

>MIPb_tr|Q4ZJI3|Q4ZJI3_DANRE_Danio

MWEFRSMMFWRAVFAEFFGTMFFVFFGMGAALRWTTGPYHVFHTALCFGFAAATLIQSIG

HISGGHINPAVTFAYLVGSQMSVFRAFFYICAQCLGAMAGAAALYGVTPNNMRGTMALNT

LQPGMSLGMATTVEVFLTMQLVVCVFAVTDERRNGRLGSAALSIGFSVTMGHLMGMYYTG

AGMNPARSFAPAVITRNFINHWVYWVGPMIGAAMGAIFYDFFLFPRMRGFSERLATLKGS

RPPEAENQQETRGEPIELKTQTL

>AQP10a_tr|A0A0R4IZM3|A0A0R4IZM3_DANRE_Danio

MCSFFLLQLFGCAAAAQVKTSRETKGQFLSGNIAFSVGVMSAMYLCRAVSGAHLNPAVSL

SFCVLGDLAWIKLLPYSLAQILGAYLASGLVYLIYHDAIMEFSGGVLTVFGPNETASIFA

TYPTDVVSVQTNFLDQVVGTAMLMLCILPLNDKRNAPAPEALLPPIVATVVLGISISMSA

NCGAAINPARDLGPRLFTFTAGWGTEVFTCYDYFFWIPLVAPMVGGVLGSIIYLVFIQWH

LPELEDESESGEMNDQTKVMEHNNKKDEIYLKMSSI

>AQP1a.1_tr|Q32ZE7|Q32ZE7_DANRE_Danio

MNELKSKAFWRAVLAELLGMTLFIFLSITAAVGNANTQNPDQEIKVALAFGLSIATLAQS

LGHISGAHLNPAVTLGLLASCQISLLRAVMYILAQMIGATVASAIVLGVSKGDALGLNQI

HTDISAGQGVGIELLATFQLVLCVLATTDKRRRDVSGSAPLAIGLSVCLGHLTAISFTGC

GINPARTFGPAMIRLDFANHWVYWVGPMCGGVAAALIYDFLLYPKMDDFPERVRVLVSGP

ATDYEVNGTDDPPAVEMSSK

>AQP9a_tr|D5FFZ1|D5FFZ1_DANRE_Danio

MEYLENIRNLRGRCVLRRDIIREFLAELLGTFVLILFGCGSVAQTVLSREAKGQLLTIHF

GFTLGVMLAVYMAGGVSGGHVNPAVSLAMVVLRKLPLKKFPVYVLAQFLGAFFGSCAVYC

LYYDAFTEFANGELAVTGPNVTAGIFASYPREGLSLLNGFIDQVIGAGALVLCILAVVDK

KNIGAPKGMEPLLVGLSILAIGVSMALNCGYPINPARDLGPRLFTAIAGWGLTVFSAGNG

WWWVPVVGPMVGGVVGAAIYFLMIEMHHPENDKNLEDDNSLKDKYELNTVN

>AQP10a_tr|D3TZW7|D3TZW7_DANRE_Danio

MSQMKKIMKRMKVKNELARQIMGEILGTFVLLLFGCAAAAQVKTSRETKGQFLSGNIAFS

VGVMSAMYLCRAVSGAHLNPAVSLSFCVLGDLAWIKLLPYSLAQILGAYLASGLVYLIYH

DAIMEFSGGVLTVFGPNETASIFATYPTDVVSVQTNFLDQVVGTAMLMLCILPLNDKRNA

PAPEALLPPIVATVVLGISISMSANCGAAINPARDLGPRLFTFTAGWGTEVFTCYDYFFW

IPLVAPMVGGVLGSIIYLVFIQWHLPELEDESESEEMNDQTKVMEHNNKKDEIYLKMSSI

>AQP10b_tr|Q1L8T0|Q1L8T0_DANRE_Danio

MFGGVFWSLCFNTVRVWISGSGHHLSEYQGRVPVNQPGLRTGNHIWHLHCKRSVRFSWTR

LPFYVCSQLFGAFLAAATVALQYYDAIMDFTGGHLTVSGATATAGIFSTYPADYLSLWGG

VVDQIIGTAALLVCVLALGDAHNTPAPAGLEPVLVGAAVLVIGISMGSNSGYAINPARDF

GPRLFSYIAGWGDEVFRAGHGWWWVPIIVTCVGALLGSLLYELLIGVHHPDSEAVDHEDP

TAALQQTVEMEGAQSFDTIKENKKSGIFSITSADVG

>AQP4_tr|Q6AZD2|Q6AZD2_DANRE_Danio

MTSCGALDTFRRCVSSCSCNNSIMAAFKGVWTQEFWRAVSGEFLAMIIFVLLSLGSTINW

GAKQENPPPADLVLISLCFGLSIATLVQCFGHISGAHINPAVTVAMVATRKLSLAKGVFY

LLAQCLGAVVGAAILYGVTPASVRGGMGVTSVNEEISAGHAIVIELIITFELVFTVFATC

DPKRNDLKGSAALAIGLSVCIGHLFAIPYTGASMNPARSFGPAVIMVKWQDHWVYWVGPL

IGGILAAAVYEYLFCPDPDLKRRYADVLSKSPFQMEPYRVVDTDSYPSDQAQLMAKQAAL

RVLDLEKKERESTGEVLSSV

>AQP7_tr|F1QYN2|F1QYN2_DANRE_Danio

FNSHQCNYSKSLNDQAGREESSAEKEVKQEVSIMEDGSIQGRMAPNVGSMLKIKNEYIRV

ALAESLCTFIMMVFGLGTVAQVVTGEGYFGEYLSINIGFGLAVAMGVHVGGKVSGAHMNA

AVSFTMCVFGRLRWKMLPLYVFAQFLGSFLAAGTIFSLYYDAINHFCGGNLTVSGPKATA

GIFATYPAPYISVYTGFFDQVAGTGLLLLCLMALSDQRNQPLVSGGEAVGVGLLVMLIGV

SMGSNSGYAINPTRDLGPRLFTLMAGWGTEVFRAGNCWWWVPLVAPFIGGVLGALIYKAL

VELHHPDLKSTITRPAVDPECIPLDKCKNGRIEIPV

>AQP1a_tr|F1QQB8|F1QQB8_DANRE_Danio

MARELKSWSFWRAVLAEFVGMTIFVFIGIASAIGNKHNRYPDQEVKVALAFGLAIATLAQ

SLGHISGAHLNPAITLGLLVSCQISFFRAFMYIIAQMLGAVLASGIMFKVSPDPDTTLGL

NMLGNGVKVGQGFAIELFATFQLVLCVLATTDKNRTDVSGSAPLAIGLSVGLGHLVAISY

TGCGINPARSFGPAVVLESFKNHWIYWIAPMCGGVAAALIYDFLLFPKREALRKRMNVLK

GTADPDPSATEALIEPRSARSGSGQWPRP

>AQP11_tr|A0A0R4IIL3|A0A0R4IIL3_DANRE_Danio

MADLTVSLSILVGIVVLSEFARRTALYLFPNREWIIYMLEFISTFQLCACTHELKLLAEL

GGLDPQTGLTLTFIISVVHGFSFRGAICNPTGALELLSRGTLPWGCALARVSCQLTAAVV

SRWVMPVAWALALSDLHQRHSLTGFRCSSSPVNAPVLQAAAVELSCAFVMHTAVNNAEKL

EEKYRVPAVAALITTLVYAGGHLTGAVFNPALAFSIQFPCPGNTFTEYSFVYWMGPILGM

SASLLLSDKLIPAISGKSTIPLQLNSNGLKKKKMK

>AQP10b_tr|F1QZV4|F1QZV4_DANRE_Danio

MDRLLRRYRIKSRLPRECLAEFFGVYVLILFGCGSVAQVTTSQNTKGEYLSINLGFALGT

TFGIYIAKGVSGAHLNPAVSVSLCVLGRFSWTRLPFYVCSQLFGAFLAAATVALQYYDAI

MDFTGGHLTVSGATATAGIFSTYPADYLSLWGGVVDQIIGTAALLVCVLALGDAHNTPAP

AGLEPVLVGAAVLVIGISMGSNSGYAINPARDFGPRLFSYIAGWGDEVFRAGHGWWWVPI

IVTCVGALLGSLLYELLIGVHHPDSEAVDHEDPTAALQQTVEMEGAQSFDTIKENKKSGI

FSITSADVG

>AQP7_tr|A0A0R4IAY6|A0A0R4IAY6_DANRE_Danio

MEPIWWMEVFGLGTVAQVVTGEGYFGEYLSINIGFGLAVAMGVHVGGKVSGAHMNAAVSF

TMCVFGRLRWKMLPLYVFAQFLGSFLAAGTIFSLYYDAINHFCGGNLTVSGPKATAGIFA

TYPAPYISVYTGFFDQVAGTGLLLLCLMALSDQRNQPLVSGGEAVGVGLLVMLIGVSMGS

NSGYAINPTRDLGPRLFTLMAGWGTEVFRAGNCWWWVPLVAPFIGGVLGALIYKALVELH

HPDLKSTITRPAVDPECIPLDKCKNGRIEIPV

>AQP7_tr|D3TZW4|D3TZW4_DANRE_Danio

MEDGSIQGRMAPNVGSMLKIKNEYIRVALAESLCTFIMMVFGLGTVAQVVTGEGYFGEYL

SINIGFGLAVAMGVHVGGKVSGAHMNAAVSFTMCVFGRLRWKMLPLYAFAQFLGSFLAAG

TIFSLYYDAINHFCGGNLTVSGPKATAGIFATYPAPYISVYTGFFDQVAGTGLLLLCLMA

LSDQRNQPLVSGGEAVGVGLLVMLIGISMGSNSGYAINPTRDLGPRLFTLIAGWGTEVFR

AGNCWWWVPLVAPFIGGVLGALIYKALVELHHPDLKNTTTRPAVDPECIPLDKCKNGRIE

IPV

>AQP8a.1_tr|A7MCF8|A7MCF8_DANRE_Danio

MTSAESKSELFTVAAGDGGDNHQNQPKKLPFFEHYIQPCLAEVVGSFLFMFVGCVSVMGN

VGISGSIQPALAHGLALAIAIAIFGEISGGHFNPAVSVCVYLIGGMEVILLVPYIISQML

GGVIAASLAKAVTTNDAFSNATGAAFNAIPSSDGIGAATMAEMIMTLFLTIVVSMGAVNG

RTKSQLAPFCIGLTVTANILAGGGISGACMNPARAFGPAVVSGHWTHHWIYWVGPLTGAL

VTVSIVRLVMGDKKVRVIFK

>AQP10b_tr|D3TI83|D3TI83_DANRE_Danio

MDRLLRRCRIKSRLPRECLAEFFGVYVLILFGCGSVAQVTTSQNTKGEYLSINLGFALGT

TFGIYIAKGVSGAHLNPAVSLSLCVLGRFSWTRLPFYVCSQLFGAFLAAATVALQYYDAI

MDFTGGHLTVSGATATAGIFSTYPADYLSLWGGVVDQIIGTAALLVCVLALGDAHNTPAP

AGLEPVLVGAAVLVIGISMGSNSGYAINPARDFGPRLFSYIAGWGDEVFRAGHGWWWVPI

IVTCVGALLGSLLYELLIGVHHPDSEAVDHEDPTAALQQTVEMEGAQSFDTIKENKKSGI

FSITSADVG

>AQP8a.1_tr|D3TZW5|D3TZW5_DANRE_Danio

MTPAESKSELFTVATGDGGDNHQNQPKKLPFFEHYIQPCLAEVVGSFLFMFVGCVSVMGN

VGISGSIQPALAHGLALAIAIAIFGEISGGHFNPAVSVCVYLIGGMEVILLVPYIISQML

GGVIAASLAKAVTTNDAFSNATGAAFNAIPSSDGIGAATMAEMIMTLFLTIVVSMGAVNG

RTKSQLAPFCIGLTVTANILAGGGISGACMNPARAFGPAVVSGHWTHHWIYWVGPLTGAL

VTVSIVRLVMGDKKVRVIFK

>AQP8b_tr|F6NNW3|F6NNW3_DANRE_Danio

MTSKDLRDIRAHTLQSPQQHTMADDKMEKAAMDKMVQETEMEEPGLFEQLVQPCMAELVG

TAFFVLMGCLCVIESAQEGHTLQAALVHGLALAVVIGCMVEISGSHFNPSFTIAVFLSGG

LELKMVLPYLISQVSGGLLGAVMAKGMTSSEKYAQAQGAAFTVLQADDHIMKALFAEAAM

TCLATLAVLLSAVNGKSKNHMFPFLVGCTVMVNVLAGANVSGACLNPVRALGPAVLTNYW

THHWIYWVGPITGGLIAAALVRLFLGDNDTRVVMK

>AQP8b_tr|D3U0R1|D3U0R1_DANRE_Danio

MADDKMEKAAMDKMVQETEMEEPGLFEQLVQPCMAELVGTAFFVLMGCLCVIESAQEGHT

LQAALVHGLALAVVIGCMVEISGSHFNPSFTIAVFLSGGLELKMVLPYLISQVSGGLLGA

VMAKGMTSSEKYAQAQGAAFTVLQADDHIMKALFAEAAMTCLATLAVLLSAVNGKSKNHM

FPFLVGCTVMVNVLAGANVSGACLNPVRALGPAVLTNYWTHHWIYWVGPITGGLIAAALV

RLFLGDNDTRVVMK

>AQP1a.2_tr|B6CMI5|B6CMI5_DANRE_Danio

MARELKSWSFWRAVLAEFVGMTIFVFIGIASAIGNKHNRYPDQEVKVALAFGLAIATLAQ

SLGHISGAHLNPAITLGLLVSCQISFFRAFMYIIAQMLGAVLASGIMFKVSPDPDTTLGL

NMLGNGVKVGQGFAIELFTTFQLVLCALATTDKNRTDVSGSAPLAIGLSVGLGHLVAISY

TGCGINPARSFGPAVVLESFKNHWIYWIAPMCGGVAAALIYDFLLFPKREALRKRMNVLK

GTADPDPSATEALIEPRSARSGSGQWPRP

>AQP12_tr|F1QWC9|F1QWC9_DANRE_Danio

MSGLNSSLGFFLAVVGLSVSGRLLLRRWTVLLELVSAFALCACRLEVDTIAEVGQWAGAL

GPDVAVTMLFLSIAVHTAVMQDVSGNPAVTLLRLLQRDVSVVVAVLSIAAQLIGAFLALE

VAGRFWAMELSDMHMITNAEGPCALCGAHAHRLHRKQLHIWICESGPGLCSDLHLPWTLV

PGVFTGLLARTTHWCVSCSLPVFGKYPTAVQQESALLQEKSIPAAERKNQ

>AQP7_tr|Q6PGW9|Q6PGW9_DANRE_Danio

MAPNVGSMLKIKNEYIRVALAESLCTFIMMVFGLGTVAQVVTGEGYFGEYLSINIGFGLA

VAMGVHVGGKVSGAHMNAAVSFTMCVFGRLRWKTLPLYVYAQFLGSFLAAGTIFSLYYDA

INHFCGGNLTVSGPKATAGIFATYPAPYISVYTGFFDQVAGTGLLLLCLMALSDQRNQPL

VSGGEAVGVGLLVMLIGISMGSNSGYAINPTRDLGPRLFTLIAGWGTEVFRAGNCWWWVP

LVAPFIGGVLGALIYKALVELHHPDLKSTTTRPAVDPECIPLDKCKNGRIEIPV

>AQP8a.2_tr|A1L2B1|A1L2B1_DANRE_Danio

MPVEAEKLELEELDKTLLNRDKPKPQSKYERIFQPCIAELVGTTFFVFIGCVSVIENVEA

AGRLQPALVHGLAVAVLVACMAEISGSHFNPPFTIAIWLCGGMQLTMVVPYLISQLIGGV

LGAAMSKVMTSDENYANATGAAFAVLKSDEQLGKVVFAEMAMTCLVTLVVLMGAVNGKSK

SPMVPFMVGCTVIVNILAGGDVSGTCLNPARAFGPALVANHWTYHWVYWVGPLGGGLVAA

ALMRLLLGDEKLRVVMK

>MIPb_tr|A9JRM7|A9JRM7_DANRE_Danio

MWEFRSMMFWRAVFAEFFGTMFFVFFGMGAALRWTTGPYHVFHTALCFGFAAATLIQSIG

HISGGHINPAVTFAYLVGSQMSVFRAFFYICAQCLGAMAGAAALYGVTPNNRRGTMALNT

LQPGMSLGMATTVEVFLTMQLVVCVFAVTDERRNGRLGSAALSIGFSVTMGHLMGMYYTG

AGMNPARSFAPAVITRNFINHWVYWVGPMIGAAMGAIFYDFFLFPRMRGFSERLATLKGS

RPPEAENQQETRGEPIELKTQTL

>MIPb_tr|Q4KMJ7|Q4KMJ7_DANRE_Danio

MWEFRSMMFWRAVFAEFFSTMFFVFFGMGAALRWTTGPYHVFHTALCFGFAAATLIQSIG

HISGGHINPAVTFAYLVGSQMSVFRAFFYICAQCLGAMAGAAALYGVTPNNMRGTMALNT

LQPGMSLGMATTVEVFLTMQLVVCVFAVTDERRNGRLGSAALSIGFSVTMGHLMGMYYTG

AGMNPARSFAPAVITRNFINHWVYWVGPMIGAAMGAIFYDFFLFPRMRGFSERLATLKGS

RPPEAENQQETRGEPIELKTQTL

>AQP12_tr|Q502T6|Q502T6_DANRE_Danio

LSVRRLIMSGLNSSLGFFLAVVGLSVSGRLLLRRWTVLLELVSAFALCACRLEVDTIAEV

GQWAGALGPDVAVTMLFLSIAVHTAVMQDVSGNPAVTLLRLLQRDVSVVVAVLSIAAQLI

GAFLALEVAGRFWAMEMSDMHMIKNLMMSECSTSLRVSTALGVSTEALAALLLLLLHLVL

KNRSQMLKVPALSVALTLIAYTANNYTSGYVNPALAYAVTLTCPGHSFLVYSLVYWLGPL

IGVFLAVFLYLGNIPLLFSKNLLYSKKNRFRLPKGKTNDEKSS

>tr|A0A0G2KRN0|A0A0G2KRN0_DANRE_Danio

ILTDILSILFLRDVFCEFLGTVFFLFISLSSAIHEPLPTLDSSLASTPDPLHVSLAFGVS

VARAGVCLGEVHLNPVITLALVAGLRVSPWRGVLLVGAQLLAALSACAILLVIAPTTQRN

KLFFNPLLHKSKCGPGGVSVSGSVSLVLCVQAATHPKSAFSSNPPAVTGLSDTLGHLMAI

GFTGCGMNPARSFGPAVLTMNFHNHWVYWVGPCSGSLLTWFLHDLLLRPCWSCFGD

>tr|A9ULS6|A9ULS6_DANRE_Danio

MDKMVQETEMEEPGLFEQLVQPCMAELVGTAFFVLMGCLCVIESAQEGHTLQAALVHGLA

LAVVIGCMVEISGSHFNPSFTIAVFLSGGLELKMVLPYLISQVSGGLLGAVMAKGMTSSE

KYAQAQGAAFTVLQADDHIMKALFAEAAMTCLATLAVLLSAVNGKSKNHMFPFLVGCTVM

VNVLAGANVSGACLNPVRALGPAVLTNYWTHHWIYWVGPITGGLIAAALVRLFLGDNDTR

VVMK

>AQP12_tr|Q6DGW2|Q6DGW2_DANRE_Danio

MSGLNSSLGFFLAVVGLSVSGRLLLRRWTVLLELVSAFALCACRLEVDTIAEVGQWAGAL

GPDVAVTMLFLSIAVHTAVMQDVSGNPAVTLLRLLQRDVSVVVAVLSIAAQLIGAFLALE

VAGRFWAMELSDMHMITNAEGPCALCGAHAHRLHREQLHIWICESGPGLRCDLHLSWTLV

PGVFTGLLARTTHWCVSCSLPVFGKYPTAVQQESALLQEKSLPAAERKNQ

>tr|A0A0G2L4D1|A0A0G2L4D1_DANRE_Danio

MAIKEELRSRQFWQGILAEVLGSLVFVSAVLGSLVPGPDGVSPGPIYPALAAGMATVVLG

YCFGEISGAQVNPAVTVALLATRKVDVLRAVVYLVAQCLGGILATGLMYLSLPLKSTAQN

YINKVPVEMNAGQALGMEMLATFLLGFTVFSVEDQRRREINEPGNLAIGLAVTTAIFIAG

RFSGASLNPARSLGPAIILGYWEHHWASKTLWP

>AQP11_tr|Q502B5|Q502B5_DANRE_Danio

FLNTIIRSEDGEMADLTVSLSILVGIVVLSEFARRTALYLFPNREWIIYMLEFISTFQLC

ACTHELKLLAELGGLDPQTGLTLTFIISVVHGFSFRGAICNPTGALELLSRGTLPWGCAL

ARVSCQLTAAVVSRWVMPVAWALALSDLHQRHSLTGFRCSSSPVNAPVLQAAAVELSCAF

VMHTAVNNAEKLEEKYRVPAVAALITTLVYAGGRLTGAVFNPALAFSIQFPCPGNTFTEY

SFVYWMGPILGMSASLLLSDKLIPAISGKSTIPQQLNSNGLKKKKMK

***Three spine stickleback (Gasterosteus aculeatus),***

>tr|G3PAN7|G3PAN7_GASAC_Gasterosteus

MINIRDSSMSGLNASLGYFLSAVVSAALVRALLGRWQRLGFVVELASSFALVACRLEVQT

IVEVGEWAGGLGPDVTLTVLFLVLLTHGVVCGAASGNPSVAALRFLQLEASFPATLLAVG

AQFLGAHLALRVAAYYWSLELTDMHMIKNLMAAECSTSLLVSLVQGFITECVCSLVLHLV

HLNLGRRSALIRVPLTAVLLTFLSHIAKGYTSAYANPSLSYGLTFHCSGFSFSEYALVYW

LGSLTGMFLALLLHAGHIPRISAKNLLYGQKTRFRVPRSDKGEKKKK

>tr|G3P7X0|G3P7X0_GASAC_Gasterosteus

MGRHKFYLDKLSRFFQIRNLLLRQALAECLGTLILVMFGCGSVAQLVLSGGSHGMFITVN

FAFGFAATLGILVCGQVSGGHLNPAVTFALCLLGREPWRKFPMYFLFQTIGGFFGAAIIF

GMYYDALWDHPGSYEVSGPNATAGIFATYPGNHLTIVNGFFDQIIGTAALIVCVLAIVDP

YNNPIPQGLEAFTVGFVVLVIGLSMGFNSGYAVNPARDLGPRLFTAIAGWGVEVFTVRNG

WFLVPVCAPFLGTIVGVVIYQLMVGFHVEGEARDFKSKLEESVALTDVPKNEKTNEKNKN

MH

>tr|G3PTE8|G3PTE8_GASAC_Gasterosteus

PSSYFSYLNNFEYTFNSQRVIIKEFLAEFLGIFVLILFGCGSVAQTVLSKGALGEPLTIH

IGFTLGVMMAVYVAGGVSGAHVNPAVSLAMVILGKLPLKKFPVYVVAQFLGAFAGSCAVY

GLYYDALMEYTNGEFTVTGANATANIFASYPSKHLSILNGFVDQVIATAALILCILAITD

KRNIGAPKGMEPLCIGLIIMAIGVSMGLNCGYPINPARDLGPRFFTAVAGWGVDVFRAGG

CWWWIPVAGPMVGGAVGAGIYFLFIELHHTEPEKQEEKNIQDKSFKVSFYRIVQASF

>tr|G3PEQ0|G3PEQ0_GASAC_Gasterosteus

HRVVRVRNALLRECMAEILGTFVLLLFGCAAGAQVKTSRESKGQFLSANMAFSVGVMSAT

YLTKGISGAHLNPAVTLSFCVLGQVPWGKLVPYCLSQVLGAYLASALVFLVYYDAIMEFS

GGVLTVYGANETASIFATYPSEYLSLGRSFLDQVVGTGMLMLCILGLEEKRNTPAPSKLI

APIVAAIVLGISISMSANCGAAINPARDLGPRLFTLTAGWGTEVFTCYNYWFWVPLVAPL

VGGVMGTFMYLVFIHWHLPDPDPAESFSSSGDVIKQPSIQQENGAELKATHL

>tr|G3PZE0|G3PZE0_GASAC_Gasterosteus

MREFMSKNFWRAVLAELVGMTLFIYLSISAAIGNKNNSSPDQEVKVSLTFGLAIATLAQS

LGHISGAHLNPAVTFGMLASCQISLFKAVMYVVAQMLGSALASGIVYGARPSTTDALGLN

ALNGVTPSQGVGIELLATFQLVLCVIAVTDKRRRDVTGSAPLAIGLSVCLGHLAAISYTG

CGINPARSFGPALILSDFTNHWVYWVGPMCGGLAAALIYDFLLFPKYDDFPERMKVLVSG

PAGNYDVNGGTDNATVEMTSK

>tr|G3P1X7|G3P1X7_GASAC_Gasterosteus

GRRKVFVERANKLLKKRNELVRVGLAESLCTYVMMVFGLGSVAQVVTGQGEFGQYISINL

GFGLGVAMGVHVGGKVSGAHMNAAVSFTMCTFGRLGWKMLPLYVFAQLFGSFLAAATIYG

VYYDAEAIHDYCGGNLTVTGPKATAGIFATYPAPYLSLHAGFIDQVIGTAMLLLCLMALS

DQKNKPAAAGSEPVAVGLLVLLIGISLGSNSGYAINPTRDIAPRVFTAIAGWGGDVFRSG

NGWWWVPLVAPPIGGILGAGLYKVLVEMHHQPEQKERLVEEFESSGIKVKDQI

>tr|G3NKH8|G3NKH8_GASAC_Gasterosteus

ILLETCGMERLLRKCRIRSQLVRECMAECLGVYVLILFGCGSVAQVTTTEDKKGQYLSIN

LGFAMGVTFGVFVSRGVSGAHLNPAVTLSLCVLGRHPWMKLPFYAFFQALGAFLAAATVG

LQYYDAIRVYGGGELTVTGPTATAGIFATYPADYLSVWGGVVDQVIGTAALLLCVLALGD

QKNSPVPDGLQPVLVGAAVLVIGISMGSNSGYALNPARDFGPRLFTYIAGWGVEVFEAGG

GWWWVPIVAPCGGALLGTLIYELMIEVHHPLRLAEEQMPRQEPTEGKMGLELEEDCE

>tr|G3PZC8|G3PZC8_GASAC_Gasterosteus

MSEIKAWGFWRAVLAECLGTIVFVFVFIGLSAAIGDRNNSYPDQEIKVALAFGLAVATLA

QCLGHVSAAHLNPAITLGLLVSCKISPLRALVYMSAQVVGAVAGSVIVYGTRPETTHSLG

VNKLNGVGPGQGFGVEFLLTLQLVLCVLAVTDQRRDVGGFAPLAVGLSVALGPLGHLAGI

SFTGCGINPAQSFGPAILQDSFDNHWVYWAGPMCAGVVAALLYDYLLAPRKDPCGEKTRG

LLCRGSKQEKDHREPLLEEVSVHDLWRARGGL

>tr|G3PIK5|G3PIK5_GASAC_Gasterosteus

LTSAGFQSSATRSLSRCNCQSIMVAFKGVWTKDFWRAVSGEYLATLIFVLLGLGSTINWA

AGEEKPPPADLVLISLCFGLSIATMVQCFGHISGGHINPAVTAAMVVTRKLSLAKGVFYV

LAQCLGAVTGAAVLYLVTPAAVRGSFGVTAVNPNISVGHGLLVELLITFQLVFTIFATCD

PKRSDLGGSASLSIGFAVAIGHLFAIPYTGASMNPARSFAPAMLTLNFENHWVYWVGPVL

GGILAAGLYEYLYCPDPVMKKRLKQVFQKAPSGKYREVEADDISKAIHSTEVEKKESTGE

VLSSV

>tr|G3PIL1|G3PIL1_GASAC_Gasterosteus

SLSRCNCQSIMVAFKGVWTKDFWRAVSGEYLATLIFVLLGLGSTINWAAGEEKPPPADLV

LISLCFGLSIATMVQCFGHISGGHINPAVTAAMVVTRKLSLAKGVFYVLAQCLGAVTGAA

VLYLVTPAAVRGSFGVTAVNPNISVGHGLLVELLITFQLVFTIFATCDPKRSDLGGSASL

SIGFAVAIGHLFAIPYTGASMNPARSFAPAMLTLNFENHWVYWVGPVLGGILAAGLYEYL

YCPDPVMKKRLKQVFQKAPSGKYREVEADDISKAIHSTEVEKKESTGEVLSSV

>tr|G3NRZ5|G3NRZ5_GASAC_Gasterosteus

MRQHCAIKHGILKEFLAEFLGTFVLVLFGCGSVAQTVLSRNSLGEPLTVHIGFSVGLMMA

AYVAGGVSGGHVNPAVSLAMVVLGKLKIWKFPFYVIAQFLGAFAGAAAVFGLYYDAFMDF

TSGILSVTGINATGHIFASYPARHLSVLGGFIDQVVGTGMLVLCILAIIDGENIGAPKGV

QPLAIGLIIMAIGVSMGLNCGYPLNPARDLGPRLFTAAAGWGMEVFSTANNWWWIPVAGP

MVGGVLAAAIYYLLIEVHHHREAPEKPHSSLKDKYEMITMS

>tr|G3PD76|G3PD76_GASAC_Gasterosteus

IFERLFQPCLGELVGTTFFVFIGCVSVIENVESTGRFQPALVHGLAVAVMVACMAEISGS

HFNPPFTIGIYLCGGMEMSMVAPYIASQLVGGVLGAAMAKAMTSSENFAKAHGAAFALLQ

PDDEVAGAVFGEVAMTCLVTMVVLLGAVNAKTRSPLVPFMVGCTVVINILAGGDVSGTCL

NPARAFGPAVVSNYWVRHWVYWMGPITGGLIAAALVRLLLGDRKMRLIFK

>tr|G3PNF8|G3PNF8_GASAC_Gasterosteus

MEKGNKPPAARPPNKYETLFQPCLAEVVGTMFFVFVGCVSVIENVPAAGRLQPALVHGLA

VAVMVAVMDNISGSHFNPPFTIAIYLCGGMKLMMVGPYLVSQLIGGVLGAGMAKMMTPRD

RYLNATGAAFDILKSESQLSGAIFGEVAMTCLITMVVLLVAVNGKTKTPLAPFLVGCTVI

INVLAGGDVSGTCLNPARAFGPALMTNYWSYHWVYWVGPIGGGLLAAALLRLILGDDKIR

VVMKS

>tr|G3P7V1|G3P7V1_GASAC_Gasterosteus

NAFMRSMKQLINTTMIRNLLLRQALAECLGTLILVMFGCGSVAQLVLSGGSHGMFITVNF

AFGFAATLGILVCGQVSGGHLNPAVTFALCLLGREPWRKFPMYFLFQTIGGFFGAAIIFG

MYYDALWDHPGSYEVSGPNATAGIFATYPGNHLTIVNGFFDQIIGTAALIVCVLAIVDPY

NNPIPQGLEAFTVGFVVLVIGLSMGFNSGYAVNPARDLGPRLFTAIAGWGVEVFTVRNGW

FLVPVCAPFLGTIVGVVIYQLMVGFH

>tr|G3PTF0|G3PTF0_GASAC_Gasterosteus

EKMEDESKRKMKERFGLRQDIFKEFLAEFLGIFVLILFGCGSVAQTVLSKGALGEPLTIH

IGFTLGVMMAVYVAGGVSGAHVNPAVSLAMVILGKLPLKKFPVYVVAQFLGAFAGSCAVY

GLYYDALMEYTNGEFTVTGANATANIFASYPSKHLSILNGFVDQVIATAALILCILAITD

KRNIGAPKGMEPLCIGLIIMAIGVSMGLNCGYPINPARDLGPRFFTAVAGWGVDVFRAGG

CWWWIPVAGPMVGGAVGAGIYFLFIELHHTEPEKQEEKNIQDKYEMITLS

>tr|G3PR51|G3PR51_GASAC_Gasterosteus

TMWEFRSMNFWRAVFAEFFGTMFFMFFGMGAALRWTTGPHHVLHVALCFGLAAATLIQSI

GHISGGHINPAVTFAYLVGSQMSLFRAVFYIVAQCLGAVAGAAVLYGVTPGNMRGNMAMN

TLQPGISLGMGTTVEVFLTMQLVVCIFAVTDERRNGRMGSAALSIGFSVTIGHLMGMYYT

GAGMNPARSFAPALLFRNFLNHWVYWVGPMIGGAMGALLYDFMLFPRMRGLSERLAALKG

SRPPENEAQQDGRGEPIELKTQAL

>tr|G3P0W1|G3P0W1_GASAC_Gasterosteus

MWEFKSMSFWRAVFAEFYGTMFFVFFGLGAALRWTTGPHNVLHVAFCFGLAAATLIQSIG

HISGGHINPAVTFAYLIGSQMSLFRAFFYIVAQCLGALAGAAVLYGVTPTNMRGNLALNT

LQPGISLGMATTMEVFLTLQLVVCVFAVTDERRNGRLGSAALAIGFSVLVGHLLGMYYTG

AGMNPARSFAPAVLVRNFVNHWVYWVGPMIGGALGALLYDFMLFPRVRGLSERLATLKGI

RPSEAEGQQETRGEPIELKTQAL

>tr|G3QCG6|G3QCG6_GASAC_Gasterosteus

TAAVRRHTTGSMSDAYVSLAVLGAAVLLSEATRVAAARLLSGVHRTRVLEAASTFQLCCC

THELKLLGEAAPLAPPLGLALTYAVTVVHLGTFRGATCSPIGAAESVCRGKLVACQFAAA

VAAQFFATAVWSLGLSEVHVRHQRFGFRCFDPLGGTLREAAAVELGCAFAVQAAAMHAGA

LDENLRAHFLAAVIPAVFSAGGGISGAVFNPVLAFSVQFPCSGHTYPEYCFVYWLGPVLG

VASCVLLFEKVVPFLSGKTPVAPDVPVVQKQKTQ

>tr|G3PR48|G3PR48_GASAC_Gasterosteus

MWEFRSMNFWRAVFAEFFGTMFFMFFGMGAALRWTTGPHHVLHVALCFGLAAATLIQSIG

HISGGHINPAVTFAYLVGSQMSLFRAVFYIVAQCLGAVAGAAVLYGVTPGNMRGNMAMNT

LQPGISLGMGTTVEVFLTMQLVVCIFAVTDERRNGRMGSAALSIGFSVTIGHLMGMYYTG

AGMNPARSFAPALLFRNFLNHWVYWVGPMIGGAMGALLYDFMLFPRMRGLSERLAALKGS

RPPENEAQQDGRGEPIELKTQAL

>tr|G3P0X0|G3P0X0_GASAC_Gasterosteus

YKKASELPLLCMWEFKSMSFWRAVFAEFYGTMFFVFFGLGAALRWTTGPHNVLHVAFCFG

LAAATLIQSIGHISGGHINPAVTFAYLIGSQMSLFRAFFYIVAQCLGALAGAAVLYGVTP

TNMRGNLALNTLQPGISLGMATTMEVFLTLQLVVCVFAVTDERRNGRLGSAALAIGFSVL

VGHLLGMYYTGAGMNPARSFAPAVLVRNFVNHWVYWVGPMIGGALGALLYDFMLFPRVRG

LSERL

***Tilapia (Oreochromis niloticus),***

>tr|I3J3X0|I3J3X0_ORENI_Oreochromis

MSGLNASVGYLVAAVTFVVSVRFLLKKWPRFSFISEFTSSFMLVACWLEVQTIVEVGEWA

GGLGPDVTLTILFVVLLTHGIICGDASGNPTLAVQKFLKLEATTVTTLLDVAAQFIGAHL

GLLAAKYYWSLELTDMHMIKNLMVSECSTSLLVSVVQGFFTEFACALVLHLIHLNLCRRS

ALIRVPLIAVLLTFLSHSAKGYTSAYMNPSLAYGLTFHCSGFTFAEYAVVYWLGSLTGMT

LAVLLYMGHIPRIFAKNLLYSQKTRFRVPKTDKAEKKKQ

>tr|I3JT38|I3JT38_ORENI_Oreochromis

VRPCGPVKSREMADLWVSVAVLGVSVLLSELIRSAARLFQRDYRIYLLEVASTFQLCCCT

HELKLLGDSAQLELSYGLALTYTVTLVHLATFQDATCNPTAALESVCRGTRSLRAAGVLI

MLQFAAAVAAQSYVASVWSQGLSDIHLRHQSSDFRCFDPLGGTLLEAAVVELACAFVFQA

ACMHVHKLDARLQVHFIAAVVTAAVYAGGEISGAIFNPVLAFSVQFPCSGHTYLEYCFVY

WLGPVLGVVSCILLFEKIIPFLFGGSTAEKED

>tr|I3JGA2|I3JGA2_ORENI_Oreochromis

MLKLRGMTVRCPLVRECMAEFLGTFVLMLFGCAAAAQVKTSNETKGQFLSGNMAFSVGVM

SAMYLTKGISGAHLNPAVTLSFCVLGKVSWSRLVPYSLSQVVGAFVASGVVYLVYYDAIM

TFSGGVLTVYGRNETASIFATYPSDYLSVGRSFFDQVVGTSMLLLCILGLEEKRNTPAPS

ELVPVVVAGVVLGISISMSANCGAAINPARDLGPRLFTLAAGWGTEVFTCYNYWFWVPVV

APLVGGVFGSLMYLIFIDWQLPDPDQLERISTISEKLAKKQPVTTLEKGDELKAMHF

>tr|I3JEP6|I3JEP6_ORENI_Oreochromis

MENEYRRKMKEKLGLRRDIFKEFLAEFLGIFFLILFGCGSVAQMVLSRAGPGDILSVHIG

FTLGVMMAVYIAGGVSGAHVNPAVSLAMLILGKLPLKKFPIYVAAQFLGAFAGSCAVYGL

YYDALMDFTKGEFIVTGENATATIFASYPAKHLSVLNGLGDQVIATAALVICILAITDRK

NIGAPKGMEPLCIGLIIAAIGVSMNLNCGYPINPARDLGPRFFTALAGWGMEVFRAGGCW

WWIPVVGPMVGGAVGAGVYLIFIELHHPEPEKQEENNVQDKYEIVTMT

>tr|I3JSM5|I3JSM5_ORENI_Oreochromis

MREFKSKAFWRAVLAELVGMTLFIFLSIATAIGNTNNANPDQEVKVSLAFGLAIATLAQS

LGHISGAHLNPAVTLGMLASCQISVFKAIMYIISQMLGSALASGIVYGTRPDNITVLGLN

ALNGVTPSQGVGIELLATFQLVLCVIAVTDKRRRDVTGSAPLAIGLSVCLGHLAAISYTG

CGINPARSFGPALILNNFTNHWVYWVGPMCGGVAAALLYDFLLSPKFDDFPDRLKVLVSG

PVGDYDVNGGNDATTVEMTSK

>tr|I3KTR0|I3KTR0_ORENI_Oreochromis

VEVGVFQQRGGKVTRPKVWLKNELIRVGLAESLSTYVMMSLGLGSVAQVVTGQGAFGQYL

SINLGFGLAVAMGSHVGGKISGAHMNGAVSFTMCVFRRLPWKMLPLYISAQLLGSFLAAG

TIYAVYYEAIHDYCGGNLTVTGEKATAGIFATYPAPYLSLIAGFFDQVFGTAMLLLCLMA

LSDQKNKPAPAGSEPAFVGFLVLLIGISLGSNSGYAINPTRDIAPRVFTAMAGWGTDVFR

VGNGWWWVPLVATPIGGVLGAGLYKAVVELQHPHLSEAGGEMVEEAVPLDKEINTSENVC

V

>tr|I3KUR4|I3KUR4_ORENI_Oreochromis

MERLLKKCQIRNQLIRECMAECLGVYVLILFGCGSVAQVVTTEDKKGQYLSINLGFALGV

TFGVFVSRGVSGAHLNPAVSLSLCVLGRHSWMKLPFYIFFQVLGAFLAAATVALQYYDAI

QAYSGGDLTVTGPKATAGIFCTYPADYLSVWGGIVDQVIGTAALLLCVLALGDQRNSSIP

HYLQPVLVGAVVLVIGISMGSNSGYALNPARDFGPRLFTYIAGWGADVFKAGSGWWWVPI

VAPCVGALLGTLIYELMVEVHHPSEQSESQASCPENTNDKMGVELEGVEADREKPT

>tr|I3KUV2|I3KUV2_ORENI_Oreochromis

MFHDKTNHLTSSSYISWEYKQLEEGGASQRKNVYVRMSLLHHWLIYRTVCRSYCSPPVEV

EGVKHRAGGFKYSMSGLESKTEVFTVTEMGEPEVEKDIGNRSKIQSKYELYVQPCLAELL

GTCLFVFVGCSSVIGNVETGGVIQPALAHGLALGVLITVFGQISGGHFNPAVSLSIYLCG

GMKLILLVPYIVAQLAGGVVGAFLSKAVYPSNNYTASLGGAFKAVSADAGRSTLVEMLLT

LILTMVVCLGAVNRKTQTDWVPFCIGLTVAANIFAGGTLSGACMNPARAFGPAMAANHWD

NHWVFWVGPVAGALLTVSIIRLLIGDHKTRVVLK

>tr|I3JSM3|I3JSM3_ORENI_Oreochromis

MTEVKSWAFWRAVAAEFVGMLLFILIGLSSIVGIGLGNDKNQMIAQEVKVSLAFALAIAT

LAQSLGHISGAHLNPAVSLGLLVNCQISALRCAFYILAQMLGAVAASAIVNGYKPGESLG

VNALNVSVRAGFAIEFFATLQLVLCIIAVTDKRRTDVTGSAPLAIGLSVGLGHLTAISFT

GCGINPARSFGPALILGKMKNHWVYWLAPMCGGIAASLIYDFLLYPQTRNFRSRIGILVH

GPQEEYVERIGEENNSPGPSHWPKQ

>tr|I3JJV4|I3JJV4_ORENI_Oreochromis

FPCTPPSQFWPPSCFTQDKAMTAFKGIWTKEFWRCVTAEFSAMLIFVLLGLGSTINWGTV

KEDPHPPDLVLISLCFGLTIATMVQCFGHISGAHINPAVTVAMVVTRKLSLAKAVFYLLA

QCVGAIVGAAVLYGITPASVRGGMGVTEVNESISVGTALVVELFITFQLIFTIFATCDHK

RKDLKGSSALAIGLSVCVGHLFAIPYTGASMNPARSFGPAMVTWSWENHWVYWVGPSMGG

TLAAALYEYLFCPDPEVKKRFSETFVKTPFTAAKQRQDSATAQEPLFTVMDVERAERRER

EVSGEVLSSV

>tr|I3KG53|I3KG53_ORENI_Oreochromis

MGWQKHYLDKLSRFFQIRNLLLRQALAECLGTLILVMFGCGSVAQLVLSGGSHGMFLTVN

FAFGFAATLGILVCGQISGGHLNPAVTFALCLLGRERWRKFPMYFLFQTIGAFFGAAIIF

GMYYDALWDHPGSFNVTGPDATAGIFATYPGTHLTLVNGFFDQIIGTAALIVCILAIVDP

YNNPIPQGLEAFTVGFVVLVIGLSMGFNSGYAVNPARDLGPRLFTAIAGWGSEVFTASPG

WFLVPVFAPFLGTLIGVMIYQLMVGFHMEGEVRDRKESTEQETVRLTNVTSKDNSREAVK

ERNEC

>tr|I3JTM6|I3JTM6_ORENI_Oreochromis

MQSQGTMSGSCSSDRPAFSNNPAHARPAAPLRLLSWCNCQNIMVAFKGIWTKDFWRAVSG

EYLATLIFVLLGLGSTINWAAGEEKPPPADLVLISLCFGLTIATMVQCFGHISGGHINPA

VTAAMVVTRKLSLAKAVFYVAAQCLGAITGAGILYLVTPTAVRGSFGVTTVNPTISVGHG

FLVELLITFELVFTVFATCDPKRTDLGGSASLAIGIAVVIGHLFAIPYTGASMNPARSFG

PAMVTLNFENHWVYWVGPILGGILAAGLYEYLYCPDPEIKQRMKQVFKKDPSGKYTEVET

GGVKPGSIHNISVEKADVMLTGEV

>tr|I3KUV1|I3KUV1_ORENI_Oreochromis

GATMGEEQIEMGDSSLMEPMKTKPVLKPQPNKYERLFQPCLAEIVGTLFFVFIGCVSVIE

NVPEAGRLQPALVHGLAVAVLVAVMDKISGSHFNPPFTVAIYLCGGMDLMMAGLYIVCQL

IGGVVGAGMAKLMTPPERFINATGAAFTILKSDSQLSRAIFGEVAMTCLITLVVLLVAVN

TKTKTPLAPFLVGCTVIINVLAGGDISGTCLNPARAFGPAVVTGYWTYHWVYWVGPIGGS

LVAAALVRIILGDDKLRLIMK

>tr|I3JNP9|I3JNP9_ORENI_Oreochromis

MEVQNKRSMRQHCALKHGIFKEFLAEFLGTFVLVLFGCGSVAQTVLSRNTLGEPLTVHIG

FSVGLMMAAYVAGGVSGGHVNPAVSLAMVILGKLKIWKFPFYIIAQFLGAFAGAAAVFGL

YYDAFMDFTSGILSVTGINATGHIFASYPARHLSVLGGFIDQVVGTGMLVLCILAITDSG

NIGAPKGIEPLAIGLIIMAIGVSMGLNCGYPLNPARDLGPRLFTAVAGWGMEVFSTADNW

WWIPVAGPMVGGVVAAVIYYLFIELHHPHAEHEKPQEEEEEDEEDEEDDDSSLKDKYEMI

TMS

>tr|I3KK79|I3KK79_ORENI_Oreochromis

TMWEFRSMNFWRAVFAEFFGTMFFVFFGMGAALRWTTGPHHVLHVALCFGLAAATLIQSI

GHISGGHINPAVTFAYLVGSQMSLFRAIFYIAAQCLGAVAGAAVLYGVTPGNMRGNMAMN

TLQPGISLGMATTVEVFLTMQLVICIFAVTDERRNGRLGSAALSIGFSVTIGHLMGMYYT

GAGMNPARSFAPAVIFRNFINHWVYWVGPMIGGAMGALLYDFMLFPRMRGLSERLATLKG

SRPPENETQQDTRGEPIELKTQAL

>tr|I3K1N6|I3K1N6_ORENI_Oreochromis

MWEFRSMSFWRAVFAEFYGTMFFVFFGLGAALRWTTGPHNVLHVAFCFGLAAATFIQSIG

HISGGHINPAVTFAYLIGSQMSLFRAFFYIIAQCLGALAGAAVLYGVTPSNMRGNLALNT

LQPGISLGMATTIEIFLTLQLVVCIFAVTDERRNGRLGSAALAIGFSVLMGHLLGMYYTG

AGMNPARSFAPAVLVRNFVNHWVYWVGPMIGGAMGALLYDFMLFPRFRGLSERLATLKGA

RPPEADGQQETRGEPIELKTQAL

>tr|A0A023ZXY1|A0A023ZXY1_ORENI_Oreochromis

MGRQKEYLDKLSRFFQIRNLLLRQALAECLGTLILVMFGCGAVAQRVLSGGSHGLFLTVN

FAFGFAAMLGILVCGQVSGGHLNPAVTFALCLLGRERWRKFPMYFLFQTIGAFFGSAIIF

GMYYDALLLRPGSFNLTSTNNTAGIFATYPARHLTLVNGFFDQIIGTTALIVCVLAIVDP

FNNPIPQGLEAFTVGFVVLVIGLSMGFNSGYAVNPARDFGPRLFTSMSGWGGAVFTARDC

WFLVPIFAPFLGSILGVVIYQLMVGFHTEGEARDKKQGTVQENLQLTNVASSNNSKEATK

EIY

>tr|I3KK78|I3KK78_ORENI_Oreochromis

MTWRELRSRQFWHAMLAELLGTLVLVSAVLGASVPGPGDAPGGPLYPAVAVGVVIVALGH

CFGEISGAQVNPAVTLALLATRRLDVLRALVYIIAQCLGASLGAGALYLALPFKTTAEHF

VNRVPLELNAAKALGIEVLCTFQMVFTVFSVEDQRRRESPEPGNLAIGLAHTAGVLIGAR

FSGASMNPARSLGPAIITGFWEDHWVYWIGPVLGAILAGMSHEFFFARSASRQKLVACLT

CKDIEIVETASMTGSSLSTVTQNAMRAKQASKQENN

***Mudskipper (Boleophthalmus pectinirostris)***

>XP_020791255.1_Boleophthalmus

MREFKSKDFWRAVLAELVGMTLFIFLSLSTAIGNVNNSNPDQEVKVALGFGLAIATLAQSLGHISGAHLNPAVTLGMLASCQISAFKAVMYIIAQMLGSALASGIVYGTKPNTTAALGLNSLAGVTPSQGVGIELLATFQLVLCVIAVTDKRRRDVTGSAPLAIGLSVCLGHLAAISYTGCGINPARSFGPAVILNEFTNHWVYWVGPMCGGVAAALIYDFLLYPKDDFPERMKVLVSGPIGDYDVNGGNDATTVEMTSK

>XP_020791256.1_Boleophthalmus

MRELKSGAFWRAVAAEFIGMLLFIFFGIGSAIGNPSKDSQEVKVSLAFGLAIATLAQSLGHVSGAHLNPAVSIGLLVSCQISFLRFLCYTVAQMLGAVAASALLNGFRPRQIGLGVNELGDISLGQGFAIEFFATLQLVLCVIAVTDKRRSDVQGSAPLAIGLSVTLGHLVAISYTGCGINPARSFGPAIIRNQMKYHWVYWLGPMCGGIAAAFIYDLILAPRSPSLSARKDILLYGPDDDVDNNLLREGESPGPSQWPKQ

>XP_020785352.1_Boleophthalmus

MMWEFRSMSFWRAVFAEFYGTMFFVFFGLGAALRWTTGPHNVLHVAFCFGLAAATLIQSIGHISGGHINPAVTFAYLIGSQMSLFRAFFYIIAQCLGALAGAAVLYGVTPTNMRGNLALNTLQPGISLGMATTMEIFLTVQLVICIFAVTDERRNGRLGSAALAIGFSVLMGHLLGMYYTGAGMNPARSFAPAVLVRNFVNHWVYWVGPMIGGAIGALVYDFMLFPRMRGLSERLATLKGIRPPEAQGQPDTRGEPIELKTQAL

>XP_020785343.1_Boleophthalmus

MWEFRSMNFWRAVFAEFFGTMFFVFFGMGAALRWTTGPHHVLHVALCFGLAAATLIQSIGHISGGHINPAVTFAYLVGSQMSLFRAVFYIAAQCLGAVAGAAVLYGVTPGNMRGNMAMNTLQPGISLGMGTTVEVFLTMQLVVCIFAVTDERRNGRLGSAALSIGFSVTIGHLMGMYYTGAGMNPARSFAPAVLFRNFINHWVYWVGPMIGGAMGALLYDFMLFPRMRGLSERLATLKGSRPPESENQQDPRGEPIELKTQAL

>XP_020777769.1_Boleophthalmus

MVAFKGVWTKGFWRAVSAEYLATLIFVFLGVGSTINWHATKENPPPADLVLISLCFGLSIATMVQCFGHISGGHINPAVTAAMVVTGKLSLAKALFYVLAQCLGAITGAGLLYLVTPPAVRGSFGVSDVNTDISVGLGLLVEIIITFQLVFTVFATCDPKRTNLSGSAALAIGVAVVIGHLFGIPYTGASMNPARSFGPAVVTVTFKNHWIYWLGPLLGGILAAGLYEYLFCPDPEVKKRLQQVFQKDQSGRYKEVDADDITIKPGSIDVEKGEKKDLFNDKGDVLSSV

>XP_020785193.1_Boleophthalmus

MKDFVPMVIHNKLFLHSRPFWRAVLAELLGTLVLVSAVLGSLVPGPGEASGGPLYPALAVGVAIVAVAHCFGEISGAQVNPAVTLALLATRRMDILRAVFFICAQCVGACLGTGALYLALPLKTTADHFINKVPLELNAAQALCVEMLCTFQMVFTVFSAEDQRRRDSPEPGNLAVGFSHTAGVLIGARFSGASMNPARTLGPAIITGFWENHWVYWIGPVLGAVLAGVSHEFFFAPSASRQKLVACLTCKDIEIVETASMTGSSLSTVTQNAARAKQSNKTEN

>XP_020780042.1_Boleophthalmus

MTKTEPEVFTISGSEMGQVNEEAECSRSRGNKHTHIYEQYVQPCLAEFLGSTLFIFVGCACVVGNSAPGAIQPALAHGLALSIVIMLFGQISGGHFNPAVTLCVYLCGGMSLALVVPYILAQMLGGIAGAGLVRAVFNTLDYQAGTGGAFTPEAIPNDLGKITLAELVLTMFLTTTVAMGAVNGKTSSQLAPFGIGLTVAADILAGGGLSGACMNPARAFGPAVVANKWDHHWIYWVGPMAGALCTVIHVRLFLGDRKTRIILRK

>XP_020780043.1_Boleophthalmus

MDVDKMQMEGEELMEDKKKIGTMFFVFVGCVSVIENVPAAGRLQPALAHGLALATLVAVMGNISGSHFNPTFTIAIYLCGGIELTMVGPYLVSQLIGGVLGAGMAKVMTPAERYMNATGAAFDILRPESNSQLYSALFGEMTMTCMLTMVVLLVAVNNKTKTPMAPFLVGCTVIANVLSGGDISGTCLNPARAFGPAVIANYWMYHWVYWAGPISGGLLAAILLRLILGDEKVRLILKS

>XP_020796975.1_Boleophthalmus

MEDENKKRMKEKFGLRRDIFKEFLAEFLGIFVLILFGCGSVAQTVLSKGALGEPLTIHIGFTLGVMMAVYMAGGVSGGHVNPAVSLAMVILGKLSVKKFPIYVAAQFLGAFVGSCAVFGLYYDALMEYTNGEFMVTGPNATANIFASYPAKHLSVLNGFIDQVIGTAALILCILAITDKRNIGAPKGMEPLCIGLAITAIAVSMGLNCGYPINPARDLGPRFFTAVAGWGMDVFRAGGCWWWIPVAGPMVGGAVGAGVYFLCVELHHPEPEKQAENNIPDKYEMVTMS

>XP_020783751.1_Boleophthalmus

MMQPRFMRVRCGLIRECMAEFLGTFVLLVFGCSAAAQVKTSRETKGQFLSVNMSFAVGVMSAMYLTKGISGAHLNPAVSLSFCILGQVPWNRLVPYSLSQLLGAYTASALVYFMYYDAIMDFSGGELTVYGANETASIFATYPTEYLSLSGSFLDQVVGTGMLMLCILCLGEKRNTPAPEQLVPPIVAAIVLGISMSMSANCGAAINPARDLGPRLFTLTAGWGTEVFTCYNYWFWVPAVAPMVGGVLGTLIYLMFIEWQLPDHPPEDPHTDISFSDKPRSSVTWNSKDELKEARF

>XP_020795939.1_Boleophthalmus

MVQSLELGSSVQRKTLKSRIQIQNQVLRLGLAETLCTYVMMVFGLCSVAQVVTGKGEFGHYISINLSFGLAVAMGVHVGGGVSGAHMNAAVSFSSCVFGTLRWRLLPLYITAQFIGSFLAAATVYAVYYEAIQSYCGGNLTVSGPGATAGIFATYPAPYLSLSGGFSDQVFGTAMLLLCLSALSDRRNQPCPAGGEPVAVGLLVLLIGMSAGSNSGYAINPTRDIGPRVFTAVAGWGPDVFRYDPRPPSGARTLTFDL

>XP_020790536.1_Boleophthalmus

MRRHCGLKYGICKEFLAEFLGTFVLVLFGCGSVAQTVLSRNQLGEPLTVHIGFSVGLMMGVYVAGGVSGGHVNPAVSLAMVILGKLKIWKFPIYVIAQFLGAFAGAAAVFGLYYDAFMDFTSGILSVTGINATGHIFASYPARHLSVLGGFIDQVVGTGMLVLCILAIIDGGNIGAPKGMEPLAIGLIIMAIGVSMGLNCGYPLNPARDLGPRLFTAVAGWGFEVFSTAGYWWWIPVAGPMVGGVVAAVLYYLLIEMHHKRDTPEKTDEEEEEEEDDEDEDSSLKDKYEMILMN

>XP_020791696.1_Boleophthalmus

MGAQKVALEKLSHFFQLRNLLLRQALAECLGTLILVMFGTGAVAQYVLSGGTHGMFITVNFAFGFAATLGILVCGQISGGHLNPAVTFALCLLGRERWRKLPMFFLFQTIGAFFGAAIIFGMYYDALWDYPGCFNMTGNSSTAGIFATYPGKHLTIVNGFFDQVIGTAALIVCILAIVDPYNNPIPQGLEAFTVGFVVLVIGLSMGFNSGYAVNPARDLGPRVFSAMAGWGVDVFTFRNGWFLVPTFAPFIGTFIGVMVYQLMVGFHVEGEVRDRKAAQQENEKVRLTNVTTNNDHSKELH

>XP_020788176.1_Boleophthalmus

MERILTKCRITNHLVRECLAECLGVYIIILFGCGSVAQVTTTQNQKGQYISINLGFALGVTFGVYVSRGVSGAHLNPAVSLSLCALGRHPWAKLPFYVFFQFLGAFLGAATVGLEYYEAIKWYSGGELTVSGPTATAGIFCTFPAEYLSLWAGFVDQVIGTAALLVCVLALGDEGQGSIPAGIQPVLVGAVVMVIGVSMGSNSGYALNPARDLAPRLFTYFAGWGDDVFR

***Homo sapiens***

>sp|Q96PS8|AQP10_HUMAN Aquaporin-10_Homo

MVFTQAPAEIMGHLRIRSLLARQCLAEFLGVFVLMLLTQGAVAQAVTSGETKGNFFTMFL

AGSLAVTIAIYVGGNVSGAHLNPAFSLAMCIVGRLPWVKLPIYILVQLLSAFCASGATYV

LYHDALQNYTGGNLTVTGPKETASIFATYPAPYLSLNNGFLDQVLGTGMLIVGLLAILDR

RNKGVPAGLEPVVVGMLILALGLSMGANCGIPLNPARDLGPRLFTYVAGWGPEVFSAGNG

WWWVPVVAPLVGATVGTATYQLLVALHHPEGPEPAQDLVSAQHKASELETPASAQMLECK

L

>sp|Q13520|AQP6_HUMAN Aquaporin-6_Homo

MDAVEPGGRGWASMLACRLWKAISRALFAEFLATGLYVFFGVGSVMRWPTALPSVLQIAI

TFNLVTAMAVQVTWKASGAHANPAVTLAFLVGSHISLPRAVAYVAAQLVGATVGAALLYG

VMPGDIRETLGINVVRNSVSTGQAVAVELLLTLQLVLCVFASTDSRQTSGSPATMIGISV

ALGHLIGIHFTGCSMNPARSFGPAIIIGKFTVHWVFWVGPLMGALLASLIYNFVLFPDTK

TLAQRLAILTGTVEVGTGAGAGAEPLKKESQPGSGAVEMESV

>sp|Q92482|AQP3_HUMAN Aquaporin-3_Homo

MGRQKELVSRCGEMLHIRYRLLRQALAECLGTLILVMFGCGSVAQVVLSRGTHGGFLTIN

LAFGFAVTLGILIAGQVSGAHLNPAVTFAMCFLAREPWIKLPIYTLAQTLGAFLGAGIVF

GLYYDAIWHFADNQLFVSGPNGTAGIFATYPSGHLDMINGFFDQFIGTASLIVCVLAIVD

PYNNPVPRGLEAFTVGLVVLVIGTSMGFNSGYAVNPARDFGPRLFTALAGWGSAVFTTGQ

HWWWVPIVSPLLGSIAGVFVYQLMIGCHLEQPPPSNEEENVKLAHVKHKEQI

>sp|P30301|MIP_HUMAN AQP0_Homo

MWELRSASFWRAIFAEFFATLFYVFFGLGSSLRWAPGPLHVLQVAMAFGLALATLVQSVG

HISGAHVNPAVTFAFLVGSQMSLLRAFCYMAAQLLGAVAGAAVLYSVTPPAVRGNLALNT

LHPAVSVGQATTVEIFLTLQFVLCIFATYDERRNGQLGSVALAVGFSLALGHLFGMYYTG

AGMNPARSFAPAILTGNFTNHWVYWVGPIIGGGLGSLLYDFLLFPRLKSISERLSVLKGA

KPDVSNGQPEVTGEPVELNTQAL

>sp|P55087|AQP4_HUMAN Aquaporin-4_Homo

MSDRPTARRWGKCGPLCTRENIMVAFKGVWTQAFWKAVTAEFLAMLIFVLLSLGSTINWG

GTEKPLPVDMVLISLCFGLSIATMVQCFGHISGGHINPAVTVAMVCTRKISIAKSVFYIA

AQCLGAIIGAGILYLVTPPSVVGGLGVTMVHGNLTAGHGLLVELIITFQLVFTIFASCDS

KRTDVTGSIALAIGFSVAIGHLFAINYTGASMNPARSFGPAVIMGNWENHWIYWVGPIIG

AVLAGGLYEYVFCPDVEFKRRFKEAFSKAAQQTKGSYMEVEDNRSQVETDDLILKPGVVH

VIDVDRGEEKKGKDQSGEVLSSV

>sp|O43315|AQP9_HUMAN Aquaporin-9_Homo

MQPEGAEKGKSFKQRLVLKSSLAKETLSEFLGTFILIVLGCGCVAQAILSRGRFGGVITI

NVGFSMAVAMAIYVAGGVSGGHINPAVSLAMCLFGRMKWFKLPFYVGAQFLGAFVGAATV

FGIYYDGLMSFAGGKLLIVGENATAHIFATYPAPYLSLANAFADQVVATMILLIIVFAIF

DSRNLGAPRGLEPIAIGLLIIVIASSLGLNSGCAMNPARDLSPRLFTALAGWGFEVFRAG

NNFWWIPVVGPLVGAVIGGLIYVLVIEIHHPEPDSVFKTEQSEDKPEKYELSVIM

>sp|O14520|AQP7_HUMAN Aquaporin-7_Homo

MVQASGHRRSTRGSKMVSWSVIAKIQEILQRKMVREFLAEFMSTYVMMVFGLGSVAHMVL

NKKYGSYLGVNLGFGFGVTMGVHVAGRISGAHMNAAVTFANCALGRVPWRKFPVYVLGQF

LGSFLAAATIYSLFYTAILHFSGGQLMVTGPVATAGIFATYLPDHMTLWRGFLNEAWLTG

MLQLCLFAITDQENNPALPGTEALVIGILVVIIGVSLGMNTGYAINPSRDLPPRIFTFIA

GWGKQVFSNGENWWWVPVVAPLLGAYLGGIIYLVFIGSTIPREPLKLEDSVAYEDHGITV

LPKMGSHEPTISPLTPVSVSPANRSSVHPAPPLHESMALEHF

>sp|P41181|AQP2_HUMAN Aquaporin-2 _Homo

MWELRSIAFSRAVFAEFLATLLFVFFGLGSALNWPQALPSVLQIAMAFGLGIGTLVQALG

HISGAHINPAVTVACLVGCHVSVLRAAFYVAAQLLGAVAGAALLHEITPADIRGDLAVNA

LSNSTTAGQAVTVELFLTLQLVLCIFASTDERRGENPGTPALSIGFSVALGHLLGIHYTG

CSMNPARSLAPAVVTGKFDDHWVFWIGPLVGAILGSLLYNYVLFPPAKSLSERLAVLKGL

EPDTDWEEREVRRRQSVELHSPQSLPRGTKA

>sp|Q8NBQ7|AQP11_HUMAN Aquaporin-11_Homo

MSPLLGLRSELQDTCTSLGLMLSVVLLMGLARVVARQQLHRPVAHAFVLEFLATFQLCCC

THELQLLSEQHPAHPTWTLTLVYFFSLVHGLTLVGTSSNPCGVMMQMMLGGMSPETGAVR

LLAQLVSALCSRYCTSALWSLGLTQYHVSERSFACKNPIRVDLLKAVITEAVCSFLFHSA

LLHFQEVRTKLRIHLLAALITFLVYAGGSLTGAVFNPALALSLHFMCFDEAFPQFFIVYW

LAPSLGILLMILMFSFFLPWLHNNHTINKKE

>sp|P55064|AQP5_HUMAN Aquaporin-5_Homo

MKKEVCSVAFLKAVFAEFLATLIFVFFGLGSALKWPSALPTILQIALAFGLAIGTLAQAL

GPVSGGHINPAITLALLVGNQISLLRAFFYVAAQLVGAIAGAGILYGVAPLNARGNLAVN

ALNNNTTQGQAMVVELILTFQLALCIFASTDSRRTSPVGSPALSIGLSVTLGHLVGIYFT

GCSMNPARSFGPAVVMNRFSPAHWVFWVGPIVGAVLAAILYFYLLFPNSLSLSERVAIIK

GTYEPDEDWEEQREERKKTMELTTR

>sp|Q8IXF9|AQ12A_HUMAN Aquaporin-12A_Homo

MAGLNVSLSFFFATFALCEAARRASKALLPVGAYEVFAREAMRTLVELGPWAGDFGPDLL

LTLLFLLFLAHGVTLDGASANPTVSLQEFLMAEQSLPGTLLKLAAQGLGMQAACTLMRLC

WAWELSDLHLLQSLMAQSCSSALRTSVPHGALVEAACAFCFHLTLLHLRHSPPAYSGPAV

ALLVTVTAYTAGPFTSAFFNPALAASVTFACSGHTLLEYVQVYWLGPLTGMVLAVLLHQG

RLPHLFQRNLFYGQKNKYRAPRGKPAPASGDTQTPAKGSSVREPGRSGVEGPHSS

>sp|P29972|AQP1_HUMAN Aquaporin-1_Homo

MASEFKKKLFWRAVVAEFLATTLFVFISIGSALGFKYPVGNNQTAVQDNVKVSLAFGLSI

ATLAQSVGHISGAHLNPAVTLGLLLSCQISIFRALMYIIAQCVGAIVATAILSGITSSLT

GNSLGRNDLADGVNSGQGLGIEIIGTLQLVLCVLATTDRRRRDLGGSAPLAIGLSVALGH

LLAIDYTGCGINPARSFGSAVITHNFSNHWIFWVGPFIGGALAVLIYDFILAPRSSDLTD

RVKVWTSGQVEEYDLDADDINSRVEMKPK

>sp|O94778|AQP8_HUMAN Aquaporin-8_Homo

MSGEIAMCEPEFGNDKAREPSVGGRWRVSWYERFVQPCLVELLGSALFIFIGCLSVIENG

TDTGLLQPALAHGLALGLVIATLGNISGGHFNPAVSLAAMLIGGLNLVMLLPYWVSQLLG

GMLGAALAKAVSPEERFWNASGAAFVTVQEQGQVAGALVAEIILTTLLALAVCMGAINEK

TKGPLAPFSIGFAVTVDILAGGPVSGGCMNPARAFGPAVVANHWNFHWIYWLGPLLAGLL

VGLLIRCFIGDGKTRLILKAR

**CLAUDINS**

***Round goby (Neogobius melanostomus)***

*(the ones with an asterix (*) are manually curated from long, fused gene models)*

>NEME_00008799_1*_Neogobius

MVSSGFQMLGAALATIGWIGAIVICALPMWNVTAFIGSNIVTSQTIWEGIWMNCVVQSTG

QMQCKVYDSMLALSSDLQAARALTIIAIIVGILGILLAVAGGQCTNCVEDKQAKAKVGIA

AGVVFIIAGVLVLVPVCWTAHTIIRSFYNPLMIDAQKRELGAALYIGWGAGALLLIGGAM

LCSGCPKTEKEGYTARY

>NEME_00008799_2*_Neogobius

MAASSEAPSTRPKDYRGETMASQGLQLLGVLLALVGDYHHLRL

PMWRVTAFVGANIVTAQVIWEGLWMNCVVQSTGQMQCKVYDSMLALPQDLQAARAMVIIS

IIVGIFGVLMAIVGGKCTNCMEDETAKAKTCIVSGVIFIIAAFLIMIPVSWSAHAVIRDF

YNPMLIAAQRRELGASLYIGWGAAGLQLLGGGLLCSNCPPKDGQPYIPAKFIPARSISNS

IADYILA

>NEME_00008798_Neogobius

MPAAGRQILAIFLATVGFIGDIIICALPMWKVSAFVGSNIVTAQTFWEGLWMNCVQQSTG

QMQCKVYDSMLALPQDLQAARALVVISILVVLLGMLLAIAGGKCTNCVEDEGAKMKVAIA

AGLLFIIGGILCLIPVSWSANQIISTFYNPMMIEAQRRELGACLFIGWGSAGLILIGGAL

LCCQCKKKRTASILSNTLLLAPLPEGPTSEGLQVCKSAH

>NEME_00013768_1*_Neogobius

MNRTPDAMQVVALTRSSSMQLVHLAPAMARSNPTIPVRMEMMTMARAACRQRPLMHSFLG

GKLCSSMYSTGLEILGIILAVAGWLGVMVACGLPMWRVTAYIGQNIVISQVIWEGLWMNC

SVQSTGQMHCKVHDSMLGLPVDLQAARALVIVSMVLSIVGICFSTAGAKCTNCSQDVNGK

PRLMVAAGVTFIVAGLLMLVAVSWTAHAIVMGFYDPMLEETGKREFGNALYFGWAASCLL

ILGGALLCCSCPPRASHAHSGSVRPR

>NEME_00013768_2*_Neogobius

MPSAGLEILGVT

LAILGWISAIVSCALPMWRVSAFIGVNIVTAQITWEGIWMNCVVQSTGQMQCKVHDSMLA

LSGDLQAARALTIISVVLGIVGVMVSTMGAKCTNCVEEEAAKARVMIAAGVCFIIASLME

LIPVSWSAHTIITEFYNPVIPEAQKREIGAALYLGWAATAFLLIGGIILCTSCPPQTEKR

YGPPSKVIYSPTRSVAPSYPERRDYVSRRQVLFR

>NEME_00013768_3*_Neogobius

MSIGLELIGISLCILGWIIAIVACAL

PMWRVTAFIGSNIVTAQIIWEGLWMTCVVQSTGQMQCKVYDSMLALSQDLQAARALTVIS

ILLAILAVLIAIAGAKCTNCIDDEASKAKVVIISGVFFIVSGVMQLIPVSWSANTIIRDF

YNPLLTDSQRRELGAALYIGWAAATLLILGGDCCAAHARHERPGTTRPGWPTQCLGPQGG

QRWRGKIT

>NEME_00005100_Neogobius

MVSAGLEIMGLSMCVIGSLLVMVACGLPMWKVTAFIEANIVVAQTMWDGLWMSCVVQSTG

QMQCKVHDSILALSHDLQAARALTVTSCVLGVVGMMVVIAGAQCTNCISKENVKVRAVNA

GGILYIISGLFVLVPLCWMANNIISDFYNPQVPASKKREIGAALYIGWAATALLLIGGSL

LCCSCPSSGNSGYSVKYAPTKARTSQNGEYDKRNYV

>NEME_00008797_1*_Neogobius

MQTQIVGIFLSIVGFLGTILICGLPMWKMSAFIGANVVTAQVFYEGIWMNCVIQSTGHAQ

CKAYDSILALPESLQAARAMICASIGVSLVAIGLTVVGGKCSNFYRHDRRMKANLGLSGG

IVFLIAGVLCMIPVSWTAHNTISGFYNPSSFEGRKGELGACIYVGWASGVLLIIGGSLWD

GLWLQCVIQATGQ

>NEME_00008797_2*_Neogobius

MINAQKRELGPRCTSGGARPVCCFWGSCPPNEENDYDVKYSKARSVATSKAYVPNPSNMV

SSGRQMLGLVLAIVGFFGSIIICALPTWRVTAFIGANIVTSQDIWEGLWMNCVTQSTGQM

QCKVYDSLLALPQDLQAARALTIIAIIAGVFGILLGVVGGKCTNFVDDERLKCRVAIASG

IVFIIAGLMVLVPVCWTANTIIQDFYNPIMINAQKRELGASLYIGWGSAGLLFLGGGLLC

SSCPPNEENDYDVKYSKARSVATSKAYVKILC

>NEME_00013771_1*_Neogobius

MVSAGLQILGIALALIGWIGVIIACAIPQWRVTAFVGQNIVTAQTTWEGIWMTCVVQSTG

QMQCKVYDSMLALGQDLQAARALTIISILVVFSAYCWPSLEENAPTVTKAKVCIAAGVIF

IISGILCLVPVCWTANSVVTEFYNPMLMNSQKNELGAALFIGWGASALLLIGGGLLCANC

PPKDDYPAKYSASRATAPRDYVR

>NEME_00013771_2*_Neogobius

MVSQGIQMIGISVSVIGWLLVVVACGLPMWRVTAFIG

ANIVTAQTIWQGMWMNCVVQSTGQMQCKVYDSMLALSQDLQAARAMVIISILTGIVGLLL

AIAGAKCTNCIEEERVKATTCIASGVLFIVSGLLCLIPVSWTANTIVSNFYNPMLLQSQR

YELGASLYIGWAAAGLLLIGGGLMCWNCPPKEQYCNYAPKFVPVKSTSTQREYPHTHCIE

>NEME_00013771_3*_Neogobius

MGKIGKEIAGQVLSFIGLVGVSVTCGVPMWRVTSFIGANIVTGQIVWDGLWMNCVIQSTG

QMQCKLNESLMSLTRDLQAARALIIISLVVGLIGFLISFLGTKCTSCLKKEASMANVVII

SGCLIILAAILDLIPVCWSAAVTIADFQSPLTITTQKREIGASIYIGWASAAFLLIGGII

LTTSCPPQRKFYGYPGYAPPPGAYPYAAPGSQGGYRSVYTPVSQPYSGPYVPNKPY

>NEME_00013771_4*_Neogobius

MLGF

AMAVIGFLGTIIVCAMPMWKVTAFIGANIVTAQVIWEGLWMNCVTQSTGQMQCKIYDSLL

ALPQDLQAARALVVIAIILAALGLLLGIAGGKCTNFVEEPRAKSRVAIAAGVMFICAGVM

MLVPVCWSANTIIRDFYNPIMTNAQRRELGAALYIGWGTAALLLLGGALLCSSCPPKEGP

EYPVKYSGARSTDSRAYNFTCTCCYFRAHLENFVA

>NEME_00013771_5*_Neogobius

MASMGMQLLGAVLALFGWIGVIVVC

AAPMWRVTAFIGNNIVTSQTIWEGIWMSCVVQSTGQMQCKVYDSMLALGTDLQGARALVI

VSIITGLVGLLIAFAGGKCTNFIPEERAKLRATIAAGVVLIISGLLCLIPVSWTASMIIR

DFYNPVMVDAQRRELGASLYVGWGAGALLVLGGALLCANCPKGEDQALSVTAECGP

>NEME_00013771_6*_Neogobius

MGGI

QELGISLSMTGMAGTILICALPMWKVTAFIGTHLVVMQVFWEGLWMTCVSEYTGQMQCKL

YDALLDLSPDLQAARGLICISLVLGCLGFLIFLLGARCTNCLRHPQAKQRMVMSSGLIFC

LDSLTTIVAVAWTANSIINDFYNPRVPDVLKKELGAAIYCGFVSSGLLFIGGVLLCMTCP

PQRAKFPSTRYTLAKVPTQSSYANKNYV

>NEME_00008801_1*_Neogobius

MVSLALEIVGITLSTLGWIMSIVCCTLPMWRVSAFIGANIVTAQVYWEGLWMSCVFQSTG

QMQCKVYDSMLALPQDLQAARALTVVSIMIGILALLIATVGAKCTNCIPDEGVKARVMAS

SGGAFIAASLALLVPVSWSANTIVVEFYSPIVPSGQKMEIGVALYLGWAAAALLMIGGCI

LCCSCPPQEEKALRYSFPPQSRVAYSAAPRSVAQSSYNRKDYRNKDFTK

>NEME_00008801_2*_Neogobius

MSMGLEIVGIA

LGAIGFIIAIVVCALPMWKVTAFIGANIVTSQTIWEGLWMNCVIQSTGQMQCKMYDSLLA

LSSELQASRAMIIIAIILGILGVMISIVGAQCTNCIEDNPTKAKVMIISGIFFLLGGLLV

LIPASWTASVIVRDFYNPLLIDAQRREIGAALYIGWAAAALLIIGGAMLCSSCPKKEQKY

KPQRMAYTAPRSSSGYDRKDYV

>NEME_00008800_Neogobius

MASLGLQILGVGLAVLGWIGNILICMLPMWKVSAFIGNNIVVAQTIWEGLWMTCVVQSTG

QMQCKVYDSLLALPPDLQAARAMVVIAILFALFGILLSVVGGKCTTCIDDKGAKARVAIS

AGIFFILSGALCLVTVSLPANTIIKDFYNPLVPDAQRRELGACLYVGGAQRD

>NEME_00012075_Neogobius

MLTACLEFLGLALSVVGSLLVMIACGLPMWKVTAFIDSNIVVAQTIWDGLWMSCVVQSTG

QMQCKVHDSVLALTQDLQTARALTVISAVLGVVGLTVTVTGAQCTNCLRDETVKARVVNA

GGTIYILSGLFVLVPLCWMSNNIIVDFHNPQIPPSKKREIGASIYIGWAATALLLLGGTL

LCCSFTQSVRGTYPVKYTPTKPVSVNGDFDKNYYV

>NEME_00008796_Neogobius

MASLGTQMIASVLCLFGWAGVIFACLLPMWTVTAFVGSTIVTSQTVWEGIWMTCVVQSTG

EIQCKPYESLLALTADLKAARALMMLAVVIGGVGLLLAFFGGKCTRFLDNEGPEKKRKVA

IAAGVLLVITGLFCLIPTAWAAGAVVKNFYSATFDAQKRELGACLYIGWGAAILLILGGA

LFIASNCHAKSDELDKHSSVRYLVVRSRMEQVRQGPTAGSHQCSPIL

>NEME_00031809_Neogobius

MVNTGMQLISFTCAVTGWIMAIAVTALPQWKVSAFIGASLLTSEIKWEGIWMTCVYQTTG

HMQCKTYDSMLALPPDIQAARALMCLAIFMGWLSCTVSCCGMKCTTCAGDDRRAKAWIAL

SGGILFILTGLCVLIPVSWTANTVVQDFYNPKVPAMHKRELGQAIYLGWASAVILMISGA

VLSSTCPAIEKGMRYRRGYMGRSFANSPATIQDVPQPITTNSVPLKEYV

>NEME_00013533_Neogobius

MGSVGVQIVCVALGVLGLIGVIVCCAIPSWSVSSFVGSNIVTAQSTYKGLWMECVIQSTG

QQQCKNYDSLLVLASDLQAARALTIISAMISCIALLILFCGADFTTCVENEDVKPKISLM

AGVGLLIAGLLVIIPASWSAHTVVRDFHSQVPGQQRMELGPCIFIGWGAGVLLMLAGGLL

CCFSRPKSSGGSAKYYSKSNSGPASKNYV

>NEME_00027886_Neogobius

MASTGLQLVGIVLSVLGGVFGALVCAAPLWRVSAFVGGELVIAQVLWEGLWMNCLSQTTG

QIQCKTYDSTLALPFSAQAARALMLAVAGAKCTHCMGATNPSSKTRVARLAGVLFLAAGV

IYIVPICWTAYGIIKDFYDPNIAAPLKRELGPALYLGWGASVLLIVGGFLLQHGSGPPMS

RVKPVITAPVKDNPKAPEVKQPEKSFV

>NEME_00010016_Neogobius

MANSGLQMLGFSLALLGVIGLIIGTILPQWKMSAYVGDNIITAIAMYEGLWMSCAFQSTG

QIQCKVYDSILQLNSALQASRALMIISIIVSVAGLGVACMGMKCTTCGGDDKAKKSKIAM

VGGIILLVGALSSIIACSWYAHNIIQAFYDPFTPVNTKYEFGSAIFIAWGGSFLDVIGAA

MLAASCSKQKSSPKYPRAPSTNKDYV

>NEME_00004176_Neogobius

MANSGLQLLGYFLALGGWIAIISTTALPQWKQLSYAGDAIITAVGLYEGLWMSCASQSTG

QVQCKIFDSMLSLDIHIQTCRALMVISVLLGLMGIIISVVGMKCTKVGDTNPTTKTRIAV

AGGALFLLAGLCTLVSVSWYATQVSYQFFSPNTPPNARYEFGSALFVGWAAASLTILGGS

FLCCSCSKDEMRGQQYYRQSQPSTTREYV

>NEME_00029461_Neogobius

MANAGIQLLGFCLAFLGFIGSIASTVMVEWKASSYAGDNIITAQALYEGLWKTCASQSTG

QVQCKVYDSLLQLPGIVQGTRGLMLASVLFSFIAMMVGMVGMRCTTCMADSPQQKDKVAL

AGGVLFIIAGFMALVGTSWFGHRVAQDFHNPLTPTNSRYEFGSALYVGWGAASLSIIGGS

ASAVTAAARAAPGRPPATLPPPRDTADPTMSEGVRRLSEKD

>NEME_00026117A_1*_Neogobius

MHGYSGYTKPPQTYVGTYYDKQPPRTDYGEYYQDSIDEQKRLKEKRNHRSAVCCEVVALVI

GFIGLIGVAAVTGLPMWKVTAFIEENIIVMETRWEGLWMNCYRQANIRMQCKVYDSLLFL

PPDLQAARGLMCASVALTTFALITSAVGMKCTTAVDHRARTKHIVLVAGGCLFLLGSVTT

LIPVSWTGHVIIRDFYNPLLIDAQRRELGEALYIGWVTGALLFSAGVILLCRHAPRTQEA

EALKYAQGQPPEYTYQPYTYQPGYQPGYAPSYTYQPAYSAPPPGTVAYTPTRD

>NEME_00026117_2*_Neogobius

MANSALE

IVGLLLTLIGLIGAAASTGMPQWRVTAFIGENIIVFETRYEGLWMNCYRQADIRMQCKVY

DSLLALPPDLQAARGLMCCALALGGLGLLISLVGLQCTSCIRNNDRAKRLVLIIAGSMIL

GACICTIIPVSWTGHVIIRDFYNPLLIDAQRRELGEALYIGWVSSAFLFAGGCFFTCCNI

KSEKEGSERYLYSRNSDYVAYPPPQQLQPQQLVFLPQPQPQFQPRPQPMLSRHPSANYSH

YSLQSQYSQPQPVTRYPS

>NEME_00010201_Neogobius

MASMALELMGFFLGLLGTLGTMVSTVLPYWQISAHIGSNIVTAVANMRGLWMECVYQSTG

AFQCETYNSMLALPADLQASRALMVMSLVLSVLAIAICCVGMQCTVCLDGSGAAKTRVAG

TGGGLFVASGFLSLIPVAWTTHEVVQTFYQNNIPQSLKFEIGECLYVGLASALVTMLGGG

LLAASCCEDQSGRGRRNGPGYPYPVGGAMAGTSTRMTSQAYRSQPVSQPGGVKGQTLQAL

VRSASESSQSTNHNAQRGKKNTGYDITGYV

>NEME_00026112_Neogobius

MPKGLIDIVAVSVGLVGLIGAATTTGLPMWKVTAFIGENIIVMETRWEGLWMNCYRHADV

RMQCKVYDSLLFLPPELQASRGLMCCSLALSGLGLLVALAGMKCISCFQNARAKRILLIV

AGAMQFMASICVFIPVSWTGHAIIRDFYNPLLIDAQRRELGEALYIGWVTGAVLFASAML

FLCRKMPSEKGLFDVYQPPVNLLRYKQQPNQVPMMMYHPSSTSSGLSPPPNSVSNGSQQT

LPFYGSQFNPAQVYNQRLPVSVPMQYHLAHQSSMRSSSHPQTMYTSGNSFYKSQSSMPMT

DTTQTGDPSMSYGSSFYPVPQTPVFVAYNRSIVRHPSASSSAMYI

>NEME_00034332_Neogobius

MSAYIGDNIITAVAMYQGLWMSCAFQSTGQLQCKIYDSILQLDSSLQATRALMIVGIIVS

IAGLGVSCMGMKCTTCGGGDKIRKSRVAMIGGIILLVGGLCCIVACSWFAHNVIRAFYNP

YTPVNTKIVCSIIVCETVRIRRAIFIAWGGSLLDVLGGTMLASSCPRKKQVSKYPTTVST

ARSGPPSSAKEYV

>NEME_00002234_Neogobius

KTRSGFSPFAAGPLGVMDPIVEVVALILGFVGWVMAGVALPNRYWRTSTVDGNVITTATI

YENLWMSCATDSIGVHNCREFPSLLALSGYIQASRALMITAIVLGSFALVAALIGIQCSK

AGGENYVLKGRIAGTAGVLFMLQGLCTMVSVSWYAFNITQEFFNPLFVGTKYEIGEGLYI

GWCSAVLAIVGGACLTCSCKMASDEKHLMPYHSRGTAYSGAARSAAASTYGRNAYV

>NEME_00005400_1*_Neogobius

MGRMATEIVAFLLTISGWILVSSTLPTDYWKVSSVDGTVITTATFWSNLWKTCVTDSTGV

SNCKDFPSMLALDAYIQVCRGLMIAAVCLGFFGAALALVGMKCTKIGGSDTTKARLTVLS

GFHFILCGLCSMTACSIYAHRITTDFFDPLFFAQKFELGAALFIGWSGSVLCIIGGLVFC

LTLSCGFTLRQVEKYSYRGAASFATARPSPCKPVSSLHKETQEHRQFGRNAYT

>NEME_000054002*_Neogobius

MKIRVVQ

IWGFLMTVFGWIFVACTMAMEGWKITSIGGMAGSSIITVAWYWSSLWRSCFTDSTSVSNC

YDFPMLWSVEGHIQIVRGLLMAALSLGMLGFVLSMLGMECTYLGGQDTSKNRKVFAGGWC

HIISGVLSTSGYAVYAQYVSMEYFNPNFDDLKYDLGTPLFLGWVGSAFHMAGGFFYLWSV

CKPLCERGEAEAITVKAQSDVEQNTSTNLNKSTSTLTTQSGVTSGTKLSSISELSSRRPD

VLEISSRTARATKTGRSAKSGRTSRSAGSRGSSVSRTESRSSSRSSRSSRTVSSVSGISR

GESSRTLV

>NEME_00003787_Neogobius

MDTCTCAMELLGMLLFIGAWVCTLVTTILPQWLTMSTALLPIESYDLGLWETCVVQDVGG

MECRAYDSLLGLSRDLKMARIFMCSSLGVGFLGILTTVPGLYVINTCKDQEEVRTKRVMT

VIGGVLGIVAGVLCLIPVCYMAHMAVVHFFDDKVPDVVPRWEFGDALYCGWVGDSCSSRP

EEPQSAVQRRYNFMNAGVAYRKRQEYV

>NEME_00014068_Neogobius

MANMAVQVLGFFLGLLGFVGTVVSTVLPYWRSTAYLGSNIITATSYMKGLWMECVWHSTG

IYQCEVYRSMLALPRDLQKLSDTPLQHTAVAFKQQHCAGGGSAELRSSGSPSSYGPVLCH

VCPGCPRLCSGDEVHLLCPNVRHQVPLALSGGVCFICAGILCLVTVIWTTNDVVMEFYDP

FLPAGFKYEIGLCVYMGYAAASLSLIGGVVILWSSSSERRSRRPEFQTQRSQPVSPPPAF

NNIYPPAPPYKPPEALKDNRAPSLTSLSSNGYRLNNYV

>NEME_00024587_Neogobius

MASTGMQIFGFVLALLGIFGAMVATVLPNWKVSADVGSNIITAISQMQGLWMDCTWYSTG

MFSCTLNCHGPLSGLSWTQVHTLGRGRRSKRHAAIASGGCFVAAGFLCLVPASWFTNEVI

TNFLDSSVPESNKFEPGGAVYVAFVSAGFLFMGGFIFCMSCSGKRHGPQDMVLLPPPDKL

LLQQQQQQLLQQELQHQYCSLSPLDNKTGYSLQDYV

>NEME_00026018_1*_Neogobius

MSCASDSTGLYNC

RDFPSLLALPGVCVLLAFLAWLMVGIALPNRYWKESSVDGNVITTSTIYENFWMSCATDS

TGPQLQRLSFFTGAEWYALQYFIDGCLSGYIQASRALMIASIVFGTFGLVATLAGMQCSK

IGGENYVLKGRIAAIGGVFFLLQGICTLIAVSWYAANITRQFFDEFFSGTKYEIGEGLYI

GWASATLAICGGSCLLFACGGKTSDGKAISVPAGLQKSCLFSNGTISVCAQQLWKKRICL

>NEME_00027152_Neogobius

MASSGLQIVGFLLALSGLSATIAATFMVEWTMESQGNHKIYEGLWMGCSVDEKTICDTHD

SLFKVSEDIQVTRSVMLLSIILTGSGLLMSILGMKCTRFLDEKIETKARTAVTGGIILIV

AGLLSFSITSWYASGIVNSFKNANHLNSYEFGQAIFVSWAGGLLSILGGLFLSYRRCSFC

SQVLTSNHLLPTSHPKSNYV

>NEME_00013772_Neogobius

LRCQLCAVRAVGPELRRRRLVTVVMLRHLELGALGLAFVGWICAILTRCLALWKVSGTVD

NTTASLPAYWDGVWLQWDHWDLAHDGSLHCSFYQTLMSLSGSFRTWRSLISAGIGAGAFA

VVISAVGAVWFPHRGQIKVFSGAVFVLSGILLLVPTAWTCHHTSQPLEGAIHLTRDWGPA

LYLGWIAFTLMTVGGVFLTTRCPTAQSQGEHTEVSTSPHQEEDSSHP

***Zebrafish (Danio rerio)***

>sp|Q9YH91|CLDY_DANRE_Danio

MASVGLQLLATVLAIIGWLGEIVICALPMWKVTAFIGNNIVTAQIFWEGLWMNCVQQSTG

QMQCKVYDSMLALPQDLQAARALVVISIIVTFMGVFLTIAGGKCTNCIEDQDAKAKVVVA

AGVFFLVGGILCLIPVCWSANSVIKDFYNPTLSDAQKRELGASLFIGWCASGLLLLGGAL

LCCQCPKNEGRAYSVKYSAPRSAPGAYV

>sp|Q9YH90|CLDZ_DANRE_Danio

MSTGLQLLGTTLGTLGWLGIIISCAIPLWRVTAFIGNNIVTAQTMWEGLWMSCVVQSTGQ

MQCKVYDSMLALAQDLQASRAILVISAIVGLIAMFASFAGGKCTNCLADNSAKALVATTG

GVAFIIAGILGLVPPSWTANTIIRDFYNPLVAEAQKREFGAAIFICWGAAVLLVIGGGLL

CSSYPKGRTSSRGRYTPASQNGRERSEYV

>tr|Q7T2E7|Q7T2E7_DANRE_Danio

MDPIIEAVALFLGFVSWTMVGITIPNRYWKVSSLDGTVITTSTLYENLWMSCATDSTGVH

QCREFPSLLALSGYIQASRALMIAAVVSGTFGVVATLIGMQCSKAGGENYVLKGRIAGTG

GVFFLLQGLCTMISVSWYAANITQEFFNPLYPGQKYEIGEALYIGWASAVLAICGGVCLM

FSCKLGTEKKTAYSYHPTRETVYSASASRRETQSTYGKNAYV

>tr|Q6NUZ6|Q6NUZ6_DANRE_Danio

MASAALELLGLILCVCGTLLEMVACGLPTWKVTAFIEANIVVAQTIWDGLWMSCAVQSTG

QMQCKVHDSMLALSHDLQAARALTVISSVLSVLALMVVIAGAQCTNCIKEDNVKARVVNV

GGVIYILSAIFVLVPLCWMANNIISDFYNPQVLPAQKREIGAALYIGWAASALLLLGGSI

LCCSCPASGSSGYSVKYAPTKRATSNGEYDKRNYV

>sp|Q9YH92|CLD7B_DANRE_Danio

MAHKGLQLLGFTLSLLGLIGLIIGTIMPQWKMSAYVGDNIITAIAMYQGLWMSCAYQSTG

QQQCKVYDSVLQLDSALQATRALMVVAILLTVAGLGVASMGMKCTNCGGDDKVKKSRIAM

TGGIILSVGALCSIVACGWFTSQIIRDFYNPFTPVNTKYEFGAAIFIAWAGAFLDIMGGG

MLASSCSKGQSSPNYPKSSRPVKSSRPPSSSKEYV

>tr|Q6DHE9|Q6DHE9_DANRE_Danio

MVLQVLGLFLGIVGWCLESSSINSSVWKSSSHGEAVVTASSQFEGLWFSCASNSLGAIHC

QRFKTTLGLPGYIQACRALMIIALILGLLSVVLASMGLKCTKLGSTSEEAKGKISLTAGI

MFILSGLCVIVAVSWYAARVVQEFNDPFYGGTKYELGAGLYLGWAAAALCILGGGTLCTS

FKGSSPAQTRGPGYNYSAAQPQKIYRSAPSDNSITKAYV

>tr|A0A0R4ID02|A0A0R4ID02_DANRE_Danio

MAVMVLQVLGLFLGIVGWCLESSSINSSVWKSSSHGEAVVTASSQFEGLWFSCASNSLGA

IHCQRFKTTLGLPGYIQACRALMIIALILGLLSVVLASMGLKCTKLGSTSEEAKGKISLT

AGIMFILSGLCVIVAVSWYAARVVQEFNDPFYGGTKYELGAGLYLGWAAAALCILGGGTL

CTSFKGSSPAQTRGPGYNYSAAQPQKIYRSAPSDNSITKAYV

>sp|Q6DHP1|CLD7A_DANRE_Danio

MANSGVQLLGFGLSLIGIIGLIVGTILPQWKMSAYVGDSIITAVATYQGLWMSCAFQSTG

QLQCKIYDSILQLDSDLQATRALMIVGIIVSIAGLGVASIGMKCTTCGADDKVRKTRTAM

TGGIILLVGALCAVVACSWFAHNVIRAFYNPFTPVNTKFEFGAAIFIAWGGSFLDVLGGA

MLAASCPRSKQVSKYPKSNSTRSANGSNKEYV

>tr|Q8QHA3|Q8QHA3_DANRE_Danio

MGSMATEIVAFLLTISGWILVSSTLPTDYWKVSSVDGTVITTATFWSNLWKTCVTDSTGV

SNCKDFPSMLALDAYIQVCRGLMISAVCLGFFGASLALVGMKCTRIGGSETTKARITCLC

GLHFILSGICSITACSLYAHRITSEFFDPLFVKQKFELGAALFIGWAGSVLSILGGLIFC

FSMTEGFSIREYSYNGATSVISTRTKITKTNKSTKSQKDPSDQMFSGRRFGRNAYV

>tr|Q90XQ8|Q90XQ8_DANRE_Danio

MAHAGLQMLGYCLGFLGLLGLIASTAMAEWKMSSYAGDNIITAQAQYEGLWQSCVSQSTG

QLQCKKYDSLLKLPGEIQGARGLMLTGIFLCGLSTLVSFVGMKCTTCLSEAPQVKSKVAL

AGGVLFITGGLFALIATSWYGEKIRQKFFDPFTPTNARYEFGKALYVGWGSSALSIIGGS

LLCCICGSEASEKPSYPPARAAGRPGTDRV

>tr|Q1RLW4|Q1RLW4_DANRE_Danio

MQNILQIFALLLGFLGCGSAVLSLYNHYWKVSTDDDSVIITSNIFENLWMSCADDSSGTF

SCRDFQSLLALPGYIQACRALIITSIVLGTFGLVATLVGMQCFKIGGGNYIMKSRIAGAG

GVFFILQGLCTMTAVSWYAFNITQEFFDPLYPGIKYEIGEGLYIGWSSATLALCSGSCLL

CSCGLCTPMGKKSFRQSHLQDFRSSHQPQSINIVPTVSTTSHYLSRQYV

>tr|A8WGR6|A8WGR6_DANRE_Danio

MRNMAREILAFILCTSGWVLISSTLPTDYWKVSSIDGTVITTATFWSNLWKTCVTDSTGV

SNCKDFPSMLALDGYIQVCRGLMITAVCLGFFGSIFALIGMKCTKIGGPDNFKARVVCFA

GFNFACTGLCSLTAYSLYAHRITSEFFDPMFVAQKYELGAALFIGWAGSTLCLLGGALFT

LSMADGSSKKSSPNRRVASLMSGAMPQSQTSHTRRTLPEYSSTPQHFDKNAYV

>sp|F1QIK8|EMP2_DANRE_Danio

MLVILAFIILFHITSAILLFIATINNAWRIKGDFSMDLWYNCNTTACYDIPKSATYDAAY

LQAVQATMILATILCCVGFFVFILQLFRLKQGERFVFTAIIQLLSAFCVMTGASIYTAEG

LTFNGQEFKNAEYGYSFVVAWVAFPMTLLSGLMYLVLRKRK

>tr|F1QH81|F1QH81_DANRE_Danio

MRNMAREILAFILCTSGWVLISSTLPTDYWKVSSIDGTVITTATFWSNLWKTCVTDSTGV

SNCKDFPSMLALDGYIQVCRGLMITAVCLGFFGSIFALIGMKCTKIGGPDKFKARVVCFA

GFNFACTGLCSLTAYSLYAHRITSEFFDPMFVAQKYELGAALFIGWAGSTLCLLGGALFT

LSMADGSSKKSSPNRRVASLMSGAMPQSQTSHTRRTLPEYSSTPQHFDKNAYV

>tr|F1R6Z2|F1R6Z2_DANRE_Danio

MAHAGLQMLGYCLGFLGLLGLIASTAMAEWKMSSYAGDNIITAQAQYEGLWQSCVSQSTG

QLQCKKYDSLLKLPGEIQGARGLMLTGIFLCGLSTLVSFVGMKCTTCLSEAPQVKSKVAL

AGGVLFITGDKLMLITRLWFESRLGQLVFLPNTCADAMYEFGKALYVGWGSSALSIIGGS

LLCCICGSEASEKPSYPPARAAGRPGTDRV

>tr|E7F357|E7F357_DANRE_Danio

MASTGTQIFAFVLALLGILGATVATLLPNWKVSADVGSNIITAISQMQGLWMDCTWYSTG

MFSCTLKYSVLALPAYLQTARTTMVLSCVLAALGLCLASLGLKCTRWGGGRRAKRHAAMA

AGTCFLASGFLCLVPASWYTNEVIVSFLDANVPESNKFEPGGAVYVAFISSGFLFVGGSI

FCLSCSGKRHGPQDLILLPPDKLLYCSLSPLDNKTGYSLQDYV

>tr|E7FAL1|E7FAL1_DANRE_Danio

MPSSTMQMFAFILALLGVVSATVATLLPNWKVSADVGSNIMTAISQMQGLWMDCTWYSTG

VFSCSLKYSVLALPAYLQTARTTMVLSCMMATMGLCLAALGLKCTRWGGSYRSKGHTAIS

AGACFVIAGILCLVPASWFTNEVITTFMDSKVPKSSKYEPGAAVYVAFVSAGFLLAAGVI

FCLSCPGRRGGALDSGSSHPEKPKQLKLRQKQPTGNQKKQQHREQQSQTETPEQEESRPE

TLHQEKPQPQQEKKSVPDRLNLEKEKQY

>tr|Q6DHL0|Q6DHL0_DANRE_Danio

MANTCLQFTGFVMSFLGWIGLIVATATNEWVFTCKYQMNTCRKMDELEAKGLWADCVIST

ALYHCITLTQILELPAYIQTSRALMVTASILGLPAVVMVLMSMSCINLGGEPESAKNKRS

VLGGILILLTAFCGIVSTVWFPIGAHHEKGLMSFGFSLYSGWVGTALCLLGGCIISCCSV

DSPATYTENNRFYYSKQGPAHTGPTSTNHAKSAHV

>tr|Q90XR9|Q90XR9_DANRE_Danio

MVSMCREILGMCLAIIGFLGAIIICALPMWKVTAFIGANIVTAQTIWEGLWMNCVMQSTG

QMQCKIYDSLLALPQDLQAARALVVIAIIVSFFALILGIAGGKCTNFVEREDAKAKVSIA

SGVIFIIAGVLVLVPVCWSANTIIRDFYNPLLTDAQRREMGASLYIGWGVAALLIIGGGI

LCSSCPPKDEKYNMKYSQPRSTATSRAYV

>tr|A0A0R4IYC1|A0A0R4IYC1_DANRE_Danio

MQNILQIFALLLGFLGCGSAVLSLYNHYWKVSTDDDSVIITSNIFENLWMSCADDSSGTF

SCRDFQSLLALPGYIQACRALIITSIVLGTFGLVATLVGMQCFKIGGGNYIMKSRIAGAG

GVFFILQGLCTMTAVSWYAFNITQEFFDPLYPGIKYEIGEGLYIGWSSATLALCSGSCLL

CSCGLCTPMGKKSFRQ

>tr|Q66HX3|Q66HX3_DANRE_Danio

MLGTLVATLLPYWATSAHVGPNIVTAVVSMKGLWMECVYQSTGAFQCETYNTLLGLTVDL

QAARAMMVMSSIFSVMACAVSTVGMQCTVCMDGSSVKTKVAGVGGSMFLLAGLLSLIPVA

WKTHEVVQTFYMPNMPASLKFEIGDCLYVGLASSLLSMLGGGLLTASCCDDIDGNRGSRR

HYPYPERNPLRGPSHSMTYQPAALHSTTNPNITNKTQTLNSHTSSGTHSIQAQDSRKMTR

QNTAAGYDVTGYV

>tr|Q503G1|Q503G1_DANRE_Danio

MRNMAREILAFILCTSGWVLISSTLPTDYWKVSSIDGTVITTATFWSNLWKTCVTDSTGV

SNCKDFPSMLALDGYIQVCRGLMITAVCLGFFGSIFALIGMKCTKIGGPDKFKARVVCFA

GFDFACTGLCSLTAYSLYAHRITSEFFDPMFVAQKYELGAALFIGWAGSTLCLLGGALFT

LSMADGSSKKSSPNRRVASLMSGAMPQSQTSHTRRTLPEYSSTPQHFDKNAYV

>tr|Q7T021|Q7T021_DANRE_Danio

MATEIVAFLLTISGWILVSSTLPTDYWKVSSVDGTVITTATFWSNLWKTCVTDSTGVSNC

KDFPSMLALDAYIQVCRGLMISAVCLGFFGASLALVGMKCTRIGGSETTKARITCLCGLH

FILSGICSITACSLYAHRITSEFFDPLFVKQKFELGAALFIGWAGSVLSILGGLIFWNTP

TMVPHLSYQQEPRSQRQTKAPSLKKTPQTRCSLADGLAGTHMSDFNFLCKAGNDNK

>tr|F1QVR6|F1QVR6_DANRE_Danio

MALQVLGITLSMIGFAGTIIICALPMWKVTAFIGTNIVVAQVFWEGLWMTCVYERIGQMQ

CKLYDALLDLDPFLQASRGLIVTTMALASLAFLIFLIGADCTNCLSNPRAKGRIVVVSGI

TFMLSGLTTVVPVSWTADSIIRDFHNPVVHEALKREMGAALYVGWLTAGFLFIGGAILCT

SCPPERDNYLPRYTLTKSGTHSGYAVKNYV

>tr|A0A0G2KYS3|A0A0G2KYS3_DANRE_Danio

MHYRTLLMHTEMVCFVVCVCGWLLVCSTVPTECWAYSEVHTSVLTSAHYFSNLWKDCLSD

STGVTDCKVFPTFLALRPYIHVCRSFLFTSILLGLFGSILALVGMKCTKLGGSETVNAKV

TFAGGINYLTSGLCGMFTYSWYGHRVVSEFMDPGFVGKRYELGPALFVGWGGSALLMLGG

LVFSFTAGNQGFQS

>tr|Q90XR8|Q90XR8_DANRE_Danio

MASTGLQMLGIALAIFGWIGVIVLCALPMWKVTAFIGANIVTSQTSWEGIWMSCVVQSTG

QMQCKVYDSMLALSSDIQAARALTVISIVIGVMGIMLSMAGGKCTNCIEEESSKAKVGIT

AGVIFIISGVLCLVPVCWTANAIIQDFYNPLVVQAQKREIGASLYIGWGASALLIIGGSL

LCCHCPEKSDSGKYTAKYNATPRSEASAPSGKNFV

>tr|A8E4Z2|A8E4Z2_DANRE_Danio

MVQGPCDVIALCLSLIGLIGVATVTALPMWRVTAFIGENIIVMETRWEGLWMNCYRQANI

RMQCKVYDSLLYLPPDLQAARGLTCCSLALCGFGIIVSLFGLRCMSCLSDQPRNKSIILM

ITGIIEILASFCIVIPVSWTAHTIIRDFYNPLLLDAQRRELGEALYIGWVTSAFLLASGV

IFLCRRVDSNEQPFYARPPRNKQVPNHFQTFNSIRRQPLLQNNSFSTPQHSSAAFSFAVT

PSSGQYSQHNVQPVVNGSLIYSPNMIPPQRVNYNGQNVMQSQILASNSQHPVSDPRDSFN

TGSAFHPVQQNPLYVGYNYSRVQSGSSNSSTGLRI

>tr|Q90XR4|Q90XR4_DANRE_Danio

MALQVLGITLSMIGFAGTIIICALPMWKVTAFIGTNIVVAQVFWEGLWMTCVYERIGQMQ

CKLYDALLDLDPFLQASRGLIVTTMALASLAFLIFLIGADCTNCLSNPRAKGRIVVVSGI

TFMLSGLTTVVPVSWTADSIIRDFHNPVVHEALKREMGAALYVGWLTAGFLFVGGAILCT

SCPPERDNYLPRYTLTKSGTHSGYAVKNYV

>tr|A8WGD4|A8WGD4_DANRE_Danio

NRMSAPALQTTGFVSGVIGTAGVFAATLMDVWCFRNQQQDQPQRVSSIYTYKGLWKDCEM

SGTVFPECQPLYSHPNYSGILQAARALMIIAILVAVIAVFIGFFCLKCLKMKNMQLSTRA

KLILSSAIIFFIAGICGIAAASVYADQLVPSFMMPQFNQKQGEKGGVQCANNVAAMDPSA

SRYTFGPALYIAWVGAALLILGGILMSIAYKRMQCKTETREGYIYNAAQCRAAEEESQCQ

TQGQSLSQCQYQCHSQVQRI

>tr|A4QNU3|A4QNU3_DANRE_Danio

MSNSCRLLCGFLMSFVGWIGIIIATSTNDWVLSCTYGSHSCRNMDDLETKGLWTECVIST

ALYHCIPLNQVRRIPAYIQACRVLMVSASLLGLPALALLLLAMPCVKVSQETEGTKHRRA

VQGGLIILVISLCGMVSTVWFPIGKLDGLMSFGFSLYAGWVGSALCFFAGSVMVCCSKDH

SPSENPESRHYYSNPDGVTSTGPSENPETRYYYSNHDGVTSSGTSENQETRFYYSNHDGV

TSYGPAKNSHAKSEHV

>tr|F1QKA9|F1QKA9_DANRE_Danio

MAILALELMGFFFGLIGMLGTLVATLLPYWATSAHVGPNIVTAVVSMKGLWMECVYQSTG

AFQCETYNTLLGLTVDLQAARAMMVMSSIFSVMACAVSTVGMQCTVCMDGSSVKTKVAGV

GGSMFLLAGLLSLIPVAWKTHEVVQTFYMPNMPASLKFEIGDCLYVGLASSLLSMLGGGL

LSASCCDDIDGNRGSRRHYPYPERNALRGPSHSMTYQPAALHSTTNPNITNKTQTLNSHT

SSGTHSIQAQDSRKMTRQNTAAGYDVTGYV

>tr|A0A0G2L7R9|A0A0G2L7R9_DANRE_Danio

MSAPALQTTGFVSGVIGTAGVFAATLMDVWCFRNQQQDQPQRVSSIYTYKGLWKDCEMSG

TVFPECQPLYSHPNYSGILQAARALMIIAILVAVIAVFIGFFCLKCLKMKNMQLSTRAKL

ILSSAIIFFIAGICGIAAASVYADQLVPSFMMPQFNQKQGEKGGVQCANNVAAMDPSASR

YTFGPALYIAWVGAALLILGGILMSIAYKRMQCKTETREGYIYNAAQCRAAEEESQCQTQ

GQSLSQCQYQCHSQVQRI

>tr|Q7ZTS2|Q7ZTS2_DANRE_Danio

MGSAGVQIVCVALGILGLIATVVTIAIPQWKTSAFIGQNIITAQVSEEGLWMQCVVQSTG

QQQCKSYDSLLILSSDLQAARAMTILCGGADFTTCIENEDVKPKVTLVSAIGLILAGLLV

IIPVSWAANNVVRDFNNPMVPEAQKRELGPCIYIGWASGVLLILAGGLLCCFSRPRSSGS

SGAAKYYSNSASAPSKNYV

>tr|E7EXG8|E7EXG8_DANRE_Danio

MASQGIQILGIMLSMIGWLGTIIACGMPMWRVTAFVGANIVTAQIIWEGLWMSCVVQSTG

QMQCKVYDSMLALSQDMQASRAMVIISIMVGIFGFLMAVVGGKCTNCLEDEAAKAKACIF

SGVVLIIAGLLILIPICWSAHTLIRDFYNPLLTASQRRELGACLYIGWGSGGLLLLGGGL

LCWNCPSNNTRPYIAAKYTHGRSMTPTINYV

>tr|Q567D5|Q567D5_DANRE_Danio

MANSGFQLLGYFLALGGWIGIISTTVLPQWKQSSYAGDAIIMAVGLYEGLWMSCASQSTG

QVQCKIFDSLLSLDVNIQTCRALMVISVLMGFMGIIVSVVGMKCTKVGDNNPSTKSKIAI

SGGSLFLLSGVCTLVAVSWYAAQVSAHFFDPNTPTNAKYDFGTALFVGWAASVLMMLGGA

FLCCSCINEDRRGQQFYRQSQPSTTREYV

>tr|B8JLG4|B8JLG4_DANRE_Danio

MSTALEVTGYFMCLIGWVLTGLAVANDYWKISSIQGNVIVSNRLYENLWHACGEDSTGKA

NCQDFQSMLALPVHIQACRALVIIALLLGLVGLVLSTMGLNCIKIGSKTDESKGKNMFIG

GIIYLIGGLCTMVGVSWYAARVVQEFNDPFYGGVRFELGSGLYIGWAGAALCMLGGGFQC

SAYQRFSKSKEKGAYYPAGKPQTIYTTAQSNAETSKAYV

>tr|Q90XR7|Q90XR7_DANRE_Danio

MSTALEVTGYFMCLIGWVLTGLAVANDYWKISSIQGNVIVSNRLYENLWHACGEDSTGKA

NCQDFQSMLALPVHIQACRALVIIALLLGLVGLVLSTMGLNCIKIGSKTEESKGKNMFIG

GIIYLIGGLCTMVGVSWYAARVVQEFNDPFYGGVRFELGSGLYIGWAGAALCMLGGGFQC

SAYQRFSKSKEKGAYYPAGKPQTIYTTAQSNAETSKAYV

>tr|F1QP10|F1QP10_DANRE_Danio

MVQGPCDVIALCLSLIGLIGVATVTALPMWRVTAFIGENIIVMETRWEGLWMNCYRQANI

RMQCKVYDSLLYLPPDLQAARGLTCCSLALCGFGIIVSLFGLRCMSCLSDQPRNKSIILM

ITGIIEILASFCIVIPVSWTAHTIIRDFYNPLLLDAQRRELGEALYIGWVTSAFLLASGV

IFLCRRVDSNEQPFYARPPRNKQVLNHFQAFNSLRRQPLLQNNSFSTPQHSSAAFSFAVT

PSSGQYSQHNVQPVVNGSLVYSPNMIPPQRVNYHGQNVMQSQILASNAQYPVSDPRDSFN

TGSAFHPVQQNPLYVGYNYSRVQSGSSNSSTGLRI

>tr|F1REC4|F1REC4_DANRE_Danio

PGSCALELLGVFFSLCACLCSLLSTMMTRWLTLSTELLPTESFELGLWMTCVVQELGVTE

CRPYDSLLGLPPDIRLARIMMCTSVAAGLSALVFAIPGINLVNSCKNRADSIEAKRTLKI

FGGILSLSSGVLGIVPVSYVAHLTVLRFFDESVPSVVPRWEFGDALFLGWTAGCLQVVAG

LLLITSCFFLQDKTRGLGESIHMDRVNGTRSPRNRTENV

>tr|A0A0R4IEG7|A0A0R4IEG7_DANRE_Danio

MRKRLIQVLGFLISTFGWLFVSCTLAMDYWRILYVGGKGGNWMVKASWYWSNLWKDCVTD

MSSISDCRDYDALWAVTPYVQAVRGLLMIAMGLGFIAAILCFIGMECTYIGGSEKNKRRV

LLAGAALHFAGGLSAAAAYCLYTNRVARAAFAPAVDTTIIRYGIGAPVFFGLVGSFLIIL

GSALYAVTHISTRKKTVSVGRTLYTRPSYGRRTASKTAYTSAYSAPSRQSSSRLSRMSQA

TDEKLPARDTFV

>tr|B8JII0|B8JII0_DANRE_Danio

MSTGLQLLGTTLGTLGWLGIIISCAIPLWRVTAFIGNNIVTAQTMWEGLWMSCVVQSTGQ

MQCKVYDSMLALAQDLQASRAILVISAIVGLIAMFASFAGGKCTNCLADNSAKALVATTG

GVAFIIAGILGLVPPSWTANTIIRDFYNPLVAEAQKREF

>tr|Q561T6|Q561T6_DANRE_Danio

MASLGMQILGVALSFVGTLGSIITCVLPMWRVTAFIGNNIVTAQMIWEGLWIVCVVQSTG

QMQCKLYESMLALEPDLQAARALLVVSILLSILAVLLAVVGGKCTTCIENKSAKARVVVS

AGVIFLLSGLLCLTPVCWTAHNTIRDFYNPLLQDAQRRELGAALYTGWGSAGLMLIGGAI

LCCQCPAKDQRAFVAKYPAPKSNGSDKEFV

>tr|A0A0G2KZC8|A0A0G2KZC8_DANRE_Danio

MRTRQFVAAFLALVGLCGTILICALPMWKVSAFVGANIVTAQVYWQGLWMNCVLQSTGHM

QCMAYYSVLALTQDLQAARGLICASIGVSAIAFGMMVVGANCTRFYREDQLKKTNIGISA

GALYIVGGVMCLVAVCWQTSIIVMNFYNPQVIAGTQGELGACIYIGWVAGFLLIIGGGLL

LSTYSNRC

>tr|Q567K6|Q567K6_DANRE_Danio

MSTALEVTGYFMCLIGWVLTGLAVANDYWKISSIQGNVIVSNRLYENLWHACGEDSTGKA

NCQDFQSMLALPVHIQACRALVIIALLLGLVGLVLSTMGLNCIKIGSKTDESKGKNMFIG

GIIYLIGGLCTMVGVSWYAARVVQEFNDPFYGGVRFELGSGLYIGWAGAALCMLGGGFQC

SAYQRFSKSKEKGAYHPAGKPQTIYTTAQSNAETSKAYV

>tr|Q6DBT9|Q6DBT9_DANRE_Danio

MATTGMQLLGLVLSIIGLVGGFLVCTLPMWRVTAFIGNNIVTAQITWEGLWMNCIWQSTG

EIQCKGYDSLLALPSDMKAARGLTVLAILICSLSLTLGILGIKCTECVGLPSLKARLARV

SGVLFVIAGFLILVPVCWTAHSIIRDFYDPYVAAPHKRELGPALYLGWGASALLLIGGSL

LYAGSNPPGIPSSPTFSSDESSPRRAGGSSQVKGYV

>tr|Q7T2P4|Q7T2P4_DANRE_Danio

MSMGLEIGGIALGIIGWIISIVACALPMWRVSAFVGANIVTAQVMWDGLWMNCVVQSTGQ

MQCKVYDSMLALGQDLQASRAMTVIAIILAVLGVMISVMGAKCTNCIEDEGAKAKVMIVS

GIMFIIAGILDLIPSAWVANQIIRDFYNPLLPGAQQRELGASIYIGFAAAALLIIGGAML

CCTCPPKEKKYKPARMGYSAPRSASAGYDKKDYV

>tr|Q5XJD5|Q5XJD5_DANRE_Danio

MISACLEIVGLCLTVTGTLLVMVACGLPMWKVSAFIEGNIVVAQNIWDGLWMSCVVQSTG

QMQCKMHDSVLALTTDLQTARALTVISAVLGVLALMITIAGAQCTNCINTESIKSKVVNA

GGVMYIVAGLFVLIPLCWMANNIISDFYDPQVPYAKKREIGAALYIGWAASAMLLVGGAI

LCFSRPHDEKSTYPLKYIPPPTKGTSINGDYDKRNYV

>tr|Q90XR6|Q90XR6_DANRE_Danio

MGSAGVQIVCVALGILGLIATVVTIAIPQWKTSAFIGQNIITAQVSEEGLWMQCVVQSTG

QQQCKSYDSLLILSSDLQAARAMTIISCMLSVLSLLILCGGADFTTCIENEDVKPKVTLV

SAIGLILAGLLVIIPVSWAANNVVRDFNNPMVPEAQKRELGPCIYIGWASGVLLILAGGL

LCCFSRPRSSGSSGAAKYYSNSASAPSKNYV

>tr|B0V349|B0V349_DANRE_Danio

MVLFTTKFVQRASLFVSFGGLVTTFVTTFLPLWKTMNSDLNEMENWYEGLWHMCIYTEEV

GIHCKAFDSFLALPPDTFAGRVLMCISIATGILGVAAAFFGLRGVEIGASRERMKRNLLI

LGGVFVVVSGVTTLAAVSFMAYVMVVKFWDDDRPEVMPGWEYGEAMFSAWFAGLLLVVGG

SFLFVAVCMGDHEVKLQTEMIARCQEQRPRSLHYRKTEII

***Three spine stickleback (Gasterosteus aculeatus)***

>tr|G3NKI0|G3NKI0_GASAC_Gasterosteus

MAATLCQVMGFILSLLGVAGIIAATGMDQWATEDLFDNPVTAVYSYSGLWNSCVRQSSGF

TECRPYFTILGLPALLQAVRALMIVGIVLGAIGGLIAIFSLKCLKMGNMEDNIKATMTLT

AGIMFLLGGVCGIAGVSAFANLIVQSFRFTTYADGGFSMMGGGGIGGLSGSLTPRYTFGP

ALFVGWIGGAVLVIGGVMMCLACRGMSSDAKQRYDGMAYKASSHHTIYRTDPKPRPAFND

SYRAHSADGRMSNQKFDYV

>tr|G3NJE4|G3NJE4_GASAC_Gasterosteus

MSGLQILAFVSGLAGLGATIGATVSNEWRATSRASSVITATWVLQGLWNNCAGNAIGALH

CRPHHTILQLEGYIQACRGLMIAAICLGFFGSISALVGMKCTKIGGNDKNKARIACFAGG

NFILSGLCSLSACSLYAHRITTEFFDPMYIAQKYELGAALFIGWSGSVLCILGGSMLCCS

ITASFPKSQSQVNYIYNGAVSHSHISSYPRGQAKSANQRPPPDYSSSSRTQHFDKNAYV

>tr|G3NJF1|G3NJF1_GASAC_Gasterosteus

TSSMFQEIFAFILSTLGWVLVSSTLPTDYWKVSSLDGTVITTATYWSNLWKTCVTDSTGV

SNCKDFASMLALDGYIQACRGLMIAAICLGFFGSISALVGMKCTKIGGNDKNKARIACFA

GGNFILSGLCSLSACSLYAHRITTEFFDPMYIAQKYELGAALFIGWSGSVLCILGGSMLC

CSITASFP

>tr|G3NJE5|G3NJE5_GASAC_Gasterosteus

MSGLQILAFVSGLAGLGATIGATVSNEWRATSRASSVITATWVLQGLWNNCAGNAIGALH

CRPHHTILQLEGYIQACRGLMIAAICLGFFGSISALVGMKCTKIGGNDKNKARIACFAGG

NFILSGLCSLSACSLYAHRITTEFFDPMYIAQNQSQVNYIYNGAVSHSHISSYPRGQAKS

ANQRPPPDYSSSSRTQHFDKNAYV

>tr|G3NKG1|G3NKG1_GASAC_Gasterosteus

AGLQVAGLLLGLVSWCLQSSCTSSQVWKVRSQQGTVGSGQWQYEGLWMSCAATSLGSVQC

SRFKTLLGLPPHLQACRALMILSLLVGLASIVVSILGLKCIKIGRTSENVKDQIALSGGV

LFILSGVFTLTATSWYASRVIQDFYNPLYGGVRFELGTGLYLGWSASCLAIVGGSLLCCS

CRRSSTTTSSRQFSYNFSSTREGQKIYRAAPTSNDGSSKAYV

>tr|G3P8Q8|G3P8Q8_GASAC_Gasterosteus

MATTGMQLLGLIMSLVGWVGGAVVCAMPLWRVTAFIGNNIVTAQIIWEGLWMNCIVQSTG

QIQCKVYDSLLALPSDMQAARGLTVFSILMCGLALALGVLGVKCTKCIGVNSVKARVARI

SGGLFAIAGFLYLVPVCWTAHSIIRDFYDPHVAAPHKRELGPALYIGWGASALLLIGGSL

LYAGSSPPGMPGSPTFSSGESSPRRAPASQVKGYV

>tr|G3QAU6|G3QAU6_GASAC_Gasterosteus

MNAIVEALAFLLGFLAWLMLGIALPNRYWKVSSVDGNVITTSTIYENLWMSCATDSTGVH

NCRDFPSLLALNGYIQASRALMIASIVFGTFGLVATLVGMKCSKIGGENYFLKGRIAAIG

GVFFLLQGISTMIAISWYAANITQQFFDQFYPGTKYEIGEGLYIGWSSAILAICGGACLI

CACKLNTPKEKIPYPYQPSSRGLVPQTVAMSQSAATNYGRNAYV

>tr|G3NJF9|G3NJF9_GASAC_Gasterosteus

VEIGCFVVCVTGWILVCSTMPTEIWTWSEVDSIVLTTSNYFSNLWKDCVSDSTGVSDCKG

IPSMLALNWDIHMCRALIIIAIILAFFGSVLVLVGMKCTKVGGSEIVNARVTFAGGMNYL

IGGLCSMIAFSYYGNKIRAEFQNPNYRAQKFEIGVGVFIGWAGSSLLVVGGLIYSTVAGR

EGCNSRYSVYLYLYFEHW

>tr|G3QB40|G3QB40_GASAC_Gasterosteus

MVSFGLELVGVLLSVLGWVLSVTSCALPMWRVSAFIGSNIVTAQVYWEGLWMNCVFQSTG

QMQCKVYDSMLALPQDLQAARALTIVSIIVGAVALLISMVGAKCTNCIEEEGVKARVMAA

SGGAFITAALAQLVPVSWSAHTIVFEFYSPLVPSGQKMEIGAALYLGWAAAALLLVGGSI

LCCSCPPREDKPARYSVHPQSRMAYSAAPRSNAPSSYNKRDYV

>tr|G3NUJ0|G3NUJ0_GASAC_Gasterosteus

MASTAVQLLGFFLGLLGFIGTVVATLLPYWRSTAYVGSNIITTTAYMKGLWMECVWHSTG

IYQCEVYRSLLALPRDLQAARALMVLSCVTSVLASLVSVMGMKCTRCARDPLVKSSLVLS

GGIGFLCAGVLCLVTVSWTTNDVILDFYDPFLPSGMKYEIGLAVYLGYASACVSLGGGMA

LCWRRRHVQRNQPLSPPPAFSHIYPPAPPYKPPEALKDNRAPSLCSLSSNGYRLNNYV

>tr|G3Q8Z2|G3Q8Z2_GASAC_Gasterosteus

MANSGIQLLGFFMSLIGIVGLIIGTILPQWKMSAYIGDNIITAVAMYQGLWMSCAFQSTG

QVQCKIYDSILQLDSSLQATRALMIVGIIVSVAGLGVACMGMKCTTCGGNEKLRKSRIAM

TGGIILLVGGLCAIVACSWFAHNVIRAFYNPYTPVNTKFEFGAAIFIAWGGSLLDVLGGA

MLAASCPRKKQVSKYPSMAPPRSGPPSSTKEYV

>tr|G3Q6F4|G3Q6F4_GASAC_Gasterosteus

VMDQILEVVALLLGFIGWVMVGVTLPNRYWRTSTVDGNVITTSTIYENLWMSCATDSTGV

HNCREFPSLLALSGYIQASRALMITSIVLGTCGLLAALIGVKCSKAGGENYVLKGRIAGT

AGVLFILQEGMCTMVAVSWYAFNITQEFFDPFHPGIRYEIGEALYIGWCSAVLAIAGGAC

LTCSCKMGTKQARLLVSFRSPLPLHSRGTVYSGAAARSQAASTYGRNAYV

>tr|G3PG36|G3PG36_GASAC_Gasterosteus

MAHLCRQVSGSAASCAGWVGIIVATATNDWVRTCDYSVATCVRMDELGSRGLWAECVISP

ALYHCVALSQVLSLPAYIQTSRALMICSCLLGLPSLLLVLMSMPCVRLQNDNAAIKQRRA

RVGGVLFLLMALCGGVSTVWFPIGAHREERLMAFGFSLYAGWVGSGLCLLGGAVILFCRG

IDPGPPSRGNSFYYYSRQGGTATPLDPPANHAKSERV

>tr|G3N8T8|G3N8T8_GASAC_Gasterosteus

MGNMATEIVAFVLTISGWILVSSTLPIDYWKVSSVDGTVITTATFWSNLWKTCVTDSTGV

SNCKDFPSMLALDAYIQVCRGLMIASVCLGFFGAVLALVGMKCTKIGGSETTKARLTVGS

GFHFVLSGLCCMTACSIYAHRITTDFFDPLFVAQKFELGAALFIGWAGSVLCILGGLIFC

FSMPEGSSARRVAGAEYSCAGAASFVTAPNKCRKTVNGLQKEPGQSGRQFGRNAYV

>tr|G3PTZ9|G3PTZ9_GASAC_Gasterosteus

MANAGIQLLGFTLAFLGFIGSIASTVMVEWKASSFSGDNIITAQAMFQGLWKSCVSQSTG

QVQCKVYDSLLQLPGIVQGTRGLMLSSILLCFISMMVCVVGMRCTTCMADQPEQKDKVAL

AGGVVFIIAGLLALVGTSWYGHRIAQEFYDPFTPTNSRYEFGSALFVGWGAACLIIIGGG

FLCCNCSSQSSGKSPRYPPPRSTGPPGKDFV

>tr|G3PD69|G3PD69_GASAC_Gasterosteus

MVEGFCEIAAVCVGLIGLIGAAATTGMPMWKVTAFIGENIVVMETRWEGLWMNCYRQANI

RMQCKIYDSLLFLPPDLQAARGLMCCSLALSGLGLLVAVVGMRCTSCIQDNDRAKNVILM

VAGGMQLAACVCVLIPVSWTAHVIIKDFYNPLLIDAQRRELGEALYIGWVTGAFLFASAM

LFLCRRVASDKASFGVYHQDNLVPYQMRYQPISSVSSHGYDGQQLGPQWQNIPQVMTNGE

VLLNPPVVYNSGLPDNVSVVYQGGTARHPSMRSSSNVGSVYAPGISFHSSQSATPYSHAY

VSDPNASYQSSFHPVPHTPVFIGYETSRIQQRGNNAGVYI

>tr|G3NPU4|G3NPU4_GASAC_Gasterosteus

MVSAGLELMGLSLCVIGTLLVMVSCGLPMWKVTAFIEANIVVAQTIWDGLWMSCVVQSTG

QMQCKVHDSVLSPHRFKTFITSLQLWLTSVSCYVIKYVKARVVNAGGVVSSSALFVLVPL

CWMANSIISDFYNPQVAPSKKREIGAAIYIGWAATALLLIGGSLLCCSCPSSGSSGYSVK

YAPTKRATPNGEYDKRNYV

>tr|G3PD61|G3PD61_GASAC_Gasterosteus

MANSALEIVGLILTLIGLIGTAASTGMPMWRVTAFIGENIIVFETRFEGLWMNCYRQADI

RMQCKVYDSLLALPPDLQAARGLMCCALALAGLGLLIALLGMQCTSCIQNNDQAKRMVLI

VAGSMIIIACICVLVAVSWTAHVIIRDFYNPLLIDAQRRELGEALYIGWVAAAFLLFGGC

MFVCCNLQSEDKGSERYVYSKTSDFMNYPPQRLQPQQLVLLPQTQPQFQPVLSRQPSTNY

SYHSRYPSVRSGVAYL

>tr|G3QB42|G3QB42_GASAC_Gasterosteus

MVSQGLQIMGVLLAFIGWLGTIITCAMPMWRVTAFVGANIVTAQVIWEGLWMTCVVQSTG

QMQCKVYDSMLALPQDLQAARAMVVISVIVGVFGVLMAVVGGKCTNCMEDEAAKAKACIV

SGVIFIIAALLIMVPVSWSAHAVIRDFYNPLVIAAQRRELGAALYIGWGSAGLLLLGGGL

LCNNCPPQNSTRHFHPTKFTPVRARQRQKTVPLSSVNGFV

>tr|G3QAD2|G3QAD2_GASAC_Gasterosteus

MANSSLQMLGFSLSLLGLIGLIVGTILPQWKMSAYVGDNIITAIAMYEGLWMSCAFQSTG

QIQCKVYDSILQLNSALQATRALMIVSIIVIVVGLGAACMGMKCTNCGGDDKTRKTRIAM

AAGIIILIGSLCALIACSWYANDIIKAFYNPFTPVNTKYEFGSAIFIAWAGSFLALVGGG

MLAASCPKGSPKSTPKYPISRPPSSSKEYV

>tr|G3Q501|G3Q501_GASAC_Gasterosteus

MASTGLQLLGLVLAVLGWVCGALVCAAPLWRVSAFVGGELVIAQVLWEGLWMNCLSQTTG

QIQCKTYDSTLALPTSAQAARGLTVLSLLLCLLALMLGVAGAKCTHCMGDANQTSKARLA

RIAGLLFVVSGLAYLIPICWTAHAVIRDFYDPTVAAPLKRELGPALYLGWLASVLLLVGG

SLLHLGSSQPGAGALAFFGGAMKNNPTKGAAGEAKQQEKSFV

>tr|G3PTZ3|G3PTZ3_GASAC_Gasterosteus

MANAGIQLLGFTLAFLGFIGSIASTVMVEWKASSFSGDNIITAQAMFQGLWKSCVSQSTG

QVQCKVYDSLLQLPGIVQGTRGLMLSSILLCFISMMVCVVGMRCTTCMADQPEQKDKVAL

AGGVVFIIAGLLALVGTSWYGHRIAQEFYDPFTPTNSRYEFGSALFVGWGAACLIIIGGG

FLCCNCSSQSSGKSPRYPPPRSTGKNADFR

>tr|G3QB43|G3QB43_GASAC_Gasterosteus

MVSQGLQIMGVLLAFIGWLGTIITCAMPMWRVTAFVGANIVTAQVIWEGLWMTCVVQSTG

QMQCKVYDSMLALPQDLQAARAMVVISVIVGVFGVLMAVVGGKCTNCMEDEAAKAKACIV

SGVIFIIAALLIMVPVSWSAHAVIRDFYNPLVIAAQRRELGAALYIGWGSAGLLLLGGGL

LCNNCPPQNSTRHFHPTKFTPVRAASSNVDYYFSRTFV

>tr|G3PD66|G3PD66_GASAC_Gasterosteus

MANSALEIVGLILTLIGLIGTAASTGMPMWRVTAFIGENIIVFETRFEGLWMNCYRQADI

RMQCKVYDSLLALPPDLQAARGLMCCALALAGLGLLIALLGMQCTSCIQNNDQAKRMVLI

VAGSMIIIACICVLVAVSWTAHVIIRDFYNPLLIDAQRRELGEALYIGWVAAAFLLFGGC

MFVCCNLQSEDKGSERYVQPSTNYSYHSRYPSVRSGVAYL

>tr|G3P405|G3P405_GASAC_Gasterosteus

MVFLTPSMMQRTALFVTLGGLVTSLITTFLPLWKTMNSDLNEVENWYSGLWHTCLYTEEV

GVQCKAYESIMGLPVDLQISRVLMLVSVGTGGLALLAAFPGLEGVAMCMGRPGLKRRLLI

LGGVLSWVSGLATLAPVSLVAYTTVVDFWDEGFPDVVPRWEYGEAMFSGWFGGLALAIGG

TLFFVAVCMADYDQRPGSVADSPQAKHRTKHYLKTEVL

>tr|G3PGS4|G3PGS4_GASAC_Gasterosteus

MAVQELGISLSMIGVAGTILICALPMWKVTAFIGTHLVVMQVFWEGLWMTCVSEYTGQMQ

CKLYDALLDLSPDLQAARGLICIGLVLGCLGFLVFILGARCTNCLDHPGLKARLVVGSGA

TFCLAALVTVVPVSWTANSVIRDFYNPRVPEVLKRELGAAVYVGFVACGLLFCGGVILCT

SCPPQGAGFPAGGYTAARTPTRSSYAIRNYV

>tr|G3NUW7|G3NUW7_GASAC_Gasterosteus

MASTGMQIFGFVLALLGIMGAMVATLLPNWKVSADVGSNIITAISQMQGLWMDCTWYSTG

MFSCTLKYSVLSLPAYLQTARTTMVLCCVMAAMGLCLASLGLKCTRWGGGRRSKRHAAIA

SGGCFVAAGFLCLVPASWFTNEVITNFLDSSVPESNKFEPGGAVYVAFVSAGFLFVGGSI

FCMSCSGKRHGPQDMILLPPPDKLLLQHQYCSLSPLDNKTGYSLQDYV

>tr|G3QB45|G3QB45_GASAC_Gasterosteus

MASAGLQIMGIFLAAIGFIGAIITCALPMWKVSAFIGSNIVEAQTYWDGLWMNCVMQSTG

QMQCKIYPSMLALASDLQAARALMVLSILVSSMGLLLAVVGGKCTNCIEDDAAKSKVAIA

AGVFFIVGGILCLIPASWTAHEVIRNFYSPIMLEAQKRELGASLFIGWGSAGLLLVGGAL

LCCQCRQREEDRFSVKYSAPRKASSGGAYV

>tr|G3PGT7|G3PGT7_GASAC_Gasterosteus

MVSMGRQMLGFALGIIGFLGTIIVCALPMWKVTAFIGANIVTAQDIWEGLWMNCVTQSTG

QMQCKIYDSLLALPQDLQAARALVVISIIVAAMAVILGVVGGKCTNFVSDESSKAKVAIA

SGIIFICAGVLILVPVCWSANTIIRDFYNPILTNPQRRELGAALYIGWGTAGVLLLGGAL

LCSSCPRKDSPEYPVKKANKAKSPQESGWASQRTGFW

>tr|G3QB39|G3QB39_GASAC_Gasterosteus

MSMGLEIVGIALGCIGFIIAIVSCALPMWRVTAFIGANIVTAQTIMEGLWMNCVTQSTGQ

MQCKIYDSLLDLSQDLQASRAMMIIAIILGVLGVMISIVGAKCTNCIEDEPSKAKVMIIA

GIFFLLAGLLVLIPCSWTASVIIRDFYNPILSSPQKREIGASLYIGWGAAALLLIGGAML

CTSCPPKEKKYKPPRMAYTAPRTASAGGGYDRKDYV

>tr|G3PGS9|G3PGS9_GASAC_Gasterosteus

MVSMGRQMLGFALGIIGFLGTIIVCAMPMWKVTAFIGANIVTAQVIWEGLWMNCVTQSTG

QMQCKIYDSLLALPQDLQAARALVVISIIVAAMAVILGVVGGKCTNFVSDESSKAKVAIA

SGIIFICAGVLILVPVCWSANTIIRDFYNPILTNPQRRELGAALYIGWGTAGVLLLGGAL

LCSSCPRKDSPEYPVKYAPARSTATSREYV

>tr|G3PT05|G3PT05_GASAC_Gasterosteus

MAPSVLQNGGFVSGLVGAAALIAATAMNNWSVKDRQGDVVTSVYTYKGLWRDCETTSSGL

TECRPLYGLLGFSGVFQAVRALMIVGVVLAVLAAVISVFSLTCLTMNSMADTTKAKMSLT

AGIMFIIGGVCGIAGASIYANQIVASFRMSNNLNYGGGGGGFGGNMQGGLGGEMGGGMPR

YTFGPALFVAWIGGGVLLIGGVLKSLAFRGMLKEDKSHYPGVVYKPQSRT

>tr|G3PT92|G3PT92_GASAC_Gasterosteus

MSTAVEATGFVMCIVSWLITGSALANDYWKMSTVSGSVIISQRQFENLWHSCAENSAGIA

ECRDFESLLALPAHISASRALMIMSLLLGLGSMVVALLGLKCIKIGSATDQSKAKIAVGG

GVLSMLAGLCCMIAVSWYAYRVVQDFHNPFFGGVKFELSTGLYMGWGGSCLAILGGAFLC

SACKRAAPKGMKR

>tr|G3PGW7|G3PGW7_GASAC_Gasterosteus

SMSVGLELIGICLCILGWIIAILACALPMWRVTAFIGSNIVTAQIIWEGLWMTCVVQSTG

QMQCKVYDSMLALSQDLQAARALTVISILLAILAVLIAIAGAKCTNCIDDEASKSKVMII

SGVFFIVAGVMQLIPVSWSANTIIRDFYNPLLTDAQRRELGAALYIGWAAAALLILGGGL

LCCSCPPRETSPAPKHTQPHLTKEVNAKMVYKAKSEYM

>tr|G3QCC9|G3QCC9_GASAC_Gasterosteus

MASAALELMGFFLGLLGMLGTLVATVLPYWQISAHIGSNIVTAVANMRGLWMECVYQSTG

AFQCETYNSMLALSPDLQASRALMVISLVLSVLAVAVAALGMQCTLCLEGAGAAKSRVAG

ASGGLFLTAGFLSLIPVSWTTHEVVQTFYRPDLPSSMKFELGECLYVGLASSLVTMLGGG

MLSASCLPATAPSPPHTEPRGPRSPPRQDMTSQKSYT

>tr|G3PJY1|G3PJY1_GASAC_Gasterosteus

MGSAAVQIVCVSLGVLGMIGVIISCAVPLWKVTSFTGSNIVTAQSSQEGLWTTCVVQSTG

QQQCKNYDSLLELPSELQARALTIISCMFCCLSLLILFCGARLTNFVENEERQSPKMSLV

AAVGLLLAGLLVIIPVSPPTLFAKTFNNPLVAATQKRELGACIYVGWGPGMLLLLSGGLL

CCFSRSKSGGSGGTAKYYSNGAASAPNKNYV

>tr|G3PD54|G3PD54_GASAC_Gasterosteus

MRAKTEVFALVLGFFGLIGTIAVTALPMWRVSAFIGANLIVMEELWEGLWMNCYRQADFR

MQCKVYDSLLILPPELQAARGLMCVSVVLVVVSLSVTACGTKRSSCCDDNTKGKNVTLAL

GGGLHLLSFLTTLIPVSWVAHTVVSNFYNPTVLDSQKRELGKALFVGWATSGVLLISGVI

LLFSYSRRRSKEEEPYADTHLMDVTDAPKDGSVYLERKPSSFYEHQQYV

>tr|G3PAF1|G3PAF1_GASAC_Gasterosteus

MANSGLQLLGYFLALGGWIGIISTTSLPQWKQSSYAGDAIITAVGLYEGLWMSCASQSTG

QVQCKIFDSMLSLDIHIQTCRALMVISVLLGFIGIIVSVVGMKCTKVGDNNPVTKTRIAV

TGGALFLLAGLCTLVSVSWYATQVSYQFFNPNTPPNARYEFGSALFVGWAAASLTVLGGS

LLCCSCSKEDIRGQQYYRQSQPSTARDSPPNREHQTCLAPTLLSQI

>tr|G3PD56|G3PD56_GASAC_Gasterosteus

MASYTAYTGYSKPPQSYPGSFYDEKQAKTYADYQDSLYEEKKRNEKKDRRDAICCEVVAL

IIGFIGLIGVAAVTGLPMWKVTAFIEENIITMETRWEGLWMNCYRQANIRMQCKVYDSLL

FLPPDLQAARGLMCSSVAVATIALVVSAVGMKCTKVVDHRARTKHIVLVSGGCLFLLACL

TTIIPVSWTAHVIIRDFYNPLLIDAQRRELGEALYIGWVTSALLFTAGVILLCRHAPRTQ

DPEEKIVYNPDGNRYPPAYT

>tr|G3QB47|G3QB47_GASAC_Gasterosteus

MASAGLQIMGIFLAAIGFIGAIITCALPMWKVSAFIGSNIVEAQTYWDGLWMNCVMQSTG

QMQCKIYPSMLALASDLQAARALMVLSILVSAMGLLLAVVGGKCTNCIEDDAAKSKVAIA

AGVFFIVGGILCLIPASWTAHEVIRNFYSPIMLEAQKRELGASLFIGWGSAGLLLVGGAL

LCCQCRQCEEDRFSVKYSAPRKASSGAYVKLSPSPKAYTRQRFL

>tr|G3QB44|G3QB44_GASAC_Gasterosteus

MVSLGLQILGVGLAVLGWIGNILICMLPLWKVSAFIGNNIVVAQTIWEGLWMSCVVQSTG

QMQCKVYDSLLALPTDLQAGRAMAVIAILLSLFGLLLSVVGGKCTTCIGDQVVKARVVIS

AGVFFVLSGALCLVTVSLPANTIIKDFYNPMVPDSQRRELGACLYVGWGASGLLMIGGAL

LCCQCPSGGDRYNGAKYSAPKSTTPAAAQQSPSRYKVQHRV

>tr|G3PGU0|G3PGU0_GASAC_Gasterosteus

MGRIGKEVAGQLISFIGFVGVAVTTGIPMWRVTSFIGANIVTGQVIWDGLWMNCVMQSTG

QMQCRLNESLQSLARDLQASRALVIISLIFGFIGFLITFIGAKCTSSLKKDSSKAKVVIL

GGCLILVSALLIVIPVTWSAVITITDFQNPLTIGSQKRELGAAIYIGWAAAAFLLIGGII

LTTSCPPQKNGYGYPGYAPAPMYPYGGPGGNQPVYGPMYAPASSQPYTGTGTYLPSKPYA

APTAYSARQYL

>tr|G3PGV1|G3PGV1_GASAC_Gasterosteus

MYSAGLEILGMILAVAGWLGVMVACGLPMWRVAAYIGQNIVISQVIWEGLWMNCSVQSTG

QMHCRVHDSMLGLPVDLQAARALVIVSMVLCVVGIGLSVAGAKCTNCSRDVSSKPRLVVT

AGVTFVVAGLLLMVAVSWTAHAIVLGFYDPLLQETGKREFGNALYFGWASSCLLLVGGAL

LCCSCPPRAAAAPSGSGSAAPRVDYSAVKMLSVNGYPRRDYV

>tr|G3NJF4|G3NJF4_GASAC_Gasterosteus

RVISEMGKRLIQVLGFLISSLGWVFVLCTMAMDFWRTSQLGGQGGSFIIKVAWYWSNLWK

DCYTDSTAVTNCRDFPVLWSVKPFVQAVRGLLMCGLTLGFVAVILCFVGMECTYIGGAQK

IKDKLLLAGALFHIAGCVSDVSAYCLYINRVARTAFAGRVGPGVLRYDLGPPIFLGLVGC

FLILLGGIFYAVTVYRVIFPKSKVVYAYGGGTYMAPRSRGRTMYSGYYRPSRQNGPYRGS

QLSKSSRISGLSQTTRTNISERDAFV

>tr|G3QB46|G3QB46_GASAC_Gasterosteus

MQTQLVGVGLAIIGFIGTILICGLPMWKVTAFVGASIITAQVFWEGLWMNCVIQSTGQSQ

CKAYDSVLALPRDLQASRALICVSIAVSVVALGLTVVGARCTNFFSRDRLAKANIGLAGG

LVFVLAGVLCIIPVSLSAYSIITGFYNPLPTSERRGELGASIYVGWASGALLVIGGGVLC

STYRC

>tr|G3PGS5|G3PGS5_GASAC_Gasterosteus

MVSAGLQIAGCAMALLGWIGVIIICGAPMWRVTAFIGSNIVTAQVTWEGLWMTCVVQSTG

QMQCKVYDSMLSLSPDLQGARALVVVSIVVGIAGIFVALAGGKCTNFIADERAKARASVA

AGVLLIISGLLCLIAVSWTASIIIRDFYNPVLLDSQKSEIGASLYIGWGAGALLVLGGAL

LCATCPPKEDKSPSVKYLVNRSGGNSKEDSTFLPDKTYI

>tr|G3Q6F3|G3Q6F3_GASAC_Gasterosteus

MDQILEVVALLLGFIGWVMVGVTLPNRYWRTSTVDGNVITTSTIYENLWMSCATDSTGVH

NCREFPSLLALSGYIQASRALMITSIVLGTCGLLAALIGVKCSKAGGENYVLKGRIAGTA

GVLFILQEGMCTMVAVSWYAFNITQEFFDPFHPGIRYEIGEALYIGWCSAVLAIAGGACL

TCSCKMGTKEKY

>tr|G3PAG1|G3PAG1_GASAC_Gasterosteus

MANSGLQLLGYFLALGGWIGIISTTSLPQWKQSSYAGDAIITAVGLYEGLWMSCASQSTG

QVQCKIFDSMLSLDIHIQTCRALMVISVLLGFIGIIVSVVGMKCTKVGDNNPVTKTRIAV

TGGALFLLAGLCTLVSVSWYATQVSYQFFNPNTPPNARYEFGSALFVGWAAASLTVLGGS

LLCCSCSKEDIRGQQYYRQSQPSTAREYV

>tr|G3PAD6|G3PAD6_GASAC_Gasterosteus

MANSGLQLLGYFLALGGWIGIISTTSLPQWKQSSYAGDAIITAVGLYEGLWMSCASQSTG

QVQCKIFDSMLSLDIHIQTCRALMVISVLLGFIGIIVSVVGMKCTKVGDNNPVTKTRIAV

TGGALFLLAGLCTLVSVSWYATQVSYQFFNPNTPPNARYEFGSALFVGWAAASLTVLGGS

LLCCSCSKEDIRAFTHGIHRYSSPNTNTSSTGGIYFGDKKNYT

>tr|G3QB41|G3QB41_GASAC_Gasterosteus

MVSAGLQMLGAVLGVLGWIGAIIVCALPMWKVSAFIGSNIVTSQITWEGIWMNCVVQSTG

QMQCKVYDSMLALSSDLQAARALTIISIVVGILAILLSVAGGQCTKCLEDKSSKTKVGIT

AGVMFIIAGVLCLVPVCWAAHTIIQDFYNPLLVGAQKRELGAALYIGWGAAALMLIGGGM

LCCNCPPKEEDSYTANTGGNLGSL

>tr|G3PAG4|G3PAG4_GASAC_Gasterosteus

MANSCLQLSGFLISFLGWLGIVIAISTNDWVIMCKYSLNTCKKTDELETKGPWAECVIST

GLYHCFSRSQILDLPAYIQTTRALMIAGSILGLPAVAMILMSMPCINFGNEPQGPKNKRT

VLGGVLILVVALCGMVSTVWFPIGAYQEHGLMSFGFSLYTGWFGSIFCLLGGCILTCCSS

ESSSSRSYQDNNRYYYSKQGGGNPPAAPSTNHAKSAHV

>tr|G3PGU9|G3PGU9_GASAC_Gasterosteus

MVSAGFQMMGTALCIIGWIGAIVVCALPMWKVTAFIGQNIVTAQTTWEGIWMTCVVQSTG

QMQCKVYDSMLALSSDLQAARALIIIAIMLGVIGILLSIAGGKCTNCVEDEVAKAKIGVG

SGVIFILAGILCIIPVCWTANTVVTEFYNQLVMASQKKELGAALFIGWGASGLLILGGAL

LCANCPPKDNYTAKYAAARPNAPKDFV

***Tilapia (Oreochromis niloticus),***

>tr|I3K1G2|I3K1G2_ORENI_Oreochromis

MGNMATEITAFLLTISGWILVSSTLPTDYWKVSSVDGTVITTATFWSNLWKTCVTDSTGV

SNCKDFPSMLALDAYIQVCRGLMIAAVCLGFFGAILALVGMKCTKIGGSKTTKARLTILS

GFHFILSGLCCMTACSIYARRITADFFDPLFVAQKFELGAALFIGWAGSVLCILGGLIFC

LSLSEGFSIRAEYSYNGAASFPTTRNKTSKPAQRPNKETQEPLRQFGRNAYV

>tr|I3J0C5|I3J0C5_ORENI_Oreochromis

MANSGLQILGFALALLGMIGLVVGTIMPQWKMSAYVGDNIITAISMYEGLWMSCAFQSTG

QMQCKVYDSILQLNSALQATRALMIVSIIVTLAGLGVACMGMKCTNCGGDDKVRKSRTAM

GGGIIILIGALCAIVACSWYANDIIRAFYNPFTPVNTKYEFGSAIFIAWAGAFLTVVGGG

ILAASCPKSKGRQSGPKYPISNPRSTGSNKDYV

>tr|I3IZV1|I3IZV1_ORENI_Oreochromis

MAASGLQLLGFFLSLVGVAATAAATVMVDWKKLVQGKHRSYEGLWMSCGGVQDRATCEIY

HSFIKLPDEIQATRIALPLSLFLSALAMLVSTVGMKCTHFMDGASQSKSVTAMIGGIMFM

LSGLLTIIITSWFVKVVLQNFYDSPPLIRSEFGNAVFVSWAGGLLTLTGGAFLSCRRCAT

TSSMSISSSRLLPTSDPKSNYV

>tr|I3L011|I3L011_ORENI_Oreochromis

MANSALEIVGLLVSLIGLIGAAATTGMPMWRVTAFIGDNIIVFETLFEGLWMNCYQQASF

RMQCKVYDSLLALTPDLQAARGLMCCSLALSGLGVLISLIGMQCTSCIQNNDRAKRLVLI

IAGSMIILGCFCVLIPVSWTGHVIITNFYNPLLIDAQRRELGAALYIGWVTSAFLFAGGC

IFICCNIQSDDKDSERYLYSKASDYSVYPQQPLQPLQPQQLMTLPPQPRSQPMLSRYPSS

IYSYQSRYPSVHSAHSGVAFL

>tr|I3KZ74|I3KZ74_ORENI_Oreochromis

MASTGLQLMGLVLALLGWVCGALVCAAPLWRVSAFVGGELVIAQVLWEGLWMNCLSQTTG

QIQCKTYDSTLALPMFTQAARSLTVLSLLLCLLAIILGVSGAKCTHCMGDNNQDTKARLG

RIAGVLFIVAGLAYLIPICGTAYAIIRDFYDPNIAAPLKRELGPALYLGWGASILLLVGG

SLLHVGSPPPGARAMPIVGRPAKDSQQTVATGEVKQQEKSFV

>tr|I3KZL5|I3KZL5_ORENI_Oreochromis

MLSAAVQILAFALALLGVLGAVVATLMPNWKVSINVWTSIMTPISQMQGLWMDCVWYSSG

IFSCTMKNSVLSLPPHLQAARAGMVLSCVVALFGLCLATLGLKCTRWGGSHRAKGHTAIA

AGGCFILASLLCLIPASWFTNEVITAFLTTDLPDSSKYQPGGALCITFISAGFLLAGGVI

FCLSCPGKRTRRPDRASPADPDRFTLHPDERRRQGLQAERNQPKNRKKTV

>tr|I3KQ06|I3KQ06_ORENI_Oreochromis

MANSGIQLLGFFLSLIGIVGLIIGTILPQWKMSAYIGDNIITAVAMYQGLWMSCAFQSTG

QLQCKIYDSILQLDSSLQATRALMIVGIIVSIAGLGVACMGMKCTTCGGSDRLRKARIAM

TGGIILLVGGLCAIVACSWFAHNVIRAFYNPYTPVNTKFEFGAAIFIAWGGSLLDVLGGA

MLAASCPRKKQVSKYPAIGSSRSGPPSSTKEYV

>tr|I3JQ63|I3JQ63_ORENI_Oreochromis

MASFGLELAGITLAVLGWALTVTSCALPMWRVSAFIGANIVTAQVYWEGLWMSCVVQSTG

QMQCKIYDSMLALPQDLQAARALTVVAILVGLIALLIAMVGAKCTNCIEEDGVKAKVMIS

SGVAFIIAGLAQLVPVSWSANTIVAEFYSPIVPAGQKMEIGASLYVGWAASALLVTGGAI

LCCSCPPKEDKPLKYNIQPQSRAAYSAAPMSAAQSSYTRRDYVHAESQM

>tr|I3IZU9|I3IZU9_ORENI_Oreochromis

MATSGLQLLGFFLSLAGVAATVAATVMVDWKKLVQGKHRSYEGLWMSCAGVQDRMTCEIY

QSILKLPTEIQVTRAVLPVSLLLSALAVLISTVGMKCTHFMDSVPKSKSMTAMIGGITFI

LSGVLTIIITSWFVNMVLSYSGHKLFRYCHRPEFGNAVFVSWVGGLLTAAGGAFLSC

>tr|I3JS66|I3JS66_ORENI_Oreochromis

MDPIVEVVAVILGFIGWIMVGVAIPNRYWRTSTVDGNVITTSTIYENLWMSCATDSTGVS

NCRDFPSMLALNGYIQASRALMIASVVFGTFGLLGSLIGIQCSKAGGENYVLKGRIAGTS

GVLFMLQGLCTMVAVSWYAFNITQNFFDPYYPGTKYEIGEGLYIGWCSAVLAIAGGAVLT

CSCKLGTEEKYPLPHARGTVYSGPAPTRSMAASTYGRNAYV

>tr|I3KT07|I3KT07_ORENI_Oreochromis

MAATLCQGLGFVLSLIGIAGIIAATGMDQWATQDLFDNVVTAVYSYSGLWRSCVRQSSGF

TECRPYFTILGLPALLQAVRALMIVGIVLGAIGLLIAIFSLKCLKMGNMEDNLKATMTLS

AGIMLLLAGVCGIAGVSAFANLIVQSFQFTTYASGFSGTANVGGLTGTLTPRYTFGPALF

VGWIGGAILFIGGILMCLACRAMTPEDHRYDGMAYKAASQNTMYRPDTRSRPVYNDSYKA

QSMDGRQTNQRFDYV

>tr|I3KMC2|I3KMC2_ORENI_Oreochromis

MSQMRRQLLGTAASGVGWVGVIVATATNDWVRTCDYTKTTCVRMDELSSRGLWAECVISP

SLYHCVGLTQILTLPAYVQTARALMICACLLGLPAMLLILMSMACVRLQNDTSAVKKRRA

RVGGVLFIFMAVCSIISTVWFPIGANKEEDLMSFGFSLYSGWVGSALCLVGGIIILCCQG

IDPGTPSRDNSFYYSGNRGTATLLDPPANHAKSARV

>tr|I3JP55|I3JP55_ORENI_Oreochromis

MSSMFQEIVAFFFSTAGWILESSTLPTDYWKVSSDDGSVITTASFWSNLWKTCVTDSTGV

SNCKEFISMLALDGYIIACRALMISAVILGFFGSIFALVGMKCTKIGGSDKNKARIACFA

GVHFILSGLCSLSSCSIYAHRITSEFFDPMFISQKYELGAALFIGWGGSVLCILGGTMLC

FSIAGSFTSSKREVNYIYRGAASHSRISSYPKGQAKSVNQRLTPDNSTSSRIQHFDKNVY

V

>tr|I3KZR4|I3KZR4_ORENI_Oreochromis

MLTACLEVLGLALCVTGSLLAMVACGLPMWKVSAFIDSNILVAQTLWEGLWMSCVVQSTG

QMQCKVHDSMLALAQDHQTARALTVISGVLGIVALAVTVAGAQCTNCIRDETIKSRVVHA

GGVIYIISGLFMLVPLCWMAHRIILDFNNPNIPSSKKREIGAAIYVGWAASVLLLLGGTL

LCCSFSQMVRGANSMKYTPTKTIAVNEEFHKNLYV

>tr|I3J9B9|I3J9B9_ORENI_Oreochromis

MASMALQLLGFFLGLLGFAGTIVATLLPHWRSRAHVDANIITATGYMKGLWMECVWRSTG

IYQCELYRSLLALPPDLQAARALMVISCLTSVLASLVSVIGMKCTRFARGSLIKSPLVLS

GGICFLCAGLLCLVTVSWTTNEVIKNFYNPLIHSGMKYEIGLAVYLGYASACLSLAGGMV

LCWSSSGDRSRQSPPHMQRGQSSHLPPVFNNLHPPAPPYRPPKALKGNHAPSLCSASSSG

YRLNNYV

>tr|I3JEW1|I3JEW1_ORENI_Oreochromis

MVSMGRQMLGFALAIIGFLGTIIICAMPMWKVTAFIGANIVTAQIIWEGLWMNCVMQSTG

QMQCKIYDSLLALPQDLQAARALVVIAIIVAFMGVILGIAGGKCTNFVEEQRAKSRVAIA

AGVVFICAGVLVLIPVCWSANTIIQNFYNPTLINAQRREMGAALYIGWGTAALLILGGAL

LCSSCPPKESPEYPVKYGGARSIATSRAYPGKKYI

>tr|I3KZ52|I3KZ52_ORENI_Oreochromis

MASQGLQILGVLLAFFGWLGAIITCSLPMWRVTAFVGANIVTAQVIWEGLWMSCVVQSTG

QMQCKMYDSMLALPPDLQAARAMVIISVIVGIFGILMAVIGGKCTNCMEDDVAKAKTCIV

AGVIFIITALLILIPVSWSAHAVISDFYNPLLVEAQRRELGSSLYIGWCSAAVLLLGGGL

LCSSCPPKDGGRPYMPAKFTPVRSVSSNVDYV

>tr|I3KYJ7|I3KYJ7_ORENI_Oreochromis

MLGTLVATVLPYWQISAHVGSNIVTAVDNMRGLWMECVYQSTGAFQCETYNSMLALPSDL

QAARALMVISIVLSILAIAMATLGMQCTLCLESSGGVKSRVAGAGGVLFVFAGFLALVPV

AWTTHEVVKTFYLPNVPQSMKYELGECLYTGLASALISMLGGGMLCVSCCEEQEGGRGRR

HGGGYPYPVGGGISSTGVRTTSQTYRNPTLQVGGVNTPSRGQTLIRSASGSSESGMHGAQ

RPRKPTAAGYDITGYV

>tr|I3JDY4|I3JDY4_ORENI_Oreochromis

MIQGASELAAMCVGLVGLIGASATTGLPMWKVTAFIGENIIVMETRWEGLWMNCYRQANI

RMQCKVYDSLLYLPPELQAARGLMCSAVALSAVGLLVALAGMRCVSCIRGNDWAKTIILM

VAGVMQFMACICVFIPVSWTGHVIITDFYNPLLIDAQRRELGEALYIGWVTGAVLFASSL

LFVCRRLPKGKGSFDVYQPPNLLSYKPMPNRPALMNYHPLSTLSSLRSTGSVGQNAAMPL

QTVIPVRTNNPTVVNPPVVYNPTHQENASLLYQGRNSSYMSQNTTPYSLTYVPTTSYQSS

FHPVPQTLIFITYKESRTESVPYSEMRGGKI

>tr|I3KXT5|I3KXT5_ORENI_Oreochromis

MHTPASMVMGIVFAPLGLVLVFTAAITPQWREGQAHLGMAGPGSLLRSGTKSHGTRSGSV

EALILLRSDGLWESCLQVERSDLRQCWPVAGPYQRDPRVRLAQGLVLTSLFLCGTGIVLA

CIGVRCWADLPLRGVAASGGLLVVTAGLLSLTALGVYTHNLKRLGMDPSQRIHNPRYPHL

NLHPAGSLYFGWLGSCLQVLGGTALLFSFKRPRCPTCPCRHELSVSPCHSCPEMTNKPDM

DVYEVSC

>tr|I3JDY2|I3JDY2_ORENI_Oreochromis

TMQAKSEILALVLGALGLCGTIVTTCLPHWRVSAFIGANIIVMEELLEGLWMDCYRQANI

NIQCKVYDSMLILPPELQAARGLMCVSIVLAFISLKSKNITYALGGSLFLLSCLTTLIPV

SWVAHTVISNFYNPETLDARKRELGASLFIGWATSGVLLITGIIVLFRYSKRSAKEDQRY

IEVYHIVKRNEQKADSVDFSRAKSSFHSNQEYV

>tr|I3KXC0|I3KXC0_ORENI_Oreochromis

MGLQELGISLSMTGVAGTLLICALPMWKVTAFIGTHLVVMQVFWEGLWMTCVSEYTGQMQ

CKLYDALLDLSPDLQAARGLICISLVLECLAFLIFLLGARCTNCLGHPQIKARVVLSSGA

IFCLAALTSIVPVSWTANSIIRDFHNPRVPEVLKRELGAAIYIGFVTSGLLFCGGAILCT

SSPPRRDRFPSSGYTLAKAPTRSSYAIKNYV

>tr|I3KZ53|I3KZ53_ORENI_Oreochromis

MASLGLQILGVGLAVLGWIGNILICMLPLWKVSAFIGNNIVVAQTIWEGLWMSCVVQSTG

QMQCKVYDSLLALPPDLQAARAMIVISILFSLFGLLLSVVGGKCTTCVGDKMAKARVAIS

AGFFFILSGALCLVTVSLPANTIIKDFYNPMVPDAQRRELGACLYVGWGAAGLLLIGGSL

LCCQCPSGDDRYSGPKYSPPKSTTPGKEFV

>tr|I3KXC5|I3KXC5_ORENI_Oreochromis

SMSIGLELIGISLCILGWLIAIVACALPMWRVTAFIGSNIVTAQIIWEGLWMTCVVQSTG

QMQCKVYDSMLALSQDLQAARALTVISILLAILALLIAITGAKCTNCIEDEASKAKVMII

SGVFFIVSGVMQLIPVSWSANTIIRDFYNPLLTDAQRRELGAALYIGWAAAALLILGGGL

LCFSCPPRETRYNPSRMAYSTPRSVGAQGLERKDYV

>tr|I3JWY5|I3JWY5_ORENI_Oreochromis

MANAGVQLLGFILAFIGFVGIIAATIMVQWKASAYVGDNIITAQAMYEGLWQTCASQSTG

QVQCKVYDSLLQLPGNVQGTRGLMVSSIVLCFISILVAVMGMKCTTAMSDSPDLKDKVAL

SGGIVLIIAGLLALGGTSWFGSNIAKEFYNPFTPTNSRYEFGSALYVGWASAILVIIGGA

FLSCHCPKQDSGPARRYPQSHSAGGKDYV

>tr|I3J5S2|I3J5S2_ORENI_Oreochromis

MANSGLQLLGYFLALGGWIGIISTTALPQWKQSSYAGDAIITAVGLYEGLWMSCASQSTG

QVQCKIFDSMLSLDIHIQTCRALMVVSVLLGFIGIIVSVVGMKCTKVGDNNPATKTRIAM

TGGALFLLAGLCTLVSVSWYATQVSYQFFNPNTPPNARYEFGSALFVGWAAASLTVLGGS

LLCCSCSKEDMRGQQYYRQSQPSTAREYV

>tr|I3KX91|I3KX91_ORENI_Oreochromis

MVSAGLEILGLSMCVIGSLLVMVACGLPMWKVTAFIEANIVVAQTLWDGLWMSCVVQSTG

QMQCKVHDSVLALSHDLQAARALTIISSVMGVLGLMVVIAGAQCTNCIRTEYIKARVVNA

GGVIYILSGLFVLVPLCWMANNIISDFYNPQVPASKKREIGAALYIGWAATALLLIGGAL

LCCSCPSSGNSGYSVKYAQTKRAATQNGDYDKRNYV

>tr|I3KYP6|I3KYP6_ORENI_Oreochromis

MATTGMQLLGLILSIVGWVGGLIVCAIPLWRVTAFIGNNIVTAQIIWEGLWMNCIVQSTG

EIQCKVYDSLLALPSDMQAARGLTVLSILLCGLALALGVLGVKCTKCIGVNSLKARIARI

SGSLFAIAGFLYLVPVCWTAHSIIRDFYDPHVAAPHKRELGPALYIGWGASALLLIGGSL

LYAGSSPPGIPGSPTFSSGESSPRRAPATQVKGYV

>tr|I3KZ54|I3KZ54_ORENI_Oreochromis

MASAGLQILGIFLAVIGFFGDIITCALPLWKVSAFIGNNIVTAQVFWEGLWMNCVMQSTG

QMQCKVYDSMLALPQDLQAARALVVISILLSLMGLLLAIAGGKCTNCIEEETAKSKVAIA

AGVVFIVGGILCLIPVSWSANEVIRNFYNPIMSDGQRRELGASLFIGWASSGLLIIGGAL

LCCQCKQPKDGGYSVKYSAPRSATGGGAYV

>tr|I3K375|I3K375_ORENI_Oreochromis

MSTAVEAVGFIMCIISWLVNGAALVNDYWKVSTVSGSVIISQRQFENLWHACAENSAGIA

ECRDFESMLALPVHVQACRALMIICLLLGLGSMIVALLGLKCIKIGSTTEQSKAKIATTG

GILSILAGLSCMVAASWYAARVVNDFNDPFYGGMKFELGTGLYMGWAGASLAMLGGAFLC

SACKRASPKKGGYYGNAPQKVYTPTAKSEPDAKAYV

>tr|I3J8P5|I3J8P5_ORENI_Oreochromis

MNTLIEVVALFLSFVSWILTLITLEDQHWRDSSQDGSVILTSNLYENLWMSCASDSTGNY

NCRNFPSLFALPGYIQASRALMITSIVFGSFGLVATLVGMKCSKIGGENYVLKGKIAAVG

GVFFLLQGLCTMIAVSWYAANITQQFFDILYQGTKYELGEALYIGWASATLAICGGSCLL

CSCGIMTDNEKIPYPVQPSSRGHVLSTVAPSQSVPSNYDRNAYV

>tr|I3L003|I3L003_ORENI_Oreochromis

MVAGCRQLLGLFLGVIGFLGSIITCGLPLWRVTAFIGANIVTSQVIWEGLWMNCVTQSTG

QMQCKVYDSMLELAQDLQAARALTVIAIIVGVFGILLGVVGGKCTNFVEDEVQKSRVAVA

AGVVFIIAGLLVLIPVCWTANTVIRDFYNPTLIAAQKRELGASLYIGWGTAGVLILGGGL

LCSSCPPNDATDYDIRYSKARSVDSHKEYV

>tr|I3L010|I3L010_ORENI_Oreochromis

MATYTAYSGYSKPPQSYVGSYYDGKPALTNGEYGEYGDSVYEEKKKIEKRKRRNAICCEV

VALIIGFIGLIGVAAVTGLPMWKVTAFIDANIIVMETRWEGLWMNCYRQANIRMQCKVYD

SLLFLPPDLQAARGLMCSSVALTTFALIVSAVGMKCTKVVDHRARTKHIVLVAGGCLFLM

GCLTTLIPVSWTGHVIIQDFYNPLLIDAQRRELGEALYIGWVTSALLFSAGVILLCRHAP

RTQDPDEMIQYNPTGSLYQPAYTNQPAYTYQPAYAYQPAYSAAPPGSVIYAPSQY

>tr|I3K1F9|I3K1F9_ORENI_Oreochromis

MKYRTVVMYVEIGCFVSCLCGWILVSSTLPTEYWTFSEVGTLVLTTSNYYSNLWMDCISD

TTGVSDCKYYPSMLALPAFLHACRALAVCAVITGFFGGVLTLIGMKCTKIGGTEIVNSRV

TFSGAITYLASGGCGMITYSWWANRVITEFKDPNFRAQKFELGAAIFVGWGGSLLLICAG

AVLAYLSGKELVPSCISSNQRPERPVSYATARTMRTYMMPPPSSRVTLVPPLYNKGMNGR

TTRATNKTDQTLGRDSFV

>tr|I3JP62|I3JP62_ORENI_Oreochromis

MRKRLIQISGFLISSLGWLFVLCTMAMDYWRITKLGGQGGSFIIKVAWYWSNLWSDCYTD

SAAVTNCRDYPVLWNVAYYVQAVRGLLMCGLTIGFFAVVCCFVGMECTYIGGSDKTKDKV

LFAGAVLHFVGGMSDIAAYCLYINRIARTAFARRVASGVLRYELGPPVFLGLAGCFFILV

GAVLYAVTVYKVIRRGSRVVQADDAHTYMAPPSRGRTIYSGYYKPSKLNGSNYMGSGRSS

AQLSKLSQATPTKPSDRDAFV

>tr|I3KXC1|I3KXC1_ORENI_Oreochromis

MGRIGKETAGQVISFIGLIGVAVACGIPMWRVTSYIGANIVSGQIVWDGLWMNCVMQSTG

QMQCKLNDSILTLTSDLQAARALVIISLVFGFIGFIISFVGAKCTSCLEKDASMANVVII

SGCLIIVSAILILIPVCWSAAFTISDFENPLTIATQKREIGAAIYIGWGSTVLLLIGGII

LTTSCPPQKQMYGYPSYQQPVYAYSRPNSAAYGRVYAPPSNQPYTVAGGYAPSKPYAAPA

SYPASQYL

>tr|I3KXC2|I3KXC2_ORENI_Oreochromis

MVSQGIQITGIAMAIVGWLLDIVACGLPMWKVTAFIGANIVTAQTFWKGLWMDCVVQSTG

QMQCKVYDSMLALESDLQAARAMIVISILVGVFGLILSVAGGKCTNCIEEEQAKAKVCIC

SGVLFIISGFLCLIPVCWSANTVITNFYNPMMMNLQRYELGSALYIGWAATALLLMGGGL

LCWNCPPKHERPHYVPKYAPGKSVSTSREYV

>tr|I3JWT6|I3JWT6_ORENI_Oreochromis

MAPSVLQNGGFILGLIGATALIAATAMNNWSVKDRQEQVVTSVYTYKGLWQDCETTSAGL

TECRPLYGLLGYSGTFQAVRALMVVGVVLSVLGALISVFSLSCLTMNKMADSAKAKMSLT

AGIMFVIAGVCGIAGASVYANQIVESFRTSTYTSYTQEMMGEGMNMGMGGVGITTYTFGP

ALFVGWIGGGVLLIGGILKSLAFREMIKDEKPRFFSAICFLCSYPGVVYKPQSNNKTEVM

DHRSEGGKKEQQYV

>tr|I3JU02|I3JU02_ORENI_Oreochromis

MGSAVVQMVCVALGVVGLIGIVVCCAVPLWRVSAFTGSNIVTAERSQEGLWTTCVVQSTG

QQQCKNFESLLDLGADRQAARALTIISGMLCCLSLLVLFCGSDFTTCLENEDVKPKISLM

AGVGMLLAGILVIIPVSWFAHNTVRDFYTPLLSSSPRMELGACIFVGWASGVLLIIAGGL

LCCFSRPKQGSSGGKAKYYSNSASAPNKNYV

>tr|I3KYD2|I3KYD2_ORENI_Oreochromis

TCACALELLGMLVYVGAWLCALTTTILPQWLTMSTALLPVESYELGLWETCVVQDVGGME

CRAYDSLLGLSTDLKLARILMCAALAVGMLGILVGIPGLDLVHSCKEDSDQQTKRILIIT

GGVLGMVSGVLCLIPVSYMAHIAVMHFFDDKVPDVVPRWEFGDALYCGWAAGFLLVVAGV

ILITSSSCSQVEPQPVLQRRYQVMGTGVNYRKRTEYV

>tr|I3JP63|I3JP63_ORENI_Oreochromis

MDYRTGVMYMEIGCFVVCVCGWILVCSTMPTDIWTWSELSNVVLATSSYYSNLWKDCISD

STGVSDCKGIPSMLALNWDIHMCRALIIICIILSFFGSILVLVGMKCTKIGGSELANARV

TFAGGMNYLIGGMCSMIAYSYYGNKIRAEFEDPNYMEQKFEIGVGVFIGWGGSTLLIVGG

LIYSIFAGREGCHSSSKRYATHQFQDAYTFVPTQKSVVSLAPTEISRSKKPRTISSSRAS

SISEINSVTKTSVTYSYV

>tr|I3KZ56|I3KZ56_ORENI_Oreochromis

MTAGVRHLLGLALAIVSFVGTIVICALPQWRIEKIFWESSNVAPQVFIEGLWKVCGPYSA

HTECKEYEEPFVSISQDLEVARALIVIAIIVGVIGILLGAAGSKFINFVPDERKRSKIAM

ASGVVCIIAGIFVLIPISWTTITTTTIQNNFNNFLPHGTLELGASLYIGWITTALLFLGG

GLLCSSFICQDGTD

>tr|I3KXC4|I3KXC4_ORENI_Oreochromis

MYSTGLEILGIILAVAGWLGVMVTCGLPMWRVAAYIGQNIVISQVIWEGLWMNCSVQSTG

QMHCKVHDSMLGLPVDLQAARALVIISLVLCIVGIVLSVAGAKCTNCSRDVSSKPRLMVA

AGVTFVVAGLLLLVAVSWTAHAIVLGFYDPMLEETGKREFGNALYFGWAASCLLILGGAL

LCCSCPRRSTTAPSGGGPAPS

>tr|I3KZ50|I3KZ50_ORENI_Oreochromis

MSACCRHVLGLLLSIIGFLGTIFICALQMWKISGFIGSSITISEIHWEGLWLKCVMRSTG

EMQCKSYNSIQDLPQDLQDARVLIVFAIIIQFFGIVLGVVGAKCTNFIPDSRRKTKAAIA

SGVLLLTAASLVALPVQWTTHAIVNDMFNPDLANNQRTVLGASLFIGWVSAGLLYLGGAL

LCCSFFCRNETECDVKSAHTNTKQQRSSRQRSPRNKEKVS

>tr|I3KZS1|I3KZS1_ORENI_Oreochromis

SSNGTLELLGVGLAIVAWICSLTTILMPSWITLSTDLLPTEETNMGLWVTCVSQEAMDRL

ECRPYDSMFDLPPDIILARILMSLVLGVGLLGVLLAIPGIRLVNGCHNPLDDFRCKRVLK

AIGGALCLVSGILGLIPVSYVAHLTLVRFFDETVPQIVPRWEFGYALFCGWIAGILHLVA

GSLLLISCLHLQKLDRPVHIPLGIVTPGPNFERTRTEYV

>tr|I3KZ55|I3KZ55_ORENI_Oreochromis

MADCKLPGLVLAIIGVLGSIITCVLPMWRVTAFIGANIVTAQIIQEGLWMSCVIQSTGQM

QCKVYDSMLALPRALQTSRALVVIAILLGFFGILFGVIGYSCSNCRMDQRKRSKVTIASG

VIVIIAGVLLLIPVSWTAHIIIHEFYNPILISAQRRELGSSLYIGWGSAGLLFLGGGLLC

SSRPRMNDTDCEVRYSNSRAV

>tr|I3L063|I3L063_ORENI_Oreochromis

LIISVLELVGVLISAGAWLCSLATTLMSSWLTLSNLLVTETLGVGLWETCVLNQQGTLEC

RPYDSLLGLRPEIKLARILMCTALGVGMLGILLAIPGLHLVNGCRQQLEDVSCKKALKAT

SGALCLVAGILDLIPVSYIAHVTVEQFFDESVPDMVPRWEFGDALFCGWTAGVLYLAAGI

LLLISCFYVQKLNSNRPV

>tr|I3IWY2|I3IWY2_ORENI_Oreochromis

MANSCLQLGGFLLSFLGWLGIVIATSTTDWVNICKYGLNTCKKMDELEARGPWAHCVIST

GLYHCVSLTQILDLPAYIQTTRALMITGSILGFPAVVMLLMSMPCISLGNEPQSSKNKRA

ILGGVLIIIAAICGLVSTIWFPIGAHREHGLMSFGFSLYSGWVGTIFSLLGGSIVTCSSD

SSTSHSYQENNHFYYSKQGGSNLPAASSTNHAKSAHV

***Mudskipper (Boleophthalmus pectinirostris)***

>XP_020793507.1 claudin-15_Boleophthalmus

MDPIVEVVAVILGFIGWVMVGVALPNRYWRTSSVDGNVITTSTIYENLWMSCATDSTGVHNCREFPSLLALSGYIQASRALMITSIVLGSFGLVAALIGIQCSKAGGENYVLKGRIAGTAGVLFLLQGLCTMVSVSWYAFNITQDFFNPLYVGTKYEIGEGLYIGWCSAVLAIAGGACLTCSCKLGLDEKRPLPYHSRGTVYSGAARSAAASTYGRNAYV

>XP_020778788.1 claudin-15-like_Boleophthalmus

MNAIVEVFAFFLGFLGWLMVGIALSNRNWKVSTVEGNVITTSTIYENLWMSCATDSTGVHNCRDFPSLLALNGYIQASRALMIASIVFGTFGLVATLVGMQCSKIGGENYVLKGRIAAIGGVFFLLQGICTMIAVSWYAANITQQFFDQFYPGTKYEIGEALYIGWSSATLAICGGACLLFACKGKTTDEKVPYPYQPSSRGHVFSVTAPSQSVPSNYGRNAYV

>XP_020772853.1 claudin-10-like_Boleophthalmus

MGSMGTEIVAFLLTISGWILVSSTLPTDYWKVSSVDGTVITTATFWSNLWKTCVTDSTGVSNCKDFPSMLALDAYIQVCRGLMIAAVCLGFFGAILALVGMKCTKIGGSDTIKARLTVLSGFHFILCGLCCMTACSIYARRITTDFFDPLFVAQKFELGAALFIGWSGSVLCIIGGFVFFLSLSDGFSLRQHSYRGAASFETSRHYPSKAAGSSQHKDTHEQRQFGRNAYV

>XP_020792553.1 claudin-19_Boleophthalmus

MANSGLQLLGYFLALGGWIGVISTTALPQWKQSSYAGDAIITAVGLYEGLWMSCASQSTGQVQCKIFDSMLSLDIHIQTCRALMVISVLLGFMGIIISVVGMKCTKVGDNNPTTKTRIAVTGGALFLLAGLCTLVSVSWYATQVSYQFFNPNTPANARYEFGSALFVGWAAASLTILGGSLLCCSCSKEEMRGQQYYRQSQPSTTRETNVKTTPPEKREQYL

>XP_020772846.1 claudin-10-like_Boleophthalmus

MKYRTVVMYVEIGCFVSCLCGWILVCSTLPTEYWTFSEVGSIVLTTANYYSNLWRDCVSDTTGVSDCKEYPSMLALPVFLHACRALAVCAVITGFFGGVLTLIGMKCTKIGGSEIVNARVTFAGALTYLASGFCGMITYSWWAHKVISEFVDPNFRAQKFELGAAVFVGWGGSILLICGGIVQGYFSGKESVPSSSTDRRRVYGTYVTSRSRRTYMAPASRVTLVPPLFYEGRKSRTRGTRSAKTINTYNRDTFV

>XP_020786823.1 claudin-15-like_Boleophthalmus

MSTAIEATGFAMCLISWLITGAALTNDYWKISTVSGSVIISQRQFENLWHACAENSAGIAECRDFESLLGLPAHIQACRALMIVALLFGLGGMIVALLGVKCIKIGSATDQSKAKMAATGGIVSALGGLCTLIAASWYAYRVVQDFYDPFTGGIKFEMGTGLYMAWAGAALDLLGGALLCSACKRASPPKKGGYIANQPKTVYTATAKSDPDSGRAYV

>XP_020794856.1 claudin-10-like_Boleophthalmus

MSGLQILAFLSGLAGLGASFAATVSNEWRATSRASSVITATWVLQGLWNNCAGNAIGALHCRPHHTILKLEGYIQACRGLMIAAVCLGFFGSIFALVGMKCTKIGGSDQNKARIACLAGVSFILSGLCSLSACSLYAHRITSEFFDPMFVAQKYELGAALFIGWAGSVLCILGGCMFCFSIAGSFSKSHSQRSYIYKGTVSHVTSFPKGQAPGYTRPAPKYSTSSRNQAFDKNEYV

>XP_020778488.1 claudin-3-like_Boleophthalmus

MVSLALEIVGVTLSTLGWILSIVCCSLPMWRVSAFIGANIVTAQVYWEGLWMSCVFQSTGQMQCKVYDSMLALPQDLQAARALSVISIIIGVLALLIATVGAKCTNCIPDEGVKARVMAASGGAFITASLALLVPVSWSANTIVVEFYSPIIPNGQKMEIGVALYLGWAAAALLMIGGCILCCSCPPQEEKGLKYSLPPQSRVAYSTAPRSVAQSSYNRKDYV

>XP_020785871.1 claudin-7-B-like_Boleophthalmus

MANSGLQILGFSLALLGLIGLIIGTILPQWKMSAYVGDNIITAVAMYEGLWMSCAFQSTGQIQCKVYDSILQLNSALQATRALMIVSIIVTVAGLGVACMGMKCTTCGGDDKTKKSKIAMTGGIVLVIGSLCSIVACSWYAHDIIQAFYNPFTPVNTKYEFGSAIFIAWAGAFLVIIGGGMLAASCPRGKSASAPRYPRAPSSSKEYV

>XP_020784320.1 claudin-4-like_Boleophthalmus

MPSAGLEILGVTLAVLGWISAIVSCALPMWRVSAFIGVNIVTAQITWEGIWMNCVVQSTGQMQCKVHDSMLALSGDLQAARALTVISIVLGIVGVMVSIMGAKCTNCVDQEVVKARVMIAAGACFIIASLMELIPVSWSAHTIIMEFYNPVTPEAHKREIGAALYLGWAATAFLLIGGIILCSSCPPQTEKRYRPPSKVVYSQTRSVAPSYPERRDYV

>XP_020796819.1 claudin-7-A-like_Boleophthalmus

MANSGIQLLGFFMSLVGIVGLIIGTILPQWKMSAYIGDNIITAVAMYQGLWMSCAFQSTGQLQCKIYDSILQLDSSLQATRALMIVGIIVSVAGLGVSCMGMKCTTCGGSDKIRKSRVAMTGGIILLVGGMCCIVACSWFAHNVIRAFYNPYTPVNTKFEFGVAIFVAWGGSLLDVLGGTMLASSCPRKKQVSKYPAPVTARSGPPSSAKEYV

>XP_020784215.1 claudin-4-like_Boleophthalmus

MVSQGIQMIGICMSVIGWLMVVVVCALPMWKVTAFIGANIITAQTIWQGMWMNCVVQSTGQMQCKVYDSMLALPQDLQAARAMVIISILAGIVGVILGIAGAKCTNCIEEERAKAKTCILSGVLFIVSGLLCLIPVSWTANTIVHNFYNPMLLETQRYELGASLYIGWAAAALLLMGGGLMCWNCPPKEQYPNYGPKFTPVKSASSQREYV

>XP_020772920.1 claudin-1-like_Boleophthalmus

MANAGLQLLGFTLAFLGFIGTIASTVMVEWKASSYAGDNIITAQALYEGLWKTCASQSTGQIQCKVYDSLLQLPGIVQGTRGLMLASVLFSFIAMMVAVVGMRCTTCMADSPQQKDKVALGGGVVFIIAGLMALVGTSWYGHRVAQDFYNPFTPTNAKYEFGSALYLGWGAACLSLIGGAFLCCNCSGQGSSGKTPRYPPTSSGQGGRDYV

>XP_020797042.1 claudin-5 isoform X1_Boleophthalmus

MVSAGLEILGLSLCVVGSLLVMVACGLPMWKVTAFIEANIVVAQTIWDGLWMSCVVQSTGQMQCKVHDSVLALSHDLQAARALTIISSVMGVLGLMVVIAGAQCTNCIRTEYVKARVVNAGGVIYIISGLFVLVPLCWMANNIISDFYNPQVPASKKREIGAALYIGWAATALLLIGGALLCCSCPSGGNSGYSAKYAPTKRTTQNGEYDKRNYV

>XP_020778519.1 claudin-4-like_Boleophthalmus

MVSTGLQMLGAALGIIGWIGAIIVCALPMWRVTAFIGSNIVTSQTIWEGIWMNCVVQSTGQMQCKVYDSMLALSSDLQAARALTIISIVVGILGILLAVAGGQCTNCVEDKQSKAKVGIAAGVVFIAAGVLCLIPVCWTAHTIVQDFYNPMLVSAQKRELGAALYIGWGAAALMLIGGGLLCCGCPEKEEGSYAARYKAARSDASAPVSSKDYV

>XP_020794860.1 claudin-10-like_Boleophthalmus

MKYRTVVMYMEIGCFVICVAGWILVCSTMPTEIWTWSEVDSIVLTTSNYFSNLWKDCISDSTGVSDCKGIPSMLALNWDIHMCRALIIISIILAFFGSILVLVGMKCTKIGGSELANARVSFAGGMNFLISGMCSMVAFSFYGNKIRSEFQDPTFRAQRFEIGVGVFVGWGGSTLLVVGGLLISIFLGKEGCRSSSERYPVFPLPDAYTVVPTRKSVVSSARSVVTQSRKSRVSSSDSRASSISGISSVTKTTATNSYV

>XP_020795461.1 claudin-19-like, partial_Boleophthalmus

MALVLQIVGLVLGLLSWGLTSSSTSNHHWKVKSQVESVSLSQWTFEGLWMSCAAAATGSTQCSRFKTVLGLPVHVQVCRGLMILALVLDLVSALVSALGLKCTRLSGISDRTKTRLSLTGGALFILSGLLSLTAVSLYASRVIQDFYNPVYPGV

>XP_020796789.1 claudin-14-like_Boleophthalmus

MASMALELMGFFLGLLGTVGTMVATVLPYWQISAHIGPNIVTAVANMRGLWMECVYQSTGAFQCETYNSMLALPADLQASRALMVIALVLSVLAIAISSVGMQCTVCLDGSGAVKSRVAGTGGGLFLASGFLALIPVAWTTHEVVQTFYQHNVPQSMKFEIGECLYVGLASALVTMLGGGLLAASCCEEQNGRGRRNGPGYPYPVGGAMTGTGTRMTSQTYRNPTSQPGGLNPSGRGQTLQALVRSASESSQSTNQNAPRNKKNTGYDITGYV

>XP_020778510.1 claudin-3-like_Boleophthalmus

MSMGMEIVGIALGAIGFIMAIVTCALPMWRVTAFIGSNIVTAQTIWEGLWMNCVVQSTGQMQCKVYDSLLALPQELQASRAMTIISIILGILGILISIMGAKCTNCIEDEASKAKVMIISGIFFILGGILVLIPVSWTASVTIRDFYNPVVISAQKRELGPSLYIGWGAAALLLIGGAMLCTSCPPKEQKYKPPRIAYSAPRSTGGYDRKDYV

>XP_020796873.1 claudin-8-like_Boleophthalmus

MANSALEIVGLLLTLIGLVGTAASTGMPQWRVTAFIAENIIVFETRYEGLWMNCFRQADIRMQCKVYDSLLALPPDLQAARGLMCCALALGGLGILVSLVGLQCTSCIQNNDRAKRMVLIIAGIMILCACICVLIPVSWTGHVIIRDFYNPLLLDAQRRELGEALYIGWVSSAFLFAGGCMFTCCNVKSEKEGSERYLYSRNSDYITYPPQQLQPQQLVFLPQPQPQLQPRDQPMLSRHPSANYSHYSLQSHHSNHQPARYPSVRSAVAYL

>XP_020778469.1 claudin-4-like_Boleophthalmus

MASVGIQMLASVLCLVGWTGIIFACVLPMWRVTAFVGSTIVTSQTVWEGIWMNCVVQSTGQLQCKPYDSMLALSTDLQAARALMVLAIITGGAGLILAFFGGKCTKFLDEEGPETKGKVAIAAGAVLIATGLLCLIPTSWAAGAVVKKFYSAAIDAQRRELGACLYIGWGSAILLMLGGGLFISTVCPLKSHESDKNPSVRYLVVRSSNGGSQAGSHYRLPSVQNHPVGAMSRAQSYEQPMNRSNVMYTRPPVEDDQPIAGESERSWAPSTKSQMKRPGSVKSDVSEASTTKSQLKRAELEEVEELAQSDNEDVSSHPAKTYL

>XP_020778485.1 claudin-4-like_Boleophthalmus

MPSLGLQILGVGLAVLGWIGNILICMLPMWKVSAFIGNNIVVAQTIWEGLWMTCVVQSTGQMQCKVYDSLLALPPDLQAARAMVVIAILFSLFGILLSVVGGKCTTCIGDKGAKARVAISAGVFFFLSGALCLVTVSLPANTIIKDFYNPLVPDAQRRELGACLYVGWGASGLLLIGGALLGCQCPTREDRYNGPKYSPKSTSPAKEFV

>XP_020796849.1 claudin-8-like_Boleophthalmus

MASVYTGYSGYTGYTKPPQTYADGSYYYDEKRPQTEYYPDTIYEEKKRLDMKKRRSAICCEVVALVIGFIGLIGVAAVTGLPMWKVTAFIEENIIVMETRWEGLWMNCYRQANIRMQCKVYDSLLFLPPDLQAARGLMCASVAVTTIALIVSAVGMKCTKTIDHHARTKHVVLVVGGALFLLGCVTTLIPVSWTGHVIIRDFYNPLLIDAQRRELGEALYIGWVTSALLFTAGVILLCRHAPRTQEAEALKYQGQPPAYTYQPYTYQPGYQPGYQPAYSYQPAYSAPPPGSVAYTPTQY

>XP_020776079.1 claudin-14-like_Boleophthalmus

MANMAVQILGFFLGMLGFVGTVVSTVLPHWRSTAYVGSNIITATSYMKGLWMECVWHSTGIYQCEVYRSMLALPRDLQAARALMVLSCVTSVLAGVVSVLGMKCTYFAHRSIIKSPLALSGGVCFICAGILCLITVIWTTNDVVMEFYDPFLPAGFKYEIGLCVYLGYAASCLSLVGGVVICWSSSRRSGSRRPEFHTQRSQPMSPPPAFNNIYPPAPPYNPPEALKDNRAPSVTSLSSNGYRLNNYV

>XP_020778480.1 claudin-like_Boleophthalmus

MASAGLQILAVFLATVGFIGDIIICALPMWKVTAFIGNNIVTAQTFWEGLWMNCVKQSTGQMQCKVYDSMLALPQDLQAARALVVISILVVLLGMLLSVAGGKCTNCVEDEAAKSKVVITAGVFFIVGGILCLIPVSWSANEIIRNFYNPVMIDAQRRELGASLFIGWGSAGLLLIGGSLLCCQCNKKKDSRYSVKYSAPRSTASAGAYV

>XP_020778465.1 claudin-4-like_Boleophthalmus

MVSTGRQMLGLVLAIIGFLGSIIICALPTWKVTAFIGANIVTSQVIWEGLWMNCVTQSTGQMQCKVYDSLLALPQDLQAARAMVVIAIIIGVFGILLGVVGGKCTNFVEDERQKCKVAIASGIVFIMAGVMVLIPVCWTANTIIRDFYNPVMTNAQRRELGASLYIGWGAAGLLFLGGGLLCSSCPPSEEYPVKYSKAGSMASSKAYV

>XP_020784179.1 claudin-4-like_Boleophthalmus

MVSMGRQMLGFALAIIGFLGTIIVCALPMWKVTAFIGANIVTAQVIWEGLWMNCVMQSTGQMQCKIYDSLLALPQDLQAARALIVIAIIVAAFGVLLGIAGGKCTNFVEDERAKARVAIAAGIVFIVAGVLVLIPVCWSANTIIRDFYNPIMTNAQRRELGAALYIGWGTAALLLLGGALLCSSCPHRNDAPEYPVKYSGARSTDSRAYV

>XP_020784216.1 claudin-4-like_Boleophthalmus

MVAAGFQIMGIALAIIGWIGVIIVCAIPQWRVTAFVGQNIVTAQTTWEGIWMTCVVQSTGQMQCKVYDSMLALGQDLQAARALIIISILVGIFGIFLAIAGGKCTNCVEDEAAKAKVCVAAGVIFIISGILCLVPVCWTANTVVREFYNPMLMNSQKKELGAALFIGWGASALLLIGGGILCANCPPKDNYNSAKYTASRATAPKDYV

>XP_020787613.1 claudin-5-like_Boleophthalmus

MLAACLEFLGLALCVAGSLLVMIACGLPMWKVTAFIDSNIVVAQTIWDGLWMSCVVQSTGQMQCKVHDSVLALTQDLQTARALTVISAVLGVIGLTVTIAGAQCTNCIKDEMVKAKVVNAGGIIYLISGLFVLVPLCWMANIIIVDFHNPQIPPSKKREIGASIYIGWAAAALLLLGGTLLCCSFSQGVRGSFPIKYTPTKPISANGDFDKKHYV

>XP_020796858.1 claudin-8-like_Boleophthalmus

MRAKLEIVALVLGSLGIIGIIAITAMPQWRVSAFIGANLIVMEDRWEGLWMNCYKQIDRMQCKVYDSLLILPPELQAARGLVCAAIVIALIAFAITISGTEVTSCGNDSPSAKTILLVVGGVFFLLACLTTLVPVCWVAHTVIKDFYNPTLMDAQRRELGPALYVGWGTAGLLLAAGVILLVRFCQRREMEEKEGVYPGAYDLAPKADSVYLQRIPSSTHKTMEYV

>XP_020778518.1 claudin-4-like_Boleophthalmus

MQTQIVGIILSIIGFLGTILICALPMWKMSAFIGSNIVTAQVFWEGIWMNCVIQSTGHAQCKAYDSILALPESLQAARALICASIGVSVVAIGLTVVGARCTNFYRHDRRMKLNIGLSGGIVFIVAGALCIIPVSWSAHNIITGFYNPMTIEGRKGELGACIYVGWASGALLVIGGGVLCSTYGC

>XP_020784214.1 claudin-4-like_Boleophthalmus

MSSTGLEILGMILAVAGWLGVMVACGMPMWRVAAYIGQNIVISQVIWEGLWMNCSVQSTGQMHCRVHDSMLGLPVDLQASRALVIISMVLCIIGIGLSVAGAKCTNCSQDVHGKPRFMVAAGVTFIVAGLLMLVAVSWTAHAIVMGFYDPMLEETGKREFGNALYFGWAASCLLILGGALLCCSCPPRAARAQSGSVPPRVDYSALKTVTVNGYTKRDYV

>XP_020772860.1 claudin-10-like_Boleophthalmus

MKIRVVQIWGFLMTVLGWIFVACTMAMEGWKITSIGGMAGSSIIKVAWYWSSLWRSCFTDSTAVSNCYDFPVLWSVEGHIQIVRGLLMAALSLGMLGFVLSMLGMECTYLGGQDLSKYKKVYAGGWCHIISGVLSTSGYAVYAQYVSVEYFNPDFQDLKYDLGTPLFLGWVGSAFHMAGGFFYLWSVCKPLCGGGEPVVVAVRGLPDVEKNTTTNLNKSTTVLSTQSDLTSQSKLSSISELSSKQSDVSEISSRSARTSKTGRTSRTSRTPRSPRAAGSQASTSRLGSESGRSSRSSLSTISGSSRSSRSSRSSRSSRSSGSSGSSGSSRTVSTISGISKSESAQFVKNSYI

>XP_020784149.1 claudin-4-like_Boleophthalmus

MGRVGKEVAGQVLSFIGLVGVSVTCGVPMWRVTSYIGANIVTGQIVWDGLWMNCVMQSTGQMQCKLNESLMKMSRDLQAARALIIISLVVGFIGFMISFLGAKCTSCLKKEASMANVVIISGCLIILAAVLDLIPVCWSAAYTITDYQNPLTIETQKREIGASIYIGWASAAFLLIGGIILTTSCPPQRSFYGYPGYPPQPGVYPYTAPPNQGGYRPVYTPVSQPYSGSYVPSKPYAAPSAYSGGQYI

>XP_020784148.1 claudin-4-like_Boleophthalmus

MGGMQELGISVSMAGVAGIILICALPMWKVTAFIGTHLVVMQVFWEGLWMTCVSEYTGQLQCKLYDAILDLSPDLQAARGLICISLFLGCLGFLIFLLGARCTNCLSHPRAKQRVVISSGFIFCLSSLTTVVAVAWTANSIINDFYNPRVPEVLKKELGAAIYCGFVTSGLLFVGGLILCITCSPQRAKFPSTRYTLSRAPTTQSSYAIKNYV

>XP_020786984.1 claudin-9-like_Boleophthalmus

MASTGLQLVGIVLSVLGWVFGALVCAAPLWRVSAFVGGELVIAQVLWEGLWMNCLSQTTGQIQCKAYDSTLALPFSAQAARGLTVLSLLLCVLALMLAVAGAKCTHCMGATNPSSKTRVARVAGVLFLLAGVIYIVPICWTAYGIIRDFYDPNIAAPLKRELGPALYLGWGASVLLIVGGFLLQHGSSPPISRVKPIITAPVKDNPKAPEVKQPEKSFV

>XP_020788752.1 claudin-4-like_Boleophthalmus

MGSVGVQIVCVALGVLGLIGVIVCCAIPRWNMSSFVGSNIVTAQTTYKGLWMECVMQSTGQQQCKSYDSLLVLPSDLQAARAMTIISAMVSTLSLLILFCGADFTTCIENEDAKPKISLVAGIGMIIAGLLVIIPASWSAHTVVREFNNPMITASAKMELGPCIFIGWGAGVLLLIAGGLLCCFSRPSSSSGGTAKYYSNSSPSAPKNYV

>XP_020796827.1 claudin-8-like_Boleophthalmus

MIQGVTEIAAVAVGLLGLIGAAATTGMPMWKVTAFIGENIIVMETRWEGLWMNCYRQANIRMQCKVYDSLLFLPPELQAARGLMCCSLALSGLGLLVACAGLRCITCFQANERVKTLILMTAGAMQLAASVCVFIPVSWTGHVIIRDFYNPLLIDAQRRELGEALYIGWVTGAVLFASAMLFLCRKFPSEKGSFDIYHPVNLLRYKQQPNHVPMMRYDPLSTVSSLSAPPNSASTRTSTHRSPINAPIVYNPGLPDTAVLQYQANLAHQSSMRSSSHPHSMYTPGNSLYASHNTVPLTYTTQTGNPSTSYQSSFYPVPQTPVFIGYNTSIVHRPSVTSSGMYI

>XP_020794871.1 claudin-10-like_Boleophthalmus

MRKRLIQIFGFLVTSLGWFFVLCTMAMDYWRISQIGGQGGSYIIKVAWYWSNLWKDCFTDSTAITNCRDYPVLWSVATYVQGVRGLLMCGLTIAFFAVVLCFVGMECTYIGGSEAVKDKLVLVGAVFHFVGGVSDICAYCLYINRVARTSFAASLAPGVLRYDLGPPIFLGLVGCFLIYLGAVLYAVTVYRVIFPKREVINVYGTNTYAAPRYRGKSLYTGYYKPSRQYGYYVGSGLSSSSKISMLSRTTEKISQRDAFV

>XP_020774615.1 claudin-18-like_Boleophthalmus

MAATLCQVLGFVLSLLGVALVVAATAMDQWATEDLFDNVVTAVYSYSGLWRSCVRQSSGFTECRPYFTILGLPALLQAVRALMIVGLVLGAIGCLVSIFALKCLKMGNMEGNIKATMTLTAGAMHILAGICAIAGVSAFANLIVQSFRFTTYTDGAYGIMGGANVGGITGALTPRYTFGPALFVGWIGGAILLIGGVMMCMACRGMMSNRNENYNGIAYKASSQHTQMYRQDHRSRPAYDSYKVPSEGRHSNQRFDYV

>XP_020780034.1 claudin-11-like_Boleophthalmus

MANSCLQLTGFVLSFVGWLGIVIATATNDWVINCKYGLNTCKKMDELGAKGPWADCVISTGINHCVFLTQLLDLPAYIQTTRALMITGSILGLPAVGIILMSLPCIKLGNEPQSSKNQRAIIGGVLILIVGACGLISTIWFPIGAYQEQALMAFGFSLYAGWFGTIFVLIGGFMLACCSSESHPRAYQDNNRFYYSKNGSSTVPAPTSANHAKSAHV

>XP_020784254.1 claudin-4-like_Boleophthalmus

MLRQLELGALALAFLGWICAILTRCLALWNVSGTVDNTTASLPAYWDGVWLQWDHWDLAHDGSLHCSFYQTLMSLSGNFRTWXXXXXXXXXAGGFAVVISAVGAVWFPHRNQIKVFSGAVFVLSGILLLVPTAWTCHHTSQSLEGATHLRRDWGPALYLGWISFVLMTVGGVFLTTRCPTTQSQGEHTEVATAPPHEEESNHPLSRINRATFTNSQLQRGAGPV

***Homo sapiens***

>sp|O95832|CLD1_HUMAN Claudin-1_Homo

MANAGLQLLGFILAFLGWIGAIVSTALPQWRIYSYAGDNIVTAQAMYEGLWMSCVSQSTG

QIQCKVFDSLLNLSSTLQATRALMVVGILLGVIAIFVATVGMKCMKCLEDDEVQKMRMAV

IGGAIFLLAGLAILVATAWYGNRIVQEFYDPMTPVNARYEFGQALFTGWAAASLCLLGGA

LLCCSCPRKTTSYPTPRPYPKPAPSSGKDYV

>sp|O95471|CLD7_HUMAN Claudin-7_Homo

MANSGLQLLGFSMALLGWVGLVACTAIPQWQMSSYAGDNIITAQAMYKGLWMDCVTQSTG

MMSCKMYDSVLALSAALQATRALMVVSLVLGFLAMFVATMGMKCTRCGGDDKVKKARIAM

GGGIIFIVAGLAALVACSWYGHQIVTDFYNPLIPTNIKYEFGPAIFIGWAGSALVILGGA

LLSCSCPGNESKAGYRVPRSYPKSNSSKEYV

>sp|O00501|CLD5_HUMAN Claudin-5_Homo

MGSAALEILGLVLCLVGWGGLILACGLPMWQVTAFLDHNIVTAQTTWKGLWMSCVVQSTG

HMQCKVYDSVLALSTEVQAARALTVSAVLLAFVALFVTLAGAQCTTCVAPGPAKARVALT

GGVLYLFCGLLALVPLCWFANIVVREFYDPSVPVSQKYELGAALYIGWAATALLMVGGCL

LCCGAWVCTGRPDLSFPVKYSAPRRPTATGDYDKKNYV

>sp|P56750|CLD17_HUMAN Claudin-17_Homo

MAFYPLQIAGLVLGFLGMVGTLATTLLPQWRVSAFVGSNIIVFERLWEGLWMNCIRQARV

RLQCKFYSSLLALPPALETARALMCVAVALSLIALLIGICGMKQVQCTGSNERAKAYLLG

TSGVLFILTGIFVLIPVSWTANIIIRDFYNPAIHIGQKRELGAALFLGWASAAVLFIGGG

LLCGFCCCNRKKQGYRYPVPGYRVPHTDKRRNTTMLSKTSTSYV

>sp|Q8N6F1|CLD19_HUMAN Claudin-19_Homo

MANSGLQLLGYFLALGGWVGIIASTALPQWKQSSYAGDAIITAVGLYEGLWMSCASQSTG

QVQCKLYDSLLALDGHIQSARALMVVAVLLGFVAMVLSVVGMKCTRVGDSNPIAKGRVAI

AGGALFILAGLCTLTAVSWYATLVTQEFFNPSTPVNARYEFGPALFVGWASAGLAVLGGS

FLCCTCPEPERPNSSPQPYRPGPSAAAREPVVKLPASAKGPLGV

>sp|O14493|CLD4_HUMAN Claudin-4_Homo

MASMGLQVMGIALAVLGWLAVMLCCALPMWRVTAFIGSNIVTSQTIWEGLWMNCVVQSTG

QMQCKVYDSLLALPQDLQAARALVIISIIVAALGVLLSVVGGKCTNCLEDESAKAKTMIV

AGVVFLLAGLMVIVPVSWTAHNIIQDFYNPLVASGQKREMGASLYVGWAASGLLLLGGGL

LCCNCPPRTDKPYSAKYSAARSAAASNYV

>sp|P78369|CLD10_HUMAN Claudin-10_Homo

MASTASEIIAFMVSISGWVLVSSTLPTDYWKVSTIDGTVITTATYWANLWKACVTDSTGV

SNCKDFPSMLALDGYIQACRGLMIAAVSLGFFGSIFALFGMKCTKVGGSDKAKAKIACLA

GIVFILSGLCSMTGCSLYANKITTEFFDPLFVEQKYELGAALFIGWAGASLCIIGGVIFC

FSISDNNKTPRYTYNGATSVMSSRTKYHGGEDFKTTNPSKQFDKNAYV

>sp|Q9Y5I7|CLD16_HUMAN Claudin-16_Homo

MTSRTPLLVTACLYYSYCNSRHLQQGVRKSKRPVFSHCQVPETQKTDTRHLSGARAGVCP

CCHPDGLLATMRDLLQYIACFFAFFSAGFLIVATWTDCWMVNADDSLEVSTKCRGLWWEC

VTNAFDGIRTCDEYDSILAEHPLKLVVTRALMITADILAGFGFLTLLLGLDCVKFLPDEP

YIKVRICFVAGATLLIAGTPGIIGSVWYAVDVYVERSTLVLHNIFLGIQYKFGWSCWLGM

AGSLGCFLAGAVLTCCLYLFKDVGPERNYPYSLRKAYSAAGVSMAKSYSAPRTETAKMYA

VDTRV

>sp|O15551|CLD3_HUMAN Claudin-3_Homo

MSMGLEITGTALAVLGWLGTIVCCALPMWRVSAFIGSNIITSQNIWEGLWMNCVVQSTGQ

MQCKVYDSLLALPQDLQAARALIVVAILLAAFGLLVALVGAQCTNCVQDDTAKAKITIVA

GVLFLLAALLTLVPVSWSANTIIRDFYNPVVPEAQKREMGAGLYVGWAAAALQLLGGALL

CCSCPPREKKYTATKVVYSAPRSTGPGASLGTGYDRKDYV

>sp|O95484|CLD9_HUMAN Claudin-9_Homo

MASTGLELLGMTLAVLGWLGTLVSCALPLWKVTAFIGNSIVVAQVVWEGLWMSCVVQSTG

QMQCKVYDSLLALPQDLQAARALCVIALLLALLGLLVAITGAQCTTCVEDEGAKARIVLT

AGVILLLAGILVLIPVCWTAHAIIQDFYNPLVAEALKRELGASLYLGWAAAALLMLGGGL

LCCTCPPPQVERPRGPRLGYSIPSRSGASGLDKRDYV

>sp|P56746|CLD15_HUMAN Claudin-15_Homo

MSMAVETFGFFMATVGLLMLGVTLPNSYWRVSTVHGNVITTNTIFENLWFSCATDSLGVY

NCWEFPSMLALSGYIQACRALMITAILLGFLGLLLGIAGLRCTNIGGLELSRKAKLAATA

GALHILAGICGMVAISWYAFNITRDFFDPLYPGTKYELGPALYLGWSASLISILGGLCLC

SACCCGSDEDPAASARRPYQAPVSVMPVATSDQEGDSSFGKYGRNAYV

>sp|P57739|CLD2_HUMAN Claudin-2_Homo

MASLGLQLVGYILGLLGLLGTLVAMLLPSWKTSSYVGASIVTAVGFSKGLWMECATHSTG

ITQCDIYSTLLGLPADIQAAQAMMVTSSAISSLACIISVVGMRCTVFCQESRAKDRVAVA

GGVFFILGGLLGFIPVAWNLHGILRDFYSPLVPDSMKFEIGEALYLGIISSLFSLIAGII

LCFSCSSQRNRSNYYDAYQAQPLATRSSPRPGQPPKVKSEFNSYSLTGYV

>sp|P56747|CLD6_HUMAN Claudin-6_Homo

MASAGMQILGVVLTLLGWVNGLVSCALPMWKVTAFIGNSIVVAQVVWEGLWMSCVVQSTG

QMQCKVYDSLLALPQDLQAARALCVIALLVALFGLLVYLAGAKCTTCVEEKDSKARLVLT

SGIVFVISGVLTLIPVCWTAHAIIRDFYNPLVAEAQKRELGASLYLGWAASGLLLLGGGL

LCCTCPSGGSQGPSHYMARYSTSAPAISRGPSEYPTKNYV

>sp|O95500|CLD14_HUMAN Claudin-14_Homo

MASTAVQLLGFLLSFLGMVGTLITTILPHWRRTAHVGTNILTAVSYLKGLWMECVWHSTG

IYQCQIYRSLLALPQDLQAARALMVISCLLSGIACACAVIGMKCTRCAKGTPAKTTFAIL

GGTLFILAGLLCMVAVSWTTNDVVQNFYNPLLPSGMKFEIGQALYLGFISSSLSLIGGTL

LCLSCQDEAPYRPYQAPPRATTTTANTAPAYQPPAAYKDNRAPSVTSATHSGYRLNDYV

>sp|O75508|CLD11_HUMAN Claudin-11_Homo

MVATCLQVVGFVTSFVGWIGVIVTTSTNDWVVTCGYTIPTCRKLDELGSKGLWADCVMAT

GLYHCKPLVDILILPGYVQACRALMIAASVLGLPAILLLLTVLPCIRMGQEPGVAKYRRA

QLAGVLLILLALCALVATIWFPVCAHRETTIVSFGYSLYAGWIGAVLCLVGGCVILCCAG

DAQAFGENRFYYTAGSSSPTHAKSAHV

>sp|P56748|CLD8_HUMAN Claudin-8_Homo

MATHALEIAGLFLGGVGMVGTVAVTVMPQWRVSAFIENNIVVFENFWEGLWMNCVRQANI

RMQCKIYDSLLALSPDLQAARGLMCAASVMSFLAFMMAILGMKCTRCTGDNEKVKAHILL

TAGIIFIITGMVVLIPVSWVANAIIRDFYNSIVNVAQKRELGEALYLGWTTALVLIVGGA

LFCCVFCCNEKSSSYRYSIPSHRTTQKSYHTGKKSPSVYSRSQYV

>sp|P56856|CLD18_HUMAN Claudin-18_Homo

MSTTTCQVVAFLLSILGLAGCIAATGMDMWSTQDLYDNPVTSVFQYEGLWRSCVRQSSGF

TECRPYFTILGLPAMLQAVRALMIVGIVLGAIGLLVSIFALKCIRIGSMEDSAKANMTLT

SGIMFIVSGLCAIAGVSVFANMLVTNFWMSTANMYTGMGGMVQTVQTRYTFGAALFVGWV

AGGLTLIGGVMMCIACRGLAPEETNYKAVSYHASGHSVAYKPGGFKASTGFGSNTKNKKI

YDGGARTEDEVQSYPSKHDYV

>sp|P56749|CLD12_HUMAN Claudin-12_Homo

MGCRDVHAATVLSFLCGIASVAGLFAGTLLPNWRKLRLITFNRNEKNLTVYTGLWVKCAR

YDGSSDCLMYDTTWYSSVDQLDLRVLQFALPLSMLIAMGALLLCLIGMCNTAFRSSVPNI

KLAKCLVNSAGCHLVAGLLFFLAGTVSLSPSIWVIFYNIHLNKKFEPVFSFDYAVYVTIA

SAGGLFMTSLILFIWYCTCKSLPSPFWQPLYSHPPSMHTYSQPYSARSRLSAIEIDIPVV

SHTT

>sp|Q9NY35|CLDN1_HUMAN Claudin domain-containing protein_Homo

MDNRFATAFVIACVLSLISTIYMAASIGTDFWYEYRSPVQENSSDLNKSIWDEFISDEAD

EKTYNDALFRYNGTVGLWRRCITIPKNMHWYSPPERTESFDVVTKCVSFTLTEQFMEKFV

DPGNHNSGIDLLRTYLWRCQFLLPFVSLGLMCFGALIGLCACICRSLYPTIATGILHLLA

GLCTLGSVSCYVAGIELLHQKLELPDNVSGEFGWSFCLACVSAPLQFMASALFIWAAHTN

RKEYTLMKAYRVA

>sp|Q96B33|CLD23_HUMAN Claudin-23_Homo

MRTPVVMTLGMVLAPCGLLLNLTGTLAPGWRLVKGFLNQPVDVELYQGLWDMCREQSSRE

RECGQTDQWGYFEAQPVLVARALMVTSLAATVLGLLLASLGVRCWQDEPNFVLAGLSGVV

LFVAGLLGLIPVSWYNHFLGDRDVLPAPASPVTVQVSYSLVLGYLGSCLLLLGGFSLALS

FAPWCDERCRRRRKGPSAGPRRSSVSTIQVEWPEPDLAPAIKYYSDGQHRPPPAQHRKPK

PKPKVGFPMPRPRPKAYTNSVDVLDGEGWESQDAPSCSTHPCDSSLPCDSDL

>sp|P56880|CLD20_HUMAN Claudin-20_Homo

MASAGLQLLAFILALSGVSGVLTATLLPNWKVNVDVDSNIITAIVQLHGLWMDCTWYSTG

MFSCALKHSILSLPIHVQAARATMVLACVLSALGICTSTVGMKCTRLGGDRETKSHASFA

GGVCFMSAGISSLISTVWYTKEIIANFLDLTVPESNKHEPGGAIYIGFISAMLLFISGMI

FCTSCIKRNPEARLDPPTQQPISNTQLENNSTHNLKDYV

>sp|Q8N7P3|CLD22_HUMAN Claudin-22_Homo

MALVFRTVAQLAGVSLSLLGWVLSCLTNYLPHWKNLNLDLNEMENWTMGLWQTCVIQEEV

GMQCKDFDSFLALPAELRVSRILMFLSNGLGFLGLLVSGFGLDCLRIGESQRDLKRRLLI

LGGILSWASGVTALVPVSWVAHKTVQEFWDENVPDFVPRWEFGEALFLGWFAGLSLLLGG

CLLHCAACSSHAPLASGHYAVAQTQDHHQELETRNTNLKH

>sp|C9JDP6|CLD25_HUMAN Putative claudin-25_Homo

MAWSFRAKVQLGGLLLSLLGWVCSCVTTILPQWKTLNLELNEMETWIMGIWEVCVDREEV

ATVCKAFESFLSLPQELQVARILMVASHGLGLLGLLLCSFGSECFQFHRIRWVFKRRLGL

LGRTLEASASATTLLPVSWVAHATIQDFWDDSIPDIIPRWEFGGALYLGWAAGIFLALGG

LLLIFSACLGKEDVPFPLMAGPTVPLSCAPVEESDGSFHLMLRPRNLVI

>sp|H7C241|CLD34_HUMAN Claudin-34_Homo

MVWFCNSADCQFSVFALTTIGWILSSTSTGLVEWRIWYMKDTSLYPPGIACVGIFRVCIY

RRRTNSTTTKFCYRYSYQDTFLPFEISMAQRFLLTASIFGFFGRAFNMFALRNMSMRMFE

EDTYNSFVVSGILNIAAGVFNLIAVLQNYDAVINSQGITFLPSLQMPFKPDVQEVGTAIQ

VAGIGVLPMLLTGMFSLFYKCPPYGQVHPGISEM

**OCCLUDINS**

***Round goby (Neogobius melanostomus)***

>NEME_00018861_Neogobius

MGILEGIISIVYIVGVNPMAQGSSSMMYNPMLMMCQNLYQTSYSQMGGVGGFPIYNQYLY

HYCFVDPQEGVAMVCGFLVVIALGVSAFYAHKTRGKIWRYGKPNIYWEEPLVGGKASEGR

DVEEWVNNVEENRSVQDAPTLLVSEKGAGLLNASANSVISYPAAKGDSSSFDGDTYNNEY

SMGVKGDVSRGTEPSHGRVSSSPSEETGGRKPSANRGKRRRRNPDWEESQYETEYTTGGE

TGNELDVEEWDRMYPAITSDAQRQDYKREFDADLKDYKRMCADMDDINDQLNKLSRQLDT

LDETSAKYQAAAEEYNQLKDLKQTPEYQSKKKQCRRLRHKLFHVKRMVKNYDKSHS

>NEME_00012039_Neogobius

MAAIIFIFCLVIFILISNQRLSQSRRLYLALIIVSAILALLMLIAAIYQNPQTLGAFVNQ

YLYHYCVVEPQEAIAIVFGFLIAVALIIILVFALKTRQKIQHFGKSNILWKRVKVTDEMA

PSQDVEAWVNNVLPAPGESSISDYPEKLRSSRSQLDDKTSDYDKPPYNYTPQPVTEEAVP

LNASVPYIRNSEAESSVGRPKPRTGGRPRRTDGYDTDYASSGDELDDDDFSNEYPPIRDD

QQRADYKREFDREHQEYKDLQAELDTINKNLSEVDEELDDLEEGSPQYQDALDEYNRIKN

MKKSADYKTKKRRCKYLKPKLNHIKQMVSNYDRSG

>NEME_00012037_Neogobius

MLRDTLYNIKDLQTDFVKEVHDFVLEQFSSSQSELQRILHDVEYLQSELSPLKLRCQANA

ACVDLMVWAVTEEQGAENLCTKLSEKLQSKTSSKVIIAHMPLLICCLQGLGRLCERFPVV

AHSVTMSLRDFLVVPSPVLVKLYKYHSQYIAGGSEIKIHVTNEHSQSTLSTTSSKKSQAS

MYEQLREISIDNICRCLKAGLTMDNVIVEAFLASLSNRLYIYQENDKDAHLIPDHTIRAL

GHISVALRDTPRVMEPILQILQQKFCQPPSQLDVLIIDQLGCMVITGNQYIYQEVWNLFQ

QISVKASSMVYSTKDYKDHGYRHCSLAVINALGNIAANLQNELMVDELLVNLLELFVQLG

LEGKRASERAPDKGPALKASSSAGNLGVLIPVIAVLTRRLPPIKEAKPRLQKLFRDFWLY

SVVMGFAVEGSGLWPEEWYEGVCEIATKSPLLTFPSGEPLRSELQYNSALKNDTVTPTEL

NDLRSTIINLLDPPPDVAVLINKLDFAMSTYLLSVYRLEYMRMLRALDSDRFQVMFRYFE

DRAIQKDKSGMMQCVIAVSDKVFEVFLQMMAEKAKTKSHEEELERHAQFLLVNFNHIHKR

IRRVADKYLSGLAETFPHLLWSGRVLKTMLDILQTLSLSLSADIHKDQPYYDIPDTPYRV

TVPDTYEARESIVKDFAARCGEILKEAMKWAPSVTKSHLQCFTYWGKPPSSDCDPQLLHH

LCWSPLKMFTEHGMETAIACWEWLLAARTGIEVPFMREMAGAWQMTVELKMGLFSDAQVE

ADPLAASEESQPVPCPPDVTPHHIWIEFLVQRFEIAKYSSADQVEIFGSLLQRSLSLNVG

GNQSGLNRHVAAIGPRFRLLTLGLALLHSDVVTNATVRNVLREKIYSTAFDYFSGTPKFP

TQCDNRLREDISVMIRFYASILSDKKYLAACQLVPPDNQDASLNSLSVMNAADLRSNIDS

STRSQQSGQGWINTYPLSSGMSTASKKSGTSKKSNRGTQLHKYYTKRRTLLLALLASEIE

RLTTWYNPLSTQELAITTEQSVETSISNWRSKYISLNEKQWKDNDAILFYIPQIVQALRY

DKMGYVREYILWAAQKSQLLAHQFIWNMKTNIYLDEEGHQKDPDIGELLEQMVEEIINSL

SGPAKDFYQREFDFFNKITNVSAIIKPVPKGEERKRACLKALSDIKVQAGCYLPSNPEAI

VLDIDYKSGTPMQSAAKAPYLAKFKVKRCGVSELEREGLRCPSDSVEDGEENPDGPRKVC

WQAAIFKVGDDCRQDMLALQIIGLFKNIFQLVGLDLFVFPYRVVATAPGCGVIECIPDCK

SRDQLGRQTDFGMYDYFRNQYGDESTLAFQKARYNFIRSMAAYSLLLFLLQIKDRHNGNI

MLDSKGHLIHIDFGFMFESSPGGNLGWEPDIKLTDEMVMIMGGKMEATPFKWFMEMCVRG

YLAVRYDSSTRITSFPMSHGGGFTGRTERARQAPFYDQGPDDSLPRDVHPLMSLAQSTDP

LPPPPLPGQPPVGPQFDPSGSEDEADAAIDPAIDIKPVHRYIPDSWKNFFRASDRSTTSK

PWFMPGSSYNNSNGTTEGVPCSPPRSPVPGSYRDPYGGSGGSYNSRKELLEVQDTHESSL

GRTCHTALTYSERVEQYHQRYAYMKSWAGLLRILGCVELLLGAAVFACVCAYIHKDNEWF

NMFGYSSPGGAYGAYGAYGAGAGGAYYTGPQTPFVLVVAGLAWLVTVILLVLGMTMYYRT

ILLDSNWWPLTEFTLNLALAVLYLAAGIVYVRDTTRGGLCSYPVFNNGINGAFCRTEAGQ

TAAIIFLFVTMVVYLIGAMVCLKLWRHEAARRYRERYGQEMQAAPVQGSKALQLAPETTT

VRKLPEQVQGSSVVPIQASPKATFKKVLHGAIPSGYIPKPVIVPDYIAKYPAIRSDEERD

QYRAVFNDQYAEYKELHLDVQTTLKKFDEMDAMMRTLPQHPTSQMEVERINRILQDYQRK

KKDPTFTEKKERCEYLKNKLSHIKQKIHEYDKVMEWNDGFSED

***zebrafish (Danio rerio)***

>tr|Q4KM13|Q4KM13_DANRE_Danio

MSSKHNGSPPPYDYEENRYNVAPQPAYSYYPDDEFQHFYRWTSPPGIIKIMCVLSIIFCV

GIFVCVASTLAWDTNAGAAGFGSANNPCSSGSYGGSSFAGTGYGMGSGYGVGFGYGILGS

QNDPRQGKGFMIAMAIITFIALMVIFIMVISHQRVSQGRKFYLSVIIVSALLAFFMFIAT

IVYLVTVYPMAQTSGSVRFNQVYMMCSAFQNSQMSGTFVNQYLYHYCVVDPQEAIALVLD

FVVIAALIIIMVFAIKTRQRINNYGKDNILWRRVKEIEDQNSLQDVEDWVNNVNGAPEGL

LADYPVKFGSRNNLDDNSTSYDKPPLSESPVEILPVRNSVPISSGSEMNSSVGRPKKRRA

GRPRTADGRDYDADYASSGDELDDDDFFSEFPPIVNTQERDDYKHLFDQDHQEYKDLQAE

MDQINKRLAEVDRELDGLQEGSPQFLDAMDEYNAIQDQKRSGEYKQKKKRCKYLKAKLNH

IKKMVSDYDRRS

>tr|E7F2I5|E7F2I5_DANRE_Danio

MFEKQPYDSPPKYTPPSNGFGPPGSFQDPRSEFDPYPAPPGSYYVGEEVPQHFYKWFSPP

GIVKIMIGTVIVLCLGIFACVASTLVWDMQYGYGYGSGYPGYGSGYGYGGGYGSYGYGSG

YGSGYGYGSYYNSYPTPYSAKTSMIAVAAINFIIALGFFAASFSKTNTFRSRKFFLAVLV

VCSIMAVIQGIINIVYIVGVNPMAQSSSSSYYNPMLMMCQNLYGSSVTGMGAGFPVYNQY

LYHYCYVDPQEAVAMVCGFMVMVALAVAAFFSYRTRSKIWQYGKPNIYWDRPLMAGTEGR

DVEDWVNHVQDGQSIQEASMVFSEKMSPVLASAPSVGSIPAKTGSVFSSENHNRSFSRPS

HCTSPSREVSSSPSDQTDTIRKPSAGRGKRRRRNPELDESQYETEYTTGGETADDFDQDL

WTRLYPEITSDPQRHDYKKEFDTDLRSYKELCAEMDDINDQINKLSRELDTLDEGTSKYQ

AVAEEYNRLKDLKRMPDYQNKKHQCRKLRHKLFHIKRMVKNYDRRQ

>tr|Q08BB8|Q08BB8_DANRE_Danio

MYETQQYDSPPQYSPHYTAPSNMYTSRSFYSQRSEFGLYSSVPDAPHALGNPQHFYRWFS

PPGIVKTMEGLTALLCFIIFACVASTLVWDMPGYEGSIGGYIAGSGGYGVGSGYYGGTYG

YQSSYMTPYSAKSAMISMAALNFVVSLTFLVVSFSKMWCVRGKAFYLTVLVVDVVLAILQ

CIIDIVFVIGVNPMAQSSQSILYNPILMMCQSSLGTPSISAGVNTGGFPGAYPSFNRYLY

HYCFMDPEEAVALVCGLFVMIALIIAAYFAYKTRSKIWRHGKPNIYWEEPPILRTNRSQN

SQDWRGTQYTPTVVLSEKASPHLKAENSFTSYTEGTVSVYSEGGYKSNVYSENVKGFSPE

PLYQNQWRSSSPVEEVEVQSRSVAEKAEVFEVEDVLCETGYTTAADSATELHTYELEDTY

SEITTDEQRRQYKKQFDVTFAVYKNLRAELDDISDQMNELSQELDTLHEESTKFQAVADE

YNRLKDLKRSPEYKTKKLQCKKLRQELCRIKQLVKNYDQGYKKNMKTYVADPQDFFV

>tr|F1QFR8|F1QFR8_DANRE_Danio

MPRKSSHPPPYGSRHQHRSSHRTVSYHPDEMLHFYSWKSPPGVMKILCIIIIIMCVAMFA

CVAATLAWDYDANNMGLGGLASLGMGSSGGGYNGGSYNGGSAGGYGGYGGGMGGYGYGGG

TYMDPKSGKGFIISIAAITFIAILIIFILVVSRQSSSQSPKFYLATIIICAILAALMLIA

TIVYLVTVNPTSQTSGSMMYNQILQLCAQYQNQDQASGIFINQYLYHYCVVDPEEAIAIV

LGVLVVIGLIILLVFAVKTRGLIRKYGRDRVLWYDVKTIKDGLTSQGIGEWINNVSGDPE

VFVNDQNDKVSAAQPMVYSQKPIYLPSSASDLTSSVSGLKGKLRAYDAGESGDELDTDEY

PPIINEQERLEYKRDFDRDHMVYKRLQAELDDINQGLADADRELDRLEEGSPQFMDVMDE

YNRLKSLKKSTDYQMKKRKCKQLKSKLSLIKRRVSDYDHRQ

***three spine stickleback (Gasterosteus aculeatus)***

>tr|G3NY69|G3NY69_GASAC_Gasterosteus

MFDKQHYESPPVYSPPFSSPSNNGFGPGGSFRGPHSDYNAYPPPPGSYYIEDKPQHFYKW

LSPPGIIKAMMASVIVLCLAIFACVASTLMWDMQYGMGMGMGMGMGMGSGYGGGSYGSGY

GGYGGYGGGYGGGYGNSYGSYVSPYSAKTTMIAMAAINFVGALVFFIATFSKSDTVHSRK

FFLAVMIGSIIMGVLQGIISIVYIVGVNPMAQGTSSMMYNPMLMMCQNLYQTSYSQMGGV

GGFPIYNQYLYHYCFVDPQEGIAMACGFLVVIAFGVAAFFAHKTRGKIWRYGKPNIYWQE

PLAGGKASEGRDVEEWVNNVEESRSAEDTPTLLVSEKGAGLLNASANSVISFPPARVDSS

SYNEDTYDNKEYSDRTTSRPSEAFSHGGRTSSSPSDETGGGRKPSANRGKRRRRNPELDE

SQYETEYTTGGETGNELDGEEWENTYPDITSDAQRQDYKREFDVDLREYKRLCTEMDDIN

DQLNKLSRLLDTLDHTSAKYQGVADEYNQLKDLKQTSDYRSKKKECRRLRHKLFHIKRIV

KDYDKKH

>tr|G3N596|G3N596_GASAC_Gasterosteus

SAPRGPAAMSSRPNGSPPPYESDTGYNVAPQPAYSYYPDDEFQHFYRWTSPPGVLKIMSI

ISIVMCVAIFACVASTLAWDSQGALGGFGGGNSGGAYGYGGYNTDPRTSKGFIIAMAAIT

FIACLVVFIVVVSHQNLTESRRFYLAVVIICAILALLMLIAAIVYLVAVNPSAQSSGSGY

GNPIAGLCAQYQQPQVSGVFVNQYLYHYCVVEPQEAIAVVLGFLVAVALIVMLVFALKTR

QKIRGYGKSNILWKKVKVVGEMEPAGYVEAWVNNVSTGPEVLPMSDFPEKMRSSRSHLDD

ESTNYDKPAYSYTPQPSEEHLPLQNGAPYAAASDTSGKPKKKDYETDYNSSGDELDDDDF

DSEFPPITDESRRIEYKRQFDREHLEYKELQGELDALNKGLSEADRALDELPEGSPQYLD

ALDEYDKIKAVKKSADYRVKRRRCKYLKSKLNHIKKMVSDYDRKA

>tr|G3NI06|G3NI06_GASAC_Gasterosteus

FCCSHSSRPRHSELVYNPAFSSYQGERMLHFYRWTSPPGVMKILCLIIIVLCVAVFACVA

STLAWDYDIDSIGYGGSSGGSYGGGFGGSYGGGAGGSYGGGAGGSYGGGAGGYGETELSP

AAAKGFLIAMAAITFIAALIIFVLVISRQNAARSPKFYLATIIICAILAFLMIIATIVYL

VSVNPTAQSTGSVYYSQVQQLCSQYQTQSQAQGIFLNQYLYHYCVVEPQEVIAIVLAILV

FVALIILLVFAVKTRTNIRRWGQDRILWDELTVVNNGLQNSVGDLQCQVSKVSGNPEVLL

NGANHGGRGSRDYVDQLDHSKPLYLPGDGCVLIMVLDPVCKSVMWFCSDSDISSSVGGPK

PRLRDYDTAVESGDDLEEEDFNVLFPPIVDEQERLNYKREFDLEHQEYKSLQAELEGINQ

DLADLDRELDRHPEGSAPYLDALDKHSRLKNLKKSPHYQVKKNRCKYLRSKLSHIKRRIS

NYDRRP

***tilapia (Oreochromis niloticus)***

>tr|I3JV07|I3JV07_ORENI_Oreochromis

MFDKQHYESPPVYSPPFTPPPNNGFGPAASFHGPQSDYNMYPPPPGSYYMEDKPQHFYKW

LSPPGIVKAMMVTVIVLCLAIFACVASTLMWDIQYGYGSGMGYYGSGMGYGSGYIGGSYG

SGYGGYGGYGNYYGSGSYISPYSAKAAMIAMAAINFVGALAFFIASFSKTNAVRGRKYFL

AVLISSVLMAVLQGIINIVYIVGVNPMAQSSSSMMYNPMLVMCQNLYQTSYSQMGGVGGF

PMYNQYLYHYCFVDPQEGIAMACGFLVVIALSVAAFFAQRTRGKIWRYGKPNIYWEEPLV

GGKASEGRDVEDWVNNVEESRSVQDAPTLLVSEKGGGPLNASVNSVVSYSPPKVESSSYN

DDAYDNNEYSERTNSRPSEMFFHSGGTSSSPSEETGGGRKPSANRGKRRRRNPELDESQY

DTEYTTGGETGNELDTEELESMYPEITSDDQRHNYKREFDADLREYKRLCAEMDDINDQL

NKLSRQLDTLDDTSAKYQIVAEEYNQLKDLKQTSDYQAKKRECRRLRHKLFHIKRMVKDY

DKNH

>tr|I3JW93|I3JW93_ORENI_Oreochromis

MPSNKHSNHSGNKRHRHSEVMSNPAFSYYPEDKMLHFYRWTSPPGVMKIMCIIIIIMCVA

VFACVASTLAWDYEMSLMGGGTGLMPGSYGSYGSGYGTGYGGSYGGSYGGGYGGSYGGAY

GSGLGGSYSGYGGTQMDPRSGKGFIIAISAITFIAVLIIFIMVVSRQGTARSSKFYLASI

IICAILAFLMIIASIVYLVAINPTAQSTGSVYYNQIVQLCAQFQNQVQPQGLLLNQYLYH

YCVVEPQEAIAVVMGFLVFIALIILLVFAVKTRQQIRRWGPERILWEEVKVVNVGHHNSV

GEWVTNVSGDPEVFVNDQNDKVGASRDYLDQLDYNKPLYLPGDSDISSYVGGLQPRLRDY

DTGVESGDDLEEEDFSVLLPPILDEQERLAYKQEFDRDHQEYKSLQAELDSINQDLADLD

QELDRHSDGSAEFMDALSDYNRLKDLKKSSEYQIKKKRCKYLRSKLSHIKRRISEYDRRP

***mudskipper (Boleophthalmus pectinirostris)***

>XP_020793842.1_Boleophthalmus

MSSRQNGSPPPYESENGYTAPPQPAYSYYPDDEFQHFYRWTSPPGIIKIMAIICIVLSVATFACVASTLAWDSQGALSGFGYGGYGGSYGGTYGGAYGGGYGSFGGGGAYGPGTNYGYGSIGGNYNDPRSGKGFIIAMAAIVFIYCLVIFIIIVSHQSLSESRRFYLAIVIISAFLALLMLVASIVYLVAVNPMSQSYNSAYGSQIAGLCAQYQQPQPSGVFVNQYLYHYCVVEPQEAIAIVFGFLIGVALIIILVFALKTRQKLQHFGKSNILWKRVKVVDELEPPQDVEAWVNNVSADPTELPISDYPEKLRNSRSHLDDESSNYDKPPYNYTPQPAVEEAIPLNASVPYSSNSEAESSAGRPKKRRAGRPRRTDGYDTDYGSSGDELDDNDFSTEFPPIRDDQERTDYKREFDRDHLEYKDLQAELDTINKNLSEVDRELDDLEEGSPQYLDALDEYNRIKDLKKTADYQMKKRRCKYLKAKLNHIKKMVSDYDRSG

>XP_020796584.1_Boleophthalmus

MRSREKDDSTSETSSLNESSNQKPIPVWVDHTPTPDDTNSTGGRSGTPPSSEPKSDPKSSLKSRLQTLLRLNKKTENDDSDASSSGIPILPNGTRVSPPVSPLAGRKHDIYQSDSSQNSDKNVLRYEQEESLLTSLHPAEYYAEKVEIYNLKYAYLKSWPGLLRLLAGMELLFGGMIFACVIAYIQKDSEWSNAYGMYNGAYNNGHGLSGYSYRGPMTPFVLAVAGVSWIITFVLLVVGMTMYYRTILLDSPWWPLTEAFVNVAMFLLYMAGGIVYVNDLNRGGLCFSTIGVNPIMASLCRVEGGQVAGTAFIFINMVLYLISFLVCLKMWRHEAARRERESFLNEPSEVPNSMPLVPQKPKQISFRDQTETDYSKSVKGPDSEKAPVKTFINQTGASTARKTRVKVIADYIMKYPEITSLEQRDEYRAVFNDQYQEYKDLHHHISLTLSKFRQLDALMDRLLRDKTQNPQRIQIILQKLEEKKNDPTFWEKKERCDYLKAKLSHLKNRIYTFDQMTENQKQKK

>XP_020793838.1_Boleophthalmus

MSYGGGNTGRTERARQAPYYDQVPQDSLPGDAQPMREHTSRSLARSADPLPPPPLPDQPPVGPEFDPSGSEATADSDIDPAIDIKPVHRFIPDSWKNFFRSSNRSSEKPWSMLGSSCNNNNSTTEGVPCSPPPSPIPGSYRDPYGGSGDSYNSHKELLGAQDTHESVSGRTYHTALTYSERVEQYHQRYAYMKSWAGLLRILGCVELLLGAAVFACVCAYIHKDNEWFNMFGYSSPGAPYGAYGAYGAGTGGAYYSGPKTPFVLVVAGLAWLVTVIMLVLGMTMYYRTILLDSNWWPLTEFTINLALAVLYLAAGIVYVRDTTRGGLCSYPLFNNSINGAFCRTEAGQTAAIIFLFVTMVVYLIGALVCLKLWRHEAARRYRERYGQEMLTAPVQRSQELHLVPETTTVRKLPEQVQGSSVVPIQTSPKATPKKVLQGAIPSGYIPKPVIVPDYIAKYPAIRTDEERDQYRAVFNDQYAEYKELHLDVQATLKKFDEMDAMMRTLPQHPTSQMEVDRINRILQEYQRKKNDPTFTEKKERCEYLKNKLSHIKQKIQEYDKVMEWNDGYS

>XP_020789581.1_Boleophthalmus

MPSNKHSSSKLHGGRFRPSEVSNPVFPYSPDDHLLHFYRWTSPPGVMKIVSIIIIIFCVAVFACVASTLAWDYDMSLMGGMGGMVPGYGGYGQSSARGHTFYLAAIVVGAVLAFLMLIATIVYLVAVNPTAQSSGSMMYNQVLQLCAQYQNQNQAQGIFVNQYLYHYCVVEPQEAIAIVLGFLVVVGLVVLVVFAAKARGQIRRWGEHRVLWEEPKVLGSATHHNSVGEWVNNVSGDPEFLVNAPNEKVGGSRDFLDRLDPGRPLYLPGDSDVSSSILIPKPREQDESGDELEDEDFSSLFPSIADESERLWYKREFDRDHDEYRSIQNDLEQINHDLQRLDHDLSQLPHGSPQYLDSLDEFERLRSFKKSAQYQMKKQRCKHLRSKLSHLKRKISEYDHRP

>XP_020789066.1_Boleophthalmus

MFDKQHYDSPPVYTPPSNNGYGPPGSFQDPQSEYNPYPPPPGSYYIEDKPQHFYKWLSPPGIIKAMMATVIVLCVAIFACVASTLMWDMQYGYGGMGYYGSGMGYGSGYPGGSYGSGYGSGYGSYGGNYGSYYGSYISPYSAKTAMIAMAAINFIISLAFFITSFSKSPTVRSRKYFLAVLITSIIMAILQGIISIVYIVGVNPMSQGTSSMMYNPMLMMCQSMYQGGYSNMGGVGGFPMYNQYLYHYCFVDPQEGVAMVCGFLVAIALGVAAFFAQKTRGKIWRYGKPNIYWDQPLVGGKASEGRDVEEWVNNVEENRSIQDAPTLLVSEKGAGLLNASANSVLSYPAARTDIGSFDGDTFNNNEYTERLTRPSVVSQGRISSRSSDETTTRKPSAHRGKRRRRNPDWEESQYETEYTTGGETGNELDMEEWDRMYPEITSDSQRHDYKAEFDADLKEYKRMCADMDDINDQLNKLSRQLDTLDDTSDKYQAVAEEYNQLKDLKRTPEYQEKKQHSRTLRHKLFHIKRMVKNYDKSHS

***Homo sapiens***

>sp|Q16625|OCLN_HUMAN_Homo

MSSRPLESPPPYRPDEFKPNHYAPSNDIYGGEMHVRPMLSQPAYSFYPEDEILHFYKWTS

PPGVIRILSMLIIVMCIAIFACVASTLAWDRGYGTSLLGGSVGYPYGGSGFGSYGSGYGY

GYGYGYGYGGYTDPRAAKGFMLAMAAFCFIAALVIFVTSVIRSEMSRTRRYYLSVIIVSA

ILGIMVFIATIVYIMGVNPTAQSSGSLYGSQIYALCNQFYTPAATGLYVDQYLYHYCVVD

PQEAIAIVLGFMIIVAFALIIFFAVKTRRKMDRYDKSNILWDKEHIYDEQPPNVEEWVKN

VSAGTQDVPSPPSDYVERVDSPMAYSSNGKVNDKRFYPESSYKSTPVPEVVQELPLTSPV

DDFRQPRYSSGGNFETPSKRAPAKGRAGRSKRTEQDHYETDYTTGGESCDELEEDWIREY

PPITSDQQRQLYKRNFDTGLQEYKSLQSELDEINKELSRLDKELDDYREESEEYMAAADE

YNRLKQVKGSADYKSKKNHCKQLKSKLSHIKKMVGDYDRQKT

>tr|A1BQX2|A1BQX2_HUMAN_Homo

MSNDGRSRNRDRRYDEVPSDLPYQDTTIRTHPTLHDSERAVSADPLPPPPLPLQPPFGPD

FYSSDTEEPAIAPDLKPVRRFVPDSWKNFFRGKKKDPEWDKPVSDIRYISDGVECSPPAS

PARPNHRSPLNSCKDPYGGSEGTFSSRKEADAVFPRDPYGSLDRHTQTVRTYSEKVEEYN

LRYSYMKSWAGLLRILGVVELLLGAGVFACVTAYIHKDSEWYNLFGYSQPYGMGGVGGLG

SIALVCLKLWRHEAARRHREYMEQQEINEPSLSSKRKMCEMATSGDRQRDSEVNFKELRT

AKMKPELLSGHIPPGHIPKPIVMPDYVAKYPVIQTDDERERYKAVFQDQFSEYKELSAEV

QAVLRKFDELDAVMSRLPHHSESRQEHERISRIHEEFKKKKNDPTFLEKKERCDYLKNKL

SHIKQRIQEYDKVMNWDVQGYS

>tr|D6RA09|D6RA09_HUMAN_Homo

MSNDGRSRNRDRRYDEVPSDLPYQDTTIRTHPTLHDSERAVSADPLPPPPLPLQPPFGPD

FYSSDTEEPAIAPDLKPVRRFVPDSWKNFFRGKKKDPEWDKPVSDIRYISDGVECSPPAS

PARPNHRSPLNSCKDPYGGSEGTFSSRKEADAVFPRDPYGSLDRHTQTVRTYSEKVEEYN

LRYSYMKSWAGLLRILGVVELLLGAGVFACVTAYIHKDSEWYNLFGYSQPYGMGGVGGLG

SMYGGYYYTGPKTPFVLVVAGLAWITTIIILVLGMSMYYRTILLDSNWWPLTEFGINVAL

FILYMAAAIVYVNDTNRGGLCYYPLFNTPVNAVFCRVEGGQIAAMIFLFVTMIVYLISAL

VCLKLWRHEAARRHREYMEQQEINEPSLSSKRKMCEMATSGDRQRDSEVNFKELRTAKMK

PELLSGHIPPGHIPKPIVMPDYVAKYPVIQTDDERERYKAVFQDQFSEYKELSAEVQAVL

RKFDELDAVMSRLPHHSESRQEHERISRIHEEFKKK

**INOSITOL MONOPHOSPHATASES (IMPA)**

***Round goby (Neogobius melanostomus)***

>NEME_00006010_Neogobius

MADPWIEAYTFAVEIAKKAGAEIRKASESEIKISTKSSTVDLVTATDVRVEKIIIGALKE

KFGEDVHCFIGEESISKKGACILTDEPTWIIDPVDGTTNFVHGFPFVAVSIAFVVKKELE

FGVVYSCFEDKLYKGRKGQGAYCDEESIQVSDNIDINKSVIISEFGTDRTPSKVKKICST

MERILKIPVHGIRGTGTAATNMCMVACGAVEAFFEIGIHCWDVAAGAVIVQEAGGVVCDV

DGGPFDLMSRRMMCANHIDIANRVIKEIEIFPSERDDGQD

>NEME_00006008_Neogobius

MSDPWQECMDFCVEVTKKAGQMIRDALEKDITIMEKSSPVDLVTETDQKVEQLIISSIRE

KYPTHSFIGEESVAAGAPSILTDDPTWIIDPVDGTTNFVHRFPFVAVSIGFTVQKVIEFG

IVYSCVEDKMYTARKGKGAFCNGAPIKVSGQQDITKSMVLTEMNFKEDVERFNTMLANVK

TILSIPVHGIRSPGSAAVNMCLVACGSADAYYHMGIHCWDMAGGAAVVTEAGGVIMDISG

GPFDLMSRRLIVASSRAIADRLVKEIREFPVGRDDQ

***Zebrafish (Danio rerio)***

>tr|Q503S9|Q503S9_DANRE_Danio

MADWSECLDVAVDIARRAGQMVSCAVQLEKRVSSKSTPTDLVTEADHQVEELIISTLREK

YPTHRFIGEESSAAGVKCELTDSPTWIIDPIDGTCNFVHSFPMVAVSIGFAVRKELEFGV

IYHCFDGTLYTARKGHGAFCNGVRLQVSKEKDVSKALILTEIGAKRDSATLDIFLGNMKK

ILSAPTHGVRIIGSSTLSLCQVASGAAEAYYQYGLHCWDIAAAAVIIREAGGCVMDTTGG

PLDLMSRRVVAAGTREIAEYVVKQLQPINYGRDDL

>tr|Q6DGB2|Q6DGB2_DANRE_Danio

MPDLWQDAMDHAVTLARKAGEIVREALQNDLKIMCKSSSVDLVTKTDQNVEQLIITSVKE

KFPEHSFIGEESVAAGEPCVLTENPTWIVDPVDGTTNFVHGYPFVAVSIGFAVNKTLEFG

VVYSCIEDKMYTARKGKGAFCNGQPLQVSDQKEINQSIIATEFGSNRDPENVEKIFSSMR

KILCLPVHGIRGAGSAAINMCLVAAGCVEAYYEIGIHCWDMAAGAVIVSEAGGVLLDVEG

GPFDLMSRRVLAANNKTIGERIVQEVEAFPAVRDDAPVNPIK

>tr|B0S5J9|B0S5J9_DANRE_Danio

MPDLWQDAMDHAVTLARKAGEIVREALQNDLKIMCKSSSVDLVTKTDQNVEQLIITSVKE

KFPEHSFIGEESVAAGEPCVLTENPTWIVDPVDGTTNFVHGYPFVAVSIGFAVNKTLEFG

VVYSCIEDKMYTARKGKGAFCNGQPLQVSDQKEINQSIIATEFGSNRDPENVEKIFSSMR

KILCLPVHGIRGAGSAAINMCLVAAGCVEAYYEIGIHCWDMAAGAVIVSEAGGVLLDVE

***Three spine stickleback (Gasterosteus aculeatus),***

>tr|G3NMA2|G3NMA2_GASAC_Gasterosteus

MADPWQTAYDFAVQVARAAGAVIRKAGEEEIKIQTKSSAVDLVTKTDERVEKIIIGSLKE

QFGDGTHCFIGEESVSKGEACILTDKPTWIIDPVDGTTNFVHGFPFVAVSIAFAVNKQLE

FGVVYSCLEDKMYKARRGQGAFCDDEQIQVSDVKDINKSIIISEHGTDRSIEKVTKIFST

MQKILRIPVHGLRGSGTAATNMCLVASGAVEAFFEIGIHCWDIAAGAVIVQEAGGLILDV

DGGPFDLMSRRMMSANNDVIARRIIKEIELFPVVRDDAPVQKK

>tr|G3ND95|G3ND95_GASAC_Gasterosteus

MADLWQNAMDHAVALARRAGEVVREALQDDRKVMTKSCSVDLVTQTDQKVEQLIIQSVKE

KFPTHSFIGEESVAAGEACVLTDCPTWIVDPIDGTTNFVHAFPFVAVSIGFSVNRQMEFA

VVYSCLEDKMYTARRGKGAFCNGEPLQVSGQSDIKQSIISTEFGSSRDPEAVEKIFLSLR

SILCIPVHGVRGAGTAAINMCHVASGCVEAYYEVGIHVWDVAAGSLLVSEAGGVLMDVDG

GAADLMSRRIVAANSRTIAERLVSEIHSFSPLRDDATP

>tr|G3NMF7|G3NMF7_GASAC_Gasterosteus

MSDPWQECLDLCVEVTKKAGEMIREALQKDIAVMQKSSPVDLVTETDQRVEQLIISSIKE

KFPAHSFIGEESVAAGAASVLTDDPTWIIDPIDGTTNFVHRFPFVSVSIGFTVKKEIEFG

VVYSCIEDKMYTARKGKGAFCNGAPIKVSGQEDISRSLVLTEMNFDKDPARFRTMLANVE

SILTIPVHGIRSPGSAAVNMCLVACGSADAYYHMGIHCWDMAGGAAVVTEAGGVIMDISG

GPFDLMSRRLIVASSRAVAERIAKQVAELRVGRDDTDG

>tr|G3NMA6|G3NMA6_GASAC_Gasterosteus

WQTAYDFAVQVARAAGAVIRKAGEEEIKIQTKSSAVDLVTKTDERVEKIIIGSLKEQFGD

GTHCFIGEESVSKGEACILTDKPTWIIDPVDGTTNFVHGFPFVAVSIAFAVNKQLEFGVV

YSCLEDKMYKARRGQGAFCDDEQIQVSDVKVLNMNIKGAERPTVAKYLLFFFPINWLRGS

GTAATNMCLVASGAVEAFFEIGIHCWDIAAGAVIVQEAGGLILDVDG

***Tilapia (Oreochromis niloticus),***

>tr|I3JGL5|I3JGL5_ORENI_Oreochromis

MEDPWQKAYDFAVAVARKAGAEIRKAGESEIRVMTKRSTVDLVTKTDERVEKIIIGSLKE

EFGEGTHCFIGEESVAKGEPCILTDKPTWIIDPVDGTTNFVHGFPFVAVSIAFAVNKELE

FGVVYSCLEDKMYKARKGKGAFCDDEPIQVSDVKDINKSIIISEHGTDRSPEKVTKIFST

MQKMLCIPVHGLRGSGTAATNMCLVATGAVEAFFEIGIHCWDIAAGAVIVREAGGILLDV

DGGPFDLMSRRMVSANNETIAKRIIKEIEIFPVVRDDAPIEKK

>tr|I3JGM2|I3JGM2_ORENI_Oreochromis

MSDPWQECMDYCIEVTKRAGKMICEALQKDIAVMHKSSAVDLVTETDQKVEQLIISSIRE

KYPTHSFIGEESVAAGAPSALSDNPTWIIDPIDGTTNFVHRFPFVSVSIGFTVKKEIEFG

IVYSCIEDKMYTARKGKGAFCNGVPIKVSGQEDISKSLVLTEMGFKKDAEQFQTMMANIR

NILSIPVHGIRSPGSAAVNMCLVACGSADAYYHMGIHCWDMAGGAAVVREAGGVIMDISG

GPFDLMSRRLIVASSRMIAERIAKEITEFHVGRDDTDD

>tr|I3K906|I3K906_ORENI_Oreochromis

VFPLSTGSSSNMADPWQNAMDHAVAVARKAGEVVCEALRGDRAVMTKSSVVDLVTQTDQK

VEQLIIHSVKERFPTHRFIGEESVAAGEPCVLSDSPTWIIDPIDGTTNFVHAFPFVAISI

GFAINKQVEFGVVFSCVEDKMFTARRGKGAFCNGEPLQVSRQEAVEQAMVATEFGSNRDT

DVVNKIFQSLRNILCLPVHGVRGAGSAAINMCLVASGCVEAYYEIGIHVWDVAAASLIVS

EAGGVLMDVEGGDVDLMSRRIIAANSKTIAERLIKAIVPFSVPRDDAA

***Mudskipper (Boleophthalmus pectinirostris)***

>XP_020786419.1_Boleophthalmus

MADPWLEAYNFAIEIAKKAGAEIRRASESEIRICTKSSTVDLVTATDVRVEKIIIGALKEKFGDDVHCFIGEESASKKGACILTDEPTWIIDPVDGTTNFVHGFPFVAVSIAFAVNKELEFGVVYSCFEDKLYKGRKGQGAYCDEESIQVSDNKEITKSVIISEFGTDRTPSKVQKICNTMKRIIEIPVHGIRGTGTAATNMCMVARGAVEAFFEIGIHCWDVAAGAVIVQEAGGIVCDVDGGPFDLMSRRMMCANHPDIANRVIKEIEIFPSDRDDAE

>XP_020786424.1_Boleophthalmus

MSDPWQECLDYCVEVTKRAGQMIRDALQKDITIMEKSSPVDLVTETDQNVEKLIISSIREKYPSHSFIGEESVAAGAPSILTDNPTWIIDPVDGTTNFVHRFPFVAVSIGFTVKKVIEFGIVYSCVEDKMYTARRGKGAFCNGDPIKVSGQQDIKQSLVLTEINFKKDTEQFNTMMDNVKTILSIPVHGIRSPGSAAINMCLVACGAADAYYHMGIHCWDMAGGAAVLTEAGGVIMDISGGPFDLMSRRLIVASSRAIADRIVKEIKEIPVGRDDQ

>XP_020796829.1_Boleophthalmus

MSKSHDPWQDAMDHAVAVARKSGEIIREAVQNQDQSSLVQTKSSAVDLVTETDQKVEQLIIQSVKSKFPSHRFIGEESVSSGELCVLTSDPTWIIDPIDGTTNFVHGYPFVGVCIGFAVNQQVEWGVVYSAIEDKMFTARRGRGAFCNGRPLQVSPQTDVHRALVATEFGSSRDPQVVDKIFSSLRKVLSLPVHGVRGAGSAALNMCAVALGSVELYYETGIHIWDFAAASVIVTEAGGVLRDTTGGAVDLMSRRVVAANNETIATRIISELDVYSPERDDGAPHGAPHGAPHGAPE

***Homo sapiens***

>tr|E5RGY4|E5RGY4_HUMAN_Homo

MGQRPGPVLPAVAVLGQVAKRKVAWLLRWKAVTRTETAGNSSGVYGFGKMKIFVKYFQKM

ADPWQECMDYAVTLARQAGEVVCEAIKNEMNVMLKSSPVDLVTATDQKVEKMLISSIKEK

YPSHSFIGEESVAAGEKSILTDNPTWIIDPIDGTTNFVHRFPFVAVSIGFAVNKKIEFGV

VYSCVEGKMYTARKGKGAFCNGQKLQVSQQE

>tr|H0YBL1|H0YBL1_HUMAN_Homo

XWKAVTRTETAGNSSGVYGFGKMKKMADPWQECMDYAVTLARQAGEVVCEAIKNEMNVML

KSSPVDLVTATDQKVEKMLISSIKEKYPSHSFIGEESVAAGEKSILTDNPTWIIDPIDGT

TNFVHRFPFVAVSIGFAVNKKIEFGVVYSCVEGKMYTARKGKGAFCNGQKLQVSQQEDIT

KSLLVTELGSSRTPETVRMVLSNMEKLFCIPVHGIRSVGTAAVNMCLVATGGADAYYEMG

IHCWDVAGAGIIVTEAGGVLMDVTG

>tr|A0A024R830|A0A024R830_HUMAN_Homo

MADPWQECMDYAVTLARQAGEVVCEAIKNEMNVMLKSSPVDLVTATDQKVEKMLISSIKE

KYPSHSFIGEESVAAGEKSILTDNPTWIIDPIDGTTNFVHRFPFVAVSIGFAVNKKIEFG

VVYSCVEGKMYTARKGKGAFCNGQKLQVSQQEDITKSLLVTELGSSRTPETVRMVLSNME

KLFCIPVHGIRSVGTAAVNMCLVATGGADAYYEMGIHCWDVAGAGIIVTEAGGVLMDVTG

GPFDLMSRRVIAANNRILAERIAKEIQVIPLQRDDED

>tr|A0A140VJL8|A0A140VJL8_HUMAN_Homo

MGQRPGPVLPAVAVLGQVAKRKVAWLLRWKAVTRTETAGNSSGVYGFGKMKIFVKYFQKM

ADPWQECMDYAVTLARQAGEVVCEAIKNEMNVMLKSSPVDLVTATDQKVEKMLISSIKEK

YPSHSFIGEESVAAGEKSILTDNPTWIIDPIDGTTNFVHRFPFVAVSIGFAVNKKIEFGV

VYSCVEGKMYTARKGKGAFCNGQKLQVSQQEDITKSLLVTELGSSRTPETVRMVLSNMEK

LFCIPVHGIRSVGTAAVNMCLVATGGADAYYEMGIHCWDVAGAGIIVTEAGGVLMDVTGG

PFDLMSRRVIAANNRILAERIAKEIQVIPLQRDDED

**MYO-INOSITOL PHOSPHATE SYNTHASES (MIPS)**

***Round goby (Neogobius melanostomus)***

>NEME_00015537_Neogobius

MPVRIHVNSPNVRYTDTHIEAKYVYQTTSVHREGHHITVTPQSTQMTFRTERWDISSLDL

GRAMERAEVLDWALQEQLRSHMSHMKPRKSIYIPDFIAANQESRADNVLTGTIAEQMEQI

RADIRDFRGSEGVDKVIVLWTANTERFCDVTPGVNDTAENLMAAIQNTFVPGAVELAVQR

GVFIAGDDFKSGQTKIKSVLVDFLVSAGIKPTSIVSYNHLGNNDGKNLSAPQQFRSKEIS

KSNVVDDMVQSNPILYQPGEKPDHCVVIKYVPYVGDSKRAMDEYTSEIMMGGTNTLAIHN

TCEDSLLASPIILDLVILTELCQRISVRVQGEKEYQSFHSVLAILAFLCKAPLVPPGAPV

INAFFSQRACIENILRACLGLPPQNHMLLEHKLQRGFLPNNKKSKSVSGVHYDVNGEAHV

NGVTYRVSDVSGPAAVDTPTHGPGPASSEGERGKRAMADTIPTTATNLLNGYRNAATDTT

AMPVRIHVNSPNVRYTDTHIEAKYVYQTTSVHREGHHITVTPQSTQMRFRTGSRVPRLGV

MLVGWGGNNGTTVTAAVLANKLGLTWRTKTGVQNANYYGSFLQSSTLCLGSGWDISSLDL

GRAMERAEVLDWALQEQLRSHMSHMKPRKSIYIPDFIAANQESRADNVLTGTIAEQMEQI

RADIRDFRGSEGVDKVIVLWTANTERFCDVTPGVNDTAENLMAAIQYTGGLRLYQRLPTE

HLCPGAVELAVQRGVFIAGDDFKSGQTKIKSVLVDFLVSAGIKPTSIVSYNHLGNNDGKN

LSAPQQFRSKEISKSNVVDDMVQSNPILYQPGEKPDHCDSLLASPIILDLVILTELCQRI

SVRVQGEKEYQSFHSVLAILAFLCKAPLVPPGAPVINAFFSQRACIGNILRACLGLAPQN

HMLLEHKLQREFLTNGKKSKSVSGVHYDINGEAHVNAGRRSITAAARSPRKALPPPLLRH

HTSTDMADNKEKRRSAVIPSAMAESKAAKGDSDREFLAQAGVGELLRGAILKMVEARSDD

PIGFLADHFCNLASGSDAGTGGSEGDEMSNGTVGGRAHEQQRLNRALWHLRLAHHSHRLA

FSNNVRAAYDLLNEPSSSPQQQPSPADLQQAPDPDPDSPAAGSGGGVRGGLYTQTLECLC

SEGGVPASTSAPLLRRLHCQDHEAVPYEVFRHGVLTCAVFSDYIRQAQRLYAEVCCPDEG

PVSRALCMAVIETLRDALETSQGNEAGSANANANMTSCLDASVKALRYLEASAKLSPDRL

AQAMAKAQTRGPGGNMDAKEFENAAAELFIARVKVLS

***Zebrafish (Danio rerio)***

none

***Three spine stickleback (Gasterosteus aculeatus),***

>tr|G3P635|G3P635_GASAC_Gasterosteus

MSANVHINSSNVKYTDTHIVSQYSYQTTSVQRDGNKVTVTPRTTEMTFRTERRVPRLGVM

LVGWGGNNGTTVTAAVLANKLGLTWRTNTGVKKANYYGSLLQASTVCLGTGVEGEVNVPI

CDLLPMVHPNDIVFDGWDISSMDLGRAMERAQVLDWSLQEQLRPHLSQMKPRPSIYIPDF

IAANQESRADNVLTGSIAEQVEQVRADIRDFRQSSGVDKVIVLWTANTERFCDVIPGVND

KAKNLLAAIQAGAEVSPSTLFAVASILEGCAYINGSPQNTFVPGAIELAVQNGVFIGGDD

FKSGQTKVKSVLVDFLVSSGIKPTSIVSYNHLGNNDGKNLSAPQQFRSKEISKSNVVDDM

VHSNSILYQPGEKPDHCVVIKYVPRVGDSKRAMDEYTSEIMMGGTNTIAMHNTCEDSLLA

TPIMLDLVILTELCQRITVRPQGEEDFQSFHSVLALLSFLCKAPLVPSGTPVINAFFRQR

AAIENIMRACLGLPPQSHMLLEHKLQKNFLPQHTTYVNGDVAALKKASLTNGVHIPLTNG

IHTHLDDATHAL

***Tilapia (Oreochromis niloticus),***

>tr|I3IXX3|I3IXX3_ORENI_Oreochromis

MSVNVHINSPNVKYTDSHIEAQYSYQTTSVHRDGNKVTVTPRTTEMTIRTERRVPRLGVM

LVGWGGNNGTTVTAAVLANKMGLTWKTKNGVKKANYFGSLLQSSTVCLGSGLEGEVNVPF

RDLLPMVHPNDIVFDGWDISSLDLGSAMERAQVLDWSLQEQLRPYMSSLKPRPSIYIPEF

IAANQESRADNVLTGTMAEQMERIRADIRDFRQASGVDKVIVLWTANTERFCDIIPGVND

SAKNLLAAIQAGAAEVSPSTLFAVASILEGCAYINGSPQNTFVPGAIELAMQRGVFIGGD

DFKSGQTKIKSVLVDFLVSAGIKPTSIVSYNHLGNNDGKNLSAPQQFRSKEISKSNVVDD

MVQSNPILYEPGEKPDHCVVIKYVPYVGDSKRAMDEYTSEIMMGGINTIALHNTCEDSLL

ATPIILDLVMLTELCQRVTVKPQGEESFQSFHSVLSLLSFLCKAPLVPSGTPVVNAFFRQ

RSSIENIMRACLGLPPQNHMLLEHKLQRNFLPPHETCVNNDVASLKKVPLVNGNHIPLTN

GVYAHMDHTACAL

***Mudskipper (Boleophthalmus pectinirostris)***

>XP_020775377.1_Boleophthalmus

MSVKIHINSPHVSYTDTHIEADYVYQSTAVHREGDHITVTPRNTQMTFRTQSHVPRLGVMLVGWGGNNGTTVTAAVLANKLQLTWRTKTGVKKANYYGSLLQSSTVCLGSDPQGDVNIPFRDLLPMVHPNDIVFDGWDISSLDLGRAMERAEVLDWALQEQLRPIMSRMKPRKSIYNPDFIAANQESRADNVLMGTIAEQMKQIRADVKDFRSSSGVDKVIVLWTANTERFCDVTPGLNDTADNLLAAIQAGAEVSPSTLFAVASILEGCAFINGSPQNTFVPGAVELAVQRGVFIGGDDFKSGQTKIKSVLVDFLVSAGIKPTSIVSYNHLGNNDGKNLSAPQQFRSKEISKSNVVDDIVQSNHVLYKPGEKPDHCVVIKYVPYVGDSKRAMDEYMSEIMMGGTNTIALHNTCEDSLLASPIILDLVILTELCERVSVRVQGEQEEQSFHSVLALLAFLCKAPLVPPGAPVVNAFFKQRACIENIMRACLGLPPQNHMLLEHKLQKGFVLLQKKKFGETALNGEARTNGHGYIDGYSYNSEPA

***Homo sapiens***

>sp|Q9NPH2|INO1_HUMAN_Homo

MEAAAQFFVESPDVVYGPEAIEAQYEYRTTRVSREGGVLKVHPTSTRFTFRTARQVPRLG

VMLVGWGGNNGSTLTAAVLANRLRLSWPTRSGRKEANYYGSLTQAGTVSLGLDAEGQEVF

VPFSAVLPMVAPNDLVFDGWDISSLNLAEAMRRAKVLDWGLQEQLWPHMEALRPRPSVYI

PEFIAANQSARADNLIPGSRAQQLEQIRRDIRDFRSSAGLDKVIVLWTANTERFCEVIPG

LNDTAENLLRTIELGLEVSPSTLFAVASILEGCAFLNGSPQNTLVPGALELAWQHRVFVG

GDDFKSGQTKVKSVLVDFLIGSGLKTMSIVSYNHLGNNDGENLSAPLQFRSKEVSKSNVV

DDMVQSNPVLYTPGEEPDHCVVIKYVPYVGDSKRALDEYTSELMLGGTNTLVLHNTCEDS

LLAAPIMLDLALLTELCQRVSFCTDMDPEPQTFHPVLSLLSFLFKAPLVPPGSPVVNALF

RQRSCIENILRACVGLPPQNHMLLEHKMERPGPSLKRVGPVAATYPMLNKKGPVPAATNG

CTGDANGHLQEEPPMPTT

**SODIUM/INOSITOL COTRANSPORTERS (SMIT)**

***Round goby (Neogobius melanostomus)***

>augustus_masked-NEME_48-processed-gene-8.6-mRNA-1 protein AED:0.14 eAED:0.14 QI:0|0|0|1|1|1|3|0|610

MGPGMEAADIAVVALYFVLVLLVGCFAMWKANRSTVSGYFLAGRSMTWIVIGASLFVSNI

GSEHFIGLAGSGAASGFAVGAWEFNSLLLLQLLGWVFIPVYIHSGVYTMPEYLSKRYGGS

RLKVYFAFLSIMLYIFTKLSVDLYAGALFIQVSLGWNMYLSIVVLICTTAILTITGGLVA

VLYTDALQAVLMIGGALTLTILSLVKVGGLEGVRTKYMQAVPNVAGITAAGNYTYSPSCR

IHPKPDSLRLLRGPLDEDIPWPGFILGQTPASVWYWCADQVIVQRVLAAKNIAHAKGATL

MAGFLKILPMFLIVIPGMISRIMFADELVCIGPEHCMAVCGSQAGCSNIAYPRLVMAIMP

VGLRGLMMAVMMAALMSDLDSIFNSASTIFTLDIYKTFRKGASQRELLIVGRVFIVFMVA

ISIAWVPAVVEMQGGQTYLYIQEVAGYLTPPIAALFLLGRCDQPDTRPLFITRVHYMYVA

AVLFWFSGFVAVFVSLCTAPPDVEQLSVSDWSNLTASLEESLTGALASLLNNWERNVTAS

SGEERDNDTDVLCYLAVMLGLFAFIIVAILVSTVKSKRQEHSNDPYHQYIKEGWTAQIQQ

SEIISNFAAK

>NEME_ORF_194840 L:2061_GC:48.7

MATSSTMEAADIVIVALYFLLVLAIGLLAMWKANHSTVSGYFLAGRSMNW

AAVGASLFVSNIGSEHFIGLAGSGAASGFSVAAWEFNALLLMQLLGWLFI

PVYILSGVYTMPEYLAKRFGGQRLKVYFATLSLVLYIFTKLSVNLYTGAL

FIQESLGWNIYLSIILLIGMTAILTVTGGLVAVIYTDVIQAILMIGGALT

LTGISLYKVGGLEGVRTKYMLATPNVTSIMLSYPNLTYSNSCLLHLDPKP

ASLKILRGPTDPDLPWPGFLLGQTPASLWYWCADQVIVQRVLAAKNISHV

KASTIMAGFLKILPMFIIVIPGMIARILFADDLICISPEHCMEVCGSPAG

CSNIAYPRLVMSVMPIGLRGLMLAVMIAALMSDLDSIFNSASTIFTLDIY

KMLRKQSTSRELLIVGRLTVVFLVVISIAWVPVVIEMQGGQMFYYTQEVS

NYLTPPIAAMFLLAVLWHRCNETGAFWGALTGFVLGTFRLTLGFIYREPL

CDQPDNRPSFIKDVHFMYVAAILFWISALVTVVVSLLTPPPAEEQIKIAT

LWGLRNRKKLKPQQMVDLKDTSNVLKPLNPPECNGNADLNGHLPLNDTSV

TVSQNGLPKQKVDCYGDAGNSRCLQFVNWFCGIQEDPVSACVQSPQEHEK

MVEELLYESPRMKMILNLALVFICGVTIFLFVYFTL

***zebrafish (Danio rerio)***

>tr|B8JHU7|B8JHU7_DANRE Solute carrier family 5 (sodium/myo-inositol cotransporter), member 3a OS=Danio rerio OX=7955 GN=slc5a3a PE=3 SV=1

MASTMEAADIIVVAMYFVLVLCFGFFSMWRANRGTVSGYFLAGRSMNWFVIGASLFVSNI

GSEHFIGLAGSGAASGLAVGAWEFNAILLLQLLGWVFIPVYIHSGVYTMPEYLSKRFGGN

RLKIYFASLTLLLYIFTKLSVDLYSGALFIQESLGWNLYLSIILLITLTALLTVTGGLVA

VIYTDTLQAFLMISGALCLMAISLVKVGGLEGLRTKYMNALPNITFITLENNLTYSNSCH

VNPKADSFKLLRSPLDKDLPWPAFLLGQTPASIWYWCADQVIVQRVLAARNIAHAKGSTL

LAGVLKVLPMFIIVIPGMISRVLFADELACIGPEHCMQVCNSRAGCTNTAYPRLVMNIMP

VGLRGLMMAVMLAALMSDLDSIFNSASTIFTLDIYKMIRHQASSKELMVVGRLFVVLIVA

ISIAWVPFVIEMQGGQMFLYIQEVSDYLTPPIAALFLLGVLWKRCNETGAFWGGFVGLIL

GIVRLILAFVYREPDCDSVDDRPAFIKDVHFMYVAAVLFWVSGFVMVVVSLCTNPPSKEQ

ISKTTLWGLRNRNQPADTKAKGGNGKLEQTLDGVEFQPLPPEMLPNGHLEAPHTHRQATE

HAARNKGRCVKCLGWFCGYNEHSEESDSPFEDDGRSVHEMLKESAQNKTALNIGLLLVCA

LGIFLYVYFSL

>tr|E9QDC2|E9QDC2_DANRE Solute carrier family 5 (sodium/myo-inositol cotransporter), member 3b OS=Danio rerio OX=7955 GN=slc5a3b PE=3 SV=1

MEAADITVVALYFVLVLVIGLLAMWKANHSTVSGYFLAGRTMNWAVIGASLFVSNIGSEH

FIGLAGSGAASGFAVAAWEFNALLLLQLLGWVFIPVYIHSGVYTMPEYLSKRYGGKRLKI

YFAGLSLLLYIFTKLSVDLYAGALFIQESLGWNLYLSIILLISMTALLTVTGGLVAVIYT

DTLQAVLMIGGALTLTIISLIKVGGLEGVREKYMEAVPNVTAILASSNFSFEYTNSCRIY

PKPDSLKLLRGPLDEDIPWPGFLLGQTPSSIWYWCADQVIVQRVLAAKNIAHAKGSTLMA

GVLKILPMFIIVIPGMISRILFPDELACIGPEHCMAVCGSQAGCSNIAYPRLVMSVMPVG

LRGLMMAVMIAALMSDLDSIFNSASTIFTLDIYKMLRARASSRELVVIGRIFVIFMVAIS

IAWVPVIIEMQGGQMYLYIQEVAGYLTPPIAALFLLGVFWKRCNEIGAFCGGMTGFVLGT

TRLLLGFIYREPKCDQPDERPAFITRIHYMYIAAGLFWISGLVAVVVSLCTTPPDEERIK

RTTMWGLRKLRRLVVTEREEMYKLTNGSGNGILKKDVPEDVQKEKCLDGADIKLLLPSPC

DQDPTTPNTEASTPMELYGNGHAKLVKQKDESQSDDESCLQVIDWICCHKEKTSVVEPKF

IKEDEETIREMLHEPPKIKLLLNICLVLICLLGIFMFVYFSL

***three spine stickleback (Gasterosteus aculeatus)***

>tr|G3NHQ8|G3NHQ8_GASAC Solute carrier family 5 (sodium/myo-inositol cotransporter), member 3b OS=Gasterosteus aculeatus OX=69293 PE=3 SV=1

SGMETVDVVVVALYFVLVLAIGFFAMWKSNRSTVSGYFLAGRSMTWIVVGASLFVSNIGS

EHFIGLAGSGAASGYAVGAWEFNALLLLQLLGWVFVPVYIHSGVYTMPEYLSKRFGGNRL

KVYFAFLSLLLYIFTKLSVDLFAGALFIQESLGWNLYVSIFLLISMTALLTVTGGLVAVL

YTDSLQAVLMICGALTLSILSLLKVGGLEGVRTKYMQAVPNVTAIMATGNFTYSPSCRID

PKPNALRLLRGPLDEDIPWPGFLLGQTPASIWYWCADQVIVQRVLAAKNLAHAKGSTLMA

GLLKILPMFLIVIPGMISRIMFVDELACIGPEHCVSVCGSQAGCSNLAYPRLVMAVMPVG

LRGLMMAVMIAALMSDLDSIFNSASTIFTLDIYKTARGEASQRELLIMGRLFVVLMVAIS

IAWVPVIIDMQGGQTYLYIQEVAGYLTPPIAALFLLGVFWKRCNEKGAFCGAMTGFALGV

TRLMLAFAYRQPRCDQPDDRPAFIVKIHYMYVATGLFWITGLVVVVVSLCTPPPDEEQVR

TTTFWGVRNMKTLPVDEKKSHCGADGSLEEEMPLNTPSTETSPVTTPPERFDSAKMEMIR

KESCHDTGETSRFVRLLECFCGCKEEAQSPKRNVLQEDARVIAERLYEPPQIKALLNMGL

LINCCLAISMYVYFS

***tilapia (Oreochromis niloticus)***

>tr|I3KXY0|I3KXY0_ORENI Solute carrier family 5 (sodium/myo-inositol cotransporter), member 3b OS=Oreochromis niloticus OX=8128 GN=slc5a3 PE=3 SV=1

MAGGMEAADIAVVALYFVLVLAIGFLAMWKANRSTVSGYFLAGRSMTWIVIGASLFVSNI

GSEHFIGLAGSGAASGYAVGAWEFNALLLLQLLGWVFIPVYIHSGVYTMPEYLSKRYGGK

RLKVYFAFLSVLLYIFTKLSVDLYAGALFIQESLGWNLYLSIVLLISMTALLTVTGGLVA

VLYTDALQAVLMISGALTLTIMSLIKVGGLEGVRTKYMQAVPNVTAILATGNYTYSPSCR

IDPKPNSLRILRGPLDEDIPWPGFIFGQTPASIWYWCADQVIVQRVLAAKNIAHAKGSTL

MAGFLKILPMFLIVIPGMISRILFADEIACIGPEHCMAVCGSQAGCSNIAYPRLVMAVMP

VGLRGLMMAVMIAALMSDLDSIFNSASTIFTLDIYQTFRTKACQRELLIVGRMFVVVMVV

ISIAWVPVIIEMQGGQTYLYIQEVAGYLTPPIAALFLLGVFWKRCNERGAFCGGMTGFTL

GTIRLILAFIYRQPRCDQPDNRPAFVVRIHYMYFAAGLFWISGLVAIVVSLCTSPPDKEQ

VRTTTLWGLYSIEMTPPKNREETYRLTEKNHCNGDTSLRKELPPDVKKERCVDGMDIKLL

VPSTDHDPATPSTETSPATTPGAQIGNRNMETIRAEEGCRSGAETSRCMRLLDWVCGYKG

ESQSAQQTGAKEDARAIAEMLYEPPRVKLLLNLGLVSVCSVGIFMFVYFSL

>tr|I3KDP7|I3KDP7_ORENI Solute carrier family 5 (sodium/myo-inositol cotransporter), member 3a OS=Oreochromis niloticus OX=8128 GN=LOC100707934 PE=3 SV=1

MVMEAADMIVVAIYFILVLGIGFLAMWKANRSTVSGYFLAGRSMNWAAVGASLFVSNIGS

EHFIGLAGSGAASGLSVAAWELNAVLLLQFLGWMFIPVYIQCGVYTMPEYLSKRYGGNRL

KVYFAALSLVLYIFTKLSVDLYSGALFIKESLGWNLYLSILLLIAMTALLTISGGLVAVI

YTDTVQAFLMIAGALCLTGISLVKVGGFEGVRLGYMEATPNISAILLSSPNLTYSESCYN

HATPKPDSLKILRAAQDPDLPWLGFLLGQTPASIWYWCADQVIVQRVLAAKNIAHAKGST

IMAGFLKILPMFIIVIPGMISRILFADDLACISPKHCVEVCGSAAGCSNVAYPRLVMSVM

PVGLRGLMMAVMIAALMSDLDSIFNSASTIFTLDIYKMLRKRASSRELMIVGRLFVVFMV

IISIAWVPVIIEMQGGQIFYYIQEVSDYLTPPIAALFLLGILWRRCNETGAFWGGMVGFF

LGALRLALAFVYREPHCDQPDERPSFIKDIHYMYVAAILFWISSLVAVVVSLCTPPPEEG

QIKTTTLWGLNKKKKLKNQEKNAEKDVNVLKPLNHSAFNGNGVLGEEKCPKNPNLLNGSE

ANLENGQSRSSNNVTATDIEAIPSVQNGGCAQSSGSKMIENEETRGVEDVEEEDESCGAG

DAKEERGCCIKTLDWFCGFKEELSKDQAITVQEQEKMVEELLYEPPTTRIILNIALVVVC

SVGIFLFIYFSL

***mudskipper (Boleophthalmus pectinirostris)***

>XP_020787565.1 sodium/myo-inositol cotransporter [Boleophthalmus pectinirostris]

MRPGMEAADIAVVALYFVLVLLIGFFAMWKANRSTVSGYFLAGRSMTWMIIGASLFVSNIGSEHFIGLAGSGAASGFAVGAWEFNALLLLQLLGWVFIPVYIHSGVYTMPEYLSKRYGGNRLKVYFAFLSILLYIFTKLSVDLYAGALFIKESLGWNLYLSIVLLISMTAILTVTGGLVAVLYTDALQAVLMIGGALTLTIMSLIKVGGLDGVRTKYMQAVPNVTAITAAGNFTYSPSCRIHPKPDSLRILRSPLDEDIPWPGFILGQTPASIWYWCADQVIVQRVLAAKNIAHAKGSTLMAGFLKILPMFIIVIPGMISRILFADEIACIGPEHCMSVCGSQAGCSNIAYPRLVMAVMPVGLRGLMMAVMIAALMSDLDSIFNSASTIFTLDIYQTVRRAASQRELLIVGRLFVVFMVGISIAWVPVIIEMQGGQTYLYIQEVAGYLTPPIAALFLLGVFWKRCNEKGAFWGGMTGFVLGTLRLILAFIYRAGRCDQPDTRPLFVSQVHYMYVAATLFWISGFVAVVVSLCTSPPDEEQVRTTTVWGLHNLEKRPVKDREEIYRLTHNGSSGGPCKEVPLDVSKQMCLDGADVKLLVLSTDHDPATPSTEASPVTSPADHFGNTVRGSEEGWLKERKTNTCVRFLDWLCGYKDETDGKQSTQQKVVQDEIRIAQELLYEPPGVKLLLNIFLVCVCCVGIFLFVYFSL

>XP_020773420.1 sodium/glucose cotransporter 1-like [Boleophthalmus pectinirostris]

MSRDYFGFSYVKNAARNGTSSLLNNAPDISVIVIYFLVVLAVGVWAMVRTNRSTVGGFFLAGRSMVWWPIGASLFASNIGSGHFVGHSGTAAAAGIAIAAFELNALIIVVLLGWVFVPIYIRAGIVTMPEYLKKRFGGQRIRIYLSVLSLCLYVFTKISADMFSGAIFIRQALGLNIYLAVIALLAITALYTVTGGLAAVIYTDTLQTIIMLAGSFVLMGFAFNEVGGYEQFQQRYMTAVPSITGNISEECYKPRADSFHLFRDAITGDLPWPGLVFGLTIQATWYFCTDQVIVQRCLSAKSLSHVKGGCILCGYLKLLPMFLMVFPGMISRILYPDEVACVEPAECERFCGTEVGCTNFAYPKLVVDLMPNGLRGLMLSVMLASLMSSLTSIFNSASTLFTMDIYTKIRRSASERELMIAGRIFILALIGVSIAWIPVVQNAQSGQLFDYVQSITSYLTPPVVAVFMLAIFCKRVNETGAFYGLMIGLVIGLCRMIPEFVYGTGSCVNPSNCPTIICGVHYLYFSIILFTISCILILGISLMTKPIDDKHLYRLCWTLRNNTEERIDIEVDDWIDNQNDKIMEIDEKEHEKEKPGLCKKAILCFCGLEQQKDTVKLTPEEEEEMKRKLTDTSEVPLWRNVVNANALILLCVAVFFHGFYA

>XP_020783330.1 sodium/glucose cotransporter 4-like [Boleophthalmus pectinirostris]

MSTTSTTDSSVAPPTAAPPSRLSLGTADIAVLVAYFVFVMVVGIWSSIRANRSTVGGYFMAGRSMTWWPIGASLMSSNVGSGLFIGLAGTGAAGGLAVGGFEWNAAWVLVALGWVFIPVYIAAGVLTMPEYLCRRFGGARIRIYISLLSLILYIFTKISTDIFSGALFIQVSLGWDLYLSTAVLLLVTAAYTIAGGLAAVMYTDALQTVIMVGGAFSLMFIAFSRVGWYEGLEQLYPKAVPSVTVPNSTCHLPRHDAFHLLRDPLTGDLPWPGLIFGLTVLATWVWCTDQVIVQRSLSAKSLSHAKGGSVLGAYLKLLPMFFVVMPGMISRALYPDEVACVVPELCQKVCGTAVGCSNLAYPKLVVELMPSGLRGLMLAVMLAALMSSLTSIFNSSSTLFTLDLYQRLRPAASERELMVVGRLFILFLVCISLLWIPVIQTANSGQLFDYIQSVTSFLAPPITAVFLLAIFWSRTNEQGAFWGLMGGLIVGGIRMALEFSYSAPSCGQPDLRPSVLAQVHYLYFSMLLFALSTLLICAVSLATEPIPPHSLHRLTWWSRHSQEPRVELPGVTSGSESPQASSSDSQAPRGWCYRALLWMCGLTGQSSSSVDPDCSPLVSLQEEPLWRHVCNINALLLLTVNIFLWGYFA

>XP_020795862.1 sodium/glucose cotransporter 1-like [Boleophthalmus pectinirostris]

MSRDYFGFSYVKNAGRNGTYAALNNPPDISVIVIYFVVVLAVGVWAMVRTNRSTVGGFFLAGRSMVWWPIGASLFASNIGSGHFVGIAGTGAAAGIAIGGFEWNALIVVVILGWLFVPIYIKAGVVTMPEYLKKRFGGQRIRIYLSVLSLCLYVFTKISADMFSGAIFINQALGLNIYIAVIALLAITALYTVTGGLAAVIYTDTLQTIIMLAGSFVLMGFAFNEVGGYEQFQQSYMTAVPSNTSNISEQCYQPRADSFHLFRDAVTGDLPWPGLVFGLTIQATWYWCTDQVIVQRCLSAKSLSHVKGGCILCGYLKLLPMFLMVFPGMISRILYPDEIACVVPEECEKYCGARVGCTNIAYPKLVVDLMPNGLRGLMLSVMLASLMSSLTSIFNSASTLFTMDIYTKIRRSASERELMIAGRIFILVLIGVSIAWIPVVQNAQSGQLFDYVQSITSYLTPPVVAVFMLAIFCKRVNETGAFYGLMIGLVIGLCRMIPEFVYGTGSCVNPSNCPTIICGVHYLYFSIILFTISCILILGISLMTKPIDDKHLYRLCWTLRNNTEERIDIEVDDWIDNQDDKKMEIEEKKHENKPGLCKRAILCFCGLEQQKETVKLTAEEEEEMKRKLTDTSEVPLWRNVVNANALILLCVAVFFHGFYA

>XP_020788177.1 sodium/myo-inositol cotransporter 2 [Boleophthalmus pectinirostris]

MWRTKRNTVDGYFLAGKSMTWWPVGASLFASNIGSGHFIGLAGSGAAAGIGAIAYEWNGMFMILLLGWIFLPIYLASGVTTMPEYLQKRFGGRRTQLLIAVLYLFIYIFTKISVDMYAGALFIQLALQWNIYLAVVLLLSITAVYTVAGGLAAVIYTDAAQTMIMLAGALTLMGFSFAEVGGWNALMGGYADAIPSVRVPNTTCGIPREDAFHLFRDPVTSDLPWPGVLIGMSIPSMWYWCSDQVIVQRSLAAKTLTHAKGGSLLAVYLKVIPFFAVMLPGMISRILYTDDVACADPDLCKEICGNPVGCTDTAYARLVMELLPAGLRGLMMAVMIAALMSSLTSIFNSASTIFTMDLWKTFRSRASEWELMIVGRVFVLVLVGVSVLWIPVVQASQGGQLFIYIQSISTYLQPPVSIVFIMGCFWKRTNEKGAFWGLALGLLIGCIRMLLDFIYPQPQCYQTDDRPGVLKHVHYLYFSMLLSAITLIVVVTVSLVTEEPRPEQISRLTWFTKSDPVVFTVEQSINGVSLQRRASTSSSVRVHQSKLVSALYWLCGMERRQEDSKPPPPVQPDVCSLEEKPHLKKLVNVNLIICLTVTVFIIGYWA

>XP_020788626.1 sodium/glucose cotransporter 5 [Boleophthalmus pectinirostris]

MSNYTINFYASPKSFSFSDIIVIVIYFLLNLAVGIWSSCRVSRNTLSGYFLAGRDMAWWPIGASLFASCEGSGLFIGLAGTGAAGGIAVAGFEWNATYALLALAWLFVPVYISSGIVTMPEYLGRRFGGERIRTYLAVLSLLLSVFTKISTDLYSGALFVQVCLGWNLYLSTVLMLVVTALYTIAGGLAAVIYTDTLQTFVMIMGAIILTITAFNKIGGFGNLAQAYSIAIPRKIIPNSTCHLPRADAMHLFRDAVTGDLPWPGMTLGLTILATWYWCTDQVIVQRSLSAKNMSHVKGASILAAYLKFLPFIFIILPGMISRALYPDSVGCVDPEECVRVCGAEVGCSNIAYPKLVIELMPSGLKGLMIAVMMAALMSSLTSIFNSSSTLFTMDIWKKHRPAASEKELLLVGRIVTVILVVVSVVWIPILQSANSGQLYVYIQSVTSYLAPPVTAVFTLAVFWKRTNEQGAFWGLMVGLVVGVCRMVLEFAFPPLRCGLEDSAPSVLRDVHYLHFAILLCGLTAGVVVVVSLVTPPPSQEQLFNLTWWTITEPPPRDIPLQKVSSVSHRSSQEPVRRGPCGLGPTSCMGRGSEQAPAPRVPSVKESVFWSRFCCANAILLVSVNIFLYAYFA

>XP_020783839.1 sodium/myo-inositol cotransporter-like, partial [Boleophthalmus pectinirostris]

VWSKYSQASPNITSIMSSRPNLSFADSCLLHLRPKPAALKILRGASDPDLPWPGFLLGQTPASLWYWCADQVIVQRVLAAKNMAHVKGSTIMAGFLKILPMFIIVIPGMISRIFFADDLICISPEHCAEVCGSPAGCTNLAYPRLVMSVLPVGLRGLMLAVMIAALMSDLDSIFNSASTIFTLDVYKMLRARATSRELLIVGRLAVVFVVVISIAWVPVIIEMQGGQMFYYTQEVSNYLTPPIAAMFLLAVLWHRCNETGAFWGALTGFVLGTVRLALAFVYREPHCDQPDTRPSFIKDVHFMYVAAVLFWISGLVTVVVSLLTPPPTEEQIQTATFWGLRNRKKRILPESNELKPLNDPEVNSNANLNGQTPTSECSAAEIQNRRPKQNDVIANSKCLRVVNWFCKINEDPASTYKPSAQEQERMVDELLYESPKMKIVLNFALALICGVGVFLFVYFTL

***Homo sapiens***

>sp|Q8WWX8|SC5AB_HUMAN Sodium/myo-inositol cotransporter 2 OS=Homo sapiens OX=9606 GN=SLC5A11 PE=1 SV=1

MESGTSSPQPPQLDPLDAFPQKGLEPGDIAVLVLYFLFVLAVGLWSTVKTKRDTVKGYFL

AGGDMVWWPVGASLFASNVGSGHFIGLAGSGAATGISVSAYELNGLFSVLMLAWIFLPIY

IAGQVTTMPEYLRKRFGGIRIPIILAVLYLFIYIFTKISVDMYAGAIFIQQSLHLDLYLA

IVGLLAITAVYTVAGGLAAVIYTDALQTLIMLIGALTLMGYSFAAVGGMEGLKEKYFLAL

ASNRSENSSCGLPREDAFHIFRDPLTSDLPWPGVLFGMSIPSLWYWCTDQVIVQRTLAAK

NLSHAKGGALMAAYLKVLPLFIMVFPGMVSRILFPDQVACADPEICQKICSNPSGCSDIA

YPKLVLELLPTGLRGLMMAVMVAALMSSLTSIFNSASTIFTMDLWNHLRPRASEKELMIV

GRVFVLLLVLVSILWIPVVQASQGGQLFIYIQSISSYLQPPVAVVFIMGCFWKRTNEKGA

FWGLISGLLLGLVRLVLDFIYVQPRCDQPDERPVLVKSIHYLYFSMILSTVTLITVSTVS

WFTEPPSKEMVSHLTWFTRHDPVVQKEQAPPAAPLSLTLSQNGMPEASSSSSVQFEMVQE

NTSKTHSCDMTPKQSKVVKAILWLCGIQEKGKEELPARAEAIIVSLEENPLVKTLLDVNL

IFCVSCAIFIWGYFA

>sp|P53794|SC5A3_HUMAN Sodium/myo-inositol cotransporter OS=Homo sapiens OX=9606 GN=SLC5A3 PE=1 SV=2

MRAVLDTADIAIVALYFILVMCIGFFAMWKSNRSTVSGYFLAGRSMTWVTIGASLFVSNI

GSEHFIGLAGSGAASGFAVGAWEFNALLLLQLLGWVFIPIYIRSGVYTMPEYLSKRFGGH

RIQVYFAALSLILYIFTKLSVDLYSGALFIQESLGWNLYVSVILLIGMTALLTVTGGLVA

VIYTDTLQALLMIIGALTLMIISIMEIGGFEEVKRRYMLASPDVTSILLTYNLSNTNSCN

VSPKKEALKMLRNPTDEDVPWPGFILGQTPASVWYWCADQVIVQRVLAAKNIAHAKGSTL

MAGFLKLLPMFIIVVPGMISRILFTDDIACINPEHCMLVCGSRAGCSNIAYPRLVMKLVP

VGLRGLMMAVMIAALMSDLDSIFNSASTIFTLDVYKLIRKSASSRELMIVGRIFVAFMVV

ISIAWVPIIVEMQGGQMYLYIQEVADYLTPPVAALFLLAIFWKRCNEQGAFYGGMAGFVL

GAVRLILAFAYRAPECDQPDNRPGFIKDIHYMYVATGLFWVTGLITVIVSLLTPPPTKEQ

IRTTTFWSKKNLVVKENCSPKEEPYQMQEKSILRCSENNETINHIIPNGKSEDSIKGLQP

EDVNLLVTCREEGNPVASLGHSEAETPVDAYSNGQAALMGEKERKKETDDGGRYWKFIDW

FCGFKSKSLSKRSLRDLMEEEAVCLQMLEETRQVKVILNIGLFAVCSLGIFMFVYFSL

**Na^+^/H^+^ EXCHANGERS (NHE)**

***Round goby (Neogobius melanostomus)***

>NEME_00002007_Neogobius

MAVLRQYALCFCVLLFIWLCQASDPHRADHADDHADDHTDDHGDNHTGTDESDGHSEHGA

PITTLPIVTWKWPHLSTPYLVALWILVCWLCKLVLLSGVEADAKSDHHSENGHHQNASSI

SGIPIVTFKWHHVETPYTVALWIIVAGLCKLVIEANHHVTNVIPESALLICSGFILGGMV

WGADKKQTFKLNPTVFFYYLLPQIILDAGYSMPNKLFFTNLGGILLHAVIGTCWNAATVG

LSLWGCHMGGAMGLLQYLLFGSLLSAVDPVAVIAVFEQVHVNEVLFIMVFGESLLNDGVT

VVLFNVFDSFVSLGGPAINAAEIVKGIISFFVVAFGGSLVGLVFGLLLSLLTRCTKNIQI

IEPGFVFVVGYLSYLTAEMLSLSAILSITFCGVCCQKYMNANMDEKSVTTVRYAMKVLAN

GSETMIFVFLGITAIDKTIWVWNTGFILLTLLFVLVYRVIGVFFLTWIQNKFRLVPLDLM

DRIIMSYGGLRGAVAYGLAALLDENKIKEKPLMIGTTLIIVYFTVVLQGITMKPLVTWLK

VKRATNNELTLIEKVQNKVFNHMLVAIEDISGQKGNNYMRDKWTHFEEKWMSWFLMKPSA

RKKHDHLFNIFHMINLKDALEYVDKGESNGTLEFIRNDTAFIDFKKNYEDDFSDVSMPDL

TIDMGDDTIANPYARQTRDAVPSISLEMHKQTNYGLRESENINTHHLLQQHLYKSRNQHR

FRYSRSHFDLNQDENEVQEIFQRTMRNRLESFKSAKLGVAPPKPITKHPRKDTQKKMPNG

KSTEKRSYYSGDEDFEFSEGDSASGYFPRVTHRPGTGIENPSFMPDLDPRAPVQIPPWLA

EGELDSTAGVAPSQRAQAGLPRTPSNLNRLAPLRISTRSTDFPTSSSSQQSNELPPPPSP

PPYHPHKI

>NEME_00018498_N-trunk_Neogobius

VTVDTSKLEHLFESKAKELPMAKKILTMVPTEEEKQQIQEAQLASPGAKGFELSYLEKVV

EVKDTVQRQSLLHHTCNLVLEKYPESTDLYSEIPAITRSAKVDFELLSENLSQLERRCKA

SWDNLKLVAKHETKAALKNKLTEFLKDCTQRIIILKVVHRRIINRFHSFLLFLGQPSCTA

RNIKVTNFCRITEKFSGAVPQDSPSPVSMAAEEEAGQQEEHENMTNLLISGGSDSNGRLS

MSQVTASKEDGGAQDDATDEIMDRLVKSVTQNPSDRQSSPKSRKRSRVNRKSRAGGAGSL

DRGGGGGARVGAEDEEDQHQDHQHGGGYKVVQWEWSYVQTPYIIATWLLVASVAKILFHF

SQRFTKVVPESCMLILLGLVLGGIVLIANKKQLYQLEPALFFLFLLPTIVGDAGYFMPAR

LFFDNLGAILTYAVVGTLWNAFCTGFCLYAAKVLNVIDAKVQADLMDFLLFGALISAVDP

VAVLAVFEEVHVNDTLFIIVFGESLVNDAVTVVLYKVYISFVQVGAANVQTADYFKGVAS

FLIVSIGGTLVGLVFAVILGFITRFTKKVRIIEPLFVFLLVYLAYLTAELFSLSAILSMT

FCGIGANKYVEANISQKSRTTVKYTMKTLASIAETIIFIFLGISVVDKSKWAWDTGLVSC

TLVFILVFRAIGVVGQTWVLNRFRLVPLDKIDQVMMSYGGLRGAVAFALVVLLDGEQVKA

KDYFVATTIVVVFFTVMFQGLTIKPLVKWLKVPRSTSRKPTINEEIHERAFDHILTAVED

IAGLQGYHHWRDKWEQFDKKYLSKLLLRKSVYTKSELWEAYQKINIRDAISVIDQYQSHY

SRHFMPLGEKERQDREVFQRNMKIRMESSKSTRHKRHKKERSIKKRRGSDSKEESSEKPR

RNVSWHDKDRSDAEKEDDEGITFVAQKVETPKPRPKSVPAALEVSSSPPARSPSPPLSAD

SHLPWKGGVGSPPPCVSVEATKIIPVDLQQAWNQSISSLESISSPPAPPVPDPLHPRISA

LSRLGAPRPVSYTPPSSSSSSTAAGTSFNTSNTQLLHGSFLFPDRRLKEEEEDDSFESSQ

ELQPLMSTLKPPAEAIPPPPHLQAPSSRGKGARRTPACTSGAWCRRPLMAARSRAAEDPP

SFRR

>NEME_00017961_Neogobius

MNADVFFLCLLPPIILDAGYFLPIRPFIENIGTILMFAVIGTLWNTFFIGGLLFAVCQIN

PGYTMDLGLLPCLLFATIISAVDPVAVLAVFEEIHINELLHILVFGESLLNDAVTVVLYH

LFEEYAGAGTVTVLDGFLGIICFLVVALGGVLVGAIYGILAAFTSRFTSHMRVIEPLFVF

VYSYMAYLSAEMFHLSGIMALIACGAVMKPYVEANISHKSHTTVKYFLKMWSSVSETLIF

IFLGVATVDGTHHSWNWIFVTATVILCLIARVIGVIGLTYLMNKVRIVKLTTKDQFIIAY

GGLRGAIAFSLGFLLNEEHFPQKKMFLTAIITVVFFTVFVQGMTIKPLVELLAVKKKQEA

KRTINEEIHTQFLDHLLTGIEDICGHYGHHHWKDKLNRFNKKYLKRYLIAGTRCEEPQLL

SFFHKMEMKHAIEQVEKGGGSNVPSALPSTVSMQNIQPKKPTVEPERMLPKLPKGKEDEI

RNILRTNLQKTRLRLHSYNRHTLVADPFEEGISDIILRKRVIQKQMKDINNYLTVPAAPP

DTPVMNPQARGAKYNADNVPTFQVDLASPQSPDSVTLLNELKKVEDQGLMMKAPSSKSME

GERDQQKLTRCLSDPGPSADEDEDEPFLK

>NEME_00035431_Neogobius

MGTFYNLGWRGRAPWCSLLVLLLCCSNGGNCEAETPKAPEFSDPQGVKPLENEMPQAFPE

EEKAYLPFFTMDYPRIQVPFEFTLWVLLASFAKIGFHIYHKITIWIPESCLLICIGLIVG

GIMYSVKEEPPAVLRSDVFFLYMLPPIVLDSGYFMPTRPFFENMGTVLWYAVVGTLWNSV

GIGMSLFAICQIDAFGLQDINLQENLLFASIISAVDPVAALNVFEDIQVNEQMYIVIFGE

GLFNDAVTVVLYNMFSFIADMPTVESADVFLGIARFMVVSLGGILFGLLFGIAAAFTTRF

THKIREIEPLFVFMYSYLAYLVAELFAISSVLSIIVCAITMKYYVEENVSQRSCTTIRHV

VKVLASISETLIFFFLGVVTITTKHEWNWAYILFTLLFAFIINPFRTIPFSKQDQFGLVY

GGLRGAVCFALVFTLPDTINRKNLFVTASIAVIIFTVFIQGISIRPIVEYINIRRSNNVK

YNINVEIHTRTMEHLVSGIEDLCGQWSHYYWKDKFKKFNDRVLRRILLRDNRAESSIVSL

YKKLELQSAIDLLESPMSDLSAAPSIVCLYDERMDTSRPKKTLSTADTRKIQDMLAKNMY

KIRQQTLAYTNKYKLPDESHTREILIRRHASHRRSLRAASFRDPTSQNIPKSQRYFSLQP

GTDLESAFVLRRRSHGRWSSSAHSAIPMSPIEERAQSPLAQNSTSTEQQENEPGEGAAAR

GWTPEHPRENTVSTNPLLRK

>NEME_00015753_Neogobius

MASLLVSLLIYACFVVGCSSLTRPVDTPEPQSQPLSLGQNGSSDGVPSVDVPMALRVFSV

DYHHVQAPFEIVLWIMLASLAKLAVVPESCLLIMVGLLVGGVIYGVRHSAPPTLSADAFF

LFLLPPIVLDAGYFLPGRLFFENLGTILWYAVLGTLWNVLGIGLSLYGVCLLCPSSLGDV

SLLHCLLFGSLIAAVDPVAVLSVFEEMHVNEQLHILVFGESLLNDAVTVVRQKQECVRVL

YQLFESFLRMPSVSGLDILLGGSRVLVVGFGGFFLGLFFGLVAALTSRFTPRVPVIAPLF

VFLYSYLSYLTSEMLHLSGIMAIVTCAVTMKQYVEANVSERSNSSIRYFLKMWSSVSETL

IFIFLGVSTIQDVHMWSWAFVCATLLLCLLWRASGVLLLTAVVNKVRRNKVTFRDQFIIA

YGGLRGAICFSLVFLIDDFPKKRLFITTTIIVILFTVFVQGMTIKPLVDLLDVKRKKRAL

PTVSEEIHSRLIDHLLSGIEDVVGYWGQHYWKDKFEQFNRKFLRRFLIREEHQDRSSILR

VHQELERREQRGDGETAITVLEPVSPSSLTLLHEGFFWRCSFQVELSAEPHLATLFTNQP

ESKVCVHGYLFPLPVSLGPVPHPMYDSTAVFRGFWTVLMILALLCAPIGGFLLVCGIPFY

SHKLYRVGGALLIAAACLFLSVLLLYVMWMELVDVKRYILQEKGESCPGADVQIHYGLSF

MVAAAAVPLVLTSGTVFMCVARALSGDKL

>NEME_00014717_Neogobius

GCLFGGFLFLIHLNVPVVINPKVFFQILLPHVILDAGYFLPVRPENLGTITMFAIVGTLW

NALFIGGGLFIVCNFWTDLSLELLPCLLFGAITSAVDPVAVLEVFEAVHINELVHILVFG

DMSSRTLLLVLVLLVGSALHSAAAEPVDHETNHNSTPTNNTHSTHKVKKPFPVLGLNYQH

VQMPFEISLWVIFGSVMKLGFHMVPCLNSIVPESCLLIIVGLLFGGFIRLAGKEVPPVMN

ADVFFLFLLPPIILDAGYFLPVRPFIANLGTILMFAVVGTLWNAFFIGGLLFAMCELLPD

YKLRLLPCLLFGSIISAVDPVAVLAVFEEIHINELLHILVFGESLLNDAVTRWHRDGIGR

LPRHHLLLCGSAGGLFVGAVYGILAAFTTRFTAHMRVIEPLFVFAFSYLAYLTAEICHLS

GIMAIIACAVVMKPYVEANISQKSHTTIKYFLKMWSGVSETLIFIFLGVATVGGKNLAWN

WIFMTATVFLLTHLMNKVRIVKLSHKDQFIIAYGGLRGAIAFSLGFLLDPDHFPEKDMFL

TAIITVIFFTVFVQGMTVKPLVECLAVRKKVEAKRTINEEIHTEFVDFLLTGMEDICGHR

YSGEENQLLKFFHKRQIKQAIALVEKGGGTSAMPSTESMQNIQPKMPEEAPKLSQGTQNK

IEAVLLTRLQKIRQKQQSYSRHNLPADPFEEDTSDVLLSKKVPKKQMNTYPTVSAIPPNT

PYPSRARTRLATGPQAQVHMEPPKAPDSMTPVDELKKDEDQKLSMKTPPSKSGE

>NEME_00032425_Neogobius

MLSLWGVRLTYLMNKFRIVKILIKDLPILSYGGLQGAIAFSLGFLLYPDHFPEKDIVPVV

INPKVFFQILLPPIILDAGRPVRPFTENLGTNTIMSSRTLLLVLVLLVGSALHSTAAMPL

DHETNHNSTPTNDTHSTHKVKKGFPVLDINYQHVQMPFEISLWVIFGSIMKLGFHMVPRL

NSIVPESCLLIIVGLLLGGFIRLAGIEVVMNADVFFLFLLPPIILDAGYFLSVRPFMENL

GTILMFAVVGTLWNAFFIGGLLFAVCKLIPGSKLQLLHCLLFGSIISAVDPVAVLAVFEE

IHINELLHVLVFGESLLNDAVTLEYAGAGTVTALDGFLGIISFFVVALGGLFVGAVYGIL

AAITTRFTSHMSVIEPLFVFAFSYMAYLTAEIFHLSGIMALIASAVVMKPYVEANISQKS

RTTIKYFLKMWSGVSETLIFIFLGVATVGGKNLTWNWIFMTATGRIVKLSNKDQFIIAYG

GLRGAIAFSLGFLLDPDHFPEKDMFLTAIITVIFFTVFVQGMTIKPLVECLAVRKKQKAK

RTINEEIHVEMEPIQREVFEEVADHRIQRGGEPAAYREMTQAIELVEKGGGTSAMPSTVS

KQNIQPKIPEVGPKLSQGTQDDIAAVLLTNLQKIRQKQPNYSRHTLPDDPDEKTSNLLLR

NRVAKEKNTYPTVSAFPPTRPTQKNQTGLSFVKSPQAQTTKSNTDNEVTVQVHMEPPRSP

DSVTLEDEDQELSMKTPPSKSGE

>NEME_00008682_Neogobius

MKSITESCPAGSAPAAAASYTRCMSVSAESSETLLSGCMYRVLWKSFQSNNAETERQAEE

SHRQDSANLLIFIMLLTLTILTIWLFKHRRFRFLHETGLAMIYGLLVGVILRFGVHVPPS

TSDVVLSCAVNASPATLLVNVSGRFYEYTLKGEVSRAKGHQVQDDEMLRKVTFDPEVFFN

ILLPPIIFHAGYSLKRRHFFRNIGSILAYAFVGTVISCFVIGLIMYGFVSFMKVVGQLGG

DFFFTDCLFFGAIVSATDPVTVLAIFNELKVDVDLYALLFGESVLNDAVAIVLSSVLLRF

QKQSRQAFVSSGDRGPVLLQGDHEMMGQNDTGTSRSIVAYQPVGDNSHSFEAMAMLKSFG

VFLGVFSGAFALGVATGVFTKLRDFPLLETALFFLMSWSTFLLAEACGFTGVVAVLFCGI

TQAHYTYNNLSPDSQDRTKQLFELLNFLAENFIFSYMGLTLFTFQSHVFNPLFIIGAFLA

VFLGRAANIYPLSFLLNLGRRNKIGSNVQHVMMFAGLRGAMTFALSIRDTATYARQMMFS

TTLLIVFFTVWVCGGGTTPMLSFMSIPVGVDSDQENTSTTVLDGSQRRNTKHESAWPFRI

WYNFDRNDSSLHLWLFMVQVNFEIHTTNTITVLVLDSVHSYLKPLLTHSGPPLTATLPSC

CGPIARCLTSPQAYENEGPLHDDDDFILNDASVSSMFADVTVSTDASGSRTVNRKRSGAG

LHDDGLDNELAQSENEIAIRGTRLVLPMDDPLDPPPQSHCPLRPPRPNLTRAGTDCRQPP

DSPNSTALHPRLHPSVPGESVKMADGGTHCLYCREDLGGKKFVRNEGRPVCVRCHTKFCA

NACHECHRPISVETKELTHKNRHWHEECFRCAKCYKPLAKEPFSTKDERILCGKCCSRED

APRCHGCYKPILAGTESMEYKGNSWHEDCFTCTNCKRPIGTQSFLSKGDDIFCSSCYDKK

FAKQCFSCKKPITSGGVTYQEQPWHGHCFVCTSCSKPLAGTSFTTHQDQAFCVDCYKSSV

AKKCAACHNPITGFGKGINVITFEGSSWHEYCFNCKRCSLSLSNKRFVANGKDILCSDCG

NK

>NEME_00003765_Neogobius"

MRRYRGSHGARLGVWRMPLVFAVLVTGLSSGGKASDAMEELATEKEAEESHRQDSVNLLT

FILLLTLTILTIWLFKHRRVRFLHETGLAMIYGLLVGVILRYGIPATSYHNQTPLSCSLK

KGPASTLLLNVSGKFFEYSLKGEINLKDIHNVEQNDMLRKVTFDPEVFFNILLPPIIFHA

GYSLKKRHFFRNLGSILTYAFIGTVVSCFVIGNLMYGVVKLMQVTGQLLDKVYYTDCLFF

GAIISATDPVTVLAIFNELHADGDLYALLFGESVMNDAVAIVLSSSIVAYQPSGANTHKF

DASALFKSIGVFIGIFSGSFVMGAANGVVTALISFTSVLHVSQITHNVFTKLHCFPLLET

ALFFLMSWSTFLLAEACGFTGVVALFEVLHFLAENFIFSYMGLALFTFQNHVFSPIFIIG

AFIAIFIGRALNIYPLSFLLNLGRQHKISGNFQHMMMFAGLRGAMAFALAIRDTATYARQ

MMFSTTLLIVFFTVWVFGGGTTPMLSWLHIRVGVDPDQDFQPTADSFQVLQGDGSQDQSR

TKQESAWFFRLWYTFDHNYLKPILTHSGPPLTSTLPSYCGPLASCLTSPQAYEDDDQVGE

PDTDSIRNDQDLTFMFGDTALTANGTSADRGRVDEGGPSWSVNGKRSASTSEEALERDLD

SRDQELLSRGTRLVFPTQDQ

>NEME_00001997_Neogobius

MAYLSAEVFHLSGIMSLISCGVMMRPYVEANISHKSYTTIKYFLKMWSSVSETLIFIFLG

VSTVVGPHSWNWTFVIVTVILCLVSRVLGVIGLTYIINKFRIVKLTKKDQFIVAYGGLRG

AIAFSLGFLLKDDDELKNMFLTAIITVIFFTVFVQTYVKRCLIAGDRSMEPQLISFYNKM

EMKQAMMLVESGSSSKLPSLVSTASLQNIQKMSKKRAMPSISKKREDEIRKVLRGNLQKT

RQRLRSYSRHDLMDPFEDNVSEIRFRKQRVEMERRRTVLFVAEHRCFIESASIIMSRLYV

GRLSYRAREKDVERFFKGYGKILEVDLKNGYGFVEFDDPRDADDAVYDLNGKELCGERVI

VEHTKGPRRDGGGGGEEGDTGAAEATVEEEEEEAGEEEEEAAAAAAEPGEADYMRQAGEV

TYADTHKGRKNEGVIEFRLYADMKRALEKLDGTEVNGRKIRLIEDRPGARRRRSYSRSRS

RSRSRRSRKSRSRSMSSSRSRSRSRAAASRSRSRSRSKKGKAKGKKEDEERSQKNKDQSR

SRSRSRSHSSKPKKSKKEAKKNKKDVSRSPSRSKSRSKSRSKSGTKDRPKKSGSEGRDAP

RSDTEGADGDRTSRSRSRSPVKSKSRARSKSRSKSMSKSRSPSPTNGKRRSVSRSPSRSV

SRSKSRSLSRSRSRS

>NEME_00008284_Neogobius

MSFPVTVGGAWRAMGTRWPFVLVSLSVCACVCRAASPQEEEDSAMENIVTEKKAEESHRQ

DSADLLIFILLLTLTILTIWLFKHRRFRFLHETGLAMIYGLLVGVVLRYGIHMPRDGTSV

MLNCHVNASPATVLVNVSGKFYEYTLKRVLSSSGNKVDDVSDNENEMLRKVTFDPEVFFN

ILLPPIIFHAGYSLKRRHFFRNMGSILAYAFLGTVVSCFIIGLLMYGTVTLMKHVGQLHG

EFFFTDCLFFGAIVSATDPVTVLAIFHELQVEPDLYALLFGESVLNDAVAVVLSSSIVAY

QPTGDNSHTFEGMALLKSFGMFLGIFSGSFALEIFGHLNCCIYVTKFTKLRDFQLLETAL

FFLMSWSTFLLAEACGFTGVVAVLFCGITQAHYTYNNLSRLRGAMTFALAIRDTATYARQ

MMFSTTLLIVFFTVWICGGGTTQMLSCQRIRYLKPILTHSGPPLTATLPPCCGPLARILT

SPQAFENEGQLKDDDSDLILTDDINLAYGDITVNTDGSGVHTSSGAGFASGMSSEDLDRE

LTCGDNEHSFFIKGPRHYMSSTMTDRFDCYYCRDNLHGKKYVKKDDKHVCPKCFDKLCAN

TCAECRRPIGADAKELHHKNRHWHEDCFRCAKCYKPLASEPFCARDDGKIMCGKCSSRED

GNRCQGCYKVVMPGSQNVEYKNKVWHEECFTCFECKQPIRTQSFLAKGDDIYCAPCHDKK

FAKKCFHCKQPITSGGISYQDQPWHSECFVCATCRKPLAGTRFTSHDNHAYCVDCYKTDV

AKKCHGCKNPITGFGHGTNVVNYEGYSWHEYCFNCKKCCLSLANKRFVISGDHVYCPDCA

KKL

>NEME_00018498_Neogobius

LSSLFTPLSGCAHAAMASITCRVQYLEDSDPFVCTNFPEPRRPPLVELGQDLPLSEQIAG

IHNLLNAPLKLEECTLQLSPSGNYLDLDSSLSEQRDELENFYADVEKGKKPIVILRTQLS

VRVHSILEKLYNSHGPELRRSLFSLKQLFEDDKDLVPEFVASEGEKPWSYIIEVLDERNL

ADTELLMFTMTLINKTLAALPEQDSFYDVTDRLEELGMEKIIQKHLNNNATEPDLRAQFT

IYENALKYEDGETDDLLSPYPRKERRKPASSSDQDTLRKSRRASSQNLPDLLPSTSSSSS

SSASPSPTSSLQNRVPSPTVTSPLPTQNNKAPTFTLLSPTQDVLSPTISSPSPTENPMSP

YVLSPSPTQDGDYLDLTNYVPSPTISTPTLTPTSPDLNANVFSPAEDTASPPHSHKSDSR

PTSPLAVNSDQSSPASSPATSQRGSPLPTHTVPNGNVATEQEAKVPQSPGRSFLSHHMSA

LGLGRKSRLFSKNSSISEEPSAAHSPASPEPSHSEQQQQEQPQSKSEGKTIFKDNFLRNL

AATQWEKRRKSKLQRSRRSSADDLAAPSPPPTEETGSAEAEEQTPQSPTNRAPDTNGHTE

PQNDTSSYQRHCSTLSDDKKFVLDMLYSKSNPGPLSPTEPSTEEGQQTERDLLDEEDIDV

LDMDTFDSSQAFTVSGVPPPPPPPPGAGPPPPPPLPPPPGLPSAAPPPPPPPPPPPPPGA

APPPPPPPPPVLNAVKASPLGDAPKKKKTVKLFWRELKHADGGDQKCRFGRGTVWASLDK

VTVDTSKLEHLFESKAKELPMAKKILTMVPTEEEKQQIQEAQLASPGAKGFELSYLEKVV

EVKDTVQRQSLLHHTCNLVLEKYPESTDLYSEIPAITRSAKVDFELLSENLSQLERRCKA

SWDNLKLVAKHETKAALKNKLTEFLKDCTQRIIILKVVHRRIINRFHSFLLFLGQPSCTA

RNIKVTNFCRITEKFSGAVPQDSPSPVSMAAEEEAGQQEEHENMTNLLISGGSDSNGRLS

MSQVTASKEDGGAQDDATDEIMDRLVKSVTQNPSDRQSSPKSRKRSRVNRKSRAGGAGSL

DRGGGGGARVGAEDEEDQHQDHQHGGGYKVVQWEWSYVQTPYIIATWLLVASVAKILFHF

SQRFTKVVPESCMLILLGLVLGGIVLIANKKQLYQLEPALFFLFLLPTIVGDAGYFMPAR

LFFDNLGAILTYAVVGTLWNAFCTGFCLYAAKVLNVIDAKVQADLMDFLLFGALISAVDP

VAVLAVFEEVHVNDTLFIIVFGESLVNDAVTVVLYKVYISFVQVGAANVQTADYFKGVAS

FLIVSIGGTLVGLVFAVILGFITRFTKKVRIIEPLFVFLLVYLAYLTAELFSLSAILSMT

FCGIGANKYVEANISQKSRTTVKYTMKTLASIAETIIFIFLGISVVDKSKWAWDTGLVSC

TLVFILVFRAIGVVGQTWVLNRFRLVPLDKIDQVMMSYGGLRGAVAFALVVLLDGEQVKA

KDYFVATTIVVVFFTVMFQGLTIKPLVKWLKVPRSTSRKPTINEEIHERAFDHILTAVED

IAGLQGYHHWRDKWEQFDKKYLSKLLLRKSVYTKSELWEAYQKINIRDAISVIDQYQSHY

SRHFMPLGEKERQDREVFQRNMKIRMESSKSTRHKRHKKERSIKKRRGSDSKEESSEKPR

RNVSWHDKDRSDAEKEDDEGITFVAQKVETPKPRPKSVPAALEVSSSPPARSPSPPLSAD

SHLPWKGGVGSPPPCVSVEATKIIPVDLQQAWNQSISSLESISSPPAPPVPDPLHPRISA

LSRLGAPRPVSYTPPSSSSSSTAAGTSFNTSNTQLLHGSFLFPDRRLKEEEEDDSFESSQ

ELQPLMSTLKPPAEAIPPPPHLQAPSSRGKGARRTPACTSGAWCRRPLMAARSRAAEDPP

SFRR

***Zebrafish (Danio rerio)***

>tr|A3KPJ8|A3KPJ8_DANRE_Danio

MAFSTLLLAFLVVSGALHEAAAGLDFYGAEGKQNYSSRSSAEGSASSGNSSHSATITTLP

IVTWKWHHVETPYLVALWILTCWLCKLVTELNHNITSVIPESGLLIILGFILGGIVWGAD

KAQTFKLIPVNFFYYLLPQIILDASYCMPNKLFFGNLGAILVHAIIGTCWNAGTVGIALW

ACYEGGAMGTLNIGCLQFLLFGALMSAVDPVAVIAVFEEVHVNEVLYILVFGESLLNDGV

TVVLYNVFDAFVSLGGPKINAAEIIKGIVSFAVVAFGGSFLGVVFAVLTSMLTRVTKNVQ

IIEAGFIIVLGYLSYLTAEMLSLSAILSLTFCGVCCQKYINANMDEKSVTCLRYSLKVLA

NGSETMIFVFLGVSAIDQTIWVWNTGFILLTLLFIVVFRMMGVFFLNWILNQSRLIPIDL

TDQLVMGYGGLRGAVAYGLAASLDENKIPEKNLMLGTTLIVVYFTVILQGITMKPLVNWL

KVKKATQSDLTLNGKLNNRVFEHTLTGMEDICGRMGDNWWTRHWNHFEEKYVCWLLMTSE

ARKRNDKILDAFHKLNVEDATKYVNEGESKGSLAFIRSYDSASVDFKKKLALEYADIIPD

IMADMSEYDFDIDSVPVTSVMKNPIPSVSLDIHEQERRGMSNDLNAHHLLEQHLYKSRRN

NRATYSRSHYNTSENPDEMQEIFQRTMRNRLESFKSAKMGVNPAKKVSKHPKKDAPHKPN

GKPADPNRDHLYGDEDFEFSADSASSSENTGQFPRRVINTTGAGVDNPAYIPELDTSPSM

RNPPWQTETGHNTAVAPSQRAQGRLPWTPTNLRRLAPLRVSSRSTDSFAMPEPAADEETP

EEKPATHHTRL

>tr|A9XP97|A9XP97_DANRE_Danio

MAFSTLLLAFLVVSGALHEAAAGLDFYGAEGKQNYSSRSSAEGSASSGNSSHSATITTLP

IVTWKWHHVETPYLVALWILTCWICKLVTELNHNITSVIPESGLLIILGFILGGIVWGAD

KAQTFKLIPVNFFFYLLPQIILDASYCMPNKLFFGNLGAILVHAIIGTCWNAGTVGIALW

ACYEGGAMGTLNIGCLQFLLFGALMSAVDPVAVIAVFEEVHVNEVLYILVFGESLLNDGV

TVVLYNVFDAFVSLGGPKINAAEIIKGIVSFAVVAFGGSFLGVVFAVLTSMLTRVTKNVQ

IIEAGFIIVLGYLSYLTAEMLSLSAILSLTFCGVCCQKYINANMDEKSVTCLRYSLKVLA

NGSETMIFVFLGVSAIDQTIWVWNTGFILLTLLFIVVFRMMGVFFLNWILNQSRLIPIDL

TDQLVMGYGGLRGAVAYGLAASLDENKIPEKNLMLGTTLIVVYFTVILQGITMKPLVNWL

KVKKATQSDLTLNGKLNNRVFEHTLTGMEDICGRMGDNWWTRHWNHFEEKYVCWLLMTSE

ARKRNDKILDAFHKLNVEDATKYVNEGESKGSLAFIRSYDSASVDFKKKLALEYADIIPD

IMADMSEYDFDIDSVPVTSVMKNPIPSVSLDIHEQERRGMSNDLNAHHLLEQHLYKSRRN

NRATYSRSHYNTSENPDEMQEIFQRTMGNRLESFKSAKMGVNPAKKVSKHPKKDAPHKPN

GKPADPNRDHLYGDEDFEFSADSASSSENTCQFPRRVINTTGAGVDNPAYIPELDTSPSM

RNPPWQTETGHNTAVAPSQRAQGRLPWTPTNLRRLAPLRVSSRSTDSFAMPEPAADEETP

EEKPATHHTRL

>tr|B7ZVJ6|B7ZVJ6_DANRE_Danio

MASSTYICLLRAAFLVCLVYPLVKGNSVEVSDPHHHQTENLTGLPIVTFKWHHVETPYLV

ALWVFVAGLAKLSVIQLNHSITGVFPESGLLIILGFILGGIVVGADKAQTFKLLPSTFFY

YLLPQIILDASYFMPNKLFFRNLGAILIYAIFGTCWNAAAVGLSLWGCHEAGAMGDLNIG

LLQFLLFGSLIAAVDPVAVIAVFEEVHVNEVLFILVFGESLLNDGVTVVLYNVFDGFVSL

GGPMIDAAEIFKGICSFVVVAFGGSFVGVAFAILLSLLTRCTKHVQIIEAGFVFLVGYLS

YLTAEMLSLSAILSITFCGVCCQKYINANMDERSVTTVRHAMKMLANGSETMIFLFLGVT

AIDTEIWVWNTGFILLTLLFIFIYRIIGVFMFTWMLNKFRLVPVELIDQVVMSYGGLRGA

VAYGLAAMLDESRIPEKKLMISTTLIVVYFTVILQGITMKPLVQWLKVKRAAHSDLKLVE

KLNNRAFDHILAAVEDICGRIGDNWWTRGWNRFEEKYISWLLMKPKARKNRDYLFNIFHK

LNLEDAMNYVSEGERQGSLAFIRNDEAANVDFKKQFDSEFADVMPDIMVDMSNYDFGIEN

ISVATIVKDTIPSVSLDIHGQDGKNVVDDINAHRLLQQHLYRSRKHHHHKYSRSHFSINQ

SEEEVQEIFQRTMRSRLESFKSAKMGVNPTKKITKFHKKDPTDKMNGHTEGNGYLYGDEA

DSGVGADGAISSTFMRDTYKMGAGIENPVFMPDMDSPVHSPVHSPPWLLETEMSVAPSQR

AQRRLPWTPNRRLAPMRISTCSTDSFLMADVSSTLEVHEEQPQSDNGESVFE

>tr|F1QSJ6|F1QSJ6_DANRE_Danio

MGSTSEMKAVAMRVWRRLPVHVFVLASSLCLCQCGAEEDSAMENIVTEKKAEESHRQDSA

DLLIFIMLLTLTILTIWLFKHRRFRFLHETGLAMIYGLLVGVVLRYAIHVPSDINNVTLS

CHVNASPATLLVNVSGKFYEYTLKGEIGANKVNDIQDNEMLRKVTFDPEVFFNILLPPII

FHAGYSLKRRHFFRNLGSILAYAFVGTVVSCFIIGALMYGCVMLMKKIGHLQDDFFFTDC

LFFGAIVSATDPVTVLAIFNELQVDADLYALLFGESVLNDAVAVVLSSSIVAYQPTGDNS

HTFEAMAMLKSFGIFLGVFSGSFALGVATGIVTALVTKFTKLRDFPLLETALFFLMSWST

FLLAEACGFTGVVAVLFCGMTQAHYTFNNLSPESQDRTKQLFELLNFLAENFIFSYMGLT

LFTFQNHVFNPIFIVCAFLAVFLGRAANIYPLSFLLNLGRRNKISSNFQHMMMFAGLRGA

MTFALSIRDTATYARQMMFSTTLLVVFFTVWICGGGTTQMLSCQKIRVGVDTDHENSIGP

DGVERRSTKQESAWLFRIWYNFDHNYLKPILTHSGPPLTATLPACCSPLARCLTSPQAYE

NEGELKNADSDLILNDGDITLTYGDIAVSTDGTGAHSSGVLMGGAAVNSDEALDRELAFG

DHELVIRGTRLVLPMDDSEPPLTLDSHRQRHRNEFMS

>tr|F1R354|F1R354_DANRE_Danio

MGCQRRRSPASGARLLPRLLCLMCFSVVILAENTEMVNVVTERKAEESHRQDSANLLIFI

TLLTITILTIWLFKHRRFRFLHETGLAMIYGLFVGVILRFGVHSLQDTSEVTLSCAVNSS

PTTILVNVSGRFYEYSLKGEVQDSKGHDVPNAEMLRKVTFDPEVFFNILLPPIIFHAGYS

LKRRHFFRNLGSILSYAFVGTMTSCFVTGLAMYSCVVFMKIIGQLGGDFFFTDCLFFGAI

VSATDPVTVLAIFNELKVDVDLYALMFGESVLNDAVAIVLSSSIVAYQPEGDNRHTFDAS

ALLKCLGVFLGVFSGSFAMGAATGVMTALVTKFTKLRDFPLLETALFFLMSWSTFLLAEA

CGFTGVVAVLFCGITQAHYTYNNLSVESKSRTKQLFELLNFLAENFIFSYMGLTLFSFQH

HVFNPFFIIGAFVAIFLGRLANIYPLSFLLNLGRKNKIPSNFQHVMMFSGLRGAMTFALS

IRDTATYARQMMFSTTLLIVFFTVWVCGGGTTPVLSLMQIRVGVDTEQDNSATAADGMTQ

RSTKHESAWPFRLWYTFDHNYLKPFLTHSGPPLTTTLPACCGPIARCLTSSQAYENAGQL

HDEDTELILNDGGGLMYSDMTVSTDVHGVPTTSSSQAGLYSLAGTSDDPLERELAFGGTR

LVLPTDEPLDPLTSFPPPPPSPPLSDPLRHRV

>tr|B0S734|B0S734_DANRE_Danio

MVLHFHVGSSFNFQVFLCICFASVGYGENVDVTTKATAGFPSKSPLWSSNDDTAGTEADK

TNPELFTLNYARIKIPFEITLWVLLASFAKIGFNVYHKITVWVPESCLLISLGFVVGGVM

HAVRKEPPAVISSNVFFLYMLPPIILESGYFMPTRPFFENVGTVMWFAVVGTLWNTMGIG

ISLFAICQIEEFGVQDINLQENLLFAAILSAVEPVAVLSVFEDISINEQLYIVMFGECLF

NDAVTVVLYNLFNHVAAMEVDEPAVIFMDIGGFFVVGLGGIFFGLLFGFLTAFTTRFTGK

VREIEPLVIFMFSYMSYLIAELFALSSIMAILICTLSMKYYVEENVSQRSCTTIRHVIKM

VGSVSETLIFFFLGVVSITTEHEWNWGYILFTLFFTLFWRFLGILVLTQIINPFRKIPFN

FQDQFGLAYGGLRGALSFALAFTLPNSIPHRKLFITDTIAVIIFTVFIQGITIRPLLKLM

NIRKTNRNLETINDEIHKRLMEHTVAGIEDLCGQWSHYRWKDMFMKFNNRFIRKILIRDN

RPESSIVSLYKKLELRNALEILDTMSGDISAAPSLMSLYEERKNASKHKKFLPSDVENMH

ELLSKNMYKIRQRTVSYTNRHSLPNENIAKEILMRRHTTIRSSLRSGSFHATNSANTSHK

FSSLPVGQSLNSGFPPGRFRHTDHIDAESETAYPSNRSTFRHQRRSSSKAGIPLRPLNDL

REDNEPNEEIPYKMSHPRSKFRHSGHQTSRQYYGTSRNCRDTDFEEQQSSLSPHGWTEEP

QNNSDVQHPLMRKPQ

>tr|F1R610|F1R610_DANRE_Danio

MRGLSLLLFITLARDAVTDLYVTGASLPPNLGVSAAPPATSPVAARYIPREYGHPGIALA

RSAEEPTTPSTEDGESSHHHGGGFKIVQWEWSYVQTPYIIASWLLVASVAKILFHFSQRF

TTAVPESCMLILLGLALGTVVLLASKKQPYQLDPGLFFLFLLPTIVGDAGYFMPARLFFD

NLGAILMYAVVGTLWNAFCTGFCLYGVKMAGVIDEKVDAGLMEFLLFGALISAVDPVAVL

AVFEEVHVNETLFIIVFGESLLNDAVTVVLYKVYISFVEVGPGNVNTADYFKGIASFLIV

SIGGTLVGLIFAVILAFFTRFTKKVRIIEPMFIFLLVYLAYLTAELFSLSAILSLTFCGI

GCNKYVEANISQKSRTTVKYTMKTLASIAETIIFIFLGISAVDKSKWAWDTGLVVSTLIF

ILVFRAMGVIGQTWFLNWFRLVPLDKIDQVVMSYGGLRGAVAFALVVLLDKEHVKAKDYF

VATTIVVVFFTVMFQGLTIKPLVKWLKVPRSTNRKPTINEEIHERAFDHILAAVEDIAGL

NGYHHWRDKWEQFDKNYLSKLLLRKSVYHKSELWEAYQKINIRDAISVIDQGGNVLTSAR

VSLPSMASRTSFPEVTNVTNYLRENGSGVCLDLQVIDTVPGAKIEEELETHHVLAGNLYK

PRRRYQSHYSRHFMTLGEKERQDREVFQRNMRNRLESFKSTRYKRHKKDRSQKKRKGSDA

KEQDTSDKPRRNVSWQDKDPLVAPLAPEEDKHDPTEVDKDDDVGIIFVARRTPETPKERP

KSVPAALDVCQSPITAPPSLTCGEKNLPWKSGMGSLPPCVSVEATKIIPVDLQQAWNQSI

SSLESLASPPAPAEPLHPRISALSRLGGPLRTPAKDKSAGVDAGLESTTSSPSSGQFKYP

MGKEEEEEGALSQQHREMQPLMSSLRPPPPPPPLAAQSAGKRRNPCVYLRSLVTVPPTGA

EPHSRKPTQL

>tr|F1Q610|F1Q610_DANRE_Danio

MEAAKAPKPWTLYLPNPSSLFPLFVFILSQCYRISADDGGIEELATEKEAEESHRQDSVN

LLTFILLLTLTILTIWLFKHRRVRFLHETGLAMIYGLLVGVILRYGIPSTSYHNKTPPSC

MLKEGPVSTVLLNVSGKFFEYTLKGEINLREIHSVEQNDMLRKLTFDPEVFFNILLPPII

FHAGYSLKKRHFFRNLGSIITYAFLGTAISCFVIGNLMYGVVKLMQVLGQLTDKFYYTDC

LFFGAIISATDPVTVLAIFNELHADGDLYALLFGESVMNDAVSIVLSSSIVAYQPSGANT

HTFDASAFFKSVGVFIGIFSGSFAMGAVTGVVTALVTKFTKLHCFPLLETALFFLMSWST

FLLAEACGFTGVVAVLFCGITQAHYTYNNLSEESTKRTKQLFEVLHFLAENFIFSYMGLA

LFTFQNHVFSPIFIVGAFLAIFIGRALNIYPLSFLINLGRRHKIRGNFQHMMMFAGLRGA

MAFALAIRDTATYARQMMFTTTLLIVFFTVWVFGGGTTPMLSWLHIRVGESSDAEKERMR

RLWNYDSRVGVDPDQDLQPCGDSFQVLQGDGSQSEGQSKTKQESAWLFRLWYTFDHNYLK

PILTHSGPPLTSTLPACCGPLARCLTSPQAYETHEPLRDNDSDLILNEGDLTLTYGDTAI

TANGASSSSGPEGSAGIWGLNGKRSDSTSEDALERELDTREHELLSRGTRLVFPIEDHA

>tr|A9XP99|A9XP99_DANRE_Danio

MPVVALFFLPLLIGSCRGIDHNETDQRVFPVISINYQHVQMPFEISLWILLASILKLGFH

LVPHVSAVVPESCVLIVVGLLVGGLIRVLGEDPPVLDSQLFFLCLLPPIILDAGYFLPIR

PFMENVGTILTFAVLGTLWSAFFMGVALYGACQLEGAELAGVDLLSCLLFGSIVSAVDPV

AVLAVFQEIHINELLHILVFGESLLNDAVTVVLYHLFEEFVHRGEVTAGDVFLGVVCFFV

VSVGGVLVGCVYGFLAAFTSRFTAHTRVIEPLFVFLYSYMAYLSAEVFHLSGIMALIACG

VVMRPYVEANISHKSYTTIKYFLKMWSSVSETLIFIFLGVSTVAGPHAWNWTFVSLTLVL

CLLSRVLGVVGLTFIINKYRMVKLSGKDQFIVAYGGLRGAIAFSLAFLLSSESCAMKDLF

LTAVITVIFFTVFVQGMTIRPLVDLLAVKKKKQSKFSVNEEIHTQFLEHLLTGIEDVCGH

YGHHHWRDKISRFNKRYMMRFLIGGERSQEPQLISFYKQMELEQTMKMLENNSTTSLSSA

ATQLHQRSKRTLNISKEREAEIRKMLRANLQKTRQRLRSYSRHTLLPDAEEDDWSETRLR

KQRMEMERRMSHFLTVPANRPESPRRARLQTDLDGRAQKASVTFAEELEEHRD

>tr|A9XPA0|A9XPA0_DANRE_Danio

MDDTAGTEADKTNPELFTLNYARIKIPFEITLWVLLASFAKIGFNVYHKITVWVPESCLL

ISLGFVVGGVMHAVRKEPPAVISSNVFFLYMLPPIILESGYFMPTRPFFENVGTVMWFAV

VGTLWNTMGIGISLFAICQIEAFGVQDINLQENLLFAAILSAVEPVAVLSVFEDISINEQ

LYIVMFGECLFNDAVTVVLYNLFNHVAAMEVDEPAVIFMDIGGFFVVGLGGIFFGLLFGF

LTAFTTRFTGKVREIEPLVIFMFSYMSYLIAELFALSSIMAILICTLSMKYYVEENVSQR

SCTTIRHVIKMVGSVSETLIFFFLGVVSITTEHEWNWGYILFTLFFTLFWRFLGILVLTQ

IINPFRKIPFNFQDQFGLAYGGLRGALSFALAFTLPNSIPHRKLFITDTIAVIIFTVFIQ

GITIRPLLKLMNIRKTNRNLETINDEIHKRLMEHTVAGIEDLCGQWSHYRWKDMFMKFNN

RFIRKILIRDNRPESSIVSLYKKLELRNALEILDTMSGDISAAPSLMSLYEERKNASKHK

KFLPSDVENMHELLSKNMYKIRQRTVSYTNRHSLPNENIAKEILMRRHTTIRSSLRSGSF

HATNSANTSHKFCSLPVGQSLNSGFPPGRFRHTDHIDAESETAYPSNRSTFRHQRRSSSK

AGIPLRPLNDLREDNEPNEEMPNKMSHPRSKFRHSGHQTSRQYYGTSRNCRDTDFEEQQS

SLSPHGWTEEPQNNSDVQHPLMRKPQ

>tr|A9XPA2|A9XPA2_DANRE_Danio

MRGLSLLLFITLARDAVTDLYVTGASLPPKLGVSAAPPATSPVAARYIPREYGHPGIALA

RSAEEPTTPSTEDGESSHHHGGGFKIVQWEWSYVQTPYIIASWLLVASVAKILFHFSQRF

TTAVPESCMLILLGLALGTVVLLASKKQPYQLDPGLFFLFLLPTIVGDAGYFMPARLFFD

NLGAILMYAVVGTLWNAFCTGFCLYGVKMAGVIDEKVDAGLMEFLLFGALISAVDPVAVL

AVFEEVHVNETLFIIVFGESLLNDAVTVVLYKVYISFVEVGPGNVNTADYFKGIASFLIV

SIGGTLVGLIFAVILAFFTRFTKKVRIIEPMFIFLLVYLAYLTAELFSLSAILSLTFCGI

GCNKYVEANISQKSRTTVKYTMKTLASIAETIIFIFLGISAVDKSKWAWDTGLVVSTLIF

ILVFRAMGVIGQTWFLNWFRLVPLDKIDQVVMSYGGLRGAVAFALVVLLDKEHVKAKDYF

VATTIVVVFFTVMFQGLTIKPLVKWLKVPRSTNRKPTINEEIHERAFDHILAAVEDIAGL

NGYHHWRDKWEQFDKNYLSKLLLRKSVYHKSELWEAYQKINIRDAISVIDQGGNVLTSAR

VSLPSMASRTSFPEVTNVTNYLRENGSGVCLDLQVIDTVPGAKIEEELETHHVLAGNLYK

PRRRYQSHYSRHFMTLGEKERQDREVFQRNMRNRLESFKSTRYKRHKKDRSQKKRKGSDA

KEQDTSDEPRRNVSWQDKDPLVAPLATEEDKHDPTEVDKDDDVGIIFVARRTPETPKERP

KSVPAALDVCQSPITAPPSLTCGEKNLPWKSGMGSLPPCVSVEATKIIPVDLQQAWNQSI

SSLESLASPPAPAEPLHPRISALSRLGGPLRTPAKDKSAGVDAGLESTTSSPSSGQFKYP

MGKEEEEEGALSQQHREMQPLMSSLRPPPPPPPLAAQSAGKRRNPCVYLRSLVTVPPTGA

EPHSRKPTQL

>tr|F1QHS0|F1QHS0_DANRE_Danio

XICIILVHLLIKFKLHFLPESVAVVSLGILMGAFIKIIESQQLANWKEEEMFRPNMFFLL

LLPPIIFESGYSLHKGNFFQNIGSITLFAVFGTAISAFIVGGGIYFLGQADVIYKMTMTD

SFAFGSLISAVDPVATIAIFNALNVDPVLNMLVFGESILNDAVSIVLTNTAEGFGGSDVS

TGWETFMQALGYFLKMFFGSAALGTLTGLISAISLKHFDLRKTPSLEFGMMIIFAYLPYG

LAEGIKLSGIMAILFSGIVMSHYTHHNLSPVTQILMQQTLRTVAFMCETCVFAFLGLSIF

SFPHKFELSFVIWCIVLVLVGRAVNIFPLSFLLNFFRDHKITPKMMFIMWFSGLRGAIPY

ALSLHLGLEPIEKRQLIGTTTIVIVLFTILLMGGGTMPLIRIMDIEESQSRRKSKKDINL

SKTEKMGNAIESEHLSELTEEEYEAQIYQRQDLKGFLWLDAKYLNPFFTRRLTQEDLLHG

RIQMKTLTNKWYEEVRQGPSGSEDEEDEAELL

>tr|F1QR72|F1QR72_DANRE_Danio

GFHLIPRISSIVPESCLLIVVGLLIGGLMKLVGEVPPVLRSDIFFLYLLPPIILDAGYFL

PIRPFTENLGTILMFAVVGTLWNSFFIGGLLYGVCQLEGVHLVHVDLLSCLLFGSIISAV

DPVAVLAVFEEIHINELLHILVFGESLLNDAVTVVLYHLFEEYSSVGTITITDVFLGIVC

FLVVSLGGIVVGAIYGILAAFTSRFTSHTRVIEPLFVFVYSYMAYLSAEVFHLSGIMALI

ACGAVMRPYVEANISHKSHTTTKYFLKMWSSVSDTLIFIFLGVSTVAGPHQWNWTFVIVT

LILCLVARVLGKIYINIYIHRFHLVYSGGSDRFLPLCIRLIIIFFISFILSNLLCLINII

YLILVITKILFRIRVRGMTIRPLVDLLAVKKKQETKRSINEEIHTQFLDHLLTGIEDICG

HYGHHHWKD

>tr|A9XPA3|A9XPA3_DANRE_Danio

MGCQRRRSPASGARLLPRLLCLMCFSVVILAENTEMVNVVTERKAEESHRQDSANLLIFI

TLLTITILTIWLFKHRRFRFLHETGLAMIYGLFVGVILRFGVHSPQDTSEVTLSCAVNSS

PTTILVNVSGRFYEYSLKGEVQDSKGHDVPNAEMLRKVTFDPEVFFNILLPPIIFHAGYS

LKRRHFFRNLGSILSYAFVGTMTSCFVTGLAMYSCVVFMKIIGQLGGDFFFTDCLFFGAI

VSATDPVTVLAIFNELKVDVDLYALMFGESVLNDAVAIVLSSSIVAYQPEGDNRHTFDAS

ALLKCLGVFLGVFSGSFAMGAATGVMTALVTKFTKLRDFPLLETALFFLMSWSTFLLAEA

CGFTGVVAVLFCGITQAHYTYNNLSVESKSRTKQLFELLNFLAENFIFSYMGLTLFSFQH

HVFNPFFIIGAFVAIFLGRLANIYPLSFLLNLGRKNKIPSNFQHVMMFSGLRGAMTFALS

IRDTATYARQMMFSTTLLIVFFTVWVCGGGTTPVLSLMQIRVGVDTEQDNSATAADGMTQ

RSTKHESAWPFRLWYTFDHNYLKPFLTHSGPPLTTTLPACCGPIARCLTSSQAYENAGQL

HDEDTELILNDGGDLMYSDMTVSTDVHGVPTTSSSQAGLYSLAGTSDDPLERELAFGGTR

LVLPTDEPLDPLTSFPPPPPSPPLSDPLRHRV

>tr|A9XPA1|A9XPA1_DANRE_Danio

MASSTYICLLRAAFLVCLVYPLVKGNSVEVPNPHHHQTENLTGLPIVTFKWHHVETPYLV

ALWVFVAGLAKLSVIQLNHSITGVFPESGLLIILGFILGGIVVGADKAQTFKLLPSTFFY

YLLPQIILDASYFMPNKLFFRNLGAILIYAIFGTCWNAAAVGLSLWGCHEAGAMGDLNIG

LLQFLLFGGLIAAVDPVAVIAVFEEVHVNEVLFILVFGESLLNDGVTVVLYNVFDGFVSL

GGPMIDAAEIFKGICSFVVVAFGGSFVGVAFAILLSLLTRCTKHVQIIEAGFVFLVGYLS

YLTVEMLSLSAILSTTFCGVCCQKYINANMDERSVTTVRHAMKMLANGSETMIFLFLGVT

AIDTEIWVWNTGFILLTLLFIFIYRIIGVFMFTWMLNKFRLVPVELIDQVVMSYGGLRGA

VAYGLAAMLDESTIPEKKLMISTTLIVVYFTVILQGTTMKPLVQWLKVKRAAHSDLKLVE

KLNNRAFDHILAAVEDICGRIGDNWWTRGWNRFEEKYISWLLMKPKARKNRDYLFNIFHK

LNLEDAMNYVSEGERQGSLAFIRNDEAANVDFKKQFDSEFADVMPDIMVDMSNYDFGIEN

ISAAAIVKDTIPSVSLDIHGQDGKNVVDDINAHRLLQQHLYRSRKHHHHKYSRSHFSINQ

SEEEVQEIFQRTMRSRLESFKSAKMGVNPTKKITKFHKKDPTDKMNGHTEGNGYLYGDEA

DSGVGADGAISSTFMRDTYKMGAGIENPVFMPDMDSPVHSPVHSPPWLLETEMSVAPSQR

AQRRLPWTPNRRLAPMRISTCSTDSFLMADVSSTLEVHEEQPQSDNGESVFE

>tr|A9XPA5|A9XPA5_DANRE_Danio

MSMMGVSNRAVVVCVCLIILTANSAPFARADDQIYSKAEGEAVNNNNNNADSDAQAQNPA

DGHAEQLTRLNETDTSSNDTAQTTTPAPKPATPEPPKNEPILPVQTGVKAQEEEQSSGMT

IFFSLLVIGICIILVHLLIKFKLHFLPESVAVVSLGILMGAFIKIIESQQLANWKEEEMF

RPNMFFLLLLPPIIFESGYSLHKGNFFQNIGSITLFAVFGTAISAFIVGGGIYFLGQADV

IYKMTMTDSFAFGSLISAVDPVATIAIFNALNVDPVLNMLVFGESILNDAVSIVLTNTAE

GFGGSDVSTGWETFMQALGYFLKMFFGSAALGTLTGLISAISLKHFDLRKTPSLEFGMMI

IFAYLPYGLAEGIKLSGIMAILFSGIVMSHYTHHNLSPVTQILMQQTLRTVAFMCETCVF

AFLGLSIFSFPHKFELSFVIWCIVLVLVGRAVNIFPLSFLLNFFRDHKITPKMMFIMWFS

GLRGAIPYALSLHLGLEPIEKRQLIGTTTIVIVLFTILLMGGGTMPLIRIMDIEESQSRR

KSKKDINLSKTEKMGNAIESEHLSELTEEEYEAQIYQRQDLKGFLWLDAKYLNPFFTRRL

TQEDLLHGRIQMKTLTNKWYEEVRQGPSGSEDEEDEAELL

>tr|Q5PR36|Q5PR36_DANRE_Danio

MGVSNRAVVVCVCLIILTANSAPFARADDQIYSKAEGEAVNNNNNNADSDAQAQNPADGH

AEQLTRLNETDTSSNDTAQTTTPAPKPATPEPPKNEPILPVQTGVKAQEEEQSSGMTIFF

SLLVIGICIILVHLLIKFKLHFLPESVAVVSLGILMGAFIKIIESQQLANWKEEEMFRPN

MFFLLLLPPIIFESGYSLHKGNFFQNIGSITLFAVFGTAISAFIVGGGIYFLGQADVIYK

MTMTDSFAFGSLISAVDPVATIAIFNALNVDPVLNMLVFGESILNDAVSIVLTNTAEGFG

GSDVSTGWETFMQALGYFLKMFFGSAALGTLTGPISAISLKHFDLRKTPSLEFGMMIIFA

YLPYGLAEGIKLSGIMAILFSGIVMSHYTHHNLSPVTQILMQQTLRTVAFMCETCVFAFL

GLSIFSFPHKFELSFVIWCIVLVLVGRAVNIFPLSFLLNFFRDHKITPKMMFIMWFSGLR

GAIPYALSLHLGLEPIEKRQLIGTTTIVIVLFTILLMGGGTMPLIRIMDIEESQSRRKSK

KDINLSKTEKMGNAIESEHLSELTEEEYEAQIYQRQDLKGFLWLDAKYLNPFFTRRLTQE

DLLHGRIQMKTLTNKWYEEVRQGPSGSEDEEDEAELL

>tr|A0A0R4IF22|A0A0R4IF22_DANRE_Danio

MKQDVFLCICFASVGYGENVDVTTKATAGFPSKSPLWSSNDDTAGTEADKTNPELFTLNY

ARIKIPFEITLWVLLASFAKIGFNVYHKITVWVPESCLLISLGFVVGGVMHAVRKEPPAV

ISSNVFFLYMLPPIILESGYFMPTRPFFENVGTVMWFAVVGTLWNTMGIGISLFAICQIE

EFGVQDINLQENLLFAAILSAVEPVAVLSVFEDISINEQLYIVMFGECLFNDAVTVVLYN

LFNHVAAMEVDEPAVIFMDIGGFFVVGLGGIFFGLLFGFLTAFTTRFTGKVREIEPLVIF

MFSYMSYLIAELFALSSIMAILICTLSMKYYVEENVSQRSCTTIRHVIKMVGSVSETLIF

FFLGVVSITTEHEWNWGYILFTLFFTLFWRFLGILVLTQIINPFRKIPFNFQDQFGLAYG

GLRGALSFALAFTLPNSIPHRKLFITDTIAVIIFTVFIQGITIRPLLKLMNIRKTNRNLE

TINDEIHKRLMEHTVAGIEDLCGQWSHYRWKDMFMKFNNRFIRKILIRDNRPESSIVSLY

KKLELRNALEILDTMSGDISAAPSLMSLYEERKNASKHKKFLPSDVENMHELLSKNMYKI

RQRTVSYTNRHSLPNENIAKEILMRRHTTIRSSLRSGSFHATNSANTSHKFSSLPVGQSL

NSGFPPGRFRHTDHIDAESETAYPSNRSTFRHQRRSSSKAGIPLRPLNDLREDNEPNEEI

PYKMSHPRSKFRHSGHQTSRQYYGTSRNCRDTDFEEQQSSLSPHGWTEEPQNNSDVQHPL

MRKPQ

>tr|F8W586|F8W586_DANRE_Danio

MGCQRRRSPASGARLLPRLLCLMCFSVVILAENTEMVNVVTERKAEESHRQDSANLLIFI

TLLTITILTIWLFKHRRFRFLHETGLAMIYGLFVGVILRFGVHSLQDTSEVTLSCAVNSS

PTTILVNVSGRFYEYSLKGEVQDSKGHDVPNAEMLRKVTFDPEVFFNILLPPIIFHAGYS

LKRRHFFRNLGSILSYAFVGTMTSCFVTGLAMYSCVVFMKIIGQLGGDFFFTDCLFFGAI

VSATDPVTVLAIFNELKVDVDLYALMFGESVLNDAVAIVLSSSIVAYQPEGDNRHTFDAS

ALLKCLGVFLGVFSGSFAMGAATGVMTALVTKFTKLRDFPLLETALFFLMSWSTFLLAEA

CGFTGVVAVLFCGITQAHYTYNNLSVESKSRTKQLFELLNFLAENFIFSYMGLTLFSFQH

HVFNPFFIIGAFVAIFLGRLANIYPLSFLLNLGRKNKIPSNFQHVMMFSGLRGAMTFALS

IRDTATYARQMMFSTTLLIVFFTVWVCGGGTTPVLSLMQIRVGVDTEQDNSATAADGMTQ

RSTKHESAWPFRLWYTFDH

>tr|A0A0R4IT04|A0A0R4IT04_DANRE_Danio

MSMMGVNNRAVVVCVCLIILTANSAPFARADDQIYSKAEGEAVNNNNNNADSDAQAQNPA

DGHAEQLTRLNETDTSSNDTAQTTTPAPKPATPEPPKNEPILPVQTGVKAQEEEQSSGMT

IFFSLLVIGICIILVHLLIKFKLHFLPESVAVVSLGILMGAFIKIIESQQLANWKEEEMF

RPNMFFLLLLPPIIFESGYSLHKGNFFQNIGSITLFAVFGTAISAFIVGGGIYFLGQADV

IYKMTMTDSFAFGSLISAVDPVATIAIFNALNVDPVLNMLVFGESILNDAVSIVLTNTAE

GFGGSDVSTGWETFMQALGYFLKMFFGSAALGTLTGLISAISLKHFDLRKTPSLEFGMMI

IFAYLPYGLAEGIKLSGIMAILFSGIVMSHYTHHNLSPVTQILMQQTLRTVAFMCETCVF

AFLGLSIFSFPHKFELSFVIWCIVLVLVGRAVNIFPLSFLLNFFRDHKITPKMMFIMWFS

GLRGAIPYALSLHLGLEPIEKRQLIGTTTIVIVLFTILLMGGGTMPLIRIMDIEESQSRR

KSKKDINLSKTEKMGNAIESEHLSELTEEEYEAQIYQRQDLKGFLWLDAKYLNPFFTRRL

TQEDLLHGRIQMKTLTNKWYEEVRQGPSGSEDEEDEAELL

>tr|A0A0R4IX40|A0A0R4IX40_DANRE_Danio

MGSTSEMKAVAMRVWRRLPVHVFVLASSLCLCQCGAEEDSAMENIVTEKKAEESHRQDSA

DLLIFIMLLTLTILTIWLFKHRRFRFLHETGLAMIYGLLVGVVLRYAIHVPSDINNVTLS

CHVNASPATLLVNVSGKFYEYTLKGEIGANKVNDIQDNEMLRKVTFDPEVFFNILLPPII

FHAGYSLKRRHFFRNLGSILAYAFVGTVVSCFIIGALMYGCVMLMKKIGHLQDDFFFTDC

LFFGAIVSATDPVTVLAIFNELQVDADLYALLFGESVLNDAVAVVLSSSIVAYQPTGDNS

HTFEAMAMLKSFGIFLGVFSGSFALGVATGILFSLTLSHVTKFTKLRDFPLLETALFFLM

SWSTFLLAEACGFTGVVAVLFCGMTQAHYTFNNLSPESQDRTKQLFELLNFLAENFIFSY

MGLTLFTFQNHVFNPIFIVCAFLAVFLGRAANIYPLSFLLNLGRRNKISSNFQHMMMFAG

LRGAMTFALSIRDTATYARQMMFSTTLLVVFFTVWICGGGTTQMLSCQKIRVGVDTDHEN

SIGPDGVERRSTKQESAWLFRIWYNFDHNYLKPILTHSGPPLTATLPACCSPLARCLTSP

QAYENEGELKNADSDLILNDGDITLTYGDIAVSTDGTGAHSSGVLMGGAAVNSDEALDRE

LAFGDHELVIRGTRLVLPMDDSEPPLTLDSHRQRHRNEFMS

>tr|B7ZV95|B7ZV95_DANRE_Danio

MDDTAGTEADKTNPELFTLNYARIKIPFEITLWVLLASFAKIGFNVYHKITVWVPESCLL

ISLGFVVGGVMHAVRKEPPAVISSNVFFLYMLPPIILESGYFMPTRPFFENVGTVMWFAV

VGTLWNTMGIGISLFAICQIKAFGVQDINLQENLLFAAILSAVEPVAVLSVFEDISINEQ

LYIVMFGECLFNDAVTVVLYNLFNHVAAMEVDEPAVIFMDIGGFFVVGLGGIFFGLLFGF

LTAFTTRFTGKVREIEPLVIFMFSYMSYLIAELFALSSIMAILICTLSMKYYVEENVSQR

SCTTIRHVIKMVGSVSETLIFFFLGVVSITTEHEWNWGYIIFTLFFTLFWRFLGILVLTQ

IINPFRKIPFNFQDQFGLAYGGLRGALSFALAFTLPNSIPHRKLFITDTIAVIIFTVFIQ

GITMRPLLKLMNIRKTNRNLETINDEIHKRLMEHTVAGIEDLCGQWSHYRWKDMFMKFNN

RFIRKILIRDNRPESSIVSLYKKLELRNALEILDTMSGDISAAPSLMSLYEERKNASKHK

KFLPSDVENMHELLSKNMYKIRQRTVSYTNRHSLPNENIAKEILMRRHTTIRSSLRSGSF

HATNSANTSHKFSSLPVGQSLNSGFPPGRFRHTDHIDAESETAYPSNRSTFRHQRRSSSK

AVIPLRPLNDLREDNEPNEEIPYKISHPRSKFRHSGHQTSRQYYGTSRNRRDTDFEEQQS

SLSPHGWTEEPQNNSDVQHPLMRKPQ

>tr|A4QP93|A4QP93_DANRE_Danio

MGSTSEMKAVAMRVWRRLPVHVFVLASSLCLCQCGAEEDSAMENIVTEKKAEESHRQDSA

DLLIFIMLLTLTILTIWLFKHRRFRFLHETGLAMIYGLLVGVVLRYAIHVPSDINNVTLS

CHVNASPATLLVNVSGKFYEYTLKGEIGANKVNDIQDNEMLRKVTFDPEVFFNILLPPII

FHAGYSLKRRHFFRNLGSILAYAFVGTVVSCFIIGALMYGCVMLMKKIGHLQDDFFFTDC

LFFGAIVSATDPVTVLAIFNELQVDADLYALLFGESVLNDAVAVVLSSPIVAYQPTGDNS

HTFEAMAMLKSFGIFLGVFSGSFALGVATGIVTALVTKFTKLRDFPLLETALFFLMSWST

FLLAEACGFTGVVAVLFCGMTQAHYTFNNLSPESQDRTKQLFELLNFLAENFIFSYMGLT

LFTFQNHVFNPIFIVCAFLAVFLGRAANIYPLSFLLNLGRRNKISSNFQHMMMFAGLRGA

MTFALSIRDTATYARQMMFSTTLLVVFFTVWICGGGTTQMLSCQKIRVGVDTDHENSIGP

DGVERRSTEQESAWLFRIWYNFDHNYLKPILTHSGPPLTATLPACCSPLARCLTSPQAYE

NEGELKNADSDLILNDGDITLTYGDIAVSTDGTGAHSSGVLMGGAAVNSDEALDRELAFG

DHELVIRGTRLVLPMDDSEPPLTLDSHRQRHRNEFMS

>tr|Q4V9A5|Q4V9A5_DANRE_Danio

MEAAKAPKPWTLYLPNHSSLFPLFVFILSQCYRISADDGGIEELATEKEAEESHRQDSVN

LLTFILLLTLTILTIWLFKHRRVRFLHETGLAMIYGLLVGVILRYGIPSTSYHNKTPPSC

MLKEGPVSTVLLNVSGKFFEYTLKGEINLREIHSVEQNDMLRKLTFDPEVFFNILLPPII

FHAGYSLKKRHFFRNLGSIITYAFLGTAISCFVIGNLMYGVVKLMQVLGQLTDKFYYTDC

LFFGAIISATDPVTVLAIFNELHADGDLYALLFGESVMNDAVSIVLSSSIVAYQPSGANT

HTFDASAFFKSVGVFIGIFSGSFAMGAVTGVVTALVTKFTKLHCFPLLETALFFLMSWST

FLLAEACGFTGVVAVLFCGITQAHYTYNNLSEESTKRTKQLFEVLHFLAENFIFSYMGLA

LFTFQNHVFSPIFIVGAFLAIFIGRALNIYPLSFLINLGRRHKIRGNFQHMMMFAGLRGA

MAFALAIRDTATYARQMMFTTTLLIVFFTVWVFGGGTTPMPSWLHIRVGESSDAEKERMR

RLWNYDRVGVDPDQDLQPCGDSFQVLQGDGSQLEGQSKTKQESAWLFRLWYTFDHNYLKP

ILTHSGPPLTSTLPACCGPLARCLTSPQAYETHEPLRDNDSDLILNEGDLTLTYGDTAIT

ANGASSSSGPEGSAGIWGLNGKRSDSTSEVALERELDTREHELLSRGTRLVFPIEDHA

>tr|A9XPA4|A9XPA4_DANRE_Danio

MEAAKAPKPWTLYLPNPSSLFPLFVFILSQCYRISADDGGIEELATEKEAEESHRQDSVN

LLTFILLLTLTILTIWLFKHRRVRFLHETGLAMIYGLLVGVILRYGIPSTSYHNKTPPSC

MLKEGPVSTVLLNVSGKFFEYTLKGEINLREIHSVEQNDMLRKLTFDPEVFFNILLPPII

FHAGYSLKKRHFFRNLGSIITYAFLGTAISCFVIGNLMYGVVKLMQVLGQLTDKFYYTDC

LFFGAIISATDPVTVLAIFNELHADGDLYALLFGESVMNDAVSIVLSSSIVAYQPSGANT

HTFDASAFFKSVGVFIGIFSGSFAMGAVTGVVTALVTKFTKLHCFPLLETALFFLMSWST

FLLAEACGFTGVVAVLFCGITQAHYTYNNLSEESTKRTKQLFEVLHFLAENFIFSYMGLA

LFTFQNHVFSPIFIVGAFLAIFIGRALNIYPLSFLINLGRRHKIRGNFQHMMMFAGLRGA

MAFALAIRDTATYARQMMFTTTLLIVFFTVWVFGGGTTPMLSWLHIRVGESSDAEKERMR

RLWNYDRVGVDPDQDLQPCGGSFQVLQGDGSQSEGQSKTKQESAWLFRLWYTFDHNYLKP

ILTHSGPPLTSTLPACCGPLARCLTSPQAYETHEPLRDNDSDLILNEGDLTLTYGDTAIT

ANGASSSSGPEGSAGIWGLNGKRSDSTSEDALERELDTREHELLSRGTRLVFPIEDHA

>tr|F1QI95|F1QI95_DANRE_Danio

MEAAKAPKPWTLYLPNPSSLFPLFVFILSQCYRISADDGGIEELATEKEAEESHRQDSVN

LLTFILLLTLTILTIWLFKHRRVRFLHETGLAMIYGLLVGVILRYGIPSTSYHNKTPPSC

MLKEGPVSTVLLNVSGKFFEYTLKGEINLREIHSVEQNDMLRKLTFDPEVFFNILLPPII

FHAGYSLKKRHFFRNLGSIITYAFLGTAISCFVIGNLMYGVVKLMQVLGQLTDKFYYTDC

LFFGAIISATDPVTVLAIFNELHADGDLYALLFGESVMNDAVSIVLSSSIVAYQPSGANT

HTFDASAFFKSVGVFIGIFSGSFAMGAVTGVVTALVTKFTKLHCFPLLETALFFLMSWST

FLLAEACGFTGVVAVLFCGITQAHYTYNNLSEESTKRTKQLFEVLHFLAENFIFSYMGLA

LFTFQNHVFSPIFIVGAFLAIFIGRALNIYPLSFLINLGRRHKIRGNFQHMMMFAGLRGA

MAFALAIRDTATYARQMMFTTTLLIVFFTVWVFGGGTTPMLSWLHIRVGESSDAEKERMR

RLWNYDRVGVDPDQDLQPCGDSFQVLQGDGSQSEGQSKTKQESAWLFRLWYTFDHNYLKP

ILTHSGPPLTSTLPACCGPLARCLTSPQAYETHEPLRDNDSDLILNEGDLTLTYGDTAIT

ANGASSSSGPEGSAGIWGLNGKRSDSTSEDALERELDTREHELLSRGTRLVFPIEDHA

>tr|A0A0R4IZF2|A0A0R4IZF2_DANRE_Danio

MEAAKAPKPWTLYLPNPSSLFPLFVFILSQCYRISADDGGIEELATEKEAEESHRQDSVN

LLTFILLLTLTILTIWLFKHRRVRFLHETGLAMIYGLLVGVILRYGIPSTSYHNKTPPSC

MLKEGPVSTVLLNVSGKFFEYTLKGEINLREIHSVEQNDMLRKLTFDPEVFFNILLPPII

FHAGYSLKKRHFFRNLGSIITYAFLGTAISCFVIGNLMYGVVKLMQVLGQLTDKFYYTDC

LFFGAIISATDPVTVLAIFNELHADGDLYALLFGESVMNDAVSIVLSSSIVAYQPSGANT

HTFDASAFFKSVGVFIGIFSGSFAMGAVTGVVTALISFCILLHCFPLLETALFFLMSWST

FLLAEACGFTGVVAVLFCGITQAHYTYNNLSEESTKRTKQLFEVLHFLAENFIFSYMGLA

LFTFQNHVFSPIFIVGAFLAIFIGRALNIYPLSFLINLGRRHKIRGNFQHMMMFAGLRGA

MAFALAIRDTATYARQMMFTTTLLIVFFTVWVFGGGTTPMLSWLHIRVGESSDAEKERMR

RLWNYDRVGVDPDQDLQPCGDSFQVLQGDGSQSEGQSKTKQESAWLFRLWYTFDHNYLKP

ILTHSGPPLTSTLPACCGPLARCLTSPQAYETHEPLRDNDSDLILNEGDLTLTYGDTAIT

ANGASSSSGPEGSAGIWGLNGKRSDSTSEDALERELDTREHELLSRGTRLVFPIEDHA

>tr|A0A0G2KZI8|A0A0G2KZI8_DANRE_Danio

MKLGFHLIPRISSIVPESCLLIVVGLLIGGLMKLVGEVPPVLRSDIFFLYLLPPIILDAG

YFLPIRPFTENLGTILMFAVVGTLWNSFFIGGLLYGVCQLEGVHLVHVDLLSCLLFGSII

SAVDPVAVLAVFEEIHINELLHILVFGESLLNDAVTVVLYHLFEEYSSVGTITITDVFLG

IVCFLVVSLGGIVVGAIYGILAAFTSRFTSHTRVIEPLFVFVYSYMAYLSAEVFHLSGIM

ALIACGAVMRPYVEANISHKSHTTTKYFLKMWSSVSDTLIFIFLGVSTVAGPHQWNWTFV

IVTLILCLVARVLGKIYINIYIHRFHLVYSGGSDRFLPLCIRLIIIFFISFILSNLLCLI

NIIYLILVITKILFRIRVRGMTIRPLVDLLAVKKKQETKRSINEEIHTQFLDHLLTGIED

ICGHYGHHHWKDKLNRFNKKYVKKCLIAGERSKEPQLIAFYHKMEMKQAIELVESGGGAK

LPSALPLVHKTADRALPQLSKAREEEIRKILRTNLQKTRMRLRSYNRHTLVADPYEDGWN

DIIKRKKMIELEKKMNNYLTVPAPPPDSP

***Three spine stickleback (Gasterosteus aculeatus),***

>tr|G3NDF8|G3NDF8_GASAC_Gasterosteus

MRGIARNVLLFSFLLLLANKSVTSEGDDVGNREAPPGDPRQTVTGHAKGDGPGSVDQKKD

GQSDGKSNGTEASSNETLHSTKPVVKPTTPAPPTAKPILPVQTGVEAQEEEQSSGLTIFF

SLLVIGICIILVHLLIKFKLHFLPESVAVVSLGILMGGFIKIIEFQELANWKEEEMFRPN

MFFLLLLPPIIFESGYSLHKYVQGNFFQNIGSITLFAVIGTAISAFIVGGGIYFLGQADV

IYKMSMTDSFAFGSLISAVDPVATIAIFNALNVDPVLNMLVFGESILNDAVSIVLTNTAE

GFSRSDNSATGWQTFVQALAYFLKMFFGSAALGTLTGLISALFLKHFDLRKTPSLEFGMM

IIFAYLPYGLAEGLKLSGIMSILFAGIVMSHYTHHNLSPVTQILMQQTLRTMAFMCETCV

FAFLGLSIFSFPHKFELSFVIWCIVLVLLGRAVNIFPLTVLLNVFRDHKITPKMMFIMWF

SGLRGAIPYALSLHLGLEPIEKRQLIGTTTIIIVLFTILLMGGGTMPLIRLMDIEESQSR

RKSKKDITLSKTEKMGNTIESEHLSELTEEEYEAHIFQRRDLKGFMWLDAKYLNPFFTRR

LTQEDLLHGRIQMKTLTNKWYEEVRHGPSGSEDDDDEAELL

>tr|G3PYY9|G3PYY9_GASAC_Gasterosteus

TLRHFNRRTAEMVSKVTVTSARRATRTTWLLVLVSLSVSICVCRASSPQEEEDSAMENIV

TEKKAEESHRQDSVDLLIFIMLLTLTILTIWLFKHRRFRFLHETGLAMIYGVLVGVVLRY

GIHVPRDISNVTMSCHVNASPATLLVNVSGKFYEYTLKGEIGANEVNDIQDNEMLRKVTF

DPEVFFNILLPPIIFHAGYSLKRRHFFRNMGSILAYAFMGTVVSCFVIGLLMYGCVLLMK

QVGQLAEDFFFTDCLFFGAIVSATDPVTVLAIFNELHVDVDLYALLFGESVLNDAVALVL

SSSIVAYQPQGDNSHTFEVTAVLKSFGMFLGVFSGSFALGVATGVVTALYVTKFTKLRDF

PLLETALFFLMSWSTFLLAEACGFTGVVAVLFCGITQAHYTFNNLSPDSQDRTKQLFELL

NFLAENFIFSYMGLTLFTFQSHVFNPMFIVGAFLAVFLGRAANIYPLSFLLNFGRRNKIK

SNFQHMMMFAGLRGAMTFALSIRDTATYARQMMFSTTLLVVFFTVWVCGGGTTQMLSCQN

IRVGVDSDQDNSVSITDGSERRSTKHESAWLFRIWYNFDHNYLKPILTHSGPPLTATLPP

CCAPFARFLTSPQAYENECQLKDDDSDLILTDGDINLAYGDITVSTDASGSHASDDPDRE

LTYGDNELVMRGTRLVLPMDDSDPP

>tr|G3PK51|G3PK51_GASAC_Gasterosteus

LLLLLVLVLLEGSVVFPLGAASSPDVDPRGHSEDRAVAEESQNYSKKAFPVLSLDYHHIH

VPFEIALWVLLASLMKLGFHLIPRLSSIVPESCLLIVVGLLVGGFIKLAGEKVPPVLDSQ

SFFLCLLPPIILDAGYFLPIRPFMENLGTILMFAVVGTLWNAFFVGGLLYAVCQIQPGNP

SDLHNLELLPCLLFASIISAVDPVAVLAVFEEIHINELLHILVFGESLLNDAVTVVLYHL

FEEYSGAGTVTILDGVLGVISFLVVALGGVLVGAIYGFLAAFTSRFTSHVRVIEPLFVFV

YSYMAYLSAEMFHLSGIMALIACGAVMRPYVEANISHKSHTTIKYFLKMWSSVSETLIFI

FLGVATVDGPHSWNWTFVTVTVLLCLVSRVIGVIGLTFVINKFRIVKLTTKDQFIVAYGG

LRGAIAFSLGFLLEEKHFPMRDMFLTAIITVIFFTVFVQGMTIKPLVELLAVKKKQEAKR

TINEEIHTQFLDHLLTGIEDICGHYGHHHWKDKLNRINKKYVKKCLIADERGREPQLIAF

YHKMEMKQAIELVESGGGFKPPSALPSTVSMQNIQPKKPAAAKPVQRALPQLPKGREEEI

RTILRSNLQRTRQRLRSYNRHTLVADPYEEGFSDFIIKKQKIIELEKKIIHMNNLTLPAA

PPDSPTL

>tr|G3Q3R6|G3Q3R6_GASAC_Gasterosteus

MTAQWTLSLLLSACLFRASCSAPAIPRPPRSLHPVRGFNNSGGSLSVDLPMALRVFSVDY

HHVQAPFEIVLWIMLASLAKLGFHWSGRLPAVVPESCLLIMVGLLVGGVIYGVRHSAPPT

LSADAFFLFLLPPIVLDAGYFLPGRLFFENLGTILWYAVVGTLWNVLGIGLSLYSLCLLA

PGSLGDISLLHCLLFGSLIAAVDPVAVLSVFQEMHVNEQLRILVFGESLLNDAVTVVLYK

LFESFLRMPSVSGLDVLLGGCRVLVVGMGGLFVGLFFGLIAALTSRFTSGAQVIAPLFVF

LYSYLSYLTSEMLHLSGIMAIVTCAVTMKQYVEANVSERSNTSIQYFLKMWSSVSETLIF

IFLGVSTIQDVHMWSWPFVCSTLLLCLVWRATGVLLLTAVVNKLRRNTVTFRDQFIIAYG

GLRGAICFSLVFLIDDFPKKRLFITTTIVVILFTVFVQGMTIKPLVELLDVKKEKGALPS

VSEEIHSRLIDHLLAGIEDVVGYWGQHYWKDRFDQFNVKYLRRILIREDHQACSSILRVY

QELERQEQRGDEDVPPPVVPRSLGRPLLPQEMDSIRCILSTNLQNFSNKQTAAYSRHTLH

QDAGTRKPLHRHQSLEERSTINTQDVLPLCQGEDGEFHDSRRVRSGLSRSHTVCSVSPRR

LLLDSSAPAASPQIADPRTAKKKLAFVLECEKQQ

>tr|G3NEX4|G3NEX4_GASAC_Gasterosteus

MPVFTMNYSRIQIPFEITLWVLLASFAKIGFHVYKKITVWVPESCLLITIGLIVGGIMHS

VHEEPPAVLRSNVFFLYMLPPIVLDSAYFMPTRPFFENIGTVLWFAVVGTLWNSIGIGMS

LYAICQIEAFGVQDINLQENLLFATIISAVDPVAVLSVFEDVSVNEQLYIVVFGECLFND

AVTVVLYNMFNFVAEMPVVEPVDVFLGVARFFVVGFGGMGFGILFGFTAAFTTRFTSRVR

EIEPLFIFMYSYLAYLVAELFSISSIMAIVTCALTMKYYVEENVSQRSCTTIRHVVKMLG

SIAETLIFFFLGVVTITTDHEWNWGYILFTLLFAFVWRGLGVLVLTQIINPFRTIPFNFK

DQFGLAYGGLRGAISFALVFTLPDSIGRKQLFVTATIAVILFTVFLQGISIRPLIEFINV

RRTNRNLDTINVEIHCRIMDHTIAGIEDLCGQWSHFYWKDKFMKFNNQILRKILIRDNRA

ESSIVALYKKLELQNAMEILDDVSGDISAAPSVVSPYEAKPKRKFLAPDLKNMHDILSKN

MYKIRQRTVAYTTKHALPNESQSKEILIRRHASIRRSLRPGSFQSSVVPKSHKYFSLPAG

KSLDSKFPPGMTSYADDEETMSEAAYPSRRSRLVQPARSSSSRAMMPLRRLDTLTEVHSV

DVVDEKVGDRRRAARGTSRTNSSSSRDPRIPAPQRHHSSADNGRGSADNFRNGHHEEEEE

EPQPLSPAPSWTAEPREDAAQNPLLRRPQWKPKK

>tr|G3NCZ3|G3NCZ3_Gasterosteus

MVSRDLANGGAAPSPATVTADPGHSTGGGQEAATNSSEGNHGATGGHETAPVTTLPIVTW

KWAHVSTPYLVALWVLVSWICKLIIEANHHVTNVIPESALLICFGFILGGIIWGADKVQT

FKLNPTVFFFYLLPQIILDTGYSMPNKLFFGNMGAILVYAVIGTCWNAASLGLSLWGCHQ

GGAMEGDLDIGLLQYLLFGSLIAAVDPVAVIAVFEQVHVNEVLFILVFGESLLNDGVTVV

LFNVFDAFVSLGGARINAVEIIKGIISFFVVAFGGSLLGLVFGLLISLLTKVTQKVKIIE

PAFVFVLGYLSYLTAEMLSLSSILSITFCGVSCQKYINANMAESSVSTVRYVMKVFANGS

ETIIFVFLGISAIDTEIWVWNTGFILLTLFFIFVYRIIGVFFLTWILNRYRLVPVEFIDQ

VIMSYGGLRGAVAYGLATLLNESKIKEKNLMICTTLLVVYFTVILQGITMKPLVNWLKVK

KAAVSELTLIEKLQSKVFDHMLVAIEDISGQIGHNYMRDKWNQFEERWMTRILMKKSVRK

SRDCVFKVFHQLNLKDAMSYVAEGERRGSLEFVRNESAFIDFKKKFVDDCSGAMPDITAC

MSDDYSAMSSMRIDQVPSVSLEMHEQNMKGVRETEDINTHHLLQQHLYKGRKQHRHRYSR

SHFNVNKDENEVQEIFQRTMRNRLESFKSAKMGVSPPKKISKHTKKDQQQKVPNGKSTEN

GKRYFSGDEDFQFPEGDASGFAASDGSHPFPMTYRAGGPAGIENPAYIPDLDPMAAEIPP

WLAEYDAGPVAPSQRAQLRLPWTPSDLGRLAPLRISTRSNDSFAQADAPAAQQKDDEPPT

PSKRDGRN

>tr|G3PZW9|G3PZW9_GASAC_Gasterosteus

GYRVVQWEWSYVQTPYIIATWLLLASVAKILFHFSQRFTTVVPESCMLILLGLVLGGVVL

IANEKQLYQLEPALFFLFLLPTIVGDAGYFMPARLFFDNLGAILTYAVVGTLWNAFCTGF

CLYAAKLLGVIDERVQADLMDFLLFGALISAVDPVAVLAVFEEVHVNDTLFIIVFGESLV

NDAVTVVLYKVYISFVEVGVDNVQTADYFKGVASFLIVSIGGTLVGLVFALILGFITRFT

KKVRIIEPLFVFLLVYLAYLTAELFSLSAILSMTFCGIGANKYVEANISQKSRTTVKYTM

KTLASIAETIIFIFLGISAVDKSKWAWDTGLVSCTLVFIFVFRAAGVIGQTWVLNRFRLV

PLDKIDQVVMSYGGLRGAVAFALVVLLDGKQVKAKDYFVATTIVVVFFTVMFQGLTIKPL

VDWLKVPRSTNRKPTINEEIHERTLSSCIDIYRDLSLPPACLPLPLRWEQFDKNYLSKLL

LRKSVYRKSELWEAYQKINIRDAISVIDQGGNVLTSARLSLPSMASRASFPEVTNVTNYL

RENGSGVCLDLQVIDNVPGAKVEEESETHHFLSGNLYKPRRRVIKTLSNIHCSDICKLQG

SLSRIQGSLKRLYGSLNKLQGSLILTSVVPPPQRRGSDSREEGGDKPRRNVSWQDKNPVV

VPVESEEDRGHSDPEKDEDVGITFIARQVETPTQQPRSGAVPATLDGGQSPPLAPSSPHL

PWKGSVGSPPPCVSVEATKIIPVDLQQAWKHSISSLERISSAP

>tr|G3QAV3|G3QAV3_GASAC_Gasterosteus

MGFTPRGFLGDKPAKAPPLFPFLLTFLLVGSRGDSNAMDNVSTERLAEESHRQDSANLLI

FITLLTLTILTIWLFKHRRFRFLHETGLAMIYGVLVGVILRFGVHVPQNMSDVILGCAVN

NSPATLLVNVSGRFYEYTLKGEVGRGKGHQVQDDEMLRKVTFDPEVFFNILLPPIIFHAG

YSLKRRHFFRNIGSILAYAFVGTVISCFVIGLIMYGFVSFMKAVGQLGGDFFFTDCLFFG

AIVSATDPVTVLAIFNELKVDVDLYALLFGESVLNDAVAIVLSSSIVAYQPAGDNSHSFE

AMAMLKSFGVFLGVFSGSFALGVATGFMTALVTKFTKLRDFPLLETALFFLMSWSTFLLA

EACGFTGVVAVLFCGITQAHYTFNNLSPDSQDRTKQLFELLNFLAENFIFSYMGLTLFSF

QSHVFNPLFIFGAFLAVFLGRAANIYPLSFLLNLGRKNKIGSKFQHVMMFAGLRGAMTFA

LSIRDTATYARQMMFSTTLLIVFFTVWICGGGTTPMLSFMSIPVGVDSDQQNSAVLDGSQ

RRDTKHESAWPFRIWYNFDHNYLKPLLTHSGPPLTATLPTCCGPLARCLTSPQAYENEGR

LFDDDSDSILNDATVSSMYADVTVSTDASGRTVNRSSGTRGLSDDCVDHDLALAEHEVAI

RGTRLVLPVDDPVEPPTTVTLPSPPSPP

>tr|G3NC67|G3NC67_GASAC_Gasterosteus

KCALLRCAPRSLPVPSSSSSSCAPLTGMLCVLVLVCVSVASASRDVGNNETRRDAAGEES

PHLTNSTSREKVFPVLALNYDHVRKPFEIALWILLALLMKLGFHIIPRVSNVVPESCLLI

GVGLLVGGIIRAIREEAPVLDTKLFFLYLLPPIILDAGYFLPIRAFTENMGTILVFAVVG

TLWNAFFIGGMIYGVCQIEGAQLASVDLLSCMLFGTIVSAVDPVAVLAVFEEIHINELLY

ILVFGESLLNDAVTVVLYHLFEEFCHAGTVTVVDILLGVVCFFVVSLGGILVGAIYGLLG

AFTSRFTSHTRVIEPLFVFLYSYMAYLSAEVFHLSGIMSLIACGVIMRPYVEANVSHKSY

TTIKYFMKMWSSVSETLIFIFLGVSTVAGPHAWNWTFVVFTVVLCLVSRVLGVIGLTFII

NKFRIVKLTKKDQFIMAYGGLRGAIAFSLGFLLTENETKSMFLTAIITVIFFTVFVQGMT

IRPLVELLAVKRKKEGKESINEEIHTQFLDHLLTGIEDVCGHYGHHHWKDKLNRFNKTYV

KKWLIAGDRSSEPQLISFYNKMEMKQAMMLAESGSAGRLPALMSTVSMQNIQLKQSARGR

AIPSISKSREQEIRKILRGNLQNTRQRLRSYSRHDLFMDPFEDNVSEVRFRKQRVEMERR

MSHYLTVPANRQEPPAVRKVCFEPEHRVYTYDDSESPRGPAARPRPGPPETVGLLKETLP

RPNQSGECRTKAREEQEAQEEQDPLKLSRCLSDPGPNKEDD

>tr|G3PQ84|G3PQ84_GASAC_Gasterosteus

LPVFTMDYPRIQVPFEFTMWVLLASFAKIGFNIYHKITIWIPESCLLISIGLIVGAIMHS

VKEEPPAVLNSNVFFLYMLPLIVLDSGYFMPTRPFFENVGTVLWYAVVGTLWNSIGIGIS

LFAICQFEVFGVQDINLQENLLFASIVSTVDPVAALNVFDDIGVNEQTYIIIFGEGLFND

AVTVVLYNMFTFLAALPVVESTDVFLGTAQFLVVAAGGVLFGLVFGFVAAFTTRFTHGVR

QIEPLFVFLYSYLAYLVAECFAISSVMALITCAITMKYYVEENVSQSSCTTIRHVVKMLA

TISETLIFFFLGVVTITTEHEWNWAYILFTLLFAFVWRGIGMLVLTQIINPFRTVPLNFT

DQFGLAYGGLRGAICFALVFTLPDSINRKDLFVTASIAVITFTVFIQGISIRPILEYMNI

RKTNKDVDTINVEIHTRTMEHVLSGVEDLCGQWSHYYWKDKFKKFNDRVLRRILLRHNRA

ESSIVSLYKRLELQSAIGLLDSPMGDLGAAPSLLPLHDERKEATRPKKKFLAADVRKMHH

ILSKNMYNIRQKTMAYTNKYDLPDDSQNREILICRHASVRRSLRAASFRQPHIPRSQKYF

SLQPGTALRPPSRGEDLSGDVQHRSAPPSRSPCRSPVPLERLDTI

>tr|G3NDF5|G3NDF5_GASAC_Gasterosteus

VDQKKDGQSDGCLGGGICIILVHLLIKFKLHFLPESVAVVSLGILMGGFIKIIEFQELAN

WKEEEMFRPNMFFLLLLPPIIFESGYSLHKYVQGNFFQNIGSITLFAVIGTAISAFIVGG

GIYFLGQADVIYKMSMTDSFAFGSLISAVDPVATIAIFNALNVDPVLNMLVFGESILNDA

VSIVLTNTAEGFSRSDNSATGWQTFVQALAYFLKMFFGSAALGTLTGLISALFLKHFDLR

KTPSLEFGMMIIFAYLPYGLAEGLKLSGIMSILFAGIVMSHYTHHNLSPVTQILMQQTLR

TMAFMCETCVFAFLGLSIFSFPHKFELSFVIWCIVLVLLGRAVNIFPLTVLLNVFRDHKI

TPKMMFIMWFSGLRGAIPYALSLHLGLEPIEKRQLIGTTTIIIVLFTILLMGGGTMPLIR

LMDIEESQSRRKSKKDITLSKTEKMGNTIESEHLSELTEEEYEAHIFQRRDLKGFMWLDA

KYLNPFFTRRLTQEDLLHGRIQMKTLTNKWYEEVRHGPSGSEDDDDEAELL

***Tilapia (Oreochromis niloticus),***

>tr|I3K858|I3K858_ORENI_Oreochromis

MATLWRFTLFLCMLLMACGGLSLASEEAETRNTQTETGTGSSSNTTGGGGHEAAPITTLP

IVTWKWHHISVQYLVALWILVCWLCKLIIEANHSVTNYIPESALLICSGFILGGMIWGAD

KMQTFSLSPTVFFYFLLPQIILDSGYHMPNKLFFTNLGAILIHAVIGTCWNAASLGLSLW

GCHMGGAMGDLDIGLLQYLLFGSLMAAVDPVAVLAVFDQVHVNEVLFIIVFGESLLNDGV

TVVLFNVFDAFVSLGGGRINAAEIVKGIVSFFVVAFGGSLVGIAFGILIALLTRCTKNIQ

IIEPGFIFVMGYLSYLTAEMLSLSAILSTTFCGVLCQKYVRANMDESSVQTVKYAMKVFA

NGSETIIFVFLGISAIDKVIWVWNTGFILLTLLFIFVYRFIGVFFLQWILNRYRLVPIEL

IDMVIMSYGGLRGAVAYGLAVLLDENKIKEKNLMICTTLIVVYFTVILQGITMKPLVMWL

KVKRAEVTEVSLLEKMTNKMFNHTLVAIEDISGQIGDNYFRSKWRHFEEKWMTPLLMKPS

ARKKQDEVFDVFHQLNLKDAMEYVTEGERSGSLEFVRNDSAILDFKKKLESEYTEVLPDI

PPYMINNHGAMFTMGRDPIPSVSLEMHEQTMKAMREAEAVDAHHLLDQHLYKSRKQYRHG

FSRSDFKINRDEKEVQEIFQRTMRNRLESFKSAKMGVSPPKTIVKHSKKEQQEKIPNGKS

MSKTKLCHSGDEDFEFSEGDSASGYDSSYPSFSMRATYRPGDGIENPAFVPDLYPTDLLQ

IPPWLAESEPDSSMVAPSLRAQANLPNSPMEMRRMGLLRTSTRSTDTKQMNNDQFPPPPT

PPPPPSGHM

>tr|I3KLH0|I3KLH0_ORENI_Oreochromis

SRLRMRGVSVKALSFLLLFISLCVSERAERENSLSLHERDDDGGNVKQQDNIIAEEGQTV

ENHEKEHGLNLDSNELQKDGSSLRSETKSNDTTDNYSNDTLHSTTTTVKPTTPAPPTVKP

ILPVQTGVKAQEEEQSSGMTIFFSLLVIGICIILVHLLIKFKLHFLPESVAVVSLGILMG

GFIKIIEFQELANWKEEEMFRPNMFFLLLLPPIIFESGYSLHKGNFFQNIGSITLFAVIG

TAISAFIVGGGIYFLGQADVIYKMTMTDSFAFGSLISAVDPVATIAIFNALNVDPVLNML

VFGESILNDAVSIVLTNTAEGFFSRSDNSTVTGWETFLQALGYFLNMFFGSAALGTLTGL

ISALFLKHFDLRNTPSLEFGMMIIFAYLPYGLAEGIKLSGIMSILFAGIVMSHYTHHNLS

PVTQILMQQTLRTVAFMCETCVFAFLGLSIFSFPHKFEISFVIWCIVLVLVGRAVNIFPL

SFLLNFCRDHKITPKMMFIMWFSGLRGAIPYALSLHLGLEPIEKRQLIGTTTIIIVLFTI

LFLGGGTMPLIRIMDIEDSQSRRKNKKDINLSKTQKMGNTIESEHLSELTEEEYEAQIYQ

RQDLKGFMWLDAKYLNPFFTRRLTQEDLLHGRIQMKTLTNKWYEEVRQGPSGSEDDEDEA

ELL

>tr|I3J5H3|I3J5H3_ORENI_Oreochromis

MLPLGLMLLLLPAAAADLPGTGSPPLRGPGLGFSPPVPASPSAPPLLLPQPGAEYGAPGV

ALARSAAEEPTGTGEEEDPGEHDHHHGGGYRVVQWEWSYVQTPYIIAIWLLVASVAKILF

HFSQRFTTVVPESCMLILLGLVLGGVVLIANKKQLYQLEPSLFFLFLLPTIVGDAGYFMP

ARLFFDNLGAILTYAVVGTLWNAFCTGFSLYTAKLLGFIDEHVQAEMLDFLLFGALISAV

DPVAVLAVFEEVHVNETLFIIVFGESLVNDAVTVVLYKVYISFVEVGAENVQTADYFKGV

ASFLIVSIGGTMVGLIFAVILGFITRFTKKVRIIEPLLVFLMVYLAYLTAELFSLSAILS

MTFCGIGANKYVEANISQKSRTTVKYTMKTLASIAETIIFIFLGISAVDKSKWAWDTGLV

SCTLVFIFIFRAVGVVGQTWVLNRFRLVPLEKIDQVVMSYGGLRGAVAFALVVLLDGEQV

KAKDYFVATTIVVVFFTVMFQGLTIKPLVKWLKVPRATNRKPTINEEIHERAFDHILTAV

EDIAGLQGYHHWRDKWEQFDKNYLSKLLLRKSVYRKSELWEAYQKINIRDAISVIDQGGN

VLTSARLSLPSMASRASFPEVTNVTNYLRENGSGVCLDLQVIDNIPGAKVEEDMETHHVL

AGNLYKPRRRYQSHYSRHFMPLGEQERQDREVFQRNMRSRMETFKSTRYKRHKKERSLKK

RRGSDVKEDGSDKPRRNVSWQDKDPVVVPVETEDKAEHSDPEKDEDVGITFIARKAETPK

QRPKSVPAALEGCQIPLLPPPPPAPPSPPPASSSPTSEDSHLPWKGSLGSPPPCVSLEAT

KIIPVDLQQAWNQSISSLESISSPPAPAEPVHPRVSALSRLGGPRPASYTPPGSAAGSTF

SIGPQVSQGSFLFPDKKKEEEEEEEDVEGGVAAEMQPLMSSLKPPGPLLPPPPPPSSAPS

SGRRRTPCVYLRSLVTSPPPEPHSRGPTQL

>tr|I3J3A8|I3J3A8_ORENI_Oreochromis

MVFKVTVSSARRATRTLWLLVLVSLPVCISVCRASSPQEDSAMENIVTEKKAEESHRQDS

ADLLIFILLLTLTILTIWLFKHRRFRFLHETGLAMIYGLIVGVILRYGIHVPRDVSNITL

NCQVNASPATLLVNVSGKFYEYTLKGEINANEVNDVQDNEMLRKVTFDPEVFFNILLPPI

IFHAGYSLKRRHFFRNMGSILAYAFLGTVISCFIIGLLMYGCVTLMKQVGQLGGDFFFTD

CLFFGAIVSATDPVTVLAIFNELQVDVDLYALLFGESVLNDAVAVVLSSSIVAYQPEGDN

SHTFEVMALLKSFGIFLGVFSGSFALGVATGIVTALVTKFTKLRDFQLLETALFFLMSWS

TFLLAEACGFTGVVAVLFCGITQAHYTYNNLSPESQDRTKQLFELLNFLAENFIFSYMGL

TLFTFQNHVFNPMFIVGAFVALFIGRAANIYPLSFLLNLGRRNKIRSNFQHMMMFAGLRG

AMTFALSIRDTATYARQMMFSTTLLVVFFTVWICGGGTTQMLSCQRIRVGVDSDQDNSIS

VGEGSERRSTKQESAWLFRIWYNFDHNYLKPILTHSGPPLTATLPPCCGPLARFLTSPQA

FENECQLKDDDSDLILRDGDISLTYGDITVSTDASGAHTSGGLAFAGASTSDDLDRELAY

GDHELVMRGTRLVLPMDDSEPSFTEPHHRMRM

>tr|I3K857|I3K857_ORENI_Oreochromis

MATLWRFTLFLCMLLMACGGLSLASEEAETRNTQTETGTGSSSNTTGGGGHEAAPITTLP

IVTWKWHHISVQYLVALWILVCWLCKLIIEANHSVTNYIPESALLICSGFILGGMIWGAD

KMQTFSLSPTVFFYFLLPQIILDSGYHMPNKLFFTNLGAILIHAVIGTCWNAASLGLSLW

GCHMGGAMGDLDIGLLQYLLFGSLMAAVDPVAVLAVFDQVHVNEVLFIIVFGESLLNDGV

TVVLFNVFDAFVSLGGGRINAAEIVKGIVSFFVVAFGGSLVGIAFGILIALLTRCTKNIQ

IIEPGFIFVMGYLSYLTAEMLSLSAILSTTFCGVLCQKYVRANMDESSVQTVKYAMKVFA

NGSETIIFVFLGISAIDKVIWVWNTGFILLTLLFIFVYRFIGVFFLQWILNRYRLVPIEL

IDMVIMSYGGLRGAVAYGLAVLLDENKIKEKNLMICTTLIVVYFTVILQGITMKPLVMWL

KVKRAEVTEVSLLEKMTNKMFNHTLVAIEDISGQIGDNYFRSKWRHFEEKWMTPLLMKPS

ARKKQDEVFDVFHQLNLKDAMEYVTEGERSGSLEFVRNDSAILDFKKKLESEYTEVLPDI

PPYMINNHGAMFTMGRDPIPSVSLEMHEQTMKAMREAEAVDAHHLLDQHLYKSRKQYRHG

FSRSDFKINRDEKEVQEIFQRTMRNRLESFKSAKMGVSPPKTIVKHSKKEQQEKIPNGKS

MSKTKLCHSGDEDFEFSEGDSASGYDSSYPSFSMRATYRPGDGIENPAFVPESEPDSSMV

APSLRAQANLPNSPMEMRRMGLLRTSTRSTDTKQMNNDQFPPPPTPPPPPSGHISL

>tr|I3K859|I3K859_ORENI_Oreochromis

WRFTLFLCMLLMACGGLSLASEEAETRNTQTETGTGSSSNTTGGGGHEAAPITTLPIVTW

KWHHISVQYLVALWILVCWLCKLIIEANHSVTNYIPESALLICSGFILGGMIWGADKMQT

FSLSPTVFFYFLLPQIILDSGYHMPNKLFFTNLGAILIHAVIGTCWNAASLGLSLWGCHM

GGAMGDLDIGLLQYLLFGSLMAAVDPVAVLAVFDQVHVNEVLFIIVFGESLLNDGVTVVL

FNVFDAFVSLGGGRINAAEIVKGIVSFFVVAFGGSLVGIAFGILIALLTRCTKNIQIIEP

GFIFVMGYLSYLTAEMLSLSAILSTTFCGVLCQKYVRANMDESSVQTVKYAMKVFANGSE

TIIFVFLGISAIDKVIWVWNTGFILLTLLFIFVYRFIGVFFLQWILNRYRLVPIELIDMV

IMSYGGLRGAVAYGLAVLLDENKIKEKNLMICTTLIVVYFTVILQGITMKPLVMWLKVKR

AEVTEVSLLEKMTNKMFNHTLVAIEDISGQIGDNYFRSKWRHFEEKWMTPLLMKPSARKK

QDEVFDVFHQLNLKDAMEYVTEGERSGSLEFVRNDSAILDFKKKLESEYTEVLPDIPPYM

INNHGAMFTMGRDPIPSVSLEMHEQTMKAMREAEAVDAHHLLDQHLYKSRKQYRHGFSRS

DFKINRDEKEVQEIFQRTMRNRLESFKSAKMGVSPPKTIVKHSKKEQQEKIPNGKSMSKT

KLCHSGDEDFEFSEGDSASGYDSSYPSFSMRATYRPGDGIENPAFVPDLYPTDLLQIPPW

LAESEPDSSMVAPSLRAQANLPNSPMEMRRMGLLRTSTRSTDTKQMNNDQFPPPPTPPPP

PSGHIYQL

>tr|I3JPC7|I3JPC7_ORENI_Oreochromis

MGFTRSRFLGGKPSKAPLLLPFLLTVLLVGSQGDGGAMDNGATERIAEESHRQDSAYLLI

FIMLLTLTILTIWLFKHRRFRFLHETGLAMIYGLLVGVILRFGIHAPQSMSDVILGCAVN

GSPATLLVNVSGRFYEYTLKGEVGQGKGHQVQDDEMLRKVTFDPEVFFNILLPPIIFHAG

YSLKRRHFFRNIGSILAYAFIGTVISCFVIGLIMYGFVSFMKVVGQLGGDFYFTDCLFFG

AIVSATDPVTVLAIFNELKVDVDLYALLFGESVLNDAVAIVLSSSIVAYQPAGDNSHSFE

AMAMLKSFGVFLGVFSGSFALGVVTGVVTALVTKFTKLRDFPLLETALFFLMSWSTFLLA

EACGFTGVVAVLFCGITQAHYTYNNLSPDSQDRTKQLFELLNFLAENFIFSYMGLTLFSF

QSHVFNPLFIVGAFVAVFLGRAANIYPLSFLLNLGRKNKIGYNFQHVMMFAGLRGAMTFA

LSIRDTATYARQMMFSTTLLIVFFTVWICGGGTTPMLSFMSIPVGVDSDQDNTNSGALDG

SQRRSTKHESAWPFRIWYNFDHNYLKPLLTHSGPPLTATMPACCGPLARCLTSPQAYENE

GQLHDDDSDFILNDASVSSMYADVTVSTDASGSRTVNPKRSSTTRAGGGVGSFDEGLDHE

LSVAEHEVAIRGTRLVLPMDDPVEAPTTVTLPPPPSPPRSEPHRHRL

>tr|I3KHH2|I3KHH2_ORENI_Oreochromis

KNKLVFIFIQGRLYFSAIMPSRTVSHGSSGRLRGLFLVMVLVFLESSVFTLAFASSDQHS

PTTAHSDGHKATDHGSHNNTGKAFPVLDLNYENVRVPFEISLWVLLASLMKLGFHLVPHL

SNIVPESCLLIVVGLLVGGIIRGTGETVPPVLDSKLFFYILLPPIILDAGYFLPIRPFME

NLGTILIFAVVGTLWNAFFVGGLLYGVCQIGTESSSILSRIGLLQCLLFGSIISAVDPVA

VLAVFEEIHINELLHILVFGESLLNDAVTVVLYHLFEEFSGIGTVTLVKGFLGVISFFVV

ALGGVLVGVLYGILAAFTSRFTSHTRVIEPLFVFLYSYMAYLSAEIFHLSGIMALIACGA

VMRPYVEANISHKSHTTIKYFLKMWSSVSETLIFIFLGVATVAGPHDWNWTFVTVTVILC

LVARVIGVVGLTFIINKFRIVKLTTKDQFIIAYGGLRGAIAFSLGFLLEKDHFPEREIFL

TAIITVIFFTVFVQGMTIKPLVELLAVKKKQEAKRSINEEIHTQFLDHLLTGIEDICGHY

GHHHWKDKLNRFNKKYVKKCLIAGERSKEPQLIAFYHKMEMKQAIQLVESGGGIKLPSAM

PSTVSMQNIQPRPPTKPAAERAIPQLSKTKEEEIRKILRTNLQRTRQRLRSYNRHTLVAD

PYEDGLSDLFLKKQKMFQIEKKLKEMNTYLTVPAPPPDSPTMCRARLASDPQAYSIKPAL

NKVPMIQVDLASPQSPDSANQQEEEEQGLMMKPPSRPREGKKPGEEDEAKDQQKLMRCLS

DPGPSADE

>tr|I3K9T8|I3K9T8_ORENI_Oreochromis

MNLVFIRQAGTLLFICMILILLHGSRGEIPSKPIAATILPSVKPDGGVQAYPEDQKNNLP

MFTMNYPRIQIPFEITLWVLLASFAKIGFHVYHKITIWVPESCLLISIGLIVGAIMHSVH

EEPPAVLTSNVFFLYMLPLIVLESGYFMPTRPFFENVGTVLWFAVVGTLWNSIGIGISLF

AICQIPAFGVQDINLQENLLFSAIISAVDPVAVLNVFEDVSVNEQLYIVVFGECLFNDAV

TVVLYNMFSFVAEMPVVEPVDVFIGVARFFIVGLGGMAFGILFGFVAAFTTRFTSKVREI

EPLFVFMYSYLAYLVAELFAISSIMAIITCALTMKYYVEENVSQRSCTTIRHVIKMLASI

SETLIFFFLGVVTITTDHEWNWGYILFTLLFAFVWRGLGILVLTQIINPFRTIPFNMKDQ

FGLAYGGLRGAVSFALAFTLPDTIGRKQLFVTATIAVILFTVFLQGISIRPLIEFINVRR

TNRNLETINVEIHGRLMEHTIAGIEDLCGQWSRFYWKDKFMKFNNRILRKILIRDSRAES

SIVALYKKLELQNAMEILDTASGDISAAPSIVSLYEEKKSSSKPQKKFLAADLKNMHDIL

SKNMYKIRERTVAYRGKHALPNDNHTREILIRRHVSIRRSLRPGSFQSSSYVPRTHKYFS

LPAGKSLESTFPAARRSYADDHETLSEVAYPSQRSRFGKPARSSSSKAMIPLRRLGTLEE

VPSTDVLNEDSSSTEQGVFRTNSSYSDSHVPRPRRLNSSFDNNNGSADDLRNVHEEPSSP

PPGWAAEPRDNTARNPLLRRPQWTPKRK

>tr|I3KMB0|I3KMB0_ORENI_Oreochromis

MSPSQVVHGCRMAALWTLSLLLSASLLGCLAVTSPDGSPLPHNLPPSLGTNSSDGSLSVD

LPMALRVFSMDYHHVQAPFEIVLWIMLASLAKLGFHWSGRVPAVVPESCVLIMVGLLVGG

VIYGVRHSAPPTLSADAFFLFLLPPIVLDAGYFLPGRLFFENLGTILWYAVLGTLWNVLG

IGLSLYGVCLLAQSSLGDVSLLHCLLFGSLIAAVDPVAVLSVFQEMQVNEQLHILVFGES

LLNDAVTVVLYKLFESFLRLPSVSGLDVLLGGCRVVVVGIGGLCVGLFFGLLAALTSRFT

SRAQVIAPLFVFLYSYLSYLTSEMLHLSGIMAIVTCAATMKQYVEANVSERSNTSIQYFL

KMWSSVSETLIFIFLGVSTIQDIHMWSWPFVCSTLLLCLIWRATGVLLLTAFVNKLRRNA

VTFRDQFIIAYGGLRGAICFSLVFLIDDFPKKRLFITTTIVVILFTVFVQGMTIKPLVEL

LYVKRKKRALPTLSEEIHSRLIDHLVAGIEDVVGYWGQHYWKDKFEQFNKKYLRRFLIRN

DRQASSSILRVYQELERREQRGVEEEAPPIKPRVDPRSQGRPLLPEEMESIRRILSRNLQ

NFNNKQTPAYSRHTLHQDTSTSKSGRKILIQHNHTL

>tr|I3JEY2|I3JEY2_ORENI_Oreochromis

MMSGAMQQKQDNVALPVFAAVTLALITILSVWKLKQFKYKLVNEAGVAMFYGLLLGLTVR

LIWTDKGEQCPGNTTATNEQENTSDISMVNITPHGHIRNDKRQFSSHPEETFDLQVLFNL

FLPPIIFHGAYTLNQKRFIANLGSVLTYAFAGTIISCMCIGACMYGFTKLMPPKEEFLLT

HCLLFGAIMSATDPVSVLGLLSDLRVDLDLHMLLFGESVLNDAVAIVLTHAITTYTQMDA

GLIFDPTAFLHAFVFFLGVLIGSFLLGLIFTVITALLTKFTRLHENPLLETSVFFLLSWS

SYLSAEACGLSGIVAVMFCGLSQARYTVLNLSSEGRARIKQLFEVFNFLGEISIFGYMGY

ILLKFTCHKFEPVFISGALLSVFLSRAFNIYPLSFLLNLGRTNKIPCKFQHFMIFAGLRG

AVAFRLAVKETDTEVTRTIFTTTLLMIGFTIWVLGTAADPMLRRLDIRVGVDPDENQEDH

SFEIATERPDTTQGLWHSLDYKYLKPVLTHCGPPLTESLPQCCGVFARVFSSPRVQEDEE

EELCEIEAGKNTLDLGKTESQQGSSESDSASEHQEDLLDGDLGLGTAPVHPREASGSSTQ

L

>tr|I3JA90|I3JA90_ORENI_Oreochromis

MAAWILLSCAWGQGMRAEDFAMEELATEKEAEESHRQDSFNMLTFILLLTLTILTIWLFK

HRRVRFLHETGLAMIYGLLVGVILRYGIPANSYHNKTPASCSLREGPTSTLLLNVSGKFF

EYTLKGEINSREIHNVEQNDMLRKVTFDPEVFFNILLPPIIFHAGYSLKKRHFFRNLGSI

ITYAFLGTAVSCFVIGNLMYGVVKLMQAVGQLTGKFYYTDCLFFGAIISATDPVTVLAIF

NELHADGDLYALLFGESVMNDAVAIVLSSSIVAYQPAGANTHQFDASAFFKSVGVFLGIF

SGSFVMGAATGVVTALISKFTKLHCFPLLETALFFLMSWSTFLLAEACGFTGVVAVLFCG

ITQAHYTYNNLSEESTKRTKQLFEVLHFLAENFIFSYMGLALFTFQNHIFSPIFIIGAFI

AIFIGRALNIYPLSFLLNLGRRHKIKGNFQHMMMFAGLRGAMAFALAIRDTATYARQMMF

TTTLLIVFFTVWVFGGGTTPMLSWLHIRVDVDPDQDLQPSADSFQVLQGDGPQAEGQSKT

KQESAWLFRLWYTFDHNYLKPILTHSGPPLTSTLPSYCGPLASCLTTPRAYENHEQLRDP

DSDLIRGDADLTFTFGDTAITANGASGSGADAAGGSGWSANGKRSGSTSEEALERELDSR

EQELLSRGTRLVFPIEDHI

>tr|I3K8I6|I3K8I6_ORENI_Oreochromis

PSDQLKMAPLRRSSTLRLTGLLCLSFLVFFSRGFASHDVVSEKSSGTIQEDSHSQANSTA

HKKTFPVLSFNYEHVRTPFEISLWILLALFMKLGFHIIPTVSHIVPESCLLIFVGLLVGG

IIKATGEQPPVLDSNLFFLYLLPPIILDAGYFLPMRSFTENIGTILMFAVVGTLWNVFFI

GGMMYAVCQIDAAELAHVDLLSCLLFGSIISAVDPVSVLAVFEEIHINELLHILVFGESL

LNDAVTVVLYHLFKEFAHEQTITVTGAILGVISFFVVSLGGVMVGVIYGIVGAFTSRFTS

YSRVIEPLFVFLYSYMAYLSAEIFHLSGIMSLIACGVIMRPYVEANISHKSYTTVKYFLK

MWSSVSDTLIFIFLGVSTVAGPHTWNWIFVIFTVIFCLVSRVLGVIGLTAFINKFRIVKL

TKKDQFIIAYGGLRGAIAFSLGFLLDNDEIKNLFLTAIITVIFFTVFVQGMTIRPLVELL

AVKKKKESKESINEEIHTQFLDHLLTGIEDICGHYGHHHWKDKLNRFNKTYVKRWLIAGE

RSTEPQLISFYNKMELKQAMMMVESGSAAKLPTIVSTVSIRNVPTKGAPKPSRAISKSRE

EEIRRLLRANLQKTRQRLRSYSRYELLNDPLEDNISEARFRKQREMERRMSHYLTVPANR

QETPPVRKVFFQPEHKVYTYDDSESPRGRASPSCADDVSELKETPEKPDQSSTPRANRDA

ELKPKEQDDQGDQLKVSRCLSDPGPNPEDDEDSTFLP

>tr|I3J3A7|I3J3A7_ORENI_Oreochromis

LFRRFILFAALNFSTGVDFNHPRSARRATRTLWLLVLVSLPVCISVCRASSPQEDSAMEN

IVTEKKAEESHRQDSADLLIFILLLTLTILTIWLFKHRRFRFLHETGLAMIYGLIVGVIL

RYGIHVPRDVSNITLNCQVNASPATLLVNVSGKFYEYTLKGEINANEVNDVQDNEMLRKV

TFDPEVFFNILLPPIIFHAGYSLKRRHFFRNMGSILAYAFLGTVISCFIIGLLMYGCVTL

MKQVGQLGGDFFFTDCLFFGAIVSATDPVTVLAIFNELQVDVDLYALLFGESVLNDAVAV

VLSSSIVAYQPEGDNSHTFEVMALLKSFGIFLGVFSGSFALGVATGIVTALVTKFTKLRD

FQLLETALFFLMSWSTFLLAEACGFTGVVAVLFCGITQAHYTYNNLSPESQDRTKQLFEL

LNFLAENFIFSYMGLTLFTFQNHVFNPMFIVGAFVALFIGRAANIYPLSFLLNLGRRNKI

RSNFQHMMMFAGLRGAMTFALSIRDTATYARQMMFSTTLLVVFFTVWICGGGTTQMLSCQ

RIRVGVDSDQDNSISVGEGSERRSTKQESAWLFRIWYNFDHNYLKPILTHSGPPLTATLP

PCCGPLARFLTSPQAFENECQLKDDDSDLILRDGDISLTYGDITVSTDASGAHTSGGLAF

AGASTSDDLDRELAYGDHELVMRGTRLVLPMDDSEP

>tr|I3K860|I3K860_ORENI_Oreochromis

SPVKGEAVVTHEGSHLKLSKSSHSNESQHGHSSNATGIPIISFKWHHVETSYLVAFWVLV

SFLGKLVIEANHHVTSVIPESALLICFGFILGGMIWAADKVQTFTLTPTVFFLYLLPQVI

LDAGYHMPNKLFFTNLGGILVHAVIGTCWNAASLGLSLWGCQMGGAMGDLDIGLLQYLLF

GSLMAAVDPVAVIAVFEQVHVNDVLFIMVFGESLLNDGVTVVLFNVFDAFVSLGGPKIDA

AEIIKGIVSFFVVAFGGSLVGLGFGLLISLLTRCTKNIQIIEPGFIFVLGYLSYLTAEML

SLSAILSIVFCGMCCQKYVRANMDENSVQTVRYAMKAFANGSETIIFVFLGISAIDKVIW

VWNTGFILLTLLFVLVYRFIGVFFLTWILNRFRLVPLEFVDQVIMSYGGLRGAVAYGLAV

MLDENKIKEKNLMVCTTLIVVYFTVILQGITMKPLVMWLKVKRAAVTEVSLLEKLTNKMF

NHTLVAIEDISGQIGHNYMRSKWKYFEENWMSWILMKPSARKKQDQLLDVFHQLNLKDAM

EYVTEGERRGSLEFVRNENSIIDFKKKSESESFDINNANISNDHGVMSNTSRDPIPSVSL

EMHEQTMKAVREAEAINAHHLLDQHLYKSRKQHRHRFSRSHFDINRDEKEVQEIFQRTMR

NRLESFTSAKMGVKPPKTIVKHTQKEQQQKMPNGKSMDKAKTSYSGDEGFKIGSEGDSAS

GYDQSYMRATYRAGAGIENPAFMPDLDPMDLLQIPPWLAESELDSSMVALSHRAQVGLPR

TPSNLRRLAPLRISTRSTDSFTLENIPDTQQMDSDQLPPPPPPRPPRDDGYV

>tr|I3KML0|I3KML0_ORENI_Oreochromis

DRCLFIVVGLVFGGLFRAGGQGVPQILHKLFFLCLLPPIILDAGYFLPIEPLFKNLGRVL

LFGFVGSIWNGFFVGGILYGVCRIGTGSTVLTNTGLLQCLLFGSITSAVDSAAVFEKIQM

KDILHILLFGESILSDAVAVVLYHLYHEFHVTASVTFMKVFLGVIAFSLAGLGGILVGSC

YGILTAFTTRFTSNTGIVEPLVVFLFCNLAYLSADMFNLSGIIALISCCVVMRPFVQANI

SCESEMTIKYFLKMWSRFNESLIFIFVGVVTVAGPQVWNWTFITLTVILCVVARVIAVVG

LTFLTNKFHSVKLTKEDQFLFVCSGLRGGVAFFLAFLLEKDHFPEKEIFVTAIITVIFFT

IFVQCMAIKPLMKLLAVKKNQLATDQEIRTQRLKNLSLHIDGVCGDNGHHHRMDKLKQFS

DLYLKKWLIAGEGKMEEQ

>tr|I3JPC6|I3JPC6_ORENI_Oreochromis

MGFTRSRFLGGKPSKAPLLLPFLLTVLLVGSQGDGGAMDNGATERIAEESHRQDSAYLLI

FIMLLTLTILTIWLFKHRRFRFLHETGLAMIYGLLVGVILRFGIHAPQSMSDVILGCAVN

GSPATLLVNVSGRFYEYTLKGEVGQGKGHQVQDDEMLRKVTFDPEVFFNILLPPIIFHAG

YSLKRRHFFRNIGSILAYAFIGTVISCFVIGLIMYGFVSFMKVVGQLGGDFYFTDCLFFG

AIVSATDPVTVLAIFNELKVDVDLYALLFGESVLNDAVAIVLSSVLLRFEKLGRHGAGAV

RSAGNESSLLPEVAEEWLGENQTVTPRSIVAYQPAGDNSHSFEAMAMLKSFGVFLGVFSG

SFALGVVTGVVTALVTKFTKLRDFPLLETALFFLMSWSTFLLAEACGFTGVVAVLFCGIT

QAHYTYNNLSPDSQDRTKQLFELLNFLAENFIFSYMGLTLFSFQSHVFNPLFIVGAFVAV

FLGRAANIYPLSFLLNLGRKNKIGYNFQHVMMFAGLRGAMTFALSIRDTATYARQMMFST

TLLIVFFTVWICGGGTTPMLSFMSIPVGVDSDQDNTNSGALDGSQRRSTKHESAWPFRIW

YNFDHNYLKPLLTHSGPPLTATMPACCGPLARCLTSPQAYENEGQLHDDDSDFILNDASV

SSMYADVTVSTDASGSRTVNPKRSSTTRAGGGVGSFDEGLDHELSVAEHEVAIRGTRLVL

PMDDPVEAPTTVTLPPPPSPPRSEPHRHRL

***Mudskipper (Boleophthalmus pectinirostris)***

>XP_020789995.1_Boleophthalmus

MATLRQYALCFYVFCMLVTVCQANTAVQETTEANSETDTGNATHEETAGGEHGAPITTLPIVSWKWHHVEVPYLVALWVLVCWLCKLVIEANHHVTAYIPESALLICSGFILGGIIWGADKVQTFSLTPTVFFFYLLPQIILDAGYFMPNKLFFSNLGGILIHAIIGTCWNAATVGVSLWGCHMGGAMGDIDIGLLQYLLFGSLLSAVDPVAVIAVFEEVHVNEVLFIMVFGESLLNDGVTVVLFNVFDAFVSLGGAEIDAVEIIKGIISFFVVAFGGALVGVIFGIMLSLLTRCTKNIQIIEPGFVFVLGYLSYLTAEMLSLSAILSITFCGVCCQKYINANMDEKSVTTVRYVMKVFANGSETMIFVFLGISAIDKTIWVWNTGFILLTLLFVLVYRVIGVFFLTWIQNMFRLVPLDFTDRVIMSYGGLRGAVAYGLAVLLDENKIKERPLMIGTTLIVVYFTVVLQGITMKPLVTWLKVKRAAVTELTLVEKVQNKVFDHALVAVEDISGQKGHNYMRDKWNNFESKWLSWILMKPSARKKHDYVFNVFHKLNMKDALDYVAEGECNGTLEFVRNETAYIDFKKKFGDDFSDISMPDITIDMGDTTITNPYARRDTVPSISMEMHEQTMNGLRESEDMNTHHLLQQHLYKGQKQHRHRYSRSHFDVNQDENEVQEIFQRTMRNRLESFKSAKMGVMPPKTITKHVKKDAQQKMPNGKSKSDEEGNSASGYTTSIPRVTYRPGAGIENPAFLPDLDPMAPTQIPPWLAEGEEDIATVSPSQRAQAELPYTPSNLRRLAPLRISTRSTDSFVLADHPPTSSSWDKDNIPPPPSPPPY

>XP_020780200.1_Boleophthalmus

MPKPGLPSALLLLGVSLVALGLLPASSADPSSGDRGTRGPVALAHSAAAAAEEPAVPGVGAEDGSGEQHQHHGHGGGYKVVQWEWSYVQTPYIIAIWLLVASVAKILFHFSQRFTTVVPESCMLILLGLVLGGIVLIANKKQLYQLEPALFFLFLLPTIVGDAGYFMPARLFFDNLGAILTYAVVGTLWNAFCTGFSLYAAKVLNVIDEKVQADLMDFLLFGALISAVDPVAVLAVFEEVHVNDTLFIIVFGESLVNDAVTVVLYKVYISFVEVGPKNVQTADYFKGVASFLIVSIGGTLVGLFFAVILGFITRFTKKVRIIEPLFVFLLVYLAYLTAELFSLSAILSMTFCGIGANKYVEANISQKSRTTVKYTMKTLASIAETIIFIFLGISAVDKSKWAWDTGLVSCTLIFIFIFRAVGVVGQTWVLNRFRLVPLDKIDQVVMSYGGLRGAVAFALVVLLDGEQVKAKDYFVATTIVVVFFTVMFQGLTIKPLVKWLKVPRSTSRKPTINEEIHERAFDHILTAVEDIAGLQGYHHWRDKWEQFDKNYLSKLLLRKSVYRKSELWEAYQKINIRDAISVIDQGGNVLTSARLSLPSMASRASFPEFTNVTNYLRENGSGVCLDLQVIDNVPGAKVEEDTETHHVLAGNLYKPRRRYQSHYSRHFMPLGEKERQDREVFQRNMKYRMESSKSTRHKRHKKERSIKKRRGSDAKEESSEKPRRNVSWHDKEPVLVPVDSEDDKGEPSEKEEDEGITFVARKVETPKPRPKSVPAATEVSSGIGSPPPCVSVEATKIIPVDLQQAWNQSISSLESISSPPAPPAAAADAVHPRISALSRIGASRPASYTPSSSSSFFSTSNTQLVQGSFLFPEKSPKEEEEDESSHELQPLMSTLKPPTATSSSSAMAPPPPPPGSAPGQGRRKHPRVFLRSLVSAPPPGSTEPRSRGPTQL

>XP_020795955.1_Boleophthalmus

MRTRTRSSSPAETGLLLALVLVLLDGLVIHTAAGASASPADATNHHSPTNDSSPGHHKKPFPVLDLNYKHVQTPFEISLWVLLASLMKLGFHLIPRLSNIVPESCLLIVVGLLVGGLIRLAGEEVPPVMNADVFFLCLLPPIILDAGYFLPIRPFMENLGTILMFAVVGTLWNAFFVGGLLYAVCQINPGYPSDLHQLELLPCLLFGSIISAVDPVAVLAVFEEIHINELLHILVFGESLLNDAVTVVLYHLFEEYAGAGTVTVLDGFLGIVSFLVVALGGVLVGAIYGLLAAFTSRFTSHMRVIEPLFVFVYSYMAYLSAEMFHLSGIMALIACGAVMKPYVEANISHKSHTTIKYFLKMWSSVSETLIFIFLGVATVDGKHHSWNWIFVTATVFLCLVARVMGVVGLTCIINKFRMVKLTTKDQFIIAYGGLRGAIAFSLGFLLSDVHFPKKDMFLTAIITVIFFTVFVQGMTIKPLVELLAVKKKQEAKRSFNEEIHTQFLDHLLTGIEDICGHYGHHHWKDKLNRFNKKYVKRCLIAGDRCEEPQLIAFFHKMEMKQAIELVEKGGANMPAIPSTVSMQNIQPRKPVSTDRMLPQLPKGKEEEIRSILRSNLQRTRQRLHSYNRHTLVADPYEDGFNDLILRKKRVMEKKIKEMNNYLTVPAPPPDSPTMCRARLASDPQAYSMKPTADDVPTIHVDLASPQSPDSVNLMDELKKKNESKPDEDQGLVMRPPAQNRSEDPDREQQKLTRCLSDPGPSADEEEDEPFLP

>XP_020790001.1_Boleophthalmus

MAVLPGFLQSPTQIGSSLHFTARFWCFLLLICLAVASSSRDVVPKNDGSTGTVHGDGSGHHNMSTIHKKAFPVLSFNYAHIRKPFEISLWILLALLMKLGFHIIPRVSNIVPESCLLIVVGLLVGGVIKAIKEEPPVLDDQLFFLYLLPPIILDAGYFLPIRPFTENLGTILTFAVVGTLWNTFFIGGTMYGVCQIAGAQLQRVDLLSCLLFGTIISAVDPVAVLAVFEEIHINELLHILVFGESLLNDAVTVVLYHLFKEFSLEGGVTVGDAFLGVVCFFVVALGGILVGAIYGILGAFTSRFTSHTRVIEPLFVFLYSYMAYLSAEVFHLSGIMSLISCGVMMRPYVEANISQTTKYFLKMWSSISETLIFIFLGVSTVVGPHTWNWTFVIVTVFLCLGSRVLGVIGLTYIINKFRIVKLTKKDQFIVAYGGLRGAIAFSLGFLLTNDDLKHMFLTAIITVIFFTVFVQGMTIRPLVELLAVKRKKETKGSINEEIHTQFLDHLLTGIEDICGHYGHHHWKDKLNRFNKTYVKRWLIAGERSTEPQLLSFYNKMEMKQAMMLVESGSTAKLPTLVSTVTLQNIQPKDLPKKRAIPSISKKREEEIRKILRANLQKTRQRLRSYSRHDLMSDPFEDNVSEMRFRRQRVEMERRMSHYLTVPAKTQERPSLRKVSFEPEHKVYTYDDSESPNRRRSEGASVSSRDPDDVFVPNETPPPTNQRAAPKKPTNQTTDEDEDDVALTMSRCLSDPGPKEEDEDDAFLP

>XP_020773382.1_Boleophthalmus

MAPLLLSLLVHTCLMVGCSTRTPPVDPSEPQPMHSSLGQNGSSEGVPLVDVPMALGVFSVDYHHVQAPFEIVLWIMLASLAKLGFHWSGRVPAVVPESCLLIMVGLLVGGVIYGVRHSAPPTLSADAFFLFLLPPIVLDAGYFLPGRLFFENLGTILWYALLGTLWNVLGIGLSLYGVCLLCPSSLGDISLLHCLLFGSLIAAVDPVAVLSVFQEMHVNEQLHILVFGESLLNDAVTVVLYQLFESFLRLPSVSGLDVLLGGCRVLLVGFGGLFVGLFFGLVAALTSRFTPRVPVIAPLFVFLYSYLSYLTSEMLHLSGITAIVTCAVTMKQYVEANVSERSNSSIQYFLKMWSSVSETLIFIFLGVSTIQDVHMWSWAFVCSTLLLCLIWRGTGVLLLTAVVNKVRRNKVTFRDQFIIAYGGLRGAICFSLVFLIDDFPKKRLFITTTIIVILFTVFVQGMTIKPLVDLLDVKRKKRALPTVSEEIHSRLIDHLLAGIEDVVGYWGQHYWKDKFEQFNTKFLRRFLIREEQQNRSSILRVYQELERREQGAHRETTIQARTVPQTASRPLLPEEMDSIRRILSRNLHHFNSRMPAYSRHTLHQDRSLQRHRSLGERDTQVEDAEFNELHILRSGLRRSHTVCAISPSCASLDSPLPDLQTAPKHGPKKHLSFDIENEKNN

>XP_020794800.1_Boleophthalmus

MDIGFSRSAGFILFICSLLTLGSNGEVPPKPTRSVTILPPIKPMDGPQAYPDAEKASLPVFTMDYPRIQIPFEVTLWVLLASFAKIGFHVYHKITVWVPESCLLISIGLIVGAIMHSVHEEPPAVLTSNVFFLYMLPPIVLDSGYFMPTRPFFENIGTVLWFAVVGTLWNSIGIGMSLFAVCQIEAFGVQDINLQENLLFATIISAVDPVAVLNVFEDVSVNEQLYIVVFGECLFNDAVTVVLYSMFTFVADMPVVEPVDVFLGVARFFVVGLGGMAFGVLFGFVSAFTTRFTSKVREIEPLFIFMFSYLAYLVAELFAISSIMAIVTCALTMKYYVEENVSQRSCTTIRHVVKMLGSISETLIFFFLGVVTITTVHEWNWGYILFTLLFAFVWRGLGVLVLTQIINPFRTIPFNLKDQFGLAYGGLRGAISFALAFTLPDNIGRKQLFITATIAVILFTVFLQGISIRPLIEFINVRKTNRNLETINVEVHCRLMEHTIAGIEDLCGQWSHFYWKDKFLKFNNRILRKILIRDSRAESSIVALYKKLELQNAMEILDTVAGDISAAPSIVSLYEEKKNPKPKMFVASDLRNMHDILSKNMYKIRQRTVAFTSKHALPNDRHTKEILIRRHASIRRSLRPGSFQSSAVSKSQKYYSLPAGQNLESRLSPRRLTQTDDRETMSEVAYPIANRRSRSSQPTRSSSSRAMMPLRRLDTLTEAVSVDMLNENKENNGSRAGTTRSNSSSSDTRLPGPRQQLSPTRDDNGLSDNLINVHHEAEEEPSAPSPPPVWVAEPRGNVNRNPLLRRPLWNPKRT

>XP_020797456.1_Boleophthalmus

MGRPSIVGVRERAAWCTLLLLLCSSGGDCESELPQPPGPTNFQGVKPFHNEPEPQPFPDKEKANLPVFTMDYPRIQVPFEFTLWILLASFAKIGFHIYQKITVWIPESCLLICIGLIVGGIMYSVKEEPPAVLSPEVFFLYMLPPIVLDSGYFMPTRPFFENVGTVLWYAVVGSLWNSVGIGVSLFAICQIEAFGVQDINLQENLLFASIIAAVDPVAALSVFEDISVNEQIYIVIFGEGLFNDVVTVALYSMLSFVADMPTVESVDVFLGVARFFVVSLGGLLIGVVFGIAAAFTTRFTHNARQIEPLFVFMYSYLAYLVAELFAISSVLSIITCAITMKYYVEENVSQRSCTTIRHVVKVLASISETLIFFFLGVVTITTEHEWNWAYILFTLLFAFVWRGLGILVLTQIINPFRTIQFNFKDQFGLAYSGLRGAVCFALVFTLPDTINRRNLFVTASIAVIIFTVFIQGISIRPIVEYMDIRRTNNEKYNINMEIHSRTMEHIVSGIEDLCGQWGHYYWKDKFKKFNDRILRKILLRDNKAESSIVSLYKKLELQSAIELLDTPMGDLSAAPSVVSLHDEMKEASRPKKKFLSADVRNMHNILTKNMYKIRQKTLAYTTKYKLPDDSRTREILIRRHTSIRRSLRATSFRHPPSENIPKSQKYYSLQPGSDLGSALALRRRSQGGSTARSIIPLSPIEERNLSPAPHRTTPTLQQSEGGDEATPHGWQPEDPEGNTVSRSPLLR

>XP_020774408.1_Boleophthalmus

MDALRSYRGALQGMLVVWLMFAAEIIRLSGVKASDVMEELATEKEAEESHRQDSVNLLTFILLLTLTILTIWLFKHRRVRFLHETGLAMIYGLLVGVILRYGIPATSYHNQTPLSCSLKEGPASTLLLNVSGKFFEYSLKGEINLREIHNVEQNDMLRKVTFDPEVFFNILLPPIIFHAGYSLKKRHFFRNLGSILTYAFIGTAVSCFVIGNLMYGVVKLMQVAGQLMDKFYYTDCLFFGAIISATDPVTVLAIFNDLHADGDLYALLFGESVMNDAVAIVLSSSIVAYQPSGANSHKFDASALFKSIGVFLGIFSGSFLMGAANGVVTALVTKFTKLHCFPLLETALFFLMSWSTFLLAEACGFTGVVAVLFCGITQAHYTYNNLSEESTKRTKQLFEVLHFLAENFIFSYMGLALFTFQNHIFSPIFIFGAFIAIFIGRALNIYPLSFLLNLGRRHKITGNFQHMMMFAGLRGAMAFALAIRDTATYARQMMFSTTLLIVFFTVWVFGGGTTPMLSWLHIRVGVDSDQDFQPTGDNQSKTKQESAWFFRLWYTFDHNYLKPLLTHSGPPLTSTLPSYCGPLASCLTSPQAYQDDDQLRDPDSDLIRNDQDLTFMFGDTSLTANGTSSAREDSDAGATNWSVNGKRSGSTSEEALERDLDSRDQELLSRGTRLVFPTQDQL

>XP_020774407.1_Boleophthalmus

MDALRSYRGALQGMLVVWLMFAAEIIRLSGVKASDVMEELATEKEAEESHRQDSVNLLTFILLLTLTILTIWLFKHRRVRFLHETGLAMIYGLLVGVILRYGIPATSYHNQTPLSCSLKEGPASTLLLNVSGKFFEYSLKGEINLREIHNVEQNDMLRKVTFDPEVFFNILLPPIIFHAGYSLKKRHFFRNLGSILTYAFIGTAVSCFVIGNLMYGVVKLMQVAGQLMDKFYYTDCLFFGAIISATDPVTVLAIFNDLHADGDLYALLFGESVMNDAVAIVLSSSIVAYQPSGANSHKFDASALFKSIGVFLGIFSGSFLMGAANGVVTALVTKFTKLHCFPLLETALFFLMSWSTFLLAEACGFTGVVAVLFCGITQAHYTYNNLSEESTKRTKQLFEVLHFLAENFIFSYMGLALFTFQNHIFSPIFIFGAFIAIFIGRALNIYPLSFLLNLGRRHKITGNFQHMMMFAGLRGAMAFALAIRDTATYARQMMFSTTLLIVFFTVWVFGGGTTPMLSWLHIRVGVDSDQDFQPTGDSFQVLQGDGAQDQSKTKQESAWFFRLWYTFDHNYLKPLLTHSGPPLTSTLPSYCGPLASCLTSPQAYQDDDQLRDPDSDLIRNDQDLTFMFGDTSLTANGTSSAREDSDAGATNWSVNGKRSGSTSEEALERDLDSRDQELLSRGTRLVFPTQDQL

>XP_020785393.1_Boleophthalmus

MFGFRGLVLFSSVALLLLNPSRLTLAQQQERDGAGEADDHKPVTAKKQSLSINNEGNDGTAEGEEPKDGQPASKSNDTDIYNNDTLHSPTTTNKPTTMSPPTAKPILPVQTGVKAQEEEQSSGLTIFFSLLVIGICIILVHLLIKFKLHFLPESVAVVSLGILMGGFIKIIEFQELANWKEEEMFRPNMFFLLLLPPIIFESGYSLHKGNFFQNIGSITLFAVIGTAISAFIVGGGIYFLGQADVIYKMTMTDSFAFGSLISAVDPVATIAIFNALNVDPVLNMLVFGESILNDAVSIVLTNTAEGFFSQSDNSTVTGWEIFIQALGYFLKMFFGSAALGTLTGLISAVFLKHFDLRKTPSLEFGMMLIFAYLPYGLAEGIQLSGIMSILFAGIVMSHYTHHNLSPVTQILMQQTLRAVAFMCETCVFAFLGLSIFSFPHKFEISFVIWCIVLVLLGRAINIFPLSFLLNFFRDHKITPKMMFIMWFSGLRGAIPYALSLHLGLEPIEKRQLIGTTTIIIVLFTILLLGGGTMPLIRIMDIEESQSRRKSKKDVNLSKTQKMGNTIESEHLSELTEEEYEAHIYQRQDLKGFMWLDAKYLNPFFTRRLTQEDLLHGRIQMKTLTNKWYEEVRQGPSGSEDDEDEAELL

>XP_020779124.1_Boleophthalmus

MSFTVTVRWARRAIGPRWLLVLVSLCVCVRVCRAASPPEEEDSAMENIVTEKRAEESHRQDSADLLIFILLLTLTILTIWLFKHRRFRFLHETGLAMIYGLLVGVVLRYGIHMPRDGTNVILNCNVNASPATVLVNVSGKFYEYTLKRVISSSGNKVDDMSDNENEMLRKVTFDPEVFFNILLPPIIFYAGYSLKRRHFFRNMGSILAYAFLGTVISAFIIGLLMYGTVTLMKHVGQLAGEFFFTDCLFFGAIVSATDPVTVLAIFHELQVDADLYALLFGESVLNDAVAVVLSSSIVAYQPAGDNSHTFEGMALLKSFGMFLGVFSGSFALGVATGIVTALVTKFTKLRDFQLLETALFFLMSWSTFLLAEACGFTGVVAVLFCGITQAHYTFNNLSPESQDRTKQLFELLNFLAENFIFSYMGLTLFTFQNHVFNPMFIVGAFLAVFLGRAANIYPLSLLLNLGRRHKIRSNFQHMMMFAGLRGAMTFALSIRDTATYARQMMFSTTLLIVFFTVWICGGGTTQMLSCQRIRVGVDSDQDNSVSDSSFQMSLAEGPERKSTKQESAWLFRIWYNFDHNYLKPILTHSGPPLTATLPPCCGPLARFLTSPQAYENEGQLKDDDSDLILTDDINLAYGDITVNTDASGVHTSSTGFASGMISDDLDRELAYGDNEVVMRGTRLVLPMDDSEPPLKDPHIHF

>XP_020779128.1_Boleophthalmus

MSFTVTVRWARRAIGPRWLLVLVSLCVCVRVCRAASPPEEEDSAMENIVTEKRAEESHRQDSADLLIFILLLTLTILTIWLFKHRRFRFLHETGLAMIYGLLVGVVLRYGIHMPRDGTNVILNCNVNASPATVLVNVSGKFYEYTLKRVISSSGNKVDDMSDNENEMLRKVTFDPEVFFNILLPPIIFYAGYSLKRRHFFRNMGSILAYAFLGTVISAFIIGLLMYGTVTLMKHVGQLAGEFFFTDCLFFGAIVSATDPVTVLAIFHELQVDADLYALLFGESVLNDAVAVVLSSSIVAYQPAGDNSHTFEGMALLKSFGMFLGVFSGSFALGVATGIVTALVTKFTKLRDFQLLETALFFLMSWSTFLLAEACGFTGVVAVLFCGITQAHYTFNNLSPESQDRTKQLFELLNFLAENFIFSYMGLTLFTFQNHVFNPMFIVGAFLAVFLGRAANIYPLSLLLNLGRRHKIRSNFQHMMMFAGLRGAMTFALSIRDTATYARQMMFSTTLLIVFFTVWICGGGTTQMLSCQRIRVGVDSDQDNSVTEGPERKSTKQESAWLFRIWYNFDHNYLKPILTHSGPPLTATLPPCCGPLARFLTSPQAYENEGQLKDDDSDLILTDDINLAYGDITVNTDASGVHTSSTGFASGMISDDLDRELAYGDNEVVMRGTRLVLPMDDSEPPLKDPHIHF

>XP_020782715.1_Boleophthalmus

MERFLDVARSKIFILLPFLSTFLLVKSQGNAMNNVETERQAEESHRQDSANLLIFIMLLTLTILTIWLFKHRRFRFLHETGLAMIYGLLVGVILRFGVHVPQSTSDVVLSCAVNASPATLLVNVSGRFYEYTLKGEVSRAKGHQVQDDEMLRKVTFDPEVFFNILLPPIIFHAGYSLKRRHFFRNIGSILAYAFMGTVVSCFVIGLIMYGFVSFMKVVGQLGGDFYFTDCLFFGAIVSATDPVTVLAIFNELKVDVDLYALLFGESVLNDAVAIVLSSSIVAYQPAGDNSHSFEAMAMLKSFGVFLGVFSGSFALGVATGVVTAFVTKFTKLRDFPLLETALFFLMSWSTFLLAEACGFTGVVAVLFCGITQAHYTYNNLSPESQDRTKQLFELLNFLAENFIFSYMGLTLFSFQSHVFNPLFIIGAFLAVFLGRAANIYPLXXXXXXXXRNKIGYNFQHVMMFAGLRGAMTFALSIRDTATYARQMMFSTTLLIVFFTVWVCGGGTMPMLSFMSIPVGVDSDQENTSATMLDGSQRRNTKHESAWPFRIWYNFDHNYLKPLLTHSGPPLTATLPSCCGPLARCLTSPQAYENEGQLHDDDDFVLNDASVSSMYADVTVSTDASGSRTINHKRSSTGAGFPDDGLDNDLALSENEVAIRGTRLVLPMDDPLDAPTTVTLPPPPSPPKSDPRRHRL

***Homo sapiens***

>sp|Q6AI14|SL9A4_HUMAN NHE4_Homo

MALQMFVTYSPWNCLLLLVALECSEASSDLNESANSTAQYASNAWFAAASSEPEEGISVF

ELDYDYVQIPYEVTLWILLASLAKIGFHLYHRLPGLMPESCLLILVGALVGGIIFGTDHK

SPPVMDSSIYFLYLLPPIVLEGGYFMPTRPFFENIGSILWWAVLGALINALGIGLSLYLI

CQVKAFGLGDVNLLQNLLFGSLISAVDPVAVLAVFEEARVNEQLYMMIFGEALLNDGITV

VLYNMLIAFTKMHKFEDIETVDILAGCARFIVVGLGGVLFGIVFGFISAFITRFTQNISA

IEPLIVFMFSYLSYLAAETLYLSGILAITACAVTMKKYVEENVSQTSYTTIKYFMKMLSS

VSETLIFIFMGVSTVGKNHEWNWAFICFTLAFCQIWRAISVFALFYISNQFRTFPFSIKD

QCIIFYSGVRGAGSFSLAFLLPLSLFPRKKMFVTATLVVIYFTVFIQGITVGPLVRYLDV

KKTNKKESINEELHIRLMDHLKAGIEDVCGHWSHYQVRDKFKKFDHRYLRKILIRKNLPK

SSIVSLYKKLEMKQAIEMVETGILSSTAFSIPHQAQRIQGIKRLSPEDVESIRDILTSNM

YQVRQRTLSYNKYNLKPQTSEKQAKEILIRRQNTLRESMRKGHSLPWGKPAGTKNIRYLS

YPYGNPQSAGRDTRAAGFSDDDSSDPGSPSITFSACSRIGSLQKQEAQEIIPMKSLHRGR

KAFSFGYQRNTSQEEYLGGVRRVALRPKPLFHAVDEEGESGGESEGKASLVEVRSRWTAD

HGHGRDHHRSHSPLLQKK

>sp|P19634|SL9A1_HUMAN NHE1_Homo

MVLRSGICGLSPHRIFPSLLVVVALVGLLPVLRSHGLQLSPTASTIRSSEPPRERSIGDV

TTAPPEVTPESRPVNHSVTDHGMKPRKAFPVLGIDYTHVRTPFEISLWILLACLMKIGFH

VIPTISSIVPESCLLIVVGLLVGGLIKGVGETPPFLQSDVFFLFLLPPIILDAGYFLPLR

QFTENLGTILIFAVVGTLWNAFFLGGLMYAVCLVGGEQINNIGLLDNLLFGSIISAVDPV

AVLAVFEEIHINELLHILVFGESLLNDAVTVVLYHLFEEFANYEHVGIVDIFLGFLSFFV

VALGGVLVGVVYGVIAAFTSRFTSHIRVIEPLFVFLYSYMAYLSAELFHLSGIMALIASG

VVMRPYVEANISHKSHTTIKYFLKMWSSVSETLIFIFLGVSTVAGSHHWNWTFVISTLLF

CLIARVLGVLGLTWFINKFRIVKLTPKDQFIIAYGGLRGAIAFSLGYLLDKKHFPMCDLF

LTAIITVIFFTVFVQGMTIRPLVDLLAVKKKQETKRSINEEIHTQFLDHLLTGIEDICGH

YGHHHWKDKLNRFNKKYVKKCLIAGERSKEPQLIAFYHKMEMKQAIELVESGGMGKIPSA

VSTVSMQNIHPKSLPSERILPALSKDKEEEIRKILRNNLQKTRQRLRSYNRHTLVADPYE

EAWNQMLLRRQKARQLEQKINNYLTVPAHKLDSPTMSRARIGSDPLAYEPKEDLPVITID

PASPQSPESVDLVNEELKGKVLGLSRDPAKVAEEDEDDDGGIMMRSKETSSPGTDDVFTP

APSDSPSSQRIQRCLSDPGPHPEPGEGEPFFPKGQ

>sp|Q92581|SL9A6_HUMAN NHE6_Homo

MARRGWRRAPLRRGVGSSPRARRLMRPLWLLLAVGVFDWAGASDGGGGEARAMDEEIVSE

KQAEESHRQDSANLLIFILLLTLTILTIWLFKHRRARFLHETGLAMIYGLLVGLVLRYGI

HVPSDVNNVTLSCEVQSSPTTLLVTFDPEVFFNILLPPIIFYAGYSLKRRHFFRNLGSIL

AYAFLGTAISCFVIGSIMYGCVTLMKVTGQLAGDFYFTDCLLFGAIVSATDPVTVLAIFH

ELQVDVELYALLFGESVLNDAVAIVLSSSIVAYQPAGDNSHTFDVTAMFKSIGIFLGIFS

GSFAMGAATGVVTALVTKFTKLREFQLLETGLFFLMSWSTFLLAEAWGFTGVVAVLFCGI

TQAHYTYNNLSTESQHRTKQLFELLNFLAENFIFSYMGLTLFTFQNHVFNPTFVVGAFVA

IFLGRAANIYPLSLLLNLGRRSKIGSNFQHMMMFAGLRGAMAFALAIRDTATYARQMMFS

TTLLIVFFTVWVFGGGTTAMLSCLHIRVGVDSDQEHLGVPENERRTTKAESAWLFRMWYN

FDHNYLKPLLTHSGPPLTTTLPACCGPIARCLTSPQAYENQEQLKDDDSDLILNDGDISL

TYGDSTVNTEPATSSAPRRFMGNSSEDALDRELAFGDHELVIRGTRLVLPMDDSEPPLNL

LDNTRHGPA

>sp|Q8IVB4|SL9A9_HUMAN NHE9_Homo

MERQSRVMSEKDEYQFQHQGAVELLVFNFLLILTILTIWLFKNHRFRFLHETGGAMVYGL

IMGLILRYATAPTDIESGTVYDCVKLTFSPSTLLVNITDQVYEYKYKREISQHNINPHQG

NAILEKMTFDPEIFFNVLLPPIIFHAGYSLKKRHFFQNLGSILTYAFLGTAISCIVIGLI

MYGFVKAMIHAGQLKNGDFHFTDCLFFGSLMSATDPVTVLAIFHELHVDPDLYTLLFGES

VLNDAVAIVLTYSISIYSPKENPNAFDAAAFFQSVGNFLGIFAGSFAMGSAYAIITALLT

KFTKLCEFPMLETGLFFLLSWSAFLSAEAAGLTGIVAVLFCGVTQAHYTYNNLSSDSKIR

TKQLFEFMNFLAENVIFCYMGLALFTFQNHIFNALFILGAFLAIFVARACNIYPLSFLLN

LGRKQKIPWNFQHMMMFSGLRGAIAFALAIRNTESQPKQMMFTTTLLLVFFTVWVFGGGT

TPMLTWLQIRVGVDLDENLKEDPSSQHQEANNLDKNMTKAESARLFRMWYSFDHKYLKPI

LTHSGPPLTTTLPEWCGPISRLLTSPQAYGEQLKEDDVECIVNQDELAINYQEQASSPCS

PPARLGLDQKASPQTPGKENIYEGDLGLGGYELKLEQTLGQSQLN

>sp|Q96T83|SL9A7_HUMAN NHE7_Homo

MEPGDAARPGSGRATGAPPPRLLLLPLLLGWGLRVAAAASASSSGAAAEDSSAMEELATE

KEAEESHRQDSVSLLTFILLLTLTILTIWLFKHRRVRFLHETGLAMIYGLIVGVILRYGT

PATSGRDKSLSCTQEDRAFSTLLVNVSGKFFEYTLKGEISPGKINSVEQNDMLRKVTFDP

EVFFNILLPPIIFHAGYSLKKRHFFRNLGSILAYAFLGTAVSCFIIGNLMYGVVKLMKIM

GQLSDKFYYTDCLFFGAIISATDPVTVLAIFNELHADVDLYALLFGESVLNDAVAIVLSS

SIVAYQPAGLNTHAFDAAAFFKSVGIFLGIFSGSFTMGAVTGVNANVTKFTKLHCFPLLE

TALFFLMSWSTFLLAEACGFTGVVAVLFCGITQAHYTYNNLSVESRSRTKQLFEVLHFLA

ENFIFSYMGLALFTFQKHVFSPIFIIGAFVAIFLGRAAHIYPLSFFLNLGRRHKIGWNFQ

HMMMFSGLRGAMAFALAIRDTASYARQMMFTTTLLIVFFTVWIIGGGTTPMLSWLNIRVG

VEEPSEEDQNEHHWQYFRVGVDPDQDPPPNNDSFQVLQGDGPDSARGNRTKQESAWIFRL

WYSFDHNYLKPILTHSGPPLTTTLPAWCGLLARCLTSPQVYDNQEPLREEDSDFILTEGD

LTLTYGDSTVTANGSSSSHTASTSLEGSRRTKSSSEEVLERDLGMGDQKVSSRGTRLVFP

LEDNA

>sp|Q86UD5|SL9B2_HUMAN NHE9B2_Homo

MGDEDKRITYEDSEPSTGMNYTPSMHQEAQEETVMKLKGIDANEPTEGSILLKSSEKKLQ

ETPTEANHVQRLRQMLACPPHGLLDRVITNVTIIVLLWAVVWSITGSECLPGGNLFGIII

LFYCAIIGGKLLGLIKLPTLPPLPSLLGMLLAGFLIRNIPVINDNVQIKHKWSSSLRSIA

LSIILVRAGLGLDSKALKKLKGVCVRLSMGPCIVEACTSALLAHYLLGLPWQWGFILGFV

LGAVSPAVVVPSMLLLQGGGYGVEKGVPTLLMAAGSFDDILAITGFNTCLGIAFSTGSTV

FNVLRGVLEVVIGVATGSVLGFFIQYFPSRDQDKLVCKRTFLVLGLSVLAVFSSVHFGFP

GSGGLCTLVMAFLAGMGWTSEKAEVEKIIAVAWDIFQPLLFGLIGAEVSIASLRPETVGL

CVATVGIAVLIRILTTFLMVCFAGFNLKEKIFISFAWLPKATVQAAIGSVALDTARSHGE

KQLEDYGMDVLTVAFLSILITAPIGSLLIGLLGPRLLQKVEHQNKDEEVQGETSVQV

>sp|Q14940|SL9A5_HUMAN NHE5_Homo

MLRAALSLLALPLAGAAEEPTQKPESPGEPPPGLELFRWQWHEVEAPYLVALWILVASLA

KIVFHLSRKVTSLVPESCLLILLGLVLGGIVLAVAKKAEYQLEPGTFFLFLLPPIVLDSG

YFMPSRLFFDNLGAILTYAVVGTLWNAFTTGAALWGLQQAGLVAPRVQAGLLDFLLFGSL

ISAVDPVAVLAVFEEVHVNETLFIIVFGESLLNDAVTVVLYKVCNSFVEMGSANVQATDY

LKGVASLFVVSLGGAAVGLVFAFLLALTTRFTKRVRIIEPLLVFLLAYAAYLTAEMASLS

AILAVTMCGLGCKKYVEANISHKSRTTVKYTMKTLASCAETVIFMLLGISAVDSSKWAWD

SGLVLGTLIFILFFRALGVVLQTWVLNQFRLVPLDKIDQVVMSYGGLRGAVAFALVILLD

RTKVPAKDYFVATTIVVVFFTVIVQGLTIKPLVKWLKVKRSEHHKPTLNQELHEHTFDHI

LAAVEDVVGHHGYHYWRDRWEQFDKKYLSQLLMRRSAYRIRDQIWDVYYRLNIRDAISFV

DQGGHVLSSTGLTLPSMPSRNSVAETSVTNLLRESGSGACLDLQVIDTVRSGRDREDAVM

HHLLCGGLYKPRRRYKASCSRHFISEDAQERQDKEVFQQNMKRRLESFKSTKHNICFTKS

KPRPRKTGRRKKDGVANAEATNGKHRGLGFQDTAAVILTVESEEEEEESDSSETEKEDDE

GIIFVARATSEVLQEGKVSGSLEVCPSPRIIPPSPTCAEKELPWKSGQGDLAVYVSSETT

KIVPVDMQTGWNQSISSLESLASPPCNQAPILTCLPPHPRGTEEPQVPLHLPSDPRSSFA

FPPSLAKAGRSRSESSADLPQQQELQPLMGHKDHTHLSPGTATSHWCIQFNRGSRL

>sp|Q4ZJI4|SL9B1_HUMAN NHE9B1_Homo

MHTTESKNEHLEDENFQTSTTPQSLIDPNNTAQEETKTVLSDTEEIKPQTKKETYISCPL

RGVLNVIITNGVILFVIWCMTWSILGSEALPGGNLFGLFIIFYSAIIGGKILQLIRIPLV

PPLPPLLGMLLAGFTIRNVPFINEHVHVPNTWSSILRSIALTIILIRAGLGLDPQALRHL

KVVCFRLAVGPCLMEASAAAVFSHFIMKFPWQWAFLLGFVLGAVSPAVVVPYMMVLQENG

YGVEEGIPTLLMAASSMDDILAITGFNTCLSIVFSSGGILNNAIASIRNVCISLLAGIVL

GFFVRYFPSEDQKKLTLKRGFLVLTMCVSAVLGSQRIGLHGSGGLCTLVLSFIAGTKWSQ

EKMKVQKIITTVWDIFQPLLFGLVGAEVSVSSLESNIVGISVATLSLALCVRILTTYLLM

CFAGFSFKEKIFIALAWMPKATVQAVLGPLALETARVSAPHLEPYAKDVMTVAFLAILIT

APNGALLMGILGPKMLTRHYDPSKIKLQLSTLEHH

>sp|Q9UBY0|SL9A2_HUMAN NHE2_Homo

MEPLGNWRSLRAPLPPMLLLLLLQVAGPVGALAETLLNAPRAMGTSSSPPSPASVVAPGT

TLFEESRLPVFTLDYPHVQIPFEITLWILLASLAKIGFHLYHKLPTIVPESCLLIMVGLL

LGGIIFGVDEKSPPAMKTDVFFLYLLPPIVLDAGYFMPTRPFFENIGTIFWYAVVGTLWN

SIGIGVSLFGICQIEAFGLSDITLLQNLLFGSLISAVDPVAVLAVFENIHVNEQLYILVF

GESLLNDAVTVVLYNLFKSFCQMKTIETIDVFAGIANFFVVGIGGVLIGIFLGFIAAFTT

RFTHNIRVIEPLFVFLYSYLSYITAEMFHLSGIMAITACAMTMNKYVEENVSQKSYTTIK

YFMKMLSSVSETLIFIFMGVSTVGKNHEWNWAFVCFTLAFCLMWRALGVFVLTQVINRFR

TIPLTFKDQFIIAYGGLRGAICFALVFLLPAAVFPRKKLFITAAIVVIFFTVFILGITIR

PLVEFLDVKRSNKKQQAVSEEIYCRLFDHVKTGIEDVCGHWGHNFWRDKFKKFDDKYLRK

LLIRENQPKSSIVSLYKKLEIKHAIEMAETGMISTVPTFASLNDCREEKIRKVTSSETDE

IRELLSRNLYQIRQRTLSYNRHSLTADTSERQAKEILIRRRHSLRESIRKDSSLNREHRA

STSTSRYLSLPKNTKLPEKLQKRRTISIADGNSSDSDADAGTTVLNLQPRARRFLPEQFS

KKSPQSYKMEWKNEVDVDSGRDMPSTPPTPHSREKGTQTSGLLQQPLLSKDQSGSEREDS

LTEGIPPKPPPRLVWRASEPGSRKARFGSEKP

>sp|P48764|SL9A3_HUMAN NHE3_Homo

MWGLGARGPDRGLLLALALGGLARAGGVEVEPGGAHGESGGFQVVTFEWAHVQDPYVIAL

WILVASLAKIGFHLSHKVTSVVPESALLIVLGLVLGGIVWAADHIASFTLTPTVFFFYLL

PPIVLDAGYFMPNRLFFGNLGTILLYAVVGTVWNAATTGLSLYGVFLSGLMGDLQIGLLD

FLLFGSLMAAVDPVAVLAVFEEVHVNEVLFIIVFGESLLNDAVTVVLYNVFESFVALGGD

NVTGVDCVKGIVSFFVVSLGGTLVGVVFAFLLSLVTRFTKHVRIIEPGFVFIISYLSYLT

SEMLSLSAILAITFCGICCQKYVKANISEQSATTVRYTMKMLASSAETIIFMFLGISAVN

PFIWTWNTAFVLLTLVFISVYRAIGVVLQTWLLNRYRMVQLEPIDQVVLSYGGLRGAVAF

ALVVLLDGDKVKEKNLFVSTTIIVVFFTVIFQGLTIKPLVQWLKVKRSEHREPRLNEKLH

GRAFDHILSAIEDISGQIGHNYLRDKWSHFDRKFLSRVLMRRSAQKSRDRILNVFHELNL

KDAISYVAEGERRGSLAFIRSPSTDNVVNVDFTPRSSTVEASVSYLLRENVSAVCLDMQS

LEQRRRSIRDAEDMVTHHTLQQYLYKPRQEYKHLYSRHELTPTEDEKQDREIFHRTMRKR

LESFKSTKLGLNQNKKAAKLYKRERAQKRRNSSIPNGKLPMESPAQNFTIKEKDLELSDT

EEPPNYDEEMSGGIEFLASVTKDTASDSPAGIDNPVFSPDEALDRSLLARLPPWLSPGET

VVPSQRARTQIPYSPGTFCRLMPFRLSSKSVDSFLQADGPEERPPAALPESTHM

>sp|Q9Y2E8|SL9A8_HUMAN NHE8_Homo

MGEKMAEEERFPNTTHEGFNVTLHTTLVVTTKLVLPTPGKPILPVQTGEQAQQEEQSSGM

TIFFSLLVLAICIILVHLLIRYRLHFLPESVAVVSLGILMGAVIKIIEFKKLANWKEEEM

FRPNMFFLLLLPPIIFESGYSLHKGNFFQNIGSITLFAVFGTAISAFVVGGGIYFLGQAD

VISKLNMTDSFAFGSLISAVDPVATIAIFNALHVDPVLNMLVFGESILNDAVSIVLTNTA

EGLTRKNMSDVSGWQTFLQALDYFLKMFFGSAALGTLTGLISALVLKHIDLRKTPSLEFG

MMIIFAYLPYGLAEGISLSGIMAILFSGIVMSHYTHHNLSPVTQILMQQTLRTVAFLCET

CVFAFLGLSIFSFPHKFEISFVIWCIVLVLFGRAVNIFPLSYLLNFFRDHKITPKMMFIM

WFSGLRGAIPYALSLHLDLEPMEKRQLIGTTTIVIVLFTILLLGGSTMPLIRLMDIEDAK

AHRRNKKDVNLSKTEKMGNTVESEHLSELTEEEYEAHYIRRQDLKGFVWLDAKYLNPFFT

RRLTQEDLHHGRIQMKTLTNKWYEEVRQGPSGSEDDEQELL

**Na^+^/K^+^-ATPase alfa-subunits (NKA)**

***Round goby (Neogobius melanostomus)***

>NEME_00025176_Neogobius

MEELNRKYGTDITKGLTGARVKEILLRDGPNSLSPPVTTAEWVKFCKHLFGGFSTLLWIG

AILCFFAYSIQAASEYEPVKDNLYLGIVLSAVVIITACFSYYQEAKSSKIMDSFKNMVPQ

QALVVRDGEKKCINAEEVVVGDLVEVKGGDRIPADLRIISAHGCKVDNSSLTGESEPQIR

SPEFSNENPLETRNIAFFSTNCVEGTARGIVISTGDRTVMGRIATLASSLGSGQTPIAIE

IEHFIHIITSVAMFLGVSSLIISLILGYGWLESVVYLIGIIVANVPEGLLATVTVCLTLT

AKRMAKKNCLVKNLEAVETLGSTSTICSDKTGTLTQNRMTVAHMWFDNQIHEADTTEDQR

GNSFDRSSATWAALSRIAGLCNRAVFLTDQNNVPILKRDVAGDASEAALLKCIELCCGSV

AGIREKYPNVAEIPFNSTNKYQLSIHKNSTSGETKHLLVMKGAPERILDRCSTIVLQGKE

QPLDDEMKDAFQNAYVELGGLGERVLGFCHFNLPDDQFPEGFAFDAEEVNFPTENLCFVG

LMAMIDPPRAAVPDAVGKCRSAGIKVIMVTGDHPITAKAIAKGVGIISEGNETVEDIAIR

LNVPVSDINPSGVGVLRRDAKACVVHGGDLKDFTSEELDEILKNHTEIVFARTSPQQKLI

IVEGCQRQGAIVAVTGDGVNDSPALKKADIGIAMGIMGSDVSKQAADMILLDDNFASIVT

GVEEGRLIFDNLKKSIAYTLTSNVPEITPFLMFIIANIPLPLGTVTILCIDLGTDMVPAI

SLAYEHAESDIMKRQPRNPKKDKLVNERMISKGYGQIGIMQAAGGFFTYFVILAENGFLP

IDLLGIRVRWDDKYNNDLEDSYGQQWTYEQRKIVEFTCHTAFFTSIVIVQWANLIICKTR

RNSIVQQGMTNRILIFGLFEETALAAFLSYCPGTDMGLRMFPLKPSWWFCALPYSLLIVL

YDEARRYLLRRNPGGECCSFCWVEMETYY

>NEME_00012929_Neogobius

LLLRRVGCGAHPSHYAQLYLGIVLSAVVIITGCFSYFQEAKSSKIMESFKDMVPQQALVI

REGEKMQINAEQVVAGDLVEVKGGDRIPADLRIISSHGCKVDNSSLTGESEPQTRSPDCT

HDNPLETRNIAFFSTNCVEGTARGIVVCTGDRTVMGRIATLTSGLETGKTPIAKEIEHFI

HIITGVAVFLGVSFFVLSIVLGYTWLEAVIFLIGIIVANVPEGLLATVTVCLTLTAKRMA

RKNCLVKNLEAVETLGSTSTICSDKTGTLTQNRMTVAHMWFDNQIHEADTTEDQSGAYEG

RKGAGRGGRVQLLLHSDVCPYLTGCSFDKSSTTWVSLARIATLCNRAVFKAGQDSLPILK

RDVAGDASESALLKCIELSCGPVKVMRDKNKKVAEIPFNSTNKYQVSIHETDDENDNRYL

MVMKGAPERILERCSTIMLQGKEQPMDDELKEAFQNAYLELGGLGERVLGFCHLFMSEEK

YPKGFAFDTDDVNFQIDDMCFVGLMSMIDPPRAAVPDAVGKCRSAGIKVIMVTGDHPITA

KAIAKGVGIISEGNETVEDIAARLNIPVNQVNPSDAKACVIHGTDLKDLSQDQMDDILKN

HTEIVFARTSPQQKLIIVEGCQRQGAIVAVTGDGVNDSPALKKADIGVAMGISGSDVSKQ

AADMILLDDNFASIVTGRLILTGRLIFDNLKKSIAYTLTSNIPEITPFLFFILVNIPLPL

GTITILCIDLGTDMVPAISLAYEAPESDIMKRQPRNPLRDKLVNERLISIAYGQIGMIQA

LGGFFAYFVIMAENGFLPSHLVGIRLKWDDRSTNDLEDSYGQQWTYEQRKIVEYTCHTAF

FVSIVVVQWADVIICKTRRNSVFQQGMRNKILIFGLFEETALAAFLSYTPGMDIALRMYP

LKPSWWFCAFPYSFLIFVYDEIRKLILRRNPGGWVEKETYY

>NEME_00002724_Neogobius

MWRPHGPWPPNFPDSLGTGRRSEPSGCGRGISEDLWVTVTNLGQKERYMLGGVSEEYLET

DLTRCHLGRIHCHSVKNQPMGVWSCLGEKRCTSKLLCHVPREGGREGGGLSERCPNPPTN

ELFTDAPKQPLLEERGDSEHVWVHEFKGSREDDHEIPIEELEMRYNTSVTKGMITTFAHQ

ILERDGPNELKPPKGTPEYVKFARQLAGGLQCLMWVAAVICFIAFGIEMAKGDITSFDNL

YLAITLIAVVVVTGCFGYYQEFKSTNIIASFKNLVPQQAVVIRDGQKNQINTNELVVGDL

VEIKGGDRVPADVRIITAQACKVDNSSLTGESEPQSRTPECTHENPLETRNIAFFSTTCL

EGVATGVIINTGDRTIIGRIASLASGVGNEKTPIAIEIEHFVDIIAGLAIFFGFTFFVVA

MFIGYAFLEAMIFFMAIVVAYVCLSLTAKRLARKNCVVKNLEAVETLGSTSVICSDKTGT

LTQNRMTVAHLWFDNQIHAADTTEDQSDTWRSLARVASLCNRATFRPDQEGIPIPRRIVV

GDASETALLKFTELTIGNIMDYRNRFKKVVEVPFNSTNKFQASCCTWTHLSVHELDDPLD

LRYLLVMKGAPERILERCSSILIQGQELPMDQQWKESFQTAYMDLGGLGSEYWKEFPRGY

SFDADEMNFTTSGLCFAGLISMIDPPRATVPDAVMKCRTAGIRVVMVTGDHPITARAIAA

NVGIISEGSETVEDIAQRKRIPVEQVNKRTSPQQKLIIVESCQRLGSIVAVTGDGVNDSP

ALKKADIGIAMGIAGSDAAKNAADMILLDDNFASIVTGVEQAGSLSHLHHRQRASASGCI

TILFIELATDIKAESDIMHLKPRNPRRDRLVNEALAAYSYFQIGVIQSFAGFTDYFAAMA

QEGWYPLLCVGLRSQWENVQMQDICQISDVLIRKTRRLSVFQQGLFRVQWWFIPVPYGIL

IFVYDEIRKLGVRRNPGNWGNKAARLQLYSTFQMAILPSPRAEPLSTYRPSAEHMVWIQS

VCACSSLSTACVSESSTSNRLLPRPSPRSPPLLKARQPSQI

>NEME_00036611_Neogobius

MVPAISLAYEAAESDIMKRQPRNPKTDKLVNERLISIAYGQIGMMQATAGFFTYFVILAE

NGFLPMDLLGIRVNWDDRYNNDLEDSYGQQWTYESRKIVEFTCHTAFFASIVIVQNRILI

FGLFEETALAAFLSYCPGMDVALRMYPLKPSWWFCAFPYSLLIFLYDEARRYLLRRNPGE

MGRTFSSGESALASGTPRLLKTRHGPGRLGGDGDVLLSPKFVSSLFLPHVTDLTNMLARP

KANVPSMDIFKKSISVFLNIVGPRGCQTNVFVLRYYLDEQTANAKRCSAEINRAPKPMFA

QAIQTEDFGEQDFSFSLNGGRRTRLFTFRFQLQRCYLKIPVLVFANKINAPSVEADGREQ

YELAATSEQGGKKKAKGKKKEKDMDELKKEVDMDDHKLTLDELNRKYSTDLTNLYLGVVL

SAVVIITGCFSYYQEAKSSKIMDSFKNLVPQQALVVRDGEKKCINAEEVVVGDLVEVKGG

DRIPADLRIISAHGCKVDNSSLTGESEPQTRTPDFSNENPLETRNIAFFSTNCVEVGRTP

ISIEIEHFIHIITGVAVFLGVSFFVLSLVLGYSWLEAVIFLIGIIVANVPEGLLATVTRM

AKKNCLVKNLEAVETLGSTSTICSDKTGTLTQNRMTVAHMWFDNQIHEADTTENQSGTSF

DRSSATWAALARIAGLCNRALSIHKNSTAEGESKRLLVMKGAPERILDRCATIMLQGKEQ

ALDDEMKDAFQNAYLELGGLGERVLGFCHYHLPDDQFPEDFAFDTEEVNFPTENLCFIGL

MSMIDPLEPLCPTPWANAEALASR

>NEME_00025762_Neogobius

MENSHTKVPAECLAYFGVNENTGLSPDQFKKNLEKYGYNGKSIWELVVEQFEDLLVRILL

LAACISFVLAWFEEGEETVTAFVEPLVILLILIANAIVGVWQERNAESAIEALKEYEPEM

GKVYRSDRKSVQMIKAREIVPGDIVEVSVGDKVPADIRLVTIKSTTLRVDQSILTGESVS

VIKHTEAVLDPRAVNQDKKNMLFSGTNIAAGKAIGVAVATGVATEIGKIRDQMAATEPEK

TPLQQKLDEFGEQLSKVITLICIAVWAINIGHFNDPVHGGSWIRGAVYYFKIAVALAVAA

IPEGLPAVITTCLALGTRRMAKKNAIVRSLPSVETLGCTSVICSDKTGTLTTNQMCVTKM

FVIKNVDGDHVDLDAFDISGSKYTPEGEVSQGGGKTNCSAYDGLVELSTICALCNDSSLD

YNESKKIFEKVGEATETALCCLVEKMNVFNSNVKNLSRIERANACCTVIKQLMKKKFTLE

FSRDRKSMSVYCTPAKGDGGAKMFVKVHICACRHHPRPLTNTIRDKILSVIKDWGTGRDT

LRCLALATRDSPLKPEEMVLEDSTKFVDYENDLTFVGCVGMLDPPRKEVTGSIQLCRDAG

IRVIMITGDNKGTAIAICRRIGIFGEDEDVEGKAYTGREFDDLPLTEQSEAVRRACCFAR

VEPAHKSKIVEFLQGHDDITAMTGDGVNDAPALKKAEIGIAMGSGTAVAKSASEMVLADD

NFSSIVAAVEEGRAIYNNMKQFIRYLISSNVGEVVCIFLTAALGLPEALIPVQLLWVNLV

TDGLPATALGFNPPDLDIMGKPPRSPKEALISGWLFFRYMAIGGYVGAATVGGAAWWFMY

DHTGPQITYYQLYSHFMQCHDENEDFAGLDCEIFEAYPPMTMALSVLVTIEMCNALNSLS

ENQSLVRMPPWSNFWLLSAMTLSILVDSVLTVLTAHPLSPQMIFKLTHLSVEQWMMVLKL

SFPVILIDEVLKFFARNYVDVSSQTAPPDINVTTEDFTSSAPENGTVDDTSLRTTTVSTG

TDDAPTVQPLITQGCLCDLTPDFCDIGCCCDTEDCGLANLSTVFTSCPQKAISGVCVEKW

LMFRANVDSSRVTVDDVILTYFSKTSARGLLRQSSPAVASALCTDRNPAKRTDEDMLRSG

SKTDVNAPSPFSLVPWWQREKDVKRNHTYGWNVYACQRREQRQQGARREEQKFCKTLLSI

SPSVVGSTCDIRVKNGSAFEGIFKTLSSRCELAVDAVHKRTENGGSTSAPPRREDITDTM

IFSPQDLVTMICRDVDLNYATRDTFTDTAISSTRVNGEHKEKVLQRWDGGENNGESYDLE

NDTSNGWDANEMFRFNEVKYGVTSTYDSSLAMYTVPLEKGNSDIYRQREARAARLASEIE

SSPQYRHRVNLENDEVKSEEDKYSAVVREGSDRERGRESPRERERGRDSPGTSSREGKYI

PLHQRQREINRDRAGGPPSHSRGGYRSTPPSSSSSSPRPSLPSASGPQPGTSPSERSSPL

SGRGGAYAPHHPQASPSPGPGSGPASPYTPASPGGSVPSTPTAAAPSSASPPAPHTNTVS

HSHSLPHSLSDGARPVNGVSTRTSPKAQRPPQSSRTVRTPNSHSQPTASRSPKSGSSQDT

PYLDTTSVSMPAQKTSGPAPLFPVDVNEILSAAAKERSSESPSGTDDGKSSKVQQRSQIE

ELRKFGKEFRLQPSGGSSLGAPATMTPSTGTDGSQQSGGKSQPDTATEPKSPLPTPVPSQ

PQPQPSPVPAEEPKDTAAIPGATTTATGSERQSPATPQPARTPGSDEGRSEPERSEGVPD

QVKKSTLNPNAKEFNPNPVKTPMPMTKPNTTPTPPRPTPPSPVVLSHAAAGQGQLYNSPY

LYQVSQIHSVQYARTKGSVVAPRSDHAASGPPLIQPASAAGAPLVASPYPQSYLQYNPQQ

YSQQQVIQAMSPYAGQHMYPMLQGGARMIGQGGGPHAQALGPPGAPSSRHKETVPRVHSK

AFMLHNPFNTTRVQSISHSHLAPRQETRPNHNMLLPVLDRMHSQVLSPVVVPFGSSLSAY

PSQHAPGHTSPQGSYSIPGYSIHQGIPPNYPLSQIAQAHVQGAMSGPTIQAATDSHS

>NEME_00025177_Neogobius

MPAPPGALNVLRGASSVRLGVFVPLLCRVSIDLRAAAVFNGLAYSKVMGGSEGGHSSWWS

SVGAVPGGLQWWAIPQTVSVRAASPGGVISLEFMATTQDGREQYELAATSEQGGKKKAKG

KKKGERHGRTEERGGHAKGQGNVSDTMFFPPFFPQDDHKLTLDELNRKYSTDLTNGLTGA

KAAEDLARDGPNALTPPPTTPEWVKFCKQLFVFPKQALVVRDGEKKCINAKEVVVGDLVE

VKGGDRIPADLRIISAHGCKVDNSSLTGTARGIVISTGDRTVMGRIATLASGLEVGRTPI

SIEHFIHIITGVAVFLGVSFFVLSLVLGYSWLEAVIFLIGIIVANVPEGLLATVTVCLTL

TAKRMAKKNCLVKNLEAVETLGSTSTICSDKTGTLTQNRMTVAHMWFDNQIHEADTTENQ

SGTSFDRSSATWAALARIAGLCNRAVFLAEQSNLPILKLSIHKNSTAEGESKRLLVMKGA

PERILDRCATIMLQGKEQPLDDEMKDAFQNAYLELGGLGERVLGFCHYHLPDDQFPEDFA

FDTEEVNFPTENLCFIGLMSMIDPPRAAVPDAVGKCRSAGIKVIMVTGDHPITAKAIAKG

VGIISEGNETVEDIAACLNIPVNEVNPRDAKACVVHGGDLKDLSAEQLDDILKYHTEIVF

ARTSPQQKFIIVEGCQRQGAIVAVTGDGVNDSSALKKADIGVAMGIADSDVSKQAADMIL

LDDNFASIGPEWKRDV

**zebrafish (Danio rerio),**

>tr|Q9DGL6|Q9DGL6_DANRE Sodium/potassium-transporting ATPase subunit alpha OS=Danio rerio OX=7955 GN=atp1a1a.1 PE=1 SV=1

MGRGEGREQYELAATSEQGGKKSKSKGKKEKDKDMDELKKEVDLDDHKLSLDELTRKYNT

DLTRGLSGTRAKEILARDGPNALTPPPTTPEWVKFCKQLFGGFSTLLWIGAILCFLAYGI

LAASEEEPANDNLYLGIVLSAVVMITGCFSYYQEAKSSKIMDSFKNLVPQQALVIRDGEK

KNINAEEVAVGDLVEVKGGDRIPADLRIISAHGCKVDNSSLTGESEPQTRTPDFSNDNPL

ETRNIAFFSTNCVEGTARGIVINTGDRTVMGRIATLASGLEVGRTPISIEIEHFIHIITG

VAVFLGVSFFILSLILGYSWLEAVIFLIGIIVANVPEGLLATVTVCLTLTAKRMAKKNCL

VKNLEAVETLGSTSTICSDKTGTLTQNRMTVAHMWFDNQIHEADTTENQSGTSFDRSSAT

WAALARVAGLCNRAVFLAEQENIPILKRDVAGDASESALLKCIELCCGSVKEMREKYNKI

SEIPFNSTNKYQLSIHQNPNSNNTESKHLLVMKGAPERILDRCSSILIQGKEQPLDDEMK

DAFQNAYLELGGLGERVLGFCHFNLPDEQFPEDFQFDTEEVNFPTENLCFIGLMSMIDPP

RAAVPDAVGKCRSAGIKVIMVTGDHPITAKAIAKGVGIISEGNETVEDIAARLNIPINEV

NPRDAKACVIHGGDLKDLSPEQLDDVLKHHTEIVFARTSPQQKLIIVEGCQRQGAIVAVT

GDGVNDSPALKKADIGVAMGIAGSDVSKQAADMILLDDNFASIVTGVEEGRLIFDNLKKS

IAYTLTSNIPEITPFLLFIIANIPLPLGTVTILCIDLGTDMVPAISLAYEAAESDIMKRQ

PRNPKTDKLVNERLISIAYGQIGMIQALAGFFTYFVILAENGFLPSSLLGIRVFWDDKYV

NDLEDSYGQQWTYEQRKIVEFTCHTAFFTSIVIVQWADLIICKTRRNSVFQQGMKNKILI

FGLFEETALAAFLSYCPGMDVALRMYPLKPNWWFCAFPYSLLIFIYDEIRKLIIRRSPGG

WVERETYY

>tr|A0A0R4IJ10|A0A0R4IJ10_DANRE Sodium/potassium-transporting ATPase subunit alpha OS=Danio rerio OX=7955 GN=atp1a1a.1 PE=1 SV=1

MGRGEGREQYELAATSEQGGKKSKSKGKKEKDKDMDELKKEVDLDDHKLSLDELTRKYNT

DLTRGLSGTRAKEILARDGPNALTPPPTTPEWVKFCKQLFGGFSTLLWIGAILCFLAYGI

LAASEEEPANDNLYLGIVLSAVVMITGCFSYYQEAKSSKIMDSFKNLVPQQALVIRDGEK

KNINAEEVAVGDLVEVKGGDRIPADLRIISAHGCKVDNSSLTGESEPQTRTPDFSNDNPL

ETRNIAFFSTNCVEGTARGIVINTGDRTVMGRIATLASGLEVGRTPISIEIEHFIHIITG

VAVFLGVSFFILSLILGYSWLEAVIFLIGIIVANVPEGLLATVTVCLTLTAKRMAKKNCL

VKNLEAVETLGSTTTICSDKTGTLTQNRMTVAHMWFDSQIHEADTTENQSGTSFDRSSPT

WAALARVAGLCNRAVFRAEQSHLPVLNRETAGDASESALLKCIELCCGSVIEMREKYRKI

CEIPFNSTNKYQLSVHKDPSSSGTKHLLVMKGAPERILDRCSTILINGKEQPMDDENKDS

FQSAYVELGGLGERVLGFCQFNLPDDQFPEGFAFDPEDVNFPTENLCFLGLMSMIDPPRA

AVPDAVAKCRSAGIKVIMVTGDHPITAKAIAKGVGIISEGNETVEDIAARMNIPVGEVNP

REAKACVVHGGELKNMNDSDLDEILRYHTEIVFARTSPQQKLIIVEGCQRQGAIVAVTGD

GVNDSPALKKADIGVAMGISGSDVSKQAADMILLDDNFASIVTGVEEGRLIFDNLKKSIA

YTLTSKIPEMSPFLMFVLVGIPLPLGTVTILCIDLGTDMVPAISLAYETAESDIMKRQPR

NAATDRLVNERLISVSYGQIGMIQAVGGFFTYFVILAENGFLPYDLVGIRVGWEDRFLND

LEDSYGQQWTYESRKIVEYTCHTAFFASIVIVQWTDLLICKTRRLSIFQQGMKNRVLTFG

LLEETALAAFLSYCPGMEVAVRMYPLKPLWWFCAFPYSLLIFIYDEVRKYILRRNPGGWV

ERETYY

>tr|Q7ZU25|Q7ZU25_DANRE Sodium/potassium-transporting ATPase subunit alpha OS=Danio rerio OX=7955 GN=atp1a1a.1 PE=2 SV=1

MGRGEGREQYELAATSEQGGKKSKSKGKKEKDKDMDELKKEVDLDDHKLSLDELTRKYNT

DLTRGLSGTRAKEILARDGPNALTPPPTTPEWVKFCKQLFGGFSTLLWIGAILCFLAYGI

LAASEEEPANDNLYLGIVLSAVVMITGCFSYYQEAKSSKIMDSFKNLVPQQALVIRDGEK

KNINAEEVAVGDLVEVKGGDRIPADLRIISAHGCKVDNSSLTGESEPQTRTPDFSNDNPL

ETRNIAFFSTNCVEGTARGIVINAGDRTVMGRIATLASGLEVGRTPISIEIEHFIHIITG

VAVFLGVSFFILSLILGYSWLEAVIFLIGIIVANVPEGLLATVTVCLTLTAKRMAKKNCL

VKNLEAVETLGSTSTICSDKTGTLTQNRMTVAHMWFDNQIHEADTTENQSGTSFDRSSAT

WAALARVAGLCNRAVFLAEQENIPILKRDVAGDASESALLKCIELCCGSVKEMREKYNKI

SEIPFNSTNKYQLSIHQNPNSNNTESKHLLVMKGAPERILDRCSSILIQGKEQPLDDEMK

DAFQNAYLELGGLGERVLGFCHFNLPDEQFPEDFQFDTEEVNFPTENLCFIGLMSMIDPP

RAAVPDAVGKCRSAGIKVIMVTGDHPITAKAIAKGVGIISEGNETVEDIAARLNIPINEV

NPRGAKACVIHGGDLKDLSPEQLDDVLKHHTEIVFARTSPQQKLIIVEGCQRQGAIVAVT

GDGVNDSPALKKADIGVAMGIAGSDVSKQAADMILLDDNFASIVTGVEEGRLIFDNLKKS

IAYTLTSNIPEITPFLLFIIANIPLPLGTVTILCIDLGTDMVPAISLAYEAAESDIMKRQ

PRNPKTDKLVNERLISIAYGQIGMIQALAGFFTYFVILAENGFLPSSLLGIRVFWDDKYV

NDLEDSYGQQWTYEQRKIVEFTCHTAFFTSIVIVQWADLIICKTRRNSVFQQGMKNKILI

FGLFEETALAAFLSYCPGMDVALRMYPLKPNWWFCAFPYSLLIFIYDEIRKLIIRRSPGG

WVERETYY

>tr|A0A0R4IRT1|A0A0R4IRT1_DANRE Sodium/potassium-transporting ATPase subunit alpha OS=Danio rerio OX=7955 GN=atp1a1a.1 PE=1 SV=1

MGLGTGNDKYKLAATSEDEAKAPKKGKKKQKDMDELKKEVDLDDHKLTLDELHRKYGTDL

TRGLSSSRAREVLARDGPNALTPPPTTPEWVKFCKQLFGGFSTLLWIGAILCFLAYGIQA

ASEDDPTNDNLYLGIVLAGVVIITGCFSYYQEAKSSKIMESFKNLVPQQALVVRDGEKKS

INAEEVVAGDLVEVKGGDRIPADLRIISAHGCKVDNSSLTGESEPQTRTPDFSNDNPLET

RNIAFFSTNCVEGTARGIVINTGDRTVMGRIATLASSLEGGQTPIAKEIEHFIHIITGVA

VFLGVSFFILSLILGYGWLEAVIFLIGIIVANVPEGLLATVTVCLTLTAKRMAKKNCLVK

NLEAVETLGSTSTICSDKTGTLTQNRMTVAHMWFDNQIHEADTTENQSGTSFDRSSATWA

ALARVAGLCNRAVFLAEQENIPILKRDVAGDASESALLKCIELCCGSVKEMREKYNKISE

IPFNSTNKYQLSIHQNPNSNNTESKHLLVMKGAPERILDRCSSILIQGKEQPLDDEMKDA

FQNAYLELGGLGERVLGFCHFNLPDEQFPEDFQFDTEEVNFPTENLCFIGLMSMIDPPRA

AVPDAVGKCRSAGIKVIMVTGDHPITAKAIAKGVGIISEGNETVEDIAARLNIPINEVNP

RDAKACVIHGGDLKDLSPEQLDDVLKHHTEIVFARTSPQQKLIIVEGCQRQGAIVAVTGD

GVNDSPALKKADIGVAMGIAGSDVSKQAADMILLDDNFASIVTGVEEGRLIFDNLKKSIA

YTLTSNIPEITPFLLFIIANIPLPLGTVTILCIDLGTDMVPAISLAYEAAESDIMKRQPR

NPKTDKLVNERLISIAYGQIGMIQALAGFFTYFVILAENGFLPSSLLGIRVFWDDKYVND

LEDSYGQQWTYEQRKVLEYTCHTAYFTSIVIMQWTTLLVCKSRRLSLAKQGMKNRVLTFS

LFEETAIAAFLSYCPGMDIAVRMYPLKPMWWFCAFPYMILIFIYDEVRKHFIRQNPGGWV

EQETYY

>tr|Q9DGL5|Q9DGL5_DANRE Sodium/potassium-transporting ATPase subunit alpha OS=Danio rerio OX=7955 GN=atp1a2a PE=2 SV=1

MGKGYGHESSPEAAPTGGKRKKKDKDLDELKKEVSLDDHKLTLDELSTRYGVDLARGLTH

KRAMEILARDGPNALTPPPTTPEWVKFCKQLFGGFSILLWIGAILCFLAYSIQAATEDEP

VNDNLYLGVVLSAVVIITGCFSYYQEAKSSRIMDSFKNMVPQQALVIRDGEKLQINAEEV

VQGDLVEIKGGDRVPADLRIISSSGCKVDNSSLTGESEPQTRSPEFTHENPLETRNISFF

STNCVEGTAHGIVIATGDHTVMGRIATLASGLEVGQTPINMEIEHFIHIITGVAIFLGMS

FFILSIILGYTWLEAVIFLIGIIVANVPEGLLATVTVCLTLTAKRMARKNCLVKNLEAVE

TLGSTSTICSDKTGTLTQNRMTVAHMWFDNQIHEADTTEDQSGCDFDKSSPTWFSLSRVG

GLCNRAVFKAGQEEIPIRTRDTAGDASESALLKCVEILSGNVETLRGNNRKVAEIPFNST

NKYQLSIHELEDSPTGHLLVMKGAPERILDRCSTIMINGVEFPIDDDWMDAFQGAYMELG

GLGERVLGFCHLFLSPSQFPRGFEFDCDDVNFPVNQLCFLGLISMVDPPRAAVPDAVGKC

RSAGIKVIMVTGDHPITAKAIAKGVGIISEGNETVEDIAERLNIPLSQVNPREAKACVVH

GSDLKDMSSEYLDDILRNHTEIVFARTSPQQKLIIVEGCQRQGAIVAVTGDGVNDSPALK

KADIGVAMGITGSDVSKQAADMILLDDNFASIVTGVEEGRLIFDNLKKSIAYTLTSNIPE

ITPFLLFIIASVPLPLGTVTILCIDLGTDMVPAISLAYESAESDIMKRQPRNPKTDKLVN

ERLISMAYGQIGMIQALGGFFTYFVIMAENGFLPQTLLGIRLDWDDRTVNDLEDGYGQQW

TYEQRKIIEFTCHTSFFVSIVVVQWADLIICKTRRNSVFQQGMKNRILIFGLFAETALAA

FLSYCPGMDVALRMYPLKIMWWFCAFPYSLLIFVYDEIRKFILRRNPGGWVEIETYY

>tr|B0S5Q5|B0S5Q5_DANRE Sodium/potassium-transporting ATPase subunit alpha OS=Danio rerio OX=7955 GN=atp1a2a PE=1 SV=1

MGKGYGHESSPEAAPTGGKRKKKDKDLDELKKEVSLDDHKLTLDELSTRYGVDLARGLTH

KRAMEILARDGPNALTPPPTTPEWVKFCKQLFGGFSILLWIGAILCFLAYSIQAATEDEP

VNDNLYLGVVLSAVVIITGCFSYYQEAKSSRIMDSFKNMVPQQALVIRDGEKLQINAEEV

VQGDLVEIKGGDRVPADLRIISSSGCKVDNSSLTGESEPQTRSPEFTHENPLETRNISFF

STNCVEGTAHGIVIATGDHTVMGRIATLASGLEVGQTPINMEIEHFIHIITGVAIFLGMS

FFILSIILGYTWLEAVIFLIGIIVANVPEGLLATVTVCLTLTAKRMARKNCLVKNLEAVE

TLGSTSTICSDKTGTLTQNRMTVAHMWFDNQIHEADTTEDQSGCDFDKSSPTWFSLSRVG

GLCNRAVFKAGQEEIPIRTRDTAGDASESALLKCVEILSGNVETLRENNRKVAEIPFNST

NKYQLSIHELEDSPTGHLLVMKGAPERILDRCSTIMINGEEFPIDDDWMDAFQGAYMELG

GLGERVLGFCHLFLSPSQFPRGFEFDCDDVNFPVNQLCFLGLISMIDPPRAAVPDAVGKC

RSAGIKVIMVTGDHPITAKAIAKGVGIISEGNETVEDIAERLNIPLSQVNPREAKACVVH

GSDLKDMSAEYLDDILRNHTEIVFARTSPQQKLIIVEGCQRQGAIVAVTGDGVNDSPALK

KADIGVAMGITGSDVSKQAADMILLDDNFASIVTGVEEGRLIFDNLKKSIAYTLTSNIPE

ITPFLLFIIASVPLPLGTVTILCIDLGTDMVPAISLAYESAESDIMKRQPRNPKTDKLVN

ERLISMAYGQIGMIQALGGFFTYFVIMAENGFLPQTLLGIRLDWDDRTVNDLEDGYGQQW

TYEQRKIIEFTCHTSFFVSIVVVQWADLIICKTRRNSVFQQGMKNRILIFGLFAETALAA

FLSYCPGMDVALRMYPLKIMWWFCAFPYSLLIFVYDEIRKFILRRNPGGWVEIETYY

>tr|Q90X34|Q90X34_DANRE Sodium/potassium-transporting ATPase subunit alpha OS=Danio rerio OX=7955 GN=atp1a2a PE=2 SV=1

MGKGYGHESSPEAAPTGGKRKKKDKDLDELKKEVSLDDHKLTLDELSTRYGVDLARGLTH

KRAMEILARDGPNALTPPPTTPEWVKFCKQLFGGFSILLWIGAILCFLAYSIQAATEDEP

VNDNLYLGVVLSAVVIITGCFSYYQEAKSSRIMDSFKNMVPQQALVIRDGEKLQINAEEV

VQGDLVEIKGGDRVPADLRIISSSGCKVDNSSLTGESEPQTRSPEFTHENPLETRNISFF

STNCVEGTAHGIVIATGDHTVMGRIATLASGLEVGQTPINMEIEHFIHIITGVAIFLGMS

FFILSIILGYTWLEAVIFLIGIIVANVPEGLLATVTVCLTLTAKRMARKNCLVKNLEAVE

TLGSTSTICSDKTGTLTQNRMTVAHMWFDNQIHEADTTEDQSGCDFDKSSPTWFSLSRVG

GLCNRAVFKAGQEEIPIRTRDTAGDASESALLKCVEILSGNVETLRENNRKVAEIPFNST

NKYQLSIHELEDSPTGHLLVMKGAPERILDRCSTIMINGEEFPIDDDWMDAFQGAYMELG

GLGERVLGFCHLFLSPSQFPRGFEFDCDDVNFPVNQLCFLGLISMIDPPRAAVPDAVGKC

RSAGIKVIMVTGDHPITAKAIAKGVGIISEGNETVEDIAERLNIPLSQVNPREAKACVVH

GSDLKDMSSEYLDDILRNHTEIVFARTSPQQKLIIVEGCQRQGAIVAVTGDGVNDSPALK

KADIGVAMGITGSDVSKQAADMILLDDNFASIVTGVEEGRLIFDNLKKSIAYTLTSNIPE

ITPFLLFIIASVPLPLGTVTILCIDLGTDMVPAISLAYESAESDIMKRQPRNPKTDKLVN

ERLISMAYGQIGMIQALGGFFTYFVIMAENGFLPQTLLGIRLDWDDRTVNDLEDGYGQQW

TYEQRKIIEFTCHTSFFVSIVVVQWADLIICKTRRNSVFQQGMKNRILIFGLFAETALAA

FLSYCPGMDVALRMYPLKIMWWFCAFPYSLLIFVYDEIRKFILRRNPGGWVEIETYY

>tr|Q6P271|Q6P271_DANRE Sodium/potassium-transporting ATPase subunit alpha OS=Danio rerio OX=7955 GN=atp1a3a PE=1 SV=1

MGYGRSDSYRVATTQDNDDKESPKKGKGGKDLDDLKKEVPLTEHKMSIEEVCRKYNTDIV

QGLTNARAAEYLARDGPNALTPPPTTPEWVKFCRQLFGGFSILLWIGAILCFLAYAIQAA

TEDEPAGDNLYLGIVLSAVVIITGCFSYFQEAKSSKIMESFKNMVPQQALVIREGEKLQI

NAEEVVGGDLVEVKGGDRIPADLRIISAHGCKVDNSSLTGESEPQTRSPDCTHDNPLETR

NIAFFSTNCVEGTARGIVVCTGDRTVMGRIATLTSGLETGKTPIAKEIEHFIHIITGVAV

FLGVTFFILSIILGYSWLEAVIFLIGIIVANVPEGLLATVTVCLTLTAKRMARKNCLVKN

LEAVETLGSTSTICSDKTGTLTQNRMTVAHMWFDNQIHEADTTEDQSGASFDKSSGTWLA

LARVAALCNRAVFKAGQESLPILKRDVAGDASESALLKCIELSCGSVKAMRDKNKKVAEI

PFNSTNKYQLSVHELDESEENHYLLVMKGAPERILDRCSTILQQGKEQPMDEELKEAFQN

AYLELGGLGERVLGFCHLVMPGDKYPKGFAFDTDDINFQTDNLCFVGLMSMIDPPRAAVP

DAVGKCRSAGIKVIMVTGDHPITAKAIAKGVGIISEGNETVEDIAARLNIPVSQVNPRDA

KACVIHGTDLKDLSQDQMDEVLKNHTEIVFARTSPQQKLIIVEGCQRQGAIVAVTGDGVN

DSPALKKADIGVAMGISGSDVSKQAADMILLDDNFASIVTGVEEGRLIFDNLKKSIAYTL

TSNIPEITPFLFFILVNIPLPLGTITILCIDLGTDMVPAISLAYEAAESDIMKRQPRNPL

RDKLVNERLISIAYGQIGMIQALGGFFSYFVILAENGFLPSVLVGIRLNWDDRSNNDLED

SYGQQWTYEQRKIVEFTCHTAFFVSIVVVQWADVIICKTRRNSVFQQGMKNKILIFGLFE

ETALAAFLSYCPGMDVALRMYPLKPSWWFCAFPYSFLIFVYDEIRKLILRRNPGGWVEKE

TYY

>tr|Q9DEU2|Q9DEU2_DANRE Sodium/potassium-transporting ATPase subunit alpha OS=Danio rerio OX=7955 GN=atp1a3b PE=1 SV=1

MGYGRSDSYRVATTQDKDEKGSPKKKKGAKDLDDLKKEVPLTEHKMSVEEVCRKFQTDIV

QGLTNAKARDFLARDGPNALTPPPTTPEWVKFCRQLFGGFSILLWIGAILCFLAYAIQAA

TEDDPAGDNLYLGIVLSAVVIITGCFSYFQEAKSSKIMESFKNMVPQQALVIREGEKMQI

NAEEVVAGDLVEVKGGDRIPADLRIISAHGCKVDNSSLTGESEPQTRSPDCTHDNPLETR

NIAFFSTNCVEGTARGVVVCTGDRTVMGRIATLTSGLETGKTPIAKEIEHFIHIITGVAV

FLGVSFFILAVILGYTWLEAVIFLIGIIVANVPEGLLATVTVCLTLTAKRMARKNCLVKN

LEAVETLGSTSTICSDKTGTLTQNRMTVAHMWFDNQIHEADTTEDQSGASFDKSSVTWVA

LARVAALCNRAVFKAGQDSLPILKRDVAGDASESALLKCIELSSGSVKAMREKNKKVAEI

PFNSTNKYQLSIHETEDNNDNRYLLVMKGAPERILDRCSTIMLQGKEQPMDEEMKEAFQN

AYLELGGLGERVLGFCHVLMPEDQYPKGFAFDTDDVNFQTDNLCFVGLMSMIDPPRAAVP

DAVGKCRSAGIKVIMVTGDHPITAKAIAKGVGIISEGNETVEDIAARLNIPVSQVNPRDA

KACVIHGTDLKDYSQEQIDEVLRNHTEIVFARTSPQQKLIIVEGCQRQGAIVAVTGDGVN

DSPALKKADIGVAMGISGSDVSKQAADMILLDDNFASIVTGVEEGRLIFDNLKKSIAYTL

TSNIPEITPFLLFIIVNIPLPLGTITILCIDLGTDMVPAISLAYEAAESDIMKRQPRNPM

RDKLVNERLISIAYGQIGMIQALGGFFAYFVILAENGFLPSLLVGIRLNWDDRAMNDLED

SYGQQWTYEQRKIVEFTCHTAFFVSIVVVQWADVIICKTRRNSVFQQGMKNKILIFGLFE

ETALAAFLSYCPGMDVALRMYPLKPSWWFCAFPYSFLIFVYDEVRKLLLRRNPGGWVEKE

TYY

>tr|Q9DEY2|Q9DEY2_DANRE Sodium/potassium-transporting ATPase subunit alpha OS=Danio rerio OX=7955 GN=atp1a3a PE=2 SV=1

MGYGRSDSYRVATTQDNDDKESPKKGKGGKDLDDLKKEVPLTEHKMSIEEVCRKYNTDIV

QGLTNARAAEYLARDGPNALTPPPTTPEWVKFCRQLFGGFSILLWIGAILCFLAYAIQAA

TEDEPAGDNLYLGIVLSAVVIITGCFSYFQEAKSSKIMESFKNMVPQQALVIREGEKLQI

NAEEVAGGDLVEVKGGDRIPADLRIISAHGCKVDNSSLTGESEPQTRSPDCTHDNPLETR

NIAFFSTNCVEGTARGIVVCTGDRTVMGRIATLTSGLETGKTPIAKEIEHFIHIITGVAV

FLGVTFFILSIILGYSWLEAVIFLIGIIVANVPEGLLATVTVCLTLTAKRMARKNCLVKN

LEAVETLGSTSTICSDKTGTLTQNRMTVAHMWFDNQIHEADTTEDQSGASFDKSSGTWLA

LARVAALCNRAVFKAGQESLPILKRDVAGDASESALLKCIELSCGSVKAMRDKNKKVAEI

PFNSTNKYQLSVHELDESEENHYLLVMKGAPERILDRCSTILQQGKEQPMDEELKEAFQN

AYLELGGLGERVLGFCHLVMPGDKYPKGFAFDTDDINFQTDNLCFVGLMSMIDPPRAAVP

DAVGKCRSAGIEVIMVTGDHPITAKAIAKGVGIISEGNETVEDIAARLNIPVSQVNPRDA

KACVIHGTDLKDLSQDQMDEVLKNHTEIVFARTSPQQKLIIVEGCQRQGAIVAVTGDGVN

DSPALKKADIGVAMGISGSDVSKQAADMILLDDNFASIVTGVEEGRLIFDNLKKSIAYTL

TSNIPEITPFLFFILVNIPLPLGTITILCIDLGTDMVPAISLAYEAAESDIMKRQPRNPL

RDKLVNERLISIAYGQIGMIQALGGFFSYFVILAENGFLPSVPVGIRLNWDDRSNNDLED

SYGQQWTYEQRKIVEFTCHTAFFVSIVVVQWADVIICKTRRNSVFQQGMKNKILIFGLFE

ETALAAFLSYCPGMDVALRMYPLKPSWWFCAFPYSFLIFVYDEIRKLILRRNPGGWVEKE

TYY

>tr|Q4KMK4|Q4KMK4_DANRE Sodium/potassium-transporting ATPase subunit alpha OS=Danio rerio OX=7955 GN=atp1a3b PE=2 SV=1

MGYGRSDSYRVATTQDKDEKGSPKKKKGAKDLDDLKKEVPLTEHKMSVEEVCRKFQTDIV

QGLTNAKARDFLARDGPNALTPPPTTPEWVKFCRQLFGGFSILLWIGAILCFLAYAIQAA

TEDDPAGDNLYLGIVLSAVVIITGCFSYFQEAKSSKIMESFKNMVPQQALVIREGEKMQI

NAEEVVAGDLVEVKGGDRIPADLRIISAHGCKVDNSSLTGESEPQTRSPDCTHDNPLETR

NIAFFSTNCVEGTARGVVVCTGDRTVMGRIATLTSGLETGKTPIAKEIEHFIHIITGVAV

FLGVSFFILAVILGYTWLEAVIFLIGIIVANVPEGLLATVTVCLTLTAKRMARKNCLVKN

LEAVETLGSTSTICSDKTGTLTQNRMTVAHKWFDNQIHEADTTEDQSGASFDKSSVTWVA

LARVAALCNRAVFKAGQDSLPILKRDVAGDASESALLKCIELSSGSVKAMREKNKKVAEI

PFNSTNKYQLSIHETEDNNDNRYLLVMKGAPERILDRCSTIMLQGKEQPMDEEMKEAFQN

AYLELGGLGERVLGFCHVLMPEDQYPKGFAFDTDDVNFQTDNLCFVGLMSMIDPPRAAVP

DAVGKCRSAGIKVIMVTGDHPITAKAIAKGVGIISEGNETVEDIAARLNIPVSQVNPRDA

KACVIHGTDLKDYSQEQIDEVLRNHTEIVFARTSPQQKLIIVEGCQRQGAIVAVTGDGVN

DSPALKKADIGVAMGISGSDVSKQAADTILLDDNFASIVTGVEEGRLIFDNLKKSIAYTL

TSNIPEITPFLLFIIVNIPLPLGTITILCIDLGTDMVPAISLAYEAAESDIMKRQPRNPM

RDKLVNERLISIAYGQIGMIQALGGFFAYFVILAENGFLPSLLVGIRLNWDDRAMNDLED

SYGQQWTYEQRKIVEFTCHTAFFVSIVVVQWADVIICKTRRNSVFQQGMKNKILIFGLFE

ETALAAFLSYCPGMDVALRMYPLKPSWWFCAFPYSFLIFVYDEVRKLLLRRNPGGWVEKE

TYY

>tr|B8JKS9|B8JKS9_DANRE Sodium/potassium-transporting ATPase subunit alpha OS=Danio rerio OX=7955 GN=atp1a1a.2 PE=1 SV=1

MGLGTGNDDYRLTPTSEDELKKTKKGKRKKDVDELKKEVELDDHKLTLDELSRKYGTDMI

KGLSSFRAKEVLDRDGPNALTPPPTTPQWVKFCKQLFGGFQTLLWFGAFLCFLAYGIQVA

SVEDAAHDNLYLGLVLAFVVIVNGWFSFYQESKSSKIMESFRNLVPQQALVVRDGEKKVI

NAEEVVVGDLIEVCGGDRIPADLRIVYAQGCKVDNSSLTGESEPQSRSPEFSHENPLETK

NIAFFSTNCVEGTARGIAISTGDRTIMGRIASLASSLEGGQTPIAREIEHFIHIISAVSI

FLGVTFFVLSLILGYAWIEAVVFLIGIIVANVPEGLPATVTVCLTLTAKRMAKKNCLVKN

LEAVETLGSTTTICSDKTGTLTQNRMTVAHMWFDSQIHEADTTENQSGTSFDRSSPTWAA

LARVAGLCNRAVFRAEQSHLPVLNRETAGDASESALLKCIELCCGSVIEMREKYRKICEI

PFNSTNKYQLSVHKDPSSSGTKHLLVMKGAPERILDRCSTILINGKEQPMDDENKDSFQS

AYVELGGLGERVLGFCQFNLPDDQFPEGFAFDPEDVNFPTENLCFLGLMSMIDPPRAAVP

DAVAKCRSAGIKVIMVTGDHPITAKAIAKGVGIISEGNETVEDIAARMNIPVGEVNPREA

KACVVHGGELKNMNDSDLDEILRYHTEIVFARTSPQQKLIIVEGCQRQGAIVAVTGDGVN

DSPALKKADIGVAMGISGSDVSKQAADMILLDDNFASIVTGVEEGRLIFDNLKKSIAYTL

TSKIPEMSPFLMFVVVGIPLPLGTVTILFIDLGTDLIPAISYAYENAENDIMKRQPRNAQ

KDRLVNERLISMAYGQIGMIQAVAGFFTYITVMAENGFRPSYLPGLRVGWEDRSIGDLED

SYGQQWTYEGRKIIESTCHTAFFISIVVVQWADLLIVKTRRNSILQQGMKNKVLIFAFFE

EGALAAFLSYCPGMDIAVRMYPLRPLWWFTALPYALIIFFYDEIRKYILRRNPGGFVEKE

TYY

>tr|Q9DGL4|Q9DGL4_DANRE Sodium/potassium-transporting ATPase subunit alpha OS=Danio rerio OX=7955 GN=atp1a1a.2 PE=2 SV=1

MGLGTGNDDYRLTPTSEDELKKTKKGKRKKDVDELKKEVELDDHKLTLDELSRKYGTGMI

KGLSSFRAKEILERDGPNALTPPPTTPQWVKFCKLLFGGFQTLLWFGAFLCFLAYGIQVA

SVEDAAHDNLYLGLVLAFVVIVNGWFSFYQESKSSKIMESFRNLVPQQALVVRDGEKKVI

NAEEVVVGDLIEVCGGDRIPADLRIVYAQGCKVDNSSLTGESEPQSRSPEFSHENPLETK

NIAFFSTNCVEGTARGIAISTGDRTIMGRIASLASSLEGGQTPIAREIEHFIHIISAVSI

FLGVTFFVLSLILGYAWIEAVVFLIGIIVANVPEGLPATVTVCLTLTAKRMAKKNCLVKN

LEAVETLGSTTTICSDKTGTLTQNRMTVAHMWFDSQIHEADTTENQSGTSFDRSSPTWAA

LARVAGLCNRAVFRAEQSHLPVLNRETAGDASESALLKCIELCCGSVIEMREKYRKICEI

PFNSTNKYQLSVHKNPSSSGTKHLLVMKGAPERILDRCSTILINGKEQPMDDENKDSFQS

AYVELGGLGERVLGFCQYNLPDDQFPEGFAFDPEDVNFPTENLCFLGLMSMIDPPRAAVP

DAVAKCRSAGIKVIMVTGDHPITAKAIAKGVGIISEGNETVEDIAARMNIPVGEVNPREA

KACVVHGGELKNMNDSDLDEILRYHTEIVFARTSPQQKLIIVEGCQRQGAIVAVTGDGVN

DSPALKKADIGVAMGISGSDVSKQAADMILLDDNFASIVTGVEEGRLIFDNLKKSIAYTL

TSKIPEMSPFLMFVVVGIPLPLGTVTILFIDLGTDLIPAISYAYENAENDIMKRQPRNAQ

KDRLVNERLISMAYGQIGMIQAVAGFFTYITVMAENGFRPSYLPGLRVGWEDRSIGDLED

SYGQQWTYEGRKIIESTCHTAFFISIVVVQWADLLIVKTRRNSILQQGMKNKVLIFAFFE

EGALAAFLSYCPGMDIAVRMYPLRPLWWFTALPYALIIFFYDEIRKYILRRNPGGFVEKE

TYY

>tr|B3DFZ2|B3DFZ2_DANRE Sodium/potassium-transporting ATPase subunit alpha OS=Danio rerio OX=7955 GN=atp1a1a.5 PE=2 SV=1

MGLLTGKDDYKVAATSEDGVKTPKKGKKNKKDMDELKKEVEMDDHKLTMEELSRKYGTDL

TKGLPVSRAMEVLMRDGPNALTPPVITPEWVRFCRQLFGGFQTLLWIGAFLCYFAFSIQA

ATEEPVNDNLYLGIVLTFVVTVNGCFSYSQEAKSCRIMDSFKNLVPQKALVVRDGEKKII

DAEEVVVGDLVEVKGGDKIPADIRIVSSHGCKVDNSSLTGESEPQIRTADMSSENPLETR

NIAFFSTNCIDGAARGIVVNTGDRTVMGRIASLASNLEGGQTPLGREIEHFIHIITGVAV

FLGTTFLIISVMLGFTWLEGIIFLIGLIVANVPEGLPCTVTVALTLTAKHMAKKNCLVKN

LEAVETLGSTSTICSDKTGTLTQNRMTVAHMWFDNQIHIADTTENQTGASFDRSSATWSA

LARVAGLCNRAVFQSNQSHLPVLRRETAGDASESALLKCIELCCGSVTEMRENYPKVAEI

PFNSISKYQLSIHENPNSSEPKHLLVMKGAPERILDRCSTILIEGKEHPLDDEMKEDFQN

AYVQLGGLGERVLGFCHFGLPDDQFPEGFAFDTEEMNFPTENLCFVGLMSMIDPPRAAVP

DAVAKCRSAGIKVIMVTGDHPITAKAIAKGVGIISEGNETVDDIAARLKIHIDEVNPRDA

KACVIHGGELKNMTDEQLDDVLQHHTEIVFARTSPQQKLIIVEGCQRQGAIVAVTGDGVN

DSPALKKADIGVAMGIAGSDVSKQAADMILLDDNFASIVTGVEEGRLIFDNLKKSICYTL

STKIPEMSPFLMFVLAGIPLPLGTVTILCIDLGTDMVPAISFAYENAESDIMKRQPRNAA

TDRLVNERLVSVSYGQIGVMNAFGGFFTYFVILAENGFLPWDLVGLRIGWNDRYFSEVED

SYGQQWTYEQRKVLEYTCHTAYFTSIVIMQWTTLLVCKSRRLSLAKQGMKNRVLTFSLFE

ETAIAAFLSYCPGMDIAVRMYPLKPMWWFCAFPYMILIFIYDEVRKHFIRQNPGGWVEQE

TYY

>tr|Q9DEU1|Q9DEU1_DANRE Sodium/potassium-transporting ATPase subunit alpha OS=Danio rerio OX=7955 GN=atp1a1b PE=2 SV=1

MGVGDGRDQYELAATSEQGGKKKNKNKKKEKDMDELKKEVDLDDHKLTLEELNRKYGTDL

NRGLTTARAAEILARDGPNALTPPPTTPEWVKFCKQMFGGFSMLLWTGALLCFLAYGIQA

AMEDEPANDNLYLGVVLSAVVIITGCFSYYQEAKSSKIMDSFKNLVPQQALVVRDGEKNH

VNAEEVVVGDLVEVKGGDRIPADLRIIASHGCKVDNSSLTGESEPQTRSPDYSNDNPLET

RNIAFFSTNCVEGTARGIVISTGDRTVMGRIATLASGLEVGRTPISIEIEHFIHIITGVA

VFLGVSFFVLSLALGYSWLEAVIFLIGIIVANVPEGLLATVTVCLTLTAKRMAKKNCLVK

NLEAVETLGSTSTICSDKTGTLTQNRMTVAHMWFDNQIHEADTTENQSGTSFDRSSATWA

SLARVAGLCNRAVFLAEQTDVPILKRDVAGDASESALLKCIELCCGSVKDMREKYTKVAE

IPFNSTNKYQLSVHKNPNGGTESKHLLVMKGAPERILDRCSTILIQGKVQALDDEMKEAF

QNAYLELGGLGERVLGFCHFCLPDEEFPEGFPFDTEDVNFPTENLCFVGLMSMIDPPRAA

VPDAVGKCRSAGIKVIMVTGDHPITAKAIAKGVGIISEGNETVEDIAARLNIPVNEVNPR

DAKACVVHGGDLKDLSCEQLDDILKHHTEIVFARTSPQQKLIIVEGCQRQGAIVAVTGDG

VNDSPALKKADIGVAMGIAGSDVSKQAADMILLDDNFASIVTGVEEGRLIFDNLKKSIAY

TLTSNIPEITPFLLFIIANIPLPLGTVTILCIDLGTDMLPAISLAYEAAESDIMKRQPRN

PKTDKLVNERLISIAYGQIGMIQALAGFFTYFVILSENGFLPSRLLGIRVYWDDKHINDL

EDSYGQQWTYEQRKIVEFTCHTAFFASIVVVQWADLIICKTRRNSVFQQGMKNKILIFGL

FEETALAAFLSYCPGMDVALRMYPLKPNWWFCAFPYSLLIFIYDEMRKLILRRNPGGWVE

RETYY

>tr|Q503J4|Q503J4_DANRE Sodium/potassium-transporting ATPase subunit alpha OS=Danio rerio OX=7955 GN=atp1a1a.2 PE=2 SV=1

MGLGTGNDDYRLTPTSEDELKKTKKGKRKKDVDELKKEVELDDHKLTLDELSRKYGTDMI

KGLSSFRAKEVLDRDGPNALTPPPTTPQWVKFCKQLFGGFQTLLWFGAFLCFLAYGIQVA

SVEDAAHDNLYLGLVLAFVVIVNGWFSFYQESKSSKIMESFRNLVPQQALVVRDGEKKVI

NAEEVVVGDLIEVCGGDRIPADLRIVYAQGCKVDNSSLTGESEPQSRSPEFSHENPLETK

NIAFFSTNCVEGTARGIAVSTGDRTIMGRIASLASSLEGGQTPIAREIEHFIHIISAVSI

FLGVTFFVLSLILGYAWIEAVVFLIGIIVANVPEGLPATVTVCLTLTAKRMAKKNCLVKN

LEAVETLGSTTTICSDKTGTLTQNRMTVAHMWFDSQIHEADTTENQSGTSFDRSSPTWAA

LARVAGLCNRAVFRAEQSHLPVLNRETAGDASESALLKCIELCCGSVIEMREKYRKICEI

PFNSTNKYQLSVHKDPSSSGTKHLLVMKGAPERILDRCSTILINGKEQPMDDENKDSFQS

AYVELGGLGERVLGFCQFNLPDDQFPEGFAFDPEDVNFPTENLCFLGLMSMIDPPRAAVP

DAVAKCRSAGIKVIMVTGDHPITAKAIAKGVGIISEGNETVEDIAARMNIPVGEVNPREA

KACVVHGGELKNMNDSDLDEILRYHTEIVFARTSPQQKLIIVEGCQRQGAIVAVTGDGVN

DSPALKKADIGVAMGISGSDVSKQAADMILLDDNFASIVTGVEEGRLIFDNLKKSIAYTL

TSKIPEMSPFLMFVVVGIPLPLGTVTILFIDLGTDLIPAISYAYENAENDIMKRQPRNAQ

KDRLVNERLISMAYGQIGMIQAVAGFFTYITVMAENGFRPSYLPGLRVGWEDRSIGDLED

SYGQQWTYEGRKIIESTCHTAFFISIVVVQWADLLIVKTRRNSILQQGMKNKVLIFAFFE

EGALAAFLSYCPGMDIAVRMYPLRPLWWFTALPYALIIFFYDEIRKYILRRNPGGFVEKE

TYY

>tr|B8JKT0|B8JKT0_DANRE Sodium/potassium-transporting ATPase subunit alpha OS=Danio rerio OX=7955 GN=atp1a1a.3 PE=1 SV=1

MGLVTGNDEYKLAATSEDGVKKPKKGKKNKKDMDELKKEVEMDDHKLTLEELSRKYGTDL

NKGLSITRAKEILARDGPNALTPPVTTPEWVKFCRQLFGGFQTLLWIGALLCFFAYSIQA

ASEEEPANDNLYLGLVLAFVVTVNGCFSYYQDAKSSRIMDSFRNLVPQKALVVRDGEKSV

IDAEDVVVGDLVEVKGGDRIPADVRIVSSQGCKVDNSSLTGESEPQTRAPEMSSDNPLET

RNIAFFSTNCVDGAARGVVVNTGDRTVMGRIASLASSLEGGQTPIAREIEHFIHIITGVA

VFLGLTFLVLSLILGYNWLEGVIFLIGIIVANVPEGLPATVTVCLTLTAKRMAKKNCLVK

NLEAVETLGSTTTICSDKTGTLTQNRMTVAHMWFDNHIHIADTTENQTGASFDRSSATWS

ALARVAGLCNRAVFQSNQSHIPVLKRDTAGDASESALLKCIELSCGSVAEMRENYTKLAE

IPFNSTNKYQVSIHKNPNSSEPKHLLVMKGAPERILERCSTILIQGKEQPMDDEMKDAFQ

NAYLELGGLGERVLGFCHFCLPDDQFPEGFAFDTEEMNFPTENLCFVGLMSMIDPPRAAV

PDAVAKCRSAGIKVIMVTGDHPITAKAIAKGVGIISEGNETVDDIAARLKIHIDEVNPRD

AKACVIHGGELKNMTDEQLDDVLQHHTEIVFARTSPQQKLIIVEGCQRQGAIVAVTGDGV

NDSPALKKADIGVAMGIAGSDVSKQAADMILLDDNFASIVTGVEEGRLIFDNLKKSIAYT

LTSKIPEMSPFLMFVLVGIPLPLGTVTILCIDLGTDMVPAISLAYETAESDIMKRQPRNA

ATDRLVNERLISVSYGQIGMIQAVGGFFTYFVILAENGFLPYDLVGIRVGWEDRFLNDLE

DSYGQQWTYESRKIVEYTCHTAFFASIVIVQWTDLLICKTRRLSIFQQGMKNRVLTFGLL

EETALAAFLSYCPGMEVAVRMYPLKPLWWFCAFPYSLLIFIYDEVRKYILRRNPGGWVER

ETYY

>tr|B0R068|B0R068_DANRE Sodium/potassium-transporting ATPase subunit alpha OS=Danio rerio OX=7955 GN=atp1a1b PE=1 SV=1

MGVGDGRDQYELAATSEQGGKKKNKNKKKEKDMDELKKEVDLDDHKLTLEELNRKYGTDL

NRGLTTARAAEILARDGPNALTPPPTTPEWVKFCKQMFGGFSMLLWTGALLCFLAYGIQA

AMEDEPANDNLYLGVVLSAVVIITGCFSYYQEAKSSKIMDSFKNLVPQQALVVRDGEKNH

VNAEEVVVGDLVEVKGGDRIPADLRIIASHGCKVDNSSLTGESEPQTRSPDYSNDNPLET

RNIAFFSTNCVEGTARGIVISTGDRTVMGRIATLASGLEVGRTPISIEIEHFIHIITGVA

VFLGVSFFVLSLALGYSWLEAVIFLIGIIVANVPEGLLATVTVCLTLTAKRMAKKNCLVK

NLEAVETLGSTSTICSDKTGTLTQNRMTVAHMWFDNQIHEADTTENQSGTSFDRSSATWA

SLARVAGLCNRAVFLAEQTDVPILKRDVAGDASESALLKCIELCCGSVKDMREKYTKVAE

IPFNSTNKYQLSVHKNPNGGTESKHLLVMKGAPERILDRCSTILIQGKVQALDDEMKEAF

QNAYLELGGLGERVLGFCDFCLPDEEFPEGFPFDTEDVNFPTENLCFVGLMSMIDPPRAA

VPDAVGKCRSAGIKVIMVTGDHPITAKAIAKGVGIISEGNETVEDIAARLNIPVNEVNPR

DAKACVVHGGDLKDLSCEQLDDILKHHTEIVFARTSPQQKLIIVEGCQRQGAIVAVTGDG

VNDSPALKKADIGVAMGIAGSDVSKQAADMILLDDNFASIVTGVEEGRLIFDNLKKSIAY

TLTSNIPEITPFLLFIIANIPLPLGTVTILCIDLGTDMLPAISLAYEAAESDIMKRQPRN

PKTDKLVNERLISIAYGQIGMIQALAGFFTYFVILSENGFLPSRLLGIRVYWDDKHINDL

EDSYGQQWTYEQRKIVEFTCHTAFFASIVVVQWADLIICKTRRNSVFQQGMKNKILIFGL

FEETALAAFLSYCPGMDVALRMYPLKPNWWFCAFPYSLLIFIYDEMRKLILRRNPGGWVE

RETYY

>tr|Q9DEY3|Q9DEY3_DANRE Sodium/potassium-transporting ATPase subunit alpha OS=Danio rerio OX=7955 GN=atp1a1a.3 PE=2 SV=1

MGLVTGNDEYKLAATSEDGVKKPKKGKKNKKDMDELKKEVEMDDHKLTLEELSRKYGTDL

NKGLSITRAKEILARDGPNALTPPVTTPEWVKFCRQLFGGFQTLLWIGALLCFFAYSIQA

ASEEEPANDNLYLGLVLAFVVTVNGCFSYYQEAKSSRIMDSFRNLVPQKALVVREGEKSV

IDAEDVVVGDLVEVKGGDRIPADIRIVSSQGCKVDNSSLTGESEPQTRAPEMSSDNPLET

RNIAFFSTNCVDGAARGVVVNTGDRTVMGRIASLASSLEGGQTPIAREIEHFIHIITGVA

VFLGLTFLVLSLILGYNWLEGVIFLIGIIVANVPEGLPATVTVCLTLTAKRMAKKNCLVK

NLEAVETLGSTTTICSDKTGTLTQNRMTVAHMWFDNHIHIADTTENQTGASFDRSSATWS

ALARVAGLCNRAVFQSNQSHIPVLKRDTAGDASESALLKCIELSCGSVAEMRENYTKLAE

IPFNSTNKYQVSIHKNPNSSEPKHLLVMKGAPERILERCSTIFIQGKEQPMDDEMKDAFQ

NAYLELGGLGERVLGFCHFCLPDDQFPEGFAFDTEEMNFPTENLCFVGLMSMIDPPRAAV

PDAVAKCRSAGIKVIMVTGDHPITAKAIAKGVGIISEGNETVDDIAARLKIHIDEVNPRD

AKACVIHGGELKNMTDEQLDDVLQHHTEIVFARTSPQQKLIIVEGCQRQGAIVAVTGDGV

NDSPALKKADIGVAMGIAGSDVSKQAADMILLDDNFASIVTGVEEGRLIFDNLKKSIAYT

LTSKIPEMSPFLMFVLVGIPLPLGTVTILCIDLGTDMVPAISLAYETAESDIMKRQPRNA

ATDRLVNERLISVSYGQIGMIQAVGGFFTYFVILAENGFLPYDLVGIRVGWEDRFLNDLE

DSYGQQWTYESRKIVEYTCHTAFFASIVIVQWTDLLICKTRRLSIFQQGMKNRVLTFGLL

EETALAAFLSYCPGMEVAVRMYPLKPLWWFCAFPYSLLIFIYDEVRKYILRRNPGGWVER

ETYY

>tr|Q8JIV1|Q8JIV1_DANRE Sodium/potassium-transporting ATPase subunit alpha OS=Danio rerio OX=7955 GN=atp1a1a.5 PE=2 SV=1

MGLLTGKDDYKVAATSEDGVKTPKKGKKNKKDMDELKKEVEMDDHKLTMEELSRKYGTDL

TKGLPVSRAMEVLMRDGPNALTPPVITPEWVRFCRQLFGGFQTLLWIGAFLCYFAFSIQA

ATEEPVNDNLYLGIVLTFVVTVNGCFSYSQEAKSCRIMDSFKNLVPQKALVVRDGEKKII

DAEEVVVGDLVEVKGGDKIPADIRIVSSHGCKVDNSSLTGESEPQIRTPDMSSENPLETR

NIAFFSTNCIDGAARGIVVNTGDRTVMGRIASLASNLEGGQTPLGREIEHFIHIITGVAV

FLGTTFLIISVMLGFTWLEGIIFLIGLIVANVPEGLPCTVTVALTLTAKHMAKKNCLVKN

LEAVETLGSTSTICSDKTGTLTQNRMTVAHMWFDNQIHIADTTENQTGASFDRSSATWSA

LARVAGLCNRAVFQSNQSHLPVLRRETAGDASESALLKCIELCCGSVTGMRENYPKVAEI

PFNSISKYQLSIHENPNSSEPKHLLVMKGAPERILDRCSTILIEGKEHPLDDEMKEDFQN

AYVQLGGLGERVLGFCHFCLPDDQFPEGFAFDTEEMNFPTENLCFVGLMSMIDPPRAAVP

DAVAKCRSAGIKVIMVTGDHPITAKAIAKGVGIISEGNETVDDIAARLKIHIDEVNPRDA

KACVIHGGELKNMTDEQLDDVLQHHTEIVFARTSPQQKLIIVEGCQRQGAIVAVTGDGVN

DSPALKKADIGVAMGIAGSDVSKQAADMILLDDNFASIVTGVEEGRLIFDNLKKSICYTL

STKIPEMSPFLMFVLAGIPLPLGTVTILCIDLGTDMVPAISFAYENAESDIMKRQPRNAA

TDRLVNERLVSVSYGQIGVMNAFGGFFTYFVILAENGFLPWDLVGLRIGWNDRYFSEVED

SYGQQWTYEQRKVLEYTCHTAYFTSIVIMQWTTLLVCKSRRLSLAKQGMKNRVLTFSLFE

ETAIAAFLSYCPGMDIAVRMYPLKPMWWFCAFPYMILIFIYDEVRKHFIRQNPGGWVERE

TYY

>tr|Q7T2D6|Q7T2D6_DANRE Sodium/potassium-transporting ATPase subunit alpha OS=Danio rerio OX=7955 GN=atp1a1a.3 PE=2 SV=1

MGLVTGNDEYKLAATSEDGVKKPKKGKKNKKDMDELKKEVEMDDHKLTLEELSRKYGTDL

NKGLSITRAKEILARDGPNALTPPVTTPEWVKFCRQLFGGFQTLLWIGALLCFFAYSIQA

ASEEEPANDNLYLGLVLAFVVTVNGCFSYYQDAKSSRIMDSFRNLVPQKALVVRDGEKSV

IDAEDVVVGDLVEVKGGDRIPADVRIVSSQGCKVDNSSLTGESEPQTRAPEMSSDNPLET

RNIAFFSTNCVDGAARGVVVNTGDRTVMGRIASLASSLEGGQTPIAREIEHFIHIITGVA

VFLGLTFLILSLILGYNWLEGVIFLIGIIVANVPEGLPATVTVCLTLTAKRMAKKNCLVK

NLEAVETLGSTTTICSDKTGTLTQNRMTVAHMWFDNHIHIADTTENQTGASFDRSSATWS

ALARVAGLCNRAVFQSNQSHIPVLKRDTAGDASESALLKCIELSCGSVAEMRENYTKLAE

IPFNSTNKYQVSIHKNPNSSEPKHLLVMKGAPERILERCSTIFIQGKEQPMDDEMKDAFQ

NAYLELGGLGERVLGFCHFCLPDDQFPEGFAFDTEEMNFPTENLCFVGLMSMIDPPRAAV

PDAVAKCRSAGIKVIMVTGDHPITAKAIAKGVGIISEGNETVDDIAARLKIHIDEVNPRD

AKACVIHGGELKNMTDEQLDDVLQHHTEIVFARTSPQQKLIIVEGCQRQGAIVAVTGDGV

NDSPALKKADIGVAMGIAGSDVSKQAADMILLDDNFASIVTGVEEGRLIFDNLKKSIAYT

LTSKIPEMSPFLMFVLVGIPLPLGTVTILCIDLGTDMVPAISLAYETAESDIMKRQPRNA

ATDRLVNERLISVSYGQIGMIQAVGGFFTYFVILAENGFLPYDLVGIRVGWEDRFLNDLE

DSYGQQWTYESRKIVEYTCHTAFFASIVIVQWTDLLICKTRRLSIFQQGMKNRVLTFGLL

EETALAAFLSYCPGMEVAVRMYPLKPLWWFCAFPYSLLIFIYDEVRKYILRRNPGGWVER

ETYY

>tr|B8JKS7|B8JKS7_DANRE Sodium/potassium-transporting ATPase subunit alpha OS=Danio rerio OX=7955 GN=atp1a1a.4 PE=1 SV=1

MGLGTGNDKYKLAATSEDEAKAPKKGKKKQKDMDELKKEVDLDDHKLTLDELHRKYGTDL

TRGLSSSRAREVLARDGPNALTPPPTTPEWVKFCKQLFGGFSTLLWIGAILCFLAYGIQA

ASEDDPTNDNLYLGIVLAGVVIITGCFSYYQEAKSSKIMESFKNLVPQQALVVRDGEKKS

INAEEVVAGDLVEVKGGDRIPADLRIISAHGCKVDNSSLTGESEPQTRTPDFSNDNPLET

RNIAFFSTNCVEGTARGIVINTGDRTVMGRIATLASSLEGGQTPIAKEIEHFIHIITGVA

VFLGVSFFILSLILGYGWLEAVIFLIGIIVANVPEGLLATVTVCLTLTAKRMAKKNCLVK

NLEAVETLGSTSTICSDKTGTLTQNRMTVAHMWFDSQIHEADTTENQSGTTFDRSSPTWS

ALARVAGLCNRAVFLADQRNVPILKRDTAGDASESALLKCIELCCGSVNEMREKYTKIAE

IPFNSTNKYQLSIHKNPNSSEPKHLLVMKGAPERILDRCSTILIQGKQQPLDEEMKDAFQ

NAYVELGGLGERVLGFCHFCLPDDQFPEGFAFDTEEVNFPTENLCFVGLMSMIDPPRAAV

PDAVGKCRSAGIKVIMVTGDHPITAKAIAKGVGIISEGNETVEDIAARLNIPVGEVNPRD

AKACVVHGGELKSMSEEELDDILKHHTEIVFARTSPQQKLIIVEGCQRQGAIVAVTGDGV

NDSPALKKADIGVAMGIAGSDVSKQAADMILLDDNFASIVTGVEEGRLIFDNLKKSIAYT

LTSNIPEISPFLLFIIANIPLPLGTVTILCIDLGTDMVPAISLAYESAESDIMKRQPRNA

KTDKLVNERLISMAYGQIGMMQAVAGFFSYFVILAENGFLPSDLVGIRVNWDDKYVNDLE

DSYGQQWTYERRKIVEFTCHTAFFASIVIVQWADLIICKTRRNSIVQQGMRNKILIFGLF

EETALAAFLSYCPGMDVALRMYPLKPCWWFCAFPYSLLIFVYDEARRYILRRSPGGWVEQ

ETYY

>tr|Q90X33|Q90X33_DANRE Sodium/potassium-transporting ATPase subunit alpha OS=Danio rerio OX=7955 GN=atp1a1a.4 PE=2 SV=1

MGLGTGNDKYKLAATSEDEAKAPKKGKKKQKDMDELKKEVDLDDHKLTLDELHRKYGTDL

TRGLSSSRAKEVLARDGPNALTPPPTTPEWVKFCKQLFGGFSTLLWIGAILCFLAYGIQA

ASEDDPTNDNLYLGIVLAGVVIITGCFSYYQEAKSSKIMESFKNLVPQQALVVRDGEKKS

INAEEVVAGDLVEVKGGDRIPADLRIISAHGCKVDNSSLTGESEPQTRTPDFSNDNPLET

RNIAFFSTNCVEGTARGIVINTGDRTVMGRIATLASSLEGGQTPIAKEIEHFIHIITGVA

VFLGVSFFILSLILGYGWLEAVIFLIGIIVANVPEGLLATVTVCLTLTAKRMAKKNCLVK

NLEAVETLGSTSTICSDKTGTLTQNRMTVAHMWFDSQIHEADTTENQSGTTFDRSSPTWS

ALARVAGLCNRAVFLADQRNVPILKRDTAGDASESALLKCIELCCGSVNEMREKYTKIAE

IPFNSTNKYQLSIHKNPNSSEPKHLLVMKGAPERILDRCSTILIQGKQQPLDEEMKDAFQ

NAYVELGGLGERVLGFCHFCLPDDQFPEGFAFDTEEVNFPTENLCFVGLMSMIDPPRAAV

PDAVGKCRSAGIKVIMVTGDHPITAKAIAKGVGIISEGNETVEDIAARLNIPVGEVNPRD

AKACVVHGGELKSMSEEELDDILKHHTEIVFARTSPQQKLIIVEGCQRQGAIVAVTGDGV

NDSPALKKADIGVAMGIAGSDVSKQAADMILLDDNFASIVTGVEEGRLIFDNLKKSIAYT

LTSNIPEISPFLLFIIANIPLPLGTVTILCIDLGTDMVPAISLAYESAESDIMKRQPRNA

KTDKLVNERLISMAYGQIGMMQAVAGFFSYFVILAENGFLPSDLVGIRVNWDDKYVNDLE

DSYGQQWTYERRKIVEFTCHTAFFASIVIVQWADLIICKTRRNSIVQQGMRNKILIFGLF

EETALAAFLSYCPGMDVALRMYPLKPCWWFCAFPYSLLIFVYDEARRYILRRSPGGWVEQ

ETYY

>tr|Q9DEU0|Q9DEU0_DANRE Sodium/potassium-transporting ATPase subunit alpha OS=Danio rerio OX=7955 GN=atp1a1a.4 PE=2 SV=1

MGLGTGNDKYKLAATSEDEAKAPRKGKKKQKDMDELKKEVDLDDHKLTLDELHRKYGTDL

TRGLSSSRAKEVLARDGPNALTPPPTTPEWVKFCKQLFGGFSTLLWIGAILCFLAYGIQA

ASEDDPTNDNLYLGIVLAGVVIITGCFSYYQEAKSSKIMESFKNLVPQQALVVRDGEKKS

INAEEVVAGDLVEVKGGDRIPADLRIISAHGCKVDNSSLTGESEPQTRTPDFSNDNPLET

RNIAFFSTNCVEGTARGIVINTGDRTVMGRIATLASSLEGGQTPIAKEIEHFIHIITGVA

VFLGVSFFILSLILGYGWLEAVIFLIGIIVANVPEGLLATVTVCLTLTAKRMAKKNCLVK

NLEAVETLGSTSTICSDKTGTLTQNRMTVAHMWFDSQIHEADTTENQSGTTFDRSSPTWS

ALARVAGLCNRAVFLADQRNVPILKRDTAGDASESALLKCIELCCGSVNEMREKYTKIAE

IPFNSTNKYQLSIHKNPNSSEPKHLLVMKGAPERILDRCSTILIQGKQQPLDEEMKDAFQ

NAYVELGGLGERVLGFCHFCLPDDQFPEGFAFDTEEVNFPTENLCFVGLMSMIDPPRAAV

PDAVGKCRSAGIKVIMVTGDHPITAKAIAKGVGIISEGNETVEDIAARLNIPVGEVNPRD

AKACVVHGGELKSMSEEELDDILKHHTEIVFARTSPQQKLIIVEGCQRQGAIVAVTGDGV

NDSPALKKADIGVAMGIAGSDVSKQAADMILLDDNFASIVTGVEEGRLIFDNLKKSIAYT

LTSNIPEISPFLLFIIANIPLPLGTVTILCIDLGTDMVPAISLAYESAESDIMKRQPRNA

KTDKLVNERLISMAYGQIGMMQAVAGFFSYFVILAENGFLPSDLVGIRVNWDDKYVNDLE

DSYGQQWTYERRKIVEFTCHTAFFASIVIVQWADLIICKTRRNSIVQQGMRNKILIFGLF

EETALAAFLSYCPGMDVALRMYPLKPCWWFCAFPYSLLIFVYDEARRYILRRSPGGWVEQ

ETYY

**three spine stickleback (Gasterosteus aculeatus),**

>tr|G3PMS8|G3PMS8_GASAC_Gasterosteus

KSKKEKGKKDMEELKKEVDLDDHKLTLDELHRKYGTDVSFKGLSNSRAKEILARDGPNAL

TPPPTTPEWVKFCKQLFGGFSMLLWIGALLCFLAYGIQAASEDEPANDNLYLGVVLSAVV

IITGCFSYYQEAKSSKIMDSFKNLVPQQALVVRDGEKKSINAEEVVIGDLVEVKGGDRIP

ADLRIVSASGCKVDNSSLTGESEPQTRTPDFSNDNPLETRNIAFFSTNCVEGTARGVVIN

TGDHTVMGRIATLASSLDGGKTPIAKEIEHFIHIITGVAVFLGVSFFVLSLILGYGWLEA

VIFLIGIIVANVPEGLLATVTVCLTLTAKRMAKKNCLVKNLEAVETLGSTSTICSDKTGT

LTQNRMTVAHMWFDNQIHEADTTENQSGASFDKSSATWNALARIAGLCNRAVFLAEQSNV

PILKFLSFASNPQRDVAGDASEAALLKCIELCCGPVGNMRDKYDKIAEIPFNSTNKYQLS

VHKNQTPGETKLLLVMKGAPERILDRCSTIMLQGKEQPLDDELKDAFQNAYVELGGLGER

VLGFCHFNLPDDQFPEDFAFDTEEVNFPTENLCFIGLMSMIDPPRAAVPDAVGKCRSAGI

KVIMVTGDHPITAKAIAKGVGIISEGKRDAKACVVHGGELKEMTAELLDNVLQHHTEIVF

ARTSPQQKLIIVEGCQRQGAIVAVTGDGVNDSPALKKADIGVAMGIAGSDVSKQAADMIL

LDDNFASIVTGVEEGRLIFDNLKKSIAYTLTSNIPEISPFLLFIIANIPLPLGTVTILCI

DLGTDMVPAISLAYEEAESDIMKRQPRNPKTDKLVNERLISIAYGQIGMMQATAGFFTYF

VILAENGFLPNDLLGIRVMWDDKNFCCCFSDRWRLKTSFQGCDTPRRLFSVTALCSAEEG

REQYELAATSEHCTEPACFPAYPSPLSKHTHTQISQTVFGARSLVSTACVWYQECNTPLK

RIWINLSLDHVLSRQSIEESVLDRESFAGFCHFSLPDDQFPDGFAFDTDDRGAVSLDFPL

FVEKRTFPKTYEQRKIVEFTCHTAFFASIVVVQWADLIICKTRRNSVFQQGMKNKILIFG

LFEETALAAFLSYCPGMDVALRMYPLKPSWWFCAFPYSLLIFIYDEIRKLILRRSPGGWV

ERETYY

>tr|G3PMU0|G3PMU0_GASAC_Gasterosteus

LRLQEGREQYELAATSEHCSKKKAKGKKKEKDMDELKKEVDMDDHKLTLDELNRKYATDL

NNGLTCAKAAEILIRDGPNALTPPPTTPEWVKFCKQMFGGFSMLLWTGAVLCFLAYGIQA

AMEDEPANDNLYLGVVLSAVVIITGCFSYYQEAKSSKIMDSFKNLVPQQALVVRDGEKKN

INAEEVVVGDLVEVKGGDRIPADLRIISAHGCKVDNSSLTGESEPQTRTPDFSNENPLET

RNIAFFSTNCVEGTARGVVISTGDRTVMGRIATLASGLEVGRTPISIEIEHFIHIITGVA

VFLGVSFFVLSLILGYSWLEAVIFLIGIIVANVPEGLLATVTVCLTLTAKRMAKKNCLVK

NLEAVETLGSTSTICSDKTGTLTQNRMTVAHMWFDNQIHEADTTENQSGTSFDKSSATWA

ALARVAGLCNRAVFLAEQSNLPILKRDVAGDASESALLKCIELCCGSVQEMRNKNSKISE

IPFNSTNKYQLSIHKDGTPETGSNHLLVMKGAPERILDRCSSILLHGKEQPLDDELKDAF

QNAYLELGGLGERVLGFCHFSLPDDQFPDGFAFDTEEVNFPTENLCFIGLMSMIDPPRAA

VPDAVGKCRSAGIKVIMVTGDHPITAKAIAKGVGIISEGNETVEDIAARLNIPINEVNPR

DAKACVVHGGDLKDLAPEQLDDILKYHTEIVFARTSPQQKLIIVEGCQRQGAIVAVTGDG

VNDSPALKKADIGVAMGIAGSDVSKQAADMILLDDNFASIVTGVEEGRLIFDNLKKSIAY

TLTSNIPEITPFLLFIIANIPLPLGTVTILCIDLGTDMVPAISLAYEAAESDIMKRQPRN

PKTDKLVNERLISIAYGQIGMTEELCLWIFPSLVMKPGKGLTTPPPPPPETFKRTSRVNS

GFFPQDSYGQQWTYEQRKIVEFTCHTAFFASIVVVQWADLIICKTRRNSVFQQGMKNKIL

IFGLFEETALAAFLSYCPGMDVALRMYPLKPSWWFCAFPYSLLIFIYDEIRKLILRRSPG

GWVERETYY

>tr|G3P4Q7|G3P4Q7_GASAC_Gasterosteus

LRQRDLLTCSVNSYRVATTQDKDDRSPKKKKGAKDMDDLKKEVPITEHKMSIEEVCRKYQ

TDIVQGLTNAKAAEYLIRDGPNALTPPPTTPEWVKFCRQLFGGFSVLLWTGAILCFLAYA

IQAATEDDPAGDNLYLGIVLTAVVVITGCFSYFQEAKSSKIMESFKNMVPQQALVIREGE

KVQINAEEVVGGDLVEVKGGDRIPADIRVVSAHGCKVDNSSLTGESEPQSRSPDCTHDNP

LETRNIAFFSTNCVEGTARGIVICTGDNTVMGRIATLTSGLETGKTPIAKEIEHFIHIIT

GVAVFLGLTFFILAIVLGYSWLEAVIFLIGIIVANVPEGLLATVTVCLTLTAKRMAKKNC

LVKNLEAVETLGSTSTICSDKTGTLTQNRMTVAHMWFDNQIHEADTTEDQSGASFDKSSI

TWISLARVAALCNRAQFKAGQDQLPILKRDVAGDASESALLKCIELSCGSVRAMRDRNKK

VAEIPFNSTNKYQLSVHETEDLNDNRYLLVMKGAPERVLECCSTILVQGKEQPLDEELKE

AFQNAYMELGGLGERVLGFCHVLLPEDQYPKGFAFDTDDVNFQTKDLCFVGLMSMIDPPR

AAVPDAVGKCRSAGIKVIMVTGDHPITAKAIAKGVGIISEGNETVEDIAARLNIPVSQVN

PRDAKACVCHGTDLKDLSQDQMDDILRNHTEIVFARTSPQQKLIIVEGCQRLGAIVAVTG

DGVNDSPALKKADIGVAMGISGSDVSKQAADMILLDDNFASIVTGVEEGRLIFDNLKKSI

AYTLTSNIPEITPFLLFIIVNIPLPLGTITILCIDLGTDMVPAISLAYEAAESDIMKRQP

RNPFRDKLVNERLISIAYGQIGMIQALGGFFAYFVILAENGFLPTTLVGIRLNWDDRSCN

DLEDTYGQQWTYEQRKIVEFTCHTAFFVSIVVVQWADVIICKTRRNSVFQQGMRNKILIF

GLFEETALAAFLSYCPGMDVALRMYPLKPSWWFCAFPYSFLIFVYDEVRKLLIRRNPGGW

VERETYY

>tr|G3P2F2|G3P2F2_GASAC_Gasterosteus

LYECERERTRQCVWDWTLVAKIPGAKKKDRVVWTSIDLVDIDDHEITIEELESRYNTNVT

KGLTSTFAAQVLERDGPNELKPPKGTPEYVKFARQLAGGLQCLMWVAAVICFIAFGIEFA

RGNVASFDNLYLAITLIAVVVVTGCFGYYQEFKSTNIIASFKNLVPQQAIVIRDGQKNQI

NARQLVVGDVVEIKGGDRVPADIRIISSQACKVDNSSLTGESEPQTRSPECTHENPLETR

NVAFFSTTCLEGIATGMIISTGDRTIIGRIASLASGVGNEKTPIAIEIEHFVDIIAGLAI

LFGFTFFVVAMFIGYQFLEAMIFFMAIVVAYVPEGLLATVTVCLSLTAKRLARKNCVVKN

LEAVETLGSTSVICSDKTGTLTQNRMTVAHLWFDNHIHAADTTEDQSGQSFDQSSETWRS

LARIATLCNRATFRPDQEDVPVPKRLVVGDASETALLKFTELTVGNVMDYRNRFKKAVEL

PFNSTNKFQQLSIHELEDPLDLRYLLVMKGAPERILERCSTILIKGQELPLDEQWSEAFQ

TAYMDLGGLGERVLGFCHLHLSEKEYPRGSQFDPDEMNFPTSGLCFAGLISMIDPPRATV

PDAVMKCRTAGIRVIMVTGDHPITARAIAANVGIITEGSETVEDIAERKRIPVEQVNKRD

ARACVISGGQLKDMSSEDLDDALRNHPEMVFARTSPQQKLIIVESCQRLGSIVAVTGDGV

NDSPALKKADIGVAMGIAGSDAAKNAADMILLDDNFASIVTGVEQVGRLIFDNLKKSIAY

TLTKNIPELTPYLIYITVSVPLPLGCITILFIELATDIFPSVSLAYEKAESDIMHLKPRN

PRCHRLVNEALAVYSYFQIGAIQSFAGFTDYFAAMAQEGWFPLLCVGLRSKWENAHLQDV

QDSYGQEWTFSQRLYQEYTCYTVFFVSIEICQISDVLIRKTRRLSVFQQGFFRNRVLVSA

IVFQLLLGNLLCYCPGMPNIFNFMPIRVQWWFIPVPYGILIFVYDEIRKLGVRRYPGSWW

DQELYY

>tr|G3PMS4|G3PMS4_GASAC_Gasterosteus

MGLGRGKDEYKLAPTSDGGRKKSKKEKGKKDMEELKKEVDLDDHKLTLDELHRKYGTDLV

SFKGLSNSRAKEILARDGPNALTPPPTTPEWVKFCKQLFGGFSMLLWIGALLCFLAYGIQ

AASEDEPANDNLYLGVVLSAVVIITGCFSYYQEAKSSKIMDSFKNLVPQQALVVRDGEKK

SINAEEVVIGDLVEVKGGDRIPADLRIVSASGCKVDNSSLTGESEPQTRTPDFSNDNPLE

TRNIAFFSTNCVEGTARGVVINTGDHTVMGRIATLASSLDGGKTPIAKEIEHFIHIITGV

AVFLGVSFFVLSLILGYGWLEAVIFLIGIIVANVPEGLLATVTVCLTLTAKRMAKKNCLV

KNLEAVETLGSTSTICSDKTGTLTQNRMTVAHMWFDNQIHEADTTENQSGASFDKSSATW

NALARIAGLCNRAVFLAEQSNVPILKRDVAGDASEAALLKCIELCCGPVGNMRDKYDKIA

EIPFNSTNKYQLSVHKNQTPGETKLLLVMKGAPERILDRCSTIMLQGKEQPLDDELKDAF

QNAYVELGGLGERVLGFCHFNLPDDQFPEDFAFDTEEVNFPTENLCFIGLMSMIDPPRAA

VPDAVGKCRSAGIKVIMVTGDHPITAKAIAKGVGIISEGNETVEDIAARLNVPVSEVNPR

DAKACVVHGGELKEMTAELLDNVLQHHTEIVFARTSPQQKLIIVEGCQRQGAIVAVTGDG

VNDSPALKKADIGVAMGIAGSDVSKQAADMILLDDNFASIVTGVEEGRLIFDNLKKSIAY

TLTSNIPEISPFLLFIIANIPLPLGTVTILCIDLGTDMVPAISLAYEEAESDIMKRQPRN

PKTDKLVNERLISIAYGQIGMMQATAGFFTYFVILAENGFLPNDLLGIRVMWDDKMVNDL

EDSYGQQWTYERRKIVEFTCHTAFFVSIVVVQWADLIICKTRRNSILQQGMKNRILIFGL

FEETALAAFLSYCPGMDIALRMYPLKPCWWFCAFPYSLLIFLYDEGRRYILRRNPGGWVE

METYY

>tr|G3NB58|G3NB58_GASAC_Gasterosteus

MGYGRSDSYRVATTQDKDEKESPKKKAGKDMDDLKKEVPITEHKMSVEEVCRKYSTDIVQ

GLTNAKAAEYLIRDGPNALTPPPTTPEWVKFCRQLFGGFSILLWIGAILCFLAYAIQAAT

EDEPAGDNLYLGIVLSAVVIITGCFSYFQEAKSSKIMESFKNMVPQQALVIREGEKMQIN

AEQVVAGDLVEVKGGDRIPADLRIVSSHGCKVDNSSLTGESEPQTRSPDGTHDNPLETRN

IAFFSTNCVEGTARGIVVCTGDRTVMGRIATLTSGLETGKTPIAKEIEHFIQIITGVAVF

LGMSFFVLSIVLGYSWLEAVIFLIGIIVANVPEGLLATVTVCLTLTAKRMARKNCLVKNL

EAVETLGSTSTICSDKTGTLTQNRMTVAHMWFDNQIHEADTTEDQSGCSFDKSSTTWVSL

ARIAALCNRAVFKAGQDALPILKRDVAGDASESALLKCIELSCGSVKAMREKNKKVAEIP

FNSTNKYQLSIHENDDENDNRYLLVMKGAPERILDRSSTIMLQGKEQPMDDEMKEAFQNS

YQELGGLGERVLGFCHVFLPEDKYPKGFAFDTDDVNFETDNLCFVGLMSMIDPPRAAVPD

AVGKCRSAGIKVIMVTGDHPITAKAIAKGVGIISEGNETVEDIAARLNIPISQVNPRDAK

ACVIHGTDLKDLDQGAMDDILKNHTEIVFARTSPQQKLIIVEGCQRQGAIVAVTGDGVND

SPALKKADIGVAMGISGSDVSKQAADMILLDDNFASIVITSFITTGRLIFDNLKKSIAYT

LTSNIPEITPFLFFILVNIPLPLGTITILCIDLGTDMVPAISLAYEAAESDIMKRQPRNP

QRDKLVNERLISIAYGQIGMIQALGGFFAYFVIMAENGFLPSELVGIRLNWDDRSNNDLE

DSYGQQWTYEQRKIVEFTCHTAFFVSIVVVQWADVIICKTRRNSVFQQGMKNKILIFGLF

EETALAALLSYCPGMDVALRMYPLKPSWWFCAFPYSFLIFVYDEIRKLILRRNPGGWVER

ETYY

>tr|G3P4S9|G3P4S9_GASAC_Gasterosteus

MLLYGRSDSYRVATTQDKDDRSPKKKKGAKDMDDLKKEVPITEHKMSIEEVCRKYQTDIV

QGLTNAKAAEYLIRDGPNALTPPPTTPEWVKFCRQLFGGFSVLLWTGAILCFLAYAIQAA

TEDDPAGDNLYLGIVLTAVVVITGCFSYFQEAKSSKIMESFKNMVPQQALVIREGEKVQI

NAEEVVGGDLVEVKGGDRIPADIRVVSAHGCKVDNSSLTGESEPQSRSPDCTHDNPLETR

NIAFFSTNCVEGTARGIVICTGDNTVMGRIATLTSGLETGKTPIAKEIEHFIHIITGVAV

FLGLTFFILAIVLGYSWLEAVIFLIGIIVANVPEGLLATVTVCLTLTAKRMAKKNCLVKN

LEAVETLGSTSTICSDKTGTLTQNRMTVAHMWFDNQIHEADTTEDQSGASFDKSSITWIS

LARVAALCNRAQFKAGQDQLPILKRDVAGDASESALLKCIELSCGSVRAMRDRNKKVAEI

PFNSTNKYQLSVHETEDLNDNRYLLVMKGAPERVLECCSTILVQGKEQPLDEELKEAFQN

AYMELGGLGERVLGFCHVLLPEDQYPKGFAFDTDDVNFQTKDLCFVGLMSMIDPPRAAVP

DAVGKCRSAGIKVIMVTGDHPITAKAIAKGVGIISEGNETVEDIAARLNIPVSQVNPRDA

KACVCHGTDLKDLSQDQMDDILRNHTEIVFARTSPQQKLIIVEGCQRLGAIVAVTGDGVN

DSPALKKADIGVAMGISGSDVSKQAADMILLDDNFASIVTGVEEGRLIFDNLKKSIAYTL

TSNIPEITPFLLFIIVNIPLPLGTITILCIDLGTDMVPAISLAYEAAESDIMKRQPRNPF

RDKLVNERLISIAYGQIGMIQALGGFFAYFVILAENGFLPTTLVGIRLNWDDRSCNDLED

TYGQQWTYEQRKIVEFTCHTAFFVSIVVVQWADVIICKTRRNSVFQQGMRNKILIFGLFE

ETALAAFLSYCPGMDVALRMYPLKPSWWFCAFPYSFLIFVYDEVRKLLIRRNPGGWVERE

TYY

>tr|G3NB56|G3NB56_GASAC_Gasterosteus

MGYGRSDSYRVATTQDKDEKESPKKKAGKDMDDLKKEVPITEHKMSVEEVCRKYSTDIVQ

GLTNAKAAEYLIRDGPNALTPPPTTPEWVKFCRQLFGGFSILLWIGAILCFLAYAIQAAT

EDEPAGDNLYLGIVLSAVVIITGCFSYFQEAKSSKIMESFKNMVPQQALVIREGEKMQIN

AEQVVAGDLVEVKGGDRIPADLRIVSSHGCKVDNSSLTGESEPQTRSPDGTHDNPLETRN

IAFFSTNCVEGTARGIVVCTGDRTVMGRIATLTSGLETGKTPIAKEIEHFIQIITGVAVF

LGMSFFVLSIVLGYSWLEAVIFLIGIIVANVPEGLLATVTVCLTLTAKRMARKNCLVKNL

EAVETLGSTSTICSDKTGTLTQNRMTVAHMWFDNQIHEADTTEDQSGCSFDKSSTTWVSL

ARIAALCNRAVFKAGQDALPILKRDVAGDASESALLKCIELSCGSVKAMREKNKKVAEIP

FNSTNKYQLSIHENDDENDNRYLLVMKGAPERILDRSSTIMLQGKEQPMDDEMKEAFQNS

YQELGGLGERVLGFCHVFLPEDKYPKGFAFDTDDVNFETDNLCFVGLMSMIDPPRAAVPD

AVGKCRSAGIKVIMVTGDHPITAKAIAKGVGIISEGNETVEDIAARLNIPISQVNPRDAK

ACVIHGTDLKDLDQGAMDDILKNHTEIVFARTSPQQKLIIVEGCQRQGAIVAVTGDGVND

SPALKKADIGVAMGISGSDVSKQAADMILLDDNFASIVTGVEEGRLIFDNLKKSIAYTLT

SNIPEITPFLFFILVNIPLPLGTITILCIDLGTDMVPAISLAYEAAESDIMKRQPRNPQR

DKLVNERLISIAYGQIGMIQALGGFFAYFVIMAENGFLPSELVGIRLNWDDRSNNDLEDS

YGQQWTYEQRKIVEFTCHTAFFVSIVVVQWADVIICKTRRNSVFQQGMKNKILIFGLFEE

TALAALLSYCPGMDVALRMYPLKPSWWFCAFPYSFLIFVYDEIRKLILRRNPGGWVERET

YY

>tr|G3Q0H8|G3Q0H8_GASAC_Gasterosteus

PPPKKKTKQCGAETGGKRNKKEKNFDELKKEVSLTSLQRTGLLFHRLKKCPTTPQGLTNS

RALEILAKDGPNALTPPPTTPEWVKFCRQLFGGFSVLLWIGAILCFLAYTIQVATEDEAV

NDNLYLGVVLAAVVIITGCFSYFQEAKSSRIMDSFKKMVPQQALVIREGEKMQINAELVV

RGDLVEIRGGDRIPADLRVVSSSGCKKVDNSSLTGESEPQTRSPQFTNDNPLETRNICFF

STNCVEGTARGVVICTGDRTVMGRIATLASELQVRRTPINIEIEHFIRLITGVAVFLGLS

FFVLSVILGYTWLEAVIFLIGIIVANVPEGLLATVTVCLTLTAKRMAKKNCLVKNLEAVE

TLGSTSTICSDKTGTLTQNRMTVAHMWFDNQIHEADTTEDQTGLCFDKSSATWTALSRVA

GLCNRAEFKAGQESFPILMQEATGDASESALLKCIELCCGGVRKMRARNPKVAEIPFNST

NKYQLSVHEVEDNPSGHILVMKGAPERILDRCSSIMVHGQEQPLDGASRDAFQTAYMELG

GLGERVLGFCHSSLSPSHFPRGFTFDSDNPNFPTERLCFLGLISMIDPPRAAVPDAVGKC

RSAGIKVIMVTGDHPITAKAIARGVGIISEGTETVEDIAERLNIPLSQVNPRDAKACVVH

GSDLKDMSPEYLDDLLRNHTEIVFARTSPQQKLIIVEGCQRQGAIVAVTGDGVNDSPALK

KADIGVAMGIAGSDVSKQAADMILLDDNFASIVTGVEEGRLIFDNLKKSIAYTLTSNIPE

ISPFLLFICLSVPLPLGTVTILCIDLGTDMVPAISLAYETAESDIMKRQPRNPRTDKLVN

ERLISMAYGQIGMIQALAGFFTYFVILAENGFLPGTLLGIRIAWDDREVNDLEDTYGQQW

TYEQRKIIEFTCHTSFFISIVVVQWADLVICKTRRNSLIQQGMKNRVLIFGLFVETGLAA

FLSYCPGMDVALRMYPLKPLWWFCAFPYSLLIFIYDEVRKLILRRCPGGWVEQETYY

>tr|G3P4R4|G3P4R4_GASAC_Gasterosteus

MGDKDDRSPKKKKGAKDMDDLKKEVPITEHKMSIEEVCRKYQTDIVQGLTNAKAAEYLIR

DGPNALTPPPTTPEWVKFCRQLFGGFSVLLWTGAILCFLAYAIQAATEDDPAGDNLYLGI

VLTAVVVITGCFSYFQEAKSSKIMESFKNMVPQQALVIREGEKVQINAEEVVGGDLVEVK

GGDRIPADIRVVSAHGCKVDNSSLTGESEPQSRSPDCTHDNPLETRNIAFFSTNCVEGTA

RGIVICTGDNTVMGRIATLTSGLETGKTPIAKEIEHFIHIITGVAVFLGLTFFILAIVLG

YSWLEAVIFLIGIIVANVPEGLLATVTVCLTLTAKRMAKKNCLVKNLEAVETLGSTSTIC

SDKTGTLTQNRMTVAHMWFDNQIHEADTTEDQSGASFDKSSITWISLARVAALCNRAQFK

AGQDQLPILKRDVAGDASESALLKCIELSCGSVRAMRDRNKKVAEIPFNSTNKYQLSVHE

TEDLNDNRYLLVMKGAPERVLECCSTILVQGKEQPLDEELKEAFQNAYMELGGLGERVLG

FCHVLLPEDQYPKGFAFDTDDVNFQTKDLCFVGLMSMIDPPRAAVPDAVGKCRSAGIKVI

MVTGDHPITAKAIAKGVGIISEGNETVEDIAARLNIPVSQVNPRDAKACVCHGTDLKDLS

QDQMDDILRNHTEIVFARTSPQQKLIIVEGCQRLGAIVAVTGDGVNDSPALKKADIGVAM

GISGSDVSKQAADMILLDDNFASIVTGVEEGRLIFDNLKKSIAYTLTSNIPEITPFLLFI

IVNIPLPLGTITILCIDLGTDMVPAISLAYEAAESDIMKRQPRNPFRDKLVNERLISIAY

GQIGMIQALGGFFAYFVILAENGFLPTTLVGIRLNWDDRSCNDLEDTYGQQWTYEQRKIV

EFTCHTAFFVSIVVVQWADVIICKTRRNSVFQQGMRNKILIFGLFEETALAAFLSYCPGM

DVALRMYPLKPSWWFCAFPYSFLIFVYDEVRKLLIRRNPGGWVERETYY

>tr|G3PMT3|G3PMT3_GASAC_Gasterosteus

MEELKKEVDLDDHKLTLDELHRKYGTDLARGLSNSRAKEILARDGPNALTPPPTTPEWVK

FCKQLFGGFSMLLWIGALLCFLAYGIQAASEDEPANDNLYLGVVLSAVVIITGCFSYYQE

AKSSKIMDSFKNLVPQQALVVRDGEKKSINAEEVVIGDLVEVKGGDRIPADLRIVSASGC

KVDNSSLTGESEPQTRTPDFSNDNPLETRNIAFFSTNCVEGTARGVVINTGDHTVMGRIA

TLASSLDGGKTPIAKEIEHFIHIITGVAVFLGVSFFVLSLILGYGWLEAVIFLIGIIVAN

VPEGLLATVTVCLTLTAKRMAKKNCLVKNLEAVETLGSTSTICSDKTGTLTQNRMTVAHM

WFDNQIHEADTTENQSGASFDKSSATWNALARIAGLCNRAVFLAEQSNVPILKRDVAGDA

SEAALLKCIELCCGPVGNMRDKYDKIAEIPFNSTNKYQLSVHKNQTPGETKLLLVMKGAP

ERILDRCSTIMLQGKEQPLDDELKDAFQNAYVELGGLGERVLGFCHFNLPDDQFPEDFAF

DTEEVNFPTENLCFIGLMSMIDPPRAAVPDAVGKCRSAGIKVIMVTGDHPITAKAIAKGV

GIISEGNETVEDIAARLNVPVSEVNPRDAKACVVHGGELKEMTAELLDNVLQHHTEIVFA

RTSPQQKLIIVEGCQRQGAIVAVTGDGVNDSPALKKADIGVAMGIAGSDVSKQAADMILL

DDNFASIVTGVEEGRLIFDNLKKSIAYTLTSNIPEISPFLLFIIANIPLPLGTVTILCID

LGTDMVPAISLAYEEAESDIMKRQPRNPKTDKLVNERLISIAYGQIGMMQATAGFFTYFV

ILAENGFLPNDLLGIRVMWDDKMVNDLEDSYGQQWTYERRKIVEFTCHTAFFVSIVVVQW

ADLIICKTRRNSILQQGMKNRILIFGLFEETALAAFLSYCPGMDIALRMYPLKPCWWFCA

FPYSLLIFLYDEGRRYILRRNPGGWVEMETYY

**tilapia (Oreochromis niloticus)**

>tr|I3K3J4|I3K3J4_ORENI_Oreochromis

LCFQEGREQYELAATSEQGGKKKGKGKKKEKDMDELKKEVDMDDHKLTLDELNRKYGTDL

TNGLTSEKAAEILARDGPNALTPPPTTPEWVKFCKQMFGGFSMLLWTGAILCFLAYSIQA

AMEEEPANDNLYLGVVLSAVVIITGCFSYYQEAKSSKIMDSFKNLVPQQALVVRGGEKKS

INAEEVVVGDLVEVKGGDRIPADLRIISAHGCKVDNSSLTGESEPQTRTPDFSNENPLET

RNIAFFSTNCVEGTARGIVISTGDRTVMGRIATLASGLEVGRTPISIEIEHFIHIITGVA

VFLGVSFFILSLILGYTWLEAVIFLIGIIVANVPEGLLATVTVCLTLTAKRMAKKNCLVK

NLEAVETLGSTSTICSDKTGTLTQNRMTVAHMWFDNQIHEADTTENQSGTSFDRSSATWA

SLARIAGLCNRAVFLAEQSNLPILKRDVAGDASESALLKCIELCCGSVQEMREKTPKIAE

IPFNSTNKYQLSIHKNSSEGESKHLLVMKGAPERILDRCSTIMMQGKEQPLDDEMKDSFQ

NAYLELGGLGERVLGFCHYHLPDEQFPEDFAFDTDEVNFPTENLCFIGLMSMIDPPRAAV

PDAVGKCRSAGIKVIMVTGDHPITAKAIAKGVGIISEGNETVEDIAARLNIPINEVNPRD

AKACVVHGGDLKDLSPEQLDDILKYHTEIVFARTSPQQKLIIVEGCQRQGAIVAVTGDGV

NDSPALKKADIGVAMGIAGSDVSKQAADMILLDDNFASIVTGVEEGRLIFDNLKKSIAYT

LTSNIPEITPFLLFIIANIPLPLGTVTILCIDLGTDMVPAISLAYEAAESDIMKRQPRNP

KTDKLVNERLISIAYGQIGMIQALAGFFTYFVILAENGFLPATLVGIRVSWDNKYINDLE

DSYGQQWTYEQRKIVEFTCHTAFFVSIVIVQWADLIICKTRRNSVFQQGMKNKILIFGLF

EETALAAFLSYCPGMDVALRMYPLKPNWWFCAFPYSLLIFIYDEIRKLILRRSPGGWVER

ETYY

>tr|I3KD16|I3KD16_ORENI_Oreochromis

FGHGRVDGCEYEEALGKCGAENGGKRNKKGKDLDELKKEVALDDHKIDLDDLGKRYAVDL

RLGLTNARAVENLARDGPNTLTPPPTTPEWVKFCRQLFGGFSILLWIGAILCFLAYSIQV

ATEEEPPNDNLYLGVVLAAVVIVTGCFSYFQEAKSSRIMDSFKKMVPQQAMVIREGEKMQ

INADLVVLGDLVEIKGGDRVPADLRVISSSGCKVDNSSLTGESEPQTRSPEFTHENPLET

RNICFFSTNCVEGTARGIVIATGDRTVMGRIATLASELEVRQTPISIEIEHFIQIITGVA

VFLGVSFFILSLILGYTWLEAVIFLIGIIVANVPEGLLATVTVCLTLTAKRMAKKNCLVK

NLEAVETLGSTSTICSDKTGTLTQNRMTVAHMWFDNQIHDADTTEDQTGLGFDKSSPTWA

ALSRVAGLCNRAVFKPGQEHLPTVMRETAGDASESALLKCIEVCCGSVRDMRAANPKVAE

IPFNSTNKYQLSIHEAEDNPSGHILVMKGAPERILDRCSTIMIQGQEEPMDEMWREAFQS

AYLELGGLGERVLGFCHLNLSSSQFPRGFTFDCDDTNFPTMQLCFLGLISMIDPPRAAVP

DAVGKCRSAGIKVIMVTGDHPITAKAIAKGVGIISEGNETVEDIAERLNIPLSQVNPRDA

KACVVHGSDLKEMSSEYLDDLLRNHTEIVFARTSPQQKLIIVEGCQRQGAIVAVTGDGVN

DSPALKKADIGVAMGIAGSDVSKQAADMILLDDNFASIVTGVEEGRLIFDNLKKSIAYTL

TSNIPEISPFLLFIIASVPLPLGTVTILCIDLGTDMVPAISLAYETAESDIMKRQPRNPK

TDKLVNERLISMAYGQIGMIQALAGFFTYFVILAENGFLPRNLVGIRIDWDDREVNDLED

SFGQQWTYEQRKIVEFTCHTAFFTSIVVVQWADLIICKTRRNSLFQQGMKNRILIFGLFV

ETALAAFLSYCPGMDVALRMYPLKILWWFCGIPYSLLIFIYDEVRKFILRRQPGDGDTQK

KIEF

>tr|I3JF83|I3JF83_ORENI_Oreochromis

MKDSYDMFEMNGEMDKKKKKKMKKKEKLEGMKKEMDIDDHEITIEELEMRYTTSVDKGLT

SSFAREILERDGLNELKPPKGTPEYVKFARQLAGGLQCLMWVAAVICFIAFGIELGRGNL

TSFDDLYLAIVLIAVVVVTGCFGYYQEFKSTNIIASFKNLVPQQALVIRDGQKNQINANE

LVVGDLVEIKGGDRVPADIRIITSQGCKVDNSSLTGESEPQTRSPECTHENPLETRNIAF

FSTTCLEGVATGVIINTGDRTIIGRIASLASGVGNEKTPIAIEIEHFVDIIAGLAIFFGF

TFFVVAMFIGYAFLEAMIFFMAIVVAYVPEGLLATVTVCLSLTAKRLARKNCVVKNLEAV

ETLGSTSVICSDKTGTLTQNRMTVAHLWFDNHIHAADTTEDQSGQSFDQSSETWRSLARV

AALCNRAIFKPDQEGIPIPKRIVVGDASETALLKFTELTVGNTMDYRNRFKKIVEVPFNS

TNKFQLSIHELEDPLDLRYLLVMKGAPERILERCSTILVKGQELPLDEQWSESFQTAYMD

LGGLGERVLGFCHLYLNEKEFPRGYHFETDDMNFPTSGLCFAGLISMIDPPRATVPDAVM

KCRTAGIRVIMVTGDHPITAKAIAANVGIISEGSETVEDIAQRKRIPVEQVNKSEARACV

ISGGQLKDMSSDELDEALRNHPEMVFARTSPQQKLIIVESCQRLGSIVAVTGDGVNDSPA

LKKADIGVAMGIAGSDAAKNAADMILLDDNFASIVTGVEQGRLIFDNLKKSIAYTLTKNI

PELTPYLVYITVSVPLPLGCITILFIELATDIFPSVSLAYEKAESDIMHLKPRNPRRDRL

VNEALAVYSYFQIGAIQSFAGFTDYFTAMAQEGWFPLLCVGLRGQWEDVKLQDLQDSYGQ

EWTYSQRLYQEYTCYTVFFVSIEICQIADVLIRKTRRLSAFQQGFFRNRVLVSAIVFQLC

LGNLLCYCPGMPNIFNFMPIRVQWWFVPVPYGILIFVYDEIRKLGVRRHPGSWWDQELYY

>tr|I3K3G3|I3K3G3_ORENI_Oreochromis

MGLGKGKDEYKLAATSEDGGKKDKKAKAKKDMDDLKKEVDLDDHKLTLDELHRKYGTDLT

RGLSSSRAKEILARDGPNALTPPPTTPEWVKFCKQLFGGFSMLLWIGAILCFLAYGIQAA

SEDEPANDNLYLGIVLSAVVIITGCFSYYQEAKSSKIMESFKNLVPQQALVIRDGEKKNI

NAEEVVLGDLVEVKGGDRIPADLRIISAHGCKVDNSSLTGESEPQTRSPDFSNENPLETR

NISFFSTNCIEGTARGIVINTGDRTVMGRIATLASSLEGGKTPIAIEIEHFIHIITGVAV

FLGVSFFILSLILGYNWLEAVIFLIGIIVANVPEGLLATVTVCLTLTAKRMAKKNCLVKN

LEAVETLGSTSTICSDKTGTLTQNRMTVAHMWFDNQIHEADTTENQSGTSFDRSSATWAN

LSRIAGLCNRAVFLADQSNIPILKRDVAGDASEAALLKCIELCCGSVNEMREKYPKIAEI

PFNSTNKYQLSIHKNTTPGETKHLLVMKGAPERILDRCSSIVLQGKVQALDDEMKDAFQN

AYVELGGLGERVLGFCHYHLPDDEFPEGFAFDTDEVNFPTENLCFVGLMAMIDPPRAAVP

DAVGKCRSAGIKVIMVTGDHPITAKAIAKGVGIISEGNETVEDIAARLNIPVSEVNPRDA

KACVIHGGELKDMTTEQIDDVLKHHTEIVFARTSPQQKLIIVEGCQRQGAIVAVTGDGVN

DSPALKKADIGVAMGIAGSDVSKQAADMILLDDNFASIVTGVEEGRLIFDNLKKSIAYTL

TSNIPEISPFLLFIIANIPLPLGTVTILCIDLGTDMVPAISLAYEKAESDIMKRQPRNPK

TDKLVNERLISIAYGQIGMMQATAGFFTYFVILAENGFLPMDLIGLRVLWDDKYVNDLED

SYGQQWTYERRKIVEYSCHTAFFASIVIVQWADLIICKTRRNSIVQQGMTNRILIFGLFE

ETALAAFLSYCPGMDVALRMYPMKPLWWFCAFPYSLLIFLYDEARRYILRRNPGGWVEKE

TYY

>tr|I3K372|I3K372_ORENI_Oreochromis

MGSKRMNDDYKLANTSDNDEKTTKKGNKSKDTEDLKKEVDLDDHKLSFDELHRKYGTDLT

RGLSSSKAKEILARDGPNALTPPPTTPEWVKFCKQLFGGFCMLLWIGAFLCFLAYAIQVA

SEENPGNDNLYLGIVLAAVVIITACFSYYQEAKSSRIMDSFKNMVPQQALVIRDGEKKSI

NTEEVVLGDLVEVKGGDRIPADLRIISAHGCKVDNSSLTGESEPQTRAPDFSHENPLETR

NIAFFSTNCVEGTARGVVINTGDNTIMGRIATLTSSLEAGKTPIAIEIEHFIHIITGVAV

FLGVTFFILSVILGYNWLDGVIFLIGIIVANVPEGLLATVTVCLTLTAKRMAKKNCLVKN

LEAVETLGSTSTICSDKTGTLTQNRMTVAHMWFDNQIHEADTTENQSGVAFQKSSPTWTA

LSRVAGLCNRAVFLAGQNDVPILKRNIAGDASEAALLKCIELCCGSVSEMREKYPKIAEI

PFNSTNKYQLSIHKNTTPGETKHLLVMKGAPERILDRCSTIMLEGKEQPLDDELKDAFNS

AYLQLGGLGERVLGFSHFHLPDDQFPEGFAFDADDVNFPTENLCFTGLMSMIDPPRAAVP

DAVGKCRSAGIKVIMITGDHPITATAIARGVGIISEGNETVEDIAARLNVPVSEVNPRDA

KACVIHGGELKDMTPEEIDDVLRNHTEIVFARTSPQQKLIIVEGCQRQGAIVAVTGDGVN

DSPALKKADIGVAMGIAGSDVSKQAADMILLDDNFASILTGVEEGRLIFDNLKKSIAYTL

TSKIPEMSPFLLFVIANIPLPLGTVTILCIDLGTDLVPAISLAYEQAESDIMKRQPRNAS

DRLVNERLISMAYGQIGMIQASAGFFTYFVVLAENGFLPLDLMGLRILWDNRFLNDLEDS

YGQEWTYESRKILEFTCHTAFFSSIVIVQVADLLICKTRRNSILQQGMKNNVLIFGMFEE

LALAVFLSYCPGMDVALKMYPMKPLWWFCGLPHGFLLFFYDEIRKYIIRHHPGGWVEKET

YY

>tr|I3JBJ7|I3JBJ7_ORENI_Oreochromis

MGYGRSDSYRVATTQDKDEKESPKKKGGKDLDDLKKEVPITEHKMSVEEVCRKYNTDIVQ

GLTNARAAEYLARDGPNALTPPPTTPEWVKFCRQLFGGFSILLWIGAILCFLAYAIQAAT

EDEPAGDNLYLGIVLSAVVIITGCFSYFQEAKSSKIMESFKNMVPQQALVIREGEKMQIN

AEQVVAGDLVEVKGGDRIPADLRIISSHGCKVDNSSLTGESEPQTRSPDCTHDNPLETRN

IAFFSTNCVEGTARGIVVCTGDRTVMGRIATLTSGLETGKTPIAKEIEHFIQIITGVAVF

LGVTFFILSIILGYSWLEAVIFLIGIIVANVPEGLLATVTVCLTLTAKRMARKNCLVKNL

EAVETLGSTSTICSDKTGTLTQNRMTVAHMWFDNQIHEADTTEDQSGSSFDKSSTTWVSL

ARIAALCNRAVFKAGQESLPILKRDVAGDASESALLKCIELSCGSVKAMRDKNKKVAEIP

FNSTNKYQLSIHETEEENDSRYLLVMKGAPERILDRCSTIMVQGKEQPMDDEMKEAFQNA

YLELGGLGERVLGFCHVFMPEDKYPKGFAFDTDDVNFQTDNLCFVGLMSMIDPPRAAVPD

AVGKCRSAGIKVIMVTGDHPITAKAIAKGVGIISEGNETVEDIAARLNIPVSQVNPRDAK

ACVIHGTDLKDLTQDQMDDILRNHTEIVFARTSPQQKLIIVEGCQRQGAIVAVTGDGVND

SPALKKADIGVAMGISGSDVSKQAADMILLDDNFASIVTGVEEGRLIFDNLKKSIAYTLT

SNIPEITPFLFFILVNIPLPLGTITILCIDLGTDMVPAISLAYEAAESDIMKRQPRNPLR

DKLVNERLISIAYGQIGMIQALGGFFAYFVIMAENGFLPSLLVGIRLNWDDRSNNDLEDS

YGQQWTYEQRKIVEFTCHTAFFVSIVVVQWADVIICKTRRNSVFQQGMRNKILIFGLFEE

TALAAFLSYCPGMDLALRMYPLKPSWWFCAFPYSFLIFVYDEIRKLILRRNPGGWVEKET

YY

>tr|I3K5Q8|I3K5Q8_ORENI_Oreochromis

MGDKDDRSPKKKKGGTKDMDDLKKEVPITEHKMSVEEVCRKFQTDVVQGLTNAKAAEFLL

RDGPNALTPPPTTPEWVKFCRQLFGGFSILLWTGAILCFLAYAIQAATEDEPAGDNLYLG

IVLTAVVVITGCFSYFQEAKSSKIMESFKNMVPQQALVIREGEKVQINAEEVVAGDLIEV

KGGDRIPADIRVTSAHGCKVDNSSLTGESEPQSRSPDCTHDNPLETRNIAFFSTNCVEGT

ARGIVICTGDRTVMGRIATLTSGLETGKTPIAVEIEHFIHIITGVAVFLGVTFFILAIIL

GYTWLEAVIFLIGIIVANVPEGLLATVTVCLTLTAKRMAKKNCLVKNLEAVETLGSTSTI

CSDKTGTLTQNRMTVAHMWFDNQIHEADTTEDQSGAAFDKSSVTWLSLSRVAALCNRAQF

KAGQDSVAILKRDVAGDASESALLKCIELSCGSVRMMRDKNKKVAEIPFNSTNKYQLSIH

ETEDPNDNRYLLVMKGAPERILDRCSTIMLQGKEQPMDEEMKEAFQNAYMELGGLGERVL

GFCHLLLPEDQYPKGFAFDTDDVNFQTDNLCFVGLMSMIDPPRAAVPDAVGKCRSAGIKV

IMVTGDHPITAKAIAKGVGIISEGNETVEDIAARLNIPVSQVNPRDAKACVIHGSDLKDL

SQDQMDDILRNHTEIVFARTSPQQKLIIVEGCQRLGAIVAVTGDGVNDSPALKKADIGVA

MGISGSDVSKQAADMILLDDNFASIVTGVEEGRLIFDNLKKSIAYTLTSNIPEITPFLFF

IIVNIPLPLGTITILCIDLGTDMVPAISLAYEAAESDIMKRQPRNPSRDKLVNERLISIA

YGQIGMIQALGGFFAYFVILAENGFLPSQLVGIRLNWDDRSLNDLEDSYGQQWTYEQRKI

VEFTCHTAFFVSIVVVQWADLIICKTRRNSVFQQGMRNKILIFGLFEETALAAFLSYCPG

MDVALRMYPLKPTWWFCAFPYSFLIFVYDEARKLILRRNPGGWVEKETYY

>tr|I3K395|I3K395_ORENI_Oreochromis

MGVTKMKDDYKLAATADHEEKRSKKGKKHRDTEDLKKEVDLDDHKLSVDELHRKYGTDLV

MGLSSFRAKEILARDGPNALTPPPTTPEWVKFCKQLFGGFCMLLWIGAFLCFVAYSIQAA

SEDEPASDNLYLGIVLSVVVMITACFSYYQEAKSSRIMDSFKNMVPQQALVIRDGEKKSI

NTEEVVLGDLVEVKGGDRIPADLRIISAHGCKVDNSSLTGESEPQTRSPEFSNENPLETR

NIAFFSTNCVEGTARGVVINTGDNTIMGRIATLASSLEAGKTPIAIEIEHFIHIITGVAV

FLGVSFFILSVILGYNWLEGIIFLIGIIVANVPEGLLATVTVCLTLTAKRMAKKNCLVKN

LEAVETLGSTSTICSDKTGTLTQNRMTVAHMWFDNQIHVADTTENQSGTSFDRSSATWAA

LSRIAGLCNRAVFLAEQNKVPVLKRNVAGDASEAALLKCIELCCGSVSDMREKYPKIAEI

PFNSTNKYQLSIHKNTTPGETKHLLVMKGAPERILDRCSTIVIQGKEQPLDAELKDSFNS

AYLELGGLGERVLGFCHYHLSDDQFPEGFAFDADDVNFPTENLCFIGLMSMIDPPRAAVP

DAVSKCRSAGIKVIMVTGDHPITAKAIARGVGIISEGNETVEDIAARLNISVSEVNPREA

KACVIHGSELKEMTTEQIDDVLKHHTEIVFARTSPQQKLIIVEGCQRQGAIVAVTGDGVN

DSPALKKADIGVAMGIAGSDVSKQAADMILLDDNFASILTGVEEGRLIFDNLKKSIAYTL

TSKIPEMSPFLFFVLFDIPLALGTVTILCIDLGTDMVPAISLAYEQAESDIMKRQPRNAQ

TDKLVNERLISMAYGQIGMMQALGGFFTYFVILSENGFLPKDLVGIRVFWDNRYLNDLED

SYGQEWTYERRKIVEFTCHTAFFTSIVIVQVADLLICKTRTNSIVKQGMKNYVLIFGIFE

ELALAAFLSYCPGMDIAIRMYPMKPWWWLCSVPYSFLIFIYDEVRKYILRRSPGGWVELE

TYY

>tr|I3K3B9|I3K3B9_ORENI_Oreochromis

MGFGRGKDNYKLAATSENGGKKSKKARKKKDMDELKKEVDLDDHRLTLEELIRKYGTDLT

RGLSSSRAKEILARDGPNALTPPPTTPEWVKFCKQLFGGFSMLLWIGAILCFLAYSIQAA

SEDEPANDNLYLGIVLSAVVIITGCFSYYQEAKSSKIMESFKNMVPQQALVIRDGEKKSI

NAEEVVLGDLVEVKGGDRIPADLRVVSAQGCKVDNSSLTGESEPQTRSPEFSNENPLETR

NIAFFSTNCVEGTARGVVINIGDNTVMGRIATLASSLEGGKTPIAIEIEHFIHIITGVAV

FLGVTFFILSLILGYGWLEAVIFLIGIIVANVPEGLLATVTVCLTLTAKRMAKKNCLVKN

LEAVETLGSTSTICSDKTGTLTQNRMTVAHMWFDNQIHEADTTENQSGTSFDRSSATWAA

LSRIAGLCNRAVFLADQSNIPILKRNVAGDASEAALLKCIELCCGSVSGMRDKYPKIAEI

PFNSTNKYQLSIHKNSTPGETKHLLVMKGAPERILDRCSTIMLQGKEQALDDEMKDSFNN

AYVELGGLGERVLGFCQYHLPDDQFPEGFAFDTDEVNFPTENLCFVGLMAMIDPPRAAVP

DAVGKCRSAGIKVIMVTGDHPITAKAIAKGVGIISEGNETVEDIAARLNIPVSEVNPRDA

KACVIHGGELKDMTTEQIDDVLKHHTEIVFARTSPQQKLIIVEGCQRQGAIVAVTGDGVN

DSPALKKADIGVAMGIAGSDVSKQAADMILLDDNFASIVTGVEEGRLIFDNLKKSIAYTL

TSNIPEISPFLLFIIANIPLPLGTVTILCIDLGTDMVPAISLAYEKAESDIMKRQPRNPK

TDKLVNERLISIAYGQIGMMQATAGFFTYFVILAENGFLPMDLIGLRVLWDDKYVNDLED

SYGQEWTYERRKIVEFTCHTAFFTSIVIVQWADLIICKTRRNSIWKQGMTNHILIFGLFE

ETALAAFLSYCPGMDVALRMYPMKPLWWFCAFPYSLLIFLYDEARRYILRRNPGGWVELE

TYY

>tr|R9WR11|R9WR11_9CICH Sodium/potassium-transporting ATPase subunit alpha OS=Oreochromis urolepis hornorum OX=161276 PE=2 SV=1

MGLGKGKDEYKLAATSEDGGKKDKKAKAKKDMDDLKKEVDLDDHKLTLDELHRKYGTDLT

RGLSSSRAKEILARDGPNALTPPPTTPEWVKFCKQLFGGFSMLLWIGAILCFLAYGIQAA

SEDEPANDNLYLGIVLSAVVIITGCFSYYQEAKSSKIMESFKNLVPQQALVIRDGEKKNI

NAEEVVLGDLVEVKGGDRIPADLRIISAHGCKVDNSSLTGESEPQTRSPDFSNENPLETR

NISFFSTNCIEGTARGIVINTGDRTVMGRIATLASSLEGGKTPIAIEIEHFIHIITGVAV

FLGVSFFILSLILGYNWLEAVIFLIGIIVANVPEGLLATVTVCLTLTAKRMAKKNCLVKN

LEAVETLGSTSTICSDKTGTLTQNRMTVAHMWFDNQIHEADTTENQSGTSFDRSSATWAN

LSRIAGLCNRAVFLADQSNIPILKRDVAGDASEAALLKCIELCCGSVNEMREKYPKIAEI

PFNSTNKYQLSIHKNTTPGETKHLLVMKGAPERILDRCNSIVLQGKVQALDDEMKDAFQN

AYVELGGLGERVLGFCHYHLPDDEFPEGFAFDTDEVNFPTENLCFVGLMAMIDPPRAAVP

DAVGKCRSAGIKVIMVTGDHPITAKAIAKGVGIISEGNETVEDIAARLNVPVSEVNPRDA

KACVVHGSELKDMTSEELDDLLKHHTEIVFARTSPQQKLIIVEGCQRQGAIVAVTGDGVN

DSPALKKADIGVAMGIAGSDVSKQAADMILLDDNFASIVTGVEEGRLIFDNLKKSIAYTL

TSNIPEISPFLLFIIANIPLPLGTVTILCIDLGTDMVPAISLAYEKAESDIMKRQPRNPK

TDKLVNERLISIAYGQIGMMQATAGFFTYFVILAENGFLPMDLIGVRVLWDDKYVNDLED

SYGQQWTYERRKIVEYSCHTAFFASIVIVQWADLIICKTRRNSIVQQGMTNRILIFGLFE

ETALAAFLSYCPGMDVALRMYPMKPLWWFCAFPYSLLIFLYDEARRYILRRNPGGWVEKE

TYY

**mudskipper (Boleophthalmus pectinirostris)**

>XP_020792240.1_Boleophthalmus

MDELKKEVDMDDHKLTLDELNRKYGTDLTNGLTSAKAAENLARDGPNALTPPPTTPEWVKFCKQMFGGFSMLLWTGAILCFLAYGIQAAMEDEPANDNLYLGVVLSAVVIITGCFSYYQEAKSSKIMDSFKNLVPQQALVVRDGEKKNINAEEVVVGDLVEVKGGDRIPADLRIISAHGCKVDNSSLTGESEPQTRTPDFSNENPLETRNIAFFSTNCVEVKLSPGTARGIVISTGDRTVMGRIATLASGLEVGRTPISIEIEHFIHIITGVAVFLGVSFFILSLILGYTWLEAVIFLIGIIVANVPEGLLATVTVCLTLTAKRMAKKNCLVKNLEAVETLGSTSTICSDKTGTLTQNRMTVAHMWFDNQIHEADTTENQSGTSFDRSSATWGALARIAGLCNRAVFLAEQSNIPILKRDVAGDASESALLKCIELCCGSVQEMREKNPKISEIPFNSTNKYQLSIHKNGTPEGESKHLLVMKGAPERILDRCSSILLQGKEQPLDDEMKDAFQNAYLELGGLGERVLGFCHYNLPDEQFPDGFAFDTDEVNFPTENLCFIGLMSMIDPPRAAVPDAVGKCRSAGIKVIMVTGDHPITAKAIAKGVGIISEGNETVEDIAARLNIPINEVNPRDAKACVVHGGDLKDLTAEQLDDILKYHTEIVFARTSPQQKLIIVEGCQRQGAIVAVTGDGVNDSPALKKADIGVAMGIAGSDVSKQAADMILLDDNFASIVTGVEEGRLIFDNLKKSIAYTLTSNIPEITPFLLFIIANIPLPLGTVTILCIDLGTDMVPAISLAYEAAESDIMKRQPRNPKTDKLVNERLISIAYGQIGMIQALAGFFTYFVILAENGFLPTTLLGIRVSWDNKYINDLEDSYGQQWTYEQRKIVEFTCHTAFFASIVIVQWADLIICKTRRNSVFQQGMKNKILIFGLFEETALAAFLSYCPGMDVALRMYPLKPNWWFCAFPYSLLIFIYDEIRKLILRRSPGGWVERETYY

>XP_020792263.1_Boleophthalmus

MGLGRGKDEYKLAATSDNDKSKKAKKAKEKKDMDDLKKEVDLDDHKLTLDELHRKYGTDLTRGLSGSRAKEILARDGPNALTPPPTTPEWVKFCKQLFGGFSMLLWIGAILCFLAYGIQAASEDEPANDNLYLGVVLSAVVIITGCFSYYQEAKSSKIMDSFKNLVPQQALVVRDGEKKNINAEEVVVGDLVEVKGGDRIPADLRIISAHGCKVDNSSLTGESEPQTRTPDFSNENPLETRNIAFFSTNCVEGTARGVVINTGDRTVMGRIATLASSLEGGKTPIAIEIEHFIHIITGVAVFLGVSFFILSLILGYGWLEAVIFLIGIIVANVPEGLLATVTVCLTLTAKRMAKKNCLVKNLEAVETLGSTSTICSDKTGTLTQNRMTVAHMWFDNQIHEADTTENQSGTSFDRSSATWGALARIAGLCNRAVFLADQNNVPILKRDVAGDASEAALLKCIELCCGSVAGMREKYPKVAEIPFNSTNKYQLSIHKNSTPGETKHLLVMKGAPERILDRCSTIVLQGKEQPLDDEMKDAFQNAYVELGGLGERVLGFCHFSLPDDQFPEGFAFDTEEVNFPTENLCFVGLMAMIDPPRAAVPDAVGKCRSAGIKVIMVTGDHPITAKAIAKGVGIISEGNETVEDIAQRLNVPVSEVNPRDAKACVVHGGELKDMTSEQLDDILKHHTEIVFARTSPQQKLIIVEGCQRQGAIVAVTGDGVNDSPALKKADIGVAMGIAGSDVSKQAADMILLDDNFASIVTGVEEGRLIFDNLKKSIAYTLTSNIPEITPFLLFIIANIPLPLGTVTILCIDLGTDMVPAISLAYEAAESDIMKRQPRNPKTDKLVNERLISIAYGQIGMMQAVAGFFTYFVILAENGFLPMDLLGIRVLWDDKYVNDLEDSYGQQWTYERRKIVEFTCHTAFFASIVIVQWADLIICKTRRNSILQQGMKNRILIFGLFEETALAAFLSYCPGMDVALRMYPLKPSWWFCAFPYSLLIFLYDEARRYILRRNPGGWVEQETYY

>XP_020796248.1_Boleophthalmus

MGYGRSDSYRVATTQDKDGRSPTKKKGGKDMDELKKEVPITEHKMSVEEVCRKYQTDIVQGLTNSRAAEFLKRDGPNALTPPPTTPEWVKFCRQLFGGFSILLWTGAILCFLAYAIQAATEDEPAGDNLYLGIVLTAVVIITGCFSYFQEAKSSKIMESFKNMVPQQALVIREGEKVQINAEEVVAGDLIEVKGGDRIPADIRVVSAHGCKVDNSSLTGESEPQNRSPDCTHDNPLETRNVAFFSTNCVEGTARGIVICTGDRTVMGRIATLTSGLETGKTPIAKEIEHFIHIITGVAVFLGVTFFILAIILGYTWLEAVIFLIGIIVANVPEGLLATVTVCLTLTAKRMAKKNCLVKNLEAVETLGSTSTICSDKTGTLTQNRMTVAHMWFDNQIHEADTTEDQSGASFDKSSITWQALSRVAGLCNRAQFKAGQDSIAILKRDVAGDASESALLKCIELSCGSVRQMRDRNKKVAEIPFNSTNKYQLSVHETEDPNDNRYLLVMKGAPERILERCSTIMIQGKEQPMDEELKDSFQNAYMELGGLGERVLGFCHLLLPEDQYPKGFAFDTDDVNFQTENLCFVGLMSMIDPPRAAVPDAVGKCRSAGIKVIMVTGDHPITAKAIAKGVGIISEGNETVEDIALRLNIPVSQVNPRDAKACVIHGTDLKELSQDQMDDILRNHTEIVFARTSPQQKLIIVEGCQRQGAIVAVTGDGVNDSPALKKADIGVAMGISGSDVSKQAADMILLDDNFASIVTGVEEGRLIFDNLKKSIAYTLTSNIPEITPFLFFIIVNIPLPLGTITILCIDLGTDMVPAISLAYEAAESDIMKRQPRNPFRDKLVNERLISIAYGQIGMIQALGGFFTYFVILAENGFLPSQLVGIRLNWDDRACNDLEDSYGQQWTYEQRKIVEFTCHTAFFVSIVVVQWADVIICKTRRNSVFQQGMKNKILIFGLFEETALAAFLSYCPGMDVALRMYPLKPTWWFCAFPYSFLIFVYDEIRKLLIRRNPGGWVERETYY

>XP_020793662.1_Boleophthalmus

MGYGRSDSYRVATTQDKDEKESPKKKGGKDLDDLKKEVPITEHKMSVEEVCRKYNTDIVQGLTNARAAEYLARDGPNALTPPPTTPEWVKFCRQLFGGFSILLWIGAILCFLAYAIQAATEDEPAGDNLYLGIVLSAVVIITGCFSYFQEAKSSKIMESFKNMVPQQALVIREGEKMQINAEQVVAGDLVEVKGGDRIPADLRIISSHGCKVDNSSLTGESEPQTRSPDCTHDNPLETRNIAFFSTNCVEGTARGIVVCTGDRTVMGRIATLTSGLETGKTPIAKEIEHFIHIITGVAVFLGVTFFVLSLILGYSWLEAVIFLIGIIVANVPEGLLATVTVCLTLTAKRMARKNCLVKNLEAVETLGSTSTICSDKTGTLTQNRMTVAHMWFDNQIHEADTTEDQSGCSFDKSSTTWVSLARIATLCNRAVFKAGQEALPILKREVAGDASESALLKCIELSCGAVKAMRDKNKKVAEIPFNSTNKYQLSIHETEDENDNRYLLVMKGAPERILDRCTTIMVQGKEQPMDDELKEAFQNAYLELGGLGERVLGFCHLFMPEDKYPKGFAFDTDDVNFQTDGLCFVGLMSMIDPPRAAVPDAVGKCRSAGIKVIMVTGDHPITAKAIAKGVGIISEGNETVEDIALRLNIPVSQVNPRDAKACVIHGTDLKDLSQEQMDDILKNHTEIVFARTSPQQKLIIVEGCQRQGAIVAVTGDGVNDSPALKKADIGVAMGISGSDVSKQAADMILLDDNFASIVTGVEEGRLIFDNLKKSIAYTLTSNIPEITPFLFFILVNIPLPLGTITILCIDLGTDMVPAISLAYEAAESDIMKRQPRNPARDKLVNERLISIAYGQIGMIQALGGFFAYFVIMAENGFLPSHLVGIRLNWDDRSNNDLEDSYGQQWTYEQRKIVEFTCHTAFFVSIVVVQWADVIICKTRRNSVFQQGMRNKILIFGLFEETALAAFLSYSPGMDVALRMYPLKPSWWFCAFPYSFLIFVYDEIRKLILRRNPGGWVEKETYY

>XP_020796901.1_Boleophthalmus

MVPSAGADTTGGRPMMFFWAEPGQAPDSRVQAPAPGLAPGSSCSRLGENGDGGKRKKRKKDQDLDELKKEVSLDDHKLSTEELAQRYGLDLAQGLTNAKAAEVLARDGPNALTPPPTTPEWVKFCRQLFGGFSILLWIGAILCFFAYSIQVGTEDEPVNDNVYLGVVLAAVVIITGCFSYFQEAKSSRIMDSFKKMVPQQALVIREGEKLQINAELVVQGDLVEIKGGDRIPADLRVISSSGCKVDNSSLTGESEPQTRSPDFTHENPLETRNICFFSTNCVEGTARGIVIATGDRTVMGRIATLASELQVRRTPINIEIEHFIHLITGVAVFLGVSFFILSLILGYTWLEAVIFLIGIIVANVPEGLLATVTVCLTLTAKRMAKKNCLVKNLEAVETLGSTSTICSDKTGTLTQNRMTVAHMWFDNMVHEADTTENQSGSSFDKTSATWLSLSRVAALCNRAVFKADQDHVPILMRDTAGDASESALLKCIEVCCGSVRKMRDRNPKVAEIPFNSTNKYQLSIHQCEENPGHILVMKGAPERILDRCSTIMLNGQEVPLDDAWKESFQNAYMELGGLGERVLGFCHLFLPPSQYPTHFSFDTDDVNFPTEGLCFVGLISMIDPPRAAVPDAVGKCRSAGIKVIMVTGDHPITAKAIAKGVGIISEGNETVEDIAERLNISLSEVNPRDAKACVVHGSDLKDMSSEYLDDLLKNHTEIVFARTSPQQKLIIVEGCQRQGAIVAVTGDGVNDSPALKKADIGVAMGITGSDVSKQAADMILLDDNFASIVTGVEEGRLIFDNLKKSIAYTLTSNIPEITPFLLFIIASVPLPLGTVTILCIDLGTDMVPAISLAYETAESDIMKRQPRNPKTDKLVNERLISMAYGQIGMIQALAGFFTYFVILAENGFLPYSLVGIRIQWDNREVNDLEDTYGQQWTYEQRKIVEFTCHTAFFSSIVIVQWADLIICKTRRNSLFQQGMKNRILIFGLFVETALAAFLSYCPGMDVALRMYPLRIFWWFCALPYSLLIFVYDEVRKYILRRSPGGKKNTRKTHEKKTRETASNQ

>XP_020792751.1_Boleophthalmus

MSKKDTYDMFEMNGEMDKKKKKKQKKKEKLETMKKEMDIDDHEITIEELEMRYNTSVTKGMTTTFATQVLERDGPNELKPPKGTPEYVKFARQLAGGLQCLMWVAAVICFIAFGIELGRGNLTSFDDLYLAITLIAVVVVTGCFGYYQEFKSTNIIASFKNLVPQQAVVIRDGQKNQINANQLVVGDLVEIKGGDRVPADVRIITSQGCKVDNSSLTGESEPQTRSPECTHENPLETRNIAFFSTTCLEGVATGVIINTGDRTIIGRIASLASGVGNEKTPIAIEIEHFVDIIAGLAIFFGFTFFVVAMFIGYAFLEAMIFFMAIVVAYVPEGLLATVTVCLSLTAKRLARKNCVVKNLEAVETLGSTSVICSDKTGTLTQNRMTVAHLWFDNQIHAADTTEDQSGQSFDQSSETWRFLARVASLCNRATFKPDQEGIPIPKRTVVGDASETALLKFTELTIGNIMDYRNRFKKVVEVPFNSTNKFQLSVHELEDPLDLRYLLVMKGAPERILERCSTILIRGQELPLDDQWKESFQTAYMDLGGLGERVLGFCHVYLNEKEYPRGYSFDPDEMNFPTSELCFAGLISMIDPPRATVPDAVMKCRTAGIRVVMVTGDHPITARAIAANVGIISEGSETVEDIAQRKRIPVEQVNKREARACVISGGQLKDMSSDELDEALRNHPEMVFARTSPQQKLIIVESCQRLGSIVAVTGDGVNDSPALKKADIGVAMGIAGSDAAKNAADMILLDDNFASIVTGVEQGRLIFDNLKKSIAYTLTKNIPELTPYLIYITVSVPLPLGCITILFIELATDIFPSVSLAYEKAESDIMHLKPRNPRRDRLVNEALAVYSYFQIGAIQSFAGFTDYFTAMAQEGWFPLLCVGLRSQWEDVHLQDLQDSYGQEWTFSQRLYQEYTCYTVFFVSIEICQIADVLIRKTRRLSVFQQGFFRNRVLVTAIVFQLCLGNLLCYCPGMPNIFNFMPIRVQWWFVPLPYGILIFVYDEIRKLGVRRHPGSWWDQELYY

>XP_020796885.1_Boleophthalmus

MSECESSERLAAEPESSHRRCSEDDIMVPVLSSKKASELPVNEVTCLLQADLQRGLTHEEVARRRRYHGWNEFDISEEEPLWKKYISQFKDPLILLLLASAVISILMHQFDDAFSITVAIIIVVTVAFVQEYRSEKSLEELGKLVPPECHCVRGGALEQRLARDLVPGDTVCLSVGERVPADLRLFEASDLSVDESSLTGETTPSSKGTAPSPSNDLASRQNIAYMGTLVRCGKAKGIVIGTGENSEFGEVFKMMQAEEAPKTPLQKSMDLLGKQLSLYSFGIIGVIMLVGWLQGKRILDMFTIGVSLAVAAIPEGLPIVVTVTLALGVMRMVKKRAIVKKLPIVETLSCCNVICSDKTGTLTKNEMTVTQIFTSDGLHAEVSGVGYNGAGEVILNGEVIHGFSCPSVSKIVEVGCVCNDSVIRNHTLLGRPTEGALIALAMKMGLESLQQEYVRVEEVPFSSEQKWMAVRCVHRTQQDKPGVFFVKGAFEQVIRFCSSYHSHGTTLPLNHQQRELYQQQISYMGSAGLRVLAFAVGSELSCLSFVGLVGMLDPPRSGVKDAVSTLIGSGVAVKMITGDSQETAVAIAGRLGLCSKGAQCLSGDEVDALDLQQLSQMVPRIAVFYRASPRHKLKIVKSLQNIGAVVAMTGDGVNDAVALKAADIGVAMGKSGTDVCKEAADMILVDDDFQTIMSAIEEGKGIYNNIKNFVRFQLSTSIAALSLISLATLMNFPNPLNAMQILWINIIMDGPPAQSLGVEPVDRDVIRKPPRNLGDSILTRSLLIKVLVSALIIVCGTLFVFWRELQDNVITPRDTTMTFTCFVFFDMFNALSSRSQTRMVYELGVCSNRAFCFAVLASIMGQLLVIYFPPLQSVFQTESLSVTDLLFLVALTSSVCIVSEMIKKVEQWRTERTPPPQNFHEV

***Homo sapiens***

>sp|P13637|AT1A3_HUMAN Sodium/potassium-transporting ATPase subunit alpha-3_Homo

MGDKKDDKDSPKKNKGKERRDLDDLKKEVAMTEHKMSVEEVCRKYNTDCVQGLTHSKAQE

ILARDGPNALTPPPTTPEWVKFCRQLFGGFSILLWIGAILCFLAYGIQAGTEDDPSGDNL

YLGIVLAAVVIITGCFSYYQEAKSSKIMESFKNMVPQQALVIREGEKMQVNAEEVVVGDL

VEIKGGDRVPADLRIISAHGCKVDNSSLTGESEPQTRSPDCTHDNPLETRNITFFSTNCV

EGTARGVVVATGDRTVMGRIATLASGLEVGKTPIAIEIEHFIQLITGVAVFLGVSFFILS

LILGYTWLEAVIFLIGIIVANVPEGLLATVTVCLTLTAKRMARKNCLVKNLEAVETLGST

STICSDKTGTLTQNRMTVAHMWFDNQIHEADTTEDQSGTSFDKSSHTWVALSHIAGLCNR

AVFKGGQDNIPVLKRDVAGDASESALLKCIELSSGSVKLMRERNKKVAEIPFNSTNKYQL

SIHETEDPNDNRYLLVMKGAPERILDRCSTILLQGKEQPLDEEMKEAFQNAYLELGGLGE

RVLGFCHYYLPEEQFPKGFAFDCDDVNFTTDNLCFVGLMSMIDPPRAAVPDAVGKCRSAG

IKVIMVTGDHPITAKAIAKGVGIISEGNETVEDIAARLNIPVSQVNPRDAKACVIHGTDL

KDFTSEQIDEILQNHTEIVFARTSPQQKLIIVEGCQRQGAIVAVTGDGVNDSPALKKADI

GVAMGIAGSDVSKQAADMILLDDNFASIVTGVEEGRLIFDNLKKSIAYTLTSNIPEITPF

LLFIMANIPLPLGTITILCIDLGTDMVPAISLAYEAAESDIMKRQPRNPRTDKLVNERLI

SMAYGQIGMIQALGGFFSYFVILAENGFLPGNLVGIRLNWDDRTVNDLEDSYGQQWTYEQ

RKVVEFTCHTAFFVSIVVVQWADLIICKTRRNSVFQQGMKNKILIFGLFEETALAAFLSY

CPGMDVALRMYPLKPSWWFCAFPYSFLIFVYDEIRKLILRRNPGGWVEKETYY

>sp|P05023|AT1A1_HUMAN Sodium/potassium-transporting ATPase subunit alpha-1_Homo

MGKGVGRDKYEPAAVSEQGDKKGKKGKKDRDMDELKKEVSMDDHKLSLDELHRKYGTDLS

RGLTSARAAEILARDGPNALTPPPTTPEWIKFCRQLFGGFSMLLWIGAILCFLAYSIQAA

TEEEPQNDNLYLGVVLSAVVIITGCFSYYQEAKSSKIMESFKNMVPQQALVIRNGEKMSI

NAEEVVVGDLVEVKGGDRIPADLRIISANGCKVDNSSLTGESEPQTRSPDFTNENPLETR

NIAFFSTNCVEGTARGIVVYTGDRTVMGRIATLASGLEGGQTPIAAEIEHFIHIITGVAV

FLGVSFFILSLILEYTWLEAVIFLIGIIVANVPEGLLATVTVCLTLTAKRMARKNCLVKN

LEAVETLGSTSTICSDKTGTLTQNRMTVAHMWFDNQIHEADTTENQSGVSFDKTSATWLA

LSRIAGLCNRAVFQANQENLPILKRAVAGDASESALLKCIELCCGSVKEMRERYAKIVEI

PFNSTNKYQLSIHKNPNTSEPQHLLVMKGAPERILDRCSSILLHGKEQPLDEELKDAFQN

AYLELGGLGERVLGFCHLFLPDEQFPEGFQFDTDDVNFPIDNLCFVGLISMIDPPRAAVP

DAVGKCRSAGIKVIMVTGDHPITAKAIAKGVGIISEGNETVEDIAARLNIPVSQVNPRDA

KACVVHGSDLKDMTSEQLDDILKYHTEIVFARTSPQQKLIIVEGCQRQGAIVAVTGDGVN

DSPALKKADIGVAMGIAGSDVSKQAADMILLDDNFASIVTGVEEGRLIFDNLKKSIAYTL

TSNIPEITPFLIFIIANIPLPLGTVTILCIDLGTDMVPAISLAYEQAESDIMKRQPRNPK

TDKLVNERLISMAYGQIGMIQALGGFFTYFVILAENGFLPIHLLGLRVDWDDRWINDVED

SYGQQWTYEQRKIVEFTCHTAFFVSIVVVQWADLVICKTRRNSVFQQGMKNKILIFGLFE

ETALAAFLSYCPGMGVALRMYPLKPTWWFCAFPYSLLIFVYDEVRKLIIRRRPGGWVEKE

TYY

>sp|P50993|AT1A2_HUMAN Sodium/potassium-transporting ATPase subunit alpha-2_Homo

MGRGAGREYSPAATTAENGGGKKKQKEKELDELKKEVAMDDHKLSLDELGRKYQVDLSKG

LTNQRAQDVLARDGPNALTPPPTTPEWVKFCRQLFGGFSILLWIGAILCFLAYGIQAAME

DEPSNDNLYLGVVLAAVVIVTGCFSYYQEAKSSKIMDSFKNMVPQQALVIREGEKMQINA

EEVVVGDLVEVKGGDRVPADLRIISSHGCKVDNSSLTGESEPQTRSPEFTHENPLETRNI

CFFSTNCVEGTARGIVIATGDRTVMGRIATLASGLEVGRTPIAMEIEHFIQLITGVAVFL

GVSFFVLSLILGYSWLEAVIFLIGIIVANVPEGLLATVTVCLTLTAKRMARKNCLVKNLE

AVETLGSTSTICSDKTGTLTQNRMTVAHMWFDNQIHEADTTEDQSGATFDKRSPTWTALS

RIAGLCNRAVFKAGQENISVSKRDTAGDASESALLKCIELSCGSVRKMRDRNPKVAEIPF

NSTNKYQLSIHEREDSPQSHVLVMKGAPERILDRCSTILVQGKEIPLDKEMQDAFQNAYM

ELGGLGERVLGFCQLNLPSGKFPRGFKFDTDELNFPTEKLCFVGLMSMIDPPRAAVPDAV

GKCRSAGIKVIMVTGDHPITAKAIAKGVGIISEGNETVEDIAARLNIPMSQVNPREAKAC

VVHGSDLKDMTSEQLDEILKNHTEIVFARTSPQQKLIIVEGCQRQGAIVAVTGDGVNDSP

ALKKADIGIAMGISGSDVSKQAADMILLDDNFASIVTGVEEGRLIFDNLKKSIAYTLTSN

IPEITPFLLFIIANIPLPLGTVTILCIDLGTDMVPAISLAYEAAESDIMKRQPRNSQTDK

LVNERLISMAYGQIGMIQALGGFFTYFVILAENGFLPSRLLGIRLDWDDRTMNDLEDSYG

QEWTYEQRKVVEFTCHTAFFASIVVVQWADLIICKTRRNSVFQQGMKNKILIFGLLEETA

LAAFLSYCPGMGVALRMYPLKVTWWFCAFPYSLLIFIYDEVRKLILRRYPGGWVEKETYY

>sp|Q13733|AT1A4_HUMAN Sodium/potassium-transporting ATPase subunit alpha-4_Homo

MGLWGKKGTVAPHDQSPRRRPKKGLIKKKMVKREKQKRNMEELKKEVVMDDHKLTLEELS

TKYSVDLTKGHSHQRAKEILTRGGPNTVTPPPTTPEWVKFCKQLFGGFSLLLWTGAILCF

VAYSIQIYFNEEPTKDNLYLSIVLSVVVIVTGCFSYYQEAKSSKIMESFKNMVPQQALVI

RGGEKMQINVQEVVLGDLVEIKGGDRVPADLRLISAQGCKVDNSSLTGESEPQSRSPDFT

HENPLETRNICFFSTNCVEGTARGIVIATGDSTVMGRIASLTSGLAVGQTPIAAEIEHFI

HLITVVAVFLGVTFFALSLLLGYGWLEAIIFLIGIIVANVPEGLLATVTVCLTLTAKRMA

RKNCLVKNLEAVETLGSTSTICSDKTGTLTQNRMTVAHMWFDMTVYEADTTEEQTGKTFT

KSSDTWFMLARIAGLCNRADFKANQEILPIAKRATTGDASESALLKFIEQSYSSVAEMRE

KNPKVAEIPFNSTNKYQMSIHLREDSSQTHVLMMKGAPERILEFCSTFLLNGQEYSMNDE

MKEAFQNAYLELGGLGERVLGFCFLNLPSSFSKGFPFNTDEINFPMDNLCFVGLISMIDP

PRAAVPDAVSKCRSAGIKVIMVTGDHPITAKAIAKGVGIISEGTETAEEVAARLKIPISK

VDASAAKAIVVHGAELKDIQSKQLDQILQNHPEIVFARTSPQQKLIIVEGCQRLGAVVAV

TGDGVNDSPALKKADIGIAMGISGSDVSKQAADMILLDDNFASIVTGVEEGRLIFDNLKK

SIMYTLTSNIPEITPFLMFIILGIPLPLGTITILCIDLGTDMVPAISLAYESAESDIMKR

LPRNPKTDNLVNHRLIGMAYGQIGMIQALAGFFTYFVILAENGFRPVDLLGIRLHWEDKY

LNDLEDSYGQQWTYEQRKVVEFTCQTAFFVTIVVVQWADLIISKTRRNSLFQQGMRNKVL

IFGILEETLLAAFLSYTPGMDVALRMYPLKITWWLCAIPYSILIFVYDEIRKLLIRQHPD

GWVERETYY

>sp|P54707|AT12A_HUMAN Potassium-transporting ATPase alpha chain 2_Homo

MHQKTPEIYSVELSGTKDIVKTDKGDGKEKYRGLKNNCLELKKKNHKEEFQKELHLDDHK

LSNRELEEKYGTDIIMGLSSTRAAELLARDGPNSLTPPKQTPEIVKFLKQMVGGFSILLW

VGAFLCWIAYGIQYSSDKSASLNNVYLGCVLGLVVILTGIFAYYQEAKSTNIMSSFNKMI

PQQALVIRDSEKKTIPSEQLVVGDIVEVKGGDQIPADIRVLSSQGCRVDNSSLTGESEPQ

PRSSEFTHENPLETKNICFYSTTCLEGTVTGMVINTGDRTIIGHIASLASGVGNEKTPIA

IEIEHFVHIVAGVAVSIGILFFIIAVSLKYQVLDSIIFLIGIIVANVPEGLLATVTVTLS

LTAKRMAKKNCLVKNLEAVETLGSTSIICSDKTGTLTQNRMTVAHLWFDNQIFVADTSED

HSNQVFDQSSRTWASLSKIITLCNRAEFKPGQENVPIMKKAVIGDASETALLKFSEVILG

DVMEIRKRNRKVAEIPFNSTNKFQLSIHEMDDPHGKRFLMVMKGAPERILEKCSTIMING

EEHPLDKSTAKTFHTAYMELGGLGERVLGFCHLYLPADEFPETYSFDIDAMNFPTSNLCF

VGLLSMIDPPRSTVPDAVTKCRSAGIKVIMVTGDHPITAKAIAKSVGIISANSETVEDIA

HRLNIAVEQVNKRDAKAAVVTGMELKDMSSEQLDEILANYQEIVFARTSPQQKLIIVEGC

QRQDAVVAVTGDGVNDSPALKKADIGIAMGIAGSDAAKNAADMVLLDDNFASIVTGVEEG

RLIFDNLKKTIAYSLTKNIAELCPFLIYIIVGLPLPIGTITILFIDLGTDIIPSIALAYE

KAESDIMNRKPRHKNKDRLVNQPLAVYSYLHIGLMQALGAFLVYFTVYAQEGFLPRTLIN

LRVEWEKDYVNDLKDSYGQEWTRYQREYLEWTGYTAFFVGILVQQIADLIIRKTRRNSIF

QQGLFRNKVIWVGITSQIIIGLILSYGLGSVTALSFTMLRAQYWFVAVPHAILIWVYDEV

RKLFIRLYPGSWWDKNMYY

>sp|O14983|AT2A1_HUMAN Sarcoplasmic/endoplasmic reticulum calcium ATPase 1_Homo

MEAAHAKTTEECLAYFGVSETTGLTPDQVKRNLEKYGLNELPAEEGKTLWELVIEQFEDL

LVRILLLAACISFVLAWFEEGEETITAFVEPFVILLILIANAIVGVWQERNAENAIEALK

EYEPEMGKVYRADRKSVQRIKARDIVPGDIVEVAVGDKVPADIRILAIKSTTLRVDQSIL

TGESVSVIKHTEPVPDPRAVNQDKKNMLFSGTNIAAGKALGIVATTGVGTEIGKIRDQMA

ATEQDKTPLQQKLDEFGEQLSKVISLICVAVWLINIGHFNDPVHGGSWFRGAIYYFKIAV

ALAVAAIPEGLPAVITTCLALGTRRMAKKNAIVRSLPSVETLGCTSVICSDKTGTLTTNQ

MSVCKMFIIDKVDGDICLLNEFSITGSTYAPEGEVLKNDKPVRPGQYDGLVELATICALC

NDSSLDFNEAKGVYEKVGEATETALTTLVEKMNVFNTDVRSLSKVERANACNSVIRQLMK

KEFTLEFSRDRKSMSVYCSPAKSSRAAVGNKMFVKGAPEGVIDRCNYVRVGTTRVPLTGP

VKEKIMAVIKEWGTGRDTLRCLALATRDTPPKREEMVLDDSARFLEYETDLTFVGVVGML

DPPRKEVTGSIQLCRDAGIRVIMITGDNKGTAIAICRRIGIFGENEEVADRAYTGREFDD

LPLAEQREACRRACCFARVEPSHKSKIVEYLQSYDEITAMTGDGVNDAPALKKAEIGIAM

GSGTAVAKTASEMVLADDNFSTIVAAVEEGRAIYNNMKQFIRYLISSNVGEVVCIFLTAA

LGLPEALIPVQLLWVNLVTDGLPATALGFNPPDLDIMDRPPRSPKEPLISGWLFFRYMAI

GGYVGAATVGAAAWWFLYAEDGPHVNYSQLTHFMQCTEDNTHFEGIDCEVFEAPEPMTMA

LSVLVTIEMCNALNSLSENQSLLRMPPWVNIWLLGSICLSMSLHFLILYVDPLPMIFKLR

ALDLTQWLMVLKISLPVIGLDEILKFVARNYLEDPEDERRK

>sp|P98194|AT2C1_HUMAN Calcium-transporting ATPase type 2C member 1_Homo

MKVARFQKIPNGENETMIPVLTSKKASELPVSEVASILQADLQNGLNKCEVSHRRAFHGW

NEFDISEDEPLWKKYISQFKNPLIMLLLASAVISVLMHQFDDAVSITVAILIVVTVAFVQ

EYRSEKSLEELSKLVPPECHCVREGKLEHTLARDLVPGDTVCLSVGDRVPADLRLFEAVD

LSIDESSLTGETTPCSKVTAPQPAATNGDLASRSNIAFMGTLVRCGKAKGVVIGTGENSE

FGEVFKMMQAEEAPKTPLQKSMDLLGKQLSFYSFGIIGIIMLVGWLLGKDILEMFTISVS

LAVAAIPEGLPIVVTVTLALGVMRMVKKRAIVKKLPIVETLGCCNVICSDKTGTLTKNEM

TVTHIFTSDGLHAEVTGVGYNQFGEVIVDGDVVHGFYNPAVSRIVEAGCVCNDAVIRNNT

LMGKPTEGALIALAMKMGLDGLQQDYIRKAEYPFSSEQKWMAVKCVHRTQQDRPEICFMK

GAYEQVIKYCTTYQSKGQTLTLTQQQRDVYQQEKARMGSAGLRVLALASGPELGQLTFLG

LVGIIDPPRTGVKEAVTTLIASGVSIKMITGDSQETAVAIASRLGLYSKTSQSVSGEEID

AMDVQQLSQIVPKVAVFYRASPRHKMKIIKSLQKNGSVVAMTGDGVNDAVALKAADIGVA

MGQTGTDVCKEAADMILVDDDFQTIMSAIEEGKGIYNNIKNFVRFQLSTSIAALTLISLA

TLMNFPNPLNAMQILWINIIMDGPPAQSLGVEPVDKDVIRKPPRNWKDSILTKNLILKIL

VSSIIIVCGTLFVFWRELRDNVITPRDTTMTFTCFVFFDMFNALSSRSQTKSVFEIGLCS

NRMFCYAVLGSIMGQLLVIYFPPLQKVFQTESLSILDLLFLLGLTSSVCIVAEIIKKVER

SREKIQKHVSSTSSSFLEV

>sp|P20020|AT2B1_HUMAN Plasma membrane calcium-transporting ATPase 1_Homo

MGDMANNSVAYSGVKNSLKEANHDGDFGITLAELRALMELRSTDALRKIQESYGDVYGIC

TKLKTSPNEGLSGNPADLERREAVFGKNFIPPKKPKTFLQLVWEALQDVTLIILEIAAIV

SLGLSFYQPPEGDNALCGEVSVGEEEGEGETGWIEGAAILLSVVCVVLVTAFNDWSKEKQ

FRGLQSRIEQEQKFTVIRGGQVIQIPVADITVGDIAQVKYGDLLPADGILIQGNDLKIDE

SSLTGESDHVKKSLDKDPLLLSGTHVMEGSGRMVVTAVGVNSQTGIIFTLLGAGGEEEEK

KDEKKKEKKNKKQDGAIENRNKAKAQDGAAMEMQPLKSEEGGDGDEKDKKKANLPKKEKS

VLQGKLTKLAVQIGKAGLLMSAITVIILVLYFVIDTFWVQKRPWLAECTPIYIQYFVKFF

IIGVTVLVVAVPEGLPLAVTISLAYSVKKMMKDNNLVRHLDACETMGNATAICSDKTGTL

TMNRMTVVQAYINEKHYKKVPEPEAIPPNILSYLVTGISVNCAYTSKILPPEKEGGLPRH

VGNKTECALLGLLLDLKRDYQDVRNEIPEEALYKVYTFNSVRKSMSTVLKNSDGSYRIFS

KGASEIILKKCFKILSANGEAKVFRPRDRDDIVKTVIEPMASEGLRTICLAFRDFPAGEP

EPEWDNENDIVTGLTCIAVVGIEDPVRPEVPDAIKKCQRAGITVRMVTGDNINTARAIAT

KCGILHPGEDFLCLEGKDFNRRIRNEKGEIEQERIDKIWPKLRVLARSSPTDKHTLVKGI

IDSTVSDQRQVVAVTGDGTNDGPALKKADVGFAMGIAGTDVAKEASDIILTDDNFTSIVK

AVMWGRNVYDSISKFLQFQLTVNVVAVIVAFTGACITQDSPLKAVQMLWVNLIMDTLASL

ALATEPPTESLLLRKPYGRNKPLISRTMMKNILGHAFYQLVVVFTLLFAGEKFFDIDSGR

NAPLHAPPSEHYTIVFNTFVLMQLFNEINARKIHGERNVFEGIFNNAIFCTIVLGTFVVQ

IIIVQFGGKPFSCSELSIEQWLWSIFLGMGTLLWGQLISTIPTSRLKFLKEAGHGTQKEE

IPEEELAEDVEEIDHAERELRRGQILWFRGLNRIQTQMDVVNAFQSGSSIQGALRRQPSI

ASQHHDVTNISTPTHIRVVNAFRSSLYEGLEKPESRSSIHNFMTHPEFRIEDSEPHIPLI

DDTDAEDDAPTKRNSSPPPSPNKNNNAVDSGIHLTIEMNKSATSSSPGSPLHSLETSL

**Na/K-ATPase beta-subunits (NKB)**

***Round goby (Neogobius melanostomus)***

>NEME_00028573-RA protein "Similar to ATP1B4 Protein ATP1B4 (Gallus gallus)"

KRPPPLYSGDQWLREGSADGSRDEELLEVHQSSTYDHEASSTHSDCKTSRLRVFFSCFHQ

PPKVLLKHGQELEEEQEELAEHQPLEQEDLNFERFKRRPLPQRTLHQKIIDVKTYLWNAE

TKEFMGRSGKSWSLILLFYTALYTFLAAMFGGCIFCLMWSISPYYPTYNDRVMPPGMTMA

PHLEGHDIHFNASERRSWKKYARSMEEYLRPYNDGAQERKNIRCAQDGKYFMQDDLQENE

ERKACQFKRSWLGECSGMQDPHYGYSQGRPCILLRMNRILGYLPGEGKPINVTCEVKKGP

PEALGQLQFFPKSIFDLKYYPYYGKRRHVNYSSPVVAVRFGSVQYDTPLQIQCKLNGKGI

INDSPTDRYLGSVTFSLVVGA

>NEME_00026002-RA protein "Similar to Atp1b2 Sodium_potassium-transporting ATPase subunit beta-2 (Mus musculus)"

VSLYDKMAKDGEKGEWKEFIWNPRTREFLGRTGSSWGLILLFYLVFYIFLAGLFALTMYV

MLQTLDDHKPTWQDRLATPGMVIRPKADDTFEIIYDLEKTETWDMYTEALNKFLDPYNNS

VQAQKNDECVPDQYFKQEDSGEVKNNPKRSCQFNRTILEDCSGLNDPLYGYQHGKPCIII

KLNRVIGMLPGKSGVAPYVTCGAKKEDSEKIGELVYFPPNGTFNLMYYPYYGKKAQVNYS

QPLVAIKFLNITQNEDVNVECKINAENIPIASERDKFAGRVSFKLRINNKN

>NEME_00022721-RA protein "Similar to atnb233 Sodium_potassium-transporting ATPase subunit beta-233 (Anguilla anguilla)"

MPKDKDDGGWKRFLWNSETGEFLGRTGGSWFKIALFYVIFYSCLAGIFIGTIQGMLLTLS

NYKPTWQDRVAPPGLSHTPKSDKAEMTFNVNDLESYLPYTKALKEFLSKYDEKEQSDPTK

FEDCGRDPADYKHRGDLESDVGSRKACRFPRTTLGPCSGLEDRDFGFGKGKPCIIVKLNR

IVNFRPKPPVTNESIPEEAKPKVLPNVIPIYCTSKKEEDVDKVGEVKYHGIGGGFPLQYY

PYYGKLLHPQYLQPLVALQFTNLTMNTEVRVECKVYGENIYYNDKDRYQGRFEIKFHVND

S

>NEME_00020908-RA protein "Similar to ATP4B Potassium-transporting ATPase subunit beta (Homo sapiens)"

MATMKEKRTCGQRCEDFGNFVWNSDNGTFMGRTPAKWVYISLYYVAFYAVMTALFSLSIW

VLMYTLSPYTPDYQDRLQSPGVMVWPNTYGEEIVEITYNRSNKQSWMKMSNILKNFLEPY

NETIQQSRNNYTCAKGQYFIQSQFTAPHHNKWSCPFCQSMLGACSGISDPTFGYNCTMPC

DGSQNIADIKYFPPNGTMDLSYFPYYGNLAQPNYVNPLVAVRFSLVNQKQATVQCRVVAD

KISYENIHDPYEGKVTFSLKAES

>NEME_00003737-RA protein "Similar to atp1b1 Sodium_potassium-transporting ATPase subunit beta-1 (Anguilla anguilla)"

LNCVCACVRVCVLQPLQLHKSGTRLPFLVSCKGQSTARNHGIAPDPPSRLEDRPPHAQRG

NLPESMSRTKENDGGWKKFVWNSEKKEFLGRTGSSWLKITTFYIIFYGCLAGIFIGTIQA

MLLTLSNYKPTHQDRVAPPGLSHTPRSLKSEISFNVNDEKSYKDYISSIEELLEPYNEDL

QTDGQNYEDCGADPQKYRERGELENGQGKRKACRFSRKWLNACSGVGDNTYGFAKGKPCL

IVKLNRIINFRPRVPTNNETIPEPITSKVKPYLIPIYCKHKRDEDEGKIGEIKYYGFDGG

FPLQYYPYYGKSLHPSTCSLWWPCTSPT

>NEME_00029449-RA protein "Similar to atp1b3 Sodium_potassium-transporting ATPase subunit beta-3 (Xenopus laevis)"

MASAEDKPASKENASSWKDSFYNPRTGEVLGRTAGSWGFNHADKSTIAAAVRQFNKKPEG

HPSSCSSTCVLLLPGRDVRTDHVVMLQTLDDYVPRYRDRVPFRNSAESTRMLLTGLFPFC

DRHDDPPNSIEIMYSKSEPKEYKAYVNQLENFLLRYNDTIQEQNMNCPTGEYFVQDDAEE

KKVCSFKRGFLSACSGLSDTNFGYSDGQPCILLKLNRIIGLKPRGEPYINCTLKKDYPLN

LQYFPHEGRIDKKYFPYYGKKAHENYVQPLVAVKLLLGKDDWNKEHALECRVDGSDLRNN

DERDKYLGRINFKVKVME

***zebrafish (Danio rerio),***

>tr|E7F9R1|E7F9R1_DANRE Sodium/potassium-transporting ATPase subunit beta OS=Danio rerio OX=7955 GN=atp1b4 PE=1 SV=1

MERDSTAGGAEDLLLEDKVSKLRAGSLHKHELAEAMEVEQEGLVEHQPLEQDDLNFEKWK

PKPKPKRTLHEKIDDLKTYLWNAETKEFMGRSGKSWSLILLFYAALYIFLAAMFAGCMCC

LMWSISPYAPTYNDRVMPPGMTMFPHVDTAHGFDIAFNASDRSSWRRYAKTLEAHLKPYD

DGLQSRRNIACKGNAYFMQEDLEESAERKACQFNRSSLGACSGLQDKDFGYSKGRPCILV

KMNRILGYLPGQGTPVNVTCGLKKGSTEVLGEVKFFPNPNFDLRYYPYYGKLRHVNYSSP

LVAVQFLNVQHDTPLHIQCKLNGKGIINDSPTDRFLGSVSFTLEVGA

>tr|F1Q6G8|F1Q6G8_DANRE Sodium/potassium-transporting ATPase subunit beta (Fragment) OS=Danio rerio OX=7955 GN=atp1b1b PE=1 SV=1

XNSLLKWTNKTLQYTMPAQNKDDGGWKKFVWNSEKKEFLGRTGGSWAKIFLFYLIFYGCL

AGIFIGTIQILLLTLSDYKPTWQDRVAPPGLTHFPRSDKSEIAINLDDEVSFLNYVKVMR

EFLTSYDQEKQLDNMQFENCGESPLDYKNRGDLESDVGVRRACQFSREWLGPCSGLDDPY

FGFKEGKPCLIAKLNRIVNFRPKPPVSNESIPEEVQHKVQPYLIPIHCTNKKEEDAGKLG

EVRYYGFGGGFPLQYYPYYGKLLHPQYLQPLVAIQFLNITPNTDMRIECKVYGENIYYHD

KDRYQGRFDVKFNIKKS

>tr|Q9DGL3|Q9DGL3_DANRE Sodium/potassium-transporting ATPase subunit beta OS=Danio rerio OX=7955 GN=atp1b1a PE=1 SV=1

MPANKDGDGGWKSFIWNSDKKEFLGRTGCSWLKIFIFYVIFYGCLAGIFIGTIQAMLLTL

SNYKPTYQDRVAPPGLSHSPRPDKAEISYNINDESTYMPYVNHIDAFLKAYNQDIQEDNT

KFEDCGDKPEFYTDRGELESDNGVRKACRFRREWLGECSGQKDEKQKNYGFDDGQPCLIV

KLNRIVNFMPRPPASNDSIPEAVRPKLQGNVIPIHCSSKREEEANLLGQIKYFGVGNGFP

LQYYPYYGKLLQPQYLQPLVAIKFYNITTDVDVRVECKVYGENIDYSEKDRSQGRFDIKF

TIKTKS

>tr|Q6NYH5|Q6NYH5_DANRE Sodium/potassium-transporting ATPase subunit beta OS=Danio rerio OX=7955 GN=atp1b1a PE=2 SV=1

MPANKDGDGGWKSFIWNSDKKEFLGRTGCSWLKIFIFYVIFYGCLAGIFIGTIQAMLLTL

SNYKPTYQDRVAPPGLSHSPRPDKAEINYNINDESTYLPYVNHIDAFLKAYNEDVQKDDT

KFEECGDKPQFYTDRGELESDNGVRKACRFRREWLGECSGQKDEKLKNYGFDDGQPCLIV

KLNRIVNFMPRPPASNDSIPEAVRPKLQGNVIPIHCSSKREEEANLLGQIKYFGLGTGFP

LQYYPYYGKLLQPQYLQPLVAIKFYNITTDVDVRVECKVYGENIDYSEKDRSQGRFDIKF

TIKTKS

>tr|Q7ZVX2|Q7ZVX2_DANRE Sodium/potassium-transporting ATPase subunit beta OS=Danio rerio OX=7955 GN=atp1b1a PE=2 SV=1

MPANKDGDGGWKSFIWNSDKKEFLGRTGCSWLKIFIFYVIFYGCLAGIFIGTIQAMLLTL

SNYKPTYQDRVAPPGLSHSPRPDKAEISYNINDESTYMPYVNHIDAFLKAYNKDIQEDNT

KFEDCGDKPQFYTDRGELESDNGVRKACRFRREWLGECSGQKDEKQKNYGFDDGQPCLIV

KLNRIVNFMPRPPASNESIPEAVRPKLQGNVIPIHCSSKREEEANLLGQIKYFGLGTGFP

LQYYPYYGKLLQPQYLQPLVAIKFYNITTDVDVRVECKVYGENIDYSEKDRSQGRFDIKF

TIKTKS

>tr|Q9DEY4|Q9DEY4_DANRE Sodium/potassium-transporting ATPase subunit beta OS=Danio rerio OX=7955 GN=atp1b1b PE=2 SV=1

MPAQNKDDGGWKKFVWNSEKKEFLGRTGGSWAKIFLFYLIFYGCLAGIFIGTIQILLLTL

SDYKPTWQDRVAPPGLTHFPRSDKSEIAINLDDEVSFLNYVKVMREFLTSYDQEKQLDNM

QFENCGESPLDYKNRGDLESDVGVRRACQFSREWLGPCSGLDDPYFGFKEGKPCLIAKLN

RIVNFRPKPPVSNESIPEEVQHKVQPYLIPIHCTNKKEEDAGKLGEVRYYGFGGGFPLQY

YPYYGKLLHPQYLQPLVAIQFLNITPNTDMRIECKVYGENIYYHDKDRYQGRFDVKFNIK

KS

>tr|A0A0R4IE14|A0A0R4IE14_DANRE Sodium/potassium-transporting ATPase subunit beta OS=Danio rerio OX=7955 GN=atp1b2a PE=1 SV=1

MAKEDEKKESGSWKDFFWNPRTHELLGRTASSWGLILLFYLVFYTFLAGVFCLTMYVMLL

TLDDYQPTWQDRLATPGMMIRPKGEALEIVYSRENTESWELYVQALDSFLKPYNNSQQAV

NNDDCTPDQFNIQEDSGNVRNNPKRSCRFNRTTLEDCSGLTDRFYGYPDGKPCILIKLNR

VIGMKPGKDGQSPYVTCGAKRYKEGDEWKEDAESIGEIAYFPPNGTFNLMYYPYYGMKAQ

VNYSQPLVAVKFMNISFNTDVNVECKINSNTITEFSERDKFAGRVSFKLRVNN

>tr|Q90Z34|Q90Z34_DANRE Sodium/potassium-transporting ATPase subunit beta OS=Danio rerio OX=7955 GN=atp1b2b PE=1 SV=1

MAKDDKNGWKEFFWNPRTREFCGRTASSWGLILLFYLAFYIFLAGLFTLTMYVMLQTLDD

HRPTYQDRLSTPGMMIRPKGEQLEIAYTTEYTETWERYVQALNNFLSPYNDTVQTQKNYE

CKPDQFFIQEDSGGLKNFPKRSCQFKRSILEKCSGITDRFYGYDEGKPCIIIKLNRVIGL

LPAKDGQPPYVTCGAKKYKVGKDEWTDDSDKLGELAYYPPNGTFNLMYYPYYGKKAQVNY

SQPLVAVKFLNITRNEDVNVECKINSNNIPEGSERDKFAGRVSFTLRINSRD

>tr|A0A0R4IAE6|A0A0R4IAE6_DANRE Sodium/potassium-transporting ATPase subunit beta OS=Danio rerio OX=7955 GN=atp1b2a PE=1 SV=1

MAKEDEKKESGSWKDFFWNPRTHELLGRTASSWGLILLFYLVFYTFLAGVFCLTMYVMLL

TLDDYQPTWQDRLATPGMMIRPKGEALEIVYSRENTESWELYVQALDSFLKPYNNSQQAV

NNDDCTPDQFNIQEDSGNVRNNPKRSCRFNRTTLEDCSGLTDRFYGYPDGKPCILIKLNR

VIGMKPGKDGQSPYVTCGAKKEDAESIGEIAYFPPNGTFNLIIHKSFDLKFCVNYSQPLV

AVKFMNISFNTDVNVECKINSNTITEFSERDKFAGRVSFKLRVNN

>tr|Q9DGL2|Q9DGL2_DANRE Sodium/potassium-transporting ATPase subunit beta OS=Danio rerio OX=7955 GN=atp1b2a PE=1 SV=1

MAKEDEKKESGSWKDFFWNPRTHELLGRTASSWGLILLFYLVFYTFLAGVFCLTMYVMLL

TLDDYQPTWQDRLATPGMMIRPKGEALEIVYSRENTESWELYVQALDSFLKPYNNSQQAV

NNDDCTPDQFNIQEDSGNVRNNPKRSCRFNRTTLEDCSGLTDRFYGYPDGKPCILIKLNR

VIGMKPGKDGQSPYVTCGAKKEDAESIGEIAYFPPNGTFNLMYYPYYGMKAQVNYSQPLV

AVKFMNISFNTDVNVECKINSNTITEFSERDKFAGRVSFKLRVNN

>tr|Q8QHI1|Q8QHI1_DANRE Sodium/potassium-transporting ATPase subunit beta OS=Danio rerio OX=7955 GN=atp1b3a PE=1 SV=1

MSKKSENPAEGKEPESSWKDAIYNPRTGELFGRTARNWGLILLFYLVFYGFLAAMFVFTL

WVMLQTLNDDTPKYRDRVASPGLVIRPNSLNIEFNRSDPLEYGQYVQHLESFLHQYNDSE

QAKNDLCMAGQYSEQDGESLKKVCQFKRSLLYSCSGMEDTTFGYAKGQPCVIVKMNRIIG

LKPSGDPYINCTSKSVKPLQMQYFPHEGTIDRMYFPYYGKKTHKGYVQPLVAVKLLLKKE

DYNSELIVECKVEGSNLKNNDERDKFLGRVTFRVLVTE

>tr|F1QYB9|F1QYB9_DANRE Sodium/potassium-transporting ATPase subunit beta OS=Danio rerio OX=7955 GN=atp1b3a PE=1 SV=2

MSKKSENQRRAKSREQLERRDLNPRTGELFGRTARNWGLILLFYLVFYGFLAAMFVFTLW

VMLQTLNDDTPKYRDRVASPGLVIRPNSLNIEFNRSDPLEYGQYVQHLESFLHQYNDSEQ

AKNDLCMAGQYSEQDGESLKKVCQFKRSLLYSCSGMEDTTFGYAKGQPCVIVKMNRIIGL

KPSGDPYINCTSKSVKPLQMQYFPHEGTIDRMYFPYYGKKTHKGYVQPLVAVKLLLKKED

YNSELIVECKVEGSNLKNNDERDKFLGRVTFRVLVTE

>tr|Q90266|Q90266_DANRE Sodium/potassium-transporting ATPase subunit beta OS=Danio rerio OX=7955 GN=atp1b3a PE=2 SV=1

MSKKSENQRRAKSREQLERRDLHPRTGELFGRTARNWGLILLFYLVFYGFLAAMFVFTLW

VMLQTLNDDTPKYRDRVASPGLVIRPNSLNIEFNRSDPLEYGQYVQHLESFLHQYNDSEQ

AKNDLCYGGTVPEQDGESLKKVCQFKRSLLYSCSGMEDTTFGYAKGQPCVIVKMNRIIGL

KPSGDPYINCTSKSVKPLQMTSISPMRTIDRMYFPYYGKKTHKGYVQPLVAVKLLLKKED

YNSELIVECKVEGSNLKNNDERDKFLGRVTFRVLVTE

>tr|A0A0R4IEL3|A0A0R4IEL3_DANRE Sodium/potassium-transporting ATPase subunit beta OS=Danio rerio OX=7955 GN=atp1b3a PE=1 SV=1

MSKKSENQRRAKSREQLERRDLNPRTGELFGRTARNWGLILLFYLVFYGFLAAMFVFTLW

VMLQTLNDDTPKYRDRVASPGLVIRPNSLNIEFNRSDPLEYGQYVQHLESFLHQYNDSEQ

AKMTCYGGTVSEQDGESLKKVCQFKRSLLYSCSGMEDTTFGYAKGQPCVIVKMNRIIGLK

PSGDPYINCTSKSVKPLQMQYFPHEGTIDRMYFPYYGKKTHKGYVQPLVAVKLLLKKEDY

NSELIVECKVEGSNLKNNDERDKFLGRVTFRVLVTE

>tr|Q9DGI4|Q9DGI4_DANRE Sodium/potassium-transporting ATPase subunit beta OS=Danio rerio OX=7955 GN=atp1b3b PE=1 SV=1

MANKEEKADEKQSSWKDFIYNPRTGEFIGRTASSWALIFLFYLVFYGFLAGMFTLTMWVM

LQTLDDHTPKYRDRVANPGLMIRPRSLDIAFNRSIPQQYSKYVQHLEAFLQSYNDSLQEA

NEPCQEGMYFEQDDVEEKKVCQFKRSQLRQCSGLSDTTFGYSEGNPCIIVKMNRVIGLKP

RGDPHIACTVKGDGTLQMQLYPDEGKIDKSFFPYYGKILHKNYVQPLVAVKLMLGENDYN

IEHTVECKVEGSDLLNNDERDKFLGRVTFRINVQK

***three spine stickleback (Gasterosteus aculeatus)***

>tr|G3QAN7|G3QAN7_GASAC Sodium/potassium-transporting ATPase subunit beta OS=Gasterosteus aculeatus OX=69293 PE=3 SV=1

MSKDGEKSGWKEYIWNPRTREFMGRTASSWGLILLFYVVFYIFLAGLFALTMYVMLQTLD

DYKPTWQDRLATPGMVIRPKSDETYEIVYDTRNTETWDLYAQALDKFLTPYNNSLQTEKN

KECTPDEYFLQEDTGDVKNNPKRSCQFNRTILEACSGINDRYYGYQIGQPCIIIKLNRVI

GMLPGKDGQAPFVTCAAKKYKIGKDQWREDTDKIGEMVYFPPNGIFNLMYYPYYGKKAQV

NYSQPLVAVKFLNITTNEDVNIECRINATNIPIGSERDKFAGRVSFKLRINTK

>tr|G3PKB5|G3PKB5_GASAC Sodium/potassium-transporting ATPase subunit beta OS=Gasterosteus aculeatus OX=69293 PE=3 SV=1

MQANKDSDGGWRTFVWNSEKKEFLGRTGSSWLKIFLFYVIFYGCLAGIFIGTIQALLLTL

SNYKPTYQDRVAPPGLSHTPRSEKSEIAFSLSDASSYKKYTDAMTKFLKKYDDDKQTDQM

KFEDCGGVPENYKDRGAPEDNNPGQRIGWRLKRSLVGVWPRLGNRSGFHTREAWLIVKLN

RIVNFSFQPPMNNSVEEFLLPKLKPNLIPIFCKNKREEDADKIGEIKFFGLGEGFPLQYY

PYYGKLLHPQYLQPLVAIQFNNLTLNEELRIECKVFGENIGYSEKDRYQGRFDVKFNIAS

>tr|G3PRI4|G3PRI4_GASAC Sodium/potassium-transporting ATPase subunit beta OS=Gasterosteus aculeatus OX=69293 GN=ATP4B PE=3 SV=1

MAALKEKRNCGQRCEDFGRFVWNSDEGKLLGRTPEKWVYISLYYVAFYVVMTSLFCLAIW

TLMYTLDPYAPDYQDRLKSPGVMVWPDTYGEEIVEISYNTSDKSSWMKMTNILNEFLEPY

NDTKQLECNKHNCTRGKYFIQDTFSAPNHTKWACPFTQSMLGACSGHEDPTFGYNCTMPC

VIIKMNRIISFLPSNNTEHPAYVNCTVLKSGQEGQHNVDKIEYFPEGGSMDLSYFPYYGR

LAQPTYINPLVAVRFSLVNGKKAKIQCRVVSKIISYENYHDPYEGKVIFQLEAVK

>tr|G3PRI3|G3PRI3_GASAC Sodium/potassium-transporting ATPase subunit beta OS=Gasterosteus aculeatus OX=69293 GN=ATP4B PE=3 SV=1

MAALKEKRNCGQRCEDFGRFVWNSDEGKLLGRTPEKWVYISLYYVAFYVVMTSLFCLAIW

TLMYTLDPYAPDYQDRLKSPGVMVWPDTYGEEIVEISYNTSDKSSWMKMTNILNEFLEPY

NDTKQLECNKHNCTRGKYFIQDTFSAPNHTKWACPFTQSMLGACSGHEDPTFGYNCTMPC

VIIKMNRIISFLPSNNTEHPAYVNCTVLEGQHNVDKIEYFPEGGSMDLSYFPYYGRLAQP

TYINPLVAVRFSLVNGKKAKIQCRVVSKIISYENYHDPYEGKVIFQLEAVK

>tr|G3PRG7|G3PRG7_GASAC Sodium/potassium-transporting ATPase subunit beta OS=Gasterosteus aculeatus OX=69293 PE=3 SV=1

TTMSSTKDDGGWKRFVWDSDKGELLGRTGGSWLKIIGFYIIFYGCLAGVFVGTIQALLLT

LSNYKPTWQDRVAPPGLSHTPRSDKSEVSFNLNDVETYLPYTKALKDFMAKYNEDIQKDQ

MKFEDCTDSPADYKNRGDLESDQGIRKACRFYRNLLGPCSGLDDPDFGFKEGKPCIIVKL

NRIVNFRPKPPTSNESIPDEAQHKVQPNVIPIYCTNKREEDAGKIGEIKYHGIGGGFPLQ

YYPYYGKLLHPQYLQPLLALQFTNLTLNTELRIECKVFGDNIGYSEKDRYQGRFEIKLSI

NS

>tr|G3PRF6|G3PRF6_GASAC Sodium/potassium-transporting ATPase subunit beta OS=Gasterosteus aculeatus OX=69293 PE=3 SV=1

MSSTKDDGGWKRFVWDSDKGELLGRTGGSWLKIIGFYIIFYGCLAGVFVGTIQALLLTLS

NYKPTWQDRVAPPGLSHTPRSDKSEVSFNLNDVETYLPYTKALKDFMAKYNEDIQKDQMK

FEDCTDSPADYKNRGDLESDQGIRKACRFYRNLLGPCSGLDDPDFGFKEGKPCIIVKLNR

IVNFRPKVMLNLQFTCYACLVSSCTCKREEDAGKIGEIKYHGIGGGFPLQYYPYYGKLLH

PQYLQPLLALQFTNLTLNTELRIECKVFGDNIGYSEKDRYQGRFEIKLSINSL

>tr|G3N891|G3N891_GASAC Sodium/potassium-transporting ATPase subunit beta OS=Gasterosteus aculeatus OX=69293 PE=3 SV=1

MSGTREEDRIGSSEWRDCFWNPRTHELLGRTASSWGLILLFYLVFYLFLAGMFVLTMYIM

LLTLDDYKPTWQDRLTTPGMMIRPKGDQLEISFTVSETESWDGFTNNLNSFLAPYNGSYQ

VQTNDKCSPDQYFIQEDSGEVRNNPKRSCQFNRTMLEECSGILDRFYGYSRGQPCILIKL

NRVIGMLPGKDGQSPYVTCGAKREDGDKIGPLAYYPPNGTFNLMYYPYYGKKAQVNYTQP

LVAVKFLNASLNTDISVECKINSNTLAEGSERDKFAGRVSFKLRIND

>tr|G3QAN6|G3QAN6_GASAC Sodium/potassium-transporting ATPase subunit beta OS=Gasterosteus aculeatus OX=69293 PE=3 SV=1

APNLHDKMSKDGEKSGWKEYIWNPRTREFMGRTASSWGLILLFYVVFYIFLAGLFALTMY

VMLQTLDDYKPTWQDRLATPGMVIRPKSDETYEIVYDTRNTETWDLYAQALDKFLTPYNN

SLQTEKNKECTPDEYFLQEDTGDVKNNPKRSCQFNRTILEACSGINDRYYGYQIGQPCII

IKLNRVIGMLPGKDGQAPFVTCAAKREDTDKIGEMVYFPPNGIFNLMYYPYYGKKAQVNY

SQPLVAVKFLNITTNEDVNIECRINATNIPIGSERDKFAGRVSFKLRINT

>tr|G3PT60|G3PT60_GASAC Sodium/potassium-transporting ATPase subunit beta OS=Gasterosteus aculeatus OX=69293 PE=3 SV=1

MASTEDKAAKQESATGWKDSFYNPRTGEVLGRTGSSWALILLFYLVFYCFLAGMFALTMW

VLLLTLDDYVPKYRDRVPYPGLVIRPNSLDLSFNKSDPLKYVQYVKHLESFLQRYNDSQQ

DKNDDCPPGEYFLQDDTEDMSKKACLFKRGFLSLCSGLSDTNFGYSEGKPCVLLKMNRII

GLKPRGDPYINCTVKRDTPIQMQYFPSEGRIDKKYFPYYGKKAHESYVQPLVAVELLLTK

DDYNKELPVECRVEGSDLRNNDERDKFLGRVTFRVTVVE

***tilapia (Oreochromis niloticus)***

>tr|I3J8Y6|I3J8Y6_ORENI Sodium/potassium-transporting ATPase subunit beta OS=Oreochromis niloticus OX=8128 GN=LOC100693295 PE=3 SV=1

MAKDGEKSDWKEFLWNPRTREFLGRTASSWGLIFLFYLIFYTCLAGMFALTMYVMLQTLD

EHTPTWQDRLSTPGLVIRPRADDTFEIVYTIEDTESWDLYAQALDKFLQPYNDSLQAQKN

HECAPDKYFIQEDSGEVKNNPKRSCQFNRTVLQNCSGIDDRYYGYREGQPCIIIKLNRVI

GLLPGKDNQAPYVTCAAKKYRVGKDQWKDDADKIGELIYFPPNGTINPMYFPYYGKKAQV

NYSQPLVAVKFLNITHNEDVNIECKINAENIPVGGERDKFAGRVSFKLRINSKN

>tr|I3KT86|I3KT86_ORENI Sodium/potassium-transporting ATPase subunit beta OS=Oreochromis niloticus OX=8128 GN=atp1b1 PE=3 SV=1

MSGNKDSDGGWKNFVWNSEKKEFLGRTGGSWFKILLFYVIFYGCLAGIFVGTIQALLLTL

SKDKPTYQDRVAPPGLSHTPRSDKSEISFTRSDQSSYSKYVQSMNEFLELYNDTKQNGDP

YEECGKFPGTYKDRSMEEKKKVCKFLRSWLKNCSGITDPDYGFMEGSPCVIIKLNRIVNF

RPKAPSNNSLLEELQAKITPNEIPIYCKPKRAEDKGQIEEIKYFGISQGFPLQYYPYYGK

QLHPDYLQPLMAVQFVNVTVGKEIRIECKAFGDNIDYSEKDRYQGRFDIKLTVS

>tr|I3KC45|I3KC45_ORENI Sodium/potassium-transporting ATPase subunit beta OS=Oreochromis niloticus OX=8128 GN=LOC100695267 PE=3 SV=1

MSSSPQVQTQKPNQDQTQEPNQDQNQDAKEEVKAEEKEEKKEEKREKKEDKKEANEANEA

KREQKKEKQGSWKDFIYNPRTGEFLGRTAGSWGLILLFYLIFYGFLGGMFSLTMWVMLQT

LDENVPRHQDRVASPGLVIRPHAVEITFNRSDPENYKHYIRQLHDLLQSYNDSIQERNEL

CMVGEYTTQDNEPVKKVCQFKRSTLRQCSGLPDPSFGFKEGKPCIIIKMNRVIGLKPEGD

PYINCTAKRATSLQMQYFPSEGRLDKMFFPYYGKKSHPDYVQPLVAVKLLFTKEDYNIEQ

TVECKVEGSNLRNDDERDKFLGRVTFRVKVSE

>tr|I3J761|I3J761_ORENI Sodium/potassium-transporting ATPase subunit beta OS=Oreochromis niloticus OX=8128 GN=LOC100694365 PE=3 SV=1

MPSDKKDDGGWKKFLWNSETGELLGRTGGSWFKITLFYVIFYGCLAGIFIGTIQAMLLTL

SEYKPTWQDRVAPPGLTHTPRSDKAELAFNPRAVETFLPHTKALREFLNNYDESKQKDQM

KYEDCGDEPADYKNRGDLDSDVGVRKACRFPRALLGPCSGLEDTEFGFKEGKPCLIVKLN

RIVNYRPRPPTSNESIPEEAQPKVQPNVIPIYCTSKKEEDADKIGEIKYYGIGEGFPLQY

YPYYGKKLHPQYLQPLVALQFTNLTRNTELRIECKVFGDNIDYSEKDRYQGRFEIKIQVD

ES

>tr|I3J546|I3J546_ORENI Sodium/potassium-transporting ATPase subunit beta OS=Oreochromis niloticus OX=8128 GN=ATP4B PE=3 SV=1

MATLKEKRTCGQRCEDFGHFVWNSENGTFMGRTPEKWVYISLYYVAFYVIMTGLFSLAIW

VLMYTISPYTPDYQDRLSSPGVMVWPDTYGEEDVEISYNTSDKASCMAMANILHDFLKPY

NDTKQLECNNYNCTKGKYFIQKTFSAPHHTKWVCPFTQSMLGPCSGIEDPTFGYNSTMPC

VIIKMNRIIDFLPSNNTALPPYVNCTILDGEDSVAKIEYFPDGGIMDPSYFPYYGKLAQP

TYVNPLVAVRFSLVGEKDAKIQCRVVSEKISYENIHDPYEGKVVFYLKAVK

>tr|I3JHR1|I3JHR1_ORENI Sodium/potassium-transporting ATPase subunit beta OS=Oreochromis niloticus OX=8128 GN=LOC102079915 PE=3 SV=1

SIMSGSKEDDRKGSSEWREFFWNPRTHELLGRTAASWGLILLFYLIFYLCLAGMFAFTMY

IMLLTLDDYKPTWQDRLATPGMMIRPKGNQLEITFSTSDTESWDGFVHNLNSFLSPYNDS

YQVQTNDECTPDEYFIQEDSGEVRNNPKRSCQFNRTILEECSGVWDRFYGYDKGQPCILI

KLNRVIGMLPGKDRQSPYVTCAAKRYRVGENEWREDSDKIGLLAYYPPNGTFNLMYYPYY

GKKAQVNYTQPLVAVKFLNASLNTDINVECKVTSNTLAAGSDRDKFAGRVSFKLRIND

>tr|I3J8Y5|I3J8Y5_ORENI Sodium/potassium-transporting ATPase subunit beta OS=Oreochromis niloticus OX=8128 GN=LOC100693295 PE=3 SV=1

MAKDGEKSDWKEFLWNPRTREFLGRTASSWGLIFLFYLIFYTCLAGMFALTMYVMLQTLD

EHTPTWQDRLSTPGLVIRPRADDTFEIVYTIEDTESWDLYAQALDKFLQPYNDSLQAQKN

HECAPDKYFIQEDSGEVKNNPKRSCQFNRTVLQNCSGIDDRYYGYREGQPCIIIKLNRVI

GLLPGKDNQAPYVTCAAKKDDADKIGELIYFPPNGTINPMYFPYYGKKAQVNYSQPLVAV

KFLNITHNEDVNIECKINAENIPVGGERDKFAGRVSFKLRINSKN

>tr|I3JHR0|I3JHR0_ORENI Sodium/potassium-transporting ATPase subunit beta OS=Oreochromis niloticus OX=8128 GN=LOC102079915 PE=3 SV=1

MSGSKEDDRKGSSEWREFFWNPRTHELLGRTAASWGLILLFYLIFYLCLAGMFAFTMYIM

LLTLDDYKPTWQDRLATPGMMIRPKGNQLEITFSTSDTESWDGFVHNLNSFLSPYNDSYQ

VQTNDECTPDEYFIQEDSGEVRNNPKRSCQFNRTILEECSGVWDRFYGYDKGQPCILIKL

NRVIGMLPGKDRQSPYVTCAAKREDSDKIGLLAYYPPNGTFNLMYYPYYGKKAQVNYTQP

LVAVKFLNASLNTDINVECKVTSNTLAAGSDRDKFAGRVSFKLRINDK

>tr|I3J2V6|I3J2V6_ORENI Sodium/potassium-transporting ATPase subunit beta OS=Oreochromis niloticus OX=8128 GN=atp1b4 PE=3 SV=1

MEPSSTEGGAEENLCYRSYSDQEACHFFPAPHKVILKHGQELEEEQEELAEHQPLEQEDL

NFERWKRRPIPKRTLHQKIDDLKKYLWNAETNEFMGRSGKSWSLILLFYAALYLFLAAMF

GGCLFCLMWSISPYHPTFNDRVMPPGMTMAPHLEGHEIAFNASDRKSWRKYARSMEEYLR

PYNDAAQQRKNIPCDKETYFMQDDLDEAAERKACQFKRSWLGHCSGLQDPNFGYSQGRPC

ILLRMNRILGYLPGQGKPINVTCGVKKGTPEVLGEMEFFPKSIFENKYYPYYGKLRHVNY

SSPVVAVRFMGVQQGTNIQVQCKLNGKGIINDSPTDRYLGSVTFSLEVGA

>tr|I3J2V5|I3J2V5_ORENI Sodium/potassium-transporting ATPase subunit beta OS=Oreochromis niloticus OX=8128 GN=atp1b4 PE=3 SV=1

MEPSSTEGGAEEKLVRSLPHKVILKHGQELEEEQEELAEHQPLEQEDLNFERWKRRPIPK

RTLHQKIDDLKKYLWNAETNEFMGRSGKSWSLILLFYAALYLFLAAMFGGCLFCLMWSIS

PYHPTFNDRVMPPGMTMAPHLEGHEIAFNASDRKSWRKYARSMEEYLRPYNDAAQQRKNI

PCDKETYFMQDDLDEAAERKACQFKRSWLGHCSGLQDPNFGYSQGRPCILLRMNRILGYL

PGQGKPINVTCGVKKGTPEVLGEMEFFPKSIFENKYYPYYGKLRHVNYSSPVVAVRFMGV

QQGTNIQVQCKLNGKGIINDSPTDRYLGSVTFSLEVGA

>tr|I3JWW2|I3JWW2_ORENI Sodium/potassium-transporting ATPase subunit beta OS=Oreochromis niloticus OX=8128 GN=atp1b3 PE=3 SV=1

TMASTEDKAANKENVSSWKDSIYNPRTGELLGRTASSWALILLFYLVFYCFLAGMFALTM

WVMLLTLDDYVPRYRDRVPEPGLVIRPNSLDITFNKSDSKNYRTYVNHLESFLQRYNDSM

QENNADCIPGEYYMQDGGEMTKKVCPFRRTSLSLCSGLSDTDFGYQEGKPCVLLKMNRII

GLKPIGDPYINCTVKKDHPVQMHYFPSEGRIDKMYFPYYGKKAHENYMQPLVAVKLLLTK

EDYNKELAVECRVEGTNLRNNDERDKFLGRVTFRVKVVE

***mudskipper (Boleophthalmus pectinirostris)***

>XP_020779213.1 protein ATP1B4 [Boleophthalmus pectinirostris]

MEPTPTEGEAEAQTQQVQPPEEEVKPVKLSRSPPHKVILKHGQELEEEQEELAEHQPLEQEDLNFERFKRRPLPQRTLHQKIDDLKTYLWNAETKEFMGRSGKSWSLILLFYAALYAFLAAMFGGCVFCLMWSISPYHPTYNDRVMPPGMTMAPHLEGHDIHFNASERRSWKKYARSMDEHLRPYNDGAQDRKNIRCTQDGKYFMQDDLREDEERKACQFKRSWLGECSGMQDPHYGYSQGRPCILLRMNRILGYLPGEGKPINVTCEVKKGPPEALGQLQFFPKSIFDLKYYPYYGKLRHVNYSSPVIAVRFASVQYDTPLQIQCKLNGKGIINDSPTDRYLGSVTFSLVVGA

>XP_020796323.1 sodium/potassium-transporting ATPase subunit beta-2-like [Boleophthalmus pectinirostris]

MSGGGAKEEPRGSSWDFIWNPRTHELMGRTATSWGLIFLFYLVFYLFLAGMFALTMYVMLLTLDDYRPTWQDRLTTPGMMIRPKGDQLHIAFSVSKAESYRDLTSNLDVFLSPYNDSVQAQTNDNCPPNQYFIQEDSGEVRNNPKRSCQFNRTALDLCSGLSDSSYGYSSGQPCVLLKLNRVIGMLPGKDGQSPYVSCGAKKEDVNVGPLAYFPPNGTFNLMYYPYYGNKAQVNYTQPLVAVKFLNASLNTDLSVECRIISNTLGDVYANDRDKFAGRVSFTLRINA

>XP_020778771.1 sodium/potassium-transporting ATPase subunit beta-2-like [Boleophthalmus pectinirostris]

MAKDGEKGEWKEFIWNPRTREFLGRTASSWGLLLLFYLIFYIFLAGLFALTMYVMLQTLDDHKPTWQDRLSTPGMVIRPKADETFEIVYDIQNTETWDLYAQALDKFLAPYNNSVQVMKNDECVPDQYYKQEDSGEVKNNPKRSCQFNRTILEACSGIGDPHYGYQEGKPCIIIKLNRVIGMLPGKNGMAPYVTCGAKRYKVGKDEWREDSDKIGELMYFPPNGTFNLMYYPYYGKKAQVNYSQPLVAVKFLNVTHNQDVNIECKINADNIPIGSERDKFAGRVSFKLRINTKN

>XP_020794724.1 potassium-transporting ATPase subunit beta [Boleophthalmus pectinirostris]

MATLKEKRTCGQRCEDFGHFVWNSDNGTFMGRTPVKWVYISLYYVAFYIVMTALFSLAIYVLMYTLSPYAPDYQDRLKSPGVMVWPDTYGEEIVEISYNSSDKKSWKKMATILRHFLEPYNDTTQLIVNNVTCTKGKYFIQSHFTGHNHTKCSCPFTQSMLGPCSGLEDPTFGYNSSMPCVIIKMNRIINFLPSNITGVAPYVNCTILDGEENVENIEYFPKNGTMDLSYFPYYGRLAQPNYVNPLVAVRFNLVKEKEAKIECRVVSDVISYQNIHDPYEGKVVFDLSAES

>XP_020772934.1 sodium/potassium-transporting ATPase subunit beta-3-like [Boleophthalmus pectinirostris]

MASTEDKPANKENTSSWKDSIYNPRTGELLGRTASSWALILLFYLVFYCFLAGMFALTMWVMLLTLDDYVPRYRDRVPFPGMVIRPNTMEITFTKSDASKYNGYVSQLESFLQRYNDTLQEQNVQDCVTGQYFLQDDVEEKKVCSFKRGSLKACSGLSDTSFGYSDGQPCVLLKLNRIIGLMPRGDPYINCTVKVNRPLQMAYYPDEGRIDKKYFPYYGKKAHENYVQPLVAVKLLLGKEDLNKEHSVECRVEGTDLRNNDERDKWLGRVSFKVKVME

>XP_020797469.1 sodium/potassium-transporting ATPase subunit beta-233-like [Boleophthalmus pectinirostris]

MPKDKDDGGWKKFLWNSDTGEFLGRTGGSWFKITLFYIIFYGCLAGVFIGTIQAMLLTLSSYKPTWQDRVAPPGLTHTPRSDKAELSFSVNDVESYLPYTKALKDFLAKYDDEKQNDPTKFEDCGDVPADFKHRGDLESDVGTRRACKFPRSLLGPCSGIEDRNFGFADGKPCIIVKLNRIVNFRPRPPVSNESVPVEAQSKVQPFVIPLYCTNKKEEDAGKVGEIKYHGIGGGFPLQYYPYYGKLLHPQYLQPLVAIQFTNLTMNTELRIECKVYGQNIDYNEKDRYQGRFDIKIQINNL

>XP_020774501.1 sodium/potassium-transporting ATPase subunit beta-1-like [Boleophthalmus pectinirostris]

MSPNKESDGGWQKFLWNPEKKEFLGRTGGSWAKITLFYIIFYGCLAGIFVGTIQALLLTLSNYKPTYQDRVAPPGLSHTPRSDKSEIAFNMNDPESYKKYINTIDDLLKKYDDDLQTKEKNYENCGVEPLPHKDRGPLLSDQGQKKACRFSRSLLGPCSGETDTNYGFKEGKPCLIVKLNRIVNFIPRAPTNDSMPESIASAFNPNLIPIYCKNKRDEDKEKVKEIKYHGIQGGFPLQYYPYYGKLLHPQYLQPLVAVQFTNLTVGEELRIECKVYGANIYYSEKDRYQGRFDVKITINNS

>XP_020796651.1 sodium/potassium-transporting ATPase subunit beta-3-like [Boleophthalmus pectinirostris]

MSHTNQNQDLNQALNQDLNQNETKPGLNQDQNETKLSVNQQQNKDLNKNQKNQRPKPARAKTQEKIQTRVPPGVQTGAQTRDQTRDQTRDQTHQNQVQDRSQETFCSFIYNPRTGAVLGRTKSSWGRILLFYLVFYSLLSAMFCFTLWMVLLTLDPDTPKYSHLLQNPGLVVRPQVLEISFNRSEPSEYSAYVEQLHRFLHKYNDSVQAANDLCLAGEFTLQDRDQDRDQDRDQDRDQDQDQGPNPAQRRVCQFRRSVLRSCSGLTDPHFGFKDGRPCVILSLNRVVGLKPSGDPYINCTSKVPTPLKLQYFPSDARLDRMFFPFYGKRTHSQYVQPLVAVKLLPSRRDVGVSLSITCRVEGSSIQNQNFRDRNQGRVSFKVRITQ

**Na^+^/K^+^/Cl^-^ COTRANSPORTER (NKCC)**

***Round goby (Neogobius melanostomus)***

>NEME_00014179_Neogobius

LPSERPPYGDADCSAVYNSSEESPVSLGGAAASTIPNASRRGGGDLLAPDAWPKPLGPTP

SQSRFQVDLVAEAAGVAQDRSPSSDTSTVPSSDKASAGGLAEADSGPGGEEAKGRFRVVN

FADPSGASGAASPEAPGDGLQNGDTVMSETSLHSSTGGQHHYHYDTHTNTYYLRTFGHNT

IDAVPNIDFYRHTAAPLGEKLIRPTLSELHDELDKEPFEDGFAADELTPAEEAAAKEAAE

PKGVVKFGWVKGVLVRCMLNIWGVMLFIRMSWIVGQAGIALSCLVVCMATVVTTITGIST

SAIATNGFVRGGGAYYLISRSLGPEFGGSIGLIFAFANAVAVAMYVVGFAETIVELLGGV

DAIMTDEINDIRIIGTITVIVLLGISVAGMEWEAKDPQLAIPRGTLLAILITGIVYLGVA

VSTGSCIVRDASGNINDTIDSLFKSNCTDAACKFDFDFSSCKSSDDGCRYGLQNDFQVMS

TVSGFGPIITAGIFSATLSSALASLVSAPKVFQDNIYPNLGIFAKGYGKNNEPLRGYILT

FGIALAFILIGELNTIAPIISNFFLASYALINFSVFHASLANSPGWRPSFKYYNMWVSLA

GAVLCCVVMFVINWWAALLTNVIVLGLYIYVSHKKPDVNWGSSTQALTYHQALTHTLHLS

GVEDHIKNFRPQCLVMTGYPNSRPALLDLVHSFTKNVGLMICGHVRTGYRRPNFKDIATE

QARYQRWLLKNETKGFYTSVFAEDIRQGAQYLLQAAGLGRLKPNTLVLGFKNDWRDGDMM

NVETYISMIHDAFDVQYGAVILRLEEGLDISHIQGQDELLSSQDKSSGTKDVIVSIDTSK

DSDADSSKPSSKATSLQNSPAIPKDKKSPTMPLNVTDQKLLEASQMFQKKQGKGTVDVWW

LFDDGGLTLLIPYLLTNKKRWKDCKIRVFIGGKINRIDHDRRAMATLLSKFRIDFSDITV

LGDVNIKPKKEQSQNIELFYSITAFEEMIEPYRLKEDDMEQDAAERLKNSEPWRITDNEL

ELYRAKTNRQIRLNELLKEHSSTANLIVMSLPLARKGAVSSALYMAWLEALSKDLPPLLL

VRGNHQSVLTFYS

>NEME_00016555_Neogobius

MDKSNGGEHVNAAYDRTLDEHPNYEETHRTVRPSVVSAFGHDTLDRVPNINFYRNEGSVS

GHRQVRPSLQELHNVFQMKQNGGISVPDTVQDDGESSAGTPCEDLESIDQTKALSGSAGY

EGTGLGIVVIALSCVVTTITALSVSAICTNGVVRGGGAYYLISRSLGPEFGGSIGLIFAF

ANAVAVAMYVVGFAETLVELLEDNNAQMVDTLNDIRIIGCITVVLLLGISVAGMEWEAKA

QIVLLVILLLAIVNVFVGTVIPATDSKKSQGFFNYNSKIFMENFTPDFRGSESFFTVFSI

FFPAATGILAGANISGDLRDAQAAIPKGTLLAILITGVTYLAVALVVSATVVRDATGNLF

DNITAGSMNCSGSSAVTACELGFNFTSCSDPATPCKYGLMNNFQVMTMVSGFGPLIIAGT

FSATLSSALASLVSAPKIFQALCKDNIYKILYFFAKGHGKNNEPLRGYVLTFLISVAFIL

VGNLNVIAPIISNFFLASYALINFSCFHASWAKSPGWRPAYKFYNMWLSLLGAILCCGVM

FVINWWAALITYAIEILLYIYVTVKKPDVNWGSSTQAVSFVSAVSNTLSLSGVADHVKNF

RPQILALTASVRTRPALLDLAHSFSKNYGLCVACEVFVGPRAEALEEMNANMEKNQMYLR

KAKRKAFYTAVVYEDFRNGTMSLLQALGLGRMKPNILMMGFKSNWRTAGAEAVQNYVGVL

HDAFDFEYGTLVLRMNEGLDVAHIVEAEVGSNLVLRQPPDEMFKAAKEAKEEMMPNGGKA

KGLFRKSRNSSQQVLTTRVSVCGPPQVVRMNENLVEASNLFKKKQPKGTIDVWWLFDDGG

LTLLLPYILTTRKKWKDSKLRIFIAGQPGRTEQDKQEMTSLLHKFRINCSDIIVIDDLHI

SPRSESLKKVDDMIAPFRLNENSKDSAQANALRKEFPWKITDEELSKFEEKTNLQVRLNE

LLQEHSKAANLIIVSMPIARKESVSDFLYMAWLDSLTKDLPPTLLIRGNHKSVLTFYSAY

DYIVGPMDKAIVKTDIQIAVPPGHYGRVAPRSGLAVKHFIDVGAGVVDEDYRGNVGVVLF

NFNKEPFEVKKGDRVAQLVCERICYPDLQELETLDETERGAGGFGSTGRN

>NEME_00022893_Neogobius

LGAPPDVFLEPDPLKSDSVSLHSTGTGHTHLSGVSHISDSHSNTYYMRTFGHNTIDAVPN

IDFYRQTAAPLGEKLTRPSLSELHDELDKELFEEGLANGDEPSAVEEAATALAAKEAKGG

TIKFGWVKGVLIRCMLNIWGVMLFIRMSWIVGQAGIGLTIVIILMATIVTTITGLSTSAI

ATNGFVRGGGAYYLISRSLGPEFGGSIGLIFAFANAVAVAMYVVGFAETVVEMLNAQIVL

LGILLAAIGNYFIGTFMPSTNKEPKGVFGYNTAIFLENLGPDFREDETFFSVFAIFFPAA

TGILAGANISGDLTVSNSRRNTSGDLGDPQTAIPKGTLLAILITGLTYVAVTISTGSCIV

RDATGDQNDTISDTVNCTDAACTLGYDFSICKEGGCQYGLMNDFQVMSLVSVFGPLISAG

IFSATLSSALASLVSAPKVFQALCKDNIYPGLSIFAKGYGKNNEPLRGYVLTFAIGLAFI

LIAELNVIAPIISNFFLASYALINFSVFHASLANSPGWRPSFKYYNMWVSLAGAVLCCVV

MFVINWWAALVTLLIVLALYIYVGYRKPDVNWGSSTQALIYNQALTHCLDLTMVPDHVKN

FRYLQLCPLQTFGSCRISRLTFLRLCYRPQCLVMAGYPNARPALLNLVNSFTKNIGLMVC

GHVKAVRRRPNFRELSAEHDRCQRWLQKKKIRAFYTPVFSDNLRYGTQLLLQAMGLGRVR

PNTLVMGFKHSWSEADMKDTELYVNTIQRAVASGQT

>NEME_00027576_Neogobius

MGIEDANTEVEVDSLPAPDEEAQSEKPPVRFGWLVGVMIRCMLNIWGVILFLRLSWLTSQ

AGIVLMWLIILMSTIVTTITALSVSAIATNGRVISGGTYFMISRSLGPEIGAPIGLVFSF

ANALACALNTVGFAEVVRDLLKDFDAVMVDDINDVRIVGVVTVTLLLLISLAGMEWEASA

QMVFFAALMLSFSNYLVGTFLPMPKEKQALGVFSYQVDILVANLTPDWRGPEGNFFRTFA

VFFPSATGILSGVNICGDLRDPMSAIPKGTLFAIFFTTLSYLGIALTSAACTVRDASGNI

TDFIGNNTEGCIGLGCRYGWNFTECIKSQSCEYGLANNINVLGQLSGFYYFITLGVFAAS

LSSALGFLVSAPKIFQCLCKDKIYPYIFFFAKDYGKNDEPLRAYVLCFIIAVAFILIAEL

NTIAALISNFFLCSYGLINFSCFHASITNSPGWRPAFKYYSKWTALFGAVISFVLMFLFT

WWAALVTFSIIIFLFGYVQYNKPQVNWGSSVQASTYNMALSYSVSLSGVEDHVKNFRPQC

LVLTGPPNKRPALVDFVGSFTKHISLMICGDILTEDRKNRAENATDWLVKWLNQRKVRSF

YTPFISDSLRVGVQYLLQASGLGKLKPNTLVMGFKANWRESSPESIEDYINTIYDTFDSN

YCLCILRMMDGQDVLDQFDFEVNQGFESDEATEIEDQQSPERDEANDVSDKGSSDQVQTV

FQTNQGKKNIDVYWIADDGGLTLLVPYLLTRRKKWGRCKVRVFIVGDEQNMEDRRNEMLA

LLKRFRLDFNDVTVMTDSENCPHAKRLHDEQQDGVGLQELRQTYPWKISDKEFEAFKPKS

ERKVRLNEIIRKNSQHAALVLVNLPVPQSDCPSVLYMAWLDTLTSGLHCPVVLIRGNQQN

VLTFYCQ

>NEME_00007878_Neogobius

MEFPPVLREGIHLSAFRPSVGANGNKAASPLGDEKFYDSYGGRCSVTSSLSDTAGSGYET

LDAPPHYDFYANTEVWGRRKRFRPSLYQLYANPQDDSSPNLYEETNAGHLQDHGDSTDEE

DEDEQKEPPPEPTRFGWIHGVMIRCMLNIWGDDLGDHSAVLYNHRNHRSLHLSHRDQWKS

QRRLVIHTYSKIHRGTYFLISRSLGPELGGSIGLIFAFANAVAVAMHTVGFAETVEHGAV

MVDQLNDKRIIGILTVTFLLAISMAGMAWESKAQVVFFILIMVSFASYIVGTIMPASPQK

QAKGFFGYKADIFAENFVPGWRGKEGSFFGMFSIFFPSATGILSGANISGDLKNPAVAIP

RGTLLAIFFTTLSYLIISATIGSCVVRDASGVLNDTLSTSSNPESCVGLACTYGWDFSDM

VSAVAPLITAGIFGATLSSALACLVSAPKVFQCLCKDRLYPLIGFFGKGYGKNDEALRAY

LLTYLIAAAFIVIAELNTIAPIISNFFLCSYTLINFSCFHASITNSPGWRPSFRFYNKWL

SLLGAVCCVVIMFLLTWWAALIAFGIVFLLLGYTLYKKPAVNWGSSVQASSYNIALNQCV

GLNRVEDHVKNYRPQCLVLTGPPSSRPALVDLVSCFTKSLSLMMCANVVESGCSPAVLET

VTSEKHVAWLNQRNVKSFYRGVISSDLRSGVNMLMQGAGLGRIKPNVLLLGFKKSWLSDS

PQAAHDYIGILHDAFELQYGVCVLRMMEGLDVSHPHQSHEPQTTTVFQKKQAKKTIDVYW

LCDDGVCLILLLPYLLTRRKRFSKCKVRVFVGGQPDNKEEQRLELLALIKKFRLGFHDVE

VLPDIYQSPQPMNVQLFENMVAQYRSDKNPKQDSGCGPTRVPDGPWVITEQDWERNRAKT

LRQIRLHEVLQDYSKEAALILVTMPVGRRGVCPTSLYLAWLDFISHGLRPPVLLVRGNQQ

NVLTLYCQ

>NEME_00027570_Neogobius

MIHAQSQACILIERRGSLQELRRGSFYSIMDPTPQLEHYTNSLPRQQVKSRPSLETLRKA

YENGEMGIEDANTDVEVDSLPAPDEEAQSEKPPVRFGWLVGVMVSRMLMWLIILMSTVIT

TITALSVSAIATNGRVVSGGTYFMISRSLGPEIGAPIGLVFSFANALACALNTVGFAEVV

RDLLKDFDSVIVDDINDVRIVGVVTITILLFITLGGMEWEASAQMVFFAALMLSFSNYLV

GTFLPMPKEKQALGIFNYQGDILVANLTPDWRGPEGNFFRTFAVFFPAATGILSGVNICG

DLRDPTSAIPKGTLLAIFFTTLSYLGIALTSAACTVRDASGNITDFIGNNTEGCIGLGCR

YGWNFTECIKSQSCGYGLANNINCLCKDKIYPYIFFFAKGYGKNDEPLRAYVLCFIIAVA

FIVIAELNTIAALISNFFLCSYGLINFSCFHASITNSPGWRPAFKYYSKWTSLFGATISF

LLMFLFTWWAALVTFSIIIFLFGYVQYNKPQVNWGSSVQAGTYNMALSYSVSLTGVEDHV

KNFRPQCLVLTGPPNKRPALVDFEEDRKNRAENATDWLVKWLNQRKVRSFYTPFISDSLR

VGVQYLLQASGLGKLKPNTLVMGFKANWRESSPESIEDYINTIYDTFDSNYCLCILRMME

GLDVLDQFDFEVNQGFESDEATEIEDQQSPERDEANDVSDQGSSDQVQTVFQTNQGKKNI

DVYWIADDGGLTLLVPYLLTRRKKWGRCKVRVFIVGDEENMEDRKNEMLALLKRFRLDFN

DVTVMTDSENRPHAKRNLPVPQSDCPSVLYMAWLDTLTSGLHCPVVLIRGNQQNVLTFYC

Q

>NEME_00003229_Neogobius

MNVQVVPEHKLSSVAPLKQCNLEKKEVELILVKDQNGVQYTSSSIMPTSGRGSSEDERET

WGKKIDFLLSVIGFAVDLANVWRFPYLCYKNGGGVGYTVILIALYVGFYYNVIIAWSLHY

LFSSMTRDLPWLNCGNSWNSPNCTDPIAINGSVLGNGTSYAKYKITPAAEYYERGVLHLY

ESRGINDLGPPRWDLSLCLIAVVFLLYFSLWKGVKSSGKVVYITATMPYVVLFVLLIRGV

TLPGSMDGIRAYLHIDFKRLNNLEVWIDAATQIFYSLGAGFGVLIAFASYNKFDNNCYRD

ALLTSTVNCVTSFFSGFAIFSVLGYMAFKHGVRIEDVATEGAGLVFIIYPEAISTLPGST

FWAIVFFIMLLTLGIDSSMGGMEAVITGLSDDFKFLKRNRKAFTFATAFGTFLVALCCVT

NGGIYVLTLLDKYAAGTSILFAVLIEAIGVSWFYVVPVKDRGQSSYDSLEGVNWVDYRDT

GQDCADNQDTVGSDGLGNQTEDSPFLNSAESGGKKNDFYDKNLALFEEELDIRPKVSSLL

SRLVSYTNITQGAKEHEEEEISQRGNAAKTMLTAISMSAIATNGVVPAGGSYFMISRSLG

PEFGGAVGLCFYLGNTFAAAMYILGAIEIFLKYIVPQAAIFHAVDHHGTDSAMLNNMRVY

GSICLILMVVVVFVGVKICMLGNRTLVNSKFDVCAKTVMQGNMTVPSQLWENFCTPGNLS

SSQCDEYFLLNNVTKIQAIPGMSSGIIRENMWSNYRYKGQLLEKPAVQSVDAHGSLEKLS

LYLTADITTSFTLLLGIFFPSATGIMAGSNRSGDLKDAQKSIPVGTILAIATTSFVFHNN

LVVGTLAWPSPWVIVIGSFFSTIGAGLQTVTGAPRLLQAIAKDNIIPFLRVFGHGKKNGE

PTWALLLTGLIAELGILIASLDMVAPILSMFFLMCYLFVNLACAVQTLLRTPNWRPRFKY

YHWTLSFLGMSICLALMFISTWYYAIVAIGIAGMIYKYIEYQGAEKEWGDGIRGLSLSAA

RYALLRLEAGPPHTKNWRPQLLVLLKLDEDLHVKYPRMLTFASQLKAGKGLTIVGTVIQG

NFLDSFGELQAAEQAVKNMMEIERVKGFCQVVVANKVRDGIVHLIQSCGLGGMKHNTVVM

GWPYGWRQSEDPRAWKTFINNVRCVTAAHLALMVPKNVSFYSSNHERFTDGHIDVWWIVH

DGGMLMLLPFC

>NEME_00004087_Neogobius"

MRVYGTILLTSMATVVFVGVKYVNKLALVFLACVILSIVAVYAGVIKTAIDPPIFPVCLL

GNRTLVWKTFDVCAKTLETSNGTITTQLWQMFCDSPYLNATCDKYFAANNVTEIQGIPGI

TSGIMTGIMAGSNRSGDLRDAQKSIPVGTICAITTTTIVYMSSVILFGACIDGVVLRDKF

GEGVHGNLVIGTLAWPSPWVIVIGSFFSTCGAGLQSLTGAPRLLQAIAKDGIVPALRIFG

HGKANGEPTWSLLLTACICEIGILIASLDAVAPILSMFFLMCYMFVNLACALQTLLRTPN

WRPRFKFYHWTLSFLGMSLCLTLMFLCSWYYAIVAMVIAGSIYKYIEFAGPQLLVLVSTD

AEQNIEQPRLLSLTNQLKAGKGLTIVGTALEGTYLANHDRAQRAEQALRKLMETEKVKGF

CQVTVSSNLRDATSHLLQASGLGGLKHNAVLVSWPCNWKQGDEHQHWRNFIGEACQRNTA

AHLALLVPKNISAYPSNGERFTEGHIDVWWIVWRKCKMRIFTVAQMDDNSIQMKKDLTTF

LYHLRIDAMVEVVEMQDSDISAYTYEKTLVMEQRSQMLKKINLTKTEREREIQSITDSSR

GSIRRKNPAAVTTQLSVTEESPAASKEEKPEEECCAFSKCITWYSTAKADSSLAQVKVDF

SPCVLPCTGTAHPRQHHSWQPCYPRHPTHPEEVQSPGTQVQMTWTEKCDGEAAKPPGACT

PEGLKDIFNMKPNQFNVRRMHTALRLNEVVVKKSSEAKLVLLNMPGPPKNRTGDENYMEF

LEVLTEGLNRVLLVRGGGREVITIYS

>NEME_00001099_Neogobius

MQNNMMDSEEGEGGSISQDGIPKESSPFINNDVEKSQHYDGKNMALFEEEMDTSPMVSSL

LSSLANYSNLPTGSKEHEEAENTEEVSRSTIKPVKAPQLGTLMGVYLPCIQNIFGVILFL

RMTWMVGIGGVMGSFIIVFMCCSTTMLTAISMSAIATNGVVPAGGSYYMISRSLGPEFGG

AVGICFYLGTTFAGAMYILGCIEILLIYIVPQAAIFKLEGLEGAEAEAALLNNMRVYGTI

VLSFMALVVFVGVKYVNKLALVFLACVILSILAVYAGVINTAIEPPLFPVCLLGNRTLLS

KAYDVCAKVIEIDNETITTKLWLSFCDSDTLNATCDEYFTNNNVTEIQGIPGVTSGILSE

NLFSKYLDKGMMLEKEGVLSEQDRDNTVTSSGRYVLADITSFFTLLVGIYFPSVTGIMAG

SNRSGDLKDAQKSIPIGTIAAITTTSIVCLQSLTGAPRLLQAISRDGIIPFLRVFGHGKA

NGEPTWALLLTACICEIGILIASLDAVAPILSMFFLMCYMFVNLACALQTLLRTPNWRPR

FKFYHWPQLLVLVSVDSDQNVEQPRLLSLTNQLKAGKGLTIVGTSVQGTFLDNYTEAQTA

EQALRRLMEIEKVKGFSQVVISSSLRDGTSHLVQVGGLGGLKHNAVMVSWPHNWKQPQYN

QQFRNFIEVVRETTVASLALLVPKNISSYPSNGERFTEGHIDVWWIVHDGGMLMLLPFLL

RQHKVWRKCKMRIFTVAQMDDNSIQMKKDLIMFLYHLRIDAIVEVVEMIQSITDSSRGSI

RRKGKTAFCSQRSTDESATVIEKPEEESSTSPNTKNSTNTPTSLTSHSSPAGDARTWTDG

KALGKGLSQAIEGGKDHFNMKPNQTDVRRMHTALKLNEVISKKSKEAKLVLLNMPGPPRN

RVGDENCILLCTGNMEFLEVLTEGLNRVLLVRGGGREVITIYS

***zebrafish (Danio rerio)***

>tr|A0A0G2KGS0|A0A0G2KGS0_DANRE_Danio

MSASPPISAGDYLSAPEPDALKPAGPTPSQSRFQVDLVTESAGDGETTVGFDSSPPEYVA

EPPPDGLRDSVSGGEEAKGRFRVVNFAASSPDAAPAETAQNGDTVMSEGSLHSSTGGQQH

HHYDTHTNTYYLRTFGHNTIDAVPKIDFYRQTAAPLGEKLIRPTLSELHDELDKEPFEDG

FANGEELTPAEESAAKDVSESKGVVKFGWIKGVLVRCMLNIWGVMLFIRMTWIVGQAGIA

YSCIIVIMATVVTTITGCSTSAIATNGFVRGGGAYYLISRSLGPEFGGSIGLIFAFANAV

AVAMYVVGFAETVVELLMDSGLLMIDQTNDIRVIGTITVILLLGISVAGMEWEAKAQIFL

LVILITAIFNYFIGSFIAVDSKKKFGFFSYDAGILAENFGPDFRGQTFFSVFSIFFPAAT

GILAGANISGDLADPQMAIPKGTLLAILITGLVYVGVAISAGACIVRDATGIESNFTLIS

NCTDAACKYGYDFSSCRPTVEGEVSSCKFGLHNDFQVMSVVSGFSPLISAGIFSATLSSA

LASLVSAPKVFQALCKDNIYPGIAIFGKGYGKNNEPLRGYFLTFGIALAFILIAELNVIA

PIISNFFLASYALINFSVFHASLANSPGWRPSFKYYNMWASLAGAILCCVVMFIINWWAA

LLTNVIVLSLYIYVSYKKPDVNWGSSTQALTYHQALTHSLQLCGVADHIKTFRPQCLVMT

GAPNSRPAILHLVHAFTKNVGLMLCGHVRISSRRPNFKELNSDMLRYQRWLLNNNSKAFY

TCVVAEDLRQGTQYMLQAAGLGRLRPNTLVIGFKNDWRTGDIKEVETYINLIHDAFDFQY

GVVILRLREGLDISHIQGQDDSSGMKDVVVSVDISKDSDGDSSKPSSKATSVQNSPAVQK

DKKSPTVPLNVADQRLLDASQQFQQKQGKGTVDVWWLFDDGGLTLLIPYLIANKKKWKDC

KIRVFIGGKINRIDHDRRAMATLLSKFRIDFSDITVLGDINTKPKSEGLTEFAEMIEPYK

LREDDMEQEAAEKLKSEEPWRITDNELELYKAKGNRQIRLNELLKEHSSTANLIVMSMPL

ARKGAVSSALYMAWLDTLSKDLPPILLVRGNHQSVLTFYS

>tr|Q6IQW8|Q6IQW8_DANRE_Danio

MSASPPISAGDYLSAPEPDALKPAGPTPSQSRFQVDLVTESAGDGETTVGFDSSPPEYVA

EPPPDGLRDSVSGGEKAKGRFRVVNFAASSPDAAPAETAQNGDTVMSEGSLHSSTGGQQH

HHYDTHTNTYYLRTFGHNTIDAVPKIDFYRQTAAPLGEKLIRPTLSELHDELDKEPFEDG

FANGEELTPAEESAAKDVSESKGVVKFGWIKGVLVRCMLNIWGVMLFIRMTWIVGQAGIA

YSCIIVIMATVVTTITGCSTSAIATNGFVRGGGAYYLISRSLGPEFGGSIGLIFAFANAV

AVAMYVVGFAETVVELLMDSGLLMIDQTNDIRVIGTITVILLLGISVAGMEWEAKAQIFL

LVILITAIFNYFIGSFIAVDSKKKFGFFSYDAGILAENFGPDFRGQTFFSVFSIFFPAAT

GILAGANISGDLADPQMAIPKGTLLAILITGLVYVGVAISAGACIVRDATGIESNFTLIS

NCTDAACKYGYDFSSCRPTVEGEVSSCKFGLHNDFQVMSVVSGFSPLISAGIFSATLSSA

LASLVSAPKVFQALCKDNIYPGIAIFGKGYGKNNEPLRGYFLTFGIALAFILIAELNVIA

PIISNFFLASYALINFSVFHASLANSPGWRPSFKYYNMWASLAGAILCCVVMFIINWWAA

LLTNVIVLSLYIYVSYKKPDVNWGSSTQALTYHQALTHSLQLCGVADHIKTFRPQCLVMT

GAPNSRPAILHLVHAFTKNVGLMLCGHVRISSRRPNFKELNSDMLRYQRWLLNNNSKAFY

TCVVAEDLRQGTQYMLQAAGLGRLRPNTLVIGFKNDWRIGDIKEVETYINLIHDAFDFQY

GVVILRLREGLDISHIQGQDDSSGMKDVVVSVDISKDSDGDSSKPSSKATSVQNSPAVQK

DEDDDGKAHTQPLLKKDKKSPTVPLNVADQRLLDASQQFQQKQGKGTVDVWWLFDDGGLT

LLIPYLIANKKKWKDCKIRVFIGGKINRIDHDRRAMATLLSKFRIDFSDITVLGDINTKP

KSEGLTEFAEMIEPYKLREDDMEQEAAEKLKSEEPWRITDNELELYKAKGNRQIRLNELL

KEHSSTANLIVMSMPLARKGAVSSALYMAWLDTLSKDLPPILLVRGNHQSVLTFYS

>tr|C7EA90|C7EA90_DANRE_Danio

MSASPPISAGDYLSAPEPDALKPAGPTPSQSRFQVDLVTESAGDGETTVGFDSSPPEYVA

EPPPDGLRDSVSGGEEAKGRFRVVNFAASSPDAAPAETAQNGDTVMSEGSLHSSTGGQQH

HHYDTHTNTYYLRTFGHNTIDAVPKIDFYRQTAAPLGEKLIRPTLSELHDELDKEPFEDG

FANGEELTPAEESAAKDVSESKGVVKFGWIKGVLVRCMLNIWGVMLFIRMTWIVGQAGIA

YSCIIVIMATVVTTITGCSTSAIATNGFVRGGGAYYLISRSLGPEFGGSIGLIFAFANAV

AVAMYVVGFAETVVELLMDSDLLMIDQTNDIRVIGTITVILLLGISVAGMEWEAKAQIFL

LVILITAIFNYFIGSFIAVDSKKKFGFFSYDAGILAENFGPDFRGQTFFSVFSIFFPAAT

GILAGANISGDLADPQMAIPKGTLLAILITGLVYVGVAISAGACIVRDATGIESNFTLIS

NCTDAACKYGYDFSSCRPTVEGEVSSCKFGLHNDFQVMSVVSGFSPLISAGIFSATLSSA

LASLVSAPKVFQALCKDNIYPGIAIFGKGYGKNNEPLRGYFLTFGIALAFILIAELNVIA

PIISNFFLASYALINFSVFHASLANSPGWRPSFKYYNMWASLAGAILCCVVMFIINWWAA

LLTNVIVLSLYIYVSYKKPDVNWGSSTQALTYHQALTHSLQLCGVADHIKTFRPQCLVMT

GAPNSRPAILHLVHAFTKNVGLMLCGHVRISSRRPNFKELNSDMLRYQRWLLNNNSKAFY

TCVVAEDLRQGTQYMLQAAGLGRLRPNTLVIGFKNDWRIGDIKEVETYINLIHDAFDFQY

GVVILRLREGLDISHIQGQDDSSGMKDVVVSVDISKDSDGDSSKPSSKATSVQNSPAVQK

DKKSPTVPLNVADQRLLDASQQFQQKQGKGTVDVWWLFDDGGLTLLIPYLIANKKKWKDC

KIRVFIGGKINRIDHDRRAMATLLSKFRIDFSDITVLGDINTKPKSEGLTEFAEMIEPYK

LREDDMEQEAAEKLKSEEPWRITDNELELYKAKGNRQIRLNELLKEHSSTANLIVMSMPL

ARKGAVSSALYMAWLDTLSKDLPPILLVRGNHQSVLTFYS

>tr|A0A0G2KTI4|A0A0G2KTI4_DANRE_Danio

MSASPPISAGDYLSAPEPDALKPAGPTPSQSRFQVDLVTESAGDGETTVGFDSSPPEYVA

EPPPDGLRDSVSGGEEAKGRFRVVNFAASSPDAAPAETAQNGDTVMSEGSLHSSTGGQQH

HHYDTHTNTYYLRTFGHNTIDAVPKIDFYRQTAAPLGEKLIRPTLSELHDELDKEPFEDG

FANGEELTPAEESAAKDVSESKGVVKFGWIKGVLVRCMLNIWGVMLFIRMTWIVGQAGIA

YSCIIVIMATVVTTITGCSTSAIATNGFVRGGGAYYLISRSLGPEFGGSIGLIFAFANAV

AVAMYVVGFAETVVELLMDSGLLMIDQTNDIRVIGTITVILLLGISVAGMEWEAKAQIFL

LVILITAIFNYFIGSFIAVDSKKKFGFFSYDAGILAENFGPDFRGQTFFSVFSIFFPAAT

GILAGANISGDLADPQMAIPKGTLLAILITGLVYVGVAISAGACIVRDATGIESNFTLIS

NCTDAACKYGYDFSSCRPTVEGEVSSCKFGLHNDFQVMSVVSGFSPLISAGIFSATLSSA

LASLVSAPKVFQALCKDNIYPGIAIFGKGYGKNNEPLRGYFLTFGIALAFILIAELNVIA

PIISNFFLASYALINFSVFHASLANSPGWRPSFKYYNMWASLAGAILCCVVMFIINWWAA

LLTNVIVLSLYIYVSYKKPDVNWGSSTQALTYHQALTHSLQLCGVADHIKTFRPQCLVMT

GAPNSRPAILHLVHAFTKNVGLMLCGHVRISSRRPNFKELNSDMLRYQRWLLNNNSKAFY

TCVVAEDLRQGTQYMLQAAGLGRLRPNTLVIGFKNDWRTGDIKEVETYINLIHDAFDFQY

GVVILRLREGLDISHIQGQDDSSGMKDVVVSVDISKDSDGDSSKPSSKATSVQNSPAVQK

DEDDDGKAHTQPLLKKDKKSPTVPLNVADQRLLDASQQFQQKQGKGTVDVWWLFDDGGLT

LLIPYLIANKKKWKDCKIRVFIGGKINRIDHDRRAMATLLSKFRIDFSDITVLGDINTKP

KSEGLTEFAEMIEPYKLREDDMEQEAAEKLKSEEPWRITDNELELYKAKGNRQIRLNELL

KEHSSTANLIVMSMPLARKGAVSSALYMAWLDTLSKDLPPILLVRGNHQSVLTFYS

***three spine stickleback (Gasterosteus aculeatus)***

>tr|G3Q307|G3Q307_GASAC_Gasterosteus

MSAPTSAPSAAAGCDSLAPDAGPKPLGPTPSQSRFQVDIVAEAAGATGDKSPSSDASTTA

PSSAGACGGEEAKGRFRVVNFADPSEAGSAASPEAPGDGLQNGDTVMSETSLHSSTGGPH

HYHYDTHTHTYYMRTFGHNTIDAVPNIDFYRQTAAPLGEKLIRPTLSELHDELDKEPFED

GFANGDELTPPEEAAAKEAADNKGVVKFGWVKGVLVRCMLNIWGVMLFIRMSWIVGQAGI

ALACLIVAMATVVTTITGLSTSAIATNGFVRGGGAYYLISRSLGPEFGGSIGLIFAFANA

VAVAMYVVGFAETVAELLAGVDANMTDELNDIRIIGTITVILLLGISLAGMEWEAKAQIF

LLIVLITAIINYFIGSFIPSASKKPLGYFGYDGAIMWENMGPDFREETFFSVFAIFFPAA

TGILAGANISGDLSDPQLAIPRGTMLAILITGIVYLGVAVSTGSCIVRDATGNLNDTVSS

QFMMNCTDAACKLGYDFSSCKSDKTACRYGLQNDFQVMSMVSGFGPVITAGIFSATLSSA

LASLVSAPKVFQALCKDNIYPGLGIFAKGYGKNNEPLRGYILTFVIALAFILIAELNIIA

PIISNFFLASYALINFSVFHASLANSPGWRPSFKYYNMWVSLVGAILCCVVMFVINWWAA

LMTNVIVLGLFIYVSYKKPDVNWGSSTQALTYHQALTHTLHLSGVEDHIKNFRPQCLVMT

GYPNSRPALLDLVHSFTKNVGLMMCGHIRTGYRRPNFKELATDQARYQRWLLKNETKAFY

TPVFAEDMRQGAQYLLQAAGLGRLKPNTLVLGFKNDWRDGDMMNVETYISMIHDSFDFQF

GAVILRLKEGLDVSHIQGPDDLLSSQEKSSGMKDVIVSIDTSKDSDADSSKPSSKATSIQ

NSPAMQKDEDEDGKATTQPLLKKDKKSPSVPLNVSDQRLLEASQQFLKKQGKGTVDVWWL

FDDGGLTLLIPYLLTNKKRWKDCKIRVFIGGKINRIDHDRRAMATLLSKFRIDFSDINVL

GDINTKPKKEHVSAFEEMIEPYRLKEDDMEQEAAERLKDSEPWRITDNELELYRAKTNRQ

IRLNELLKEHSSTANLIVMSLPLARKGAVSSALYMAWLEVLSKELPPILLVRGNHQSVLT

FYS

>tr|G3PPB7|G3PPB7_GASAC_Gasterosteus

MRTFGHNTIDAVPNIDFYRHTAATLGEKLTRPSLSELHDELDKDPFEDGLANGEEPSVAE

EAAAALRAKEAKGGTIKFGWVKGVLIRCMLNIWGVMLFIRMSWIVGQAGIGLTIAIILMA

TVVTTITGLSTSAIATNGFVRGGGAYYLISRSLGPEFGGSIGLIFAFANAVAVAMYVVGF

AETVVEMLSDVDALMTDELNDIRIVGTLTVILLLGISVAGMEWEAKAQIVLLVILLAAIA

NYFIGSFISTESKEPKGFFGYQSAIFLENLGPDFRDEETFFSMFAIFFPAATGILAGANI

SGDLTDPQSAIPKGTLLAILITGITYVAITISAGSCIVRDATGDHNDTVSDTVNCTDAAC

TLGYDFSICKEGGCQYGLINDFQVLGLVSGFGPLISAGIFSATLSSALASLVSAPKVFQA

LCKDNIYPGLGVFAKGYGKNNEPLRGYVLTFCIGLAFILIAAELNIIAPIISNFFLASYA

LINFSVFHASLANSPGWRPSFKYFNKWVSLAGAILCCVVMFVINWWAALVTLLIVLALYI

YVSYKKPDVNWGSSTQALIYNQALTHCMNLTGVDHVKNFRPQCLVLAGYPNSRPALLQLV

NSFTKNVGLMVCAHVRMVTRRPNLKELSQEHGRCQRWLNKKRIKAFYTHVFSDNLRHGAQ

FLMQAVGLGRLKPNTLVMGFKNNWSDGDMREVEIYINTIHDAFDLQYGVVILRLRDGLDV

SHIQGQGAVFFSPPPPPCPSGRRPPHVLALCVPPSLQTRSSSPRRSPPAAFKKKQGKGTI

DVWWLFDDAGLTLLIPFLLTNGSKWGDCRIRVFIGGKINRIRATTAEPLSLTLKLPAMLH

GLRAAVMMSLKWRCGRSKLSFRELVEPYRLREDDMEQEAAERLKAQEPWRITDNELELYK

AKNNRQIRLNELLKEHSAAAKLIVMSMPLARKGTVSSALYMCWLETLSRDVPPLLLVKAN

SESVADTVLR

***tilapia (Oreochromis niloticus)***

>tr|I3K8E1|I3K8E1_ORENI_Oreochromis

MSGQKPSLRSDDSAQSRFQVDVVMEASSAAPGPAAPSAPASTDGSRPPEESKGRFRVVGF

IDPGAMDIGLPPDVSHEPDTAKSDSVSLHSTGTGHTHISDSHSNTYYMRTFGHNTIDAVP

NIDFYRQTAAPFGEKLTRPSLSELHDELDKEPFEDGLANGEEPSAAEEAAALAAKEAKGG

TVKFGWVKGVLIRCMLNIWGVMLFIRMSWIVGQAGIGLTIAIILMATVVTTITGLSTSAI

ATNGFVRGGGAYYLISRSLGPEFGGSIGLIFAFANAVAVAMYVVGFAETVVELLNDVDAL

MTDELNDIRIVGTLTIILLLGISVAGMEWEAKAQIVLLVILLAAIVNFFFGSFMPSESKE

PKGFFGYHTAILLENFGPEFRDGETFFSVFAIFFPAATGILAGANISGDLTDPQSAIPKG

TLLAILITGLTYVAVAISAGSCIVRDATGDQNDTVSPTVNCTDAACTLGYDFSICKEGGC

QFGLMNDFQVMSLVSAFGPLITAGIFSATLSSALASLVSAPKVFQALCKDNIYPGLGMFA

KGYGRNNEPLRGYVLTFCIGLAFILIADLNIIAPIISNFFLASYALINFSVFHASLASSP

GWRPSFKYYNMWVSLVGAILCCVVMFVINWWAALVTLLIVLALYIYVSYKKPDVNWGSST

QALIYNQALTHCLNLTGVEDHVKNFRPQCLVLAGYPNARPALLQLVNSFTKNVSLMVCSH

VRTVSRRSNFRELYQDYARCQRYLNKKRIKAFYAPVFSDNLRHGAQLLLQAVGLGRLKPN

TLVMGFKNNWSDGNMRDVENYINTIHDAFDLLFGVVILRLQEGLDISHIQGQDELLSSHE

KPPVITKDVLISVIPAKDSDSDSCPSKTTSNQSSPLITRETKSPLSLTDQRLLESSQQFK

KKQGKGTIDVWWLFDDGGLTLLIPYLLTNRSKWGDCRIRVFIGGKINRIDHDRRAMATLL

SRFRIDFSDINVLGDINTKPKKHNKLTFKELIEPYRLKEDDMEQEAAERLKAQEPWRITD

NELELYKAKTNRQIRLNELLKEHSSTAKLIVMSMPLARKGTVSSALYMCWLETLSKGLPP

LLLVRGNHQSVLTFYS

>tr|I3JZF4|I3JZF4_ORENI_Oreochromis

MSAPSSASSAPAENSATEHDLLAPDAGQKPPGPTPSQSRFQVDLVAEAAGAADDKTPSSD

ASAAAPEAPSSDPAVSGEEAKGRFRVVNFADPTGEGSVASPEAAPTEGMQNGDTVMSETS

LHSSTGGQHHYHYDTHTNTYYLRTFGHNTIDAVPNIDFYRQTAAPLGEKLTRPTLSELHD

ELDKEPFEDGFANGDELTPAEEAAAKEAAESKGVVKFGWIKGVLVRCMLNIWGVMLFIRM

SWIVGQAGIALACVIVAMATVVTTITGLSTSAIATNGFVKGGGAYYLISRSLGPEFGGSI

GLIFAFANAVAVAMYVVGFAETVVELLVGIDAVMTDEINDIRIIGTITIIILLGISVAGM

EWEAKAQIFLLVVLITAIFNYFIGSFIPVKSKEAKGFMGYDASIMWENIGPDFRGETFFS

VFAIFFPAATGILAGANISGDLADPQMAIPKGTLLAILITGIVYLGVAVSTGSCILRDAS

GNVNDTISSQFMANCSTAACKFGYDFSTCKNEDTCRYGLHRDFQVMSLVSGFGPIITAGI

FSATLSSALASLVSAPKVFQALCKDNIYPGLQMFAKGYGKNNEPLRGYILTFGIALAFIL

IAELNTIAPIISNFFLASYALINFSVFHASLANSPGWRPSFKYYNMWVSLAGAILCCGVM

FVINWAAALLTNVIVMALYIYVSHKKPDVNWGSSTQALTYHQALTHTLHLSGVEDHVKNF

RPQCLVMTGYPNSRPALLDLVHSFTKNVGLMICGHIRTGYRRPNFKELATDQARYQRWLL

KNETKAFYTPVFAEDLKQGSQYLLQAAGLGRLKPNTLVLGFKNDWRDGDMMNVETYISMI

HDAFDFQFGAVILRLKEGLDVSHIQGQDESLSSQEKPSGMKDVIVSIDTSKDSDADSSKP

SSKATSLQNSPAVQKDDEDDGKATTQPLLKKDKRSPTVPLNVSDQRLLEASQQFQKKQGK

GTVDVWWLFDDGGLTLLIPYLLTNKKRWKECKIRVFIGGKINRIDHDRRAMATLLSKFRI

DFSDITVLGDINTKPKKEHMAAFEEMIEPYRLKEDDMEQEAAERLKNSEPWRITDNELEL

YRTKTHRQIRLNELLKEHSSTANLIVISLPLARKGAVSSALYMAWLEALSKDLPPILLVR

GNHQSVLTFYS

>tr|I3K8E2|I3K8E2_ORENI_Oreochromis

FPGNRLSLGASLTEMSGQKPSLRSDDSAQSRFQVDVVMEASSAAPGPAAPSAPASTDGSR

PPEESKGRFRVVGFIDPGAMDIGLPPDVSHEPDTAKSDSVSLHSTGTGHTHISDSHSNTY

YMRTFGHNTIDAVPNIDFYRQTAAPFGEKLTRPSLSELHDELDKEPFEDGLANGEEPSAA

EEAAALAAKEAKGGTVKFGWVKGVLIRCMLNIWGVMLFIRMSWIVGQAGIGLTIAIILMA

TVVTTITGLSTSAIATNGFVRGGGAYYLISRSLGPEFGGSIGLIFAFANAVAVAMYVVGF

AETVVELLNDVDALMTDELNDIRIVGTLTIILLLGISVAGMEWEAKAQIVLLVILLAAIV

NFFFGSFMPSESKEPKGFFGYHTAILLENFGPEFRDGETFFSVFAIFFPAATGILAGANI

SGDLTDPQSAIPKGTLLAILITGLTYVAVAISAGSCIVRDATGDQNDTVSPTVNCTDAAC

TLGYDFSICKEGGCQFGLMNDFQVMSLVSAFGPLITAGIFSATLSSALASLVSAPKVFQA

LCKDNIYPGLGMFAKGYGRNNEPLRGYVLTFCIGLAFILIADLNIIAPIISNFFLASYAL

INFSVFHASLASSPGWRPSFKYYNMWVSLVGAILCCVVMFVINWWAALVTLLIVLALYIY

VSYKKPDVNWGSSTQALIYNQALTHCLNLTGVEDHVKNFRPQCLVLAGYPNARPALLQLV

NSFTKNVSLMVCSHVRTVSRRSNFRELYQDYARCQRYLNKKRIKAFYAPVFSDNLRHGAQ

LLLQAVGLGRLKPNTLVMGFKNNWSDGNMRDVENYINTIHDAFDLLFGVVILRLQEGLDI

SHIQGQDELLSSHEKPPVITKDVLISVIPAKDSDSDSCPSKTTSNQSSPLITRVYFNFCF

STFRRQLIISSEQKKTKSPLSLTDQRLLESSQQFKKKQGKGTIDVWWLFDDGGLTLLIPY

LLTNRSKWGDCRIRVFIGGKINRIDHDRRAMATLLSRFRIDFSDINVLGDINTKPKKHNK

LTFKELIEPYRLKEDDMEQEAAERLKAQEPWRITDNELELYKAKTNRQIRLNELLKEHSS

TAKLIVMSMPLARKGTVSSALYMCWLETLSKGLPPLLLVRGNHQSVRVFCS

>tr|B1Q041|B1Q041_OREMO NCC_Oreochromis

MGQFNSKNKGSGPGIHQANPEEGPPSALGQKDSNMEPRKTSIYNTIDTAPQLEYYANTLP

HQEIKIRPSLDVLRKTFEEMEAEKAAADDATADDENEEPQGPQGKPPTRFGWFIAVWMRC

MLNIWGVILFLRLTWITSQAGIVLGLVIIAMAVSVTTTTALSISAIATNGRVKSGGTYFM

ISRTLGPELGASIGLIFSIANALAVALHTVGFSEVVRDLMRSYGTIMVDALNDVRIIGVI

TVTILLFITFGGMDWEAKAQIFFFIVLMASFADYLVGTLIPPSLQKKSQGFFGYNRDIFM

ENLTPSWRGPDGNFFRQFAIFFPACTGILSGANISGDLKDPSTAIPKGTLMAIFCTTLSY

VVIVVTSGASVVRDASGNMTDLMIGNSTDGCLGPACKLGWNFTKCVQSQACSEGLANYSQ

VMGLMSGSYYLIVAGIFAATLSSALGFLVSAPKIFQCLCKDKVYPYIEFFAKGYGKNNEP

LRGYILTYLIAVAIILVAQLNIIAPIISNFFLCSYALINLSCFHASIVNSPGWRPAFKYY

NKWTALYGALASIALMFAFTWWAALITWTVISLLFLYITYIKKPNVNWGSTIQASSYNMA

LSFSVSLTDVKDHVKNFRPQCLVMTGPPQQRPALVDFVGCFTKHVSLMICGNIIMEPEKQ

TQFQDSTDQCVKWLNKRKVCSFYTQFTADSLRDGVRYLMQASGLGKLKPNTLVLGFKSNW

MESSPKSIEDYIHVIYDTFDSNYCLCILRMMDGLDITDHSDFKENQGFEPDEAIETNDHQ

LPEKESANDISENINSDQIKTVFKNDGGKKTIDIYWIADDGGLILLVPYLLTRRKRWRSG

KIRVFILGDEENMEESRDAMIALLKRFRIDVTDVVVMTDAERSPQPKNMTRFLESVAPYR

LYDEQQEGVSVQEQKQNEPWKISDKQLEAFRLKSERKVRLNEIIRRNSQNTTLVLVSLPV

PHSNCPSALYIAWLDALTCGLHCPVVLVRGNQENVLTFYCQ

***mudskipper (Boleophthalmus pectinirostris)***

>XP_020794684.1_Boleophthalmus

MEKYKSNGGEHVNAAYDPTLDEPPNYEEPARAVRPSVVSAFGHDTLDRVPNIDFYRNAGSVSGHRQVRPSLQELHDVFQQKQNGRISVPDTVQDDSEGSDDGTPCEDLESNPPVDPNEGVVKFGWIRGVLVRCMLNIWGVMLFIRLSWVFGQAGWGLGIVVIALSCVVTTITALSMSAICTNGVVRGGGAYYLISRSLGPEFGGSIGLIFAFANAVAVAMYVVGFAETVVELLQDNNAIMVDPINDIRIIGCITVVLLLGISVAGMEWEAKAQIVLLVILLVAIVNVFVGTFIPATDSKKAQGFFNYNIKIFMENFAPDFRGPENFFTVFSIFFPAATGILAGANISGDLRDAQAAIPKGTLLAILITGITYLGVALVVSATVVRDATGNLFDNITAGSMDCAGSSAVAACELGFNFTSCTDTTNTCKYGLMNNFQVMTMVSGFGPLIIAGTFSATLSSALASLVSAPKVFQALCKDNIYKILHFFAKGHGKNNEPIRGYILTFIISVAFILIGNLNTIAPIISNFFLASYALINFSCFHASWAKSPGWRPAYKYYNMWLSLLGAILCCVVMFVINWWAALLTYGIEILLYIYVTVKKPDVNWGSSTQAVTFVSAVGNALSLSGVDDHVKNFRPQILALTASVRTRPALLDLAHSFTKTYGLCVTCEVFVGPRSEALEEMNANMAKNQLYLRKAKRKAFYTAVVSESFREGAVALLQASGLGRMKPNTLMMGFKTNWRTVGAEAVQSYVGLLHDAFDFEYGTVVLRMDEGLDVAHIVEAEDEMLKAAKEQQALEDEMTPNGKAKGLFRKSRNSSQQVLTTRVSVCGPPPPQVAQMNEKLVQASNLFKKKQPKGTIDVWWLFDDGGLTLLLPYILTTRKKWKDSKLRIFIAGQPGRCEQDKIEMKSLLQKFRINCSDITVIDDLHISPRSDSLKKLDEMIAPFRLKENSKDSAQAEALRKQFPWKITDEELSTFEEKTNLQVRLNELLQEHSRSANLIIVSMPIARKGSVSDFLYMAWLDSLTKDLPPTLLIRGNHKSVLTFYS

>XP_020782054.1_Boleophthalmus

MSAPSSTPPSTTEGDLLAPDAGPKPAGPTPSQSRFQVDLVAEAAGATEEKSPSSDTSTVPSSDKASAGGLAETDSGPVGEEAKGRFRVVNFADPSGASGAASPEGDGVQNGDTVMSETSLHSSTGGQHHYHYDTHTNTYYLRTFGHNTIDAVPNIDFYRQTAAPLGEKLIRPTLSELHDELDKEPFEDGFAXXXXXXXXXXAAAEEAAESKGVVKFGWIKGVLVRCMLNIWGVMLFIRMSWIVGQAGIALSCVIVCMATVVTTITGLSTSAIATNGFVRGGGAYYLISRSLGPEFGGSIGLIFAFANAVAVAMYVVGFAETVVELLAGADAIMTDEINDIRIIGTITVILLLGISVAGMEWEAKAQIFLLVVLITAIVNYFIGTFIAVKAKETFGFFGYDGSIMWENMGPDFRGETFFSVFAIFFPAATGILAGANISGDLADPQLAIPRGTLLAILITGIVYLGVAVSTGSCIVRDASGSINDTISSRFMANCTDAACKFGFDFSSCKSSATDCRYGLHHDFQVMSLVSGFGPIITAGIFSATLSSALASLVSAPKVFQALCKDNIYPGLGMFAKGYGKNNEPLRGYILTFGIALAFILIAELNVIAPIISNFFLASYALINFSVFHASLANSPGWRPSFKYYNMWVSLAGAVLCCVVMFVINWWAALLTNVIVLGLYIYVSHKKPDVNWGSSTQALTYQQALTHTLHLSGVEDHVKNFRPQCLVMTGYPNSRPALLDLVQNFTKNVGLMICGHVRTGYRRPNFKDMATEQARYQRWLLKNETKGFYTSVFAEDIRQGAQYLLQAAGLGRLKPNTLVLGFKNDWRDGDMMNVETYISMIHDAFDFQYGAVILRLQEGLDVSHIQGQDELLSSQEKSSGTKDVIVSIDTSKDSDADSSKPSSKTTSLQNSPAIPKDDEDDGKAPTQPLLKKDKKSPTMPLNVVDQRLLEASQMFQRKQGKGTVDVWWLFDDGGLTLLIPYLLTNKKRWKDCKIRVFIGGKINRIDHDRRAMATLLSKFRIDFSDITVLGDINVKPKKEHVTAFEEMIEPYRLKEDDMEQDAAERLKNSEPWRITDNELELYRAKTNRQIRLNELLKEHSSTANLIVMSLPLARKGAVSSALYMAWLEALSKDLPPILLVRGNHQSVLTFYS

>XP_020797507.1_Boleophthalmus

MGDPDEMVNDEHDVDALMTDELNDIRIVGTLTVILLLGISVAGMEWEAKAQIVLLVILLAAIGNYFIGTFMPTKNKEPKGFFGYNTAIFLENLGPDFREDETFFSVFAIFFPAATGILAGANISGDLADPQSAIPKGTLLAILITGVTYVAVAISTGSCIVRDATGDHNDTVSDTVNCTDAACSLGFDFSICREGGCQYGLMNDFQVMSLVSAFGPLITAGIFSATLSSALASLVSAPKVFQALCKDNIYPSLEVFAKGYGKNNEPLRGYVLTFCIGLAFILIAELNIIAPIISNFFLASYALINFSVFHASLANSPGWRPSFKYYNMWVSLGGAVLCCAVMFIINWWAALVTLLIVLALYVYVGYKKPDVNWGSSTQALIYNQALTHCLDLTGVPEHIKNFRPQCLVMAGFPNSRPALLHLVNSFTKNVGLMICGHVRTLSRRPNLKDLTLEHGRCQRWLQKKNIRAFYSPVLADNLRHGAQLLLQASGLGQLKPNTLVMGFKKNWSDGDMRHTEIYINTIHDAFDLQFGVVILRLQDGLDISHIQGQDELSTVDKPKDTSVISDPNPPNNHNIPLITKERSPPLSLTDQRLLESSQIFKKKQGKGTIDVWWLFDDGGLTLLIPYLLTNRGKWGDCRIRVFMGGKINRIDHDRRAMAMLLSRFRIDFSDIIVLGDINTKPKKHNKLSFKELMDPYRLREDDMQQDEAERLKALEPWRITDNELELYKAKTNRQIRLNELLQEHSNTAKLIVISMPLARKGTVSSALYMCWLETLSKDLPPVLLVRGNHQSVLTFYS

>XP_020773468.1_Boleophthalmus

MGQVTSRLRRPAGVPRVHFSTDEGPPRYSSHHFDGRSPRHGDPPTITVSEIDASSLSERRGSRMELRRASFYSTMDLAPQLEHYASSLPRQQTRSRPSLEALRKAYEDGELGIEAAGAQADVDSLPPPPDEEAADGSKENAPVRFGWIIGVMVRCMLNIWGVILFLRLSWITSQAGIVLVWVIILMSSAVTTVTALSVSAIATNGRVISGGAYFMISRSLGPEIGGPIGMVFSFANALACALNTVGFAEVVRDLLKDFDAQMVDDVNDVRIIGVITVTVLLLISLAGMEWESKAQILFFIVLLVSFANYFVGTFIPAPKEKQAVGIFSYQADIFVTNLMPDWRGPDGNFFQMFAIFFPSATGILSGVNICGDLKDPMSAIPKGTLMAIFWTTLSYLAIALTSAATTVRDASGNLTDFMTGNSTEGCVGLACKYGWNFTECIVSQTCEYGLANSVKLLGQLSGFYYLITAGVFAASLSSALGFLVSAPKVFQCLCKDNIYPYIGFFAKGYGKNNEPLRAYLLCFIISIAFILIAELNIIAALISNFFLCSYGLINFSCFHASITNSPGWRPAFHYYSKWTALFGAVISIVLMFLFTWWAALVTFGIIFFLFGYVNYTKPKVNWGSSVQAGTYNMALSYSVSLSGVEDHVKNFRPQCMVLTGPPNQRPALVDFVGSFTKHVSLMICGDIIMEQDRKHRAPNATDWLVKWMNQRKVRSFYTPFSSDSLRAGARYLLQASGLGKLKPNTLVLGFKANWRESSPESIEDYITTIYDTFDSNYCLCILRMMDGQDVLDQFDSEVNSGFEPDEPTENDDQQSPERDSADDVSDKGSRDQVKTVFQNDQGKKTIDVYWIADDGGLTLLVPYLLTRRKRWRRSKVRVFIVGDEQNMEDGRNEMLALLKRFRLDFNDVIVMTDSEKRPQTKNLNRFVESVAPFRLYDEQQDGVSLQGLRQDYPWKISDKEFEAFKLKSERKVRLNEIIRKNSQHASLVLVSLPVPQIDCPSALYMAWLDTLTCGLHCPAVLIRGNQQNVLTFYCQ

>XP_020790949.1_Boleophthalmus

MELPVVNEGVRLNAFRPSVVSNGKAASSSGESEYYGYFGDRCSVASSRADSSITGYETLDAPPHYDFYANTEVWGRRRKSRPSLYQLYANPQEDFKPPVYEESPVGHSGETTDEDDEDEQKEPPAEPTRFGWVQGVLIRCMLNIWGVILYLRLPWITAQAGIGLTWVIILLSSTITGITGLSTSAIATNGKVKGGGTYFLISRSLGPELGGSIGLIFAFANAVAVAMHTVGFAETVTDLMREHGAVMVDKTNDIRIIGIITVTCLLGISLAGMSWESKAQVVFFFVIMVSFASYIVGTIMPASVEKQSKGFFSYRADIFAQNFVPGWRGKEGNFFGMFSIFFPSATGILAGANISGDLKNPTVAIPKGTLLAIFFTTFSYLIISATIGACVVRDASGVLNDTLVNSSNPESCVGTACQYGWDFSECTQNGTCTFGLINYYQTMSLVSAFAPLITAGIFGATLSSALACLVSAPKVFQCLCKDKLYPFIGFFGKGYGKNDEPLRGYLLTYIIASCFILIAELNTIAPIISNFFLCSYTLINFSCFHASITNSPGWRPSFRFYSKWLSLLGAVCCVAIMFLLTWWAALIAFGVVFVLLGYTLYKKPAVNWGSSVQASSYNLALNQCVGLNQVEDHVKNYRPQCLVLTGPPSSRPALVDLVSCFTKGLSLMMCADVMTSGPSASVVQKENGEKHVSWLNERKVKSFYRGVVSSDLRTGVNVLMQGAGLGRIKPNVLLLGFKKSWCSDSPQALHEYIGILHDAFELQYGVCMLRMKEGLDVSHPPQSHVNPAFDEGPEISYYLSTCYTTITSSPEPQTTTVFRKKQEKKTIDVYWLCDDGGLILLLPYLLTRRNRWSRCKVRVFVSGRPDKKEEQKQEVLALIKKFRLGFHDVEVLTDVYDNPQPHNVQLFENMVAPYRKDQSPKQDLGCTPTKVCDGPWMITEQDWERNKAKTLRQIRLNEILLDYSREAALILVTMPVGRKGGCPSSLYMAWLDFISRDLRPPVLLVRGNQENVLTFYCQ

>XP_020791773.1_Boleophthalmus

MPTNFTVVPVEDAEAPSSSNATSSPEAKPISLGQIFERSGQDTPLDSPQEAQIDDAEGQKESTPFLNADNSSEPDYDGKNMALFEEEMDSAPMVSSLLSKLANYTNLTQGVREHEEYEDGVRKVTVTVPPMGTFIGVYLPCMQNILGVILFIRLSWIVGTAGILGSFAIVSMCCICTLLTAISMSAIATNGVVPAGGSYYMISRSLGPEFGGAVGLCFYLGTTFAGSMYILGTIEILLTYIIPYTRVFEAENREGEAWANNMRVFGTGCLLIMALIVFVGVKYVNKLALVFLSCVILSILATYAGVIKTLIEPPEVNVCLVGNRSLKNNMFETCAKTAMVENVTVMTQLWDLFCSNETCDEYFEFNNLTQIKAIPGLLSGVIKDNLWGDYGPAGTMLEKKTLPSVPATDTSKEIVKHYVFNDITTYFTLLVGIYFPSVTGIMAGSNRSGDLRDAQRSIPVGTIMAILTTSFIYISCVVFFGACIEGVVLRDKFGDSIGQIPVIGILAWPSPWVIVIGSFFSCCGAGLQSLTGAPRLLQAIARDGIIPFLQVFGHGKSNGEPTWALLLTVGICEIGILIASLDDVAPILSMFFLMCYLFVNLACAVQTLLRAPNWRPRFKFYHWTLSFLGMSLCLSLMFVSSWYYALVAMVIAGCIYKYIEYRGAEKEWGDGIRGLSLNAARYALIHLEEAPLHTKNWRPQLLVLCKPDTDLSVKHPRLLSFTSQLKAGKGLTIVCSVLQGTYMTRGQDVKTAEKSLKAAMVAEKTKGFHHVVVSSNLRDGFSVLIQSAGLGGMKHNSVLMAWPAAWQQASEPESRKNFIETIRETTSAHQALLVAKNIDHFPDNSQRLKEGTIDVWWIVHDGGLLMLLPFLLRQHKVWRKCRMRIFTVAQMDDNSIQMKKDLQMFVYHLRIDAVVEVVEMHDSDISAFTYEKTLVMEQRSQMLKQMQLSRTEREREAQLIHDRNTASHSDKTANSAPPDRVHMTWTKDKLISERAKQKEGMAVKDMFNMRPEWETLNQSNVRRMHTAVKLNEVVKKKSKESQLVLLNMPGPPKNKKGDENYMEFLEVLTEGLDRVLLVRGGGREIITIYS

***Homo sapiens***

>sp|P55011|S12A2_HUMAN NKCC1_Homo

MEPRPTAPSSGAPGLAGVGETPSAAALAAARVELPGTAVPSVPEDAAPASRDGGGVRDEG

PAAAGDGLGRPLGPTPSQSRFQVDLVSENAGRAAAAAAAAAAAAAAAGAGAGAKQTPADG

EASGESEPAKGSEEAKGRFRVNFVDPAASSSAEDSLSDAAGVGVDGPNVSFQNGGDTVLS

EGSSLHSGGGGGSGHHQHYYYDTHTNTYYLRTFGHNTMDAVPRIDHYRHTAAQLGEKLLR

PSLAELHDELEKEPFEDGFANGEESTPTRDAVVTYTAESKGVVKFGWIKGVLVRCMLNIW

GVMLFIRLSWIVGQAGIGLSVLVIMMATVVTTITGLSTSAIATNGFVRGGGAYYLISRSL

GPEFGGAIGLIFAFANAVAVAMYVVGFAETVVELLKEHSILMIDEINDIRIIGAITVVIL

LGISVAGMEWEAKAQIVLLVILLLAIGDFVIGTFIPLESKKPKGFFGYKSEIFNENFGPD

FREEETFFSVFAIFFPAATGILAGANISGDLADPQSAIPKGTLLAILITTLVYVGIAVSV

GSCVVRDATGNVNDTIVTELTNCTSAACKLNFDFSSCESSPCSYGLMNNFQVMSMVSGFT

PLISAGIFSATLSSALASLVSAPKIFQALCKDNIYPAFQMFAKGYGKNNEPLRGYILTFL

IALGFILIAELNVIAPIISNFFLASYALINFSVFHASLAKSPGWRPAFKYYNMWISLLGA

ILCCIVMFVINWWAALLTYVIVLGLYIYVTYKKPDVNWGSSTQALTYLNALQHSIRLSGV

EDHVKNFRPQCLVMTGAPNSRPALLHLVHDFTKNVGLMICGHVHMGPRRQAMKEMSIDQA

KYQRWLIKNKMKAFYAPVHADDLREGAQYLMQAAGLGRMKPNTLVLGFKKDWLQADMRDV

DMYINLFHDAFDIQYGVVVIRLKEGLDISHLQGQEELLSSQEKSPGTKDVVVSVEYSKKS

DLDTSKPLSEKPITHKVEEEDGKTATQPLLKKESKGPIVPLNVADQKLLEASTQFQKKQG

KNTIDVWWLFDDGGLTLLIPYLLTTKKKWKDCKIRVFIGGKINRIDHDRRAMATLLSKFR

IDFSDIMVLGDINTKPKKENIIAFEEIIEPYRLHEDDKEQDIADKMKEDEPWRITDNELE

LYKTKTYRQIRLNELLKEHSSTANIIVMSLPVARKGAVSSALYMAWLEALSKDLPPILLV

RGNHQSVLTFYS

>sp|Q13621|S12A1_HUMAN NKCC2_Homo

MSLNNSSNVFLDSVPSNTNRFQVSVINENHESSAAADDNTDPPHYEETSFGDEAQKRLRI

SFRPGNQECYDNFLQSGETAKTDASFHAYDSHTNTYYLQTFGHNTMDAVPKIEYYRNTGS

ISGPKVNRPSLLEIHEQLAKNVAVTPSSADRVANGDGIPGDEQAENKEDDQAGVVKFGWV

KGVLVRCMLNIWGVMLFIRLSWIVGEAGIGLGVLIILLSTMVTSITGLSTSAIATNGFVR

GGGAYYLISRSLGPEFGGSIGLIFAFANAVAVAMYVVGFAETVVDLLKESDSMMVDPTND

IRIIGSITVVILLGISVAGMEWEAKAQVILLVILLIAIANFFIGTVIPSNNEKKSRGFFN

YQASIFAENFGPRFTKGEGFFSVFAIFFPAATGILAGANISGDLEDPQDAIPRGTMLAIF

ITTVAYLGVAICVGACVVRDATGNMNDTIISGMNCNGSAACGLGYDFSRCRHEPCQYGLM

NNFQVMSMVSGFGPLITAGIFSATLSSALASLVSAPKVFQALCKDNIYKALQFFAKGYGK

NNEPLRGYILTFLIAMAFILIAELNTIAPIISNFFLASYALINFSCFHASYAKSPGWRPA

YGIYNMWVSLFGAVLCCAVMFVINWWAAVITYVIEFFLYVYVTCKKPDVNWGSSTQALSY

VSALDNALELTTVEDHVKNFRPQCIVLTGGPMTRPALLDITHAFTKNSGLCICCEVFVGP

RKLCVKEMNSGMAKKQAWLIKNKIKAFYAAVAADCFRDGVRSLLQASGLGRMKPNTLVIG

YKKNWRKAPLTEIENYVGIIHDAFDFEIGVVIVRISQGFDISQVLQVQEELERLEQERLA

LEATIKDNECEEESGGIRGLFKKAGKLNITKTTPKKDGSINTSQSMHVGEFNQKLVEAST

QFKKKQEKGTIDVWWLFDDGGLTLLIPYILTLRKKWKDCKLRIYVGGKINRIEEEKIVMA

SLLSKFRIKFADIHIIGDINIRPNKESWKVFEEMIEPYRLHESCKDLTTAEKLKRETPWK

ITDAELEAVKEKSYRQVRLNELLQEHSRAANLIVLSLPVARKGSISDLLYMAWLEILTKN

LPPVLLVRGNHKNVLTFYS

>tr|Q53ZR1|Q53ZR1_HUMAN_Homo

MEPRPTAPSSGAPGLAGVGETPSAAALAAARVELPGTAVPSVPEDAAPASRDGGGVRDEG

PAAAGDGLGRPLGPTPSQSRFQVDLVSENAGRAAAAAAAAAAAAAAAGAGAGAKQTPADG

EASGESEPAKGSEEAKGRFRVNFVDPAASSSAEDSLSDAAGVGVDGPNVSFQNGGDTVLS

EGSSLHSGGGGGSGHHQHYYYDTHTNTYYLRTFGHNTMDAVPRIDHYRHTAAQLGEKLLR

PSLAELHDELEKEPFEDGFANGEESTPTRDAVVTYTAESKGVVKFGWIKGVLVRCMLNIW

GVMLFIRLSWIVGQAGIGLSVLVIMMATVVTTITGLSTSAIATNGFVRGGGAYYLISRSL

GPEFGGAIGLIFAFANAVAVAMYVVGFAETVVELLKEHSILMIDEINDIRIIGAITVVIL

LGISVAGMEWEAKAQIVLLVILLLAIGDFVIGTFIPLESKKPKGFFGYKSEIFNENFGPD

FREEETFFSVFAIFFPAATGILAGANISGDLADPQSAIPKGTLLAILITTLVYVGIAVSV

GSCVVRDATGNVNDTIVTELTNCTSAACKLNFDFSSCESSPCSYGLMNNFQVMSMVSGFT

PLISAGIFSATLSSALASLVSAPKIFQALCKDNIYPAFQMFAKGYGKNNEPLRGYILTFL

IALGFILIAELNVIAPIISNFFLASYALINFSVFHASLAKSPGWRPAFKYYNMWISLLGA

ILCCIVMFVINWWAALLTYVIVLGLYIYVTYKKPDVNWGSSTQALTYLNALQHSIRLSGV

EDHVKNFRPQCLVMTGAPNSRPALLHLVHDFTKNVGLMICGHVHMGPRRQAMKEMSIDQA

KYQRWLIKNKMKAFYAPVHADDLREGAQYLMQAAGLGRMKPNTLVLGFKKDWLQADMRDV

DMYINLFHDAFDIQYGVVVIRLKEGLDISHLQGQEELLSSQEKSPGTKDVVVSVEYSKKS

DLDTSKPLSEKPITHKVEEEDGKTATQPLLKKESKGPIVPLNVADQKLLEASTQFQKKQG

KNTIDVWWLFDDGGLTLLIPYLLTTKKKWKDCKIRVFIGGKINRIDHDRRAMATLLSKFR

IDFSDIMVLGDINTKPKKENIIAFEEIIEPYRLHEDDKEQDIADKMKEDEPWRITDNELE

LYKTKTYRQIRLNELLKEHSSTANIIVMSLPVARKGAVSSALYMAWLEALSKDLPPILLV

RGNHQSVLTFYS

>tr|B7ZM24|B7ZM24_HUMAN SLC12A2_Homo

MEPRPTAPSSGAPGLAGVGETPSAAALAAARVELPGTAVSSVPEDAAPASRDGGGVRDEG

PAAAGDGLGRPLGPTPSQSRFQVDLVSENAGRAAAAAAAAAAAAAAAGAGAGAKQTPADG

EASGESEPAKGSEEAKGRFRVNFVDPAASSSAEDSLSDAAGVGVDGPNVSFQNGGDTVLS

EGSSLHSGGGGGSGHHQHYYYDTHTNTYYLRTFGHNTMDAVPRIDHYRHTAAQLGEKLLR

PSLAELHDELEKEPFEDGFANGEESTPTRDAVVTYTAESKGVVKFGWIKGVLVRCMLNIW

GVMLFIRLSWIVGQAGIGLSVLVIMMATVVTTITGLSTSAIATNGFVRGGGAYYLISRSL

GPEFGGAIGLIFAFANAVAVAMYVVGFAETVVELLKEHSILMIDEINDIRIIGAITVVIL

LGISVAGMEWEAKAQIVLLVILLLAIGDFVIGTFIPLESKKPKGFFGYKSEIFNENFGPD

FREEETFFSVFAIFFPAATGILAGANISGDLADPQSAIPKGTLLAILITTLVYVGIAVSV

GSCVVRDATGNVYDTIVTELTNCTSAACKLNFDFSSCESSPCSYGLMNNFQVMSMVSGFT

PLISAGIFSATLSSALASLVSAPKIFQALCKDNIYPAFQMFAKGYGKNNEPLRGYILTFL

IALGFILIAELNVIAPIISNFFLASYALINFSVFHASLAKSPGWRPAFKYYNMWISLLGA

ILCCIVMFVINWWAALLTYVIVLGLYIYVTYKKPDVNWGSSTQALTYLNALQHSIRLSGV

EDHVKNFRPQCLVMTGAPNSRPALLHLVHDFTKNVGLMICGHVHMGPRRQAMKEMSIDQA

KYQRWLIKNKMKAFYAPVHADDLREGAQYLMQAAGLGRMKPNTLVLGFKKDWLQADMRDV

DMYINLFHDAFDIQYGVVVIRLKEGLDISHLQGQEELLSSQEKSPGTKDVVVSVEYSKKS

DLDTSKPLSEKPITHKESKGPIVPLNVADQKLLEASTQFQKKQGKNTIDVWWLFDDGGLT

LLIPYLLTTKKKWKDCKIRVFIGGKINRIDHDRRAMATLLSKFRIDFSDIMVLGDINTKP

KKENIIAFEEIIEPYRLHEDDKEQDIADKMKEDEPWRITDNELELYKTKTYRQIRLNELL

KEHSSTANIIVMSLPVARKGAVSSALYMAWLEALSKDLPPILLVRGNHQSVLTFYS

>tr|G3XAL9|G3XAL9_HUMAN_Homo

MEPRPTAPSSGAPGLAGVGETPSAAALAAARVELPGTAVPSVPEDAAPASRDGGGVRDEG

PAAAGDGLGRPLGPTPSQSRFQVDLVSENAGRAAAAAAAAAAAAAAAGAGAGAKQTPADG

EASGESEPAKGSEEAKGRFRVNFVDPAASSSAEDSLSDAAGVGVDGPNVSFQNGGDTVLS

EGSSLHSGGGGGSGHHQHYYYDTHTNTYYLRTFGHNTMDAVPRIDHYRHTAAQLGEKLLR

PSLAELHDELEKEPFEDGFANGEESTPTRDAVVTYTAESKGVVKFGWIKGVLVRCMLNIW

GVMLFIRLSWIVGQAGIGLSVLVIMMATVVTTITGLSTSAIATNGFVRGGGAYYLISRSL

GPEFGGAIGLIFAFANAVAVAMYVVGFAETVVELLKEHSILMIDEINDIRIIGAITVVIL

LGISVAGMEWEAKAQIVLLVILLLAIGDFVIGTFIPLESKKPKGFFGYKSEIFNENFGPD

FREEETFFSVFAIFFPAATGILAGANISGDLADPQSAIPKGTLLAILITTLVYVGIAVSV

GSCVVRDATGNVNDTIVTELTNCTSAACKLNFDFSSCESSPCSYGLMNNFQVMSMVSGFT

PLISAGIFSATLSSALASLVSAPKIFQALCKDNIYPAFQMFAKGYGKNNEPLRGYILTFL

IALGFILIAELNVIAPIISNFFLASYALINFSVFHASLAKSPGWRPAFKYYNMWISLLGA

ILCCIVMFVINWWAALLTYVIVLGLYIYVTYKKPDVNWGSSTQALTYLNALQHSIRLSGV

EDHVKNFRPQCLVMTGAPNSRPALLHLVHDFTKNVGLMICGHVHMGPRRQAMKEMSIDQA

KYQRWLIKNKMKAFYAPVHADDLREGAQYLMQAAGLGRMKPNTLVLGFKKDWLQADMRDV

DMYINLFHDAFDIQYGVVVIRLKEGLDISHLQGQEELLSSQEKSPGTKDVVVSVEYSKKS

DLDTSKPLSEKPITHKVEEEDGKTATQPLLKKESKGPIVPLNVADQKLLEASTQFQKKQG

KNTIDVWWLFDDGGLTLLIPYLLTTKKKWKDCKIRVFIGGKINRIDHDRRAMATLLSKFR

IDFSDIMVLGDINTKPKKENIIAFEEIIEPYRLHEDDKEQDIADKMKEDEPWRITDNELE

LYKTKFYEPC

>sp|Q9UP95|S12A4_HUMAN KCC1_Homo

MPHFTVVPVDGPRRGDYDNLEGLSWVDYGERAELDDSDGHGNHRESSPFLSPLEASRGID

YYDRNLALFEEELDIRPKVSSLLGKLVSYTNLTQGAKEHEEAESGEGTRRRAAEAPSMGT

LMGVYLPCLQNIFGVILFLRLTWMVGTAGVLQALLIVLICCCCTLLTAISMSAIATNGVV

PAGGSYFMISRSLGPEFGGAVGLCFYLGTTFAAAMYILGAIEILLTYIAPPAAIFYPSGA

HDTSNATLNNMRVYGTIFLTFMTLVVFVGVKYVNKFASLFLACVIISILSIYAGGIKSIF

DPPVFPVCMLGNRTLSRDQFDICAKTAVVDNETVATQLWSFFCHSPNLTTDSCDPYFMLN

NVTEIPGIPGAAAGVLQENLWSAYLEKGDIVEKHGLPSADAPSLKESLPLYVVADIATSF

TVLVGIFFPSVTGIMAGSNRSGDLRDAQKSIPVGTILAIITTSLVYFSSVVLFGACIEGV

VLRDKYGDGVSRNLVVGTLAWPSPWVIVIGSFFSTCGAGLQSLTGAPRLLQAIAKDNIIP

FLRVFGHGKVNGEPTWALLLTALIAELGILIASLDMVAPILSMFFLMCYLFVNLACAVQT

LLRTPNWRPRFKYYHWALSFLGMSLCLALMFVSSWYYALVAMLIAGMIYKYIEYQGAEKE

WGDGIRGLSLSAARYALLRLEEGPPHTKNWRPQLLVLLKLDEDLHVKYPRLLTFASQLKA

GKGLTIVGSVIQGSFLESYGEAQAAEQTIKNMMEIEKVKGFCQVVVASKVREGLAHLIQS

CGLGGMRHNSVVLGWPYGWRQSEDPRAWKTFIDTVRCTTAAHLALLVPKNIAFYPSNHER

YLEGHIDVWWIVHDGGMLMLLPFLLRQHKVWRKCRMRIFTVAQMDDNSIQMKKDLAVFLY

HLRLEAEVEVVEMHNSDISAYTYERTLMMEQRSQMLRQMRLTKTEREREAQLVKDRHSAL

RLESLYSDEEDESAVGADKIQMTWTRDKYMTETWDPSHAPDNFRELVHIKPDQSNVRRMH

TAVKLNEVIVTRSHDARLVLLNMPGPPRNSEGDENYMEFLEVLTEGLERVLLVRGGGREV

ITIYS

>sp|Q9H2X9|S12A5_HUMAN KCC2_Homo

MSRRFTVTSLPPAGPARSPDPESRRHSVADPRHLPGEDVKGDGNPKESSPFINSTDTEKG

KEYDGKNMALFEEEMDTSPMVSSLLSGLANYTNLPQGSREHEEAENNEGGKKKPVQAPRM

GTFMGVYLPCLQNIFGVILFLRLTWVVGIAGIMESFCMVFICCSCTMLTAISMSAIATNG

VVPAGGSYYMISRSLGPEFGGAVGLCFYLGTTFAGAMYILGTIEILLAYLFPAMAIFKAE

DASGEAAAMLNNMRVYGTCVLTCMATVVFVGVKYVNKFALVFLGCVILSILAIYAGVIKS

AFDPPNFPICLLGNRTLSRHGFDVCAKLAWEGNETVTTRLWGLFCSSRFLNATCDEYFTR

NNVTEIQGIPGAASGLIKENLWSSYLTKGVIVERSGMTSVGLADGTPIDMDHPYVFSDMT

SYFTLLVGIYFPSVTGIMAGSNRSGDLRDAQKSIPTGTILAIATTSAVYISSVVLFGACI

EGVVLRDKFGEAVNGNLVVGTLAWPSPWVIVIGSFFSTCGAGLQSLTGAPRLLQAISRDG

IVPFLQVFGHGKANGEPTWALLLTACICEIGILIASLDEVAPILSMFFLMCYMFVNLACA

VQTLLRTPNWRPRFRYYHWTLSFLGMSLCLALMFICSWYYALVAMLIAGLIYKYIEYRGA

EKEWGDGIRGLSLSAARYALLRLEEGPPHTKNWRPQLLVLVRVDQDQNVVHPQLLSLTSQ

LKAGKGLTIVGSVLEGTFLENHPQAQRAEESIRRLMEAEKVKGFCQVVISSNLRDGVSHL

IQSGGLGGLQHNTVLVGWPRNWRQKEDHQTWRNFIELVRETTAGHLALLVTKNVSMFPGN

PERFSEGSIDVWWIVHDGGMLMLLPFLLRHHKVWRKCKMRIFTVAQMDDNSIQMKKDLTT

FLYHLRITAEVEVVEMHESDISAYTYEKTLVMEQRSQILKQMHLTKNEREREIQSITDES

RGSIRRKNPANTRLRLNVPEETAGDSEEKPEEEVQLIHDQSAPSCPSSSPSPGEEPEGEG

ETDPEKVHLTWTKDKSVAEKNKGPSPVSSEGIKDFFSMKPEWENLNQSNVRRMHTAVRLN

EVIVKKSRDAKLVLLNMPGPPRNRNGDENYMEFLEVLTEHLDRVMLVRGGGREVITIYS

>sp|Q9UHW9|S12A6_HUMAN KCC3_Homo

MHPPETTTKMASVRFMVTPTKIDDIPGLSDTSPDLSSRSSSRVRFSSRESVPETSRSEPM

SEMSGATTSLATVALDPPSDRTSHPQDVIEDLSQNSITGEHSQLLDDGHKKARNAYLNNS

NYEEGDEYFDKNLALFEEEMDTRPKVSSLLNRMANYTNLTQGAKEHEEAENITEGKKKPT

KTPQMGTFMGVYLPCLQNIFGVILFLRLTWVVGTAGVLQAFAIVLICCCCTMLTAISMSA

IATNGVVPAGGSYFMISRALGPEFGGAVGLCFYLGTTFAAAMYILGAIEIFLVYIVPRAA

IFHSDDALKESAAMLNNMRVYGTAFLVLMVLVVFIGVRYVNKFASLFLACVIVSILAIYA

GAIKSSFAPPHFPVCMLGNRTLSSRHIDVCSKTKEINNMTVPSKLWGFFCNSSQFFNATC

DEYFVHNNVTSIQGIPGLASGIITENLWSNYLPKGEIIEKPSAKSSDVLGSLNHEYVLVD

ITTSFTLLVGIFFPSVTGIMAGSNRSGDLKDAQKSIPIGTILAILTTSFVYLSNVVLFGA

CIEGVVLRDKFGDAVKGNLVVGTLSWPSPWVIVIGSFFSTCGAGLQSLTGAPRLLQAIAK

DNIIPFLRVFGHSKANGEPTWALLLTAAIAELGILIASLDLVAPILSMFFLMCYLFVNLA

CALQTLLRTPNWRPRFRYYHWALSFMGMSICLALMFISSWYYAIVAMVIAGMIYKYIEYQ

GAEKEWGDGIRGLSLSAARFALLRLEEGPPHTKNWRPQLLVLLKLDEDLHVKHPRLLTFA

SQLKAGKGLTIVGSVIVGNFLENYGEALAAEQTIKHLMEAEKVKGFCQLVVAAKLREGIS

HLIQSCGLGGMKHNTVVMGWPNGWRQSEDARAWKTFIGTVRVTTAAHLALLVAKNISFFP

SNVEQFSEGNIDVWWIVHDGGMLMLLPFLLKQHKVWRKCSIRIFTVAQLEDNSIQMKKDL

ATFLYHLRIEAEVEVVEMHDSDISAYTYERTLMMEQRSQMLRHMRLSKTERDREAQLVKD

RNSMLRLTSIGSDEDEETETYQEKVHMTWTKDKYMASRGQKAKSMEGFQDLLNMRPDQSN

VRRMHTAVKLNEVIVNKSHEAKLVLLNMPGPPRNPEGDENYMEFLEVLTEGLERVLLVRG

GGSEVITIYS

>sp|P55017|S12A3_HUMAN NCC_Homo

MAELPTTETPGDATLCSGRFTISTLLSSDEPSPPAAYDSSHPSHLTHSSTFCMRTFGYNT

IDVVPTYEHYANSTQPGEPRKVRPTLADLHSFLKQEGRHLHALAFDSRPSHEMTDGLVEG

EAGTSSEKNPEEPVRFGWVKGVMIRCMLNIWGVILYLRLPWITAQAGIVLTWIIILLSVT

VTSITGLSISAISTNGKVKSGGTYFLISRSLGPELGGSIGLIFAFANAVGVAMHTVGFAE

TVRDLLQEYGAPIVDPINDIRIIAVVSVTVLLAISLAGMEWESKAQVLFFLVIMVSFANY

LVGTLIPPSEDKASKGFFSYRADIFVQNLVPDWRGPDGTFFGMFSIFFPSATGILAGANI

SGDLKDPAIAIPKGTLMAIFWTTISYLAISATIGSCVVRDASGVLNDTVTPGWGACEGLA

CSYGWNFTECTQQHSCHYGLINYYQTMSMVSGFAPLITAGIFGATLSSALACLVSAAKVF

QCLCEDQLYPLIGFFGKGYGKNKEPVRGYLLAYAIAVAFIIIAELNTIAPIISNFFLCSY

ALINFSCFHASITNSPGWRPSFQYYNKWAALFGAIISVVIMFLLTWWAALIAIGVVLFLL

LYVIYKKPEVNWGSSVQAGSYNLALSYSVGLNEVEDHIKNYRPQCLVLTGPPNFRPALVD

FVGTFTRNLSLMICGHVLIGPHKQRMPELQLIANGHTKWLNKRKIKAFYSDVIAEDLRRG

VQILMQAAGLGRMKPNILVVGFKKNWQSAHPATVEDYIGILHDAFDFNYGVCVMRMREGL

NVSKMMQAHINPVFDPAEDGKEASARVDPKALVKEEQATTIFQSEQGKKTIDIYWLFDDG

GLTLLIPYLLGRKRRWSKCKIRVFVGGQINRMDQERKAIISLLSKFRLGFHEVHILPDIN

QNPRAEHTKRFEDMIAPFRLNDGFKDEATVNEMRRDCPWKISDEEITKNRVKSLRQVRLN

EIVLDYSRDAALIVITLPIGRKGKCPSSLYMAWLETLSQDLRPPVILIRGNQENVLTFYC

Q
