## Supplementary material for "Evolved for success in novel environments: The round goby genome": Supp Materials: Supplemental_Material_S9.docx

### NLR methods

In order to identify members of the large multigenic family of fish-specific NACHT and Leucine-Rich Repeats containing genes (NLRs; the fish-specific subset is also known as NLR-C; Laing et al. 2008), an alignment of 368 zebrafish NLR-C proteins was obtained from Howe *et al*. 2016 (Howe et al. 2016). Sequences from that alignment were used for tblastn queries as described above. Matching sequences from the round goby assembly were used as queries for a second round of BLAST, resulting in an initial set of 357 candidate regions (Supplementary file NLR_blast_357_candidate_regions.fas ). In addition, the “canonic” NLRs conserved in all vertebrates (*NLRC3*, *NLRC5*, *NLRX1*, *NOD1*, *NOD2*, *CIITA*, *NWD1*, *NWD2* and *APAF1*) were identified with a TBLASTN-alignment approach using queries from multiple species as described above (Supplementary Table Immune gene queries). A set of sequences for the largest NLR exon (~600 amino acids, contains the signature NACHT domain) was inferred from the “candidate” dataset and aligned to corresponding exon of the zebrafish alignment using MAFFT v7.310 (Katoh & Standley 2013) with the --add option. This alignment was then used to generate a Hidden Markov profile model using HMMER (Eddy 2011) version 3.2.1. Additional profiles were generated for the zebrafish alignment of this exon itself (without goby sequences) and for the second largest exon in the zebrafish alignment (containing a B30.2 domain). A fourth model was created for the DNA sequence of Leucine-rich repeats from zebrafish NLRs. All new models are available in the Supplementary folder HMM_models. Finally, hmmsearch from HMMER was used to scan the round goby assembly for the presence of both the new models and the NLR-related domain models (CARD, FIIND, FISNA, NACHT, PYD/PYRIN, PRY, SPRY, all LRR models) from the Pfam-A database (version 31.0; Finn et al. 2014a), using default parameters and an e-value cut-off of 1e-10^3^ (1e-10^1^ for the leucine-rich repeats). BEDtools was used to merge adjacent model hits of the same type (usually resulting from frameshifts); the allowed distance between model hits was 200 bp for NACHT (new models, FISNA, NACHT) and B30.2 (new model, PRY, SPRY) exons; 20 bp for all the others. The obtained dataset was cross-compared with tblastn results and used to generate a final annotation of the domains in round goby NLR-C family members (Supplementary file NLR-C_and_PYD-associated domains.xlsx), consisting of 25 PYRIN , 1 N-terminal CARD, 12 C-terminal CARD, 343 FISNA-NACHT and 178 B30.2 domains. Interestingly, most of the identified NACHT exons contained frameshifts or were split in two. This may indicate that they could be pseudogenes. However, most cases contained a single insertion and otherwise resembled the NLRs predicted as intact. Accordingly, the pseudogenization of all of them must have been a fairly recent event, or pseudogenes themselves might be duplicating (Wang et al. 2018b), or many of these frameshifts result from actual intronization, or from sequencing errors at homopolymer stretches. Indeed, automatic gene annotation with Maker predicted a complicated exon-intron structure for most NACHT exons (data not shown), which is why we favor a scenario where the frameshift-containing regions are spliced out. For the phylogenetic analysis, only clearly intact NLRs were used. All 61 intact (no large indels, frameshifts or stop codons in the assembly) round goby NACHT-containing exons were manually curated and extended to the nearest putative splice sites. These sequences were then aligned to the 368 zebrafish NACHT exons using MAFFT --add. Next, all predicted NACHT-containing proteins of NCBI *B. pectinirostris* annotation release 100 were downloaded from NCBI and aligned to the NACHT alignment using MAFFT with the --add and --keeplength options, effectively removing all parts of the proteins that did not align with the NACHT-containing exon. Finally, human NLRs and canonic NLRs from zebrafish and the round goby were added to the alignment as above, and all sequences with a length of 0 (mostly zebrafish partial NLRs missing NACHT) were removed from the alignment. Maximum likelihood trees were produced with RAxML-PTHREADS, PROTCATJTT, rapid bootstrap and 500 bootstrap replicates (Stamatakis 2006). The final trees were imported into FigTree (http://tree.bio.ed.ac.uk/software/figtree/), and subsequently Adobe Illustrator. The alignments were inspected manually for presence of conserved elements and sequence logos for these were generated with WebLogo (Crooks et al. 2004). Finally, we performed a survey of the PYD domains, Peptidase_C14 domains (Caspases) and CARD. All cases of a PYD domain followed by an adjacent CARD in the round goby (putative apoptosis-associated speck-like protein containing a CARD (ASC), also known as PYD-CARD or PYCARD) were identified from the HMMER3 dataset. The open reading frames containing these were translated, concatenated, and aligned with similarly structured proteins from human, mouse, lizard, frog and all the fish in Ensembl, and with PYD-CARDs identified from the other available goby assemblies. A phylogenetic tree was generated as described above.
