## Supplementary material for "Evolved for success in novel environments: The round goby genome": Supp Materials: Supplemental_Material_S13.docx

**Methods for 3’ and 5’ RACE**

Isolation of RNA

A juvenile specimen of *Neogobius melanostomus* was ground using liquid nitrogen and mortar and pestle. The resulting frozen powder was added to Trizol (2 x 10 ml) and incubated for 5 min on ice. Debris was removed by centrifugation and the lysed sample transferred to fresh tubes and incubated another 5 min on ice. 2 ml of Chloroform was added (each tube) and samples incubated on ice for 5 min. Phases were separated by centrifugation (45 min, 4 °C, 4000 rpm), the aqueous phase transferred to a new tube, 10 ml isopropanol added per sample and placed at -80°C for a couple of hours. RNA was pelleted by centrifugation (45 min, 4°C, 4000 rpm). Pellets were washed twice with 75% EtOH (in nuclease free water) and dried on ice. RNA was dissolved in nuclease free water.

Removal of DNA

20 µg of RNA for 5`RACE and 2.7 µg for 3`RACE, respectively, were subjected to DNAse treatment (RQ1 RNAse free DNAse from Promega according to manufacturer’s instructions).

Amplification of cDNA ends with RACE

DNAse treated RNA was transcribed to cDNA with the FirstChoice RLM-RACE kit from Invitrogen according to the manufacturers instructions. PCR reactions were done using FastStart from Roche in a 20µl reaction volume in a nested design, with primer 1 used in a first round of amplification and primer 2 in a second round of amplification.

PCR conditions were:

94°C	4min

94°C	30sec

X (primer specific)°C	30sec	35 x

72°C	Y (gene specific)

72°C	10min

4°C for ever

| gene | fw primer 1 (3’RACE) | Annealing temp/extension time(mm:ss) | Fw primer 2 (3`RACE) | Annealing temp/extension time | rev primer 1 (5’RACE) | Annealing temp/extension time | rev primer 2  (5’RACE) | Annealing temp/extension time |
| --- | --- | --- | --- | --- | --- | --- | --- | --- |
| DNMT1 | SG068 | 58°C/ 01:40 | SG069 | 58°C/ 01:40 |  |  |  |  |
| DNMT3_181 | SG134 | 58°C/ 02:00 | SG135 | 58°C/ 02:00 |  |  |  |  |
| DNMT3_331 | SG132 | 58°C/ 02:00 | SG133 | 58°C/ 02:00 |  |  |  |  |
| DNMT3_445 |  |  |  |  | SG128 | 58°C/ 02:00 | SG129 | 58°C/ 02:00 |
| AEBP2_258 | SG098 | 58°C/ 02:00 | SG099 | 58°C/ 02:00 |  |  |  |  |
| AEBP2_420 | SG100 | 58°C/ 02:00 | SG101 | 58°C/ 02:00 |  |  |  |  |
| EED | SG044 | 56°C / 01:30 | SG045 | 56°C / 01:30 |  |  |  |  |
| EZH12_55 | SG113 | 58°C/ 02:00 | SG114 | 58°C/ 02:00 | SG111 | 58°C/ 02:00 | SG112 | 58°C/ 02:00 |
| EZH12_275 | SG117 | 58°C/ 02:00 | SG118 | 58°C/ 02:00 |  |  |  |  |
| RBBP4 | SG060 | 58°C/ 02:00 | SG061 | 58°C/ 02:00 |  |  |  |  |
| SUZ12_4 | SG094 | 58°C/ 02:00 | SG095 | 58°C/ 02:00 | SG090 | 58°C/ 02:00 | SG091 | 58°C/ 02:00 |
| SUZ12_386 | SG088 | 58°C/ 02:00 | SG089 | 58°C/ 02:00 | SG107 | 58°C/ 02:00 | SG108 | 58°C/ 02:00 |

| **Primer name** | **sequence** |
| --- | --- |
| SG044 | CCAACCTCCTGCTTTCAGTC |
| SG045 | TCAGTCAGCAAAGACCATGC |
| SG060 | CTCTTTGGGACCTGAGGAAC |
| SG061 | TCCAGTGGCACAGACAGAAG |
| SG068 | GCAGACAGGCAGTTCAACAC |
| SG069 | CTGATCCCATGGTGTCTGC |
| SG088 | TCCAGGAGAAGACCGTCAAG |
| SG089 | TCATGAAGAACGGGTTTATCG |
| SG090 | CGAGTCATGCTGTCCCTTC |
| SG091 | TTGTTCTGGCATTTCTGTGAG |
| SG094 | GTGGCTCAGGGAGAAAACTG |
| SG095 | TTATGAAGAGCGGGTTCATTG |
| SG098 | TATCCCAGTTACCGCAGGAC |
| SG099 | CATGCTGCTAGACCCCAAC |
| SG100 | TGCTACACTGGAGCCCTGAG |
| SG101 | AGCCCTGAGGACATTCTGC |
| SG107 | ACTGCTTCACGGGACAACTC |
| SG108 | AAGAAGCCGGTGAAAGTCAG |
| SG111 | GAACGGACTGAGGCTTGAAC |
| SG112 | GATGGGAAACCAGACTCCAC |
| SG113 | TCCAGTTTCCTTTTCAACCTG |
| SG114 | GAACAATGACTTTGTCGTGGAC |
| SG117 | GATGATGGTGAACGGAGACC |
| SG118 | GCCAAAAGAGCGATTCAGAC |
| SG128 | TCGTACTCGGGCTCTGTGTC |
| SG129 | GTCTTGGGCGAGCTCTTCTC |
| SG132 | CCAGTCCTACTGCACTGTGTG |
| SG133 | GCTGGAGGTCATCCTCTGTG |
| SG134 | TGTTGGTGCTGAAGGACTTG |
| SG135 | CCATAACTGTGGGGATGGTC |
