## Supplementary material for "Evolved for success in novel environments: The round goby genome": Supp Materials: Supplemental_Table_S3.docx

| **Pattern recognition** | | |
| --- | --- | --- |
| Toll-like | TLR1/6 | Peptidoglycan/lipoproteins |
|  | TLR2 | Peptidoglycan/lipoproteins |
|  | TLR3 | dsRNA |
|  | TLR4 | Lipopolysaccharide |
|  | TLR5 | Flagellin |
|  | TLR7 | ssRNA |
|  | TLR8 | G-rich oligonucleotides |
|  | TLR9 | CpG dinucleotides |
|  | TLR14 | ? |
|  | TLR21 | ? |
|  | TLR22 | ? |
|  | TLR23 | ? |
|  | TLR25 | ? |
|  | TLR26 | ? |
| Other PRRs | CRP/APCS | C-reactive protein / Amyloid P component, serum |
|  | NOD1 | Nucleotide Binding Oligomerization Domain Containing 1 |
|  | NOD2 | Nucleotide Binding Oligomerization Domain Containing 2 |
| **Cytokines and chemokines** | | |
| Interleukins | IL1 | Inflammation |
|  | IL4/13 | Differentiation of T- and B-cells |
|  | IL6 | Regulation of inflammation and cell maturation |
|  | IL8 | Chemotactic factor for neutrophils |
|  | IL10 | Regulation of the adaptive response, antiinflammatory |
| Interferon | IFNg | Regulation of the immune response, also antiviral |
| Tumor necrosis factor | TNFa | Regulation of cell differentiation, apoptosis, metabolism etc. |
| **Key factors related to adaptive immunity** | | |
|  | AICDA | Somatic hypermutation, class switch and gene conversion |
|  | AIRE | Transcriptional regulator involved in T-cell selection |
|  | B2M | MHCI co-receptor, beta-2-microglobulin |
|  | CD4 | T-helper cell co-receptor |
|  | CD74 | Invariant chain, blocks MHCII binding pocket |
|  | CD8a | T-helper cell co-receptor |
|  | CD8b | T-helper cell co-receptor |
|  | CIITA | MHCII transactivator, transcription factor |
|  | IgH | Immunoglobulin heavy chain locus |
|  | MHCI | Major Histocompatibility Complex Class I |
|  | MHCII | Major Histocompatibility Complex Class II |
|  | RAG1 | Activates immunoglobulin V(D)J recombmination |
|  | RAG2 | Activates immunoglobulin V(D)J recombmination |
|  | TAP1/2/2T | Transports peptides across the ER membrane for MHCI loading |
|  | Tapasin | Mediates interaction between MHCI and the TAP complex |
| **Caspases** | | |
|  | CASP1-9 | Proteases involved in cell death and inflammation |
