## Supplementary material for "Evolved for success in novel environments: The round goby genome": Supp Materials: Supplemental_Table_S5.docx

| **Genome contig** | **Start core -500 bp** | **Stop core +500 bp** | **MHCI subtype** | **Strand** |
| --- | --- | --- | --- | --- |
| Contig_1105 | 41502 | 45359 | U-lineage | sense |
| Contig_4227 | 384413 | 386998 | U-lineage | sense |
| Contig_1105 | 367874 | 374038 | U-lineage | rev |
| Contig_1105 | 379314 | 383186 | U-lineage | rev |
| Contig_1105 | 389652 | 393489 | U-lineage | rev |
| Contig_1105 | 398857 | 402936 | U-lineage | rev |
| Contig_2107 | 3007487 | 3011566 | U-lineage | rev |
| Contig_2107 | 3035637 | 3040830 | U-lineage | rev |
| Contig_2107 | 3066825 | 3070867 | U-lineage | rev |
| Contig_2107 | 3097761 | 3103266 | U-lineage | rev |
| Contig_2107 | 3106129 | 3110656 | U-lineage | rev |
| Contig_2426 | 2038102 | 2040551 | U-lineage | rev |
| Contig_2426 | 2043798 | 2047326 | U-lineage | rev |
| Contig_3034 | 582725 | 585724 | U-lineage | rev |
| Contig_3034 | 632051 | 635014 | U-lineage | rev |
| Contig_3086 | 193828 | 198923 | U-lineage | rev |
| Contig_3086 | 214042 | 217138 | U-lineage | rev |
| Contig_3396 | 1033257 | 1039264 | U-lineage | rev |
| Contig_3396 | 407187 | 411970 | U-lineage | rev |
| Contig_3567 | 10956 | 16526 | U-lineage | rev |
| Contig_3567 | 25897 | 28329 | U-lineage | rev |
| Contig_3567 | 31217 | 36390 | U-lineage | rev |
| Contig_3958 | 58409 | 61913 | U-lineage | rev |
| Contig_3958 | 79735 | 83515 | U-lineage | rev |
| Contig_3958 | 94426 | 97813 | U-lineage | rev |
| Contig_4115 | 15846 | 21659 | U-lineage | rev |
