## Supplementary material for "Evolved for success in novel environments: The round goby genome": Supp Materials: Supplemental_Table_S6.docx

| **Genome contig** | **Start core -500 bp** | **Stop core +500 bp** | **MHCII subtype** | **Strand** | **Comment** |
| --- | --- | --- | --- | --- | --- |
| Contig_1748 | 279618 | 287164 | alpha | rev |  |
| Contig_541 | 7022014 | 7023644 | alpha | rev |  |
| Contig_447 | 2890257 | 2891827 | alpha | sense |  |
| Contig_4958 | 43169 | 44713 | alpha | sense |  |
| Contig_5344 | 24148 | 25705 | alpha | sense |  |
| Contig_541 | 7031331 | 7032958 | alpha | sense |  |
| Contig_4349 | 28226 | 29687 | alpha | sense | Short fragment, frameshifts |
| Contig_4413 | 50536 | 51844 | alpha | sense | Short fragment |
| Contig_1748 | 269829 | 271863 | beta | rev |  |
| Contig_4325 | 30835 | 34427 | beta | rev |  |
| Contig_4711 | 38466 | 40294 | beta | rev |  |
| Contig_541 | 7020573 | 7022154 | beta | rev |  |
| Contig_4349 | 58314 | 61051 | beta | sense |  |
| Contig_447 | 2886999 | 2889456 | beta | sense |  |
| Contig_447 | 2893878 | 2896525 | beta | sense |  |
| Contig_4958 | 23343 | 25695 | beta | sense |  |
| Contig_541 | 7032821 | 7034398 | beta | sense |  |
